# Autocrine insulin pathway signaling regulates actin dynamics in cell wound repair

**DOI:** 10.1101/2020.04.06.028662

**Authors:** Mitsutoshi Nakamura, Jeffrey M. Verboon, Tessa E. Allen, Maria Teresa Abreu-Blanco, Raymond Liu, Andrew N.M. Dominguez, Jeffrey J. Delrow, Susan M. Parkhurst

## Abstract

Cells are exposed to frequent mechanical and/or chemical stressors that can compromise the integrity of the plasma membrane and underlying cortical cytoskeleton. The molecular mechanisms driving the immediate repair response launched to restore the cell cortex and circumvent cell death are largely unknown. Using microarrays and drug-inhibition studies to assess gene expression, we find that initiation of cell wound repair in the *Drosophila* model is dependent on translation, whereas transcription is required for subsequent steps. We identified 253 genes whose expression is up-regulated (80) or down-regulated (173) in response to laser wounding. A subset of these genes were validated using RNAi knockdowns and exhibit aberrant actomyosin ring assembly and/or actin remodeling defects. Strikingly, we find that the canonical insulin signaling pathway controls actin dynamics through the actin regulators Girdin and Chickadee (profilin), and its disruption leads to abnormal wound repair. Our results provide new insight for understanding how cell wound repair proceeds in healthy individuals and those with diseases involving wound healing deficiencies.

## Introduction

Numerous cell types in the body are subject to high levels of stress daily. These stresses— physiological and/or environmental—can cause ruptures in the plasma membrane and its underlying cytoskeleton, requiring a rapid repair program to avert further damage, prevent infection/death, and restore normal function [1-11]. Injuries to individual cells also occur as a result of accidents/trauma, clinical interventions, and disease conditions, including diabetes, skin blistering disorders, and muscular dystrophies, as well as in response to pore forming toxins secreted by pathogenic bacteria [12-16]. Repair of these cell cortex lesions can be particularly troublesome when occurring alongside these fragile cell disease states or in a non-renewing and/or irreplaceable cell type. Thus, the importance of cell cortex continuity and delineating the molecular mechanisms regulating cell wound repair is of considerable clinical relevance, and important for advancing our knowledge of the many critical cell behaviors and fundamental regulations underpinning normal biological events that are co-opted for this repair process.

Aspects of single cell wound repair dynamics have been studied in *Xenopus* oocytes, sea urchin eggs, Dictyostelium, mammalian tissue culture cells, and the genetically-amenable *Drosophila* syncytial embryo [3, 17-23]. This repair is generally conserved among these organisms and occurs in four main phases (Fig. 1A). In the first phase, the wound expands as the cell recognizes the membrane breach, releases resting membrane tension, and subsequently forms a membranous plug to neutralize any flux between the extracellular space and cytoplasm. Second, the cell constructs an actomyosin ring that underlies the plasma membrane at the wound edge. Third, the actomyosin ring translocates inward to draw the wound area closed. Mechanistic variations exist during this step wherein the actomyosin ring in some models translocates through actin treadmilling (actin simultaneously polymerizes at the inner edge and depolymerizes at the outer edge of the actin ring), while others use myosin II for sarcomere-like contraction (anti-parallel actin filaments are directed past each other in opposing directions) [3, 22-26]. In the final step of wound repair, the plasma membrane and the underlying cortical cytoskeleton are remodeled returning them to their pre-wounded composition and organization. The mechanisms deployed by the cell for this remodeling have not yet been delineated.

**Fig 1.**
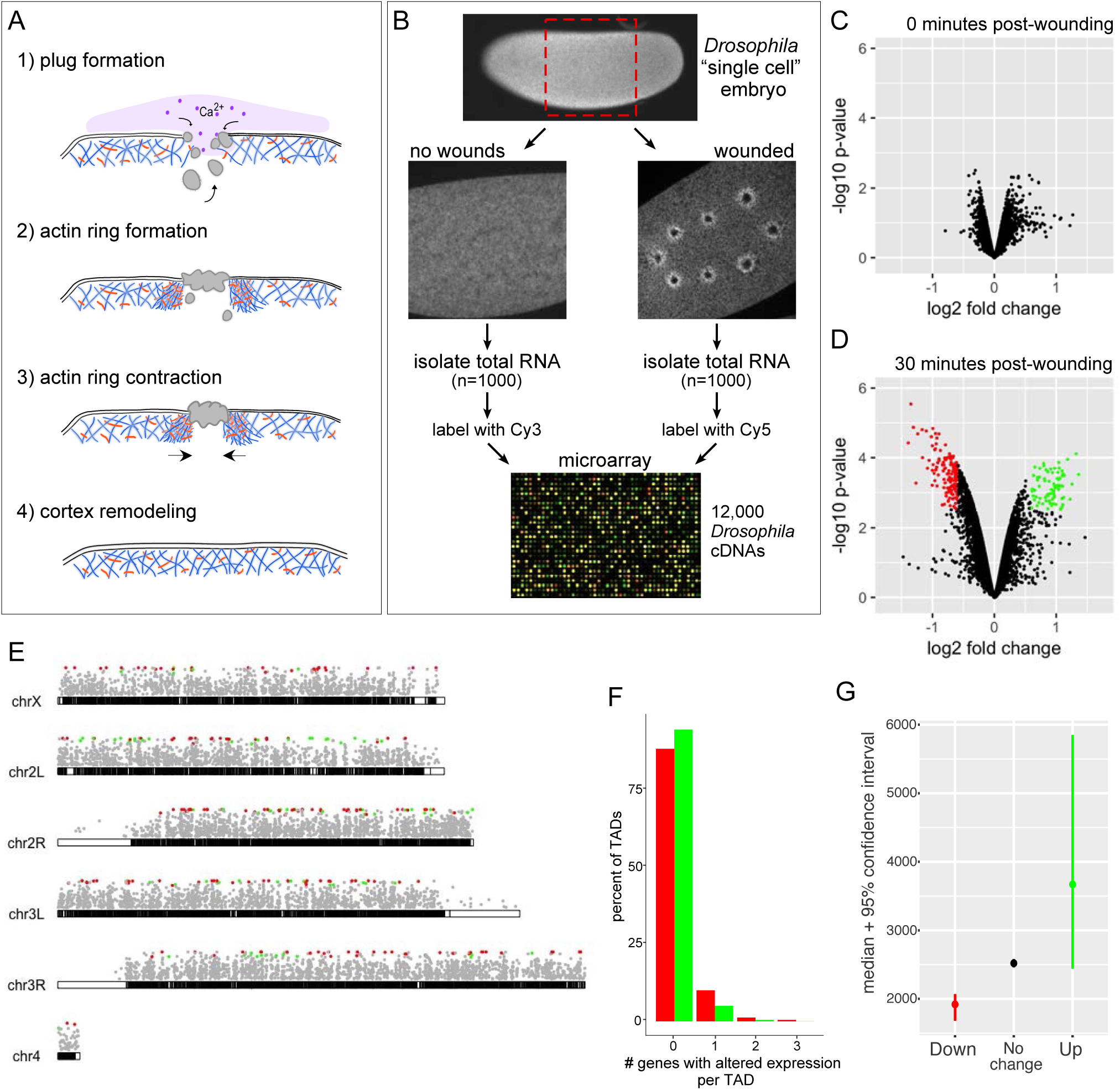
Analysis of differential gene expression following wounding in the *Drosophila* cell wound repair model. (**A**) Schematic of the four major phases of cell wound repair. (**B**) Flow chart depicting the steps involved in microarray processing for examining the transcriptional response to cell wound repair. These analyses were performed for two-timepoints post laser wounding: immediate (0-5 mpw) and near completion (∼30 mpw). (**C-D**) Volcano plots showing the differential gene expression for each of the two timepoints. Each dot represents a cDNA corresponding to its fold-change and p-value. Insignificant hits are depicted in black, whereas up-regulated and down-regulated genes are depicted in green and red, respectively. (**E**) *Drosophila* chromosome maps with each of the 4 chromosomes represented by euchromatic regions in black, heterochromatic regions in white, and with the left and right arms of chromosomes 2 and 3 depicted separately. Dots representing genes hits from the late-period microarray with significant up-regulated genes (green) and down-regulated genes (red) placed at their respective location within the genome. (**F**) Percentage of TADs containing the indicated number of up-or down-regulated genes per TAD. (**G**) Average gene size of significantly expressed genes from the ∼30 min time point microarray.

Previous studies have shown that Ca^2+^ is required for the initiation of cell wound repair and serves as a messenger to trigger downstream processes such as transcription: release of internal and/or external Ca^2+^ stores activates a number of intracellular pathways resulting in an uptick of gene expression [27-30]. Studies carried out in rat embryos and cultured bovine aortic endothelial cells showed a rapid increase in expression of the Ca^2+^-responsive element containing c-Fos protein as a direct result of plasma membrane damage [27, 31, 32]. c-Fos, a component of Activator protein 1 (AP-1), serves as a transcription factor responsible for expressing a number of cytokines and growth factors required to drive the appropriate cellular responses necessary for epithelial (tissue) wound recovery [33-37].

Interestingly, though the *Drosophila* syncytial embryo functions under the developmental control of maternally-contributed mRNAs and proteins with minimal levels of zygotic transcription, it is still able to immediately recognize and repair breaches to its cortex. Here we show that translation, rather than transcription, is required for the initial stages of repair in this cell wound repair model. Although transcription does not serve as a “start” signal, disrupting transcription leads to impaired repair in subsequent steps of the process. Using microarrays to assess gene expression changes post-wounding, we have identified 253 genes with a potential role in cell wound repair, indicated by changes in their expression—either up or down—in response to laser wounding. A subset of these genes were analyzed using RNAi knockdowns to visualize spatio-temporal patterns that verified their involvement. Strikingly, we find that the canonical insulin signaling pathway is required for proper cell wound repair where it controls actin dynamics through the actin regulators Girdin (Hook-like protein family) and Chickadee (profilin). Thus, our study provides insight into the roles of transcription, translation, and insulin signaling in cell wound repair and provides new avenues for understanding how wound healing proceeds in healthy individuals and disease sufferers with wound healing impairments.

## Results

### Assessment of transcriptional contribution to cell wound repair

To investigate the role of transcription in cell wound repair using the *Drosophila* syncytial (nuclear cycle 4-6) embryo model, we performed a microarray screen on full-length cDNA arrays to compare changes of gene expression between laser wounded and non-wounded states at two time points: immediately after wounding (0-5 minutes post-wounding (mpw)) and at the end of the repair process (∼30 mpw) (Fig. 1B). We found that at the immediate timepoint, wounded embryos exhibited no significant changes in their expression profiles when compared to their non-wounded counterparts (Fig. 1C). Interestingly, the later timepoint, which was expected to identify any repair requirements post-initiation, showed significant changes of gene expression in both the up and down directions (Fig. 1D). Using a false discovery rate of 0.05, we identified 253 genes with statistically significant changes: 80 that are up-regulated and 173 that are down-regulated (Table 1; Table S1). The robustness of the differences we observe is striking given that only ∼5-10% of the cell surface is wounded and undergoing repair.

**Table 1.** List of top 16 Up-or 16 Down-regulated genes at t=30 minutes

| FB Gene ID | Name | logFC | P-Val | Molecular/Biological function |
| --- | --- | --- | --- | --- |
| FBgn0001257 | ImpL2 | 1.363 | 0.022 | - Insulin-like growth factor binding; Insulin signaling |
| FBgn0033855 | link | 1.324 | 0.022 | - Unknown; Involved in neurogenesis |
| FBgn0261560 | Thor | 1.257 | 0.026 | - EIF4E binding protein; Insulin signaling |
| FBgn0038071 | Dtg | 1.226 | 0.036 | - Unknown; Involved in gastrulation |
| FBgn0020300 | geko | 1.187 | 0.022 | - Unknown; Involved in olfaction |
| FBgn0263776 | CG43693 | 1.133 | 0.022 | - Amino acid transmembrane transporter |
| FBgn0038028 | CG10035 ** | 1.121 | 0.035 | - Unknown |
| FBgn0003731 | Egfr | 1.112 | 0.022 | - EGF receptor; Involved in growth regulation and development patterning |
| FBgn0000071 | Ama | 1.108 | 0.043 | - Immunoglobulin-like protein domains; Involved in cell adhesion |
| FBgn0086910 | l(3)neo38 | 1.105 | 0.026 | - Zinc finger (C2H2-type); Regulation of transcription/chromatin silencing |
| FBgn0010109 | dpn | 1.100 | 0.023 | - basic Helix-Loop-Helix protein; Transcriptional regulation of sex determination and neurogenesis |
| FBgn0013272 | Gp150 | 1.096 | 0.036 | - Transmembrane glycoprotein; Regulates Notch signaling |
| FBgn0004143 | nullo | 1.095 | 0.022 | - Actin binding; Regulation of epithelial morphogenesis and actomyosin contractile ring assembly |
| FBgn0039283 | danr | 1.094 | 0.046 | - Homeobox domain; Transcriptional regulation of eye development and CNS formation |
| FBgn0028371 | jbug | 1.073 | 0.050 | - Filamin; Involved in cytoskeleton dynamics, PCP pathway, and mechanical stimulus response |
| FBgn0265274 | Inx3 | 1.062 | 0.026 | - Innexin; gap junction protein; Involved in dorsal closure; intercellular transport; and phototransduction |
| FBgn0034259 | P32 | -1.001 | 0.022 | - Mitochondrial protein; Functions in presynaptic calcium signaling and neurotransmitter release; Chromatin metabolism |
| FBgn0033191 | CG1598 | -1.001 | 0.022 | - ATP binding; ATPase activity; Transport to ER |
| FBgn0031600 | CG3652 | -1.004 | 0.026 | - Unknown (Contains Yip1 domain) |
| FBgn0039371 | CG4960 | -1.018 | 0.022 | - Unknown (TB2/DP1/HVA22-related protein); Involved with regulation of intracellular transport |
| FBgn0033906 | ReepB | -1.018 | 0.022 | - Unknown (TB2/DP1/HVA22-related protein); Involved with ER organization and regulation of intracellular transport |
| FBgn0040602 | CG14545 | -1.054 | 0.022 | - Unknown |
| FBgn0034058 | Pex11 | -1.081 | 0.022 | - Unknown; Involved in peroxisome fission and organization |
| FBgn0010078 | RpL23 | -1.099 | 0.022 | - Myosin binding; Structural constituent of ribosome; Involved in translation |
| FBgn0013771 | Cyp6a9 | -1.161 | 0.022 | - Iron/heme binding; Involved in oxidation-reduction processes |
| FBgn0000615 | exu | -1.167 | 0.022 | - Single strand RNA binding; Protein homodimerization; Involved in embryonic pole axis specification, localization of bicoid and oskar mRNA |
| FBgn0086904 | Nacα | -1.238 | 0.022 | - Protein binding; Involved in neurogenesis, oogenesis, oskar mRNA localization |
| FBgn0051075 | CG31075 | -1.266 | 0.025 | - Aldehyde dehydrogenase (NAD) activity; Involved in metabolism and oxidation-reduction processes |
| FBgn0010038 | GstD2 | -0.993 | 0.022 | - Glutathione peroxidase activity, glutathione transferase activity |
| FBgn0001149 | GstD1 | -1.307 | 0.022 | - Glutathione transferase activity; DDT-dehydrochlorinase activity |
| FBgn0011761 | dhd | -1.348 | 0.022 | - Protein disulfide oxidoreductase activity (Thioredoxin domain); Involved in glycerol ether metabolism, cell redox homeostasis, and responses to DNA damage |
| FBgn0033979 | Cyp6a19 | -1.392 | 0.022 | - Cytochrome P450; electron carrier activity and heme binding; Involved in oxidation-reduction process |

\*\* Knockdown (RNAi) lines not available.

We next determined if these genes were being co-differentially expressed by shared activating or regulatory elements within a localized region of the genome in response to wounding. Genome mapping of the 253 genes show no obvious clustering upon visual inspection (Fig. 1E). Concomitantly, we mapped the 253 differentially-expressed genes onto the 1169 unique topologically associated domains (TADs) previously characterized in flies [38], and found no difference in overall differentially-expressed genes between TADs (p=0.22), as well as when comparing just the down-regulated genes (p=0.81) (Fig. 1F). Interestingly, we detected a slight difference in differentially-expressed genes by TAD for up-regulated genes (p=0.01), however the majority of this signal appears to be driven by there being less up-regulated genes and many of these falling into TADs that were missing genes due to their poorer coverage on our arrays. The results from this TAD analysis suggest that the 253 genes are being regulated independently and deliberately in response to wound repair. Intriguingly, the 80 upregulated genes were, on-average, larger than gene products previously recorded during this stage of development (Fig. 1G) [39-42], implicating the existence of a wound-repair specific program (see Discussion).

### Transcription is not required to initiate cell wound repair

We expected that if transcription served as an initiator for wound repair as previously proposed, then inhibition of transcriptional activity would result in altered repair assessable by visualizing actin dynamics throughout the wound repair process. To confirm our microarray results that transcription is unlikely to initiate repair in the *Drosophila* system, we wounded nuclear cycle 4-6 *Drosophila* embryos that were injected with *α*-amanitin, a transcription inhibitor that targets RNA polymerase thereby halting transcritional activity. Using time lapse microscopy and a fluorescent actin reporter, we find that in control embryos, where only buffer was injected, actin became enriched in two distinct locations: 1) adjacent to the wound edge, forming a robust actin ring, and 2) in a “halo” or diffuse accumulation along the outer periphery of the ring and identical to previous findings in uninjected embryos (Fig. 2A-A’, 2E-G; Video 1) [22, 43]. Consistent with our microarray results, *α*-amanitin injected embryos initially showed actin dynamics similar to those observed in control embryos, however they exhibited disruptions to the repair process during the subsequent actin remodeling phases (Fig. 2B-B’, 2E-G; Video 1). To ensure efficient transcriptional knockdown, we verified the efficacy and duration of the *α*-amanitin treatment using the MS2-MCP system, a visual reporter of active transcription (see Methods) (Fig. 2H-K’) [44, 45]. In *Drosophila* syncytial embryos, GFP appears as puncta within the nuclei of control embryos indicative of active transcription, whereas these GFP puncta are absent in *α*-amanitin injected embryos indicating that *α*-amanitin is effectively inhibiting transcription even beyond our initial wounding window (Fig. 2H-I’; Video1). Thus, our results indicate that a transcriptional response is dispensable for the initiation of cell wound repair in the *Drosophila* model, but becomes important subsequently, potentially for replenishing and/or maintaining various factors necessary for establishing the wound repair response.

**Fig 2.**
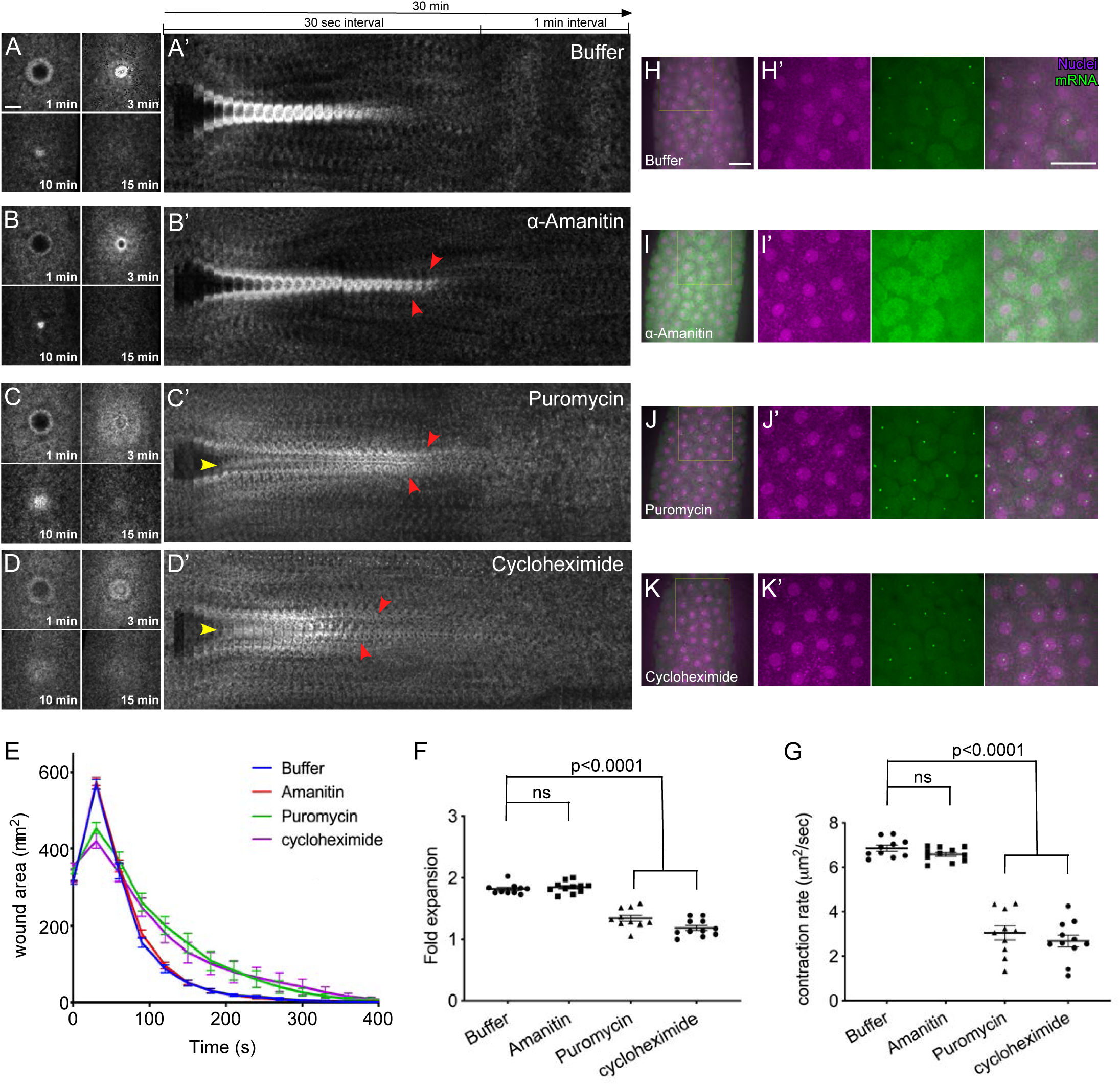
Translation, rather than transcription, is needed for the initiation of cell wound repair. (**A-C**) Confocal projection stills from time-lapse imaging of actin dynamics (sGMCA) during cell wound repair in control (buffer only) (A), alpha-amanitin injected (B), and puromycin injected (C) embryos. (**A’-C’**) XY kymographs across the wound areas depicted in A-C, respectively. Note extended actin remodeling (red arrowheads in b’,c’) and internal actin accumulation (yellow arrowhead in c’). (**D**) Quantification of wound area over time for (A-C’). Error bars represent ± SEM. (**E-F**) Quantification of wound expansion time (E) and wound closure speed (F) for conditions indicated. Student’s t-test; all p-values indicated. (**G-I**) Confocal projections of a NC 10 embryo expressing the MS2-MCP system injected with: buffer (G), alpha-amanitin (H), or puromycin (I). (**G’-I’**) higher magnification images of the respective regions in (G-I) demarcated by the yellow box, showing nuclei (magenta) and nascent mRNA (green). Scale bars: 20 µm.

### The initial steps of cell wound repair are translation dependent

*Drosophila* early embryonic development is mostly driven by maternally deposited mRNA and protein until the maternal-to-zygotic genome transition (MZT) at nuclear cyle 14 (cf. [42]). To explore the role of translation in driving the wound repair process, embryos expressing a fluorescent actin reporter (sGMCA) were injected with the translation inhibitors puromycin (causes premature chain termination) or cycloheximide (blocks translational elongation) prior to laser wound induction (Fig. 2C-D’). While the wound fails to expand, some actin was recruited to the wound periphery, however, the actin ring/halo was not properly assembled and/or maintained resulting in aberrant spatiotemporal enrichment of actin (i.e. inside the wound area) (Fig. 2C-E). Quantitative measurements show a prolonged wound healing process compared to controls (Fig. 2E), with significantly less wound expansion and slower wound closure (Fig. 2F-G). Taken together, our results suggest that the *Drosophila* embryo requires active translation to initiate wound repair, as well as to regulate actin dynamics throughout the repair process.

### Knockdown of differentially expressed genes results in wound over-expansion, abnormal actin dynamics, and remodeling defects upon wounding

We next examined the effects of removing the differentially-expressed genes on cell wound repair. We generated knockdown embryos for 15 of the top 16 up-regulated genes (Fig. 3, Fig. 4, Fig. S1) and the 16 top down-regulated genes (Fig. 4, Fig. 5, Fig. S1Q, Fig. S2) based on their fold-change (Table 1) by expressing RNAi constructs in the female germline using the GAL4-UAS system [46, 47]. We then observed actin dynamics following laser wounding using a fluorescent actin reporter (sGMCA). In all 31 cases, the wounded knockdown embryos exhibited disruptions at various post-initiation steps of the cell wound repair process, including wound over-expansion (Fig. 4A, 4E), delayed/altered rates of wound contraction (Fig. 4B, 4F), aberrant actin dynamics (Fig. 4C-D, 4G-H), and/or remodeling defects (Fig. 3, Fig. 5, Fig. S1, Fig. S2). Examples of these phenotypes are described below.

**Fig 3.**
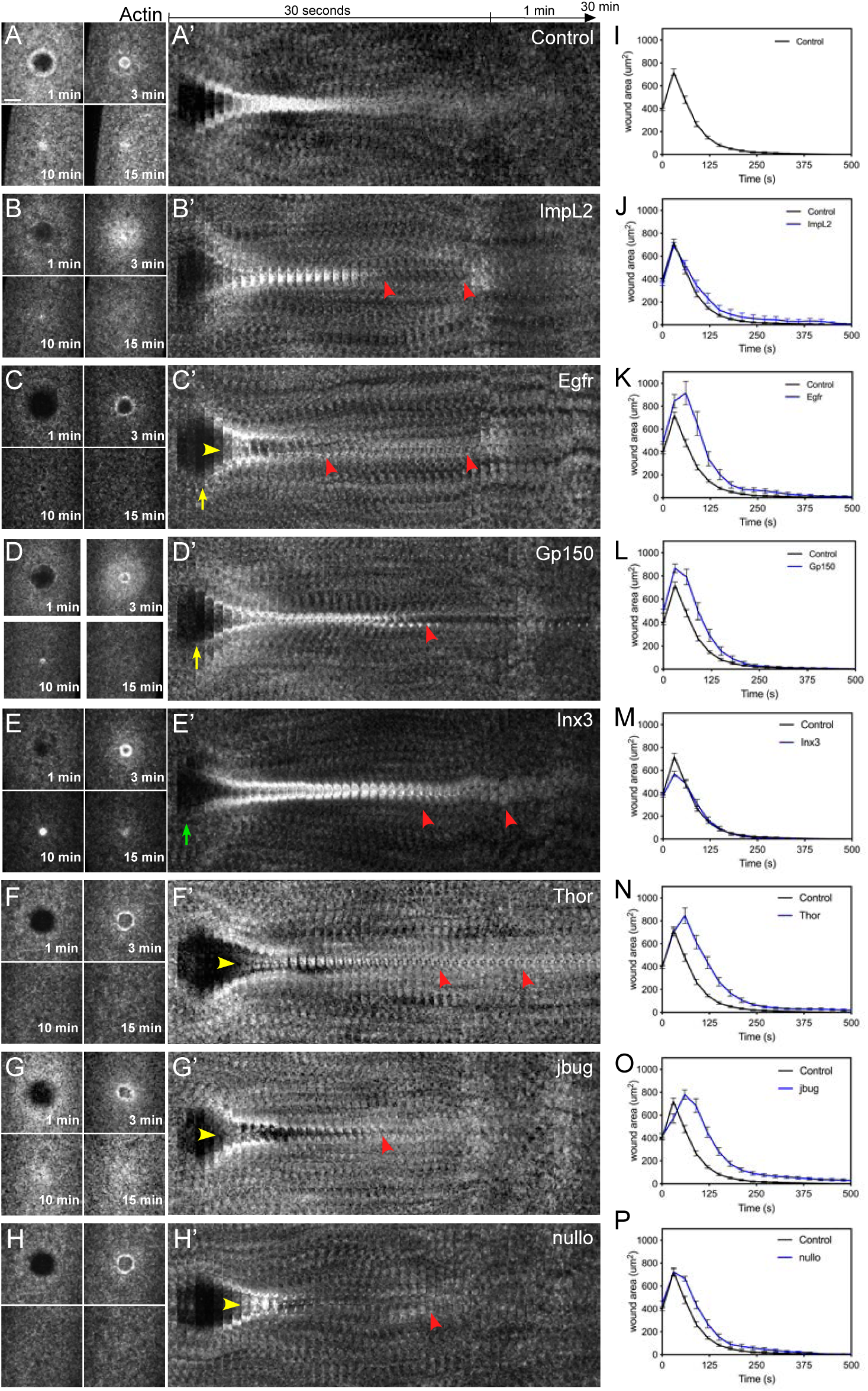
Knockdown of up-regulated genes results in wound over-expansion and abnormal actin dynamics. (**A-H**) Confocal XY projections of actin dynamics at 1, 3, 10, and 15 mpw from *Drosophila* NC4-6 embryos coexpressing sGMCA and a UAS-RNAi transgene during cell wound repair for control (w^1118^/+; sGMCA, 7063/+) (A), ImpL2^RNAi^/+; sGMCA, 7063/+ (B), EGFR^RNAi^/+; sGMCA, 7063/+ (C), Gp150^RNAi^/sGMCA, 7063 (D), Inx3^RNAi^/sGMCA, 7063 (E), Thor^RNAi^/sGMCA, 7063 (F), Jbug^RNAi^/sGMCA, 7063 (G), Nullo^RNAi^/sGMCA, 7063 (H). (A’-H’) XY kymographs across the wound areas depicted in (A-H), respectively. Note wound overexpansion (yellow arrows), wound underexpansion (green arrows), internal actin accumulation (yellow arrowhead), and remodeling defect/open wound (red arrowhead). (**I-P**) Quantification of wound area over time for (A-H’), respectively. Error bars represent ± SEM; n ≥ 10. Scale bars: 20 µm.

**Fig 4.**
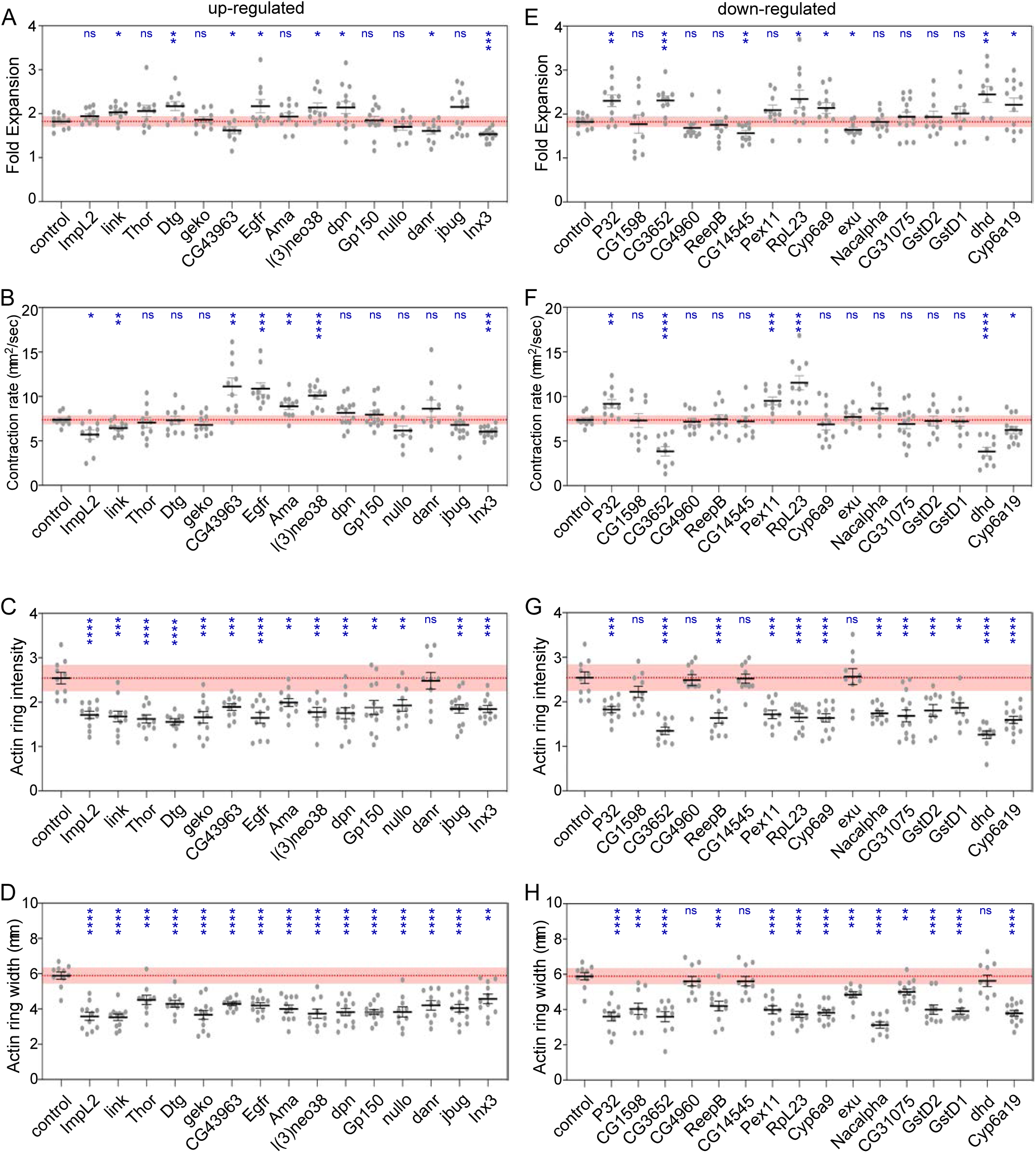
Quantification of wound and actin dynamics in control and knockdowns for upregulated and downregulated genes. (**A-D**) Quantification of wound expansion (A), contraction rate (B), actin ring intensity (C), and actin ring width (D) from control (sGMCA, 7063/+) and knockdowns for all 15 up-regulated genes (RNAi/+; sGMCA, 7063/+ or sGMCA, 7063/RNAi). (**E-H**) Quantification of wound expansion (E), contraction rate (F), actin ring intensity (G), and actin ring width (H) from control (sGMCA, 7063/+) and knockdowns for all 16 down-regulated genes (RNAi/+; sGMCA, 7063/+ or sGMCA, 7063/RNAi). Black line and error bars represent mean ± SEM. Red line and square represent mean ± 95% CI from control. n ≥ 10. Student’s t-test is performed to compare control with knockdowns. * is p<0.05, ** is p<0.01, *** is p<0.001, **** is p<0.0001, and ns is not significant.

**Fig 5.**
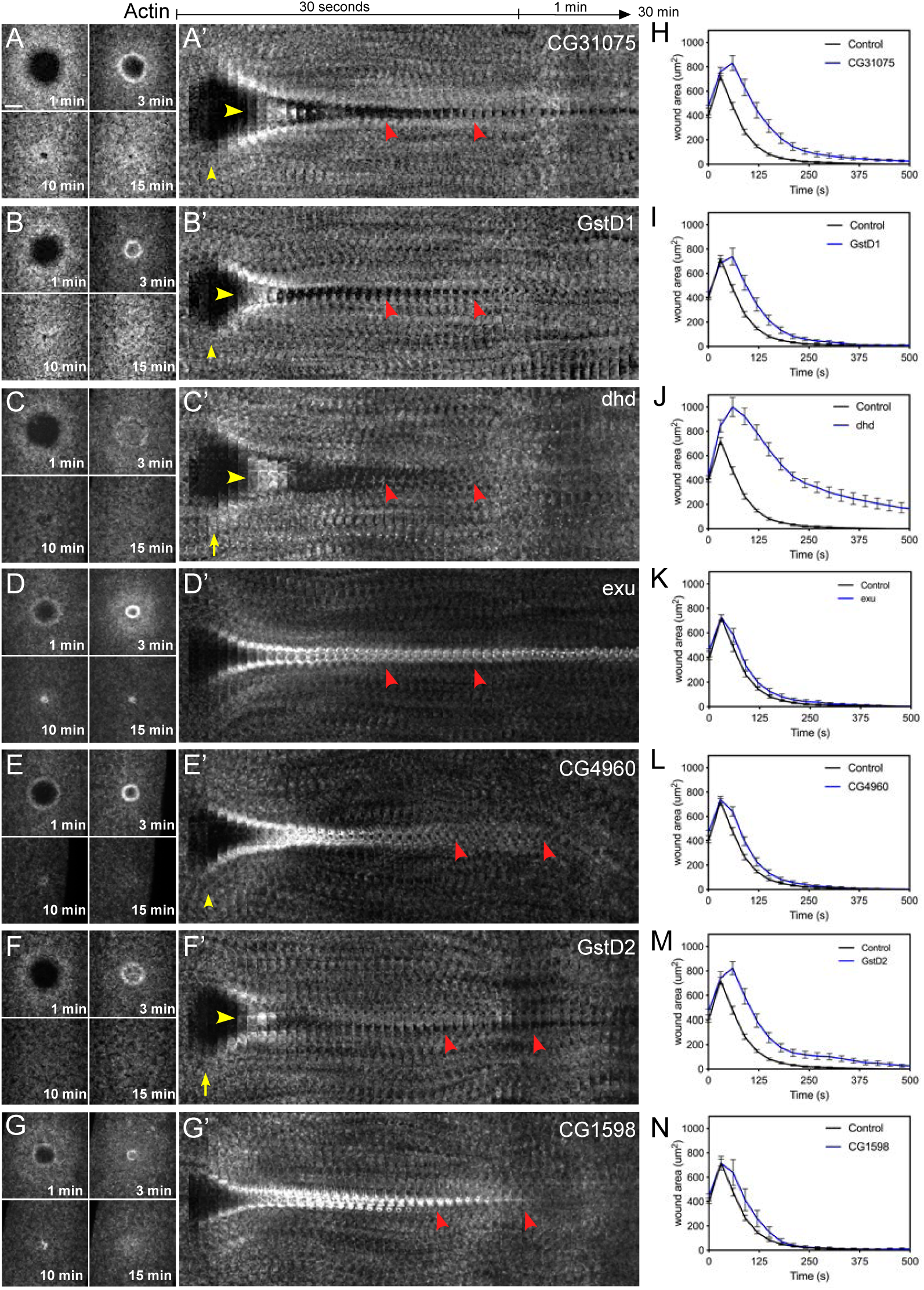
Knockdown of down-regulated genes results in wound over-expansion and abnormal actin dynamics. (**A-G**) Confocal XY projections of actin dynamics at 1, 3, 10, and 15 mpw from *Drosophila* NC4-6 embryos coexpressing sGMCA and a UAS-RNAi transgene during cell wound repair for CG31075^RNAi^/+; sGMCA, 7063/+ (A), GstD1^RNAi^/+; sGMCA, 7063/+ (B), dhd^RNAi^/+; sGMCA, 7063/+ (C), Exu^RNAi^/+; sGMCA, 7063/+ (D), CG4960^RNAi^/+; sGMCA, 7063/+ (E), GstD2^RNAi^/+; sGMCA, 7063/+ (F), CG1598^RNAi^/+; sGMCA, 7063/+ (G). (**A’-G’**) XY kymographs across the wound areas depicted in (A-G), respectively. Note wound overexpansion (yellow arrows), wound underexpansion (green arrows), internal actin accumulation (yellow arrowhead), and remodeling defect/open wound (red arrowhead). (**H-N**) Quantification of wound area over time for (A-G’), respectively. Error bars represent ± SEM; n ≥ 10. Scale bars: 20 µm.

#### Up-regulated Genes

The *Drosophila* embryo is under tension such that when it is wounded, the plasma membrane and cortical actin cytoskeleton recoil slightly leading to an expansion of the wound [22, 48]. Interestingly, wounds generated in knockdowns of three of the up-regulated genes (*Inx3, CG43963, danr*) failed to expand, whereas others (*Dtg, link, l(3)neo38, Egfr, dpn*) exhibited wound over-expansion (Fig. 3, Fig. 4A, Fig. S1). Similarly, wounds generated in knockdowns of three of the up-regulated genes (*Inx3, ImpL2, link*) exhibited slower wound contraction rates, whereas others (*l(3)neo38, CG43963, Egfr, Ama*) exhibited faster wound contraction rates compared to control wounds (Fig. 3, Fig. 4B, Fig. S1).

In all 15 cases of RNAi knockdown for up-regulated genes, wounded embryos exhibited abnormal actin dynamics, including premature actin ring/halo disassembly, failure of actin ring/halo dissassembly, and/or abnormal actin ring/halo disassembly with concomitant accumulation of actin within the wound. (Fig. 3, Fig. 4C-D; Video 2; Fig. S1). Wounds generated in knockdowns of *Imaginal morphogenesis protein-Late 2* (*ImpL2*) and *Epidermial growth factor receptor* (*Egfr*) are exemplified by their incomplete formation and premature dissassembly of the actomyosin ring causing rifts at the initial injury site that remained open for the entire time of repair (Fig. 3B-C’, 3J-K; Video 2). ImpL2 has been proposed to work antagonisitically to the insulin/insulin-like (IIS) signaling pathway by interacting with receptor/ligand interactions to inhibit downstream signal transduction [49-51]. Egfr encodes a receptor tyrosine kinase that works upstream of the c-jun N-terminal kinase (JNK) and decapentaplegic (dpp) pathways. Loss of Egfr results in down-regulation of JNK activity leading to the impairment of dorsal closure, a process sharing many features with epithelial (multicellular) wound repair [52]. Wounds generated in knockdowns of *jitterbug* (*jbug*) and *nullo*, are characteristically defined by the pronounced formation of actin inside the wound area (Fig. 3G-H’, 3O-P; Video 3). Jbug is a filamin-type protein that serves as an F-actin crosslinker providing stability to the cytoskeleton, a system that has been proposed to utilize mechanical cues such as tension to modulate cellular processes [53, 54]. Nullo has been shown to establish cortical compartments during cellularization of the *Drosophila* embryo, suggesting an important role regulating actin stability at the cortex [55, 56].

Following wound closure, extensive remodeling of the cortical cytoskeleton and its overlying plasma membrane is necessary to re-establish normal architectures and activities. Wounds generated in knockdowns of Gp150, Inx3, and Thor, are unable to resolve actin structures and/or properly remodel cortical actin after wound closure (Fig. 3D-F’, 3L-N; Video 2). Gp150 encodes a transmembrane glycoprotein that regulates Notch signaling during normal eye development in *Drosophila* [57], whereas Inx3 encodes a gap junction protein involved in morphogenesis and nervous system development [58, 59]. Thor encodes a translation inhibitor functioning downstream of insulin signaling that is sensitive to reactive oxygen species [60]. Interestingly, like ImpL2, Thor is a IIS pathway constituent and Gp150 has also been shown to physically interact with components of this pathway (Pten and S6k) [61].

#### Down-regulated Genes

Interestingly, in all 16 cases of RNAi knockdown for the down-regulated genes examined, wounded embryos exhibited abnormal cell wound repair dynamics that included the same major, but non-mutually exclusive, steps as described above for the up-regulated genes. A number of the genes that were downregulated have an unknown molecular function and/or associated biological processes (Table 1; Fig. 4E-H, Fig. 5; Video 3; Fig. S1Q, Fig. S2). Of these unknown genes, CG31075 underwent a mild expansion followed by a contraction rate similar to that in wildtype, albeit with incomplete wound closure (Fig. 4E, Fig. 5A-A’, 5H; Video 3), CG4960 exhibited a slight delay in wound repair dynamics but retained noticeably enriched actin structures after closure (Fig. 5E-E’, 5L; Video 3), and CG1598 developed a visually distinct, but transient, enrichment of actin inside the wound area prior to closure (Fig. 5G-G’, 5N; Video 3). Of genes with known motifs/functions, Glutatione S transferases D2 (GstD2) and D1 (GstD1) RNAi knockdowns showed similar phenotypes exhibiting a short-lived accumulation of actin inside the wound area and delayed closure dynamics during the initial steps of repair (Fig. 5B-B’, 5F-F’, 5I, 5M; Video 3), and in later steps, both are unable to completely close (Figure 5B-B’, 5I; Video 3). Wound repair begins normally in *exu* knockdowns, however the leading edge and surrounding actin structures soon become static resulting in an open wound area and prolonged actin accumulation (Fig. 5D-D’, 5K; Video 3). In addition to the phenotypes described above, many of these knockdowns exhibit wound over-expansion (*CG3652, P32, dhd, RpL23, Cyp6a9, Cyp6a19*) (Fig. 4E, Fig. 5; Videos 3, 4; Fig. S2) and nearly all exhibit remodeling defects (Fig. 5, red arrowheads; Videos 3, 4; Fig. S2, red arrowheads). Thus, in all 31 cases of up-or down-regulated genes examined, knockdown using RNAi transgenes resulted in abnormal cell wound repair. Despite the molecular functions of many of these genes being unknown, they have been implicated in various cellular processes, but most notably a subset are involved in insulin signaling.

### Activation of insulin/insulin-like (IIS) constituents during normal wound repair

The fact that ImpL2 and Thor, two of the most upregulated genes in our analyses, are constituents of the insulin/insulin-like growth factor signaling (IIS) pathway in *Drosophila* (Fig. 6A) was somewhat unexpected. Deficiencies in insulin signaling have been implicated in multicellular (tissue/epithelial) repair, where it is thought to impede growth factor production, angiogenic response, and epidermal barrier function [62-65], functions that might not normally be expected to govern regulation within individual cells.

**Fig 6.**
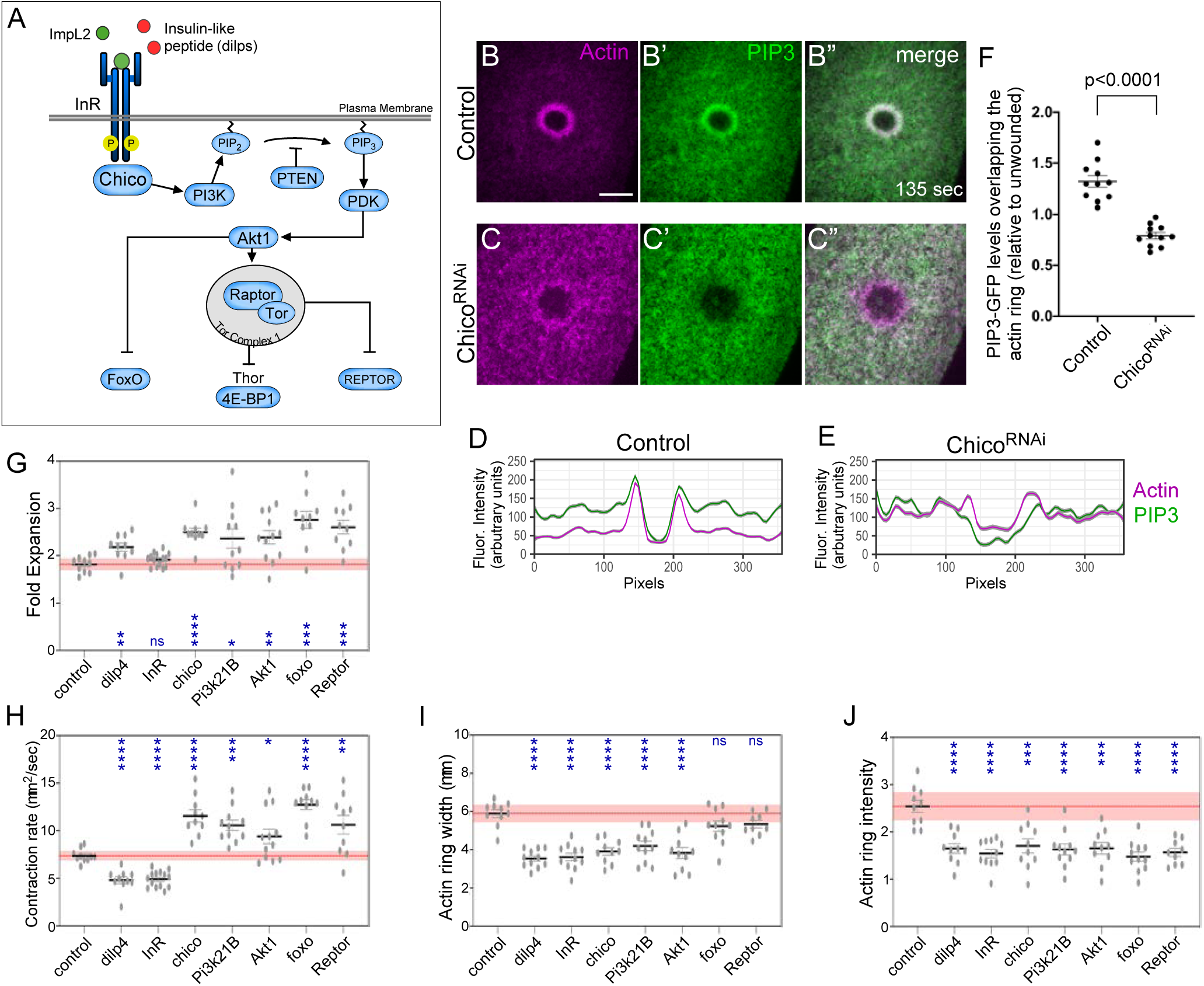
Localization of IIS pathway components. (**A**) Simplified diagram of the IIS pathway in *Drosophila* showing the components tested using GFP reporters and RNAi transgenes. (**B-C”**) Confocal xy projection images from *Drosophila* NC4-6 staged embryos co-expressing an actin marker (sChMCA) and GFP-tagged PIP3 in a control (B-B”) or chico RNAi (C-C’’). (**D-E**) Smoothened fluorescence intensity (arbitrary units) profiles derived from averaged fluorescence intensity values over a 10 pixel width across the wound area in the embryo shown (B-C”), respectively. Gray area represents the 95% CI. Scale bars: 20 µm. (**G-J**) Quantification of wound expansion (G), contraction rate (H), actin ring intensity (I), and actin ring width (J) from control (sGMCA, 7063/+) and knockdowns for IIS pathway genes (RNAi/+; sGMCA, 7063/+ or sGMCA, 7063/RNAi). Black line and error bars represent mean ± SEM. Red line and square represent mean ± 95% CI from control. n ≥ 10. Student’s t-test is performed to compare control with knockdowns. * is p<0.05, ** is p<0.01, *** is p<0.001, **** is p<0.0001, and ns is not significant.

To determine if the canonical IIS pathway was involved in individual cell wound repair, we first examined the recruitment pattern of a PIP_3_ (phosphatidylinositol (3,4,5)-triphosphate)-GFP reporter construct used as a reporter of insulin signaling activity [66], co-expressed with a Cherry fluorescently-tagged actin reporter (sChMCA) in a wildtype and *chico* RNAi knockdown background (Fig. 6B-F). PIP_3_ is a phospholipid that composes a subset of specialized plasma membrane with various trafficking and signaling related functions [67]. PIP_3_-GFP is recruited to same region as the actomyosin ring in wildtype embryos (Fig. 6B-B”, 6D, 6F), confirming the requirement for autocrine insulin pathway signaling. Importantly, this recruitment is dependent on the upstream activation of the insulin receptor (InR), as PIP_3_-GFP recruitment is disrupted in a *chico* RNAi background (Fig. 6C-C”, 6E-F).

We next examined the wound repair phenotypes in knockdown backgrounds for components spanning the IIS pathway by expressing RNAi constructs for pathway components in the female germline using the GAL4-UAS system [46, 47], then observing actin dynamics using a fluorescent actin reporter (sGMCA). The one ligand and six of the major IIS pathway components tested — Ilp4 (Insulin-like peptide), InR (Insulin receptor), Chico (IRS homolog), Pi3K21B (Phosphoinositide3-Kinase), Akt1 (Kinase), FoxO (transcription factor), and Reptor (transcription factor) — exhibited abherrant wound repair with overlapping phenotypes reflecting involvement at several steps in the repair process (Fig. 6A, 6G-J, Fig. 7; Video 4; Fig. S1Q). With the exception of ImpL2, Ilp4, and InR (components at the top of the pathway), mutants for IIS pathway components exhibited wound overexpansion immediately after laser ablation that was visible as the outward retraction of the wound edge (Fig. 6G-H, Fig.7; Video 4). Following this overexpansion, actin structures became transiently enriched inside the wound area, but dissassembled prior to complete wound closure (Fig. 6I-J, Fig. 7; Video 4). Lastly, progression of wound closure was signficantly delayed and/or incomplete, leaving openings around the actin ring as it translocated (Fig. 7, red arrowheads; Video 4). While we can not rule out contributions from non-canonical insulin signaling pathways, our results show that key components of the canonical insulin signaling pathway are not only called to a wound, but have detrimental effects on actin and wound dynamics upon knockdown. Collectively, our results suggesting that there exists a tight association between the factors that regulate both insulin signaling and cell wound repair in the *Drosophila* model.

**Fig 7.**
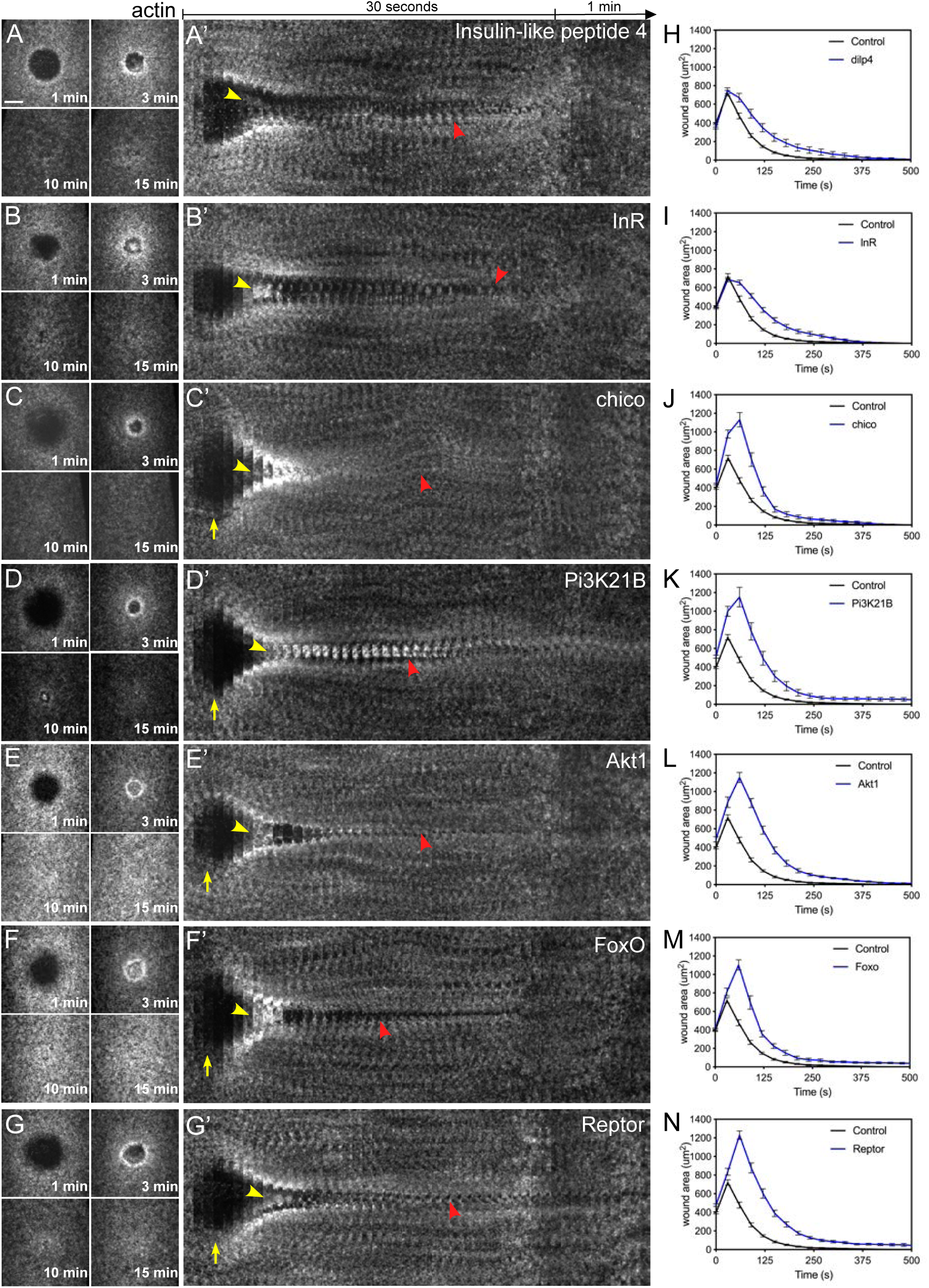
Actin dynamics of insulin/insulin-like (IIS) pathway mutants. (**A-G**) Confocal XY projections of actin dynamics at 1, 3, 10, and 15 mpw during cell wound repair in *Drosophila* NC4-6 embryos expressing sGMCA and a mutant for insulin-like peptide 4 (*Ilp4*^*1*^; A), or a UAS-RNAi transgene for InR^RNAi(1)^/+; InR^RNAi(2)^/sGMCA, 7063 (B), Chico^RNAi^/sGMCA, 7063 (C), Pi3K21B^RNAi^/sGMCA, 7063 (D), Akt1^RNAi^/sGMCA, 7063 (E), FoxO^RNAi^/sGMCA, 7063 (F), and Reptor^RNAi^/sGMCA, 7063 (G). (**A’-G’**) XY kymographs across the wound areas depicted in (A-G), respectively. Note wound overexpansion (yellow arrows), wound underexpansion (green arrows), internal actin accumulation (yellow arrowhead), and remodeling defect/open wound (red arrowhead). (**H-N**) Quantification of wound area over time for (A-G’), respectively. Error bars represent ± SEM; n ≥ 10. Scale bars: 20 µm.

### The IIS pathway effectors Profilin (Chickadee) and Girdin are required for cell wound repair

The IIS pathway has recently been shown to control actin dynamics independently of its role in growth control [68]. In particular, the IIS pathway has been found to activate the expression of the *Drosophila* profilin homolog (*Chickadee*), as well as the Akt substrate Girdin (GIRDers of actIN; also known as GIV) [68-70]. To determine if these actin regulators function as IIS pathway effectors during cell wound repair, we stained wounded embryos that expressed a GFP-tagged actin reporter (sGMCA) in a wildtype or *chico* RNAi knockdown background with antibodies to Profilin/Chickadee and Girdin (Fig. 8A-D). Both proteins are recruited to wounds, although their spatial recruitment patterns are not the same. Girdin exhibits a punctate recuitment at wounds with the highest accumulation overlapping the membrane plug inside the actin ring and with lower level diffuse accumulation overlapping the actin ring and the innermost part of the actin halo (Fig. 8A-B). Profilin/Chickadee recruitment is internal to the actin ring and appears to be excluded from the actin ring region (Fig. 8A-B). Importantly, the accumulation of both Profilin/Chickadee and Girdin at wounds requires a functioning IIS pathway as these accumulations are lost in a *chico* RNAi background (Fig. 8C-D).

**Fig 8.**
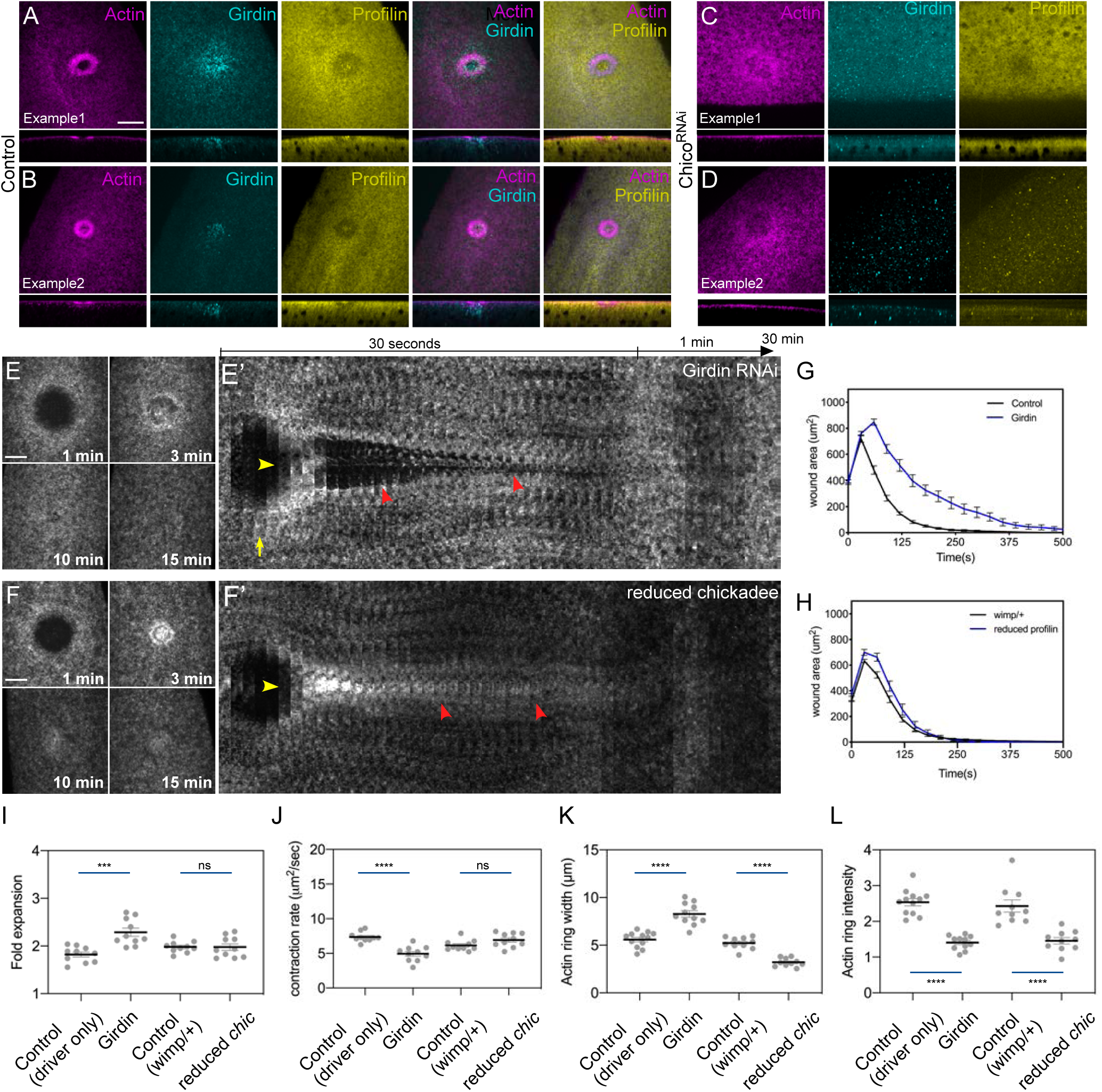
Chickadee (profilin) and Girdin are insulin/insulin-like (IIS) pathway effectors during cell wound repair. (**A-D**) Confocal XY projections of laser wounded Drosophila NC4-6 wildtype (A-B) or *chico* RNAi knockdown (C-D) embryos stained for Girdin (Girdin), Chickadee/profilin (Profilin), and F-actin/phalloidin (Actin). (**E-F’**) Confocal XY projections of actin dynamics at 1, 3, 10, and 15 mpw from *Drosophila* NC4-6 embryos coexpressing sGMCA and a UAS-RNAi transgene during cell wound repair for Girdin^RNAi^ (Girdin^RNAi^/+; sGMCA, 7063/+) (E) and reduced chickadee (sGMCA; *chickadee*^*221*^/+ sGMCA, *wimp*/+) (F). (E’-F’) XY kymographs across the wound areas depicted in E-F, respectively. Note wound overexpansion (yellow arrows), wound underexpansion (green arrows), internal actin accumulation (yellow arrowhead), and remodeling defect/open wound (red arrowhead). (**G-H**) Quantification of wound area over time for (E-F’), respectively. Error bars represent ± SEM; n ≥ 10. (**I-L**) Quantification of wound expansion (G), contraction rate (H), actin ring intensity (I), and actin ring width (J) from control (sGMCA, 7063/+) and knockdowns for IIS pathway genes (RNAi/+; sGMCA, 7063/+ or sGMCA, 7063/RNAi). Error bars represent ± SEM; n ≥ 10. Student’s t-test is performed to compare control with knockdowns. *** is p<0.001, **** is p<0.0001, and ns is not significant. Scale bars: 20 µm.

We next examined the effects of removing Girdin and Profilin/Chickadee on cell wound repair. Similar to knockdown of IIS pathway components described above, Girdin RNAi knockdown embryos exhibited aberrant wound repair including wound overexpansion, enrichment of actin structures inside the wound area, and signficantly delayed wound closure (Fig. 8E-E’, 8G, 8I-L; Video 4; Fig. S1Q). Unfortunately, Profilin/Chickadee RNAi knockdown females do not produce eggs. We therefore used the *wimp* mutation [71, 72] to generate reduced Profilin/Chickadee expression in both the germline and soma (*wimp* reduces maternal gene expression such that, when *in trans* to the *chickadee*^*221*^ allele, it effectively generates a strong *chickadee* hypomorph, referred to as reduced Profilin). Similar to knockdown of Girdin and IIS pathway components, reduced Profilin/Chickadee embryos exhibited wound overexpansion, enrichment of actin structures inside the wound area, and signficantly delayed wound closure (Fig. 8F-F’, 8H, 8I-L; Video 4; Fig. S1Q). Thus, our results indicate that Girdin and Profilin/Chickadee are actin regulatory downstream effectors of the IIS pathway in cell wound repair.

## Discussion

Our study shows that cellular wound repair is not dependent on transcriptional activity to initiate wound repair programs, that dormant transcription pathways are activated in response to wounds, and that the insulin signaling pathway is an essential component of the repair process. A calcium influx-triggered transcriptional response was thought to be important to lead off the cell wound repair process, eliciting a downstream wound repair program. However, this proposed mechanism was at odds with the *Drosophila* syncytial embryo cell wound model that faithfully recapitulates the majority of features associated with other single cell wound repair models (*Xenopus* oocytes, tissue culture cells, sea urchin eggs) [2, 3, 17, 22, 26, 43, 73-75], yet represents a special system running mostly off of maternally contributed products, highlighted by rapid cell cycles (∼10 minutes/cycle) and minimal zygotic transcription [41, 42, 76].

Consistent with the closed nature of the *Drosophila* syncytial embryo cell wound model, we find no altered gene expression immediately upon wounding either as assayed by microarray analysis of laser wounded versus unwounded embryos or following injection of the *α*-amanitin transcriptional inhibitor. We do detect alterations in gene expression at subsequent stages in the repair process: we identified 253 genes (out of ∼8000 genes assayed) whose expression is significantly up (80 genes) or significantly down (173 genes) following laser wounding.

Polymerase rates in the early *Drosophila* embryo were reported to be 1.1-1.5 kb/min, leading to the suggestion that any genes transcribed in the early *Drosophila* embryo prior to the mid-blastula transition must be small with minimal introns due to the rapid (∼10 min) cell cycles and limited transcription time [41, 42, 77-82]. Recent studies have revised this rate to 2.4-3.0 kb/min, lowering the size constraints on the zygotic genes that can be successfully transcribed prior to the mid-blastula transition[40]. Therefore, genes up to ∼20-25 kb could theoretically be transcribed during the early and rapid *Drosophila* embryo cell cycles. In this case however, the number of mRNA molecules would be likely limited by the lower number of nuclei present and thus copies of DNA.

We find that the average size of transcripts in syncytial *Drosophila* embryos is 2.5 kb, similar to the previously reported size of 2.2 kb (compared to the overall average length of coding genes in *Drosophila* of 6.1 kb) [39, 83]. Genes whose expression goes down during wound repair are, on average, 1.9 kb. It is intriguing that these actively down-regulated genes negatively impact the wound repair process when knocked-down. These genes likely represent RNAs stored in the embryo that are used up during the repair process and not replaced. Alternatively, it is possible that wound repair itself may slightly delay development leading to a subset of zygotically expressed genes whose expression is lagging behind in wounded versus unwounded embryos such that this delayed developmental upregulation is read out as a down-regulation of genes.

Surprisingly, we find that genes whose expression is higher after wounding are much larger on average (3.7 kb) than the average sized transcript at that stage (2.5 kb). These genes likely encode cellular components that were expended during the repair process and are being replenished for normal developmental events to proceed, or that are activated specifically for the repair process. This subset of “up-regulated” genes includes genes that are not usually expressed in the early embryo (e.g., CG43693). Thus, our results suggest that, when wounded, the embryo may be able to activate a transcriptional program that is usually dormant during these stages.

Interestingly, 2 of the top 3 genes whose expression is significantly higher following wounding—ImpL2 and Thor—are components of the Insulin signaling pathway. While it has been shown that defective insulin signaling impairs epithelial (multicellular) wound repair [62-65], this result was less expected for wound repair within single cells. Using a combination of RNAi knockdowns and GFP reporters, we have shown that all major components of the IIS pathway are involved in cellular wound repair, and upon knockdown, display similar phenotypes, suggesting that in this context the canonical IIS pathway activation occurs in an autocrine-like manner. Previous studies have highlighted the necessity of calcium influx to facilitate vesicle exocytosis and subsequent fusion of the plasma membrane during wound repair [17, 18, 73, 74]. Similarly, this influx has also been shown to modulate insulin secretion in β-islet cells via the opening of L-type channels by establishing calcium microdomains along the cortex [84, 85]. Insulin/insulin-like peptides are secreted into the extracellular space where they bind to InR thereby activating the heavily conserved IIS pathway that is known to regulate a number of downstream processes that range from transcription via phosphorylation events on the FOXO family of transcription factors to translation via the regulation of the 4E-binding protein, Thor [64, 86-89]. Recently emerging evidence has also shown that the activated IIS pathway can control actin dynamics through activation of actin regulators including Chickadee (profilin) and Girdin [68-70, 90, 91].

Observation of actin dynamics in mutants for a number of the IIS pathway components show common phenotypes of impaired cytoskeleton dynamics, most notably an immediate over-expansion of the wound leading edge and a transient actin structure forming inside the wound area suggesting that normal wound repair processes are heavily reliant on a functioning IIS pathway. We propose that the initial inrush of calcium generates microdomains that trigger the secretion of the *Drosophila* insulin-like peptide 4 (Ilp4) into the perivitelline space where it recognizes and binds to the extracellular face of the Insulin receptor (Fig. 9, steps 1-4). Subsequently, the InR is activated and initiates a signaling cascade that regulates a number of downstream processes, including cytoskeletal dynamics (Fig. 9, steps 5-7). Chickadee/profilin binds to actin and affects the formation/remodeling of actin-rich structures [68]. Girdin also binds to actin, as well as the catenin-cadherin complex and the Exo-70 subunit of the exocyst complex, where it has been proposed to coordinate cytoskeleton organization, cell adhesion, membrane trafficking events, and serves as an indicator for poor prognosis with invasive breast cancers [69, 70, 90, 92, 93]. Interestingly, *girdin* and Profilin knockdown embryos exhibit wound repair phenotypes consistent with defects in actin structure assembly/remodeling, actomyosin ring attachment to the overlying plasma membrane, and membrane trafficking. In addition to the genes involved in the IIS pathway, our microarray analyses identified numerous other genes that show phenotypes associated with actin dynamics regulation. For example, Nullo is a known regulator of actin-myosin stability and has been proposed to affect actin-actin and actin-membrane interactions at the cortex, suggesting a role in cortical remodeling during actomyosin ring contraction [55, 56].

**Fig 9.**
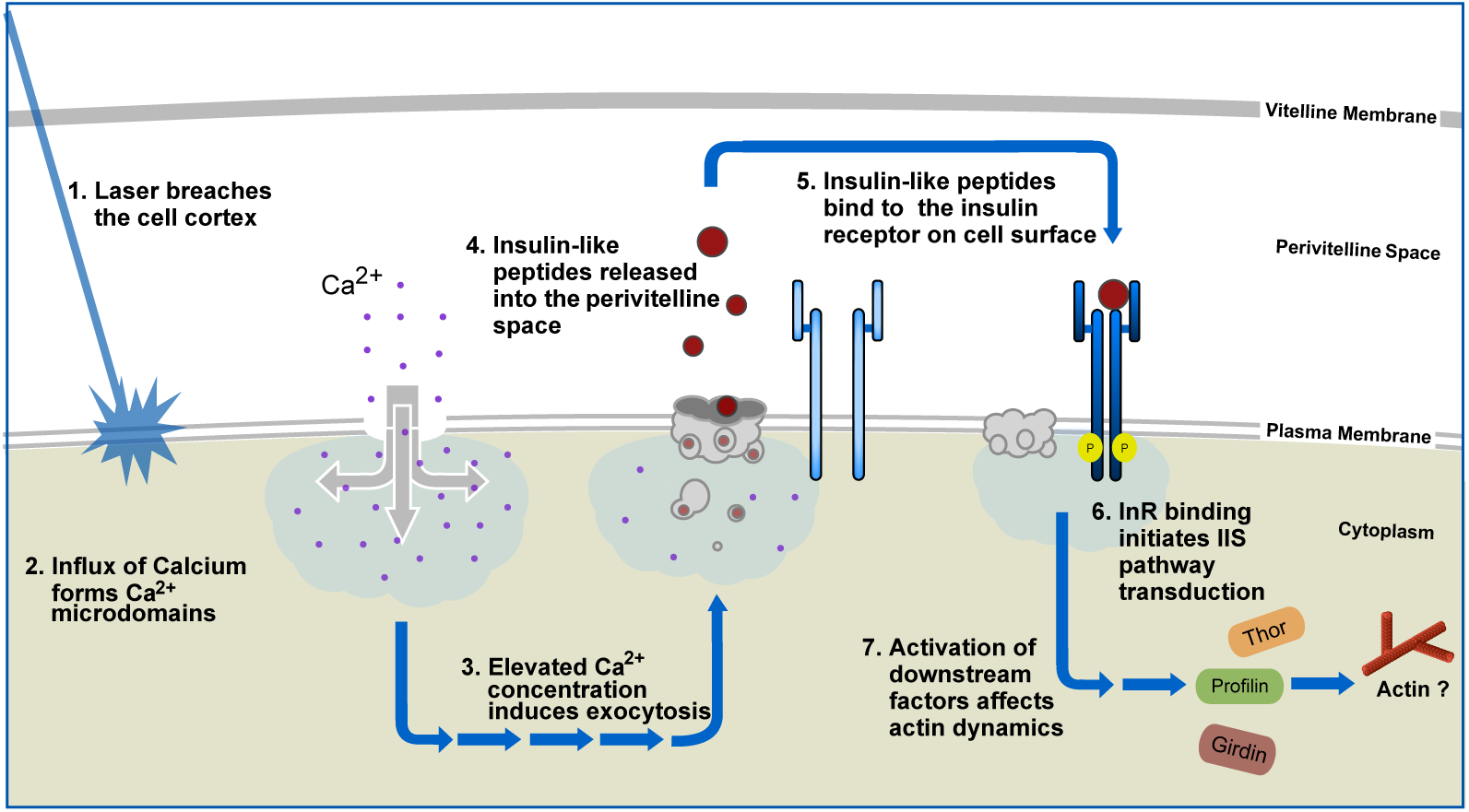
Model for insulin/insulin-like (IIS) pathway function in cell wound repair. Wounding of the cell cortex leads an rush of calcium into the cytoplasm developing locales of increased calcium concentration known as microdomains. To plug the hole, elevated calcium levels initiate exocytotic programs to recruit vescles to the wound area, forming a plug while simultaneously releasing insulin-like peptide 4 (ilp4) into the perivitelline space. Ilp4 is recognized by the insulin receptor (InR) where it binds and activates the IIS pathway. Subsequent phosphorylation events downstream of InR, ultimately activate downstream effectors including the actin remodelers Chickadee and Girdin to repair and restore the cortex back to its normal state.

In summary, our understanding of the mechanisms that trigger cell wound repair remain incomplete, but here we show functional translation is essential for initiating a normal and processive wound repair process, suggesting that the first responders are likely mRNA and protein already present in the cell. While transcription is not immediately necessary in the *Drosophila* cell wound model, it is needed for the repair process. The requirement for insulin signaling in the single cell wound repair context highlights the conservation of repair mechanisms employed. Given its prominence in the single cell, as well as multicellular (tissue), repair pathways, it is not surprising that impaired insulin signaling leads to major wound repair defects in diseases such as diabetes where chronic wounds are symptomatically observed. As many of the top up-and down-regulated genes we identified are evolutionarily conserved genes, but of currently unknown function, the challenge for the future is to determine their roles in normal cellular maintenance and/or development, in addition to their effects in a cell wound repair context, thereby allowing the establishment of a network of cellular processes involved to better aid in treatments of disease involving wound healing impairments, or in disciplines such as regenerative medicine.

## Materials and Methods

### Fly stocks and genetics

Flies were cultured and crossed at 25°C on yeast-cornmeal-molasses-malt extract medium. The flies used in this study are listed in Table S2A. RNAi lines were driven using the GAL4-UAS system using the maternally expressed driver, Pmatalpha-GAL-VP16V37. All genetic fly crosses were performed at least twice. All RNAi experiments were performed at least twice from independent genetic crosses and ≥10 embryos were examined unless otherwise noted.

An actin reporter, sGMCA (spaghetti squash driven, moesin-alpha-helical-coiled and actin binding site bound to GFP reporter) [48] or the Cherry fluorescent equivalent, sChMCA [22], was used to follow wound repair dynamics of the cortical cytoskeleton.

*wimp* (*RpL140*^*wimp*^) reduces maternal gene expression of a specific subset of genes in the early *Drosophila* embryo [71, 72, 94]. Reduced *chickadee* embryos were obtained from trans-heterozygous females generated by crossing *chickadee*^*221*^ to *RpL140*^*wimp*^.

We attempted InR knockdown in three ways: 1) expressing one shRNA (GL00139) using one maternal-GAL4 driver (BDSC #7063), 2) expressing two shRNAs (HMS03166 and GL00139) using one maternal-GAL4 driver (BDSC #70637063), and 3) expressing one shRNA (GL00139) using one maternal-GAL4 (BDSC #7063) in an *InR*^*05545*^ heterozygous mutant backgrounds. We achieved only 50% knockdown with approach (1), and no eggs were produced by approach (3). We achieved 87% knockdown with approach (2) and this condition was used for the phenotypic analyses included here.

For the MS2-MCP system [44, 45], female virgins maternally expressing MCP-GFP and Histone-RFP were crossed with males expressing 24xMS2 stem loops and lacZ driven by hunchback P2 enhancer and promoter. F1 embryos (MCP-GFP, Histone-RFP/+; 24xMS2-lacZ/+) at NC9-10 stages were used for imaging where the 24xMS2-lacZ mRNA is contributed zygotically. Localization patterns and mutant analyses were performed at least twice from independent genetic crosses and ≥10 embryos were examined unless otherwise noted. Images representing the average phenotype were selected for figures.

### Quantification of mRNA levels in RNAi mutants

To harvest total RNA, 100-150 embryos were collected after a 30 min incubation at 25°C, treated with TRIzol (Invitrogen/Thermo Fisher Scientific) and then with DNase I (Sigma). 1 µg of total RNA and oligo (dT) primers were reverse transcribed using the iScript gDNA Clear cDNA Synthesis Kit (Bio-Rad). RT-PCR was performed using the iTaq Universal SYBR Green Supermix (Bio-Rad) and primers obtained from the Fly Primer Bank listed on Table S2B. We were unable to identify primer sets that would work for qPCR for Geko, Ama, l(3)neo38, danr, and CG4960.

Each gene in question was derived from two individual parent sets and run in two technical replicates on the CFX96TM Real Time PCR Detection System (Bio-Rad) for a total of four samples per gene. RpL32 (RP-49) or GAPDH were used as reference genes and the knockdown efficency (%) was obtained using the ΔΔCq calculation method compared to the control (GAL4 only).

### Embryo handling and preparation

NC4-6 embryos were collected for 30 min at 25°C, then harvested at room temperature (22°C). Collected embryos were dechorionated by hand, desiccated for 5 min, mounted onto No. 1.5 coverslips coated with glue, and covered with Series 700 halocarbon oil (Halocarbon Products Corp.) as previously described [22].

### Drug Injections

Pharmacological inhibitors were injected into NC4-6 staged *Drosophila* embryos, incubated at room temperature (22°C) for 5 min, and then subjected to laser wounding. The following inhibitors were used: *α*-amanitin (1 mg/ml; Sigma-Aldrich); puromycin (10 mg/ml; Sigma-Aldrich); and cycloheximide (1 mg/ml; Sigma-Aldrich). The inhibitors were prepared in injection buffer (5 mM KCl, 0.1 mM NaP pH6.8). Injection buffer alone was used as the control.

### Laser Wounding

All wounds were generated with a pulsed nitrogen N2 micropoint laser (Andor Technology Ltd.) set to 435nm and focused at the lateral surface of the embryo. A circular targeted region of 16×15.5 µm was selected along the lateral midsection of the embryo, and ablation was controlled by MetaMorph software (Molecular Devices). Average ablation time was less than 3 seconds and time-lapse image acquisition was initiated immediately after ablation. Upon ablation, a grid-like pattern is sometimes observed (fluorescent dots within the wound area), as a result of the laser scoring the vitelline membrane that envelops the embryo. This vitelline membrane scoring has no effect on wound repair dynamics.

### Immunostaining of wounded embryos

Embryos (1-2 min post-wounding) were fixed in formaldehyde saturated heptane for 40 min. The vitelline membrane was removed by hand and the embryos were then washed 3 times with PAT [1x PBS, 0.1% Tween-20, 1% bovine serum albumin (BSA), 0.05% azide], then blocked in PAT for 2h at 4°C. Embryos were incubated with mouse anti-chickadee antibody (chi 1J; 1:10; Developmental Studies Hybridoma Bank) and guinea pig anti-Girdin antibody (1:500; provided by Patrick Laprise) [90] and for 24h at 4°C. Embryos were then washed 3 times with XNS (1x PBS, 0.1% Tween-20, 0.1% BSA, 4% normal goat serum) for 40 min each. Embryos were incubated with Alexa Fluor 568-and Alexa Fluor 633-conjugated secondary antibodies (1:1000; Invitrogen) overnight at 4°C. Embryos were washed with PTW (1x PBS, 0.1% Tween-20), incubated with Alexa Fluor 488-conjugated Phalloidin at 0.005 units/μl (Molecular Probes/Invitrogen, Rockford, IL) at room temperature for 1 h, washed with PTW, and then imaged.

### Live Image Acquisition

All imaging was done using a Revolution WD systems (Andor Technology Ltd.) mounted on a Leica DMi8 (Leica Microsystems Inc.) with a 63x/1.4 NA objective lens under the control of MetaMorph software (Molecular devices). Images were acquired using a 488 nm, 561 nm, and 633 nm Lasers and Andor iXon Ultra 897 EMCCD camera (Andor Technology Ltd.). All time-lapse images were acquired with 17-20 µm stacks/0.25 µm steps. For single color, images were acquired every 30 sec for 15 min and then every 60 sec for 25 min. For dual green and red colors, images were acquired every 45 sec for 30-40 min.

### Image processing, analysis, and quanitification

Image processing was performed using FIJI software [95]. Kymographs were generated using the crop feature to select ROIs of 5.3 × 94.9 µm. To generate fluorescent profile plots by R, 10 pixel sections across the wound from a single embryo were generated using Fiji as we described previously [43]. For dynamic lineplots, we generated fluorescent profile plots from each timepoint and then concatenated them. The lines represent the averaged fluorescent intensity and gray area is the 95% confidence interval. Line profiles from the left to right correspond to the top to bottom of the images unless otherwise noted. Wound area was manually measured using Fiji and the values were imported into Prism 8.2.1 (GraphPad Software Inc.) to construct corresponding graphs. Figures were assembled in Canvas Draw 6 for Mac (Canvas GFX, Inc.).

Quantification of the width and average intensity of actin ring, wound expansion, and closure rate was performed as follows: the width of actin ring was calculated with two measurement, the ferret diameters of the outer and inner edge of actin ring at 120 sec post-wounding. Using these measurements, the width of actin ring was calculated with (outer ferret diameter – inner ferret dimeter)/2. The average intensity of actin ring was calculated with two measurement. Instead of measuring ferret diameters, we measured area and integrated intensity in same regions as described in ring width. Using these measurements, the average intensity in the actin ring was calculated with (outer integrated intensity - inner integrated intensity)/(outer area - inner area). To calculate relative intensity for unwounded (UW) time point, average intensity at UW was measured with 50×50 pixels at the center of embryos and then averaged intensity of actin ring at each timepoint was divided by average intensity of UW. Wound expansion was calculated with max wound area/initial wound size. Contraction rate was calculated with two time points, one is t_max_ that is the time of reaching maximum wound area, the other is t<half that is the time of reaching 50-35% size of max wound since the slope of wound area curve changes after t<half. Using these time points, average speed was calculated with (wound area at t_max_ – wound area at t<half)/t_max_-t<half. To quantify the level of PIP3-GFP in the actin ring, we used the same method for the measurement of averaged actin ring intensity at 135 sec post-wounding image. Generation of all graphs and student’s t test were performed with Prism 8.2.1 (GraphPad Software Inc.).

### Microarray Preparation and Processing

Expression profiles were obtained using the FHCRC Fly 12k spotted array (GEO platform, GPL 1908). Embryos, prepared for wounding, were either wounded 8 times or left unwounded, then collected for total RNA extraction. Sample labeling and hybridization protocols were performed as described by Fazzio et al [96]. Specifically, cDNA targets were generated from total RNA using a standard amino-allyl labelling protocol where 30 ug of total RNA from each wounding condition (wounded vs non-wounded) were coupled to either Cy3 or Cy5 fluorophores. Targets were co-hybridized to microarrays for 16 hours at 63C and sequentially washed at room temperature (22C) in: 1 x SSC and 0.03% SDS for 2 mins, 1 x SSC for 2 mins, 0.2 x SSC with agitation for 20 mins, and 0.05 x SSC with agitation for 10 mins. Arrays were immediately centrifuged until dry and scanned using a GenePix 4000B scanner (Molecular Devices, Sunnyvale, CA). Image analysis was performed using GenePix Pro 6.0.

### Microarray Analysis

Wounded and non-wounded samples were independently replicated 4 times each at the 0 min and 30 min time point. For each array, spot intensity signals were filtered and removed if the values did not exceed 3 standard deviations above the background signal, if the background subtracted signal was <100 in both channels, or if a spot was flagged as questionable by the GenePix Pro Software. Spot-levels ratios were log_2_ transformed and loess normalized using the Bioconductor package *limma[97]*. Differential gene expression between wounded and non-wounded states was determined using the Bioconductor package *limma*, and a false discovery rate (FDR) method was used to correct for multiple testing [98]. Significant differential gene expression was defined as |log_2_ (ratio)| ≥ 0.585 (± 1.5-fold) with FDR set to 5%. Gene ontology enrichment scores were determined using DAVID with significance based on EASE scores corrected for multiple testing [99, 100]. The microarray datasets are available at GEO (NCBI Gene Expression Omnibus) under accession numbers: GSE39481, GSE39482, and GSE39483.

### TAD analysis

Genes were mapped to previously described TADs [38]. A TAD by up/down regulated gene versus unaffected gene expressed on the microarray contigency table was assembled. Fisher’s exact test of independence was used to test the null hypothesis that porportion of differentially expressed genes was different per TAD.

### Gene Size Analysis

Gene size was determined as the size of the largest expressed transcript per gene (dm6 build) expressed on the arrays. The median plus 95% CI was determined using the bootstrap procedure and 1000 iterations.

### Statistical analysis

All statistical analysis was done using Prism 8.2.1 (GraphPad, San Diego, CA). Gene knockdowns were compared to the appropriate control, and statistical significance was calculated using a Student’s t-test with p<0.05 considered significant.

## Acknowledgements

We thank Ryan Basom, Patrick Laprise, Scott Somers, the Bloomington Stock Center, the Kyoto Stock Center, the Harvard Transgenic RNAi Project, the Vienna Drosophila RNAi Center, the Drosophila Genomics Resource Center, and the Developmental Studies Hybridoma Bank for advice, antibodies, DNAs, flies, and other reagents used in this study. This research was supported by NIH GM111635 (to SMP) and NCI Cancer Center Support Grant P30 CA015704 (for Cellular Imaging and Genomics Shared Resources).

## Author Contributions

All authors contributed to the design of the experiments, performed experiments, and analyzed data. JJD and JMV performed the bioinformatic analyses. MN, JMV, and SMP wrote the manuscript with input from all authors.

## Competing Interests

The authors declare no competing or financial interests.

## Supplementary Video Legends

**Video 1** | **Translation and transcription are needed for different aspects of cell wound repair**. (A-D) Time-lapse confocal xy images from Drosophila NC4-6 staged embryos expressing an actin marker (sGMCA): control (buffer only) (A), alpha-amanitin injected (D), puromycin injected (C), and cycloheximide injected (D). Dynamic smoothened fluorescence intensity profiles (arbitrary units) derived from averaged fluorescence intensity values over a 10 pixel width across the wound area in each timepoint are shown below the image. Gray area represents the 95% CI. Time post-wounding is indicated. UW: unwounded.

**Video 2** | **Knockdown of up-regulated genes results in wound over-expansion and abnormal actin dynamics**. (A-H) Time-lapse confocal xy images from Drosophila NC4-6 staged embryos expressing an actin marker (sGMCA): control (w^1118^/+; sGMCA, 7063/+) (A), ImpL2^RNAi^/+; sGMCA, 7063/+ (B), EGFR^RNAi^/+; sGMCA, 7063/+ (C), Gp150^RNAi^/sGMCA, 7063 (D), Inx3^RNAi^/sGMCA, 7063 (E), Thor^RNAi^/sGMCA, 7063 (F), Jbug^RNAi^/sGMCA, 7063 (G), Nullo^RNAi^/sGMCA, 7063 (H). Dynamic smoothened fluorescence intensity profiles (arbitrary units) derived from averaged fluorescence intensity values over a 10 pixel width across the wound area in each timepoint are shown below the image. Gray area represents the 95% CI. Time post-wounding is indicated. UW: unwounded.

**Video 3** | **Knockdown of down-regulated genes results in wound over-expansion and abnormal actin dynamics**. (A-G) Time-lapse confocal xy images from Drosophila NC4-6 staged embryos expressing an actin marker (sGMCA): CG31075^RNAi^/+; sGMCA, 7063/+ (A), GstD1^RNAi^/+; sGMCA, 7063/+ (B), dhd^RNAi^/+; sGMCA, 7063/+ (C), Exu^RNAi^/+; sGMCA, 7063/+ (D), CG4960^RNAi^/+; sGMCA, 7063/+ (E), GstD2^RNAi^/+; sGMCA, 7063/+ (F), CG1598^RNAi^/+; sGMCA, 7063/+ (G). Dynamic smoothened fluorescence intensity profiles (arbitrary units) derived from averaged fluorescence intensity values over a 10 pixel width across the wound area in each timepoint are shown below the image. Gray area represents the 95% CI. Time post-wounding is indicated. UW: unwounded.

**Video 4** | **Actin dynamics of IIS pathway mutants**. (A-I) Time-lapse confocal xy images from Drosophila NC4-6 staged embryos expressing an actin marker (sGMCA): *insulin-like peptide 4*^*1*^ (*Ilp4*^*1*^; A), InR^RNAi(1)^/+; InR^RNAi(2)^/sGMCA, 7063 (B), Chico^RNAi^/sGMCA, 7063 (C), Pi3K21B^RNAi^/sGMCA, 7063 (D), Akt1^RNAi^/sGMCA, 7063 (E), FoxO^RNAi^/sGMCA, 7063 (F), Reptor^RNAi^/sGMCA, 7063 (G), Girdin^RNAi^/+; sGMCA, 7063/+ (H), and sGMCA; *chickadee*^*221*^/+ sGMCA, *wimp*/+ (reduced chickadee) (I). Dynamic smoothened fluorescence intensity profiles (arbitrary units) derived from averaged fluorescence intensity values over a 10 pixel width across the wound area in each timepoint are shown below the image. Gray area represents the 95% CI. Time post-wounding is indicated. UW: unwounded.

**Supplementary Figure S1.**
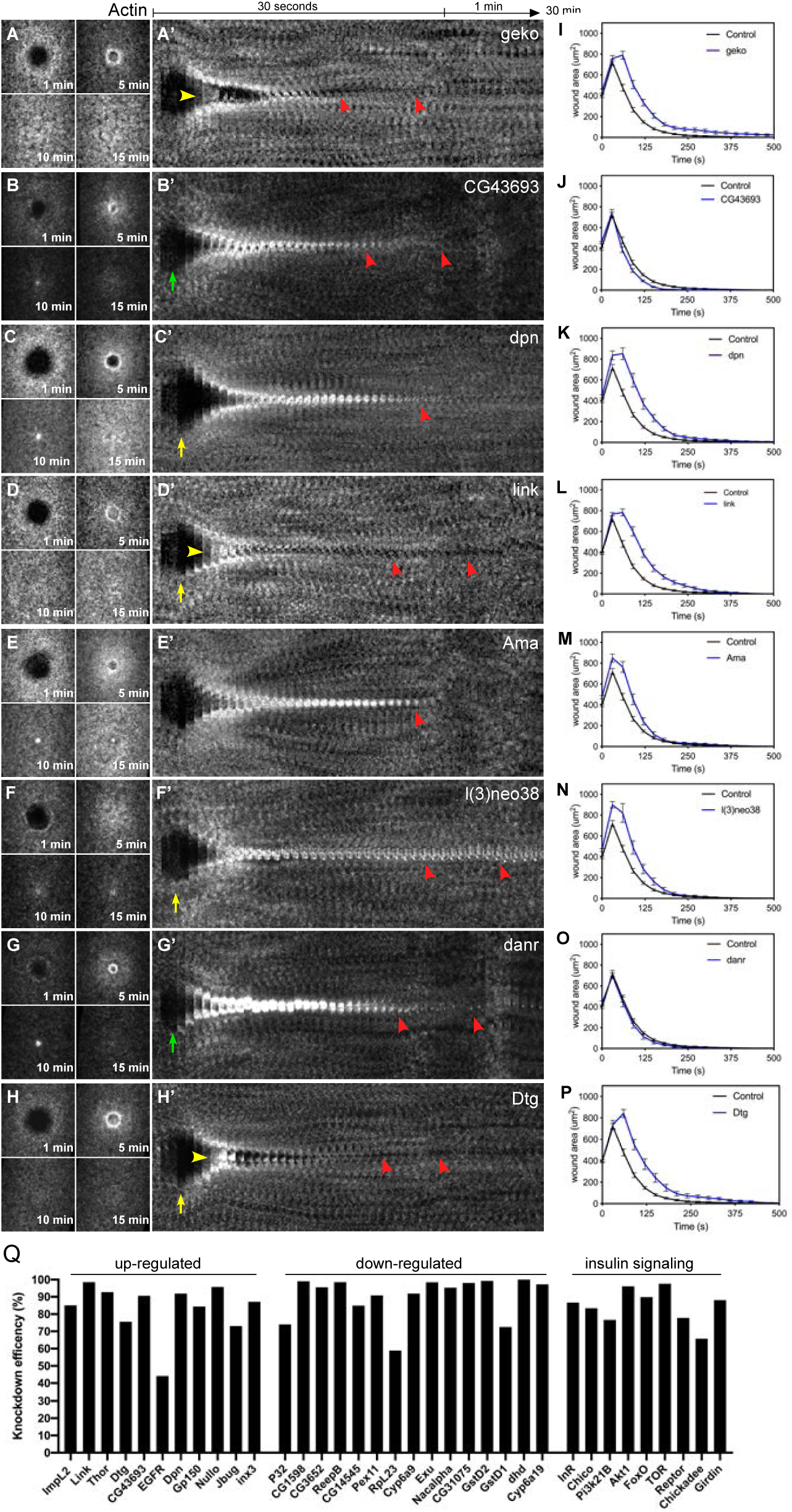
Knockdown of up-regulated genes results in wound overexpansion and abnormal actin dynamics. (**A-H**) Confocal XY projections of actin dynamics at 1, 5, 10, and 15 mpw from *Drosophila* NC4-6 embryos co-expressing sGMCA and a UAS-RNAi transgene during cell wound repair for Geko^RNAi^/+; sGMCA,7063/+ (**A**), CG43693^RNAi^/+; sGMCA,7063/+ (**B**), Dpn^RNAi^/+; sGMCA,7063/+ (**C**), Link^RNAi^/+; sGMCA,7063/+ (**D**), Ama^RNAi^/+; sGMCA,7063/+ (**E**), l(3)neo38^RNAi^/+; sGMCA,7063/+ (**F**), Danr^RNAi^/+; sGMCA,7063/+ (**G**), and Dtg^RNAi^/+; sGMCA,7063/+ (**H**). (**A’-H’**) XY kymographs across the wound areas depicted in (**A-H**), respectively. Note wound overexpansion (yellow arrows), wound under-expansion (green arrows), internal actin accumulation (yellow arrowhead), and remodelling defect/open wound (red arrowhead). (**I-P**) Quantification of wound area over time for (**A-H’**), respectively. (**Q**) Quantification of RNAi efficiencies for each RNAi mutant background (2 biological and 2 technical replicates were performed). Error bars represent ± SEM; n ≥10. Scale bars: 20 μm.

**Supplementary Fig. S2.**
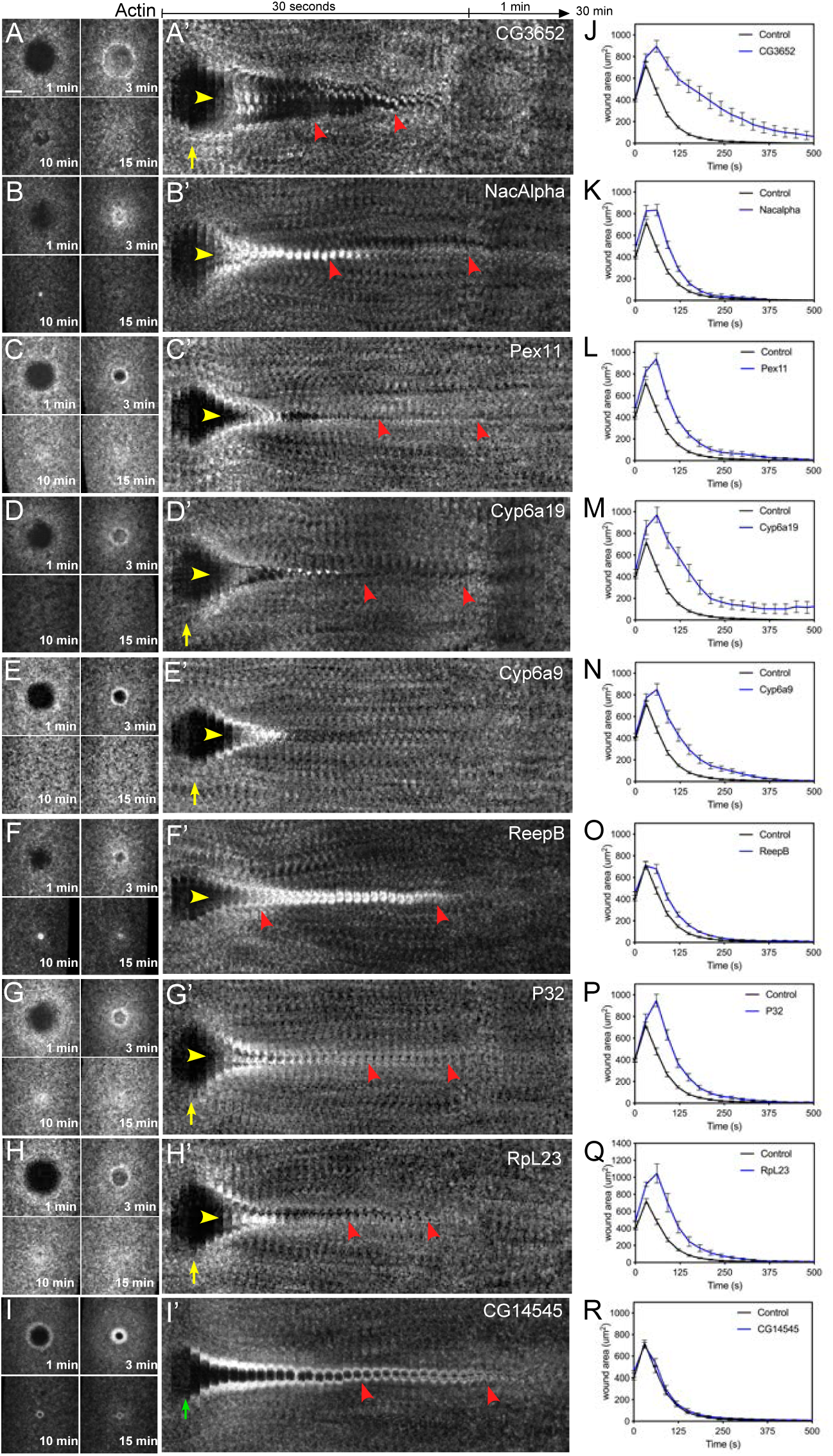
Knock-down of down-regulated genes results in wound overexpansion and abnormal actin dynamics. **a-i**, Confocal XY projections of actin dynamics at 1, 5, 10, and 15 mpw from *Drosophila* NC4-6 embryos co-expressing sGMCA and a UAS-RNAi transgene during cell wound repair for CG3652RNAi/+; sGMCA,7063/+ (**a**), NacAlphaRNAi/+; sGMCA,7063/+ (**b**), Pex11RNAi/+; sGMCA,7063/+ (**c**), Cyp6a19RNAi/+; sGMCA,7063/+ (**d**), Cyp6a9RNAi/+; sGMCA,7063/+ (**e**), ReepBRNAi/+; sGMCA,7063/+ (**f**), P32RNAi/+; sGMCA,7063/+ (**g**), RpL23RNAi/+; sGMCA,7063/+ (**h**), and CG14545RNAi/+; sGMCA,7063/+ (**i**). **a’-i’**, XY kymographs across the wound areas depicted in (**a-i**), respectively. Note wound over-expansion (yellow arrows), wound under-expansion (green arrows), internal actin accumulation (yellow arrowhead), and remodeling defect/open wound (red arrowhead). **j-r**, Quantification of wound area over time for (**a-i’**), respectively. Error bars represent ± SEM; n ≥10. Scale bars: 20 μm.

## Notes

https://www.ncbi.nlm.nih.gov/geo/query/acc.cgi?acc=GSE39481

https://www.ncbi.nlm.nih.gov/geo/query/acc.cgi?acc=GSE39482

https://www.ncbi.nlm.nih.gov/geo/query/acc.cgi?acc=GSE39483

### Competing Interest Statement

The authors have declared no competing interest.

### Summary of Updates

Additional data and quantifications included.

## References

1. Bement WM, Yu HY, Burkel BM, Vaughan EM, Clark AG. Rehabilitation and the single cell. Current opinion in cell biology. 2007;19(1):95-100. Epub 2006/12/19. doi: 10.1016/j.ceb.2006.12.001. PubMed PMID: 17174083; PubMed Central PMCID: PMCPMC4364133.

2. Sonnemann KJ, Bement WM. Wound repair: toward understanding and integration of single-cell and multicellular wound responses. Annu Rev Cell Dev Biol. 2011;27:237-63. Epub 2011/07/05. doi: 10.1146/annurev-cellbio-092910-154251. PubMed PMID: 21721944; PubMed Central PMCID: PMCPMC4878020.

3. Nakamura M, Dominguez ANM, Decker JR, Hull AJ, Verboon JM, Parkhurst SM. Into the breach: how cells cope with wounds. Open Biol. 2018;8(10). Epub 2018/10/05. doi: 10.1098/rsob.180135. PubMed PMID: 30282661; PubMed Central PMCID: PMCPMC6223217.

4. McNeil PL, Terasaki M. Coping with the inevitable: how cells repair a torn surface membrane. Nature cell biology. 2001;3(5):E124-9. Epub 2001/05/02. doi: 10.1038/35074652. PubMed PMID: 11331898.

5. McNeil PL, Kirchhausen T. An emergency response team for membrane repair. Nature reviews Molecular cell biology. 2005;6(6):499-505. Epub 2005/06/02. doi: 10.1038/nrm1665. PubMed PMID: 15928713.

6. Gurtner GC, Werner S, Barrandon Y, Longaker MT. Wound repair and regeneration. Nature. 2008;453(7193):314-21. Epub 2008/05/16. doi: 10.1038/nature07039. PubMed PMID: 18480812.

7. Velnar T, Bailey T, Smrkolj V. The wound healing process: an overview of the cellular and molecular mechanisms. J Int Med Res. 2009;37(5):1528-42. Epub 2009/11/26. doi: 10.1177/147323000903700531. PubMed PMID: 19930861.

8. Tang SKY, Marshall WF. Self-repairing cells: How single cells heal membrane ruptures and restore lost structures. Science. 2017;356(6342):1022-5. Epub 2017/06/10. doi: 10.1126/science.aam6496. PubMed PMID: 28596334; PubMed Central PMCID: PMCPMC5664224.

9. Moe AM, Golding AE, Bement WM. Cell healing: Calcium, repair and regeneration. Seminars in cell & developmental biology. 2015;45:18-23. Epub 2015/10/31. doi: 10.1016/j.semcdb.2015.09.026. PubMed PMID: 26514621; PubMed Central PMCID: PMCPMC4849125.

10. Draeger A, Schoenauer R, Atanassoff AP, Wolfmeier H, Babiychuk EB. Dealing with damage: plasma membrane repair mechanisms. Biochimie. 2014;107 Pt A:66-72. Epub 2014/09/04. doi: 10.1016/j.biochi.2014.08.008. PubMed PMID: 25183513.

11. Andrews NW, Corrotte M. Plasma membrane repair. Current biology : CB. 2018;28(8):R392-R7. Epub 2018/04/25. doi: 10.1016/j.cub.2017.12.034. PubMed PMID: 29689221.

12. Galan JE, Bliska JB. Cross-talk between bacterial pathogens and their host cells. Annu Rev Cell Dev Biol. 1996;12:221-55. Epub 1996/01/01. doi: 10.1146/annurev.cellbio.12.1.221. PubMed PMID: 8970727.

13. Clarke MS, Caldwell RW, Chiao H, Miyake K, McNeil PL. Contraction-induced cell wounding and release of fibroblast growth factor in heart. Circ Res. 1995;76(6):927-34. Epub 1995/06/01. doi: 10.1161/01.res.76.6.927. PubMed PMID: 7538917.

14. Coulombe PA, Hutton ME, Letai A, Hebert A, Paller AS, Fuchs E. Point mutations in human keratin 14 genes of epidermolysis bullosa simplex patients: genetic and functional analyses. Cell. 1991;66(6):1301-11. Epub 1991/09/20. doi: 10.1016/0092-8674(91)90051-y. PubMed PMID: 1717157.

15. Petrof BJ, Shrager JB, Stedman HH, Kelly AM, Sweeney HL. Dystrophin protects the sarcolemma from stresses developed during muscle contraction. Proceedings of the National Academy of Sciences of the United States of America. 1993;90(8):3710-4. Epub 1993/04/15. doi: 10.1073/pnas.90.8.3710. PubMed PMID: 8475120; PubMed Central PMCID: PMCPMC46371.

16. Cooper ST, McNeil PL. Membrane Repair: Mechanisms and Pathophysiology. Physiol Rev. 2015;95(4):1205-40. Epub 2015/09/04. doi: 10.1152/physrev.00037.2014. PubMed PMID: 26336031; PubMed Central PMCID: PMCPMC4600952.

17. Steinhardt RA, Bi G, Alderton JM. Cell membrane resealing by a vesicular mechanism similar to neurotransmitter release. Science. 1994;263(5145):390-3. Epub 1994/01/21. doi: 10.1126/science.7904084. PubMed PMID: 7904084.

18. Terasaki M, Miyake K, McNeil PL. Large plasma membrane disruptions are rapidly resealed by Ca2+-dependent vesicle-vesicle fusion events. The Journal of cell biology. 1997;139(1):63-74. Epub 1997/10/06. doi: 10.1083/jcb.139.1.63. PubMed PMID: 9314529; PubMed Central PMCID: PMCPMC2139822.

19. Bement WM, Mandato CA, Kirsch MN. Wound-induced assembly and closure of an actomyosin purse string in Xenopus oocytes. Current biology : CB. 1999;9(11):579-87. Epub 1999/06/09. doi: 10.1016/s0960-9822(99)80261-9. PubMed PMID: 10359696.

20. Kono K, Saeki Y, Yoshida S, Tanaka K, Pellman D. Proteasomal degradation resolves competition between cell polarization and cellular wound healing. Cell. 2012;150(1):151-64. Epub 2012/06/26. doi: 10.1016/j.cell.2012.05.030. PubMed PMID: 22727045.

21. Yumura S, Hashima S, Muranaka S. Myosin II does not contribute to wound repair in Dictyostelium cells. Biol Open. 2014;3(10):966-73. Epub 2014/09/23. doi: 10.1242/bio.20149712. PubMed PMID: 25238760; PubMed Central PMCID: PMCPMC4197445.

22. Abreu-Blanco MT, Verboon JM, Parkhurst SM. Cell wound repair in Drosophila occurs through three distinct phases of membrane and cytoskeletal remodeling. The Journal of cell biology. 2011;193(3):455-64. Epub 2011/04/27. doi: 10.1083/jcb.201011018. PubMed PMID: 21518790; PubMed Central PMCID: PMCPMC3087011.

23. Abreu-Blanco MT, Verboon JM, Parkhurst SM. Single cell wound repair: Dealing with life’s little traumas. Bioarchitecture. 2011;1(3):114-21. Epub 2011/09/17. doi: 10.4161/bioa.1.3.17091. PubMed PMID: 21922041; PubMed Central PMCID: PMCPMC3173964.

24. Benink HA, Bement WM. Concentric zones of active RhoA and Cdc42 around single cell wounds. The Journal of cell biology. 2005;168(3):429-39. Epub 2005/02/03. doi: 10.1083/jcb.200411109. PubMed PMID: 15684032; PubMed Central PMCID: PMCPMC2171735.

25. Burkel BM, Benink HA, Vaughan EM, von Dassow G, Bement WM. A Rho GTPase signal treadmill backs a contractile array. Developmental cell. 2012;23(2):384-96. Epub 2012/07/24. doi: 10.1016/j.devcel.2012.05.025. PubMed PMID: 22819338; PubMed Central PMCID: PMCPMC3549422.

26. Abreu-Blanco MT, Verboon JM, Parkhurst SM. Coordination of Rho family GTPase activities to orchestrate cytoskeleton responses during cell wound repair. Current biology : CB. 2014;24(2):144-55. Epub 2014/01/07. doi: 10.1016/j.cub.2013.11.048. PubMed PMID: 24388847; PubMed Central PMCID: PMCPMC3925435.

27. Grembowicz KP, Sprague D, McNeil PL. Temporary disruption of the plasma membrane is required for c-fos expression in response to mechanical stress. Molecular biology of the cell. 1999;10(4):1247-57. Epub 1999/04/10. doi: 10.1091/mbc.10.4.1247. PubMed PMID: 10198070; PubMed Central PMCID: PMCPMC25264.

28. Togo T. Long-term potentiation of wound-induced exocytosis and plasma membrane repair is dependent on cAMP-response element-mediated transcription via a protein kinase C- and p38 MAPK-dependent pathway. The Journal of biological chemistry. 2004;279(43):44996-5003. Epub 2004/08/20. doi: 10.1074/jbc.M406327200. PubMed PMID: 15317814.

29. Fein A, Terasaki M. Rapid increase in plasma membrane chloride permeability during wound resealing in starfish oocytes. J Gen Physiol. 2005;126(2):151-9. Epub 2005/07/27. doi: 10.1085/jgp.200509294. PubMed PMID: 16043775; PubMed Central PMCID: PMCPMC2266568.

30. Joost S, Jacob T, Sun X, Annusver K, La Manno G, Sur I, et al. Single-Cell Transcriptomics of Traced Epidermal and Hair Follicle Stem Cells Reveals Rapid Adaptations during Wound Healing. Cell reports. 2018;25(3):585-97 e7. Epub 2018/10/18. doi: 10.1016/j.celrep.2018.09.059. PubMed PMID: 30332640.

31. Verrier B, Muller D, Bravo R, Muller R. Wounding a fibroblast monolayer results in the rapid induction of the c-fos proto-oncogene. The EMBO journal. 1986;5(5):913-7. Epub 1986/05/01. PubMed PMID: 3522222; PubMed Central PMCID: PMCPMC1166882.

32. Martin P, Nobes CD. An early molecular component of the wound healing response in rat embryos--induction of c-fos protein in cells at the epidermal wound margin. Mech Dev. 1992;38(3):209-15. Epub 1992/09/01. doi: 10.1016/0925-4773(92)90054-n. PubMed PMID: 1457382.

33. Yates S, Rayner TE. Transcription factor activation in response to cutaneous injury: role of AP-1 in reepithelialization. Wound Repair Regen. 2002;10(1):5-15. Epub 2002/05/02. doi: 10.1046/j.1524-475x.2002.10902.x. PubMed PMID: 11983002.

34. Shaulian E, Karin M. AP-1 as a regulator of cell life and death. Nature cell biology. 2002;4(5):E131-6. Epub 2002/05/04. doi: 10.1038/ncb0502-e131. PubMed PMID: 11988758.

35. Geiger JA, Carvalho L, Campos I, Santos AC, Jacinto A. Hole-in-one mutant phenotypes link EGFR/ERK signaling to epithelial tissue repair in Drosophila. PloS one. 2011;6(11):e28349. Epub 2011/12/06. doi: 10.1371/journal.pone.0028349. PubMed PMID: 22140578; PubMed Central PMCID: PMCPMC3226689.

36. Brock AR, Wang Y, Berger S, Renkawitz-Pohl R, Han VC, Wu Y, et al. Transcriptional regulation of Profilin during wound closure in Drosophila larvae. Journal of cell science. 2012;125(Pt 23):5667-76. Epub 2012/09/15. doi: 10.1242/jcs.107490. PubMed PMID: 22976306; PubMed Central PMCID: PMCPMC3575702.

37. Losick VP, Jun AS, Spradling AC. Wound-Induced Polyploidization: Regulation by Hippo and JNK Signaling and Conservation in Mammals. PloS one. 2016;11(3):e0151251. Epub 2016/03/10. doi: 10.1371/journal.pone.0151251. PubMed PMID: 26958853; PubMed Central PMCID: PMCPMC4784922.

38. Li Q, Tjong H, Li X, Gong K, Zhou XJ, Chiolo I, et al. The three-dimensional genome organization of Drosophila melanogaster through data integration. Genome biology. 2017;18(1):145. Epub 2017/08/02. doi: 10.1186/s13059-017-1264-5. PubMed PMID: 28760140; PubMed Central PMCID: PMCPMC5576134.

39. Artieri CG, Fraser HB. Transcript length mediates developmental timing of gene expression across Drosophila. Mol Biol Evol. 2014;31(11):2879-89. Epub 2014/07/30. doi: 10.1093/molbev/msu226. PubMed PMID: 25069653; PubMed Central PMCID: PMCPMC4209130.

40. Fukaya T, Lim B, Levine M. Rapid Rates of Pol II Elongation in the Drosophila Embryo. Current biology : CB. 2017;27(9):1387-91. Epub 2017/05/02. doi: 10.1016/j.cub.2017.03.069. PubMed PMID: 28457866; PubMed Central PMCID: PMCPMC5665007.

41. Sandler JE, Irizarry J, Stepanik V, Dunipace L, Amrhein H, Stathopoulos A. A Developmental Program Truncates Long Transcripts to Temporally Regulate Cell Signaling. Developmental cell. 2018;47(6):773-84 e6. Epub 2018/12/19. doi: 10.1016/j.devcel.2018.11.019. PubMed PMID: 30562515; PubMed Central PMCID: PMCPMC6506262.

42. Vastenhouw NL, Cao WX, Lipshitz HD. The maternal-to-zygotic transition revisited. Development. 2019;146(11). Epub 2019/06/14. doi: 10.1242/dev.161471. PubMed PMID: 31189646.

43. Nakamura M, Verboon JM, Parkhurst SM. Prepatterning by RhoGEFs governs Rho GTPase spatiotemporal dynamics during wound repair. The Journal of cell biology. 2017;216(12):3959-69. Epub 2017/09/20. doi: 10.1083/jcb.201704145. PubMed PMID: 28923977; PubMed Central PMCID: PMCPMC5716286.

44. Forrest KM, Gavis ER. Live imaging of endogenous RNA reveals a diffusion and entrapment mechanism for nanos mRNA localization in Drosophila. Current biology : CB. 2003;13(14):1159-68. Epub 2003/07/18. doi: 10.1016/s0960-9822(03)00451-2. PubMed PMID: 12867026.

45. Weil TT, Forrest KM, Gavis ER. Localization of bicoid mRNA in late oocytes is maintained by continual active transport. Developmental cell. 2006;11(2):251-62. Epub 2006/08/08. doi: 10.1016/j.devcel.2006.06.006. PubMed PMID: 16890164.

46. Brand AH, Perrimon N. Targeted gene expression as a means of altering cell fates and generating dominant phenotypes. Development. 1993;118(2):401-15. Epub 1993/06/01. PubMed PMID: 8223268.

47. Rorth P. Gal4 in the Drosophila female germline. Mech Dev. 1998;78(1-2):113-8. Epub 1998/12/22. doi: 10.1016/s0925-4773(98)00157-9. PubMed PMID: 9858703.

48. Kiehart DP, Galbraith CG, Edwards KA, Rickoll WL, Montague RA. Multiple forces contribute to cell sheet morphogenesis for dorsal closure in Drosophila. The Journal of cell biology. 2000;149(2):471-90. Epub 2000/04/18. doi: 10.1083/jcb.149.2.471. PubMed PMID: 10769037; PubMed Central PMCID: PMCPMC2175161.

49. Figueroa-Clarevega A, Bilder D. Malignant Drosophila tumors interrupt insulin signaling to induce cachexia-like wasting. Developmental cell. 2015;33(1):47-55. Epub 2015/04/09. doi: 10.1016/j.devcel.2015.03.001. PubMed PMID: 25850672; PubMed Central PMCID: PMCPMC4390765.

50. Kwon Y, Song W, Droujinine IA, Hu Y, Asara JM, Perrimon N. Systemic organ wasting induced by localized expression of the secreted insulin/IGF antagonist ImpL2. Developmental cell. 2015;33(1):36-46. Epub 2015/04/09. doi: 10.1016/j.devcel.2015.02.012. PubMed PMID: 25850671; PubMed Central PMCID: PMCPMC4437243.

51. Amoyel M, Hillion KH, Margolis SR, Bach EA. Somatic stem cell differentiation is regulated by PI3K/Tor signaling in response to local cues. Development. 2016;143(21):3914-25. Epub 2016/11/03. doi: 10.1242/dev.139782. PubMed PMID: 27633989; PubMed Central PMCID: PMCPMC5117146.

52. Kushnir T, Mezuman S, Bar-Cohen S, Lange R, Paroush Z, Helman A. Novel interplay between JNK and Egfr signaling in Drosophila dorsal closure. PLoS genetics. 2017;13(6):e1006860. Epub 2017/06/20. doi: 10.1371/journal.pgen.1006860. PubMed PMID: 28628612; PubMed Central PMCID: PMCPMC5495517.

53. Nakamura F, Stossel TP, Hartwig JH. The filamins: organizers of cell structure and function. Cell Adh Migr. 2011;5(2):160-9. Epub 2010/12/21. doi: 10.4161/cam.5.2.14401. PubMed PMID: 21169733; PubMed Central PMCID: PMCPMC3084982.

54. Martino F, Perestrelo AR, Vinarsky V, Pagliari S, Forte G. Cellular Mechanotransduction: From Tension to Function. Front Physiol. 2018;9:824. Epub 2018/07/22. doi: 10.3389/fphys.2018.00824. PubMed PMID: 30026699; PubMed Central PMCID: PMCPMC6041413.

55. Sokac AM, Wieschaus E. Local actin-dependent endocytosis is zygotically controlled to initiate Drosophila cellularization. Developmental cell. 2008;14(5):775-86. Epub 2008/05/15. doi: 10.1016/j.devcel.2008.02.014. PubMed PMID: 18477459; PubMed Central PMCID: PMCPMC2517610.

56. Sokac AM, Wieschaus E. Zygotically controlled F-actin establishes cortical compartments to stabilize furrows during Drosophila cellularization. Journal of cell science. 2008;121(11):1815-24. Epub 2008/05/08. doi: 10.1242/jcs.025171. PubMed PMID: 18460582; PubMed Central PMCID: PMCPMC2728442.

57. Li Y, Fetchko M, Lai ZC, Baker NE. Scabrous and Gp150 are endosomal proteins that regulate Notch activity. Development. 2003;130(13):2819-27. Epub 2003/05/21. doi: 10.1242/dev.00495. PubMed PMID: 12756167.

58. Giuliani F, Giuliani G, Bauer R, Rabouille C. Innexin 3, a new gene required for dorsal closure in Drosophila embryo. PloS one. 2013;8(7):e69212. Epub 2013/07/31. doi: 10.1371/journal.pone.0069212. PubMed PMID: 23894431; PubMed Central PMCID: PMCPMC3722180.

59. Lautemann J, Bohrmann J. Relating proton pumps with gap junctions: colocalization of ductin, the channel-forming subunit c of V-ATPase, with subunit a and with innexins 2 and 3 during Drosophila oogenesis. BMC Dev Biol. 2016;16(1):24. Epub 2016/07/15. doi: 10.1186/s12861-016-0124-y. PubMed PMID: 27412523; PubMed Central PMCID: PMCPMC4944501.

60. Toshniwal AG, Gupta S, Mandal L, Mandal S. ROS Inhibits Cell Growth by Regulating 4EBP and S6K, Independent of TOR, during Development. Developmental cell. 2019;49(3):473-89 e9. Epub 2019/05/08. doi: 10.1016/j.devcel.2019.04.008. PubMed PMID: 31063760.

61. Vinayagam A, Kulkarni MM, Sopko R, Sun X, Hu Y, Nand A, et al. An Integrative Analysis of the InR/PI3K/Akt Network Identifies the Dynamic Response to Insulin Signaling. Cell reports. 2016;16(11):3062-74. Epub 2016/09/15. doi: 10.1016/j.celrep.2016.08.029. PubMed PMID: 27626673; PubMed Central PMCID: PMCPMC5033061.

62. Galiano RD, Tepper OM, Pelo CR, Bhatt KA, Callaghan M, Bastidas N, et al. Topical vascular endothelial growth factor accelerates diabetic wound healing through increased angiogenesis and by mobilizing and recruiting bone marrow-derived cells. Am J Pathol. 2004;164(6):1935-47. Epub 2004/05/27. doi: 10.1016/S0002-9440(10)63754-6. PubMed PMID: 15161630; PubMed Central PMCID: PMCPMC1615774.

63. Brem H, Tomic-Canic M. Cellular and molecular basis of wound healing in diabetes. J Clin Invest. 2007;117(5):1219-22. Epub 2007/05/04. doi: 10.1172/JCI32169. PubMed PMID: 17476353; PubMed Central PMCID: PMCPMC1857239.

64. Kakanj P, Moussian B, Gronke S, Bustos V, Eming SA, Partridge L, et al. Insulin and TOR signal in parallel through FOXO and S6K to promote epithelial wound healing. Nature communications. 2016;7:12972. Epub 2016/10/08. doi: 10.1038/ncomms12972. PubMed PMID: 27713427; PubMed Central PMCID: PMCPMC5059774.

65. Manzano-Nunez F, Arambul-Anthony MJ, Galan Albinana A, Leal Tassias A, Acosta Umanzor C, Borreda Gasco I, et al. Insulin resistance disrupts epithelial repair and niche-progenitor Fgf signaling during chronic liver injury. PLoS biology. 2019;17(1):e2006972. Epub 2019/01/30. doi: 10.1371/journal.pbio.2006972. PubMed PMID: 30695023; PubMed Central PMCID: PMCPMC6368328.

66. Britton JS, Lockwood WK, Li L, Cohen SM, Edgar BA. Drosophila’s insulin/PI3-kinase pathway coordinates cellular metabolism with nutritional conditions. Developmental cell. 2002;2(2):239-49. Epub 2002/02/08. doi: 10.1016/s1534-5807(02)00117-x. PubMed PMID: 11832249.

67. Rameh LE, Cantley LC. The role of phosphoinositide 3-kinase lipid products in cell function. The Journal of biological chemistry. 1999;274(13):8347-50. Epub 1999/03/20. doi: 10.1074/jbc.274.13.8347. PubMed PMID: 10085060.

68. Ghiglione C, Jouandin P, Cerezo D, Noselli S. The Drosophila insulin pathway controls Profilin expression and dynamic actin-rich protrusions during collective cell migration. Development. 2018;145(14). Epub 2018/07/08. doi: 10.1242/dev.161117. PubMed PMID: 29980565.

69. Hartung A, Ordelheide AM, Staiger H, Melzer M, Haring HU, Lammers R. The Akt substrate Girdin is a regulator of insulin signaling in myoblast cells. Biochimica et biophysica acta. 2013;1833(12):2803-11. Epub 2013/07/28. doi: 10.1016/j.bbamcr.2013.07.012. PubMed PMID: 23886629.

70. Lopez-Sanchez I, Ma GS, Pedram S, Kalogriopoulos N, Ghosh P. GIV/girdin binds exocyst subunit-Exo70 and regulates exocytosis of GLUT4 storage vesicles. Biochem Biophys Res Commun. 2015;468(1-2):287-93. Epub 2015/10/31. doi: 10.1016/j.bbrc.2015.10.111. PubMed PMID: 26514725; PubMed Central PMCID: PMCPMC4659757.

71. Parkhurst SM, Ish-Horowicz D. wimp, a dominant maternal-effect mutation, reduces transcription of a specific subset of segmentation genes in Drosophila. Genes & development. 1991;5(3):341-57. Epub 1991/03/01. doi: 10.1101/gad.5.3.341. PubMed PMID: 2001838.

72. Verboon JM, Decker JR, Nakamura M, Parkhurst SM. Wash exhibits context-dependent phenotypes and, along with the WASH regulatory complex, regulates Drosophila oogenesis. Journal of cell science. 2018;131(8). Epub 2018/03/20. doi: 10.1242/jcs.211573. PubMed PMID: 29549166; PubMed Central PMCID: PMCPMC5963843.

73. Bi GQ, Alderton JM, Steinhardt RA. Calcium-regulated exocytosis is required for cell membrane resealing. The Journal of cell biology. 1995;131(6 Pt 2):1747-58. Epub 1995/12/01. doi: 10.1083/jcb.131.6.1747. PubMed PMID: 8557742; PubMed Central PMCID: PMCPMC2120667.

74. Miyake K, McNeil PL. Vesicle accumulation and exocytosis at sites of plasma membrane disruption. The Journal of cell biology. 1995;131(6 Pt 2):1737-45. Epub 1995/12/01. doi: 10.1083/jcb.131.6.1737. PubMed PMID: 8557741; PubMed Central PMCID: PMCPMC2120668.

75. Verboon JM, Parkhurst SM. Rho family GTPases bring a familiar ring to cell wound repair. Small GTPases. 2015;6(1):1-7. Epub 2015/04/12. doi: 10.4161/21541248.2014.992262. PubMed PMID: 25862160; PubMed Central PMCID: PMCPMC4601322.

76. Foe VE, Alberts BM. Studies of nuclear and cytoplasmic behaviour during the five mitotic cycles that precede gastrulation in Drosophila embryogenesis. Journal of cell science. 1983;61:31-70. Epub 1983/05/01. PubMed PMID: 6411748.

77. Ardehali MB, Yao J, Adelman K, Fuda NJ, Petesch SJ, Webb WW, et al. Spt6 enhances the elongation rate of RNA polymerase II in vivo. The EMBO journal. 2009;28(8):1067-77. Epub 2009/03/13. doi: 10.1038/emboj.2009.56. PubMed PMID: 19279664; PubMed Central PMCID: PMCPMC2683705.

78. Shermoen AW, O’Farrell PH. Progression of the cell cycle through mitosis leads to abortion of nascent transcripts. Cell. 1991;67(2):303-10. Epub 1991/10/18. doi: 10.1016/0092-8674(91)90182-x. PubMed PMID: 1680567; PubMed Central PMCID: PMCPMC2755073.

79. Thummel CS, Burtis KC, Hogness DS. Spatial and temporal patterns of E74 transcription during Drosophila development. Cell. 1990;61(1):101-11. Epub 1990/04/06. doi: 10.1016/0092-8674(90)90218-4. PubMed PMID: 1690603.

80. Yao J, Ardehali MB, Fecko CJ, Webb WW, Lis JT. Intranuclear distribution and local dynamics of RNA polymerase II during transcription activation. Molecular cell. 2007;28(6):978-90. Epub 2007/12/27. doi: 10.1016/j.molcel.2007.10.017. PubMed PMID: 18158896.

81. Garcia HG, Tikhonov M, Lin A, Gregor T. Quantitative imaging of transcription in living Drosophila embryos links polymerase activity to patterning. Current biology : CB. 2013;23(21):2140-5. Epub 2013/10/22. doi: 10.1016/j.cub.2013.08.054. PubMed PMID: 24139738; PubMed Central PMCID: PMCPMC3828032.

82. O’Farrell PH. Developmental biology. Big genes and little genes and deadlines for transcription. Nature. 1992;359(6394):366-7. Epub 1992/10/01. doi: 10.1038/359366a0. PubMed PMID: 1406945; PubMed Central PMCID: PMCPMC2754300.

83. Hoskins RA, Landolin JM, Brown JB, Sandler JE, Takahashi H, Lassmann T, et al. Genome-wide analysis of promoter architecture in Drosophila melanogaster. Genome research. 2011;21(2):182-92. Epub 2010/12/24. doi: 10.1101/gr.112466.110. PubMed PMID: 21177961; PubMed Central PMCID: PMCPMC3032922.

84. Rutter GA, Tsuboi T, Ravier MA. Ca2+ microdomains and the control of insulin secretion. Cell Calcium. 2006;40(5-6):539-51. Epub 2006/10/13. doi: 10.1016/j.ceca.2006.08.015. PubMed PMID: 17030367.

85. Lee K, Kim J, Kohler M, Yu J, Shi Y, Yang SN, et al. Blocking Ca(2+) Channel beta3 Subunit Reverses Diabetes. Cell reports. 2018;24(4):922-34. Epub 2018/07/26. doi: 10.1016/j.celrep.2018.06.086. PubMed PMID: 30044988; PubMed Central PMCID: PMCPMC6083041.

86. Teleman AA, Chen YW, Cohen SM. 4E-BP functions as a metabolic brake used under stress conditions but not during normal growth. Genes & development. 2005;19(16):1844-8. Epub 2005/08/17. doi: 10.1101/gad.341505. PubMed PMID: 16103212; PubMed Central PMCID: PMCPMC1186183.

87. Puig O, Tjian R. Transcriptional feedback control of insulin receptor by dFOXO/FOXO1. Genes & development. 2005;19(20):2435-46. Epub 2005/10/19. doi: 10.1101/gad.1340505. PubMed PMID: 16230533; PubMed Central PMCID: PMCPMC1257398.

88. Haeusler RA, McGraw TE, Accili D. Biochemical and cellular properties of insulin receptor signalling. Nature reviews Molecular cell biology. 2018;19(1):31-44. Epub 2017/10/05. doi: 10.1038/nrm.2017.89. PubMed PMID: 28974775; PubMed Central PMCID: PMCPMC5894887.

89. Brunet A, Bonni A, Zigmond MJ, Lin MZ, Juo P, Hu LS, et al. Akt promotes cell survival by phosphorylating and inhibiting a Forkhead transcription factor. Cell. 1999;96(6):857-68. Epub 1999/04/02. doi: 10.1016/s0092-8674(00)80595-4. PubMed PMID: 10102273.

90. Houssin E, Tepass U, Laprise P. Girdin-mediated interactions between cadherin and the actin cytoskeleton are required for epithelial morphogenesis in Drosophila. Development. 2015;142(10):1777-84. Epub 2015/05/15. doi: 10.1242/dev.122002. PubMed PMID: 25968313.

91. Wang S, Lei Y, Cai Z, Ye X, Li L, Luo X, et al. Girdin regulates the proliferation and apoptosis of pancreatic cancer cells via the PI3K/Akt signalling pathway. Oncol Rep. 2018;40(2):599-608. Epub 2018/06/15. doi: 10.3892/or.2018.6469. PubMed PMID: 29901184; PubMed Central PMCID: PMCPMC6072288.

92. Choi JS, Kim KH, Oh E, Shin YK, Seo J, Kim SH, et al. Girdin protein expression is associated with poor prognosis in patients with invasive breast cancer. Pathology. 2017;49(6):618-26. Epub 2017/08/19. doi: 10.1016/j.pathol.2017.05.010. PubMed PMID: 28818465.

93. Wang X, Enomoto A, Weng L, Mizutani Y, Abudureyimu S, Esaki N, et al. Girdin/GIV regulates collective cancer cell migration by controlling cell adhesion and cytoskeletal organization. Cancer Sci. 2018;109(11):3643-56. Epub 2018/09/09. doi: 10.1111/cas.13795. PubMed PMID: 30194792; PubMed Central PMCID: PMCPMC6215880.

94. Liu R, Abreu-Blanco MT, Barry KC, Linardopoulou EV, Osborn GE, Parkhurst SM. Wash functions downstream of Rho and links linear and branched actin nucleation factors. Development. 2009;136(16):2849-60. Epub 2009/07/28. doi: 10.1242/dev.035246. PubMed PMID: 19633175; PubMed Central PMCID: PMCPMC2730411.

95. Schindelin J, Arganda-Carreras I, Frise E, Kaynig V, Longair M, Pietzsch T, et al. Fiji: an open-source platform for biological-image analysis. Nature methods. 2012;9(7):676-82. Epub 2012/06/30. doi: 10.1038/nmeth.2019. PubMed PMID: 22743772; PubMed Central PMCID: PMCPMC3855844.

96. Fazzio TG, Kooperberg C, Goldmark JP, Neal C, Basom R, Delrow J, et al. Widespread collaboration of Isw2 and Sin3-Rpd3 chromatin remodeling complexes in transcriptional repression. Molecular and cellular biology. 2001;21(19):6450-60. Epub 2001/09/05. doi: 10.1128/mcb.21.19.6450-6460.2001. PubMed PMID: 11533234; PubMed Central PMCID: PMCPMC99792.

97. Smyth GK, Michaud J, Scott HS. Use of within-array replicate spots for assessing differential expression in microarray experiments. Bioinformatics. 2005;21(9):2067-75. Epub 2005/01/20. doi: 10.1093/bioinformatics/bti270. PubMed PMID: 15657102.

98. Reiner A, Yekutieli D, Benjamini Y. Identifying differentially expressed genes using false discovery rate controlling procedures. Bioinformatics. 2003;19(3):368-75. Epub 2003/02/14. doi: 10.1093/bioinformatics/btf877. PubMed PMID: 12584122.

99. Huang da W, Sherman BT, Lempicki RA. Systematic and integrative analysis of large gene lists using DAVID bioinformatics resources. Nature protocols. 2009;4(1):44-57. Epub 2009/01/10. doi: 10.1038/nprot.2008.211. PubMed PMID: 19131956.

100. Huang da W, Sherman BT, Lempicki RA. Bioinformatics enrichment tools: paths toward the comprehensive functional analysis of large gene lists. Nucleic acids research. 2009;37(1):1-13. Epub 2008/11/27. doi: 10.1093/nar/gkn923. PubMed PMID: 19033363; PubMed Central PMCID: PMCPMC2615629.

