## Supplementary Tables 1 & 2 for "Autocrine insulin pathway signaling regulates actin dynamics in cell wound repair" (from another version of this preprint: version 1, 2020-04-07)

**Table S1.** Changes in gene expression at 30 minutes post-wounding.

| clone | flybase_gene_id | symbol | logFC_t30 | adj_p_t30 | sig_t30 |
|---|---|---|---|---|---|
| LD02462 | FBgn0033979 | Cyp6a19 | -1.391813323 | 0.02156494 | 1 |
| LD23983 | FBgn0011761 | dhd | -1.347598265 | 0.02156494 | 1 |
| RH47312 | FBgn0010038 | GstD2 | -1.307493407 | 0.02156494 | 1 |
| RH47312 | FBgn0001149 | GstD1 | -1.307493407 | 0.02156494 | 1 |
| RE18078 | FBgn0051075 | CG31075 | -1.26599667 | 0.02546743 | 1 |
| GH11940 | FBgn0086904 | Nacalpha | -1.238225026 | 0.02156494 | 1 |
| LD26657 | FBgn0000615 | exu | -1.166681288 | 0.02156494 | 1 |
| SD12535 | FBgn0013771 | Cyp6a9 | -1.160813804 | 0.02156494 | 1 |
| RH29296 | FBgn0010078 | RpL23 | -1.098851652 | 0.02156494 | 1 |
| RE39562 | FBgn0034058 | Pex11 | -1.080708951 | 0.02156494 | 1 |
| RE55691 | FBgn0040602 | CG14545 | -1.053904844 | 0.02156494 | 1 |
| GM14577 | FBgn0039371 | CG4960 | -1.018319157 | 0.02156494 | 1 |
| GM14577 | FBgn0033906 | CG8331 | -1.018319157 | 0.02156494 | 1 |
| GH04132 | FBgn0031600 | CG3652 | -1.00358416 | 0.02618235 | 1 |
| RH73327 | FBgn0033191 | CG1598 | -1.001068877 | 0.02156494 | 1 |
| LD29590 | FBgn0034259 | P32 | -1.000565767 | 0.02156494 | 1 |
| RE70889 | FBgn0010040 | GstD4 | -0.993393239 | 0.02156494 | 1 |
| RH08992 | FBgn0030704 | CG15916 | -0.989885417 | 0.02156494 | 1 |
| GH10864 | FBgn0086355 | Tpi | -0.974419596 | 0.02176486 | 1 |
| GH27411 | FBgn0037580 | DppIII | -0.963145591 | 0.02156494 | 1 |
| RE20434 | FBgn0026439 | Eaat1 | -0.960951224 | 0.02404547 | 1 |
| LP11595 | FBgn0012036 | Aldh | -0.951216025 | 0.02156494 | 1 |
| RH55882 | FBgn0001092 | Gapdh2 | -0.950783528 | 0.02156494 | 1 |
| HL01062 | FBgn0036316 | CG10960 | -0.946991757 | 0.02156494 | 1 |
| GM03716 | FBgn0038190 | CG9926 | -0.942095831 | 0.02156494 | 1 |
| LD11379 | FBgn0001225 | Hsp26 | -0.939133243 | 0.02156494 | 1 |
| RE67941 | FBgn0037435 | CG18048 | -0.932094561 | 0.02156494 | 1 |
| LD44652 | FBgn0028563 | sut1 | -0.928872812 | 0.02156494 | 1 |
| LD06393 | FBgn0086254 | CG6084 | -0.9252896 | 0.02618235 | 1 |
| RE50650 | FBgn0039909 | ND-49 | -0.914126931 | 0.02156494 | 1 |
| RE26822 | FBgn0038788 | Sirt2 | -0.904690503 | 0.02225 | 1 |
| RE21095 | FBgn0063494 | GstE6 | -0.894942434 | 0.02156494 | 1 |
| RE42508 | FBgn0039966 | Rab21 | -0.894843842 | 0.02156494 | 1 |
| LD07673 | FBgn0039869 | CG1890 | -0.892523924 | 0.02156494 | 1 |
| RE43665 | FBgn0034564 | CG9344 | -0.89129561 | 0.02156494 | 1 |
| LD44201 | FBgn0040207 | kat80 | -0.870531926 | 0.02156494 | 1 |
| RE19545 | FBgn0035725 | Mis12 | -0.840965312 | 0.04280714 | 1 |
| SD13688 | FBgn0004087 | Dhfr | -0.837829394 | 0.02156494 | 1 |
| SD18375 | FBgn0038692 | CG3773 | -0.824048869 | 0.02156494 | 1 |
| SD14726 | FBgn0031267 | Ipk2 | -0.822572547 | 0.02156494 | 1 |
| RE66240 | FBgn0029929 | CG4593 | -0.815120858 | 0.02667517 | 1 |
| RE01312 | FBgn0033127 | Tsp42Ef | -0.81348301 | 0.02156494 | 1 |
| SD06613 | FBgn0046214 | vig2 | -0.812576725 | 0.02363639 | 1 |
| GH24869 | FBgn0030411 | CG2540 | -0.807461673 | 0.04244772 | 1 |
| LP11313 | FBgn0010039 | GstD3 | -0.804470437 | 0.02156494 | 1 |
| GH13575 | FBgn0003074 | Pgi | -0.801832132 | 0.02156494 | 1 |

| | | | | | |
|---|---|---|---|---|---|
| LP09705 | FBgn0032233 | dpr19 | -0.795375758 | 0.02530044 | 1 |
| RE60511 | FBgn0263600 | DNApol-delta | -0.795004004 | 0.02156494 | 1 |
| GH09647 | FBgn0033373 | CG8080 | -0.79425312 | 0.02156494 | 1 |
| GH07925 | FBgn0036824 | CG3902 | -0.790341419 | 0.02156494 | 1 |
| RH24992 | FBgn0036007 | path | -0.789515809 | 0.02156494 | 1 |
| GM14134 | FBgn0023537 | CG17896 | -0.780772986 | 0.02156494 | 1 |
| AT31109 | FBgn0038858 | CG5793 | -0.779005181 | 0.03033006 | 1 |
| RH61928 | FBgn0030552 | mRpL38 | -0.777263139 | 0.03313349 | 1 |
| GH04104 | FBgn0040778 | CG17977 | -0.769950086 | 0.02156494 | 1 |
| GH02396 | FBgn0010288 | Uch | -0.760459901 | 0.02176486 | 1 |
| SD02122 | FBgn0040283 | SMC1 | -0.755476351 | 0.02677632 | 1 |
| LD47530 | FBgn0029174 | FKBP59 | -0.750605865 | 0.02156494 | 1 |
| RE19263 | FBgn0037022 | CG11396 | -0.746903506 | 0.02680023 | 1 |
| GM14514 | FBgn0038608 | WRNexo | -0.746565432 | 0.02948687 | 1 |
| LP02120 | FBgn0038577 | CG12321 | -0.745833175 | 0.03829112 | 1 |
| SD05329 | FBgn0037141 | DNApol-eta | -0.744861048 | 0.02363639 | 1 |
| RE68515 | FBgn0039404 | CG14543 | -0.740559359 | 0.02176486 | 1 |
| GH20044 | FBgn0024230 | Hs2st | -0.740451896 | 0.03746048 | 1 |
| RH01212 | FBgn0266666 | Sem1 | -0.733595541 | 0.02156494 | 1 |
| RH61539 | FBgn0010638 | Sec61beta | -0.732672802 | 0.02847382 | 1 |
| SD19278 | FBgn0267976 | Tim23 | -0.728787883 | 0.02677632 | 1 |
| SD08776 | FBgn0035312 | CG15820 | -0.725856296 | 0.02326108 | 1 |
| SD11637 | FBgn0033928 | Arc2 | -0.724620624 | 0.02156494 | 1 |
| RE29207 | FBgn0038950 | CG5382 | -0.722522202 | 0.0352673 | 1 |
| RE62833 | FBgn0035911 | CG6638 | -0.720274479 | 0.02728078 | 1 |
| RE29468 | FBgn0259143 | CG42258 | -0.71721742 | 0.02156494 | 1 |
| GM01936 | FBgn0032205 | CG4957 | -0.715657959 | 0.02363639 | 1 |
| LP01332 | FBgn0037368 | CG1239 | -0.714751741 | 0.04339964 | 1 |
| RE49388 | FBgn0035334 | CG8993 | -0.708688154 | 0.02156494 | 1 |
| GM10268 | FBgn0034614 | CG9752 | -0.708512725 | 0.02434869 | 1 |
| SD11293 | FBgn0032244 | RfC3 | -0.708361072 | 0.02461562 | 1 |
| LD30505 | FBgn0000629 | E(z) | -0.707172037 | 0.02945798 | 1 |
| RE44641 | FBgn0087040 | alphaTub67C | -0.70690029 | 0.02156494 | 1 |
| LD33701 | FBgn0039065 | Rad60 | -0.705339332 | 0.02351948 | 1 |
| GH01554 | FBgn0030695 | PGRP-LE | -0.701555859 | 0.02644978 | 1 |
| GH27264 | FBgn0265575 | yin | -0.697934949 | 0.02156494 | 1 |
| RE53816 | FBgn0031979 | CCDC53 | -0.697921024 | 0.02156494 | 1 |
| GH10148 | FBgn0262732 | mbf1 | -0.694583272 | 0.02363639 | 1 |
| GM15115 | FBgn0034729 | CG10344 | -0.693936488 | 0.02156494 | 1 |
| GH14435 | FBgn0015240 | Hr96 | -0.693257154 | 0.02869814 | 1 |
| RE56531 | FBgn0027538 | beta4GalNAcTA | -0.69118706 | 0.02351948 | 1 |
| RE60471 | FBgn0262559 | Mdh2 | -0.689720386 | 0.02176486 | 1 |
| GH05320 | FBgn0035150 | Rev1 | -0.688961515 | 0.03098436 | 1 |
| GH25425 | FBgn0032453 | CG6180 | -0.681567739 | 0.02618235 | 1 |
| LD38554 | FBgn0034021 | CG8180 | -0.679948418 | 0.02156494 | 1 |
| RH09920 | FBgn0035529 | Fdx2 | -0.676084831 | 0.03129058 | 1 |
| AT29239 | FBgn0014023 | mRpL47 | -0.673402115 | 0.0358459 | 1 |
| RE49550 | FBgn0034210 | CG6568 | -0.671776678 | 0.02164287 | 1 |

| | | | | | |
|---|---|---|---|---|---|
| LD07278 | FBgn0029733 | CG6927 | -0.670267006 | 0.02156494 | 1 |
| RE52148 | FBgn0036775 | CG5147 | -0.669216161 | 0.02156494 | 1 |
| LD12075 | FBgn0030010 | CG10959 | -0.667886009 | 0.03143603 | 1 |
| RE03116 | FBgn0034059 | CG8320 | -0.66579826 | 0.02644978 | 1 |
| RE29270 | FBgn0004926 | eIF2beta | -0.663574332 | 0.02156494 | 1 |
| GH12681 | FBgn0027619 | eIF3j | -0.662714817 | 0.02156494 | 1 |
| SD08058 | FBgn0038376 | Hmt-1 | -0.662106091 | 0.02156494 | 1 |
| RE07468 | FBgn0028694 | Rpn11 | -0.661114326 | 0.02156494 | 1 |
| RE71331 | FBgn0037890 | CG17734 | -0.660069648 | 0.02156494 | 1 |
| RE60377 | FBgn0020248 | stet | -0.656809295 | 0.02156494 | 1 |
| GH06781 | FBgn0034390 | CG15093 | -0.654214618 | 0.04434426 | 1 |
| RE59232 | FBgn0051249 | CG31249 | -0.651198338 | 0.02156494 | 1 |
| RH57257 | FBgn0030968 | CG7322 | -0.650882031 | 0.02404547 | 1 |
| RE21270 | FBgn0040477 | cid | -0.648680753 | 0.0273865 | 1 |
| AT12688 | FBgn0030660 | CG8097 | -0.647744446 | 0.02589993 | 1 |
| GH16593 | FBgn0036911 | Fibp | -0.647638561 | 0.02618235 | 1 |
| LD34181 | FBgn0020616 | SA | -0.647472131 | 0.026577 | 1 |
| GM01572 | FBgn0005390 | fs(1)M3 | -0.644130935 | 0.02677632 | 1 |
| RE14873 | FBgn0016053 | pgc | -0.643761918 | 0.02156494 | 1 |
| RH48308 | FBgn0250848 | 26-29-p | -0.638255625 | 0.02156494 | 1 |
| SD10674 | FBgn0260388 | CG42514 | -0.637789508 | 0.02333304 | 1 |
| GH25181 | FBgn0032006 | Pvr | -0.636860522 | 0.0470559 | 1 |
| GH25181 | FBgn0000056 | Adhr | -0.636860522 | 0.0470559 | 1 |
| GH25181 | FBgn0000055 | Adh | -0.636860522 | 0.0470559 | 1 |
| GM13388 | FBgn0032451 | spict | -0.636704181 | 0.02176486 | 1 |
| GH20064 | FBgn0031987 | CG12375 | -0.634241772 | 0.02156494 | 1 |
| AT26759 | FBgn0034982 | Naa35 | -0.63173949 | 0.03511727 | 1 |
| SD01519 | FBgn0027094 | AlaRS | -0.631617619 | 0.02156494 | 1 |
| GM08654 | FBgn0033209 | CG12107 | -0.630885104 | 0.02176486 | 1 |
| RE75106 | FBgn0035558 | CG11357 | -0.630801198 | 0.02326108 | 1 |
| GH01314 | FBgn0266268 | FeCH | -0.629631303 | 0.02225 | 1 |
| GM04207 | FBgn0022344 | CG10340 | -0.626584345 | 0.04137628 | 1 |
| RE43986 | FBgn0033374 | CG13741 | -0.625435143 | 0.02238464 | 1 |
| GH09241 | FBgn0038407 | CG6126 | -0.623151839 | 0.02677632 | 1 |
| RE36140 | FBgn0036476 | sstn | -0.623013435 | 0.02912716 | 1 |
| LD42724 | FBgn0020622 | Pi3K21B | -0.622437389 | 0.02156494 | 1 |
| LD27313 | FBgn0033635 | Prip | -0.622295778 | 0.02618235 | 1 |
| RE29825 | FBgn0036769 | Tsp74F | -0.620583339 | 0.02156494 | 1 |
| AT27586 | FBgn0027903 | CG12018 | -0.620206009 | 0.02941292 | 1 |
| RH60845 | FBgn0263773 | fok | -0.620108967 | 0.02156494 | 1 |
| RE69336 | FBgn0033871 | CG13339 | -0.619572476 | 0.04850883 | 1 |
| GH03577 | FBgn0034314 | nopo | -0.618187516 | 0.02333304 | 1 |
| RE24422 | FBgn0039929 | CG11076 | -0.617715956 | 0.0258032 | 1 |
| RE20606 | FBgn0283442 | vas | -0.617711841 | 0.02156494 | 1 |
| AT13263 | FBgn0036486 | Msh6 | -0.617333325 | 0.02156494 | 1 |
| LD28264 | FBgn0022702 | Cht2 | -0.617187808 | 0.026577 | 1 |
| LD35238 | FBgn0086757 | cbs | -0.616472652 | 0.02404547 | 1 |
| SD02282 | FBgn0002948 | nod | -0.616015208 | 0.02176486 | 1 |

| | | | | | |
|---|---|---|---|---|---|
| RH07572 | FBgn0260857 | Bet1 | -0.615290492 | 0.02351948 | 1 |
| LP07553 | FBgn0010497 | dmGlut | -0.61466814 | 0.02595333 | 1 |
| RH09070 | FBgn0029648 | CG3603 | -0.614310865 | 0.02156494 | 1 |
| GH12002 | FBgn0033246 | ACC | -0.613932627 | 0.02225 | 1 |
| LD25466 | FBgn0032450 | CG5776 | -0.613738638 | 0.02156494 | 1 |
| GH07346 | FBgn0033075 | Pld | -0.612508771 | 0.03557713 | 1 |
| RH45308 | FBgn0029854 | CG3566 | -0.612206521 | 0.02351948 | 1 |
| RH56961 | FBgn0034245 | UQCR-6.4 | -0.611703308 | 0.02618235 | 1 |
| AT30160 | FBgn0035691 | CG7386 | -0.610118948 | 0.02156494 | 1 |
| SD07269 | FBgn0086251 | del | -0.605331537 | 0.02176486 | 1 |
| LD41588 | FBgn0011704 | RnrS | -0.60523296 | 0.03143603 | 1 |
| SD05591 | FBgn0034315 | CG5721 | -0.603163568 | 0.02618235 | 1 |
| LD35215 | FBgn0030008 | CG2129 | -0.603052889 | 0.02176486 | 1 |
| GM32035 | FBgn0053502 | CG33502 | -0.60291974 | 0.02156494 | 1 |
| GM32035 | FBgn0285970 | CG32500 | -0.60291974 | 0.02156494 | 1 |
| GM32035 | FBgn0052857 | CG32857 | -0.60291974 | 0.02156494 | 1 |
| LD10778 | FBgn0027348 | bgm | -0.602455008 | 0.02156494 | 1 |
| SD01793 | FBgn0004432 | Cyp1 | -0.6009445 | 0.03070192 | 1 |
| GH06596 | FBgn0003527 | stil | -0.600822098 | 0.02156494 | 1 |
| RE21178 | FBgn0003082 | phr | -0.59921631 | 0.02869814 | 1 |
| LD35753 | FBgn0026143 | CDC45L | -0.597572591 | 0.02156494 | 1 |
| LD47223 | FBgn0038326 | CG5044 | -0.597172205 | 0.02618235 | 1 |
| GH09650 | FBgn0030671 | CG8134 | -0.595598012 | 0.03406155 | 1 |
| GM16085 | FBgn0035532 | CG15014 | -0.59532425 | 0.02363639 | 1 |
| GM02245 | FBgn0260468 | CG7950 | -0.592437018 | 0.02176486 | 1 |
| GM02245 | FBgn0085346 | CG34317 | -0.592437018 | 0.02176486 | 1 |
| LD27370 | FBgn0266441 | CG45071 | -0.591473935 | 0.03739132 | 1 |
| RE75113 | FBgn0038869 | Smyd5 | -0.589439586 | 0.02941292 | 1 |
| LP11514 | FBgn0038747 | RhoGAP92B | -0.588227919 | 0.02287033 | 1 |
| LP11514 | FBgn0264442 | ab | -0.588227919 | 0.02287033 | 1 |
| SD20045 | FBgn0011566 | Smyd3 | -0.587833873 | 0.02351948 | 1 |
| RE73396 | FBgn0029931 | CG14427 | 0.606734637 | 0.0470559 | 1 |
| RE70031 | FBgn0261873 | sdt | 0.61330926 | 0.02817741 | 1 |
| AT24863 | FBgn0034075 | Asph | 0.613742482 | 0.0291047 | 1 |
| SD10012 | FBgn0004512 | Mdr49 | 0.61747775 | 0.0273865 | 1 |
| LD30945 | FBgn0005590 | scw | 0.628649058 | 0.03303411 | 1 |
| SD18286 | FBgn0033483 | egr | 0.636129992 | 0.02238464 | 1 |
| RE57705 | FBgn0038134 | wntD | 0.646980871 | 0.03667876 | 1 |
| RE13908 | FBgn0003486 | spo | 0.654013428 | 0.02844246 | 1 |
| RE26473 | FBgn0031393 | CG15382 | 0.656454702 | 0.02156494 | 1 |
| LD40883 | FBgn0039286 | dan | 0.677238024 | 0.03403195 | 1 |
| GM19621 | FBgn0004859 | ci | 0.692025986 | 0.04354859 | 1 |
| GH08155 | FBgn0028523 | CG5888 | 0.698714516 | 0.02363639 | 1 |
| RH08872 | FBgn0003067 | Pepck | 0.699216459 | 0.02333304 | 1 |
| RH08872 | FBgn0266488 | CG45087 | 0.699216459 | 0.02333304 | 1 |
| LD22004 | FBgn0004108 | Nrt | 0.720221235 | 0.04041964 | 1 |
| RE16123 | FBgn0040984 | CG4440 | 0.727164119 | 0.02618235 | 1 |
| AT02631 | FBgn0000411 | D | 0.731297979 | 0.02326108 | 1 |

| | | | | | |
|---|---|---|---|---|---|
| LD07701 | FBgn0262739 | AGO1 | 0.736172794 | 0.02156494 | 1 |
| RE22018 | FBgn0051721 | Trim9 | 0.73800084 | 0.03557713 | 1 |
| RE14414 | FBgn0032629 | beat-IIIc | 0.782422636 | 0.04163343 | 1 |
| LD06131 | FBgn0004635 | rho | 0.798845246 | 0.02156494 | 1 |
| RE05137 | FBgn0020257 | Ppa | 0.800570158 | 0.02156494 | 1 |
| GH22922 | FBgn0034602 | Lapsyn | 0.814151981 | 0.02351948 | 1 |
| RE73577 | FBgn0031718 | CG14014 | 0.814348182 | 0.02404547 | 1 |
| LD44491 | FBgn0032693 | Cyp310a1 | 0.826436834 | 0.03449764 | 1 |
| RE26062 | FBgn0001174 | halo | 0.836297923 | 0.02156494 | 1 |
| GH10432 | FBgn0019661 | lncRNA:roX1 | 0.838100367 | 0.03751067 | 1 |
| RE09713 | FBgn0036810 | CG6885 | 0.855009563 | 0.031989 | 1 |
| RE64266 | FBgn0001981 | esg | 0.866465455 | 0.02176486 | 1 |
| RE59904 | FBgn0032377 | CG14937 | 0.867042165 | 0.04850883 | 1 |
| RE03641 | FBgn0060296 | pain | 0.867298429 | 0.03303411 | 1 |
| LD06565 | FBgn0266203 | lncRNA:CR44898 | 0.87166795 | 0.03135973 | 1 |
| RH64455 | FBgn0285951 | mRpS34 | 0.884797803 | 0.02425734 | 1 |
| LD44526 | FBgn0259173 | corn | 0.885447958 | 0.03779407 | 1 |
| RH18694 | FBgn0038243 | CG8066 | 0.904221788 | 0.04094414 | 1 |
| RE48077 | FBgn0037007 | CG5059 | 0.91066904 | 0.03085619 | 1 |
| GH20944 | FBgn0000120 | Arr1 | 0.911020247 | 0.02156494 | 1 |
| RE27042 | FBgn0044049 | Ilp4 | 0.923684901 | 0.02785858 | 1 |
| LD15796 | FBgn0023214 | edl | 0.926745103 | 0.04314341 | 1 |
| RE45895 | FBgn0020377 | Sr-CII | 0.937134387 | 0.03608693 | 1 |
| RE51662 | FBgn0026061 | Mipp1 | 0.937340884 | 0.04202986 | 1 |
| RE30918 | FBgn0001325 | Kr | 0.939791108 | 0.02176486 | 1 |
| GH18470 | FBgn0261836 | Msp300 | 0.956730713 | 0.02948687 | 1 |
| LD16058 | FBgn0003254 | rib | 0.957162873 | 0.04154118 | 1 |
| RE54375 | FBgn0036349 | SNCF | 0.963180924 | 0.02581069 | 1 |
| GH13441 | FBgn0085412 | CG34383 | 0.963567128 | 0.03098436 | 1 |
| RE18236 | FBgn0003683 | term | 0.965607935 | 0.02869814 | 1 |
| AT13327 | FBgn0026063 | KP78b | 0.969425783 | 0.03459731 | 1 |
| AT13327 | FBgn0026064 | KP78a | 0.969425783 | 0.03459731 | 1 |
| LD20987 | FBgn0026319 | Traf4 | 0.973963403 | 0.03303411 | 1 |
| RH55307 | FBgn0013725 | phyl | 0.976440099 | 0.02156494 | 1 |
| RE11206 | FBgn0015773 | NetA | 0.979091299 | 0.02618235 | 1 |
| LD42284 | FBgn0031621 | Elba3 | 0.984595446 | 0.02677632 | 1 |
| LD43816 | FBgn0037992 | CG4702 | 0.996735749 | 0.0424876 | 1 |
| RE24665 | FBgn0000227 | Bsg25A | 1.016080813 | 0.02351948 | 1 |
| LD28132 | FBgn0028519 | hll | 1.02460284 | 0.02618235 | 1 |
| RE35237 | FBgn0003448 | sna | 1.027732194 | 0.02176486 | 1 |
| RE40347 | FBgn0003510 | Sry-alpha | 1.037376595 | 0.026577 | 1 |
| RE40347 | FBgn0003511 | Sry-beta | 1.037376595 | 0.026577 | 1 |
| LD05688 | FBgn0026320 | Tom | 1.039626656 | 0.02363639 | 1 |
| RE50156 | FBgn0032422 | atilla | 1.039741881 | 0.02753015 | 1 |
| GH04704 | FBgn0003430 | slp1 | 1.043285324 | 0.02156494 | 1 |
| RE38281 | FBgn0003326 | sca | 1.049671416 | 0.02156494 | 1 |
| RE59988 | FBgn0028789 | Doc1 | 1.059872121 | 0.02351948 | 1 |
| LD34202 | FBgn0265274 | Inx3 | 1.06244478 | 0.02618235 | 1 |

| | | | | | |
|---|---|---|---|---|---|
| GH03118 | FBgn0028371 | jbug | 1.073043882 | 0.04954315 | 1 |
| RE72284 | FBgn0039283 | danr | 1.094238426 | 0.0459028 | 1 |
| RE47733 | FBgn0004143 | nullo | 1.094757092 | 0.02156494 | 1 |
| RE46351 | FBgn0013272 | Gp150 | 1.095936262 | 0.03576608 | 1 |
| RE34993 | FBgn0010109 | dpn | 1.099895294 | 0.02261744 | 1 |
| GM07658 | FBgn0265276 | l(3)neo38 | 1.104794461 | 0.02618235 | 1 |
| LD39923 | FBgn0000071 | Ama | 1.107575918 | 0.04253873 | 1 |
| LP11484 | FBgn0003731 | Egfr | 1.111813167 | 0.02176486 | 1 |
| RE20251 | FBgn0038028 | CG10035 | 1.12134186 | 0.03459731 | 1 |
| GM01221 | FBgn0263776 | CG43693 | 1.132619695 | 0.02238464 | 1 |
| RE30284 | FBgn0020300 | geko | 1.186797973 | 0.02156494 | 1 |
| LP04345 | FBgn0038071 | Dtg | 1.22622079 | 0.03586406 | 1 |
| HL08053 | FBgn0261560 | Thor | 1.256540568 | 0.02581069 | 1 |
| LD15563 | FBgn0033855 | link | 1.324130933 | 0.02156494 | 1 |
| SD07266 | FBgn0001257 | ImpL2 | 1.363444124 | 0.02200972 | 1 |

| | | | | | |
|---|---|---|---|---|---|
| LP10528 | FBgn0036394 | CG9040 | -1.751197141 | 0.23226005 | 0 |
| LP10764 | FBgn0030600 | hiw | -1.47471468 | 0.21724847 | 0 |
| RE46170 | FBgn0034468 | Obp56a | 1.4655519 | 0.11662204 | 0 |
| RE09034 | FBgn0036336 | ste14 | -1.376071863 | 0.27886405 | 0 |
| LD12604 | FBgn0058191 | CG40191 | -1.230544453 | 0.31858065 | 0 |
| LD26105 | FBgn0035621 | CG10591 | 1.221926579 | 0.37533608 | 0 |
| RE36278 | FBgn0039678 | Obp99a | 1.210852521 | 0.14126395 | 0 |
| HL01248 | FBgn0004646 | ogre | 1.068004314 | 0.06296618 | 0 |
| RE58815 | FBgn0001258 | Ldh | 1.067492393 | 0.13685586 | 0 |
| LP06330 | FBgn0052638 | CG32638 | -1.006885163 | 0.33781529 | 0 |
| LP06330 | FBgn0011284 | RpS4 | -1.006885163 | 0.33781529 | 0 |
| GH10871 | FBgn0010452 | trn | 0.991512124 | 0.06810473 | 0 |
| RH49330 | FBgn0038038 | CG5167 | 0.987534443 | 0.07851049 | 0 |
| LD17549 | FBgn0000182 | BicC | -0.97958328 | 0.23204888 | 0 |
| LD17549 | FBgn0266459 | CG45084 | -0.97958328 | 0.23204888 | 0 |
| GM03559 | FBgn0021967 | ND-PDSW | -0.952715396 | 0.36724764 | 0 |
| LP07339 | FBgn0004425 | LysB | -0.946870436 | 0.25511933 | 0 |
| LP07339 | FBgn0004428 | LysE | -0.946870436 | 0.25511933 | 0 |
| LP07339 | FBgn0004430 | LysS | -0.946870436 | 0.25511933 | 0 |
| LP07339 | FBgn0004427 | LysD | -0.946870436 | 0.25511933 | 0 |
| LD21828 | FBgn0031907 | CG5171 | 0.93410531 | 0.13505454 | 0 |
| GH12078 | FBgn0001090 | bnb | 0.933033703 | 0.05589463 | 0 |
| RE27850 | FBgn0263256 | CG43394 | 0.932021368 | 0.05876215 | 0 |
| GH13134 | FBgn0015777 | nrv2 | 0.927588282 | 0.26152324 | 0 |
| SD16292 | FBgn0025391 | Scgdelta | 0.879445022 | 0.08031814 | 0 |
| SD16292 | FBgn0033458 | CG18446 | 0.879445022 | 0.08031814 | 0 |
| SD16292 | FBgn0036681 | Obp73a | 0.879445022 | 0.08031814 | 0 |
| SD16292 | FBgn0052512 | CG32512 | 0.879445022 | 0.08031814 | 0 |
| SD16292 | FBgn0266847 | lncRNA:CR45309 | 0.879445022 | 0.08031814 | 0 |
| LP10853 | FBgn0036947 | obst-F | -0.846446394 | 0.23906935 | 0 |
| RE45036 | FBgn0262867 | Ptr | 0.844413103 | 0.05893535 | 0 |

| | | | | | |
|---|---|---|---|---|---|
| RH01620 | FBgn0004629 | Cys | 0.842785467 | 0.0817644 | 0 |
| SD09407 | FBgn0033159 | Dscam1 | 0.836907419 | 0.0744522 | 0 |
| LD11258 | FBgn0012037 | Ance | 0.83668105 | 0.1115611 | 0 |
| GH06660 | FBgn0260747 | Chchd2 | -0.825725654 | 0.17394337 | 0 |
| GH06660 | FBgn0031853 | TTLL3B | -0.825725654 | 0.17394337 | 0 |
| LD24387 | FBgn0002526 | LanA | 0.816821519 | 0.46599859 | 0 |
| RE42467 | FBgn0050080 | CG30080 | 0.814307546 | 0.06713041 | 0 |
| RE42467 | FBgn0261538 | CG42662 | 0.814307546 | 0.06713041 | 0 |
| RE07220 | FBgn0002772 | Mlc1 | -0.812726596 | 0.17898253 | 0 |
| GH18004 | FBgn0087002 | apolpp | 0.792872781 | 0.36870325 | 0 |
| LP09874 | FBgn0035040 | CG4741 | -0.790472829 | 0.39339733 | 0 |
| LP08442 | FBgn0029716 | CG3546 | -0.790358257 | 0.09679205 | 0 |
| GH22686 | FBgn0003145 | prd | 0.774818516 | 0.05737483 | 0 |
| GH18014 | FBgn0033188 | Drat | 0.771969008 | 0.1626251 | 0 |
| RE27477 | FBgn0035674 | CG13295 | -0.770251009 | 0.05942862 | 0 |
| LD34564 | FBgn0046253 | CG3502 | 0.767984329 | 0.06286082 | 0 |
| LD26182 | FBgn0001123 | Galphas | 0.767850184 | 0.07559498 | 0 |
| RH45971 | FBgn0260011 | NimC4 | 0.751329929 | 0.14294744 | 0 |
| RH52220 | FBgn0030641 | CG6299 | -0.7403703 | 0.41943816 | 0 |
| LD09584 | FBgn0036381 | CG8745 | 0.738634797 | 0.22082195 | 0 |
| AT18408 | FBgn0038217 | CG14840 | 0.737782412 | 0.27303944 | 0 |
| GM12753 | FBgn0037213 | CG12581 | 0.735461798 | 0.05589463 | 0 |
| LD35237 | FBgn0015380 | drl | 0.735415475 | 0.18965477 | 0 |
| SD05564 | FBgn0039776 | PH4alphaEFB | 0.731657046 | 0.18589461 | 0 |
| RE29477 | FBgn0013953 | Esp | 0.727337324 | 0.07367722 | 0 |
| LD28078 | FBgn0005771 | noc | 0.719497373 | 0.17593513 | 0 |
| GH23040 | FBgn0037238 | CG1090 | 0.714571665 | 0.05944444 | 0 |
| RH08582 | FBgn0002940 | ninaE | -0.711419412 | 0.19895429 | 0 |
| GM12291 | FBgn0021795 | Tapdelta | -0.702769041 | 0.05907437 | 0 |
| RH03540 | FBgn0030040 | CG15347 | 0.698397784 | 0.42981349 | 0 |
| LD19609 | FBgn0001254 | ImpE2 | 0.698040968 | 0.32282288 | 0 |
| GH06693 | FBgn0031824 | CG9547 | -0.697179475 | 0.05025857 | 0 |
| GH11902 | FBgn0004607 | zfh2 | 0.696485108 | 0.14406727 | 0 |
| RE25290 | FBgn0086675 | fne | 0.693685429 | 0.38427112 | 0 |
| RE33779 | FBgn0035969 | CG4476 | -0.685892048 | 0.0757301 | 0 |
| RE40068 | FBgn0003313 | sala | 0.684339153 | 0.05329829 | 0 |
| RE40068 | FBgn0263087 | CG43355 | 0.684339153 | 0.05329829 | 0 |
| GH01880 | FBgn0035762 | Rint1 | -0.678265492 | 0.06455982 | 0 |
| GH01880 | FBgn0085784 | CR41379 | -0.678265492 | 0.06455982 | 0 |
| LP11584 | FBgn0000639 | Fbp1 | -0.678251951 | 0.24617972 | 0 |
| RE25177 | FBgn0033721 | CG13159 | 0.663510099 | 0.58456723 | 0 |
| GH09168 | FBgn0262736 | Vha16-1 | -0.649794637 | 0.25454796 | 0 |
| RE40937 | FBgn0035956 | Doc2 | 0.649596908 | 0.0527752 | 0 |
| LD13210 | FBgn0003520 | stau | 0.648312736 | 0.34069725 | 0 |
| SD09067 | FBgn0039508 | CG3368 | -0.64554703 | 0.08620617 | 0 |
| GM09444 | FBgn0285926 | Imp | 0.645073073 | 0.11638186 | 0 |
| GH08821 | FBgn0266129 | lov | 0.643979639 | 0.06731995 | 0 |
| RH01044 | FBgn0003248 | Rh2 | -0.634383346 | 0.26375028 | 0 |

| | | | | | |
|---|---|---|---|---|---|
| RH08410 | FBgn0040813 | Nplp2 | 0.628417655 | 0.5551644 | 0 |
| LD31354 | FBgn0032683 | kon | 0.623261465 | 0.19464502 | 0 |
| RE30730 | FBgn0035103 | Vdup1 | 0.622216215 | 0.14854672 | 0 |
| RE54320 | FBgn0036352 | CG14110 | 0.621311039 | 0.45220244 | 0 |
| RE05093 | FBgn0033518 | Prx2540-2 | 0.619170375 | 0.07374924 | 0 |
| RE05093 | FBgn0033520 | Prx2540-1 | 0.619170375 | 0.07374924 | 0 |
| RE05093 | FBgn0033521 | CG12896 | 0.619170375 | 0.07374924 | 0 |
| LD42676 | FBgn0037447 | Neurochondrin | 0.613125641 | 0.50166434 | 0 |
| LD45181 | FBgn0038290 | CG6912 | -0.611013967 | 0.25748944 | 0 |
| SD25109 | FBgn0050169 | Brca2 | -0.608386297 | 0.08612406 | 0 |
| LD27656 | FBgn0033337 | CG8272 | -0.605850333 | 0.05383453 | 0 |
| GH01721 | FBgn0031745 | rau | 0.599100251 | 0.14525232 | 0 |
| RE10012 | FBgn0001983 | wor | 0.597457769 | 0.08525374 | 0 |
| RE18076 | FBgn0035257 | CG12011 | 0.596623984 | 0.24794568 | 0 |
| GH08635 | FBgn0063261 | CG31275 | 0.59074178 | 0.11742225 | 0 |
| RH67635 | FBgn0010531 | Ccs | -0.586249081 | 0.09532226 | 0 |
| GH08847 | FBgn0003308 | ry | 0.584341023 | 0.06094564 | 0 |
| GM06015 | FBgn0036815 | HipHop | -0.583137064 | 0.06286082 | 0 |
| RE44714 | FBgn0034797 | nahoda | 0.583073679 | 0.21208966 | 0 |
| LD12257 | FBgn0026430 | Grip84 | -0.582978125 | 0.02156494 | 0 |
| RE70645 | FBgn0002542 | lds | -0.582962797 | 0.02156494 | 0 |
| RE61805 | FBgn0030311 | CG11699 | -0.581917448 | 0.05942862 | 0 |
| LD34147 | FBgn0031313 | CG5080 | 0.581676204 | 0.27527304 | 0 |
| RE09091 | FBgn0035272 | mRpL46 | -0.581100828 | 0.04968819 | 0 |
| RE12569 | FBgn0034356 | CG10924 | 0.581019118 | 0.16751341 | 0 |
| RE30055 | FBgn0040529 | COX7A | -0.580996779 | 0.02618235 | 0 |
| LD42113 | FBgn0004901 | Prat | -0.578485659 | 0.09535834 | 0 |
| RE28551 | FBgn0015946 | grim | -0.578127318 | 0.04041964 | 0 |
| RE28551 | FBgn0267637 | lncRNA:CR45975 | -0.578127318 | 0.04041964 | 0 |
| GH10595 | FBgn0022160 | Gpo1 | 0.578033745 | 0.04079919 | 0 |
| RE11345 | FBgn0004567 | slp2 | 0.57725151 | 0.04749119 | 0 |
| GH11432 | FBgn0034724 | babos | 0.577147699 | 0.10797536 | 0 |
| SD01527 | FBgn0267975 | vib | -0.576413587 | 0.02948687 | 0 |
| RE70353 | FBgn0264855 | AP-2alpha | -0.575266174 | 0.0259526 | 0 |
| LD47348 | FBgn0262029 | d | 0.575172684 | 0.04011285 | 0 |
| GM01136 | FBgn0035443 | CG12010 | -0.574305041 | 0.03143603 | 0 |
| GM14286 | FBgn0029958 | Pdp | -0.573580638 | 0.02761233 | 0 |
| SD14666 | FBgn0036648 | CG4098 | -0.57327885 | 0.0878206 | 0 |
| GH14509 | FBgn0004456 | mew | 0.572136112 | 0.05677394 | 0 |
| RE31178 | FBgn0037492 | CG10050 | -0.571294108 | 0.04066054 | 0 |
| GH08192 | FBgn0029801 | CG15771 | -0.571252811 | 0.02326108 | 0 |
| LD15705 | FBgn0053095 | CG33095 | -0.57046433 | 0.43839377 | 0 |
| LD15705 | FBgn0053096 | CG33096 | -0.57046433 | 0.43839377 | 0 |
| RH55216 | FBgn0035144 | Kah | -0.566429013 | 0.43026786 | 0 |
| RH18728 | FBgn0031629 | Clect27 | 0.566307828 | 0.44538287 | 0 |
| RE56869 | FBgn0033210 | U2A | -0.565875122 | 0.03776809 | 0 |
| RE26521 | FBgn0037917 | wkd | -0.565706056 | 0.04434426 | 0 |
| LD23532 | FBgn0037728 | CG16817 | -0.565366942 | 0.02176486 | 0 |

| | | | | | |
|---|---|---|---|---|---|
| SD10769 | FBgn0034447 | CG7744 | -0.565308145 | 0.0258032 | 0 |
| RE03155 | FBgn0004959 | phm | 0.564736035 | 0.07367722 | 0 |
| SD20009 | FBgn0029689 | CG6428 | -0.564678684 | 0.0273865 | 0 |
| RE26327 | FBgn0002466 | sti | 0.563232489 | 0.04790739 | 0 |
| GH08782 | FBgn0267972 | Der-1 | -0.561586392 | 0.0273865 | 0 |
| LD25202 | FBgn0001291 | Jra | -0.560834188 | 0.10108162 | 0 |
| GM23116 | FBgn0029986 | CG15332 | 0.560526718 | 0.14921756 | 0 |
| SD20823 | FBgn0015000 | betaggt-I | -0.560122649 | 0.02156494 | 0 |
| RE70695 | FBgn0062978 | CG31808 | 0.559051421 | 0.56419565 | 0 |
| GH05723 | FBgn0029818 | GAA1 | -0.558778256 | 0.02363639 | 0 |
| RH34413 | FBgn0036334 | CG11267 | -0.55816146 | 0.0470559 | 0 |
| GH10876 | FBgn0000477 | DNaseII | 0.557857456 | 0.07410495 | 0 |
| LP07775 | FBgn0038881 | CG16791 | -0.557747738 | 0.02425094 | 0 |
| LD47410 | FBgn0259682 | Jabba | -0.557491658 | 0.02618235 | 0 |
| RH19667 | FBgn0030178 | CG2974 | -0.556858511 | 0.026577 | 0 |
| LD35155 | FBgn0030787 | CG9609 | -0.556234895 | 0.02176486 | 0 |
| LP01642 | FBgn0030326 | CG2444 | -0.556079198 | 0.02363639 | 0 |
| SD20696 | FBgn0262719 | CG43163 | 0.55550935 | 0.04154118 | 0 |
| GH13810 | FBgn0035153 | ebd1 | -0.554419255 | 0.02933736 | 0 |
| RE58433 | FBgn0031217 | CG11377 | -0.553887301 | 0.024443 | 0 |
| AT26381 | FBgn0026666 | MagR | -0.55303385 | 0.0470559 | 0 |
| RE14241 | FBgn0038166 | CG9588 | -0.552503536 | 0.0544057 | 0 |
| GM08252 | FBgn0267704 | lncRNA:flam | 0.552421043 | 0.13970977 | 0 |
| LP10995 | FBgn0002865 | Mst98Ca | -0.552217152 | 0.27703693 | 0 |
| LP10995 | FBgn0004171 | Mst98Cb | -0.552217152 | 0.27703693 | 0 |
| RE48052 | FBgn0030400 | CG11138 | -0.547354138 | 0.19554083 | 0 |
| LD24746 | FBgn0003401 | shu | -0.546865331 | 0.02519839 | 0 |
| RE28342 | FBgn0261284 | bou | 0.546009398 | 0.10774879 | 0 |
| GM02347 | FBgn0026718 | fu12 | -0.54571575 | 0.11882441 | 0 |
| SD02150 | FBgn0030242 | sofe | -0.545496402 | 0.02662448 | 0 |
| RE45331 | FBgn0263112 | Mitf | -0.545129732 | 0.07088605 | 0 |
| RE61294 | FBgn0037612 | CG8112 | -0.545124181 | 0.02563869 | 0 |
| LD32130 | FBgn0010389 | htl | 0.544716311 | 0.08865723 | 0 |
| RE14803 | FBgn0038150 | yellow-e3 | 0.544658366 | 0.14854672 | 0 |
| GM02466 | FBgn0014455 | Ahcy | -0.54360772 | 0.03619285 | 0 |
| LP08967 | FBgn0032473 | kmg | -0.541206348 | 0.36885296 | 0 |
| RE15818 | FBgn0030451 | CG15717 | -0.539977873 | 0.0222489 | 0 |
| GM13640 | FBgn0259676 | DNApol-alpha60 | -0.539938752 | 0.02753015 | 0 |
| RE06205 | FBgn0026634 | ldlCp | -0.539208457 | 0.02644978 | 0 |
| AT02038 | FBgn0032047 | CG13088 | -0.536823273 | 0.02644644 | 0 |
| LD21844 | FBgn0000228 | Bsg25D | 0.536819847 | 0.03303411 | 0 |
| RH09887 | FBgn0039682 | Obp99c | 0.536674058 | 0.24056835 | 0 |
| RH06707 | FBgn0028978 | trbl | 0.53579928 | 0.07367722 | 0 |
| RE27241 | FBgn0037960 | mthl5 | -0.53489285 | 0.05623648 | 0 |
| RH73529 | FBgn0261068 | LSm7 | -0.534574719 | 0.03039639 | 0 |
| SD07362 | FBgn0033476 | oys | -0.534247834 | 0.02396533 | 0 |
| LD46156 | FBgn0034583 | CG10527 | -0.533940943 | 0.02363639 | 0 |
| HL08023 | FBgn0030317 | CG1561 | -0.533871221 | 0.03751067 | 0 |

| | | | | | |
|---|---|---|---|---|---|
| RE01888 | FBgn0038128 | Ravus | -0.533047447 | 0.04641137 | 0 |
| LD13052 | FBgn0028421 | Kap3 | -0.532053313 | 0.02703303 | 0 |
| LD37189 | FBgn0003744 | trc | -0.530453032 | 0.0358459 | 0 |
| RH59033 | FBgn0033451 | CG1665 | -0.530371727 | 0.02326108 | 0 |
| GH13824 | FBgn0015270 | Orc2 | -0.52981007 | 0.0246024 | 0 |
| RH69145 | FBgn0024251 | bbx | -0.52887516 | 0.02434869 | 0 |
| HL01706 | FBgn0259111 | Ndae1 | 0.52800427 | 0.0797578 | 0 |
| RE69770 | FBgn0002631 | E(spl)m5-HLH | 0.527807072 | 0.12070859 | 0 |
| LP12032 | FBgn0014848 | Eig71Eh | -0.527682647 | 0.24875122 | 0 |
| RE08009 | FBgn0005696 | DNApol-alpha73 | -0.527581309 | 0.02176486 | 0 |
| SD10961 | FBgn0038429 | CG14882 | -0.527481508 | 0.02200659 | 0 |
| RE25878 | FBgn0015949 | hrg | 0.527033133 | 0.03322374 | 0 |
| RE68083 | FBgn0038609 | Nup43 | -0.525653962 | 0.02817741 | 0 |
| LD37241 | FBgn0037203 | slif | -0.524776681 | 0.02293673 | 0 |
| GM08848 | FBgn0037810 | sle | -0.524112309 | 0.06006631 | 0 |
| AT13028 | FBgn0032390 | dgt2 | -0.523698952 | 0.04311416 | 0 |
| RE25548 | FBgn0030257 | SmydA-4 | -0.523551374 | 0.16222819 | 0 |
| LD41062 | FBgn0000454 | Dip-B | -0.522806778 | 0.02373881 | 0 |
| SD14915 | FBgn0037305 | CG12173 | -0.522399526 | 0.02618235 | 0 |
| LP04238 | FBgn0266719 | stac | -0.52144273 | 0.16681899 | 0 |
| LP04238 | FBgn0031653 | Jon25Biii | -0.52144273 | 0.16681899 | 0 |
| RE39556 | FBgn0038099 | CG7091 | -0.521091073 | 0.03303411 | 0 |
| RE22242 | FBgn0260653 | serp | 0.520367618 | 0.43941393 | 0 |
| GM14814 | FBgn0035593 | CG4603 | -0.519774072 | 0.036157 | 0 |
| SD25546 | FBgn0266282 | jnj | -0.519407272 | 0.02477928 | 0 |
| LD27255 | FBgn0086785 | Vps36 | -0.518973527 | 0.03751067 | 0 |
| GH14974 | FBgn0036366 | JMJD7 | -0.518936919 | 0.04354859 | 0 |
| RH36582 | FBgn0036292 | CG10646 | -0.517524513 | 0.0567557 | 0 |
| LP07247 | FBgn0010316 | dap | -0.516597083 | 0.04339964 | 0 |
| RE28509 | FBgn0032682 | grnd | 0.516097103 | 0.13624799 | 0 |
| GH07940 | FBgn0034464 | CG11018 | 0.516028677 | 0.06712449 | 0 |
| LD34406 | FBgn0264694 | mgr | -0.515719907 | 0.02920323 | 0 |
| GH04557 | FBgn0030679 | CG8206 | -0.514922585 | 0.08612406 | 0 |
| RE59335 | FBgn0002561 | l(1)sc | 0.514883018 | 0.09008791 | 0 |
| RE72863 | FBgn0032031 | CG13390 | -0.514391677 | 0.0394073 | 0 |
| GM13608 | FBgn0026263 | bip1 | -0.514157772 | 0.03672939 | 0 |
| LD40483 | FBgn0260985 | RfC4 | -0.514138084 | 0.02353851 | 0 |
| RE55344 | FBgn0035904 | GstO3 | -0.513224181 | 0.10126494 | 0 |
| LP12374 | FBgn0261283 | SREBP | -0.512831778 | 0.02644978 | 0 |
| AT28119 | FBgn0039258 | beta4GalT7 | -0.511599426 | 0.02363639 | 0 |
| LD22087 | FBgn0027512 | CG10254 | -0.510877143 | 0.02425734 | 0 |
| RE04047 | FBgn0030792 | CG4789 | -0.510170535 | 0.06178355 | 0 |
| RE17330 | FBgn0022343 | CG3760 | -0.50981141 | 0.03376695 | 0 |
| SD15934 | FBgn0032781 | RtcB | -0.50919211 | 0.02618235 | 0 |
| RE27943 | FBgn0034924 | CG17658 | -0.50910106 | 0.02625237 | 0 |
| LD47919 | FBgn0010431 | mtrm | -0.509096848 | 0.05130412 | 0 |
| LD08841 | FBgn0036494 | Toll-6 | 0.508611085 | 0.0780263 | 0 |
| GH22341 | FBgn0039135 | CG13603 | -0.508288236 | 0.02595333 | 0 |

| | | | | | |
|---|---|---|---|---|---|
| SD22445 | FBgn0083123 | Uhg5 | 0.506713829 | 0.02417405 | 0 |
| SD05389 | FBgn0029003 | mab-21 | 0.506636783 | 0.19562813 | 0 |
| RE32504 | FBgn0038989 | CG6937 | -0.50653736 | 0.05630422 | 0 |
| RE67340 | FBgn0032897 | CG9336 | 0.50501488 | 0.4754937 | 0 |
| RE63412 | FBgn0058045 | CG40045 | -0.503328525 | 0.05495124 | 0 |
| LD39537 | FBgn0035955 | CG5194 | -0.503247839 | 0.0973297 | 0 |
| GH17740 | FBgn0034655 | CG10307 | -0.503000072 | 0.11742225 | 0 |
| LD23664 | FBgn0030817 | CG4991 | -0.502552102 | 0.02333304 | 0 |
| GH16429 | FBgn0033272 | RagC-D | -0.502302749 | 0.04355235 | 0 |
| RE28533 | FBgn0011774 | Irbp | -0.501379279 | 0.02351948 | 0 |
| LD05592 | FBgn0263240 | Coop | 0.501345751 | 0.0358459 | 0 |
| RE27552 | FBgn0062449 | CG13197 | -0.501287178 | 0.0817644 | 0 |
| RE04770 | FBgn0035208 | CG9184 | 0.500999048 | 0.41832775 | 0 |
| RH09491 | FBgn0036393 | CG17362 | 0.500203223 | 0.02618235 | 0 |
| LD19034 | FBgn0061515 | endos | -0.499579589 | 0.02644978 | 0 |
| LD19710 | FBgn0001075 | ft | 0.498429252 | 0.19264462 | 0 |
| GH13924 | FBgn0037378 | CG2046 | -0.497777401 | 0.02618235 | 0 |
| SD09860 | FBgn0031020 | Naa15-16 | -0.496932349 | 0.02333304 | 0 |
| AT26957 | FBgn0029853 | CG3781 | -0.495818035 | 0.11465069 | 0 |
| LP01956 | FBgn0050159 | CG30159 | -0.494750042 | 0.05942862 | 0 |
| LD09568 | FBgn0050020 | CG30020 | -0.493463461 | 0.46436889 | 0 |
| RE19603 | FBgn0015614 | CanB2 | -0.492773551 | 0.05096655 | 0 |
| RH48922 | FBgn0083991 | CG34155 | 0.491417425 | 0.02937161 | 0 |
| RE13621 | FBgn0283681 | Tcs3 | -0.491285629 | 0.0414665 | 0 |
| GH27372 | FBgn0003891 | tud | -0.491009199 | 0.24875122 | 0 |
| GM14788 | FBgn0040309 | Jafrac1 | -0.490305207 | 0.0258032 | 0 |
| RE38137 | FBgn0086679 | p | -0.490009443 | 0.02477928 | 0 |
| LD47993 | FBgn0038889 | Fancm | -0.489824901 | 0.05841216 | 0 |
| SD17927 | FBgn0026738 | CG7857 | -0.489569803 | 0.03751067 | 0 |
| RE07481 | FBgn0035203 | CG9149 | -0.487704597 | 0.02618235 | 0 |
| RE03177 | FBgn0032910 | CG9265 | -0.487640542 | 0.02363639 | 0 |
| GH03573 | FBgn0264975 | Nrg | 0.486102395 | 0.1246311 | 0 |
| GH12341 | FBgn0003896 | tup | 0.485665514 | 0.10316195 | 0 |
| RE41485 | FBgn0036547 | CG17032 | 0.484361324 | 0.29746573 | 0 |
| AT13922 | FBgn0266449 | Kr-h2 | -0.483866643 | 0.02678065 | 0 |
| RE09158 | FBgn0268063 | ltl | 0.483806956 | 0.06995214 | 0 |
| GM27569 | FBgn0033528 | trsn | -0.483663692 | 0.04305124 | 0 |
| LD07878 | FBgn0036240 | CG6928 | -0.482849205 | 0.02530044 | 0 |
| SD13634 | FBgn0024194 | rasp | -0.48257754 | 0.04740787 | 0 |
| GH14316 | FBgn0038115 | CG7966 | -0.480926301 | 0.05025857 | 0 |
| RH36186 | FBgn0035982 | CG4461 | -0.479306107 | 0.03585553 | 0 |
| RE30433 | FBgn0036787 | CG4306 | 0.478576541 | 0.33272747 | 0 |
| LD24495 | FBgn0284436 | CCT8 | -0.478220641 | 0.03376695 | 0 |
| RE55814 | FBgn0035907 | GstO1 | -0.477940774 | 0.06025942 | 0 |
| SD04231 | FBgn0033980 | Cyp6a20 | -0.477682235 | 0.02454623 | 0 |
| LD28935 | FBgn0026361 | Sep5 | 0.476993603 | 0.0575047 | 0 |
| SD16707 | FBgn0041147 | ida | -0.476699631 | 0.22082195 | 0 |
| RH47419 | FBgn0033970 | CG10205 | -0.475965625 | 0.02703303 | 0 |

| | | | | | |
|---|---|---|---|---|---|
| GH10845 | FBgn0030251 | CG2145 | 0.475937653 | 0.06025942 | 0 |
| SD15916 | FBgn0037206 | CG12768 | -0.475777454 | 0.33954823 | 0 |
| GH23075 | FBgn0004381 | Klp68D | -0.475298666 | 0.03431035 | 0 |
| GH22488 | FBgn0005616 | msl-2 | 0.475053035 | 0.02844246 | 0 |
| RE40431 | FBgn0036606 | CG13060 | 0.474265414 | 0.12499919 | 0 |
| RE40431 | FBgn0036605 | CG13041 | 0.474265414 | 0.12499919 | 0 |
| AT09986 | FBgn0031644 | CG15625 | 0.474034845 | 0.03508936 | 0 |
| RE21078 | FBgn0031969 | pes | -0.473857701 | 0.02563869 | 0 |
| SD05592 | FBgn0037619 | CG8159 | -0.473795252 | 0.09541205 | 0 |
| GH18603 | FBgn0036136 | Ufd1-like | -0.472940511 | 0.02991809 | 0 |
| GM14334 | FBgn0002069 | AspRS | -0.47285563 | 0.0273865 | 0 |
| LP07340 | FBgn0263120 | Acsl | -0.472714902 | 0.04850883 | 0 |
| RE21038 | FBgn0004872 | piwi | -0.471959424 | 0.03511727 | 0 |
| LP11234 | FBgn0031717 | Oscillin | 0.471955951 | 0.09260753 | 0 |
| LD09231 | FBgn0038476 | kuk | 0.469859813 | 0.06925767 | 0 |
| LD32196 | FBgn0038304 | CG12241 | -0.468409093 | 0.02792151 | 0 |
| SD03863 | FBgn0039175 | beta-PheRS | -0.468250912 | 0.07367722 | 0 |
| GH12235 | FBgn0003353 | sei | -0.467911685 | 0.03557713 | 0 |
| RE68138 | FBgn0031662 | CG3792 | -0.467881561 | 0.0424876 | 0 |
| RE58665 | FBgn0039965 | CG13865 | -0.467677005 | 0.06826096 | 0 |
| RE39082 | FBgn0001253 | ImpE1 | 0.466664773 | 0.17136716 | 0 |
| RH17365 | FBgn0030638 | CG11655 | -0.466432035 | 0.02991809 | 0 |
| SD06673 | FBgn0035295 | Cnb | -0.465533691 | 0.02792151 | 0 |
| RE06857 | FBgn0030786 | mRpL22 | -0.465416388 | 0.03689027 | 0 |
| GH13901 | FBgn0034173 | CG9010 | -0.465059747 | 0.07000688 | 0 |
| LD30448 | FBgn0035630 | CG10576 | -0.464590386 | 0.02700894 | 0 |
| LD12611 | FBgn0053099 | CG33099 | 0.464312059 | 0.04777097 | 0 |
| RE28781 | FBgn0031182 | Cyp6t1 | -0.463151406 | 0.04253873 | 0 |
| GH05885 | FBgn0035889 | mkg-p | -0.462929002 | 0.07367722 | 0 |
| GH05885 | FBgn0053057 | CG33057 | -0.462929002 | 0.07367722 | 0 |
| AT26639 | FBgn0265082 | Cdep | 0.462921917 | 0.1366984 | 0 |
| RE26319 | FBgn0038315 | CG14866 | -0.462568255 | 0.13966772 | 0 |
| RE01566 | FBgn0032598 | ChLD3 | -0.461731688 | 0.0312799 | 0 |
| GH16740 | FBgn0030484 | GstT4 | 0.461495803 | 0.22390724 | 0 |
| LP07668 | FBgn0035656 | CG10479 | 0.46127198 | 0.04311416 | 0 |
| LP08858 | FBgn0034897 | Sesn | -0.461134816 | 0.30100331 | 0 |
| LP08858 | FBgn0036362 | CG10725 | -0.461134816 | 0.30100331 | 0 |
| LD24639 | FBgn0259749 | mmy | -0.460833378 | 0.02792151 | 0 |
| LD20271 | FBgn0010314 | Cks30A | -0.460098499 | 0.05121332 | 0 |
| RE04029 | FBgn0086695 | hd | -0.459820664 | 0.03445935 | 0 |
| SD10334 | FBgn0010812 | unc-45 | -0.459101747 | 0.0757301 | 0 |
| RE09535 | FBgn0004914 | Hnf4 | -0.458717669 | 0.06022567 | 0 |
| SD06635 | FBgn0030073 | CG10962 | -0.458588147 | 0.0358459 | 0 |
| RE34149 | FBgn0265767 | zyd | -0.458580529 | 0.03119151 | 0 |
| SD06594 | FBgn0015245 | Hsp60A | -0.458181336 | 0.0273865 | 0 |
| RH65902 | FBgn0031189 | CG14618 | -0.457627207 | 0.02662448 | 0 |
| GH14454 | FBgn0019650 | toy | 0.457552768 | 0.07853066 | 0 |
| RE14758 | FBgn0015615 | SMC3 | -0.457078853 | 0.04079919 | 0 |

| | | | | | |
|---|---|---|---|---|---|
| RE13606 | FBgn0036271 | Pbgs | -0.456613791 | 0.03956255 | 0 |
| AT14009 | FBgn0283536 | Vha13 | -0.456147053 | 0.05589463 | 0 |
| RE34385 | FBgn0086779 | step | -0.455386331 | 0.06712449 | 0 |
| LD15043 | FBgn0025388 | CG12179 | -0.455122126 | 0.026577 | 0 |
| GH02075 | FBgn0031263 | Tspo | -0.454197993 | 0.05329829 | 0 |
| RE36967 | FBgn0035954 | Doc3 | 0.454030808 | 0.06713041 | 0 |
| RE32988 | FBgn0039156 | CG6178 | -0.453064492 | 0.0259526 | 0 |
| LD38576 | FBgn0034275 | CG5002 | 0.452825351 | 0.03135973 | 0 |
| RE61780 | FBgn0037949 | CG17360 | -0.452799275 | 0.07380561 | 0 |
| AT12613 | FBgn0031620 | CG11929 | 0.452680988 | 0.0312799 | 0 |
| RE48347 | FBgn0015793 | Rab19 | -0.451708117 | 0.04279742 | 0 |
| RE06142 | FBgn0004510 | Ets97D | -0.451346653 | 0.04603275 | 0 |
| LD33460 | FBgn0010238 | Lac | 0.449292665 | 0.04094414 | 0 |
| AT09679 | FBgn0030206 | CG2889 | -0.448885517 | 0.08836172 | 0 |
| GH09538 | FBgn0036763 | TrpRS-m | -0.448609117 | 0.02728078 | 0 |
| SD25906 | FBgn0036028 | CG16717 | -0.448492964 | 0.02948687 | 0 |
| GH11174 | FBgn0052412 | QC | -0.448420036 | 0.04154118 | 0 |
| RE61427 | FBgn0011592 | fra | 0.448327664 | 0.05025857 | 0 |
| LD24323 | FBgn0001091 | Gapdh1 | -0.448216318 | 0.02677632 | 0 |
| SD19372 | FBgn0031489 | CG17224 | 0.447922025 | 0.02933736 | 0 |
| GH13039 | FBgn0028552 | gammaSnap1 | -0.447872258 | 0.0862875 | 0 |
| RH73277 | FBgn0032160 | CG4598 | -0.447832811 | 0.04079919 | 0 |
| GH22170 | FBgn0266801 | CG45263 | 0.447583485 | 0.10152861 | 0 |
| GH26851 | FBgn0038610 | CG7675 | 0.44715246 | 0.5539731 | 0 |
| LD47602 | FBgn0040475 | SH3PX1 | -0.446589163 | 0.03181293 | 0 |
| GM31575 | FBgn0039337 | MED28 | -0.446266045 | 0.24417029 | 0 |
| LD46938 | FBgn0039013 | CG4813 | -0.446224107 | 0.11005477 | 0 |
| GM09534 | FBgn0267912 | CanA-14F | 0.446086676 | 0.09460365 | 0 |
| RE01016 | FBgn0051040 | Cog7 | -0.445655816 | 0.10434168 | 0 |
| LD47649 | FBgn0053526 | PNUTS | -0.445405251 | 0.03484105 | 0 |
| RE50384 | FBgn0051028 | CG31028 | 0.444998609 | 0.10646825 | 0 |
| GM23988 | FBgn0263873 | sick | 0.444295553 | 0.18882961 | 0 |
| LD21403 | FBgn0027529 | tapas | -0.44391455 | 0.03039639 | 0 |
| RE09661 | FBgn0032803 | CG13082 | 0.443847758 | 0.44121583 | 0 |
| LD33749 | FBgn0037549 | CG7878 | -0.443609231 | 0.05422973 | 0 |
| SD13619 | FBgn0035608 | blanks | 0.442027898 | 0.03602995 | 0 |
| GH06479 | FBgn0031637 | mxt | -0.441276579 | 0.036157 | 0 |
| GH28557 | FBgn0036918 | Pfdn6 | -0.440897256 | 0.08769971 | 0 |
| RH47329 | FBgn0033754 | Ak6 | -0.44087862 | 0.13466737 | 0 |
| SD03267 | FBgn0260463 | Unc-115b | 0.440875936 | 0.10277034 | 0 |
| SD03267 | FBgn0053105 | p24-2 | 0.440875936 | 0.10277034 | 0 |
| RH40148 | FBgn0035964 | Dhpr | -0.440869224 | 0.04154118 | 0 |
| RE58281 | FBgn0038592 | CG18599 | -0.440757751 | 0.20926155 | 0 |
| GH03305 | FBgn0032350 | CG6287 | -0.440656073 | 0.02618235 | 0 |
| GH11366 | FBgn0053470 | CG33470 | -0.440143559 | 0.25618484 | 0 |
| GH11366 | FBgn0283462 | IMPPP | -0.440143559 | 0.25618484 | 0 |
| RH46018 | FBgn0036290 | CG10638 | -0.439698514 | 0.03576608 | 0 |
| GH08370 | FBgn0035538 | DopEcR | -0.439559204 | 0.28912309 | 0 |

| | | | | | |
|---|---|---|---|---|---|
| RH57745 | FBgn0036853 | mRpL21 | -0.439400478 | 0.06296618 | 0 |
| LP11415 | FBgn0030357 | Sclp | -0.43934023 | 0.03536528 | 0 |
| GH15220 | FBgn0036620 | CG4842 | -0.4388406 | 0.13645262 | 0 |
| HL01707 | FBgn0028695 | Rpn1 | -0.438620332 | 0.06968542 | 0 |
| AT29216 | FBgn0039731 | Sas-6 | -0.438375041 | 0.02869814 | 0 |
| AT03366 | FBgn0261552 | ps | 0.43741826 | 0.10529056 | 0 |
| AT30361 | FBgn0026259 | eIF5B | -0.435693607 | 0.07853066 | 0 |
| AT29938 | FBgn0052407 | CG32407 | 0.435207893 | 0.06885709 | 0 |
| CK02229 | FBgn0261563 | wb | 0.434885349 | 0.19446156 | 0 |
| GM03596 | FBgn0052218 | lncRNA:CR32218 | 0.434664246 | 0.02941292 | 0 |
| LP06006 | FBgn0011653 | mas | 0.434146565 | 0.17147856 | 0 |
| RE40129 | FBgn0063497 | GstE3 | -0.433802199 | 0.05876215 | 0 |
| RH02936 | FBgn0029828 | CG6067 | -0.433792328 | 0.05737143 | 0 |
| LD20383 | FBgn0038012 | CG10013 | 0.433566689 | 0.09751361 | 0 |
| RE18811 | FBgn0030510 | CG12177 | 0.433139595 | 0.06731995 | 0 |
| GH17932 | FBgn0035044 | CG3663 | -0.433023472 | 0.07570279 | 0 |
| LP10092 | FBgn0038810 | Srp72 | -0.432966394 | 0.09635085 | 0 |
| GH12050 | FBgn0022153 | l(2)k05819 | -0.432906508 | 0.04850883 | 0 |
| RE57810 | FBgn0037061 | CG12975 | -0.43279262 | 0.12781374 | 0 |
| AT27571 | FBgn0063298 | asRNA:CR31429 | 0.432493701 | 0.0358459 | 0 |
| LP12257 | FBgn0003328 | scb | 0.432353407 | 0.22807506 | 0 |
| RE66051 | FBgn0285911 | Vps16A | -0.43139006 | 0.05942862 | 0 |
| GM09416 | FBgn0037565 | CG9626 | 0.43012406 | 0.07570279 | 0 |
| RE68977 | FBgn0053052 | CG33052 | -0.42986044 | 0.02811916 | 0 |
| RE13914 | FBgn0029002 | miple2 | 0.429766434 | 0.18253104 | 0 |
| GM22653 | FBgn0024234 | gbb | 0.429707701 | 0.1088733 | 0 |
| GM04147 | FBgn0033486 | dmpd | -0.429611087 | 0.02948687 | 0 |
| GM07659 | FBgn0010470 | Fkbp14 | 0.429018649 | 0.11005477 | 0 |
| LD35474 | FBgn0036752 | Adgf-A | 0.428850633 | 0.11551359 | 0 |
| GM07660 | FBgn0262734 | eIF4H1 | 0.428602544 | 0.15519681 | 0 |
| RE07829 | FBgn0003089 | pip | 0.427746678 | 0.54627099 | 0 |
| RE34508 | FBgn0264493 | rdx | -0.427493386 | 0.07038909 | 0 |
| AT02348 | FBgn0250814 | UQCR-C2 | -0.426337056 | 0.0603552 | 0 |
| GH15747 | FBgn0029137 | Patsas | -0.425880766 | 0.06810473 | 0 |
| RE06257 | FBgn0023515 | CG14814 | -0.425385402 | 0.04154118 | 0 |
| GH04080 | FBgn0283437 | PPO1 | 0.425164596 | 0.3939846 | 0 |
| RE51892 | FBgn0030305 | Uba5 | -0.425105961 | 0.02948687 | 0 |
| RH47542 | FBgn0043012 | AP-2sigma | -0.424943492 | 0.15949841 | 0 |
| RH61816 | FBgn0029608 | CG3091 | -0.424623296 | 0.0634831 | 0 |
| LD29542 | FBgn0261064 | Rbsn-5 | -0.423696113 | 0.11930713 | 0 |
| RE30726 | FBgn0053303 | CG33303 | -0.423674348 | 0.04094414 | 0 |
| LD43023 | FBgn0033677 | CG8321 | -0.423204292 | 0.06885709 | 0 |
| LD10908 | FBgn0031435 | Elba2 | 0.423020724 | 0.05068929 | 0 |
| LD27564 | FBgn0039589 | CG9986 | -0.422402651 | 0.02869814 | 0 |
| LD21576 | FBgn0015268 | Nap1 | -0.421760765 | 0.03070192 | 0 |
| SD02086 | FBgn0035033 | CG3548 | -0.42173155 | 0.02844246 | 0 |
| LD10135 | FBgn0028965 | A16 | -0.420725034 | 0.0780263 | 0 |
| LD24105 | FBgn0029969 | CG10932 | -0.419776397 | 0.09204205 | 0 |

| | | | | | |
|---|---|---|---|---|---|
| RE18302 | FBgn0087007 | bbg | 0.419144508 | 0.04285491 | 0 |
| AT03386 | FBgn0036502 | CG7841 | -0.418865679 | 0.0683056 | 0 |
| LD20423 | FBgn0031483 | CG9641 | -0.418819792 | 0.03037893 | 0 |
| LD24485 | FBgn0004507 | GlyP | -0.417694469 | 0.03770571 | 0 |
| RE68626 | FBgn0011555 | thetaTry | -0.417664201 | 0.05632121 | 0 |
| GH11618 | FBgn0031948 | CG7149 | -0.417555596 | 0.10804827 | 0 |
| RE53130 | FBgn0267824 | PRAS40 | -0.417484413 | 0.02948687 | 0 |
| HL07910 | FBgn0035720 | CG10077 | 0.417017558 | 0.13368437 | 0 |
| AT19154 | FBgn0262467 | Scox | -0.416566611 | 0.03770571 | 0 |
| RE34731 | FBgn0039640 | CG14516 | -0.416518461 | 0.05942862 | 0 |
| LD32760 | FBgn0039851 | mey | 0.416463452 | 0.30013801 | 0 |
| AT13666 | FBgn0004956 | upd1 | 0.415698656 | 0.11005477 | 0 |
| AT31036 | FBgn0027514 | CG1024 | -0.415138889 | 0.05096655 | 0 |
| GH07804 | FBgn0039738 | Mgat2 | 0.415040819 | 0.07367722 | 0 |
| AT18066 | FBgn0031451 | CG9961 | -0.414882826 | 0.03751067 | 0 |
| GH07286 | FBgn0038961 | CG13850 | -0.414831484 | 0.03358705 | 0 |
| GM03811 | FBgn0038683 | CG11779 | -0.414781523 | 0.03376695 | 0 |
| GH01842 | FBgn0002733 | E(spl)mbeta-HLH | 0.414606377 | 0.18472354 | 0 |
| LD07154 | FBgn0028480 | CG17841 | -0.414604171 | 0.33802905 | 0 |
| AT03803 | FBgn0034849 | CG3500 | -0.414603789 | 0.05495124 | 0 |
| RH51533 | FBgn0033591 | CG13216 | 0.414480427 | 0.1115611 | 0 |
| RE63573 | FBgn0051080 | TwdlH | -0.413988503 | 0.09698692 | 0 |
| GH24142 | FBgn0029783 | Sirt4 | -0.413734353 | 0.04803177 | 0 |
| RE14076 | FBgn0036786 | skl | 0.413577745 | 0.15677902 | 0 |
| GH13089 | FBgn0024371 | E2f2 | -0.412382505 | 0.04253873 | 0 |
| SD02665 | FBgn0267792 | rgr | -0.411963677 | 0.13624799 | 0 |
| RH26067 | FBgn0023534 | CG17778 | -0.411582688 | 0.06025942 | 0 |
| RE12105 | FBgn0010399 | Nmdar1 | -0.411165145 | 0.36885296 | 0 |
| GM15148 | FBgn0013733 | shot | 0.411001775 | 0.03751067 | 0 |
| LP08712 | FBgn0037684 | CG8129 | 0.410859685 | 0.0414665 | 0 |
| RE24790 | FBgn0261803 | CG42749 | -0.410331656 | 0.33063114 | 0 |
| RH47993 | FBgn0004864 | hop | -0.409913396 | 0.080407 | 0 |
| GH18634 | FBgn0027948 | msps | -0.409758815 | 0.03930978 | 0 |
| HL01173 | FBgn0023540 | CG3630 | -0.409286639 | 0.1115611 | 0 |
| SD04586 | FBgn0031817 | CG9531 | -0.409157672 | 0.07367722 | 0 |
| RH02643 | FBgn0010014 | CanB | -0.409031976 | 0.06226887 | 0 |
| AT25528 | FBgn0262109 | lncRNA:CR42862 | 0.409025119 | 0.07000688 | 0 |
| RE27587 | FBgn0033981 | Cyp6a21 | -0.408725869 | 0.05002903 | 0 |
| LD21194 | FBgn0259483 | Mob4 | -0.408296597 | 0.03380971 | 0 |
| GH14562 | FBgn0029752 | TrxT | -0.408290178 | 0.43855477 | 0 |
| LD04815 | FBgn0010173 | RpA-70 | -0.408203438 | 0.04434426 | 0 |
| GM31979 | FBgn0039486 | caix | -0.408099152 | 0.03033006 | 0 |
| RE08339 | FBgn0010434 | cora | 0.407737214 | 0.1911631 | 0 |
| GH28550 | FBgn0035542 | DOR | 0.407514715 | 0.20639193 | 0 |
| AT13969 | FBgn0036893 | PIG-F | -0.40666032 | 0.02948687 | 0 |
| RE19838 | FBgn0034301 | CG5756 | -0.406247733 | 0.10417336 | 0 |
| LD28822 | FBgn0025742 | mtm | -0.406124604 | 0.09257793 | 0 |
| SD04868 | FBgn0261274 | Ero1L | -0.405963122 | 0.03143603 | 0 |

| | | | | | |
|---|---|---|---|---|---|
| LD02352 | FBgn0000283 | Cp190 | 0.404989437 | 0.06808125 | 0 |
| RH01747 | FBgn0031646 | CG2837 | 0.404876072 | 0.05956874 | 0 |
| GH04039 | FBgn0031245 | CG3625 | 0.404266591 | 0.17961462 | 0 |
| HL08104 | FBgn0040384 | CG32795 | -0.403857917 | 0.03494848 | 0 |
| GH23035 | FBgn0036198 | crim | -0.403604268 | 0.0578828 | 0 |
| LD45021 | FBgn0261239 | Hr39 | -0.403505277 | 0.0470559 | 0 |
| RE18087 | FBgn0000658 | fj | 0.403236033 | 0.28758766 | 0 |
| GH27834 | FBgn0033232 | CG12159 | -0.402656172 | 0.0299109 | 0 |
| GH06335 | FBgn0023524 | CG3078 | -0.402531569 | 0.26307647 | 0 |
| GH12068 | FBgn0028684 | Rpt5 | -0.402083297 | 0.03667876 | 0 |
| GH05563 | FBgn0025633 | CG13366 | -0.401584441 | 0.09890059 | 0 |
| SD04973 | FBgn0034504 | CG8929 | 0.401450412 | 0.06995214 | 0 |
| RE10132 | FBgn0038147 | CCHa2 | -0.401044621 | 0.2091573 | 0 |
| RE05438 | FBgn0052626 | AMPdeam | -0.400932493 | 0.03829112 | 0 |
| AT07973 | FBgn0002787 | Rpn8 | -0.400787304 | 0.02991809 | 0 |
| LP03459 | FBgn0010473 | tutl | -0.400379991 | 0.04137628 | 0 |
| GM01690 | FBgn0032924 | Nbr | -0.400156195 | 0.05956874 | 0 |
| SD27954 | FBgn0051800 | CG31800 | -0.399668571 | 0.48214468 | 0 |
| GM08788 | FBgn0030479 | Rbp1-like | 0.39946429 | 0.06712449 | 0 |
| GH08610 | FBgn0038475 | Keap1 | 0.399317569 | 0.04163343 | 0 |
| RH67089 | FBgn0029823 | Shmt | -0.399289611 | 0.04311416 | 0 |
| LD46767 | FBgn0004868 | Gdi | -0.398779388 | 0.031989 | 0 |
| LD25748 | FBgn0029903 | pod1 | 0.39823799 | 0.0544057 | 0 |
| AT17959 | FBgn0034467 | CG15128 | 0.397835505 | 0.06539324 | 0 |
| GH26222 | FBgn0260442 | rhea | 0.397342352 | 0.0358459 | 0 |
| LD24195 | FBgn0264962 | Pcf11 | 0.397170371 | 0.03544275 | 0 |
| GH19528 | FBgn0035661 | CG10477 | -0.396977543 | 0.52750038 | 0 |
| SD05215 | FBgn0034180 | Ehbp1 | 0.396273685 | 0.04896084 | 0 |
| AT03867 | FBgn0265296 | Dscam2 | -0.39615978 | 0.05096655 | 0 |
| LP01207 | FBgn0037852 | Tpc1 | -0.395987985 | 0.03785439 | 0 |
| LD47143 | FBgn0028688 | Rpn7 | -0.395832127 | 0.04253873 | 0 |
| RH47711 | FBgn0030659 | CG9215 | -0.395407487 | 0.09603242 | 0 |
| LD23608 | FBgn0023477 | Taldo | -0.395352179 | 0.05689849 | 0 |
| GH15138 | FBgn0040334 | Tsp3A | -0.395240683 | 0.06501956 | 0 |
| GH14535 | FBgn0260458 | PGRP-LD | -0.395090488 | 0.09541205 | 0 |
| GH14535 | FBgn0044419 | Pmi | -0.395090488 | 0.09541205 | 0 |
| LD23371 | FBgn0027504 | CG8878 | 0.39504545 | 0.0757301 | 0 |
| RH44346 | FBgn0002905 | mus308 | -0.394610352 | 0.32095126 | 0 |
| GM04312 | FBgn0085432 | pan | 0.394455552 | 0.05142607 | 0 |
| GH02722 | FBgn0030039 | CG12123 | -0.39427887 | 0.1470603 | 0 |
| LD40177 | FBgn0034488 | CG11208 | -0.393738237 | 0.05942862 | 0 |
| RH27794 | FBgn0000150 | awd | -0.393675097 | 0.21784977 | 0 |
| LD24434 | FBgn0034650 | NC2alpha | -0.393189217 | 0.09715278 | 0 |
| GH23814 | FBgn0029746 | CG15465 | 0.391910978 | 0.03181293 | 0 |
| GH23814 | FBgn0266098 | rg | 0.391910978 | 0.03181293 | 0 |
| GH09045 | FBgn0028487 | f-cup | 0.391901384 | 0.04018448 | 0 |
| LD23309 | FBgn0013732 | sced | -0.391829057 | 0.55218915 | 0 |
| LD37025 | FBgn0263352 | Unr | 0.391384508 | 0.23434738 | 0 |

| | | | | | |
|---|---|---|---|---|---|
| AT06251 | FBgn0038386 | CG5478 | 0.39129874 | 0.07570279 | 0 |
| RH36765 | FBgn0036666 | TSG101 | -0.39111519 | 0.23906935 | 0 |
| GH10582 | FBgn0038966 | pinta | -0.390823968 | 0.04422997 | 0 |
| GH21134 | FBgn0000008 | a | -0.390599969 | 0.37074408 | 0 |
| GH19970 | FBgn0031998 | SLC5A11 | -0.390472977 | 0.0544057 | 0 |
| HL08057 | FBgn0011693 | Pdh | -0.390249922 | 0.10360568 | 0 |
| HL02693 | FBgn0023023 | CRMP | 0.389950718 | 0.27659084 | 0 |
| SD06593 | FBgn0027359 | Tim8 | -0.389772232 | 0.03322164 | 0 |
| RE41071 | FBgn0024236 | foi | -0.389656568 | 0.03576608 | 0 |
| LP01328 | FBgn0004885 | tok | 0.387957728 | 0.1325229 | 0 |
| GH19479 | FBgn0085434 | NaCP60E | -0.38788871 | 0.27774828 | 0 |
| LD42783 | FBgn0019890 | Smg5 | 0.387724091 | 0.0847017 | 0 |
| RE37593 | FBgn0036446 | CG9384 | -0.387566931 | 0.24512465 | 0 |
| RE02878 | FBgn0032217 | CG4972 | -0.387422356 | 0.07570279 | 0 |
| LD43280 | FBgn0023181 | Orc4 | -0.38729143 | 0.10426614 | 0 |
| GH21817 | FBgn0037897 | CG5270 | -0.386849999 | 0.05493179 | 0 |
| LD23155 | FBgn0034585 | Rbpn-5 | -0.385888631 | 0.06885709 | 0 |
| SD19419 | FBgn0033428 | Updo | -0.385685442 | 0.03327429 | 0 |
| GH01724 | FBgn0033179 | p47 | -0.385641433 | 0.05294781 | 0 |
| RH10518 | FBgn0027111 | miple1 | 0.384916559 | 0.51901403 | 0 |
| SD02276 | FBgn0286198 | LKRSDH | 0.384692947 | 0.10966157 | 0 |
| SD03655 | FBgn0035236 | CG12004 | -0.384577844 | 0.0759903 | 0 |
| RE45066 | FBgn0034808 | CG9896 | -0.384508857 | 0.03379152 | 0 |
| RH42041 | FBgn0042083 | Mccc2 | -0.384316972 | 0.04301801 | 0 |
| GH01635 | FBgn0037637 | IscU | -0.383792426 | 0.07570279 | 0 |
| SD03439 | FBgn0031286 | CG3862 | -0.383780787 | 0.0892183 | 0 |
| SD19562 | FBgn0030686 | mRpL3 | -0.383673947 | 0.0358459 | 0 |
| LD07444 | FBgn0031652 | jet | -0.383616015 | 0.05643298 | 0 |
| RH54517 | FBgn0038080 | CG12279 | -0.383565408 | 0.03303411 | 0 |
| RE70806 | FBgn0029082 | hbs | -0.383063049 | 0.15072325 | 0 |
| LD45505 | FBgn0002932 | neur | 0.382890465 | 0.05234953 | 0 |
| RE06388 | FBgn0052194 | lncRNA:CR32194 | 0.382586146 | 0.08743204 | 0 |
| LD22570 | FBgn0027108 | Inx2 | 0.38254958 | 0.06640401 | 0 |
| RE50740 | FBgn0266581 | pit | -0.382501345 | 0.04311416 | 0 |
| GH21316 | FBgn0039349 | Ssadh | -0.382476871 | 0.04790739 | 0 |
| RH07911 | FBgn0039924 | PIP4K | -0.382196645 | 0.05942862 | 0 |
| SD10603 | FBgn0031449 | CG31689 | -0.382145848 | 0.03776809 | 0 |
| LD18769 | FBgn0032161 | CG4594 | -0.382133898 | 0.05942862 | 0 |
| AT09724 | FBgn0037811 | CG12592 | -0.382132479 | 0.0459028 | 0 |
| AT23193 | FBgn0037921 | CG6808 | -0.381426756 | 0.21444299 | 0 |
| RH03206 | FBgn0034405 | Jheh2 | -0.381335813 | 0.09173843 | 0 |
| RH21090 | FBgn0038043 | CG17202 | -0.381186054 | 0.06743902 | 0 |
| RE14415 | FBgn0036716 | CG13728 | -0.381164909 | 0.24204398 | 0 |
| AT15604 | FBgn0050161 | CG30161 | 0.380776707 | 0.06178355 | 0 |
| LD31554 | FBgn0031573 | CG3407 | 0.380273256 | 0.0603552 | 0 |
| LD15332 | FBgn0035900 | ZC3H3 | 0.38017891 | 0.10942324 | 0 |
| SD14112 | FBgn0033087 | Hsepi | -0.380056837 | 0.0610838 | 0 |
| LD16320 | FBgn0262560 | wcd | 0.379929139 | 0.05096655 | 0 |

| | | | | | |
|---|---|---|---|---|---|
| HL05860 | FBgn0030960 | Atg101 | 0.379592538 | 0.04279742 | 0 |
| LD10058 | FBgn0283508 | nw | 0.378961736 | 0.51424726 | 0 |
| LD40453 | FBgn0266570 | NO66 | -0.378918056 | 0.03751067 | 0 |
| GH05536 | FBgn0027586 | CG5867 | 0.378839159 | 0.1701014 | 0 |
| RE48393 | FBgn0029662 | CG12206 | -0.37853964 | 0.16546644 | 0 |
| RH31013 | FBgn0024956 | Mat1 | -0.37841585 | 0.04790739 | 0 |
| SD27452 | FBgn0023094 | cyc | -0.378145404 | 0.04584051 | 0 |
| GM01353 | FBgn0037760 | FBXO11 | -0.378042667 | 0.06027383 | 0 |
| GH01729 | FBgn0034644 | CG10082 | 0.377477057 | 0.05108182 | 0 |
| LP12382 | FBgn0024182 | waw | -0.376889517 | 0.04079919 | 0 |
| RE04903 | FBgn0032231 | CG5056 | -0.375937247 | 0.05025857 | 0 |
| GH28351 | FBgn0037432 | CG10298 | -0.375819643 | 0.09762485 | 0 |
| SD06504 | FBgn0010278 | Ssrp | -0.375204945 | 0.06643508 | 0 |
| RE02759 | FBgn0033349 | CG8243 | -0.374790798 | 0.04220711 | 0 |
| RE22626 | FBgn0261565 | Lmpt | 0.374744333 | 0.24386685 | 0 |
| AT07857 | FBgn0037942 | CG14721 | -0.374601626 | 0.03615988 | 0 |
| RE32485 | FBgn0037000 | ZnT77C | 0.374515424 | 0.05780892 | 0 |
| LD17463 | FBgn0261648 | salm | -0.374468866 | 0.32174828 | 0 |
| GH27809 | FBgn0050421 | Usp15-31 | -0.374388504 | 0.1241655 | 0 |
| LD44327 | FBgn0031157 | CG1503 | 0.373983045 | 0.1246311 | 0 |
| SD03330 | FBgn0035987 | CG3689 | -0.373633617 | 0.04951619 | 0 |
| GH20325 | FBgn0033792 | CG13325 | -0.373312934 | 0.036157 | 0 |
| LD13807 | FBgn0029882 | CG3226 | -0.373291262 | 0.03751067 | 0 |
| LD14179 | FBgn0035249 | CG17249 | 0.373160961 | 0.08612406 | 0 |
| GM14633 | FBgn0032445 | CG6153 | -0.373156155 | 0.0706372 | 0 |
| SD01014 | FBgn0035026 | Fcp1 | -0.372994841 | 0.03645446 | 0 |
| LD09509 | FBgn0261535 | l(2)34Fd | 0.372871171 | 0.03747115 | 0 |
| HL01025 | FBgn0034265 | Snx16 | -0.372744828 | 0.06614686 | 0 |
| LD35209 | FBgn0030871 | CG8142 | -0.372705282 | 0.05476173 | 0 |
| GH10821 | FBgn0037646 | CAHbeta | -0.372563789 | 0.05364672 | 0 |
| GH15453 | FBgn0261797 | Dhc64C | -0.372449086 | 0.13551782 | 0 |
| GM09415 | FBgn0034230 | CG4853 | -0.37226673 | 0.06614686 | 0 |
| RE37074 | FBgn0016687 | Nurf-38 | -0.372234596 | 0.0358459 | 0 |
| LD47350 | FBgn0003028 | ovo | -0.372183176 | 0.06712449 | 0 |
| RE55111 | FBgn0038196 | CG9922 | -0.372178164 | 0.05142607 | 0 |
| GH11680 | FBgn0031805 | CG9505 | 0.371931861 | 0.07367722 | 0 |
| HL08006 | FBgn0020513 | Paics | -0.371729081 | 0.03895492 | 0 |
| RE20574 | FBgn0033673 | CG8298 | -0.371017254 | 0.03557713 | 0 |
| RH61522 | FBgn0025634 | CG13367 | -0.370997375 | 0.03973242 | 0 |
| GH15647 | FBgn0266083 | ocm | 0.370689741 | 0.07810369 | 0 |
| LD20463 | FBgn0033785 | Sans | 0.370484685 | 0.18418073 | 0 |
| GH19932 | FBgn0004652 | fru | -0.370480068 | 0.04079919 | 0 |
| RE48269 | FBgn0032194 | CG4901 | -0.370431221 | 0.05028731 | 0 |
| RE44836 | FBgn0037248 | srl | 0.370266176 | 0.23695841 | 0 |
| RH54067 | FBgn0031801 | CG9498 | -0.370006807 | 0.18096666 | 0 |
| AT22870 | FBgn0011361 | ND-ACP | -0.369428491 | 0.04641137 | 0 |
| LD35517 | FBgn0033690 | EndoG | -0.368991861 | 0.04253873 | 0 |
| RE31047 | FBgn0000259 | CkIIbeta | -0.368521314 | 0.06712449 | 0 |

| | | | | | |
|---|---|---|---|---|---|
| LD40565 | FBgn0031961 | CG7102 | 0.368402424 | 0.08423184 | 0 |
| AT24411 | FBgn0250788 | beta-Spec | 0.367415558 | 0.10417336 | 0 |
| RE64786 | FBgn0032726 | CG10621 | 0.367362671 | 0.44392819 | 0 |
| LP07417 | FBgn0029529 | CG13365 | -0.366847628 | 0.46041263 | 0 |
| GM01004 | FBgn0024846 | p38b | -0.366788822 | 0.04988163 | 0 |
| GH12495 | FBgn0011236 | ken | -0.366654481 | 0.06885709 | 0 |
| RH70372 | FBgn0030648 | CG6340 | -0.366593658 | 0.04339964 | 0 |
| GH09771 | FBgn0036165 | chrb | 0.366417728 | 0.17917133 | 0 |
| LD36516 | FBgn0033413 | prel | -0.36636263 | 0.05096655 | 0 |
| RE46338 | FBgn0261534 | l(2)34Fc | 0.366200362 | 0.13436074 | 0 |
| RE65127 | FBgn0261808 | cu | -0.36595527 | 0.04220711 | 0 |
| GH20093 | FBgn0037356 | CG12170 | -0.365675635 | 0.06095715 | 0 |
| RE10485 | FBgn0010414 | SerT | -0.365362497 | 0.1332109 | 0 |
| LP07906 | FBgn0052447 | CG32447 | 0.36530925 | 0.3610233 | 0 |
| RE58687 | FBgn0038516 | P5cr-2 | 0.365159021 | 0.19648812 | 0 |
| AT13296 | FBgn0037616 | CG8136 | 0.365104581 | 0.04079919 | 0 |
| GH03377 | FBgn0027597 | CG17712 | -0.364238094 | 0.09008791 | 0 |
| GH24245 | FBgn0034735 | CG4610 | -0.364174314 | 0.06614686 | 0 |
| LD38109 | FBgn0043791 | phu | 0.363744452 | 0.07570279 | 0 |
| RE66244 | FBgn0004228 | mex1 | 0.363576973 | 0.37804643 | 0 |
| RE40480 | FBgn0040465 | Dlip3 | 0.363450128 | 0.06286082 | 0 |
| LD24726 | FBgn0037186 | CG11241 | 0.362963762 | 0.09784391 | 0 |
| RE70560 | FBgn0003676 | CCT1 | -0.362217741 | 0.04422997 | 0 |
| GM05003 | FBgn0260642 | Antp | 0.361908362 | 0.16163549 | 0 |
| RH55360 | FBgn0266421 | RPA3 | -0.361641522 | 0.06296618 | 0 |
| RE55538 | FBgn0031434 | insv | 0.361479543 | 0.09362642 | 0 |
| HL08109 | FBgn0040064 | yip2 | -0.361243925 | 0.04354859 | 0 |
| GH06744 | FBgn0036760 | CG5567 | -0.361092052 | 0.05780892 | 0 |
| RH57813 | FBgn0015296 | Shc | -0.360974919 | 0.07088605 | 0 |
| LD12084 | FBgn0001120 | gnu | -0.360791834 | 0.06178355 | 0 |
| SD02625 | FBgn0034068 | casp | -0.360555321 | 0.04079919 | 0 |
| AT11552 | FBgn0020399 | Mst89B | 0.359852522 | 0.04597426 | 0 |
| RE58537 | FBgn0260632 | dl | -0.359620482 | 0.05294781 | 0 |
| LD45056 | FBgn0031399 | mio | 0.359479414 | 0.07000688 | 0 |
| RE57663 | FBgn0036501 | CG7272 | -0.359210857 | 0.10774879 | 0 |
| AT14183 | FBgn0267891 | CR46179 | -0.358898946 | 0.04253873 | 0 |
| AT14183 | FBgn0267877 | CR46165 | -0.358898946 | 0.04253873 | 0 |
| SD10541 | FBgn0025802 | Sbf | -0.358796892 | 0.05876215 | 0 |
| AT30052 | FBgn0265606 | Prosalpha4T1 | -0.358701268 | 0.03751067 | 0 |
| GH10714 | FBgn0029866 | CG3842 | -0.358043453 | 0.06173793 | 0 |
| RE47909 | FBgn0044510 | mRpS5 | 0.357869675 | 0.43501544 | 0 |
| RE47909 | FBgn0085810 | CR41597 | 0.357869675 | 0.43501544 | 0 |
| RE47909 | FBgn0261639 | CR42722 | 0.357869675 | 0.43501544 | 0 |
| RE47909 | FBgn0261640 | CR42723 | 0.357869675 | 0.43501544 | 0 |
| GM01344 | FBgn0037613 | Cks85A | 0.357730264 | 0.50766723 | 0 |
| LD32860 | FBgn0033616 | CG7745 | -0.35762213 | 0.09008791 | 0 |
| LD43541 | FBgn0031091 | Phf7 | -0.357359025 | 0.04930642 | 0 |
| RH51491 | FBgn0259715 | CG42369 | -0.357255102 | 0.34494921 | 0 |

| | | | | | |
|---|---|---|---|---|---|
| GM28660 | FBgn0033086 | CG9410 | -0.35662795 | 0.06410905 | 0 |
| RH43429 | FBgn0032281 | CG17107 | -0.355878165 | 0.32650577 | 0 |
| GH22838 | FBgn0261625 | GLS | -0.355834541 | 0.09008791 | 0 |
| AT14673 | FBgn0250755 | CG42233 | -0.355611788 | 0.05737109 | 0 |
| LD21041 | FBgn0031990 | PAPLA1 | 0.355607414 | 0.08415149 | 0 |
| LD43341 | FBgn0260959 | MCPH1 | -0.355596509 | 0.04041964 | 0 |
| AT10581 | FBgn0029153 | Menl-2 | -0.355571115 | 0.28784599 | 0 |
| AT10581 | FBgn0029154 | Menl-1 | -0.355571115 | 0.28784599 | 0 |
| RH10647 | FBgn0053516 | dpr3 | -0.35532379 | 0.17793042 | 0 |
| SD08659 | FBgn0024277 | trio | 0.355315236 | 0.04079919 | 0 |
| GH22790 | FBgn0033654 | Sobp | -0.354819149 | 0.3931041 | 0 |
| AT04454 | FBgn0031115 | CG11710 | -0.354403076 | 0.04354859 | 0 |
| LD47517 | FBgn0037360 | CG2182 | -0.354338252 | 0.05883095 | 0 |
| RE45235 | FBgn0029502 | COQ7 | -0.353973363 | 0.05630422 | 0 |
| RE12242 | FBgn0032264 | Lip4 | 0.353967506 | 0.31861085 | 0 |
| GH04201 | FBgn0010246 | Myo61F | -0.353870542 | 0.07367722 | 0 |
| SD15313 | FBgn0038353 | CG5399 | 0.353792598 | 0.25348968 | 0 |
| SD15313 | FBgn0266124 | ghi | 0.353792598 | 0.25348968 | 0 |
| SD01679 | FBgn0041100 | park | -0.353641273 | 0.13585443 | 0 |
| LD18889 | FBgn0047205 | lncRNA:CR32385 | 0.35329661 | 0.08383538 | 0 |
| GH20156 | FBgn0034975 | enok | 0.35323368 | 0.06712449 | 0 |
| LD22351 | FBgn0004107 | Cdk2 | -0.353023513 | 0.06654512 | 0 |
| GH11843 | FBgn0035246 | CG13928 | -0.352911595 | 0.34255324 | 0 |
| SD03119 | FBgn0016977 | spen | 0.3529067 | 0.09204205 | 0 |
| GH04269 | FBgn0041706 | CG3253 | 0.352808461 | 0.14477381 | 0 |
| SD06874 | FBgn0267385 | PyK | -0.352709731 | 0.04041964 | 0 |
| LP07248 | FBgn0024841 | Pcd | -0.352585186 | 0.06885709 | 0 |
| AT14053 | FBgn0015283 | Rpn10 | -0.352270857 | 0.04254487 | 0 |
| RE53634 | FBgn0052227 | gogo | 0.35221565 | 0.05571776 | 0 |
| RE67487 | FBgn0034627 | eEFSec | -0.352088877 | 0.06640401 | 0 |
| LD10878 | FBgn0036194 | CG11652 | 0.351837414 | 0.1018806 | 0 |
| RE28239 | FBgn0033668 | exp | -0.35138263 | 0.2219358 | 0 |
| GH04046 | FBgn0263354 | CG42784 | -0.351286282 | 0.33882199 | 0 |
| GH04046 | FBgn0284224 | CG46308 | -0.351286282 | 0.33882199 | 0 |
| GH09840 | FBgn0039561 | mfrn | -0.350748855 | 0.07152571 | 0 |
| GH11519 | FBgn0260745 | mfas | 0.350728575 | 0.13551782 | 0 |
| GH28308 | FBgn0035849 | ERR | -0.35072685 | 0.19895429 | 0 |
| RE02884 | FBgn0024980 | Syx4 | -0.35041448 | 0.04339964 | 0 |
| RH23394 | FBgn0039916 | Ekar | -0.349888536 | 0.04626457 | 0 |
| RH23394 | FBgn0264820 | CR44028 | -0.349888536 | 0.04626457 | 0 |
| RE32163 | FBgn0031534 | Snx1 | -0.349547474 | 0.04339964 | 0 |
| GH28335 | FBgn0038826 | Syp | -0.349463734 | 0.09008791 | 0 |
| RE14039 | FBgn0053970 | CG33970 | -0.349367518 | 0.0817644 | 0 |
| AT07434 | FBgn0033382 | Hydr1 | -0.349239921 | 0.04803177 | 0 |
| GH11143 | FBgn0035169 | CG13890 | -0.349233844 | 0.09325764 | 0 |
| SD04227 | FBgn0086690 | Plp | 0.349137675 | 0.10151555 | 0 |
| RE03785 | FBgn0040260 | Ugt36Bc | 0.349071746 | 0.31005535 | 0 |
| GH10149 | FBgn0031106 | Syx16 | -0.349015721 | 0.06410905 | 0 |

| | | | | | |
|---|---|---|---|---|---|
| RE01365 | FBgn0028997 | nmdyn-D7 | -0.348845181 | 0.04988163 | 0 |
| LP08674 | FBgn0033183 | CG1620 | -0.348514875 | 0.04253873 | 0 |
| AT15314 | FBgn0037098 | Wnk | -0.348321511 | 0.07367722 | 0 |
| LD39377 | FBgn0029094 | asf1 | -0.348240398 | 0.07380561 | 0 |
| RH40632 | FBgn0052647 | CG32647 | -0.34788697 | 0.18644325 | 0 |
| LP03320 | FBgn0016984 | sktl | -0.347740872 | 0.04434426 | 0 |
| AT25977 | FBgn0033883 | CG16935 | -0.34767258 | 0.0494494 | 0 |
| LD46744 | FBgn0027360 | Tim10 | -0.347338503 | 0.10804827 | 0 |
| LD46744 | FBgn0260223 | CG42497 | -0.347338503 | 0.10804827 | 0 |
| GH22837 | FBgn0039704 | neo | 0.347271121 | 0.61780569 | 0 |
| RE47056 | FBgn0085409 | smal | 0.347101883 | 0.11055415 | 0 |
| LD13050 | FBgn0037092 | M6 | 0.347083676 | 0.17434219 | 0 |
| GH10494 | FBgn0035390 | scramb2 | -0.34684934 | 0.05811678 | 0 |
| RE38146 | FBgn0033903 | CG8323 | -0.34670645 | 0.07367722 | 0 |
| RE63174 | FBgn0035158 | CG13895 | 0.346697902 | 0.09890059 | 0 |
| RE11339 | FBgn0002283 | l(3)73Ah | -0.346507103 | 0.06347849 | 0 |
| GH04560 | FBgn0042138 | CG18815 | -0.346390525 | 0.04253873 | 0 |
| RE18536 | FBgn0024177 | zpg | -0.345498943 | 0.05794022 | 0 |
| LD26982 | FBgn0031314 | IntS14 | 0.345367706 | 0.07570279 | 0 |
| LD35990 | FBgn0036480 | Cep135 | 0.345191346 | 0.0527752 | 0 |
| RE05342 | FBgn0035626 | lin-28 | 0.345134163 | 0.37538642 | 0 |
| RE06653 | FBgn0035228 | CG12091 | -0.344126388 | 0.06640401 | 0 |
| HL03430 | FBgn0001247 | Ide | -0.343850029 | 0.05329829 | 0 |
| RE32705 | FBgn0035944 | CG5021 | -0.343303285 | 0.10087976 | 0 |
| LD43096 | FBgn0052428 | CG32428 | -0.342547328 | 0.07853441 | 0 |
| GH22222 | FBgn0035833 | CG7565 | -0.342277631 | 0.17107296 | 0 |
| RE05911 | FBgn0259233 | CG42331 | -0.342241554 | 0.04202135 | 0 |
| AT22836 | FBgn0051055 | CG31055 | -0.341823755 | 0.19881267 | 0 |
| LP03106 | FBgn0039795 | Spn100A | 0.341789993 | 0.57265237 | 0 |
| GH11288 | FBgn0011559 | Acp36DE | 0.341536353 | 0.13232211 | 0 |
| GH06573 | FBgn0262636 | dati | -0.34130265 | 0.11235733 | 0 |
| LD16826 | FBgn0033752 | CG8569 | 0.341155606 | 0.1609697 | 0 |
| RE34782 | FBgn0085424 | nub | 0.341123622 | 0.14951145 | 0 |
| RE15841 | FBgn0033096 | Zip42C.1 | -0.340969645 | 0.13597632 | 0 |
| RE20777 | FBgn0033549 | mms4 | -0.340938659 | 0.04369271 | 0 |
| SD02428 | FBgn0026402 | NiPp1 | -0.340932513 | 0.06221166 | 0 |
| LD11130 | FBgn0262740 | Evi5 | -0.34079283 | 0.11637996 | 0 |
| AT01533 | FBgn0053129 | CG33129 | -0.340662203 | 0.07081446 | 0 |
| GH08220 | FBgn0259166 | CG42271 | -0.340434351 | 0.17309475 | 0 |
| LD19219 | FBgn0259923 | Sep4 | -0.340308803 | 0.04306004 | 0 |
| LD46058 | FBgn0041342 | Pcyt1 | 0.340122683 | 0.17863263 | 0 |
| GH16590 | FBgn0037561 | CG9630 | -0.339845488 | 0.11842639 | 0 |
| RE39441 | FBgn0028573 | prc | -0.33973469 | 0.17147856 | 0 |
| GH13437 | FBgn0029155 | Men-b | -0.339535379 | 0.04253873 | 0 |
| GH14157 | FBgn0029892 | CG3184 | -0.339168918 | 0.09672974 | 0 |
| LD05244 | FBgn0035518 | CG15011 | -0.338945392 | 0.10417336 | 0 |
| LD21247 | FBgn0004419 | me31B | -0.338732033 | 0.0878206 | 0 |
| GH06759 | FBgn0038039 | CG5196 | -0.338685135 | 0.11638186 | 0 |

| | | | | | |
|---|---|---|---|---|---|
| SD04291 | FBgn0000482 | dor | 0.338602289 | 0.07058212 | 0 |
| LP01634 | FBgn0032087 | CG9568 | -0.338190929 | 0.16091105 | 0 |
| RE12330 | FBgn0040071 | tara | -0.338182944 | 0.21538541 | 0 |
| GH22722 | FBgn0035767 | Cln7 | 0.338116942 | 0.12947861 | 0 |
| LD18295 | FBgn0052037 | CG32037 | 0.338052006 | 0.1848493 | 0 |
| LD24529 | FBgn0010417 | Taf6 | 0.337636847 | 0.11597957 | 0 |
| RE55932 | FBgn0023395 | Chd3 | -0.337074196 | 0.04666005 | 0 |
| RE55932 | FBgn0262519 | Mi-2 | -0.337074196 | 0.04666005 | 0 |
| SD01878 | FBgn0051352 | Unc-115a | 0.337073064 | 0.12320472 | 0 |
| LD47550 | FBgn0033454 | CG1671 | -0.33700479 | 0.0445191 | 0 |
| SD17750 | FBgn0261262 | CG42613 | -0.33682689 | 0.42975739 | 0 |
| RE05264 | FBgn0032225 | CG5022 | 0.336667074 | 0.12628732 | 0 |
| RE33063 | FBgn0020642 | Lcp65Ac | -0.33637084 | 0.2323332 | 0 |
| SD08060 | FBgn0039654 | Brd8 | 0.335670484 | 0.07158299 | 0 |
| LD19823 | FBgn0031752 | CG9044 | -0.335607868 | 0.11505352 | 0 |
| SD02173 | FBgn0053196 | dpy | 0.3355425 | 0.40221267 | 0 |
| GH03649 | FBgn0050392 | CG30392 | -0.335446771 | 0.0878206 | 0 |
| GH21352 | FBgn0030823 | TTLL1A | -0.335356917 | 0.40746238 | 0 |
| RH42718 | FBgn0019982 | Gs1l | -0.33511683 | 0.10451896 | 0 |
| GH15272 | FBgn0029947 | CG1999 | -0.334965111 | 0.31044547 | 0 |
| GH05422 | FBgn0264672 | Eogt | 0.334865181 | 0.15019038 | 0 |
| GH26048 | FBgn0261477 | slim | -0.334863934 | 0.05096655 | 0 |
| GH13422 | FBgn0002593 | RpLP1 | -0.334640369 | 0.09671981 | 0 |
| RE08857 | FBgn0259482 | Mob3 | -0.334464666 | 0.22622872 | 0 |
| RE01432 | FBgn0029771 | CG12730 | 0.334462531 | 0.26307647 | 0 |
| LP05054 | FBgn0031590 | CG3702 | -0.334298821 | 0.04989377 | 0 |
| LD30287 | FBgn0039602 | CG1647 | 0.334269058 | 0.0878206 | 0 |
| AT21437 | FBgn0010803 | TrpRS | -0.334041154 | 0.05075947 | 0 |
| RH62365 | FBgn0031700 | CG14022 | 0.333313301 | 0.17008461 | 0 |
| RE26342 | FBgn0031913 | CG5958 | 0.333308805 | 0.39662446 | 0 |
| RE63452 | FBgn0029693 | CG6379 | -0.333291621 | 0.06420995 | 0 |
| GH03617 | FBgn0039348 | Npl4 | -0.332913884 | 0.07367722 | 0 |
| SD05282 | FBgn0035023 | ITP | -0.33286083 | 0.39374565 | 0 |
| RE70722 | FBgn0035402 | Usp5 | -0.332770062 | 0.0470559 | 0 |
| RE55430 | FBgn0266674 | Sec15 | -0.332372411 | 0.04434426 | 0 |
| GM04690 | FBgn0035401 | Alg2 | 0.332041209 | 0.13417211 | 0 |
| RE09295 | FBgn0027093 | ArgRS | -0.331954506 | 0.07058212 | 0 |
| SD03001 | FBgn0002781 | mod(mdg4) | 0.331831785 | 0.05025857 | 0 |
| SD03001 | FBgn0266172 | pre-mod(mdg4)-E | 0.331831785 | 0.05025857 | 0 |
| SD03001 | FBgn0266171 | pre-mod(mdg4)-AB | 0.331831785 | 0.05025857 | 0 |
| SD03001 | FBgn0266170 | pre-mod(mdg4)-C | 0.331831785 | 0.05025857 | 0 |
| RE59252 | FBgn0016047 | nompA | -0.331811064 | 0.06708516 | 0 |
| GH17724 | FBgn0038617 | CG12333 | -0.331679048 | 0.24204398 | 0 |
| SD28232 | FBgn0035424 | CG11505 | -0.331256804 | 0.11351359 | 0 |
| LP03571 | FBgn0014417 | CG13397 | -0.331216434 | 0.06713041 | 0 |
| LD25575 | FBgn0037718 | P58IPK | -0.330543953 | 0.05256243 | 0 |
| RE10407 | FBgn0037063 | CG9391 | -0.329789861 | 0.06713041 | 0 |
| RE11240 | FBgn0030590 | CG9518 | -0.328997357 | 0.21693903 | 0 |

| | | | | | |
|---|---|---|---|---|---|
| GH26265 | FBgn0250849 | CG32388 | -0.328793004 | 0.29835009 | 0 |
| GH26265 | FBgn0263387 | CG43439 | -0.328793004 | 0.29835009 | 0 |
| LD41494 | FBgn0030734 | CG9911 | -0.328745262 | 0.07639145 | 0 |
| AT15807 | FBgn0038725 | CG6184 | -0.328689426 | 0.11035759 | 0 |
| AT09365 | FBgn0263131 | CG43373 | -0.328583526 | 0.14765386 | 0 |
| LD14261 | FBgn0261872 | scaf6 | 0.328295815 | 0.04790739 | 0 |
| RH70249 | FBgn0001234 | lncRNA:Hsromega | 0.327924759 | 0.04535599 | 0 |
| GH09541 | FBgn0052311 | zormin | -0.327522053 | 0.24417029 | 0 |
| GH24627 | FBgn0052380 | SMSr | -0.327316092 | 0.06761917 | 0 |
| GM09668 | FBgn0000286 | Cf2 | -0.326988174 | 0.08054283 | 0 |
| GM13032 | FBgn0035896 | CG6983 | -0.326959536 | 0.13843226 | 0 |
| LD10094 | FBgn0026533 | Dek | 0.326835852 | 0.06590178 | 0 |
| SD12744 | FBgn0259823 | CG42404 | 0.326820013 | 0.13976606 | 0 |
| SD06427 | FBgn0038146 | CG9799 | -0.326726508 | 0.07806278 | 0 |
| GH10726 | FBgn0001311 | kkv | 0.326467335 | 0.19904473 | 0 |
| AT28183 | FBgn0051948 | CG31948 | 0.326422674 | 0.18777495 | 0 |
| LD41718 | FBgn0030000 | CG2260 | 0.326092139 | 0.07570279 | 0 |
| SD10374 | FBgn0010379 | Akt1 | -0.325640885 | 0.07570279 | 0 |
| LD37702 | FBgn0036147 | Plod | 0.325621605 | 0.0892183 | 0 |
| SD04694 | FBgn0053158 | CG33158 | -0.325132308 | 0.06696875 | 0 |
| GH04054 | FBgn0037818 | CG6465 | 0.325130795 | 0.16385793 | 0 |
| GH22314 | FBgn0011754 | PhKgamma | -0.325047939 | 0.04850212 | 0 |
| AT26455 | FBgn0004654 | Pgd | -0.324930715 | 0.13506052 | 0 |
| GM21739 | FBgn0034726 | Mes4 | -0.324724762 | 0.07367722 | 0 |
| AT10348 | FBgn0013995 | Calx | -0.32463142 | 0.12977103 | 0 |
| LD25651 | FBgn0010391 | Gtp-bp | -0.32444282 | 0.05957516 | 0 |
| LD33040 | FBgn0030500 | Ndc80 | -0.324090102 | 0.09445121 | 0 |
| GH22074 | FBgn0003302 | rux | -0.324003414 | 0.07810369 | 0 |
| GM18993 | FBgn0004359 | T48 | -0.323855944 | 0.12810427 | 0 |
| GH28062 | FBgn0028996 | onecut | 0.323789692 | 0.44707004 | 0 |
| RE04220 | FBgn0017545 | RpS3A | -0.323729739 | 0.08339651 | 0 |
| RE13149 | FBgn0031309 | Tfb4 | 0.323696327 | 0.15705273 | 0 |
| RE55422 | FBgn0021768 | nudC | -0.323357428 | 0.05643298 | 0 |
| AT18545 | FBgn0030196 | Psf3 | -0.323256173 | 0.05025857 | 0 |
| GH08865 | FBgn0283467 | Pol32 | -0.32298127 | 0.0487029 | 0 |
| GM03068 | FBgn0033741 | CG8545 | 0.322480593 | 0.14275424 | 0 |
| LD24540 | FBgn0041588 | eIF2D | 0.322338561 | 0.06474917 | 0 |
| LD30288 | FBgn0036196 | CG11658 | 0.322036404 | 0.07806278 | 0 |
| RE35686 | FBgn0034893 | mRpL43 | -0.322021742 | 0.05329829 | 0 |
| LD08233 | FBgn0038981 | CG5346 | 0.32112451 | 0.13844439 | 0 |
| RE09305 | FBgn0051036 | CG31036 | -0.321122685 | 0.1115611 | 0 |
| AT14039 | FBgn0036723 | CG12229 | -0.321064133 | 0.09451622 | 0 |
| SD01796 | FBgn0003041 | pbl | -0.320767954 | 0.14961497 | 0 |
| RH32180 | FBgn0038846 | CG5697 | -0.320686406 | 0.09249667 | 0 |
| RE02070 | FBgn0015924 | crq | -0.320546097 | 0.0892183 | 0 |
| GM08734 | FBgn0014022 | Rlb1 | 0.320170684 | 0.06531909 | 0 |
| SD10846 | FBgn0261049 | Vps45 | -0.319888726 | 0.05630422 | 0 |
| GH20817 | FBgn0033446 | CG1648 | 0.319689683 | 0.1325715 | 0 |

| | | | | | |
|---|---|---|---|---|---|
| LD01951 | FBgn0051998 | CG31998 | -0.319616956 | 0.05002903 | 0 |
| RE26721 | FBgn0038832 | CG15695 | -0.319458955 | 0.12044726 | 0 |
| RE40104 | FBgn0001138 | grn | 0.319406015 | 0.28911598 | 0 |
| LD05680 | FBgn0033235 | CG8728 | -0.319342957 | 0.0710745 | 0 |
| LD37787 | FBgn0003162 | Pu | 0.319186826 | 0.09698692 | 0 |
| SD01505 | FBgn0040087 | p115 | 0.318440526 | 0.06712449 | 0 |
| LP04371 | FBgn0036273 | INPP5E | -0.318439496 | 0.06530718 | 0 |
| GH20785 | FBgn0013983 | imd | -0.318305078 | 0.10108162 | 0 |
| GH07892 | FBgn0030841 | CG8568 | -0.318244777 | 0.29543934 | 0 |
| GM09708 | FBgn0266731 | lncRNA:CR45204 | 0.318222911 | 0.14614309 | 0 |
| LD23573 | FBgn0053156 | CG33156 | -0.317928551 | 0.05816357 | 0 |
| GH25655 | FBgn0033431 | CG1827 | -0.317831567 | 0.1829606 | 0 |
| GH22139 | FBgn0033391 | CG8026 | -0.317816641 | 0.06544177 | 0 |
| GH22139 | FBgn0266486 | CG45085 | -0.317816641 | 0.06544177 | 0 |
| LD02329 | FBgn0026441 | ear | 0.317544035 | 0.07367722 | 0 |
| SD01380 | FBgn0013756 | Mtor | 0.317318485 | 0.05142607 | 0 |
| GH08447 | FBgn0038332 | CG6136 | -0.317207987 | 0.08729505 | 0 |
| LD21452 | FBgn0038487 | TwdlW | -0.317165461 | 0.21426982 | 0 |
| HL08141 | FBgn0031094 | CG9578 | -0.316933801 | 0.06259049 | 0 |
| RH61984 | FBgn0039494 | grass | -0.316831737 | 0.06455982 | 0 |
| LD40692 | FBgn0036754 | CG5589 | 0.316765964 | 0.05662207 | 0 |
| RE59650 | FBgn0031639 | mRpS2 | -0.31671495 | 0.16314895 | 0 |
| RE25242 | FBgn0034231 | APC10 | -0.316543944 | 0.06234185 | 0 |
| RE21674 | FBgn0040297 | Nhe2 | 0.31625476 | 0.49331415 | 0 |
| RE21674 | FBgn0284223 | CG46307 | 0.31625476 | 0.49331415 | 0 |
| GH08974 | FBgn0037955 | CG6950 | -0.31596771 | 0.07246806 | 0 |
| GH13403 | FBgn0033924 | CG8613 | -0.315799471 | 0.10942324 | 0 |
| GH20986 | FBgn0038870 | Oga | -0.315616767 | 0.13551782 | 0 |
| GH04507 | FBgn0030316 | CG11695 | -0.31545678 | 0.05907437 | 0 |
| LD43867 | FBgn0032791 | CG18094 | -0.315399743 | 0.13642205 | 0 |
| RE02195 | FBgn0039084 | CG10175 | -0.315382368 | 0.28553785 | 0 |
| RH02445 | FBgn0015765 | p38a | -0.315199275 | 0.10009191 | 0 |
| RH25557 | FBgn0039008 | CG6972 | 0.315185503 | 0.49390745 | 0 |
| LP05991 | FBgn0040321 | GNBP3 | -0.315099417 | 0.06808125 | 0 |
| LP05991 | FBgn0002622 | RpS3 | -0.315099417 | 0.06808125 | 0 |
| RE03249 | FBgn0030975 | SdhBL | -0.314831402 | 0.20048325 | 0 |
| AT25108 | FBgn0267624 | lncRNA:CR45962 | 0.314419852 | 0.10505574 | 0 |
| RE25179 | FBgn0010194 | Wnt5 | 0.314182681 | 0.1037592 | 0 |
| LD28275 | FBgn0036500 | CG7275 | 0.314164441 | 0.09361444 | 0 |
| SD03428 | FBgn0031392 | AIF | -0.313991891 | 0.05411029 | 0 |
| GH17694 | FBgn0263846 | CG43707 | -0.313908091 | 0.38917984 | 0 |
| GH08385 | FBgn0027565 | CG5498 | -0.313660826 | 0.06234185 | 0 |
| SD24313 | FBgn0030407 | Fpgs | -0.313534067 | 0.09672974 | 0 |
| LD02906 | FBgn0015799 | Rbf | -0.313323613 | 0.08423184 | 0 |
| RE23430 | FBgn0259245 | DIP-beta | -0.313152703 | 0.22929842 | 0 |
| RE16159 | FBgn0052791 | DIP-alpha | -0.312954384 | 0.4047532 | 0 |
| LD42739 | FBgn0005674 | GluProRS | -0.312831593 | 0.06981122 | 0 |
| AT30755 | FBgn0263048 | Gpdh3 | -0.31280018 | 0.1317984 | 0 |

| | | | | | |
|---|---|---|---|---|---|
| CK01017 | FBgn0028539 | CG31731 | 0.312371244 | 0.14693767 | 0 |
| SD02824 | FBgn0024245 | dnt | 0.312163516 | 0.07464414 | 0 |
| SD04959 | FBgn0020887 | Su(z)12 | -0.312083631 | 0.07097747 | 0 |
| RH01162 | FBgn0033321 | CG8738 | -0.311857719 | 0.32785535 | 0 |
| LD46263 | FBgn0033569 | CG12942 | -0.311690145 | 0.06095715 | 0 |
| RE70810 | FBgn0052105 | Lmx1a | -0.311319216 | 0.23906935 | 0 |
| LD13846 | FBgn0028325 | Pdha | -0.310916274 | 0.05794022 | 0 |
| GH25855 | FBgn0035231 | Pcyt2 | 0.310901517 | 0.08419286 | 0 |
| RE52190 | FBgn0083940 | RhoU | -0.310890302 | 0.12207297 | 0 |
| GH26152 | FBgn0036398 | upSET | 0.310792778 | 0.06713041 | 0 |
| RH16520 | FBgn0263258 | chas | -0.310781564 | 0.09167912 | 0 |
| GH18717 | FBgn0011659 | Mlh1 | -0.31071335 | 0.05871745 | 0 |
| RE23178 | FBgn0262647 | Nup160 | 0.310368226 | 0.07191509 | 0 |
| GH07626 | FBgn0283427 | FASN1 | 0.310279727 | 0.13642205 | 0 |
| GM13274 | FBgn0260456 | CG4806 | 0.310272892 | 0.08974365 | 0 |
| RE31408 | FBgn0261065 | Cpsf73 | -0.309874884 | 0.07038909 | 0 |
| LD30829 | FBgn0034098 | krimp | -0.309723853 | 0.11359747 | 0 |
| RE37385 | FBgn0005638 | slbo | -0.309554407 | 0.18472354 | 0 |
| GH08760 | FBgn0034915 | eIF6 | -0.309506402 | 0.06066763 | 0 |
| SD05719 | FBgn0032775 | CG17544 | -0.30923439 | 0.13744341 | 0 |
| LD32979 | FBgn0026418 | Hsc70Cb | -0.309058733 | 0.05573152 | 0 |
| GM04951 | FBgn0032906 | RPA2 | -0.308908819 | 0.10748572 | 0 |
| GM07702 | FBgn0261709 | lncRNA:CR42746 | 0.308819788 | 0.11525613 | 0 |
| RE68619 | FBgn0016075 | vkg | 0.308512428 | 0.14921756 | 0 |
| SD15892 | FBgn0261545 | CG42663 | -0.308382457 | 0.22542256 | 0 |
| SD05789 | FBgn0035436 | CG12016 | -0.308237597 | 0.05956874 | 0 |
| SD05789 | FBgn0259933 | CG42456 | -0.308237597 | 0.05956874 | 0 |
| RH33561 | FBgn0000279 | CecC | -0.307391887 | 0.10942324 | 0 |
| RE57978 | FBgn0039189 | CG18528 | -0.307195173 | 0.08612406 | 0 |
| RE05579 | FBgn0032705 | Grip71 | -0.306904333 | 0.12034993 | 0 |
| LD29131 | FBgn0002924 | ncd | 0.306831023 | 0.10942324 | 0 |
| LP09908 | FBgn0030528 | CG11095 | -0.306802889 | 0.07152571 | 0 |
| HL01609 | FBgn0031945 | CG7191 | 0.306593938 | 0.1545559 | 0 |
| RE23259 | FBgn0039829 | CG15561 | 0.306441353 | 0.05589463 | 0 |
| SD02803 | FBgn0035688 | fmt | -0.306310585 | 0.08647308 | 0 |
| RE27723 | FBgn0052365 | CG32365 | -0.306215758 | 0.22447805 | 0 |
| RE38703 | FBgn0024293 | Spn43Ab | -0.306192622 | 0.06712449 | 0 |
| LP01183 | FBgn0030837 | CG8661 | 0.30605445 | 0.12213767 | 0 |
| GH02475 | FBgn0032068 | LManV | 0.30586991 | 0.26011542 | 0 |
| GM30016 | FBgn0039405 | PIG-P | -0.305355589 | 0.07731902 | 0 |
| GM30016 | FBgn0260224 | CG42498 | -0.305355589 | 0.07731902 | 0 |
| LD15680 | FBgn0259234 | Camta | 0.30495861 | 0.07374924 | 0 |
| GH06563 | FBgn0038947 | Sar1 | -0.30456042 | 0.07731902 | 0 |
| RE50273 | FBgn0026084 | cib | -0.304493764 | 0.09008791 | 0 |
| SD08609 | FBgn0032296 | CG6729 | -0.304488841 | 0.131545 | 0 |
| RE15579 | FBgn0284435 | tyn | 0.304415163 | 0.35079094 | 0 |
| GH20068 | FBgn0053147 | Hs3st-A | -0.304347221 | 0.25068834 | 0 |
| RE59473 | FBgn0031150 | bves | -0.304326325 | 0.11689839 | 0 |

| | | | | | |
|---|---|---|---|---|---|
| LD35248 | FBgn0028382 | cyp33 | -0.304269398 | 0.08025093 | 0 |
| RE30084 | FBgn0050009 | lncRNA:CR30009 | 0.304100605 | 0.5946724 | 0 |
| RE65108 | FBgn0031855 | CG11221 | -0.303969438 | 0.09008791 | 0 |
| LP04696 | FBgn0250907 | Cht10 | 0.303930888 | 0.0973297 | 0 |
| RE65864 | FBgn0052438 | SMC5 | -0.303911584 | 0.06713041 | 0 |
| GM03482 | FBgn0034029 | eIF2Bgamma | -0.303873299 | 0.09008791 | 0 |
| LP03565 | FBgn0028537 | CG31775 | 0.303846886 | 0.37551613 | 0 |
| LP03565 | FBgn0260954 | CG42586 | 0.303846886 | 0.37551613 | 0 |
| LD28544 | FBgn0034988 | cN-IIIB | 0.303645435 | 0.08253963 | 0 |
| LP03982 | FBgn0031657 | CG3756 | -0.303579801 | 0.09362642 | 0 |
| RH09340 | FBgn0034935 | Orcokinin | 0.303568922 | 0.07570279 | 0 |
| RE18871 | FBgn0038331 | Ccm3 | -0.303535884 | 0.12487471 | 0 |
| GH12174 | FBgn0042085 | Bap170 | 0.303470839 | 0.12207297 | 0 |
| GH13383 | FBgn0086707 | ncm | 0.303064074 | 0.07021123 | 0 |
| LD16361 | FBgn0020309 | crol | -0.302969458 | 0.09915304 | 0 |
| LD05267 | FBgn0025700 | CG5885 | -0.302938202 | 0.09963256 | 0 |
| RE35395 | FBgn0031604 | Elp3 | -0.302670805 | 0.06843333 | 0 |
| LD09852 | FBgn0027885 | Aac11 | 0.30221422 | 0.0603552 | 0 |
| GM10241 | FBgn0025832 | Fen1 | -0.302039804 | 0.06025942 | 0 |
| LD42723 | FBgn0000326 | clt | 0.302024592 | 0.0603552 | 0 |
| GH26459 | FBgn0017429 | CG5989 | -0.301767472 | 0.20099097 | 0 |
| LD24919 | FBgn0038057 | CG12267 | 0.301181958 | 0.05780892 | 0 |
| RE19601 | FBgn0051612 | CG31612 | 0.300979157 | 0.16751341 | 0 |
| GH08521 | FBgn0038706 | CG3517 | -0.300964139 | 0.09890059 | 0 |
| GH08521 | FBgn0033609 | Fbl6 | -0.300964139 | 0.09890059 | 0 |
| RH40310 | FBgn0003941 | RpL40 | -0.300866082 | 0.10434168 | 0 |
| GH18921 | FBgn0038868 | CG5862 | 0.300550778 | 0.12693221 | 0 |
| RH55123 | FBgn0038504 | Sur-8 | -0.3002918 | 0.36865559 | 0 |
| RE03238 | FBgn0050195 | CG30195 | -0.300262391 | 0.08647308 | 0 |
| GH17893 | FBgn0003149 | Prm | -0.300214799 | 0.29250904 | 0 |
| GH20954 | FBgn0031335 | CG5565 | -0.300193169 | 0.12320472 | 0 |
| GH20954 | FBgn0051924 | CG31924 | -0.300193169 | 0.12320472 | 0 |
| LD17982 | FBgn0032691 | Atac2 | -0.300180248 | 0.12685445 | 0 |
| LD40802 | FBgn0024366 | CG11409 | -0.300101415 | 0.05593259 | 0 |
| RH38755 | FBgn0039241 | CG11089 | -0.299790134 | 0.07759767 | 0 |
| LP11612 | FBgn0036436 | CG4914 | -0.299424274 | 0.23635904 | 0 |
| HL03538 | FBgn0032172 | CG5850 | -0.299423605 | 0.05630422 | 0 |
| HL03538 | FBgn0266166 | asRNA:CR44872 | -0.299423605 | 0.05630422 | 0 |
| GH01087 | FBgn0038274 | Nup93-2 | 0.299326145 | 0.11842639 | 0 |
| RE54151 | FBgn0034054 | CG8366 | -0.299285619 | 0.13618504 | 0 |
| RH34904 | FBgn0034364 | CG5493 | -0.29908391 | 0.19412854 | 0 |
| LD25641 | FBgn0085451 | htk | 0.298937171 | 0.11462142 | 0 |
| GH18858 | FBgn0040398 | CG14629 | 0.298743558 | 0.59982904 | 0 |
| GH22096 | FBgn0034191 | CG6984 | -0.298719977 | 0.11235733 | 0 |
| LD46817 | FBgn0030457 | CG12096 | -0.298714873 | 0.07306891 | 0 |
| LD44769 | FBgn0038279 | Sdr | 0.298468346 | 0.06551701 | 0 |
| RE03265 | FBgn0040255 | Ugt86De | -0.298455717 | 0.0797578 | 0 |
| GH25683 | FBgn0030745 | CG4239 | -0.298365045 | 0.11235733 | 0 |

| | | | | | |
|---|---|---|---|---|---|
| LD39110 | FBgn0263605 | l(3)72Dn | -0.298350037 | 0.056513 | 0 |
| LD41575 | FBgn0030582 | CG14411 | -0.298300582 | 0.06156526 | 0 |
| RE73906 | FBgn0020618 | Rack1 | 0.298264038 | 0.26514466 | 0 |
| GH17715 | FBgn0263864 | Dark | 0.298091837 | 0.17551784 | 0 |
| AT10967 | FBgn0030109 | CG12121 | 0.29800512 | 0.09325764 | 0 |
| SD07191 | FBgn0010830 | l(3)04053 | -0.297891398 | 0.17147856 | 0 |
| SD27341 | FBgn0024360 | MTPAP | -0.297772136 | 0.05825492 | 0 |
| RE28991 | FBgn0032066 | LManIII | -0.297365994 | 0.13466737 | 0 |
| LD44494 | FBgn0033088 | PGAP3 | -0.297121707 | 0.07367722 | 0 |
| AT15369 | FBgn0261086 | Syt14 | 0.297071476 | 0.12963797 | 0 |
| AT19250 | FBgn0031608 | CG15435 | -0.296917977 | 0.1115611 | 0 |
| LD15250 | FBgn0005640 | Eip63E | 0.29689049 | 0.07251516 | 0 |
| LD02432 | FBgn0034436 | CG11961 | 0.296834632 | 0.06712449 | 0 |
| AT11334 | FBgn0032431 | CG5435 | -0.296737313 | 0.0878206 | 0 |
| AT24025 | FBgn0033813 | fsd | -0.296456684 | 0.13738364 | 0 |
| LD24363 | FBgn0033178 | CG11127 | 0.296184602 | 0.08731422 | 0 |
| LD19162 | FBgn0031250 | Ent1 | -0.296103576 | 0.09679205 | 0 |
| AT12489 | FBgn0043457 | CG5180 | 0.295873156 | 0.11813587 | 0 |
| LD08259 | FBgn0086898 | dgo | 0.295862531 | 0.14164577 | 0 |
| GH24078 | FBgn0023520 | CG3857 | 0.295800976 | 0.06708516 | 0 |
| LD37006 | FBgn0267408 | AOX1 | 0.295788044 | 0.06022567 | 0 |
| AT19071 | FBgn0032111 | CG13110 | -0.295764236 | 0.36263426 | 0 |
| GH09086 | FBgn0015797 | Rab6 | -0.295659034 | 0.07553203 | 0 |
| GH20023 | FBgn0036479 | CG13458 | -0.295460145 | 0.1382545 | 0 |
| RE07815 | FBgn0040091 | Ugt58Fa | 0.295425567 | 0.38278109 | 0 |
| RH48810 | FBgn0039269 | veli | -0.295419032 | 0.0892183 | 0 |
| RH48588 | FBgn0050185 | AIMP3 | -0.295340579 | 0.09132812 | 0 |
| HL01056 | FBgn0260658 | CG42541 | -0.295198237 | 0.06025942 | 0 |
| LP08647 | FBgn0024294 | Spn43Aa | 0.29503135 | 0.35396165 | 0 |
| SD10416 | FBgn0038098 | CG7381 | 0.295002852 | 0.07380561 | 0 |
| LD46360 | FBgn0263198 | Acn | 0.294831646 | 0.05942862 | 0 |
| GH07643 | FBgn0027570 | Nep2 | 0.294463761 | 0.44088664 | 0 |
| LD30655 | FBgn0019637 | Atu | 0.294455029 | 0.10942324 | 0 |
| RE64686 | FBgn0027330 | l(1)G0020 | -0.294358853 | 0.09260753 | 0 |
| LP01562 | FBgn0034225 | veil | 0.294070318 | 0.12320472 | 0 |
| GM13546 | FBgn0021875 | Zfrp8 | -0.293867724 | 0.1037534 | 0 |
| AT14886 | FBgn0263974 | qin | -0.293760866 | 0.06025942 | 0 |
| GH12504 | FBgn0046874 | Pif1B | 0.293329436 | 0.07026664 | 0 |
| GH12504 | FBgn0261015 | Pif1A | 0.293329436 | 0.07026664 | 0 |
| LD05179 | FBgn0250874 | ttm50 | -0.293317546 | 0.10102147 | 0 |
| RE15854 | FBgn0002592 | E(spl)m2-BFM | -0.29331404 | 0.13675153 | 0 |
| GH07390 | FBgn0051445 | CG31445 | 0.293258475 | 0.08320475 | 0 |
| GH02419 | FBgn0027611 | LManII | 0.29324243 | 0.06981122 | 0 |
| GM02605 | FBgn0014427 | CG11899 | -0.293188225 | 0.05942862 | 0 |
| LD29234 | FBgn0033665 | Zip48C | 0.293108923 | 0.10196944 | 0 |
| RE17513 | FBgn0037743 | CG8412 | 0.292587842 | 0.09617792 | 0 |
| RH74035 | FBgn0001565 | Hlc | 0.292521734 | 0.05942862 | 0 |
| AT12204 | FBgn0030359 | CG18130 | -0.292509611 | 0.06455982 | 0 |

| | | | | | |
|---|---|---|---|---|---|
| GH12958 | FBgn0022097 | Vha36-1 | -0.292462228 | 0.10088317 | 0 |
| GH12958 | FBgn0040377 | Vha36-3 | -0.292462228 | 0.10088317 | 0 |
| LD26157 | FBgn0010282 | TfIIFalpha | 0.292429509 | 0.12947861 | 0 |
| GH01039 | FBgn0038693 | unc79 | -0.292332596 | 0.14701542 | 0 |
| LD09414 | FBgn0038272 | CG7265 | 0.292095359 | 0.11995584 | 0 |
| RE60063 | FBgn0040005 | CG17883 | -0.292015763 | 0.17551784 | 0 |
| LD26426 | FBgn0039600 | CG1646 | 0.291883773 | 0.11695164 | 0 |
| RE33814 | FBgn0030859 | CG12990 | 0.291670385 | 0.26189303 | 0 |
| LD39815 | FBgn0031769 | CG9135 | -0.291605684 | 0.06640401 | 0 |
| RH71439 | FBgn0260486 | Ziz | -0.291524997 | 0.12213767 | 0 |
| GH08189 | FBgn0003651 | svp | -0.291405719 | 0.26375792 | 0 |
| HL02501 | FBgn0010905 | Spn | 0.291334584 | 0.09361444 | 0 |
| GM13171 | FBgn0005666 | bt | 0.291183765 | 0.35769494 | 0 |
| GH14043 | FBgn0017551 | Rca1 | -0.291076617 | 0.07806278 | 0 |
| LD06441 | FBgn0035374 | mRpS35 | -0.290656079 | 0.15519681 | 0 |
| GH27496 | FBgn0039296 | CG10420 | -0.29052426 | 0.13844439 | 0 |
| RE44143 | FBgn0030474 | CG15747 | 0.290258066 | 0.06713041 | 0 |
| SD17036 | FBgn0023479 | teq | -0.290071393 | 0.28600853 | 0 |
| RE69454 | FBgn0034299 | CG5757 | -0.290022476 | 0.07439805 | 0 |
| GH11368 | FBgn0032881 | CG9319 | 0.290008817 | 0.09798082 | 0 |
| GH02902 | FBgn0261004 | asl | 0.289957414 | 0.08468448 | 0 |
| GH10112 | FBgn0002930 | nec | -0.289883943 | 0.29250904 | 0 |
| LP10725 | FBgn0035056 | spz6 | 0.289585711 | 0.43087554 | 0 |
| LD30050 | FBgn0037705 | mura | -0.289455158 | 0.15810151 | 0 |
| RE13973 | FBgn0030361 | CG1492 | -0.289433381 | 0.17597258 | 0 |
| GH23384 | FBgn0040609 | CG3348 | 0.289185515 | 0.0780263 | 0 |
| LD25801 | FBgn0261397 | didum | -0.289156338 | 0.07863671 | 0 |
| LD38705 | FBgn0035850 | Atg18a | 0.289138384 | 0.08975176 | 0 |
| LD24788 | FBgn0067102 | GlcT | -0.289067184 | 0.15131052 | 0 |
| AT03020 | FBgn0050115 | GEFmeso | 0.288655572 | 0.06417958 | 0 |
| AT20865 | FBgn0003943 | Ubi-p63E | -0.288558594 | 0.06712449 | 0 |
| GH04973 | FBgn0032790 | CG10194 | -0.288465407 | 0.12466267 | 0 |
| LD33453 | FBgn0015568 | alpha-Est1 | 0.288327725 | 0.08802717 | 0 |
| LD23336 | FBgn0027505 | Rab3-GAP | 0.288232082 | 0.09715278 | 0 |
| LD26240 | FBgn0033427 | Smyd4-1 | 0.288163159 | 0.13664789 | 0 |
| RE29447 | FBgn0051002 | CG31002 | 0.288060468 | 0.30080363 | 0 |
| RH09582 | FBgn0262976 | lawc | -0.288028002 | 0.17341214 | 0 |
| RE55639 | FBgn0052732 | CG32732 | -0.287926137 | 0.09618109 | 0 |
| RE09533 | FBgn0000826 | png | -0.287623468 | 0.17434219 | 0 |
| LD47396 | FBgn0037632 | CCT7 | -0.287533141 | 0.07367722 | 0 |
| LD21405 | FBgn0027528 | goe | 0.287245875 | 0.09679205 | 0 |
| LD15851 | FBgn0000308 | chic | -0.286705744 | 0.17551784 | 0 |
| LD19061 | FBgn0030499 | CG11178 | -0.286562908 | 0.07269309 | 0 |
| GM04427 | FBgn0034902 | CG5532 | 0.286526311 | 0.13597632 | 0 |
| LD21416 | FBgn0023511 | Edem1 | 0.286497531 | 0.06158456 | 0 |
| LD34829 | FBgn0034187 | CG6967 | -0.286407281 | 0.10163701 | 0 |
| LD23788 | FBgn0036144 | GlcAT-P | -0.286356985 | 0.07329076 | 0 |
| LD45754 | FBgn0026079 | Nsun2 | 0.2858554 | 0.08749629 | 0 |

| | | | | | |
|---|---|---|---|---|---|
| RE35942 | FBgn0038053 | CG18549 | -0.285516001 | 0.06259049 | 0 |
| RH01253 | FBgn0038400 | CG5903 | -0.285440473 | 0.06627111 | 0 |
| SD01812 | FBgn0034110 | Atg9 | 0.285160072 | 0.11689839 | 0 |
| LP03706 | FBgn0030360 | CG1806 | 0.284905498 | 0.24417029 | 0 |
| RE55091 | FBgn0033351 | AIMP1 | -0.284864766 | 0.11700678 | 0 |
| LD41958 | FBgn0032246 | Wdfy2 | -0.284739551 | 0.06178355 | 0 |
| GH14650 | FBgn0265084 | CG44195 | 0.28458479 | 0.53937384 | 0 |
| LD22513 | FBgn0033897 | Rcd1 | 0.284486397 | 0.07885761 | 0 |
| SD24551 | FBgn0038742 | Arc42 | -0.28446398 | 0.06627111 | 0 |
| LD18904 | FBgn0040918 | schlank | -0.284363245 | 0.06712449 | 0 |
| LD32761 | FBgn0028541 | TM9SF4 | -0.284262389 | 0.07838006 | 0 |
| GH12701 | FBgn0052432 | CG32432 | -0.284187388 | 0.34682994 | 0 |
| LP01841 | FBgn0000479 | dnc | -0.284132001 | 0.18784701 | 0 |
| RE27784 | FBgn0041630 | Hexo1 | 0.284056763 | 0.09486298 | 0 |
| RE48016 | FBgn0264753 | Rgk1 | -0.283952607 | 0.17637114 | 0 |
| GM13306 | FBgn0025885 | Inos | -0.283848592 | 0.14548021 | 0 |
| GH19029 | FBgn0032477 | CG5287 | 0.2838009 | 0.13844439 | 0 |
| RH03914 | FBgn0037307 | Tim17a2 | -0.283722911 | 0.24946171 | 0 |
| LP06209 | FBgn0034443 | cer | 0.283636193 | 0.11035759 | 0 |
| RE74364 | FBgn0029568 | CG11381 | -0.283507158 | 0.41681002 | 0 |
| GH24511 | FBgn0023143 | Uba1 | -0.283503976 | 0.0631004 | 0 |
| LD45906 | FBgn0027296 | temp | 0.283454392 | 0.09593355 | 0 |
| LD22661 | FBgn0032393 | CG12264 | -0.283239542 | 0.06713041 | 0 |
| SD02533 | FBgn0025185 | az2 | -0.282925672 | 0.06420995 | 0 |
| LD35885 | FBgn0034406 | Jheh3 | -0.282887127 | 0.11359747 | 0 |
| SD08771 | FBgn0264389 | opm | -0.282521174 | 0.07056036 | 0 |
| AT08308 | FBgn0037101 | CG7634 | -0.282313853 | 0.06885709 | 0 |
| RE46661 | FBgn0037754 | CG8500 | -0.282267196 | 0.1670928 | 0 |
| SD07276 | FBgn0039863 | CG1815 | 0.282249752 | 0.09541205 | 0 |
| LP07441 | FBgn0032171 | CG5846 | -0.282238015 | 0.15049835 | 0 |
| RE45450 | FBgn0036135 | mRpL2 | -0.28218778 | 0.16148597 | 0 |
| AT14262 | FBgn0038233 | HtrA2 | 0.282133163 | 0.07000688 | 0 |
| LD29228 | FBgn0046258 | CG12880 | 0.282114197 | 0.09890059 | 0 |
| SD08921 | FBgn0032026 | CG7627 | -0.282098739 | 0.11813587 | 0 |
| SD08921 | FBgn0051792 | CG31792 | -0.282098739 | 0.11813587 | 0 |
| LD23954 | FBgn0001098 | Gdh | 0.28206529 | 0.15892824 | 0 |
| LD43495 | FBgn0011771 | Hem | -0.28180951 | 0.10417336 | 0 |
| LD29172 | FBgn0036678 | CG11905 | 0.281627901 | 0.25905255 | 0 |
| LD41395 | FBgn0036761 | MED19 | -0.281600039 | 0.08647308 | 0 |
| LD25280 | FBgn0031413 | CG9967 | 0.281470447 | 0.11690063 | 0 |
| RH69634 | FBgn0034470 | Obp56d | 0.281373764 | 0.09230766 | 0 |
| RE22261 | FBgn0004892 | sob | -0.281224129 | 0.24657701 | 0 |
| RE34634 | FBgn0036485 | FucTA | -0.281200193 | 0.14126395 | 0 |
| RH51767 | FBgn0037419 | Osi12 | -0.281199259 | 0.45761209 | 0 |
| RE49489 | FBgn0030341 | p24-1 | -0.281121018 | 0.12558174 | 0 |
| RH66281 | FBgn0005664 | Crys | -0.281115028 | 0.23048196 | 0 |
| LD07143 | FBgn0034742 | CG4294 | 0.281071036 | 0.11597957 | 0 |
| SD07741 | FBgn0035253 | CG7971 | 0.281061735 | 0.16776507 | 0 |

| | | | | | |
|---|---|---|---|---|---|
| RE26840 | FBgn0053108 | CG33108 | 0.281057643 | 0.0847017 | 0 |
| GH08881 | FBgn0041210 | HDAC4 | -0.281026521 | 0.11228226 | 0 |
| GH04118 | FBgn0039925 | Kif3C | 0.280961168 | 0.14005329 | 0 |
| LD24267 | FBgn0026876 | CG11403 | -0.280896996 | 0.06640401 | 0 |
| RE25571 | FBgn0031306 | CG4577 | -0.280770653 | 0.23994281 | 0 |
| AT29108 | FBgn0002183 | dre4 | -0.280497393 | 0.06501956 | 0 |
| RE11615 | FBgn0039876 | CG2126 | -0.280416335 | 0.09751361 | 0 |
| LD24471 | FBgn0025336 | CG4882 | -0.280279066 | 0.10466062 | 0 |
| LD27706 | FBgn0264492 | CkIIalpha | -0.280273582 | 0.10804827 | 0 |
| GH18144 | FBgn0261641 | CG42724 | 0.280235285 | 0.06561597 | 0 |
| LD25351 | FBgn0020389 | Papss | -0.280183061 | 0.12610062 | 0 |
| RE27864 | FBgn0085444 | mute | -0.280032439 | 0.10434168 | 0 |
| GH05812 | FBgn0022986 | qkr58E-1 | 0.27959694 | 0.07380561 | 0 |
| RE11411 | FBgn0033524 | Cyp49a1 | -0.279560101 | 0.09460365 | 0 |
| GM02004 | FBgn0086674 | Tango13 | -0.279549982 | 0.30429754 | 0 |
| GM02004 | FBgn0284227 | CG46311 | -0.279549982 | 0.30429754 | 0 |
| LD36356 | FBgn0000212 | brm | 0.279490089 | 0.08982271 | 0 |
| LD38130 | FBgn0039227 | polybromo | 0.279410372 | 0.0973297 | 0 |
| LD46453 | FBgn0031390 | tho2 | 0.279395053 | 0.07415285 | 0 |
| SD05938 | FBgn0036772 | CG5290 | -0.27914213 | 0.06981122 | 0 |
| RE16687 | FBgn0015271 | Orc5 | -0.279027127 | 0.09008791 | 0 |
| GH05088 | FBgn0035978 | UGP | -0.278986092 | 0.07314481 | 0 |
| LD06106 | FBgn0038967 | CG13847 | -0.278884631 | 0.19562813 | 0 |
| RH32172 | FBgn0037164 | CG11438 | -0.278728975 | 0.06602777 | 0 |
| GM08679 | FBgn0028734 | Fmr1 | -0.278727462 | 0.17551784 | 0 |
| GH16587 | FBgn0020385 | pug | 0.278393813 | 0.10929121 | 0 |
| RE21160 | FBgn0034035 | CG8207 | 0.27837487 | 0.09615727 | 0 |
| LD29830 | FBgn0024285 | Srp54 | 0.277997397 | 0.12541392 | 0 |
| RE04530 | FBgn0035318 | CG9018 | 0.2778746 | 0.11235733 | 0 |
| RH37405 | FBgn0045064 | bwa | -0.277714148 | 0.14632387 | 0 |
| GH23387 | FBgn0032701 | CG10341 | 0.277566129 | 0.14164577 | 0 |
| RE32624 | FBgn0015338 | CG5861 | -0.277526476 | 0.07663489 | 0 |
| RE14985 | FBgn0035423 | eIF1 | -0.2773913 | 0.08386612 | 0 |
| GM30371 | FBgn0260660 | Mp | -0.277249626 | 0.44489043 | 0 |
| GM07526 | FBgn0052636 | lncRNA:CR32636 | 0.27712355 | 0.15721637 | 0 |
| LD43196 | FBgn0036988 | CG5262 | -0.276989416 | 0.11813587 | 0 |
| GH12017 | FBgn0261393 | alpha-Est5 | 0.276798571 | 0.07038909 | 0 |
| LD21546 | FBgn0031456 | Tnpo-SR | 0.276758858 | 0.06885709 | 0 |
| GH23506 | FBgn0037275 | CG14655 | 0.276540789 | 0.1115611 | 0 |
| RE53159 | FBgn0003888 | betaTub60D | 0.276436163 | 0.14521543 | 0 |
| RE53159 | FBgn0284243 | betaTub56D | 0.276436163 | 0.14521543 | 0 |
| LD34822 | FBgn0020272 | mst | -0.276341066 | 0.10108162 | 0 |
| LD33103 | FBgn0030946 | CG6659 | 0.276184606 | 0.1246311 | 0 |
| GH08896 | FBgn0035283 | CG12024 | -0.276165779 | 0.06807467 | 0 |
| RE43278 | FBgn0035122 | mRpL17 | -0.276057107 | 0.14406727 | 0 |
| AT20771 | FBgn0030395 | Mks1 | -0.276042655 | 0.1246311 | 0 |
| LP12120 | FBgn0261016 | clos | -0.275931562 | 0.06943885 | 0 |
| RE72948 | FBgn0035825 | CG8111 | -0.275918761 | 0.07367722 | 0 |

| | | | | | |
|---|---|---|---|---|---|
| RH41322 | FBgn0034817 | Art7 | -0.275899573 | 0.12417497 | 0 |
| LD40339 | FBgn0260936 | scny | 0.275861007 | 0.07731902 | 0 |
| LD05259 | FBgn0026077 | Gasp | 0.2757507 | 0.73117965 | 0 |
| LD28447 | FBgn0037282 | CG14657 | 0.275562065 | 0.09890059 | 0 |
| LD44072 | FBgn0029906 | xit | 0.275456744 | 0.15705273 | 0 |
| GH08607 | FBgn0020278 | loco | 0.275293852 | 0.06885709 | 0 |
| RH45940 | FBgn0038396 | CG5013 | 0.275205482 | 0.09715278 | 0 |
| RH10688 | FBgn0032916 | CG9257 | -0.275096483 | 0.17147856 | 0 |
| GH08708 | FBgn0023516 | Pex5 | -0.275043708 | 0.06712449 | 0 |
| LD34157 | FBgn0039094 | CG10184 | -0.27502977 | 0.0757301 | 0 |
| LD43270 | FBgn0000351 | cort | -0.274994241 | 0.06960081 | 0 |
| LD38742 | FBgn0030874 | Spt7 | 0.274625151 | 0.09361444 | 0 |
| RH02912 | FBgn0032321 | YL-1 | 0.274545244 | 0.16321212 | 0 |
| LD01527 | FBgn0032723 | ssp3 | 0.274450372 | 0.08408769 | 0 |
| LD23810 | FBgn0034572 | CG9346 | 0.274270047 | 0.12227708 | 0 |
| LD46713 | FBgn0033584 | CG7737 | 0.274247429 | 0.11014912 | 0 |
| AT08391 | FBgn0036962 | CG17122 | 0.274194442 | 0.07573639 | 0 |
| RE28575 | FBgn0039069 | CG6763 | 0.273885438 | 0.1115611 | 0 |
| LD39559 | FBgn0036004 | Jarid2 | 0.273577391 | 0.14961497 | 0 |
| SD13756 | FBgn0039488 | CG6066 | 0.273450755 | 0.08688202 | 0 |
| SD09884 | FBgn0037207 | Mes2 | -0.272942095 | 0.09668679 | 0 |
| LD07162 | FBgn0024183 | vig | 0.272801995 | 0.0995265 | 0 |
| LD33443 | FBgn0028530 | mTTF | 0.272700627 | 0.14124168 | 0 |
| RH13166 | FBgn0264979 | CG4267 | -0.272553933 | 0.12070859 | 0 |
| RH33338 | FBgn0036146 | CG14141 | -0.272501792 | 0.11027617 | 0 |
| GH27933 | FBgn0260817 | gkt | -0.272480744 | 0.09826972 | 0 |
| GH27933 | FBgn0051953 | CR31953 | -0.272480744 | 0.09826972 | 0 |
| LD37523 | FBgn0010750 | atms | 0.272395817 | 0.09618247 | 0 |
| LD31286 | FBgn0261592 | RpS6 | 0.27238289 | 0.18250817 | 0 |
| LD31286 | FBgn0082924 | snoRNA:Or-aca2 | 0.27238289 | 0.18250817 | 0 |
| LD13706 | FBgn0036433 | CG9628 | 0.272219441 | 0.36027893 | 0 |
| LD41750 | FBgn0035947 | Srp68 | -0.272132474 | 0.07570279 | 0 |
| LD40271 | FBgn0283525 | l(2)10685 | 0.272105548 | 0.1137627 | 0 |
| GH01524 | FBgn0001248 | Idh | -0.271985899 | 0.08244431 | 0 |
| RE72803 | FBgn0036622 | CG4753 | -0.271842538 | 0.06713041 | 0 |
| GH26159 | FBgn0031305 | Iris | -0.271829074 | 0.20077053 | 0 |
| RE09471 | FBgn0039768 | CG15533 | -0.271791815 | 0.43026786 | 0 |
| RE45155 | FBgn0032020 | strat | -0.271494659 | 0.10316195 | 0 |
| RE36860 | FBgn0003187 | qua | -0.271377563 | 0.07570279 | 0 |
| RE73545 | FBgn0050265 | CG30265 | 0.271157911 | 0.1193711 | 0 |
| RE42502 | FBgn0034523 | Nnf1a | 0.271074416 | 0.16476127 | 0 |
| GH01661 | FBgn0058263 | MFS17 | -0.270979373 | 0.11055415 | 0 |
| LD33358 | FBgn0039631 | Sirt7 | 0.270907499 | 0.10325439 | 0 |
| RE34879 | FBgn0031813 | CG9527 | 0.270900844 | 0.1888044 | 0 |
| GH08511 | FBgn0051231 | CG31231 | -0.270802141 | 0.21267823 | 0 |
| LD09942 | FBgn0037344 | CG2926 | 0.270785859 | 0.09173843 | 0 |
| LD17962 | FBgn0003459 | stwl | 0.270782619 | 0.1241051 | 0 |
| LD08432 | FBgn0038391 | GATAe | -0.270773488 | 0.31912746 | 0 |

| | | | | | |
|---|---|---|---|---|---|
| GM04742 | FBgn0285917 | sbb | 0.2707327 | 0.10890456 | 0 |
| GH12043 | FBgn0050389 | CG30389 | -0.270684683 | 0.13997939 | 0 |
| RE28066 | FBgn0035812 | CG7457 | -0.270650109 | 0.07816593 | 0 |
| AT16284 | FBgn0037310 | Tim17b1 | 0.270643928 | 0.33074938 | 0 |
| LD41866 | FBgn0032646 | mEFTs | -0.270491138 | 0.08731422 | 0 |
| LP06948 | FBgn0261053 | Cad86C | 0.270448215 | 0.08146267 | 0 |
| LD24920 | FBgn0026598 | Apc2 | 0.270442497 | 0.09445121 | 0 |
| LD47625 | FBgn0266918 | CG32486 | 0.270339381 | 0.1822614 | 0 |
| GH04763 | FBgn0034921 | DCP1 | 0.270319446 | 0.12320472 | 0 |
| RE09053 | FBgn0032732 | CG15168 | -0.270231257 | 0.10151555 | 0 |
| RE64534 | FBgn0039118 | CG10208 | 0.270198375 | 0.16476127 | 0 |
| RE22905 | FBgn0032856 | CG16798 | -0.270148747 | 0.38135654 | 0 |
| RH48004 | FBgn0038405 | CG8927 | -0.270139088 | 0.09698692 | 0 |
| LD40766 | FBgn0039690 | Gnpnat | 0.270133938 | 0.11102126 | 0 |
| LD46585 | FBgn0061200 | Nup153 | 0.270129972 | 0.12487471 | 0 |
| GM02829 | FBgn0037877 | CG6689 | 0.270103451 | 0.07263677 | 0 |
| SD10554 | FBgn0031012 | CG8051 | -0.269996991 | 0.37074408 | 0 |
| SD10554 | FBgn0065084 | snmRNA:763 | -0.269996991 | 0.37074408 | 0 |
| SD02216 | FBgn0030086 | CCT2 | -0.269824698 | 0.09715278 | 0 |
| LD15640 | FBgn0266418 | wake | 0.269806505 | 0.08975176 | 0 |
| GH03058 | FBgn0036992 | CG11796 | -0.269544592 | 0.09259409 | 0 |
| LP04613 | FBgn0038804 | CG10877 | 0.26952419 | 0.17917133 | 0 |
| RE65495 | FBgn0025739 | pon | -0.269446212 | 0.14477381 | 0 |
| RE67590 | FBgn0017577 | Mcm5 | -0.26939067 | 0.0757301 | 0 |
| LD15362 | FBgn0267967 | corolla | 0.269311495 | 0.08865723 | 0 |
| RE18955 | FBgn0024920 | Ts | -0.269283321 | 0.07854661 | 0 |
| LD24467 | FBgn0028990 | Spn27A | 0.269259029 | 0.1388353 | 0 |
| AT15146 | FBgn0261396 | Rpn3 | -0.26917167 | 0.11813587 | 0 |
| RE24967 | FBgn0034859 | CG3520 | 0.269054034 | 0.07823232 | 0 |
| GH01043 | FBgn0038597 | CG8064 | 0.269023779 | 0.14709825 | 0 |
| GH08163 | FBgn0037908 | dpr5 | -0.26896967 | 0.13597632 | 0 |
| GM06595 | FBgn0029899 | CG14438 | 0.268952373 | 0.19610699 | 0 |
| LD36653 | FBgn0033199 | CG17985 | 0.268934803 | 0.12042901 | 0 |
| LD07688 | FBgn0038760 | MED25 | -0.268814775 | 0.19666058 | 0 |
| GM09620 | FBgn0263599 | l(3)72Ab | 0.268759721 | 0.06885709 | 0 |
| RH40291 | FBgn0038914 | fit | -0.268581043 | 0.16096423 | 0 |
| RH40291 | FBgn0266406 | asRNA:CR45046 | -0.268581043 | 0.16096423 | 0 |
| LD42169 | FBgn0037978 | KLHL18 | 0.268421962 | 0.11070818 | 0 |
| RE34115 | FBgn0033108 | CG15236 | -0.268409551 | 0.1389816 | 0 |
| GH09064 | FBgn0033273 | Gasz | -0.268266687 | 0.0711582 | 0 |
| RE02452 | FBgn0025683 | CG3164 | -0.26822996 | 0.15390186 | 0 |
| HL01538 | FBgn0037315 | Cerk | -0.268189807 | 0.10243236 | 0 |
| RE23429 | FBgn0032129 | jp | -0.268136944 | 0.08974365 | 0 |
| LD46083 | FBgn0028336 | l(1)G0255 | -0.268100615 | 0.09204205 | 0 |
| LD23868 | FBgn0029820 | CG16721 | 0.267979511 | 0.07365015 | 0 |
| GM14202 | FBgn0038183 | CG9286 | 0.26791167 | 0.17147856 | 0 |
| RH55324 | FBgn0039622 | eIF4E6 | -0.267796962 | 0.07045958 | 0 |
| LD21533 | FBgn0004556 | Dbp73D | -0.267727181 | 0.59873026 | 0 |

| | | | | | |
|---|---|---|---|---|---|
| SD14289 | FBgn0033810 | CG4646 | -0.26757282 | 0.14039527 | 0 |
| RH68528 | FBgn0032283 | CG7296 | 0.267521158 | 0.09762485 | 0 |
| SD01201 | FBgn0262517 | Ltn1 | -0.267381624 | 0.09362642 | 0 |
| GH12850 | FBgn0050121 | lncRNA:CR30121 | -0.267315481 | 0.34575348 | 0 |
| RE36610 | FBgn0038734 | CG11453 | -0.26726579 | 0.40921741 | 0 |
| LD37470 | FBgn0038206 | twf | -0.26721131 | 0.26375792 | 0 |
| GH04007 | FBgn0033751 | CG8818 | -0.267174236 | 0.08575996 | 0 |
| GH11113 | FBgn0263199 | Galk | -0.267034937 | 0.12052351 | 0 |
| LD36863 | FBgn0035247 | metl | 0.266757939 | 0.09890059 | 0 |
| LD27718 | FBgn0002719 | Men | -0.266626574 | 0.07088605 | 0 |
| GH27708 | FBgn0016694 | Pdp1 | -0.26642897 | 0.29393334 | 0 |
| SD10041 | FBgn0086657 | IKKepsilon | -0.266417939 | 0.42280478 | 0 |
| LD40310 | FBgn0031759 | lid | 0.266273005 | 0.07780594 | 0 |
| RE07731 | FBgn0020443 | eRF3 | -0.266091294 | 0.0752271 | 0 |
| RE52740 | FBgn0023180 | Orc6 | -0.265784276 | 0.0973297 | 0 |
| GH14884 | FBgn0033426 | CG1814 | 0.265707646 | 0.07410927 | 0 |
| LD47893 | FBgn0028982 | Spt6 | 0.265426561 | 0.07895628 | 0 |
| LD15217 | FBgn0015278 | Pi3K68D | 0.26521241 | 0.22660325 | 0 |
| LD46483 | FBgn0026582 | Hmg-2 | 0.265201611 | 0.1115611 | 0 |
| GH23271 | FBgn0082582 | tmod | -0.265086706 | 0.11670221 | 0 |
| LD02793 | FBgn0261119 | Prp19 | 0.264935581 | 0.12320472 | 0 |
| LP06502 | FBgn0034542 | Fem-1 | -0.264931558 | 0.07570279 | 0 |
| GH01265 | FBgn0029131 | Debcl | -0.264804428 | 0.12020594 | 0 |
| GM24121 | FBgn0086613 | Ino80 | 0.264767414 | 0.17341214 | 0 |
| AT14896 | FBgn0011817 | nmo | 0.264726225 | 0.17110273 | 0 |
| AT14896 | FBgn0032131 | CG3841 | 0.264726225 | 0.17110273 | 0 |
| RE29005 | FBgn0016076 | vri | -0.264535213 | 0.25022018 | 0 |
| LD37409 | FBgn0037467 | Ufl1 | 0.264445016 | 0.16096423 | 0 |
| GH07469 | FBgn0259481 | Mob2 | -0.264171754 | 0.08974365 | 0 |
| RE09079 | FBgn0037379 | CG10979 | -0.264164995 | 0.09915304 | 0 |
| SD10560 | FBgn0035149 | MED30 | -0.264104514 | 0.16010887 | 0 |
| LD45348 | FBgn0010772 | Xe7 | 0.264021812 | 0.08743204 | 0 |
| LD18634 | FBgn0028999 | nerfin-1 | 0.263989496 | 0.5354438 | 0 |
| RH56744 | FBgn0032694 | MESR3 | 0.263975081 | 0.13825583 | 0 |
| GM29503 | FBgn0032727 | CG10623 | 0.263888664 | 0.1088733 | 0 |
| LD32677 | FBgn0029888 | ND-ASHI | 0.263411055 | 0.1246311 | 0 |
| AT06648 | FBgn0030290 | CG11756 | 0.263410536 | 0.40923721 | 0 |
| AT19506 | FBgn0038532 | CG14322 | -0.263211833 | 0.08367965 | 0 |
| GH01208 | FBgn0032029 | CG17292 | -0.263105171 | 0.0892183 | 0 |
| LD08718 | FBgn0038769 | CG10889 | -0.263050042 | 0.23906935 | 0 |
| RE16941 | FBgn0263038 | CG43333 | -0.262684692 | 0.25511933 | 0 |
| LD30488 | FBgn0036887 | CG9231 | -0.262678195 | 0.31044547 | 0 |
| LD39460 | FBgn0033994 | CG7544 | -0.262340788 | 0.08761806 | 0 |
| RE10847 | FBgn0024807 | DIP1 | 0.262310505 | 0.13478189 | 0 |
| GH15213 | FBgn0030052 | CG12065 | -0.262282431 | 0.12397101 | 0 |
| LD21701 | FBgn0000377 | crn | 0.26217256 | 0.11689839 | 0 |
| LD09732 | FBgn0000330 | cm | 0.262127962 | 0.17107296 | 0 |
| LD16340 | FBgn0259704 | Nsun5 | -0.262024022 | 0.1156087 | 0 |

| | | | | | |
|---|---|---|---|---|---|
| RE64548 | FBgn0010768 | sqz | -0.26190826 | 0.17523824 | 0 |
| RH14671 | FBgn0053178 | CG33178 | -0.261877428 | 0.07573639 | 0 |
| GH08778 | FBgn0028987 | Spn28F | -0.261727011 | 0.09698692 | 0 |
| GH08778 | FBgn0028986 | Spn38F | -0.261727011 | 0.09698692 | 0 |
| LD17530 | FBgn0028691 | Rpn9 | -0.261652578 | 0.09256191 | 0 |
| RE28679 | FBgn0035736 | Cpr65Eb | -0.261588437 | 0.42553425 | 0 |
| AT29333 | FBgn0264953 | Piezo | 0.26156541 | 0.21689011 | 0 |
| GH04024 | FBgn0265089 | eIF4E3 | -0.261544861 | 0.12834476 | 0 |
| RE33242 | FBgn0000261 | Cat | 0.261505609 | 0.09008791 | 0 |
| LD21941 | FBgn0014868 | Ost48 | 0.261492338 | 0.11690063 | 0 |
| GM02209 | FBgn0266557 | kis | 0.261261017 | 0.15704603 | 0 |
| GH28416 | FBgn0011270 | Pglym87 | -0.261108928 | 0.08646407 | 0 |
| GH28416 | FBgn0014869 | Pglym78 | -0.261108928 | 0.08646407 | 0 |
| SD07279 | FBgn0032147 | IP3K1 | -0.260983719 | 0.13478189 | 0 |
| LD14646 | FBgn0031597 | CG17612 | 0.260983116 | 0.15567205 | 0 |
| LP02352 | FBgn0038953 | CG18596 | -0.260873035 | 0.12924475 | 0 |
| LD10754 | FBgn0000119 | arr | 0.260773989 | 0.07851049 | 0 |
| RE17985 | FBgn0033816 | CG4679 | -0.260763063 | 0.10180505 | 0 |
| RE28596 | FBgn0010228 | HmgZ | 0.260759239 | 0.32237304 | 0 |
| AT09333 | FBgn0033258 | CG8712 | -0.260731594 | 0.09751361 | 0 |
| LD31018 | FBgn0015553 | tos | -0.260686501 | 0.07684257 | 0 |
| GH14576 | FBgn0039478 | Nep5 | -0.260568096 | 0.335791 | 0 |
| LD29726 | FBgn0034310 | Nup75 | -0.26054927 | 0.09204205 | 0 |
| SD05424 | FBgn0034728 | rad50 | -0.260338394 | 0.10291795 | 0 |
| RE10833 | FBgn0085391 | trv | -0.260237816 | 0.11005477 | 0 |
| GM14501 | FBgn0035600 | Cyt-c1 | 0.260154249 | 0.11070818 | 0 |
| GH16320 | FBgn0030304 | Cyp4g15 | -0.260139985 | 0.07570279 | 0 |
| RE72594 | FBgn0000114 | bru1 | -0.260092182 | 0.08749629 | 0 |
| RE25730 | FBgn0039993 | CG17691 | 0.260048741 | 0.32785535 | 0 |
| RE27026 | FBgn0037108 | Alg11 | -0.260001279 | 0.07851049 | 0 |
| RE46159 | FBgn0035431 | CG14968 | 0.259938473 | 0.10816559 | 0 |
| RE12154 | FBgn0032945 | CG8665 | 0.259891111 | 0.1876974 | 0 |
| LD07994 | FBgn0021825 | DCTN2-p50 | 0.259634751 | 0.15232792 | 0 |
| GH24947 | FBgn0039324 | CG10553 | 0.259521092 | 0.18075619 | 0 |
| RH49854 | FBgn0000536 | eas | -0.259454923 | 0.35435657 | 0 |
| RE61341 | FBgn0261791 | SNRPG | -0.259386688 | 0.11842639 | 0 |
| GH10766 | FBgn0046301 | CG7148 | -0.25935994 | 0.09762485 | 0 |
| RE33408 | FBgn0000562 | egl | -0.259258953 | 0.07968675 | 0 |
| LD41216 | FBgn0038666 | Smu1 | 0.259211735 | 0.0973297 | 0 |
| LD21334 | FBgn0020236 | ATPCL | -0.259195138 | 0.07700924 | 0 |
| SD26153 | FBgn0036300 | Pmm2 | -0.259047505 | 0.08025093 | 0 |
| LD43883 | FBgn0037382 | Hpr1 | 0.258979147 | 0.09618109 | 0 |
| LD31988 | FBgn0033507 | CG12909 | 0.258943058 | 0.10929121 | 0 |
| LD43989 | FBgn0263289 | scrib | 0.258689708 | 0.12070859 | 0 |
| LD29482 | FBgn0031118 | RhoGAP19D | 0.258617001 | 0.0973297 | 0 |
| RH41996 | FBgn0086367 | t | -0.258612399 | 0.2767948 | 0 |
| LD25543 | FBgn0086346 | ALiX | 0.258540943 | 0.16763613 | 0 |
| RE14068 | FBgn0029830 | Grip | -0.258309084 | 0.33403681 | 0 |

| | | | | | |
|---|---|---|---|---|---|
| SD02746 | FBgn0040777 | CG14767 | -0.258229219 | 0.0878206 | 0 |
| GH04732 | FBgn0037747 | CG8481 | -0.258036061 | 0.10572161 | 0 |
| GH25768 | FBgn0031536 | Cog3 | -0.257920495 | 0.09260753 | 0 |
| RE09732 | FBgn0037332 | Hcs | -0.257876893 | 0.09541205 | 0 |
| LD23965 | FBgn0026189 | prominin-like | 0.257667889 | 0.10163701 | 0 |
| RE31184 | FBgn0036774 | mRpS26 | -0.257659104 | 0.08031814 | 0 |
| SD01515 | FBgn0020764 | Alas | -0.257647535 | 0.14632387 | 0 |
| LP06455 | FBgn0037412 | Osi4 | -0.257500902 | 0.43735269 | 0 |
| RE33994 | FBgn0036899 | tey | -0.257386349 | 0.41907068 | 0 |
| RE04051 | FBgn0015929 | dpa | -0.257371427 | 0.12213767 | 0 |
| RE16426 | FBgn0039453 | CG6403 | -0.257343573 | 0.09325764 | 0 |
| LD43175 | FBgn0032601 | yellow-b | -0.257257486 | 0.25718929 | 0 |
| LP03341 | FBgn0050035 | Tret1-1 | 0.257240137 | 0.40921741 | 0 |
| GH12365 | FBgn0051004 | mesh | 0.257118691 | 0.34693537 | 0 |
| LD32609 | FBgn0031360 | CG31937 | 0.257043588 | 0.22005916 | 0 |
| GH23783 | FBgn0027376 | rha | -0.256849255 | 0.49472135 | 0 |
| GH23617 | FBgn0036684 | CG3764 | -0.256791937 | 0.0878206 | 0 |
| LD46870 | FBgn0037220 | CG14641 | 0.256784852 | 0.12497934 | 0 |
| RE34084 | FBgn0033708 | CG8850 | -0.256737272 | 0.12166498 | 0 |
| GM21109 | FBgn0033526 | Caf1-105 | -0.256693858 | 0.14614309 | 0 |
| GH23546 | FBgn0086687 | Desat1 | -0.256639452 | 0.07729214 | 0 |
| AT21555 | FBgn0025692 | Lfg | 0.256597473 | 0.30935965 | 0 |
| GH23906 | FBgn0085446 | CG34417 | -0.256548117 | 0.2412154 | 0 |
| GH16431 | FBgn0039491 | CG6059 | -0.256516994 | 0.1325229 | 0 |
| LD05563 | FBgn0011802 | Gem3 | 0.256510845 | 0.13968045 | 0 |
| GH07964 | FBgn0260462 | CG12163 | -0.256502314 | 0.45761209 | 0 |
| SD06439 | FBgn0036844 | Mkp3 | -0.256460551 | 0.0934684 | 0 |
| RE14858 | FBgn0031114 | cactin | 0.256439292 | 0.12590333 | 0 |
| LD48056 | FBgn0033806 | FLASH | 0.256401009 | 0.08622161 | 0 |
| RE57138 | FBgn0030217 | CG2124 | 0.256327867 | 0.19136517 | 0 |
| LD22863 | FBgn0011705 | rost | -0.256278548 | 0.18669852 | 0 |
| GH28782 | FBgn0025456 | CREG | 0.256240241 | 0.18571847 | 0 |
| RE50415 | FBgn0037981 | Spt3 | 0.256062608 | 0.10102147 | 0 |
| GH20759 | FBgn0027556 | CG4928 | 0.255954057 | 0.08320475 | 0 |
| LD25952 | FBgn0031478 | CG8814 | -0.255802397 | 0.08612406 | 0 |
| RH48056 | FBgn0037686 | RpL34b | 0.255757547 | 0.14854672 | 0 |
| LP06066 | FBgn0032289 | CG6431 | 0.255755812 | 0.13597632 | 0 |
| GH10751 | FBgn0003429 | slo | -0.255613309 | 0.38902664 | 0 |
| GH11193 | FBgn0015794 | Rab18 | -0.255535565 | 0.12421942 | 0 |
| LD26477 | FBgn0039733 | CG11504 | 0.255486405 | 0.15810151 | 0 |
| GH03217 | FBgn0052095 | CG32095 | 0.255461045 | 0.0797578 | 0 |
| GH10421 | FBgn0000721 | for | 0.255456304 | 0.08019083 | 0 |
| GH15037 | FBgn0037518 | CG2641 | 0.255314369 | 0.1246311 | 0 |
| GH07076 | FBgn0036159 | CG7557 | -0.255280437 | 0.1939671 | 0 |
| RE67480 | FBgn0039339 | CG5116 | -0.255279217 | 0.08432958 | 0 |
| LD12631 | FBgn0003963 | ush | 0.25522792 | 0.14709825 | 0 |
| SD05004 | FBgn0010339 | 128up | -0.25518335 | 0.08743204 | 0 |
| LD21785 | FBgn0036449 | bmm | 0.255160175 | 0.13966772 | 0 |

| | | | | | |
|---|---|---|---|---|---|
| RE32772 | FBgn0015803 | RtGEF | 0.255080884 | 0.14601911 | 0 |
| LD21192 | FBgn0027535 | botv | 0.254987295 | 0.10151555 | 0 |
| LD44207 | FBgn0261436 | DhpD | -0.254929676 | 0.1115611 | 0 |
| LD24929 | FBgn0034432 | CG7461 | -0.254886291 | 0.08180731 | 0 |
| LD10183 | FBgn0051457 | CG31457 | 0.254656263 | 0.0862875 | 0 |
| GH14572 | FBgn0034200 | Gbp2 | 0.254644906 | 0.22531609 | 0 |
| LP10040 | FBgn0001226 | Hsp27 | -0.254527393 | 0.12968005 | 0 |
| GH06348 | FBgn0027580 | PCB | -0.254142338 | 0.08743204 | 0 |
| RE56731 | FBgn0034408 | sano | -0.254097333 | 0.11099012 | 0 |
| SD27354 | FBgn0002354 | l(3)87Df | -0.254051061 | 0.16394152 | 0 |
| GH01941 | FBgn0262166 | calypso | -0.253936409 | 0.11009579 | 0 |
| LD32019 | FBgn0035798 | frac | 0.253876851 | 0.19968924 | 0 |
| GH21935 | FBgn0019949 | Cdk9 | 0.253781862 | 0.10037077 | 0 |
| GH06673 | FBgn0034796 | CG3700 | 0.253616718 | 0.11235733 | 0 |
| RE01471 | FBgn0036450 | Tdrd3 | -0.253515013 | 0.07810369 | 0 |
| RH34107 | FBgn0037801 | CG3999 | -0.253388488 | 0.24557335 | 0 |
| HL01076 | FBgn0260746 | Ect3 | 0.253247691 | 0.08647308 | 0 |
| RH54985 | FBgn0032021 | CG7781 | -0.253100602 | 0.1328197 | 0 |
| RH11203 | FBgn0047038 | ND-13B | -0.253095548 | 0.08337318 | 0 |
| RE36568 | FBgn0025635 | CG17829 | 0.25305392 | 0.1037592 | 0 |
| LD07113 | FBgn0263047 | CG43342 | 0.252956702 | 0.08025093 | 0 |
| RE36502 | FBgn0039544 | CG12877 | 0.252845078 | 0.09679205 | 0 |
| RE29421 | FBgn0011768 | Fdh | -0.25274785 | 0.10055182 | 0 |
| GM04108 | FBgn0038269 | Rrp6 | 0.252700385 | 0.09593355 | 0 |
| LD28839 | FBgn0015331 | abs | 0.252691071 | 0.10466062 | 0 |
| LP11031 | FBgn0028518 | CG18480 | -0.252666418 | 0.39340434 | 0 |
| LD28101 | FBgn0052344 | CG32344 | 0.252614614 | 0.15688323 | 0 |
| LD24983 | FBgn0038576 | Arp5 | 0.252581442 | 0.10942324 | 0 |
| RE62969 | FBgn0029711 | Usf | 0.252475032 | 0.14348413 | 0 |
| GH05842 | FBgn0032704 | Jwa | -0.252468925 | 0.10434168 | 0 |
| LD13436 | FBgn0033206 | DCTN4-p62 | -0.252462262 | 0.1018806 | 0 |
| RE37904 | FBgn0020880 | CG7179 | -0.252413502 | 0.26188951 | 0 |
| LD35285 | FBgn0086694 | Bre1 | 0.252264049 | 0.08974365 | 0 |
| AT04375 | FBgn0032457 | CG15483 | -0.25216682 | 0.09542047 | 0 |
| RH44301 | FBgn0052076 | Alg10 | -0.252097805 | 0.20926155 | 0 |
| RE52572 | FBgn0038722 | Nup58 | 0.251868713 | 0.24662826 | 0 |
| RE48939 | FBgn0015818 | Spx | 0.251816521 | 0.08974365 | 0 |
| SD02315 | FBgn0014269 | prod | 0.251801232 | 0.12988469 | 0 |
| LD45339 | FBgn0038303 | SIDL | 0.251730613 | 0.08423184 | 0 |
| GH10523 | FBgn0051388 | CG31388 | 0.251688516 | 0.19815676 | 0 |
| RE39820 | FBgn0023167 | SmD3 | -0.25167424 | 0.40236089 | 0 |
| GM08588 | FBgn0052280 | CG32280 | 0.251625554 | 0.13478189 | 0 |
| LD10776 | FBgn0017590 | klg | 0.25162298 | 0.46935885 | 0 |
| GH19318 | FBgn0001320 | kni | 0.251496363 | 0.16201098 | 0 |
| AT09846 | FBgn0085206 | CG34177 | -0.251442399 | 0.14560979 | 0 |
| AT09846 | FBgn0085207 | CG34178 | -0.251442399 | 0.14560979 | 0 |
| SD11994 | FBgn0035603 | Pfdn4 | -0.251246201 | 0.15677902 | 0 |
| RH43267 | FBgn0039665 | CG2310 | 0.251238108 | 0.19593722 | 0 |

| | | | | | |
|---|---|---|---|---|---|
| GM09489 | FBgn0010747 | Srp54k | 0.251182671 | 0.08141315 | 0 |
| GH03085 | FBgn0033819 | CG4714 | -0.251102671 | 0.16620492 | 0 |
| RE53941 | FBgn0040079 | pkaap | -0.250835157 | 0.09075684 | 0 |
| LD45449 | FBgn0030569 | CG9411 | -0.250798485 | 0.1325715 | 0 |
| LD19052 | FBgn0262475 | bru2 | -0.250785292 | 0.19806393 | 0 |
| RE64695 | FBgn0035854 | CG8005 | 0.25060837 | 0.11055415 | 0 |
| GH07275 | FBgn0033631 | Sod3 | -0.250514242 | 0.13131524 | 0 |
| RE55542 | FBgn0041711 | yellow-e | 0.250478557 | 0.28754289 | 0 |
| LD06317 | FBgn0035148 | CG3402 | -0.250416558 | 0.12320472 | 0 |
| LP02994 | FBgn0038647 | CG14302 | -0.250386274 | 0.43855477 | 0 |
| SD10657 | FBgn0024754 | Flo1 | 0.250373081 | 0.09059039 | 0 |
| SD03546 | FBgn0032050 | CG13096 | 0.250112262 | 0.12020594 | 0 |
| AT25633 | FBgn0035407 | Asciz | -0.25006459 | 0.09008791 | 0 |
| GM01935 | FBgn0019938 | RpI1 | -0.250011294 | 0.09484121 | 0 |
| LP09632 | FBgn0035047 | Pof | -0.249787865 | 0.09992866 | 0 |
| GH09663 | FBgn0026147 | TTLL4A | -0.249761879 | 0.0878206 | 0 |
| LD13945 | FBgn0010501 | Dcp-1 | 0.249746112 | 0.08830214 | 0 |
| LD44262 | FBgn0034270 | PIG-A | -0.249716523 | 0.11697673 | 0 |
| LD15072 | FBgn0033842 | cbc | 0.249707142 | 0.1088733 | 0 |
| LD42274 | FBgn0026323 | Tak1 | 0.249517578 | 0.1018806 | 0 |
| RE09582 | FBgn0013759 | CASK | -0.249443776 | 0.20152257 | 0 |
| RE09582 | FBgn0266181 | lncRNA:CRG | -0.249443776 | 0.20152257 | 0 |
| GH07323 | FBgn0263390 | Nha2 | -0.249338659 | 0.31688128 | 0 |
| GH12831 | FBgn0051199 | CG31199 | -0.249323605 | 0.10291795 | 0 |
| LD30575 | FBgn0031450 | Hrs | 0.249213582 | 0.10986156 | 0 |
| LD27857 | FBgn0019686 | lok | -0.24906051 | 0.11099012 | 0 |
| LD40415 | FBgn0031739 | CG14005 | 0.249021982 | 0.09542047 | 0 |
| LD47736 | FBgn0022359 | Sodh-2 | 0.248879489 | 0.18582273 | 0 |
| RE69515 | FBgn0031049 | Sec61gamma | -0.24871794 | 0.14097266 | 0 |
| GH10091 | FBgn0052445 | CG32445 | -0.248670196 | 0.0973297 | 0 |
| RE19842 | FBgn0028663 | VhaM9.7-b | -0.248651312 | 0.16192191 | 0 |
| GM04669 | FBgn0034365 | CG5335 | -0.248636811 | 0.12685445 | 0 |
| GH03685 | FBgn0015795 | Rab7 | -0.248578084 | 0.10942324 | 0 |
| GH09711 | FBgn0051547 | NKCC | -0.248513369 | 0.12227708 | 0 |
| GH12331 | FBgn0052600 | dpr8 | 0.248468051 | 0.12506741 | 0 |
| RE61257 | FBgn0036390 | CG13738 | -0.248427908 | 0.51270727 | 0 |
| LP08894 | FBgn0038180 | Cht5 | 0.248415036 | 0.31903254 | 0 |
| SD11922 | FBgn0028983 | Spn55B | 0.248332778 | 0.10942324 | 0 |
| LP07049 | FBgn0262685 | CG43155 | -0.248249399 | 0.43189074 | 0 |
| GH06740 | FBgn0010398 | Lrr47 | -0.248221493 | 0.09242663 | 0 |
| SD07726 | FBgn0284253 | LeuRS | -0.248202668 | 0.1115611 | 0 |
| LD42037 | FBgn0037664 | CG8420 | 0.248083494 | 0.09576847 | 0 |
| LD02463 | FBgn0259745 | wech | 0.247939795 | 0.2353666 | 0 |
| LD15405 | FBgn0039740 | ZIPIC | 0.247851661 | 0.11640281 | 0 |
| SD02225 | FBgn0032922 | CG9249 | -0.247758127 | 0.08432958 | 0 |
| LD18389 | FBgn0026737 | CG6171 | -0.247710087 | 0.11055415 | 0 |
| SD02424 | FBgn0036124 | CG7839 | 0.247644356 | 0.09698692 | 0 |
| RH54112 | FBgn0037788 | CG3940 | -0.247454271 | 0.25086746 | 0 |

| | | | | | |
|---|---|---|---|---|---|
| RE66907 | FBgn0000636 | Fas3 | 0.247370176 | 0.33567361 | 0 |
| LD02427 | FBgn0032193 | CG5727 | 0.24728232 | 0.20302695 | 0 |
| RE16337 | FBgn0030743 | CG9921 | -0.246993023 | 0.12791277 | 0 |
| RE23670 | FBgn0034307 | CG10914 | -0.246983977 | 0.12008921 | 0 |
| LD23875 | FBgn0037051 | CG10565 | 0.246857262 | 0.13505454 | 0 |
| GH21171 | FBgn0267277 | asRNA:CR45713 | 0.246773983 | 0.13597632 | 0 |
| AT09112 | FBgn0033865 | CG6220 | -0.246764756 | 0.48135302 | 0 |
| GH13185 | FBgn0031455 | Snapin | 0.246753059 | 0.12052351 | 0 |
| GH13185 | FBgn0031454 | CG9960 | 0.246753059 | 0.12052351 | 0 |
| LD05671 | FBgn0036354 | Poc1 | 0.246732243 | 0.11216202 | 0 |
| GH17516 | FBgn0033769 | CG8768 | 0.246427737 | 0.12149771 | 0 |
| LD21669 | FBgn0086130 | Dbp21E2 | 0.246139099 | 0.09715278 | 0 |
| RH43965 | FBgn0021765 | scu | -0.246046121 | 0.14601911 | 0 |
| SD11542 | FBgn0030740 | CG9917 | -0.245896234 | 0.10632134 | 0 |
| RE51204 | FBgn0024361 | Tsp2A | -0.245878077 | 0.1246311 | 0 |
| LD22520 | FBgn0002638 | Rcc1 | 0.245709507 | 0.11813587 | 0 |
| GM02885 | FBgn0261014 | TER94 | -0.245658718 | 0.09542047 | 0 |
| AT02866 | FBgn0263377 | asRNA:CR43429 | -0.245642495 | 0.08661355 | 0 |
| GH04826 | FBgn0051550 | CG31550 | 0.245475593 | 0.13597632 | 0 |
| GH27479 | FBgn0031850 | Tsp | 0.245430759 | 0.41419763 | 0 |
| LD43045 | FBgn0039302 | Nup358 | 0.245257592 | 0.14079552 | 0 |
| LD35279 | FBgn0032169 | CG4709 | 0.245216934 | 0.14248667 | 0 |
| LD46084 | FBgn0035754 | MED4 | 0.245110799 | 0.14646374 | 0 |
| LD36945 | FBgn0032488 | CG16812 | 0.245042041 | 0.10434168 | 0 |
| LD09025 | FBgn0263740 | eIF2gamma | -0.244994834 | 0.10622702 | 0 |
| SD02309 | FBgn0029778 | RhoGAP5A | 0.24497027 | 0.12048033 | 0 |
| LD26912 | FBgn0036570 | IntS9 | 0.244669185 | 0.11102126 | 0 |
| GH01794 | FBgn0263832 | Rcd5 | 0.244539316 | 0.23170768 | 0 |
| SD01656 | FBgn0030486 | Set2 | 0.244423022 | 0.14236342 | 0 |
| RH66493 | FBgn0033692 | wash | -0.244388212 | 0.18668503 | 0 |
| GH12334 | FBgn0052099 | CG32099 | -0.24430348 | 0.11070818 | 0 |
| RE40851 | FBgn0039437 | TwdlL | -0.244260123 | 0.11967427 | 0 |
| RE40851 | FBgn0039436 | TwdlB | -0.244260123 | 0.11967427 | 0 |
| LP04891 | FBgn0036335 | mRpL20 | -0.244068935 | 0.09603242 | 0 |
| RE54080 | FBgn0028516 | ZnT35C | -0.24400683 | 0.0878206 | 0 |
| GM03296 | FBgn0041781 | SCAR | -0.243833564 | 0.25352966 | 0 |
| AT07720 | FBgn0023076 | Clk | -0.243678506 | 0.31861085 | 0 |
| RE27458 | FBgn0001285 | Jon44E | 0.24366773 | 0.1670928 | 0 |
| RH04549 | FBgn0033268 | Obp44a | 0.243603216 | 0.28146115 | 0 |
| GH09573 | FBgn0036986 | CG5282 | -0.243602318 | 0.14961497 | 0 |
| LD36024 | FBgn0030870 | CG6398 | 0.24344058 | 0.14236342 | 0 |
| LD11027 | FBgn0031495 | GABPI | 0.243433223 | 0.11440784 | 0 |
| RE15519 | FBgn0029768 | SPR | -0.243221742 | 0.18965477 | 0 |
| RE34808 | FBgn0039972 | CG17018 | -0.243166951 | 0.09361444 | 0 |
| RE10560 | FBgn0029762 | NAAT1 | -0.243147326 | 0.19005122 | 0 |
| LD38375 | FBgn0036686 | CG7728 | 0.243113305 | 0.0862875 | 0 |
| GM14617 | FBgn0039877 | Mccc1 | -0.243106763 | 0.10316195 | 0 |
| GH06351 | FBgn0016762 | angel | -0.243081251 | 0.16322389 | 0 |

| | | | | | |
|---|---|---|---|---|---|
| AT07490 | FBgn0035198 | CG9133 | 0.243037445 | 0.11638186 | 0 |
| HL05735 | FBgn0261549 | rdgA | -0.242845335 | 0.44538287 | 0 |
| LD29802 | FBgn0024510 | dlt | 0.242837817 | 0.11005477 | 0 |
| LD03471 | FBgn0024509 | Sec13 | 0.242704954 | 0.09232796 | 0 |
| GM03282 | FBgn0031488 | CG17265 | -0.242548405 | 0.27156398 | 0 |
| GM10765 | FBgn0263831 | Gen | 0.2423582 | 0.16321212 | 0 |
| GM05421 | FBgn0030406 | CG1463 | 0.242318061 | 0.10228764 | 0 |
| SD06513 | FBgn0045842 | yuri | 0.242154825 | 0.0969233 | 0 |
| GM10051 | FBgn0023213 | eIF4G1 | -0.241936215 | 0.131545 | 0 |
| GM14141 | FBgn0005355 | Su(fu) | 0.24166984 | 0.16757371 | 0 |
| GH08636 | FBgn0028579 | phtf | 0.24162947 | 0.22734671 | 0 |
| RE61396 | FBgn0085224 | CG34195 | -0.241422584 | 0.08645599 | 0 |
| LD24465 | FBgn0011672 | Mvl | -0.241219843 | 0.09617792 | 0 |
| GH17801 | FBgn0031356 | CG17660 | -0.241132184 | 0.44868134 | 0 |
| LD20207 | FBgn0015391 | glu | 0.241000432 | 0.11070818 | 0 |
| GH27759 | FBgn0035490 | CG1136 | -0.240996676 | 0.08813039 | 0 |
| LD35296 | FBgn0263236 | SP1029 | 0.240867503 | 0.2009534 | 0 |
| LD40136 | FBgn0036289 | CG10657 | -0.240861993 | 0.1115611 | 0 |
| LD29822 | FBgn0031764 | CG9107 | 0.240771901 | 0.12860148 | 0 |
| RH61467 | FBgn0030055 | CG12772 | -0.240744365 | 0.20102435 | 0 |
| LD25963 | FBgn0033735 | CG8525 | 0.240689739 | 0.17104539 | 0 |
| RE72235 | FBgn0024733 | RpL10 | -0.240650326 | 0.19037782 | 0 |
| LP04037 | FBgn0031562 | CG3604 | -0.240593719 | 0.09698692 | 0 |
| RH44652 | FBgn0036900 | CG8765 | -0.240546331 | 0.13551782 | 0 |
| SD07518 | FBgn0031260 | Spp | 0.240460276 | 0.10986902 | 0 |
| RE35228 | FBgn0030274 | Lint-1 | -0.240281702 | 0.1115611 | 0 |
| LD23953 | FBgn0003330 | Sce | 0.240252167 | 0.17863263 | 0 |
| LP02658 | FBgn0052772 | CG32772 | -0.240240362 | 0.11551359 | 0 |
| SD09738 | FBgn0004055 | uzip | 0.24022439 | 0.19136517 | 0 |
| LD32088 | FBgn0038548 | CG17806 | 0.24020848 | 0.13241972 | 0 |
| RH13984 | FBgn0004782 | Ccp84Ab | 0.24017937 | 0.25865893 | 0 |
| LD02884 | FBgn0016930 | Dyrk2 | 0.240124833 | 0.09679205 | 0 |
| LD41932 | FBgn0266019 | rudhira | 0.240044407 | 0.13466737 | 0 |
| GH03482 | FBgn0027596 | Kank | 0.239848297 | 0.17513682 | 0 |
| RH36507 | FBgn0032336 | AstC | 0.239843783 | 0.19232116 | 0 |
| GH11670 | FBgn0000116 | Argk | -0.239795091 | 0.19894242 | 0 |
| LD42301 | FBgn0030932 | Ggt-1 | -0.239711879 | 0.08860951 | 0 |
| LD27182 | FBgn0010340 | 140up | -0.239439751 | 0.15106807 | 0 |
| LD34105 | FBgn0031540 | Pif1 | 0.239434846 | 0.09233185 | 0 |
| LD15650 | FBgn0034878 | pita | 0.239329259 | 0.14751003 | 0 |
| GH22984 | FBgn0033969 | CG10202 | 0.239302874 | 0.25683932 | 0 |
| LD45266 | FBgn0029999 | CG1575 | -0.239242575 | 0.11502125 | 0 |
| GM01008 | FBgn0038583 | CG7183 | 0.239189992 | 0.08769971 | 0 |
| AT02886 | FBgn0037862 | CG4706 | 0.239171056 | 0.24719359 | 0 |
| HL02087 | FBgn0033872 | CG6329 | 0.238960089 | 0.43160337 | 0 |
| GH14278 | FBgn0032864 | CG2493 | 0.238933434 | 0.20769647 | 0 |
| LD24910 | FBgn0031008 | CG8010 | 0.238708631 | 0.09826972 | 0 |
| LD45742 | FBgn0037719 | bocks | -0.238522214 | 0.08974365 | 0 |

| | | | | | |
|---|---|---|---|---|---|
| RE61793 | FBgn0243586 | Tb | -0.238308789 | 0.24946171 | 0 |
| RH17678 | FBgn0002774 | mle | 0.238299993 | 0.09939583 | 0 |
| RE28931 | FBgn0032354 | CG4788 | 0.238290195 | 0.10528283 | 0 |
| LD27873 | FBgn0028471 | Nab2 | 0.238288386 | 0.18577346 | 0 |
| LD21529 | FBgn0027525 | LTV1 | 0.238285365 | 0.14275424 | 0 |
| RH12289 | FBgn0031453 | Bacc | 0.238192343 | 0.13597632 | 0 |
| LD27480 | FBgn0015321 | Ubc4 | -0.238150802 | 0.09204205 | 0 |
| LD06278 | FBgn0013799 | Deaf1 | 0.237990435 | 0.12153789 | 0 |
| RE19693 | FBgn0038549 | CG17802 | -0.237833609 | 0.09044826 | 0 |
| GH22661 | FBgn0028644 | beat-Ic | -0.237802343 | 0.22136191 | 0 |
| AT26185 | FBgn0039889 | Arl4 | -0.237795642 | 0.11070818 | 0 |
| GH05783 | FBgn0042182 | CG18749 | -0.237718649 | 0.22128894 | 0 |
| GH05783 | FBgn0064126 | CG33722 | -0.237718649 | 0.22128894 | 0 |
| RE18459 | FBgn0037526 | ArgRS-m | -0.237669427 | 0.09008791 | 0 |
| GH08048 | FBgn0039856 | CG1774 | -0.237520396 | 0.25159252 | 0 |
| GH11850 | FBgn0013301 | ProtB | -0.237361566 | 0.0973297 | 0 |
| GH11850 | FBgn0013300 | ProtA | -0.237361566 | 0.0973297 | 0 |
| LD47929 | FBgn0038320 | Sra-1 | 0.237207667 | 0.18468714 | 0 |
| AT13424 | FBgn0031626 | CG15631 | 0.23706174 | 0.19843877 | 0 |
| LD17191 | FBgn0263197 | Patronin | 0.236874969 | 0.19562813 | 0 |
| GH14412 | FBgn0001145 | Gs2 | 0.236858301 | 0.53504606 | 0 |
| SD07766 | FBgn0266696 | Svil | 0.236553174 | 0.21322625 | 0 |
| RE05806 | FBgn0034918 | Pym | -0.236518424 | 0.10642008 | 0 |
| LD23595 | FBgn0035372 | CG12093 | -0.236499485 | 0.17404184 | 0 |
| LD44979 | FBgn0039261 | Ythdf | 0.236497056 | 0.14478603 | 0 |
| RE37651 | FBgn0029959 | Rab39 | -0.236496616 | 0.10151555 | 0 |
| RE62222 | FBgn0035049 | Mmp1 | -0.236361534 | 0.35914071 | 0 |
| LD16356 | FBgn0003346 | RanGAP | 0.236327408 | 0.1609697 | 0 |
| RE49992 | FBgn0030570 | CG12540 | -0.235992832 | 0.17133976 | 0 |
| AT08919 | FBgn0036183 | CG6083 | -0.235929706 | 0.17889929 | 0 |
| GH05406 | FBgn0030692 | mRpS30 | 0.235881806 | 0.09008791 | 0 |
| GH01709 | FBgn0039537 | CG5590 | 0.235771977 | 0.16764444 | 0 |
| RE41137 | FBgn0052850 | CG32850 | -0.235748505 | 0.2459181 | 0 |
| LD31805 | FBgn0010397 | LamC | 0.23555076 | 0.09671981 | 0 |
| AT28259 | FBgn0013813 | Dhc98D | 0.235390768 | 0.4655033 | 0 |
| LD18090 | FBgn0000047 | Act88F | 0.235127799 | 0.175776 | 0 |
| LD18090 | FBgn0000043 | Act42A | 0.235127799 | 0.175776 | 0 |
| LD18090 | FBgn0000042 | Act5C | 0.235127799 | 0.175776 | 0 |
| HL02140 | FBgn0028577 | hfp | 0.234993299 | 0.13608475 | 0 |
| LD15094 | FBgn0032136 | Apoltp | 0.234966523 | 0.53118114 | 0 |
| AT26853 | FBgn0034837 | RpL22-like | 0.234859935 | 0.17917133 | 0 |
| LD42420 | FBgn0035151 | CG17129 | 0.234700617 | 0.1037592 | 0 |
| LD36843 | FBgn0043792 | CG30427 | 0.234587819 | 0.22276303 | 0 |
| GH13725 | FBgn0027329 | CCT6 | -0.234541321 | 0.16597708 | 0 |
| SD01285 | FBgn0035085 | CG3770 | 0.234506182 | 0.13418045 | 0 |
| LD24316 | FBgn0026573 | ADD1 | 0.234501828 | 0.14693767 | 0 |
| LD06138 | FBgn0051712 | CG31712 | 0.234400849 | 0.11860764 | 0 |
| SD06668 | FBgn0039215 | CG6695 | 0.234308274 | 0.09075684 | 0 |

| | | | | | |
|---|---|---|---|---|---|
| SD09402 | FBgn0035235 | CG7879 | 0.234287321 | 0.17859077 | 0 |
| RE02355 | FBgn0033341 | MrgBP | 0.234254465 | 0.20459845 | 0 |
| LD02616 | FBgn0030503 | Tango2 | 0.234141098 | 0.11540796 | 0 |
| GH10427 | FBgn0263260 | sel | -0.234054084 | 0.09784391 | 0 |
| LD24213 | FBgn0011289 | TfIIA-L | 0.234041803 | 0.15744362 | 0 |
| GM13228 | FBgn0037926 | Elp1 | 0.233976936 | 0.14301342 | 0 |
| LP05645 | FBgn0264307 | orb2 | -0.233944451 | 0.31903254 | 0 |
| GH07905 | FBgn0032162 | CG4592 | -0.233845005 | 0.29332909 | 0 |
| SD04710 | FBgn0034691 | Synj | -0.233725043 | 0.11355395 | 0 |
| LD33804 | FBgn0031119 | CG1812 | -0.233645202 | 0.10448925 | 0 |
| RE72043 | FBgn0265598 | Bx | -0.233527147 | 0.12947861 | 0 |
| LD14064 | FBgn0035838 | ldbr | 0.233449809 | 0.17793042 | 0 |
| RH62559 | FBgn0034140 | Lst | -0.233375585 | 0.43120083 | 0 |
| AT11810 | FBgn0030566 | betaNACtes4 | 0.233370746 | 0.33869695 | 0 |
| LD05576 | FBgn0032750 | CG10495 | -0.233295711 | 0.26375792 | 0 |
| GH02705 | FBgn0035101 | p130CAS | 0.233106161 | 0.21426982 | 0 |
| GH05823 | FBgn0003435 | sm | 0.233104147 | 0.25628861 | 0 |
| RE10106 | FBgn0036008 | CG3408 | -0.233099968 | 0.11007224 | 0 |
| LD45662 | FBgn0029941 | CG1677 | 0.232937394 | 0.14055868 | 0 |
| GH04745 | FBgn0042179 | CG18869 | 0.232674498 | 0.23534046 | 0 |
| LD08906 | FBgn0038768 | CG4936 | 0.232608445 | 0.1872571 | 0 |
| LD36157 | FBgn0030851 | CG8326 | 0.232581789 | 0.11689839 | 0 |
| GM08857 | FBgn0031003 | CG7889 | -0.232571676 | 0.1241051 | 0 |
| RH40504 | FBgn0030773 | CG9676 | -0.232386766 | 0.11962826 | 0 |
| RE75088 | FBgn0025839 | ND-B14.5A | -0.232372798 | 0.13597632 | 0 |
| RE54049 | FBgn0015801 | Reg-5 | 0.232334056 | 0.31842271 | 0 |
| GH10745 | FBgn0037744 | CG8417 | -0.232149376 | 0.09470904 | 0 |
| RE02696 | FBgn0264907 | CG44098 | -0.231954748 | 0.20639193 | 0 |
| LD25448 | FBgn0036856 | CG9666 | 0.231936074 | 0.11440784 | 0 |
| LD25448 | FBgn0259720 | CG42374 | 0.231936074 | 0.11440784 | 0 |
| LD20635 | FBgn0034109 | CG7747 | 0.231870378 | 0.11690063 | 0 |
| SD24562 | FBgn0004449 | Ten-m | 0.231794443 | 0.20716724 | 0 |
| RH71315 | FBgn0031265 | CG2794 | -0.231756864 | 0.20074155 | 0 |
| AT31405 | FBgn0263995 | cpo | -0.231510969 | 0.23340342 | 0 |
| GH08432 | FBgn0041182 | Tep2 | 0.231416921 | 0.20926237 | 0 |
| GH17661 | FBgn0053056 | CG33056 | -0.231321455 | 0.12227708 | 0 |
| RE22690 | FBgn0264357 | SNF4Agamma | 0.231316472 | 0.12020594 | 0 |
| GH15428 | FBgn0051706 | CG31706 | -0.23131525 | 0.2340838 | 0 |
| LD45766 | FBgn0037794 | CG6254 | 0.231066216 | 0.1366043 | 0 |
| LD08669 | FBgn0283468 | slmb | -0.231044127 | 0.11597957 | 0 |
| LD15586 | FBgn0021979 | l(2)k09913 | 0.230866605 | 0.1321855 | 0 |
| GH02536 | FBgn0027607 | CG8230 | -0.230810287 | 0.20102435 | 0 |
| AT17695 | FBgn0260461 | Rcd7 | -0.230685023 | 0.41427119 | 0 |
| RE11660 | FBgn0016054 | phr6-4 | -0.23065905 | 0.09754135 | 0 |
| RE50383 | FBgn0035157 | CG13894 | -0.230572434 | 0.32008803 | 0 |
| GM13788 | FBgn0001220 | Hsc70-5 | -0.230555454 | 0.11359747 | 0 |
| GH13672 | FBgn0034057 | CG8314 | 0.23028689 | 0.17884847 | 0 |
| GH22460 | FBgn0031689 | Cyp28d1 | -0.23006869 | 0.37459627 | 0 |

| | | | | | |
|---|---|---|---|---|---|
| RE73632 | FBgn0020910 | RpL3 | 0.230063783 | 0.25865893 | 0 |
| RE73632 | FBgn0086056 | snoRNA:Me18S-A425 | 0.230063783 | 0.25865893 | 0 |
| RH04305 | FBgn0001187 | Hex-C | -0.23004274 | 0.12409596 | 0 |
| LD35157 | FBgn0051357 | CG31357 | 0.229940064 | 0.12487471 | 0 |
| LD35157 | FBgn0051118 | RabX4 | 0.229940064 | 0.12487471 | 0 |
| SD10403 | FBgn0034748 | CG17807 | 0.229761512 | 0.1241051 | 0 |
| GH03311 | FBgn0284252 | Letm1 | -0.229542563 | 0.12968005 | 0 |
| LD38333 | FBgn0016756 | Usp47 | 0.229439438 | 0.1115611 | 0 |
| LD05335 | FBgn0025693 | ZnT41F | 0.22940072 | 0.18703013 | 0 |
| SD04735 | FBgn0010280 | Taf4 | 0.229322227 | 0.12048033 | 0 |
| GH23156 | FBgn0037050 | ICA69 | 0.229302122 | 0.13445757 | 0 |
| LD27520 | FBgn0004390 | RasGAP1 | 0.229301131 | 0.15281795 | 0 |
| LD30408 | FBgn0033925 | CG8617 | 0.229137668 | 0.10466062 | 0 |
| RE28166 | FBgn0002878 | mus101 | -0.228537125 | 0.1018806 | 0 |
| GM01975 | FBgn0026479 | Drp1 | -0.228520652 | 0.13058533 | 0 |
| LD10786 | FBgn0001986 | Mtr4 | 0.228477096 | 0.09483457 | 0 |
| LD40833 | FBgn0030054 | Caf1-180 | 0.228470295 | 0.21444299 | 0 |
| GH15958 | FBgn0001624 | dlg1 | 0.228459622 | 0.14548021 | 0 |
| SD02943 | FBgn0032051 | CG13097 | 0.228419239 | 0.11372124 | 0 |
| LD22754 | FBgn0014366 | noi | 0.228187086 | 0.10942324 | 0 |
| LD05118 | FBgn0024689 | fws | 0.228180006 | 0.13460968 | 0 |
| LP02909 | FBgn0011230 | poe | 0.228165872 | 0.17147856 | 0 |
| RE68041 | FBgn0284221 | Sema5c | 0.228133541 | 0.09679205 | 0 |
| LD40589 | FBgn0037633 | CG9839 | 0.228105065 | 0.24417029 | 0 |
| AT23490 | FBgn0039254 | Nmnat | -0.228061584 | 0.22678785 | 0 |
| AT16985 | FBgn0031728 | Hsp60C | -0.227994026 | 0.12943086 | 0 |
| SD04463 | FBgn0260742 | CG12213 | 0.227954982 | 0.10466062 | 0 |
| RE06111 | FBgn0039955 | CG41099 | -0.227907924 | 0.22929842 | 0 |
| RE64430 | FBgn0015625 | CycB3 | 0.227898154 | 0.17434219 | 0 |
| LD32255 | FBgn0001137 | grk | 0.227690075 | 0.11372124 | 0 |
| RE31307 | FBgn0031059 | CG14229 | -0.227553813 | 0.11597957 | 0 |
| LD10479 | FBgn0000221 | brn | 0.227537339 | 0.16360447 | 0 |
| RE13562 | FBgn0030037 | Miga | -0.227516758 | 0.10316195 | 0 |
| GH01786 | FBgn0035714 | CG8549 | -0.227387669 | 0.10144287 | 0 |
| AT13332 | FBgn0011206 | bol | 0.227280488 | 0.11742225 | 0 |
| LD26976 | FBgn0010611 | Hmgs | -0.227058522 | 0.12227708 | 0 |
| GM03260 | FBgn0002906 | Blm | -0.227010672 | 0.09679205 | 0 |
| GH22160 | FBgn0039223 | CG5805 | -0.226949867 | 0.11842639 | 0 |
| RH17388 | FBgn0036294 | CG10654 | -0.226919136 | 0.11842639 | 0 |
| AT12326 | FBgn0036032 | CG16711 | -0.22683018 | 0.0975717 | 0 |
| RE03627 | FBgn0032818 | CG10628 | 0.226823396 | 0.17008461 | 0 |
| RE09129 | FBgn0032435 | Oatp33Eb | -0.226790263 | 0.11210428 | 0 |
| GH12763 | FBgn0032883 | Rhau | -0.226777503 | 0.1567743 | 0 |
| LD24903 | FBgn0037135 | CG7414 | 0.226729323 | 0.19242599 | 0 |
| LD13611 | FBgn0029664 | CG10802 | 0.226722786 | 0.14124168 | 0 |
| GH22734 | FBgn0050015 | CG30015 | 0.226611707 | 0.18253104 | 0 |
| AT12853 | FBgn0031343 | CG18131 | -0.226607388 | 0.35405702 | 0 |
| RH44722 | FBgn0031099 | CG17065 | -0.226579657 | 0.19136517 | 0 |

| | | | | | |
|---|---|---|---|---|---|
| LD23445 | FBgn0037232 | Suv3 | 0.226496641 | 0.12227708 | 0 |
| RE39877 | FBgn0051386 | lncRNA:CR31386 | -0.226457098 | 0.38552174 | 0 |
| RE31492 | FBgn0250785 | vari | 0.22642206 | 0.23291098 | 0 |
| SD05496 | FBgn0015371 | chn | -0.22636195 | 0.32084759 | 0 |
| RE11315 | FBgn0040359 | CG11380 | -0.226359217 | 0.30993577 | 0 |
| RE61939 | FBgn0032444 | CCT4 | -0.226352384 | 0.12213767 | 0 |
| LD15864 | FBgn0004587 | B52 | 0.22633108 | 0.1762235 | 0 |
| LD45081 | FBgn0260397 | Su(var)3-3 | 0.226287055 | 0.10942324 | 0 |
| LD23884 | FBgn0052649 | CG32649 | 0.226204896 | 0.12487471 | 0 |
| GM01879 | FBgn0037093 | Cdk12 | 0.226180979 | 0.11255406 | 0 |
| GM09639 | FBgn0283712 | LIMK1 | 0.226158277 | 0.12506457 | 0 |
| RE15887 | FBgn0039417 | CG6073 | -0.226097622 | 0.13175758 | 0 |
| LD32687 | FBgn0032363 | Dlg5 | 0.225928545 | 0.11210428 | 0 |
| LD43035 | FBgn0037918 | CG6791 | 0.225752776 | 0.15019147 | 0 |
| RH01814 | FBgn0035753 | RpL18 | 0.225740802 | 0.35347303 | 0 |
| RE21150 | FBgn0035851 | MED24 | 0.225738324 | 0.11809235 | 0 |
| GH08833 | FBgn0015269 | Nf1 | 0.225712077 | 0.09781851 | 0 |
| GH19637 | FBgn0050291 | CG30291 | 0.225607962 | 0.13966772 | 0 |
| GH13213 | FBgn0052392 | CG32392 | -0.225512663 | 0.4814305 | 0 |
| RH23514 | FBgn0032322 | CG16743 | -0.225454444 | 0.11502125 | 0 |
| GH21984 | FBgn0086913 | Rab26 | -0.225136899 | 0.18612584 | 0 |
| RH09496 | FBgn0051719 | RluA-1 | 0.225043981 | 0.15489812 | 0 |
| LD30467 | FBgn0037621 | M1BP | 0.224897029 | 0.16549337 | 0 |
| GH17087 | FBgn0029708 | CG3556 | -0.224891114 | 0.35619541 | 0 |
| LD29525 | FBgn0038535 | alt | 0.224811345 | 0.13983076 | 0 |
| SD10038 | FBgn0038578 | MED17 | -0.224806509 | 0.1595985 | 0 |
| LD08555 | FBgn0040078 | pont | 0.224797743 | 0.14625566 | 0 |
| GH13170 | FBgn0015277 | Pi3K59F | -0.224790857 | 0.39339733 | 0 |
| GH13170 | FBgn0050184 | CG30184 | -0.224790857 | 0.39339733 | 0 |
| SD09792 | FBgn0039209 | REPTOR | -0.22472533 | 0.20555225 | 0 |
| LD14127 | FBgn0038601 | CG18600 | 0.224693298 | 0.13001995 | 0 |
| LD04007 | FBgn0039169 | Spps | -0.224543855 | 0.17861543 | 0 |
| AT29337 | FBgn0042126 | CG18788 | -0.224432187 | 0.10397702 | 0 |
| RE01153 | FBgn0052453 | CG32453 | -0.22435663 | 0.09880238 | 0 |
| RE01153 | FBgn0037177 | CG14454 | -0.22435663 | 0.09880238 | 0 |
| AT08432 | FBgn0250842 | CG17575 | 0.224216641 | 0.45210484 | 0 |
| RE62913 | FBgn0034419 | CG15111 | 0.224194903 | 0.11242653 | 0 |
| SD06851 | FBgn0034049 | bdg | 0.224174904 | 0.15024149 | 0 |
| LD25286 | FBgn0002783 | mor | 0.223954022 | 0.11813587 | 0 |
| GH26553 | FBgn0044452 | Atg2 | 0.223873209 | 0.10434168 | 0 |
| GH04846 | FBgn0037647 | RagA-B | -0.22375787 | 0.20048325 | 0 |
| GM13220 | FBgn0020392 | Nmt | -0.223732443 | 0.09851079 | 0 |
| GM02612 | FBgn0035254 | CG7974 | 0.223703312 | 0.11351359 | 0 |
| LD21622 | FBgn0003392 | shi | 0.223630713 | 0.09890059 | 0 |
| RE52015 | FBgn0034999 | CG3394 | -0.223520333 | 0.11440784 | 0 |
| GH15359 | FBgn0040273 | Spt5 | 0.223457431 | 0.1370972 | 0 |
| RE20036 | FBgn0033750 | CG13151 | 0.223411821 | 0.13368437 | 0 |
| LP11136 | FBgn0266708 | Cep89 | 0.223410622 | 0.12227708 | 0 |

| | | | | | |
|---|---|---|---|---|---|
| LD35784 | FBgn0000996 | dup | 0.223377147 | 0.1366984 | 0 |
| RE44178 | FBgn0027343 | fz3 | -0.22325469 | 0.18274693 | 0 |
| SD02978 | FBgn0010591 | Sply | -0.223254004 | 0.15906113 | 0 |
| LD18949 | FBgn0031606 | CG15439 | 0.223140089 | 0.14152822 | 0 |
| GH12163 | FBgn0020521 | pio | 0.222894527 | 0.16213341 | 0 |
| LD26135 | FBgn0043796 | CG12219 | 0.222878635 | 0.11715236 | 0 |
| GH07340 | FBgn0030990 | CG7556 | -0.222756778 | 0.11695164 | 0 |
| GH02435 | FBgn0027609 | morgue | -0.22267033 | 0.14709825 | 0 |
| RH49423 | FBgn0039358 | CG5028 | -0.222583595 | 0.11597957 | 0 |
| RE01860 | FBgn0020762 | Atet | 0.222537179 | 0.41123404 | 0 |
| RH54711 | FBgn0039159 | mRpS24 | -0.222483147 | 0.37479604 | 0 |
| SD06560 | FBgn0025628 | CG4199 | -0.222402105 | 0.11428562 | 0 |
| GH04604 | FBgn0015031 | cype | 0.222226846 | 0.17147856 | 0 |
| RE15336 | FBgn0011676 | Nos | -0.222198719 | 0.18472354 | 0 |
| RE64884 | FBgn0035886 | Jon66Ci | -0.221860568 | 0.17434219 | 0 |
| RE64884 | FBgn0035887 | Jon66Cii | -0.221860568 | 0.17434219 | 0 |
| RE60807 | FBgn0024996 | eIF2Bbeta | -0.22184321 | 0.21998928 | 0 |
| RE19835 | FBgn0030855 | CG5800 | 0.221831744 | 0.11359747 | 0 |
| GH16531 | FBgn0051451 | lncRNA:CR31451 | 0.221767862 | 0.29178686 | 0 |
| RE21703 | FBgn0058198 | CG40198 | -0.221678016 | 0.21464103 | 0 |
| GH08684 | FBgn0015622 | Cnx99A | -0.221673288 | 0.15034977 | 0 |
| GH23825 | FBgn0024985 | CG11448 | -0.221667878 | 0.11962826 | 0 |
| LD43674 | FBgn0039626 | Slu7 | 0.221587518 | 0.15310733 | 0 |
| LD15947 | FBgn0010926 | l(3)07882 | 0.221437087 | 0.15565223 | 0 |
| LD47455 | FBgn0036314 | Sf3a2 | 0.22140023 | 0.12841406 | 0 |
| GH26090 | FBgn0261564 | Reep1 | 0.221326404 | 0.11014912 | 0 |
| LD10220 | FBgn0035181 | CG9205 | 0.221308391 | 0.16394152 | 0 |
| RE53253 | FBgn0020445 | E23 | -0.221247621 | 0.11951953 | 0 |
| LD38886 | FBgn0011785 | BRWD3 | 0.221058201 | 0.13597632 | 0 |
| AT28382 | FBgn0032338 | CG16854 | 0.221009037 | 0.49200485 | 0 |
| LD22183 | FBgn0041188 | Atx2 | -0.220995817 | 0.11962826 | 0 |
| RE14657 | FBgn0004513 | Mdr65 | 0.220937995 | 0.20102435 | 0 |
| LD43392 | FBgn0261286 | Mat89Ba | 0.220928459 | 0.12048033 | 0 |
| LD04103 | FBgn0266521 | stai | -0.220926337 | 0.20926155 | 0 |
| RE13893 | FBgn0037430 | Osi20 | -0.220895673 | 0.35327229 | 0 |
| LP03067 | FBgn0036416 | CG7924 | 0.220888404 | 0.27904632 | 0 |
| RE22259 | FBgn0030001 | cyr | -0.22081163 | 0.18165 | 0 |
| AT05565 | FBgn0003140 | PpY-55A | 0.220779748 | 0.21862256 | 0 |
| RE68603 | FBgn0024993 | CG2662 | -0.220775849 | 0.1115611 | 0 |
| RE54525 | FBgn0037012 | Rcd2 | -0.220766931 | 0.33551415 | 0 |
| SD15546 | FBgn0000158 | bam | -0.220726998 | 0.15613854 | 0 |
| LD13869 | FBgn0040493 | grsm | -0.220697879 | 0.13767226 | 0 |
| LD11008 | FBgn0000547 | ed | 0.220559891 | 0.1246311 | 0 |
| GH28553 | FBgn0039282 | Bili | -0.220354618 | 0.37005256 | 0 |
| AT03755 | FBgn0030797 | CG13004 | -0.220254763 | 0.1060808 | 0 |
| RE71565 | FBgn0034073 | CG8414 | -0.220200546 | 0.10857776 | 0 |
| LP02993 | FBgn0026787 | Nhe1 | -0.220077065 | 0.10623561 | 0 |
| LD24894 | FBgn0037028 | CG3618 | 0.220051607 | 0.20282151 | 0 |

| | | | | | |
|---|---|---|---|---|---|
| RH48586 | FBgn0031312 | Tango14 | -0.219902626 | 0.12559593 | 0 |
| GH28601 | FBgn0029814 | CG15765 | -0.219717732 | 0.30584465 | 0 |
| LD45751 | FBgn0000520 | dwg | 0.219660162 | 0.1050392 | 0 |
| GH07914 | FBgn0053307 | CG33307 | 0.219568086 | 0.21487795 | 0 |
| SD19774 | FBgn0040268 | Top3alpha | -0.219450754 | 0.22128894 | 0 |
| LD30155 | FBgn0263594 | lost | -0.219300211 | 0.11070818 | 0 |
| GH04674 | FBgn0029990 | CG2233 | 0.21922946 | 0.20687266 | 0 |
| RE22741 | FBgn0031959 | spz3 | -0.219219001 | 0.25359687 | 0 |
| HL01112 | FBgn0039726 | eIF2Balpha | -0.219070163 | 0.13767226 | 0 |
| RH74717 | FBgn0039864 | CG11550 | 0.219051153 | 0.24611788 | 0 |
| GH12419 | FBgn0064123 | stg1 | -0.218995134 | 0.26066785 | 0 |
| GH12419 | FBgn0031159 | CG11566 | -0.218995134 | 0.26066785 | 0 |
| HL05730 | FBgn0038515 | CG5823 | -0.21887885 | 0.11440784 | 0 |
| LD40873 | FBgn0015288 | RpL22 | 0.218837826 | 0.16096423 | 0 |
| LD33046 | FBgn0020407 | asun | 0.218825332 | 0.10229429 | 0 |
| GM02242 | FBgn0029897 | RpL17 | 0.218749174 | 0.29122914 | 0 |
| RE60512 | FBgn0261434 | hkb | 0.218719974 | 0.27004964 | 0 |
| RE39339 | FBgn0029706 | CG3626 | -0.218702099 | 0.11638186 | 0 |
| RE67111 | FBgn0028400 | Syt4 | -0.218623953 | 0.30946795 | 0 |
| AT31826 | FBgn0039773 | CG2224 | 0.218590404 | 0.11359747 | 0 |
| AT01386 | FBgn0053340 | CG33340 | -0.218458963 | 0.29858549 | 0 |
| AT01386 | FBgn0030376 | CG2750 | -0.218458963 | 0.29858549 | 0 |
| RE15880 | FBgn0031650 | CG14044 | -0.218363482 | 0.13005317 | 0 |
| LP03286 | FBgn0034697 | GM130 | 0.218257979 | 0.12487471 | 0 |
| AT08344 | FBgn0021874 | Nle | -0.218246859 | 0.19876252 | 0 |
| LD41005 | FBgn0034631 | TAF1C-like | 0.218242308 | 0.11700678 | 0 |
| CK00436 | FBgn0020414 | Idgf3 | -0.21823483 | 0.18942072 | 0 |
| LP10814 | FBgn0052690 | lncRNA:CR32690 | 0.218044718 | 0.29671934 | 0 |
| GH22974 | FBgn0034959 | CG11388 | 0.218027151 | 0.22141553 | 0 |
| GH03119 | FBgn0283478 | und | -0.217952926 | 0.1246311 | 0 |
| RE56074 | FBgn0031732 | CG11149 | -0.217807844 | 0.37533608 | 0 |
| LD06553 | FBgn0031418 | CG3609 | 0.21775826 | 0.15565223 | 0 |
| SD20094 | FBgn0051158 | Efa6 | -0.217711261 | 0.30224264 | 0 |
| SD09769 | FBgn0052177 | Ndfip | -0.217693092 | 0.12680002 | 0 |
| HL06663 | FBgn0085350 | inaF-C | 0.217562853 | 0.21464103 | 0 |
| HL06663 | FBgn0085351 | inaF-A | 0.217562853 | 0.21464103 | 0 |
| HL06663 | FBgn0259918 | inaF-B | 0.217562853 | 0.21464103 | 0 |
| HL06663 | FBgn0260812 | inaF-D | 0.217562853 | 0.21464103 | 0 |
| RH01238 | FBgn0032810 | CG13077 | -0.217467552 | 0.35798574 | 0 |
| GH25089 | FBgn0013342 | nSyb | 0.21745252 | 0.25086746 | 0 |
| LP02637 | FBgn0038750 | CG4465 | -0.217436795 | 0.42092985 | 0 |
| RE18409 | FBgn0032447 | PICK1 | -0.217299056 | 0.26088254 | 0 |
| LD21719 | FBgn0021847 | l(2)k14710 | -0.217218528 | 0.10524592 | 0 |
| RE24457 | FBgn0028342 | ATPsyndelta | 0.217121999 | 0.17593513 | 0 |
| RE32803 | FBgn0250839 | CG2016 | 0.21707168 | 0.55926524 | 0 |
| GH26345 | FBgn0032261 | CG6094 | -0.217022914 | 0.11428562 | 0 |
| SD07983 | FBgn0034346 | PIG-O | 0.216941632 | 0.21418818 | 0 |
| AT16851 | FBgn0039152 | Root | 0.216919363 | 0.11962826 | 0 |

| | | | | | |
|---|---|---|---|---|---|
| RH63361 | FBgn0040606 | CG6503 | 0.216870318 | 0.10942324 | 0 |
| RE67616 | FBgn0032744 | Ttc19 | 0.216832593 | 0.13966772 | 0 |
| SD21944 | FBgn0034184 | CG9646 | 0.216775421 | 0.11070818 | 0 |
| GM01253 | FBgn0051320 | HEATR2 | 0.216691221 | 0.1246311 | 0 |
| LP02042 | FBgn0030244 | CG2157 | 0.216623238 | 0.25336796 | 0 |
| LP07079 | FBgn0033158 | CG12164 | -0.216566384 | 0.2071339 | 0 |
| GH01001 | FBgn0033484 | CG2269 | 0.216496359 | 0.21426982 | 0 |
| GH09271 | FBgn0261794 | kcc | 0.216464671 | 0.1545559 | 0 |
| GH21853 | FBgn0261822 | Bsg | -0.21644993 | 0.17806104 | 0 |
| LD23830 | FBgn0028473 | Non1 | 0.216360466 | 0.18537449 | 0 |
| RE43488 | FBgn0285952 | eEF5 | -0.216331337 | 0.18274693 | 0 |
| LP06602 | FBgn0037993 | dpr15 | -0.216320495 | 0.4690427 | 0 |
| RH42110 | FBgn0033169 | CG11123 | 0.216115277 | 0.16822308 | 0 |
| GH09112 | FBgn0034517 | Cpr57A | 0.216069923 | 0.29443051 | 0 |
| GH09112 | FBgn0043536 | Obp57d | 0.216069923 | 0.29443051 | 0 |
| LD06095 | FBgn0034179 | CG6805 | 0.216056382 | 0.10603766 | 0 |
| GH10631 | FBgn0034145 | CG5065 | 0.216020872 | 0.27774828 | 0 |
| RE55916 | FBgn0038619 | CG7685 | -0.215946633 | 0.17877971 | 0 |
| LD26915 | FBgn0000546 | EcR | 0.215895358 | 0.10973434 | 0 |
| RE29502 | FBgn0030941 | wgn | 0.215844089 | 0.24657701 | 0 |
| LD35228 | FBgn0034537 | DMAP1 | 0.215843131 | 0.13597632 | 0 |
| GH20931 | FBgn0029891 | Pink1 | 0.215819585 | 0.14704897 | 0 |
| LD12309 | FBgn0030850 | stas | -0.215753526 | 0.12852347 | 0 |
| LD21537 | FBgn0027524 | CG3909 | 0.215742375 | 0.1241051 | 0 |
| SD19962 | FBgn0260990 | yata | -0.215736587 | 0.2476695 | 0 |
| RH19830 | FBgn0037410 | Osi2 | -0.2157352 | 0.21538541 | 0 |
| GH19483 | FBgn0031093 | CG9581 | 0.215728477 | 0.20639193 | 0 |
| GH09073 | FBgn0031773 | Fbw5 | 0.215549574 | 0.12014527 | 0 |
| LD05535 | FBgn0034380 | Vps51 | 0.215516804 | 0.13541238 | 0 |
| GH25379 | FBgn0040308 | Jafrac2 | -0.215494154 | 0.13252957 | 0 |
| LD38592 | FBgn0037255 | Fip1 | 0.215347027 | 0.11597957 | 0 |
| RE54004 | FBgn0047095 | CG33939 | -0.215322166 | 0.12360916 | 0 |
| GH14720 | FBgn0030731 | Mfe2 | 0.215301134 | 0.17775002 | 0 |
| LD40412 | FBgn0039680 | Cap-D2 | 0.21527473 | 0.13203002 | 0 |
| GH19076 | FBgn0263593 | Lpin | -0.215133533 | 0.11035759 | 0 |
| RE24065 | FBgn0001321 | knk | 0.214917762 | 0.42145924 | 0 |
| GH01164 | FBgn0037763 | CG16904 | -0.214892746 | 0.11055415 | 0 |
| GH09832 | FBgn0034745 | CG4329 | -0.214852523 | 0.11027617 | 0 |
| GM13256 | FBgn0033055 | Tbce | -0.214799121 | 0.36967389 | 0 |
| AT18965 | FBgn0039104 | CG10252 | -0.214727128 | 0.12692358 | 0 |
| LD37466 | FBgn0033352 | PAN2 | 0.214706132 | 0.11951953 | 0 |
| RH07711 | FBgn0040396 | CG3939 | -0.214658499 | 0.11055415 | 0 |
| GH12111 | FBgn0020306 | dom | 0.214651031 | 0.1366984 | 0 |
| LD12265 | FBgn0031457 | CG3077 | 0.214633346 | 0.1194013 | 0 |
| GH03927 | FBgn0031872 | ihog | -0.214566248 | 0.1115611 | 0 |
| AT17882 | FBgn0031384 | CG4238 | -0.214415839 | 0.1246311 | 0 |
| GH04071 | FBgn0037778 | mtTFB2 | 0.214384638 | 0.15489812 | 0 |
| SD03524 | FBgn0283503 | Neurl4 | 0.214273766 | 0.1749869 | 0 |

| | | | | | |
|---|---|---|---|---|---|
| AT23571 | FBgn0063485 | Lasp | -0.214270361 | 0.23695841 | 0 |
| LD14717 | FBgn0039449 | CG6425 | 0.214258297 | 0.12052351 | 0 |
| LD10532 | FBgn0000575 | emc | 0.214220096 | 0.24754354 | 0 |
| LD02541 | FBgn0262614 | pyd | 0.214167831 | 0.11351359 | 0 |
| RH24988 | FBgn0038645 | CG7714 | -0.214143503 | 0.12884403 | 0 |
| RE52411 | FBgn0045866 | bai | -0.214066035 | 0.14883745 | 0 |
| LD25827 | FBgn0038948 | PSR | 0.214017521 | 0.19745672 | 0 |
| LP07209 | FBgn0039860 | CG1792 | -0.2138852 | 0.17593513 | 0 |
| LD29349 | FBgn0027660 | blot | -0.213842724 | 0.10942324 | 0 |
| LD41491 | FBgn0036484 | Pex3 | 0.213796609 | 0.12327888 | 0 |
| RH12258 | FBgn0035676 | ssp6 | -0.213790752 | 0.34806685 | 0 |
| GH26082 | FBgn0033261 | udd | 0.213732379 | 0.11638186 | 0 |
| AT25123 | FBgn0034825 | Gpdh2 | 0.213730872 | 0.20247456 | 0 |
| LP05112 | FBgn0035488 | CG11593 | -0.213724627 | 0.12016929 | 0 |
| RE33713 | FBgn0261553 | CG42671 | -0.213634035 | 0.20556671 | 0 |
| RE74472 | FBgn0086768 | Pcmt | 0.213603895 | 0.12834476 | 0 |
| GM13041 | FBgn0038275 | CG3817 | 0.21358186 | 0.11210428 | 0 |
| LD19584 | FBgn0000463 | Dl | 0.213572326 | 0.12227708 | 0 |
| RE06926 | FBgn0043002 | Chrac-14 | 0.213499786 | 0.16322389 | 0 |
| LD21931 | FBgn0038927 | CG6015 | 0.213438198 | 0.12481361 | 0 |
| LD21931 | FBgn0266538 | CG45099 | 0.213438198 | 0.12481361 | 0 |
| LD28592 | FBgn0037780 | ohgt | 0.213364347 | 0.12487471 | 0 |
| LD03819 | FBgn0039881 | CG1971 | 0.213342345 | 0.12065005 | 0 |
| RE60671 | FBgn0024998 | CG2685 | 0.213337004 | 0.19904473 | 0 |
| SD10018 | FBgn0015040 | Cyp9c1 | 0.213228334 | 0.19110445 | 0 |
| GM13868 | FBgn0034804 | CG3831 | -0.213218453 | 0.2088032 | 0 |
| RE11111 | FBgn0036843 | Sfxn2 | -0.213139687 | 0.11689839 | 0 |
| LD10773 | FBgn0032882 | Ns4 | 0.213135327 | 0.12680002 | 0 |
| RE19005 | FBgn0036180 | Duba | -0.213076524 | 0.11860764 | 0 |
| RE70378 | FBgn0262714 | Sap130 | 0.213069569 | 0.10804827 | 0 |
| LD44987 | FBgn0030421 | CG3812 | -0.213044666 | 0.13664789 | 0 |
| LD24704 | FBgn0040394 | CG16903 | 0.21294176 | 0.12787314 | 0 |
| RE50774 | FBgn0039941 | CG17167 | 0.212935654 | 0.13703221 | 0 |
| GH09390 | FBgn0035035 | CG3570 | -0.212927409 | 0.11099012 | 0 |
| LD17738 | FBgn0000416 | Sap-r | 0.212889084 | 0.13175758 | 0 |
| RE73017 | FBgn0052625 | CG32625 | 0.212774113 | 0.17449442 | 0 |
| RE01946 | FBgn0034440 | CG10073 | -0.212764224 | 0.19264462 | 0 |
| RE29349 | FBgn0037843 | CG4511 | -0.212700337 | 0.20926237 | 0 |
| GH13729 | FBgn0261673 | nemy | 0.212682036 | 0.26375792 | 0 |
| LD23906 | FBgn0034182 | SmydA-7 | 0.21265913 | 0.23204888 | 0 |
| LP10445 | FBgn0029937 | CG8300 | -0.212597487 | 0.21296194 | 0 |
| GM14838 | FBgn0021944 | Coprox | -0.212582439 | 0.10942324 | 0 |
| GH21874 | FBgn0085433 | CG34404 | 0.212580561 | 0.20723253 | 0 |
| LP01187 | FBgn0050104 | NT5E-2 | 0.212433657 | 0.12372805 | 0 |
| RE70209 | FBgn0066304 | Rpp20 | -0.21235132 | 0.1683216 | 0 |
| RE70209 | FBgn0066303 | CG33932 | -0.21235132 | 0.1683216 | 0 |
| GH02340 | FBgn0028504 | CG12182 | 0.21231414 | 0.13738364 | 0 |
| RE10231 | FBgn0032170 | CG4658 | -0.212313429 | 0.1115611 | 0 |

| | | | | | |
|---|---|---|---|---|---|
| GH24095 | FBgn0046225 | CG17230 | -0.212304137 | 0.14248667 | 0 |
| RE33093 | FBgn0034379 | CG15073 | -0.212235872 | 0.14938916 | 0 |
| RE14441 | FBgn0000045 | Act79B | 0.212212025 | 0.25519816 | 0 |
| RE14441 | FBgn0000046 | Act87E | 0.212212025 | 0.25519816 | 0 |
| RE14441 | FBgn0000044 | Act57B | 0.212212025 | 0.25519816 | 0 |
| LD32520 | FBgn0262737 | mub | -0.212204403 | 0.17697967 | 0 |
| LD23669 | FBgn0017558 | Pdk | 0.211998114 | 0.21464103 | 0 |
| RH04764 | FBgn0053080 | CG33080 | -0.211969748 | 0.17859077 | 0 |
| RE67794 | FBgn0030081 | CG7246 | 0.211886393 | 0.13642205 | 0 |
| LP07175 | FBgn0030574 | CG9413 | 0.211884818 | 0.2091573 | 0 |
| GH01984 | FBgn0022774 | Oat | 0.211853782 | 0.24657701 | 0 |
| GM06264 | FBgn0000307 | chif | 0.21184611 | 0.13906458 | 0 |
| RE33866 | FBgn0040660 | CG13551 | -0.211799398 | 0.45468609 | 0 |
| LD37855 | FBgn0020633 | Mcm7 | 0.211798664 | 0.1115611 | 0 |
| LD22446 | FBgn0011232 | scat | -0.211795428 | 0.13597632 | 0 |
| SD23876 | FBgn0264707 | RhoGEF3 | 0.211746178 | 0.42856874 | 0 |
| RE50926 | FBgn0033761 | CG8778 | -0.211581012 | 0.14248667 | 0 |
| RH71862 | FBgn0003885 | alphaTub84D | 0.211221569 | 0.18075619 | 0 |
| RH71862 | FBgn0003886 | alphaTub85E | 0.211221569 | 0.18075619 | 0 |
| LD26007 | FBgn0040089 | meso18E | 0.211191186 | 0.26820425 | 0 |
| RE55124 | FBgn0000533 | ea | 0.211176155 | 0.19399445 | 0 |
| LD31680 | FBgn0000139 | ash2 | 0.211137622 | 0.11071414 | 0 |
| GH26789 | FBgn0036663 | CG9674 | -0.210996543 | 0.1595985 | 0 |
| RE32663 | FBgn0000337 | cn | 0.210991781 | 0.13003152 | 0 |
| LD33759 | FBgn0031050 | Arp10 | 0.210970748 | 0.131545 | 0 |
| GH02431 | FBgn0027610 | Dic1 | 0.210854284 | 0.14164577 | 0 |
| AT10182 | FBgn0085390 | Dgk | -0.210782391 | 0.49334127 | 0 |
| LD44205 | FBgn0034051 | Mlf | 0.210575971 | 0.14614473 | 0 |
| RH44874 | FBgn0031397 | CG15385 | -0.210521385 | 0.20920946 | 0 |
| GH07317 | FBgn0050295 | Ipk1 | 0.210421187 | 0.12008921 | 0 |
| RE18651 | FBgn0034853 | Ice1 | -0.210416528 | 0.16052533 | 0 |
| RH39533 | FBgn0010387 | Dbi | -0.210367312 | 0.17341214 | 0 |
| GH15485 | FBgn0030412 | Tomosyn | -0.210298705 | 0.15957938 | 0 |
| RE05480 | FBgn0031937 | CG13795 | 0.210122175 | 0.26520084 | 0 |
| LD24749 | FBgn0015396 | jumu | -0.210082808 | 0.13884665 | 0 |
| LP10960 | FBgn0020508 | Ag5r2 | 0.209798972 | 0.31254573 | 0 |
| LD45549 | FBgn0267791 | HnRNP-K | 0.209679777 | 0.11099012 | 0 |
| GH20240 | FBgn0264598 | PsGEF | -0.209597914 | 0.25821219 | 0 |
| GH24926 | FBgn0259707 | CG42361 | 0.209519358 | 0.25612477 | 0 |
| RE31064 | FBgn0039064 | CG4467 | 0.209513552 | 0.52262648 | 0 |
| RH51833 | FBgn0039495 | CG5909 | -0.20944448 | 0.22147922 | 0 |
| AT10584 | FBgn0260750 | Mulk | -0.209350329 | 0.11428562 | 0 |
| HL01254 | FBgn0036909 | CG15881 | -0.209210075 | 0.15810151 | 0 |
| GH14088 | FBgn0019929 | Ser7 | -0.209206889 | 0.14164577 | 0 |
| RE59131 | FBgn0285949 | RpL31 | -0.209205453 | 0.19822213 | 0 |
| LD09343 | FBgn0037838 | TTLL15 | 0.209175348 | 0.14033577 | 0 |
| GH16763 | FBgn0046793 | CG32236 | -0.209159297 | 0.28228402 | 0 |
| RH09503 | FBgn0031760 | Tsp26A | 0.208987221 | 0.12320472 | 0 |

| | | | | | |
|---|---|---|---|---|---|
| RE66512 | FBgn0267336 | Glut4EF | 0.208910001 | 0.26665758 | 0 |
| AT24727 | FBgn0033061 | SmydA-5 | 0.208612037 | 0.19101365 | 0 |
| RH09469 | FBgn0025640 | CG13369 | -0.20844903 | 0.14133622 | 0 |
| RE19958 | FBgn0030992 | CG33253 | -0.208409036 | 0.33399727 | 0 |
| AT27450 | FBgn0033770 | wuc | 0.20838569 | 0.38251295 | 0 |
| GM09104 | FBgn0040290 | RecQ4 | -0.208371764 | 0.25250442 | 0 |
| GH10244 | FBgn0038662 | Mpc1 | -0.208263796 | 0.1241051 | 0 |
| GH18329 | FBgn0031401 | papi | 0.2082459 | 0.11622472 | 0 |
| AT01633 | FBgn0036498 | CG13455 | -0.208212661 | 0.18212145 | 0 |
| LP07312 | FBgn0039114 | Lsd-1 | 0.208184751 | 0.26673415 | 0 |
| LP07312 | FBgn0004117 | Tm2 | 0.208184751 | 0.26673415 | 0 |
| RH09844 | FBgn0086915 | Mst77F | 0.208067161 | 0.35994702 | 0 |
| RH09844 | FBgn0267490 | Mst77Y-4 | 0.208067161 | 0.35994702 | 0 |
| RH09844 | FBgn0267411 | CG45764 | 0.208067161 | 0.35994702 | 0 |
| RH09844 | FBgn0267412 | CG45765 | 0.208067161 | 0.35994702 | 0 |
| RH09844 | FBgn0267427 | CR45780 | 0.208067161 | 0.35994702 | 0 |
| LD08622 | FBgn0013718 | nuf | 0.208043298 | 0.11638186 | 0 |
| AT25481 | FBgn0051956 | Pgant4 | 0.208007952 | 0.23530121 | 0 |
| LD41311 | FBgn0030141 | Gga | 0.207929243 | 0.16546644 | 0 |
| LP03404 | FBgn0030522 | CG11103 | 0.207886581 | 0.13209767 | 0 |
| RE45482 | FBgn0014930 | CG2846 | -0.207850712 | 0.23926556 | 0 |
| LD09009 | FBgn0027889 | ball | 0.207823359 | 0.17646871 | 0 |
| SD25575 | FBgn0024987 | ssx | 0.207763819 | 0.186503 | 0 |
| LD23072 | FBgn0039861 | pasha | 0.207720543 | 0.13597632 | 0 |
| RH59211 | FBgn0016120 | ATPsynD | 0.207674841 | 0.2946539 | 0 |
| GH07529 | FBgn0010651 | MFS14 | 0.207612483 | 0.32172684 | 0 |
| LD42146 | FBgn0085436 | Not1 | 0.207578494 | 0.11742225 | 0 |
| RE17389 | FBgn0031740 | CG7239 | -0.207482008 | 0.2088032 | 0 |
| AT19440 | FBgn0033644 | Tret1-2 | 0.207442173 | 0.31596834 | 0 |
| GH08934 | FBgn0004861 | ph-p | -0.207430357 | 0.31253644 | 0 |
| LD29184 | FBgn0038300 | Mau2 | 0.207150562 | 0.21487795 | 0 |
| RE55427 | FBgn0029858 | mldr | 0.207147693 | 0.18784701 | 0 |
| LD26292 | FBgn0031298 | Atg4a | 0.207124097 | 0.13241972 | 0 |
| LD01519 | FBgn0034455 | CG11007 | -0.206977654 | 0.13070185 | 0 |
| GH14622 | FBgn0030277 | CG1394 | -0.206803563 | 0.19142717 | 0 |
| SD03340 | FBgn0029955 | CG15478 | -0.206793422 | 0.28758766 | 0 |
| LD45317 | FBgn0038860 | Ice2 | 0.206779122 | 0.131545 | 0 |
| AT23067 | FBgn0038473 | Ns1 | 0.206747999 | 0.11597957 | 0 |
| LD23187 | FBgn0032388 | CG6686 | 0.206718629 | 0.12320472 | 0 |
| RE25737 | FBgn0010602 | lwr | 0.206618758 | 0.1495244 | 0 |
| LD43558 | FBgn0003277 | RpII215 | 0.206585531 | 0.11359747 | 0 |
| RE28658 | FBgn0040842 | CG15212 | -0.20652034 | 0.26140837 | 0 |
| RE15573 | FBgn0034177 | AsnRS-m | -0.206445419 | 0.45923276 | 0 |
| GH02984 | FBgn0038641 | CG7708 | -0.206438075 | 0.34865874 | 0 |
| GH02984 | FBgn0284234 | CG46318 | -0.206438075 | 0.34865874 | 0 |
| LD15481 | FBgn0021995 | Rs1 | 0.20639849 | 0.1246311 | 0 |
| LD32037 | FBgn0264075 | tgo | 0.206350944 | 0.14005329 | 0 |
| GM14452 | FBgn0032833 | COX4 | 0.206343984 | 0.26339056 | 0 |

| | | | | | |
|---|---|---|---|---|---|
| LD08328 | FBgn0037656 | CG11986 | 0.20630693 | 0.14294744 | 0 |
| LD18706 | FBgn0013720 | pck | 0.206270009 | 0.131545 | 0 |
| RE13780 | FBgn0038830 | CG17272 | -0.206246782 | 0.1966553 | 0 |
| RH63657 | FBgn0038330 | CG14868 | 0.206180163 | 0.12685445 | 0 |
| LD06965 | FBgn0034030 | CG8192 | -0.206143451 | 0.18702248 | 0 |
| GH05108 | FBgn0283477 | SF2 | 0.206105568 | 0.17512775 | 0 |
| SD09890 | FBgn0038257 | smp-30 | -0.20606926 | 0.43160337 | 0 |
| LD43003 | FBgn0030301 | HP5 | 0.206059825 | 0.15957938 | 0 |
| RE12123 | FBgn0003884 | alphaTub84B | 0.206025423 | 0.31903254 | 0 |
| LD44515 | FBgn0036053 | iPLA2-VIA | -0.205934915 | 0.11597957 | 0 |
| LD23843 | FBgn0039141 | spas | -0.205833346 | 0.13695182 | 0 |
| SD07027 | FBgn0031220 | CG4822 | 0.205805415 | 0.49027635 | 0 |
| GH01073 | FBgn0027617 | CG5808 | 0.20573684 | 0.12693221 | 0 |
| RE05022 | FBgn0014026 | RpL7A | 0.205718598 | 0.16192191 | 0 |
| GH24886 | FBgn0004369 | Ptp99A | -0.205710317 | 0.24315669 | 0 |
| GH22749 | FBgn0036483 | CG12316 | -0.205638136 | 0.17995926 | 0 |
| GM01152 | FBgn0029925 | CG3040 | -0.205629562 | 0.13832386 | 0 |
| RH63918 | FBgn0050033 | CG30033 | -0.205605065 | 0.15810151 | 0 |
| LD08952 | FBgn0038584 | mTerf5 | -0.205589711 | 0.15288834 | 0 |
| RH17284 | FBgn0262139 | trh | -0.205355694 | 0.17635197 | 0 |
| RH25118 | FBgn0025682 | scf | 0.205336326 | 0.22734671 | 0 |
| LD03455 | FBgn0000017 | Abl | 0.205335156 | 0.15810151 | 0 |
| LD29918 | FBgn0040337 | CG3021 | 0.205246009 | 0.13745011 | 0 |
| GH10229 | FBgn0027554 | CG8042 | 0.205169788 | 0.12541392 | 0 |
| LD47544 | FBgn0052633 | CG32633 | -0.204990666 | 0.20926237 | 0 |
| LD47544 | FBgn0051323 | CG31323 | -0.204990666 | 0.20926237 | 0 |
| LD47544 | FBgn0051086 | CG31086 | -0.204990666 | 0.20926237 | 0 |
| LP06704 | FBgn0035976 | PGRP-LC | 0.204969033 | 0.4085175 | 0 |
| CK00026 | FBgn0028527 | CG18507 | -0.204912751 | 0.56089775 | 0 |
| GH16267 | FBgn0032771 | CG17349 | 0.204851718 | 0.26370926 | 0 |
| RH61147 | FBgn0020765 | Acp65Aa | -0.204810109 | 0.23695841 | 0 |
| LD34757 | FBgn0283510 | Pal1 | 0.204761075 | 0.24960785 | 0 |
| SD24069 | FBgn0035152 | CG3386 | 0.204759023 | 0.2481764 | 0 |
| AT28810 | FBgn0263441 | asRNA:CR43464 | 0.204703133 | 0.27050445 | 0 |
| LD33339 | FBgn0033688 | Prp8 | 0.204702941 | 0.19503434 | 0 |
| GM14158 | FBgn0015805 | HDAC1 | 0.204687238 | 0.13675153 | 0 |
| GM06547 | FBgn0250867 | CG42238 | 0.204629386 | 0.20693481 | 0 |
| GH18123 | FBgn0028969 | deltaCOP | -0.204538158 | 0.11882441 | 0 |
| SD03282 | FBgn0266572 | ebo | 0.204478767 | 0.12482369 | 0 |
| RH59280 | FBgn0052726 | CG32726 | -0.204457566 | 0.20810798 | 0 |
| HL08042 | FBgn0283471 | wupA | 0.204442369 | 0.74625318 | 0 |
| RE72980 | FBgn0002570 | Mal-A1 | 0.204431328 | 0.11882441 | 0 |
| GH15731 | FBgn0025387 | CG12184 | 0.204403096 | 0.1332109 | 0 |
| GM05521 | FBgn0086906 | sls | 0.204371018 | 0.23140423 | 0 |
| LD40824 | FBgn0005642 | wdn | 0.20436594 | 0.1652292 | 0 |
| GH11385 | FBgn0038465 | Irc | 0.204356981 | 0.31861085 | 0 |
| GH04922 | FBgn0028496 | CG30116 | -0.204274823 | 0.18468714 | 0 |
| GH15081 | FBgn0037758 | CG9467 | 0.204269723 | 0.22830906 | 0 |

| | | | | | |
|---|---|---|---|---|---|
| GH10635 | FBgn0015714 | Cyp6a17 | -0.204257225 | 0.12487471 | 0 |
| GH24548 | FBgn0029940 | CG1958 | 0.204253762 | 0.33491792 | 0 |
| LD14121 | FBgn0039638 | dgt6 | 0.204161591 | 0.12972622 | 0 |
| LP08842 | FBgn0039154 | Npc2f | 0.204095364 | 0.2767948 | 0 |
| GH21114 | FBgn0035154 | CG3344 | -0.204091625 | 0.21079071 | 0 |
| RE45108 | FBgn0035591 | CG4611 | 0.204059911 | 0.15310733 | 0 |
| RE05521 | FBgn0035926 | CG5804 | -0.203962464 | 0.24999794 | 0 |
| LD23326 | FBgn0028744 | CG5033 | 0.203944107 | 0.15489812 | 0 |
| RE08109 | FBgn0036207 | CG10907 | 0.203944042 | 0.11975128 | 0 |
| LD24662 | FBgn0030365 | Tango4 | 0.203943176 | 0.14743733 | 0 |
| LP12301 | FBgn0024289 | Sodh-1 | 0.203728427 | 0.16307179 | 0 |
| GH03286 | FBgn0032485 | CG9426 | -0.203698146 | 0.19635709 | 0 |
| SD03724 | FBgn0005386 | ash1 | 0.203677553 | 0.12320472 | 0 |
| AT21921 | FBgn0035693 | CG8219 | 0.203649217 | 0.15810151 | 0 |
| LD31448 | FBgn0037700 | CG8149 | 0.203614682 | 0.23906935 | 0 |
| LD41072 | FBgn0033998 | row | 0.20359609 | 0.11650145 | 0 |
| RE09223 | FBgn0263863 | mRpL45 | 0.203568487 | 0.17110273 | 0 |
| LD02349 | FBgn0001994 | crp | 0.203538291 | 0.12216626 | 0 |
| RE65203 | FBgn0037249 | eIF3a | 0.203528603 | 0.17512775 | 0 |
| LP01553 | FBgn0265052 | St3 | 0.203509909 | 0.2626156 | 0 |
| LP09564 | FBgn0004435 | Galphaq | 0.203488579 | 0.15310733 | 0 |
| SD01848 | FBgn0035383 | CPT2 | 0.203465111 | 0.1510648 | 0 |
| SD06626 | FBgn0044324 | Chro | 0.203443918 | 0.12487471 | 0 |
| SD06626 | FBgn0267630 | asRNA:CR45968 | 0.203443918 | 0.12487471 | 0 |
| GH07253 | FBgn0086704 | stops | -0.203388351 | 0.24554412 | 0 |
| LD30439 | FBgn0030630 | CG12608 | 0.203367222 | 0.13243618 | 0 |
| LD30439 | FBgn0030629 | CG9123 | 0.203367222 | 0.13243618 | 0 |
| LD36241 | FBgn0035793 | CG7546 | 0.203104377 | 0.14956142 | 0 |
| LD21694 | FBgn0034939 | thoc5 | 0.20306436 | 0.14864871 | 0 |
| AT09449 | FBgn0003978 | vls | -0.203036763 | 0.12020594 | 0 |
| GH08125 | FBgn0034000 | CG11808 | 0.202903922 | 0.17733893 | 0 |
| AT18937 | FBgn0031997 | PGAP5 | -0.202726163 | 0.14248667 | 0 |
| RE15157 | FBgn0038422 | CG14880 | -0.202610779 | 0.20639193 | 0 |
| LD32066 | FBgn0001085 | fz | 0.202589658 | 0.2091573 | 0 |
| LP07547 | FBgn0031782 | WDR79 | 0.202584472 | 0.17110273 | 0 |
| LD16161 | FBgn0031873 | Gas41 | 0.202490517 | 0.21754743 | 0 |
| GH07301 | FBgn0035189 | CG9119 | 0.202464839 | 0.16476127 | 0 |
| LD12812 | FBgn0014037 | Su(Tpl) | 0.202423834 | 0.13453026 | 0 |
| GH07283 | FBgn0053653 | Cadps | 0.202384917 | 0.1332109 | 0 |
| GH12489 | FBgn0046296 | CG11534 | -0.202327598 | 0.1241051 | 0 |
| GM05887 | FBgn0017482 | T3dh | -0.202248257 | 0.18456907 | 0 |
| LP11495 | FBgn0033212 | LRR | 0.202227577 | 0.20639193 | 0 |
| RE15719 | FBgn0001987 | Gli | 0.20213461 | 0.17693712 | 0 |
| SD01953 | FBgn0000244 | by | 0.202097808 | 0.1370972 | 0 |
| RE54552 | FBgn0010015 | CanA1 | -0.202066399 | 0.23621839 | 0 |
| LP02729 | FBgn0030495 | CG11816 | -0.201970918 | 0.50722792 | 0 |
| GH17891 | FBgn0041194 | Prat2 | -0.201843533 | 0.33312037 | 0 |
| RE64176 | FBgn0263355 | CG31688 | -0.201784748 | 0.12841406 | 0 |

| | | | | | |
|---|---|---|---|---|---|
| SD26182 | FBgn0022936 | CycH | -0.201584451 | 0.13933311 | 0 |
| LD19244 | FBgn0026577 | CG8677 | 0.201552138 | 0.15557434 | 0 |
| RE05601 | FBgn0031770 | CG13995 | -0.201551551 | 0.42117179 | 0 |
| GH05665 | FBgn0037688 | CG9356 | 0.201490681 | 0.19192961 | 0 |
| LD42626 | FBgn0010416 | TH1 | 0.201479837 | 0.13904355 | 0 |
| LD02709 | FBgn0010342 | Map60 | 0.201431834 | 0.19101365 | 0 |
| LD15107 | FBgn0037218 | aux | 0.201363427 | 0.24392475 | 0 |
| GH14506 | FBgn0030706 | Lrp4 | -0.201343257 | 0.11742225 | 0 |
| SD09607 | FBgn0038746 | Surf6 | 0.201312735 | 0.16856818 | 0 |
| RE58505 | FBgn0011225 | jar | -0.201306782 | 0.12034993 | 0 |
| LD23505 | FBgn0284421 | Psn | 0.201246058 | 0.19304214 | 0 |
| LD21421 | FBgn0034114 | CG4282 | 0.201233227 | 0.12366389 | 0 |
| AT07608 | FBgn0034833 | CG13539 | 0.201220369 | 0.21976131 | 0 |
| LP10918 | FBgn0250815 | Jon65Aiv | -0.201100806 | 0.4047529 | 0 |
| RE39008 | FBgn0030970 | CG7326 | -0.201093721 | 0.11860764 | 0 |
| LD27256 | FBgn0033350 | CG8237 | 0.200962721 | 0.15971906 | 0 |
| RE07981 | FBgn0266717 | Bruce | -0.200934928 | 0.39339733 | 0 |
| GH23453 | FBgn0050344 | CG30344 | 0.200887565 | 0.12837374 | 0 |
| RE07178 | FBgn0041004 | CG17715 | -0.200773206 | 0.20642679 | 0 |
| LD23951 | FBgn0038735 | CG4662 | 0.200686603 | 0.13624799 | 0 |
| GH12838 | FBgn0034118 | Nup62 | 0.200614128 | 0.1762235 | 0 |
| LD02947 | FBgn0037881 | GCC88 | 0.200600567 | 0.13356813 | 0 |
| LD27336 | FBgn0032138 | CG4364 | 0.200569021 | 0.18472354 | 0 |
| LD46862 | FBgn0032517 | CG7099 | 0.200567471 | 0.13473737 | 0 |
| SD05726 | FBgn0043364 | cbt | -0.200442651 | 0.21956473 | 0 |
| AT13706 | FBgn0039651 | Cyt-c1L | 0.200401075 | 0.26814996 | 0 |
| LD23509 | FBgn0027084 | LysRS | 0.200361728 | 0.21524805 | 0 |
| LD24490 | FBgn0038639 | CG7705 | 0.20028806 | 0.12147699 | 0 |
| GH18658 | FBgn0029859 | CG15892 | -0.200260739 | 0.12931559 | 0 |
| GH18658 | FBgn0029860 | CG15891 | -0.200260739 | 0.12931559 | 0 |
| LD02639 | FBgn0027363 | Stam | 0.200004517 | 0.14027093 | 0 |
| LD22726 | FBgn0001978 | stc | 0.199835227 | 0.14079552 | 0 |
| LD47774 | FBgn0260741 | CG3281 | 0.199825734 | 0.16214404 | 0 |
| LD21116 | FBgn0027080 | TyrRS | 0.199816021 | 0.13791478 | 0 |
| SD05212 | FBgn0014380 | RhoL | 0.199720009 | 0.32095126 | 0 |
| AT06220 | FBgn0036311 | CG17666 | 0.199684534 | 0.30377784 | 0 |
| LD20253 | FBgn0015509 | Cul1 | 0.199641165 | 0.14294744 | 0 |
| LD26519 | FBgn0067779 | dbr | 0.199480529 | 0.27578495 | 0 |
| GH08388 | FBgn0027564 | CG3149 | 0.199428824 | 0.20380583 | 0 |
| RE20371 | FBgn0035186 | CG13912 | -0.199378023 | 0.37234905 | 0 |
| GH12067 | FBgn0263916 | Ent2 | 0.199229117 | 0.18212145 | 0 |
| GH19331 | FBgn0052459 | CG32459 | -0.199213725 | 0.14708774 | 0 |
| LD24492 | FBgn0028737 | eEF1beta | 0.199174423 | 0.25900832 | 0 |
| LD42119 | FBgn0030306 | Spase25 | 0.199131399 | 0.21444299 | 0 |
| SD02285 | FBgn0261444 | CG3638 | -0.19901461 | 0.28082823 | 0 |
| GH10272 | FBgn0033184 | mEFTu2 | 0.198966357 | 0.13207685 | 0 |
| GH09576 | FBgn0052268 | Drsl6 | 0.1989414 | 0.37074408 | 0 |
| GH09576 | FBgn0035434 | Drsl5 | 0.1989414 | 0.37074408 | 0 |

| | | | | | |
|---|---|---|---|---|---|
| LD27111 | FBgn0285892 | tea | -0.198868294 | 0.13453026 | 0 |
| GH27720 | FBgn0037985 | ssp5 | -0.198858287 | 0.19968924 | 0 |
| RE60337 | FBgn0033028 | CG11665 | -0.198805479 | 0.11985845 | 0 |
| GH18278 | FBgn0034396 | CG15097 | -0.198794849 | 0.17136716 | 0 |
| LD22387 | FBgn0262955 | RpII140 | 0.198791334 | 0.15744362 | 0 |
| LD37736 | FBgn0030096 | Zpr1 | 0.198749342 | 0.18882961 | 0 |
| LD09973 | FBgn0003964 | usp | -0.198716817 | 0.12320472 | 0 |
| RE60387 | FBgn0016672 | Ipp | -0.198693375 | 0.13005317 | 0 |
| LD01482 | FBgn0037473 | CG10068 | -0.19863284 | 0.12020594 | 0 |
| LD34793 | FBgn0020388 | Gcn5 | 0.19858858 | 0.16309185 | 0 |
| AT01351 | FBgn0033463 | CG1513 | 0.19853682 | 0.13968045 | 0 |
| GH03273 | FBgn0030668 | CG8128 | 0.198520951 | 0.20512018 | 0 |
| LD47533 | FBgn0004698 | Xpc | 0.198454448 | 0.15721637 | 0 |
| RE70650 | FBgn0035155 | RabX6 | -0.198436121 | 0.21862256 | 0 |
| GH16072 | FBgn0028926 | NC2beta | 0.198378224 | 0.22894331 | 0 |
| GH07269 | FBgn0028490 | CG31705 | 0.198374167 | 0.23906935 | 0 |
| LD21715 | FBgn0259175 | ome | 0.198336379 | 0.27578495 | 0 |
| GH12153 | FBgn0022787 | Hel89B | 0.19831546 | 0.14693767 | 0 |
| GH27944 | FBgn0033809 | CG4630 | -0.198310804 | 0.22266915 | 0 |
| LD47536 | FBgn0001142 | Gs1 | -0.198283961 | 0.12497934 | 0 |
| RE03728 | FBgn0001319 | kn | -0.198207271 | 0.63765433 | 0 |
| LD23870 | FBgn0037707 | RnpS1 | 0.198107824 | 0.26550114 | 0 |
| GH04194 | FBgn0031497 | SerRS | -0.197944816 | 0.25865893 | 0 |
| RH26548 | FBgn0086909 | CG31751 | -0.197870681 | 0.15799279 | 0 |
| AT17586 | FBgn0051245 | CG31245 | 0.197818903 | 0.1366984 | 0 |
| RE39106 | FBgn0038426 | mRpS33 | -0.197809317 | 0.29410644 | 0 |
| AT18239 | FBgn0036465 | Rpn12R | -0.197766504 | 0.15063867 | 0 |
| GH15728 | FBgn0035237 | CG13917 | -0.197724079 | 0.21322625 | 0 |
| AT13707 | FBgn0259238 | CG42336 | 0.197436096 | 0.1366984 | 0 |
| LD28893 | FBgn0030504 | CG2691 | 0.197410672 | 0.17147856 | 0 |
| RE59356 | FBgn0037672 | sage | 0.197370764 | 0.18836653 | 0 |
| GM12121 | FBgn0038723 | CG6195 | -0.197361997 | 0.16250591 | 0 |
| LD22082 | FBgn0032005 | Snx6 | 0.1973418 | 0.2171281 | 0 |
| LD19311 | FBgn0035252 | CG7970 | 0.197316532 | 0.14419172 | 0 |
| GH03227 | FBgn0038052 | CG5538 | -0.197188121 | 0.39615894 | 0 |
| LD47792 | FBgn0037270 | eIF3f1 | 0.197078826 | 0.2088032 | 0 |
| RE32047 | FBgn0034931 | CG2812 | 0.196978493 | 0.25872145 | 0 |
| RE37186 | FBgn0030506 | DNAlig4 | -0.196972917 | 0.12227708 | 0 |
| GM08802 | FBgn0029867 | CG3847 | 0.196943741 | 0.131545 | 0 |
| SD02116 | FBgn0026309 | aft | -0.196938982 | 0.13052046 | 0 |
| RE15974 | FBgn0261446 | CG13377 | 0.196919997 | 0.13316275 | 0 |
| GH23990 | FBgn0033949 | CG10131 | 0.196877967 | 0.37005256 | 0 |
| RE56212 | FBgn0032847 | Taf13 | 0.196800182 | 0.17551784 | 0 |
| RE17089 | FBgn0036973 | Rbbp5 | 0.196800112 | 0.17140922 | 0 |
| RE38619 | FBgn0010808 | Chchd3 | 0.196769741 | 0.14961497 | 0 |
| GH02265 | FBgn0039816 | CG11317 | -0.19673743 | 0.25086746 | 0 |
| LD33878 | FBgn0033186 | CG1602 | 0.196655946 | 0.12937686 | 0 |
| LD08814 | FBgn0039691 | IntS11 | 0.196631521 | 0.20879833 | 0 |

| | | | | | |
|---|---|---|---|---|---|
| AT02159 | FBgn0035239 | CG18170 | 0.196611818 | 0.20879833 | 0 |
| AT02159 | FBgn0035240 | CG33791 | 0.196611818 | 0.20879833 | 0 |
| RE20389 | FBgn0032746 | CG10470 | -0.196600374 | 0.28147259 | 0 |
| RE49565 | FBgn0039461 | CG5500 | -0.19659503 | 0.13983076 | 0 |
| RE14640 | FBgn0032877 | CG2617 | 0.196576781 | 0.12276344 | 0 |
| LD36812 | FBgn0039234 | Nct | 0.196565428 | 0.21862256 | 0 |
| GH25878 | FBgn0042693 | wrd | 0.196541847 | 0.19608741 | 0 |
| LD35592 | FBgn0032476 | CG5439 | 0.19647636 | 0.15024149 | 0 |
| GH06193 | FBgn0027581 | CG6191 | -0.196462678 | 0.19635709 | 0 |
| GH06193 | FBgn0266489 | CG45088 | -0.196462678 | 0.19635709 | 0 |
| RE18169 | FBgn0031190 | CG12576 | -0.196426789 | 0.12693221 | 0 |
| RH43343 | FBgn0010411 | RpS18 | -0.196314852 | 0.20831958 | 0 |
| LP07366 | FBgn0033234 | MFS12 | -0.196212679 | 0.16903264 | 0 |
| RH45234 | FBgn0032835 | CG16772 | -0.196183624 | 0.48186414 | 0 |
| GH05505 | FBgn0037689 | CG8135 | -0.196005394 | 0.12320472 | 0 |
| GH05505 | FBgn0260237 | CG42511 | -0.196005394 | 0.12320472 | 0 |
| RE70136 | FBgn0026371 | SAK | 0.195965784 | 0.27806064 | 0 |
| RE60627 | FBgn0014006 | Ask1 | -0.195822146 | 0.14412264 | 0 |
| RH01588 | FBgn0034218 | CG18467 | -0.195811925 | 0.26595429 | 0 |
| LD33215 | FBgn0031062 | CG14230 | 0.195794096 | 0.16476127 | 0 |
| RE28848 | FBgn0001079 | fu | 0.195675477 | 0.24336599 | 0 |
| GH05942 | FBgn0035995 | CG3529 | -0.195670317 | 0.13624799 | 0 |
| LD30318 | FBgn0031256 | shv | 0.195590422 | 0.14854672 | 0 |
| LD30725 | FBgn0031078 | Nup205 | 0.195577713 | 0.12541392 | 0 |
| GH02003 | FBgn0086708 | stv | -0.19556995 | 0.35183241 | 0 |
| LD31569 | FBgn0086908 | egg | 0.195394841 | 0.1683216 | 0 |
| LD30543 | FBgn0037150 | CG7133 | 0.195370845 | 0.12611609 | 0 |
| SD04906 | FBgn0283741 | prage | -0.195320998 | 0.38011332 | 0 |
| GH09638 | FBgn0032030 | Wdr82 | 0.195301985 | 0.14876696 | 0 |
| LD28117 | FBgn0030655 | CG9213 | 0.195208545 | 0.13682108 | 0 |
| GH04031 | FBgn0014032 | Sptr | -0.195194104 | 0.25656875 | 0 |
| RE45749 | FBgn0051106 | CG31106 | 0.195144704 | 0.20123711 | 0 |
| GH06023 | FBgn0283652 | Rpp30 | 0.195054098 | 0.25696915 | 0 |
| RE13344 | FBgn0015618 | Cdk8 | -0.195038386 | 0.25489759 | 0 |
| LP11479 | FBgn0264574 | Glut1 | 0.194906569 | 0.24048374 | 0 |
| LD21294 | FBgn0030607 | dob | 0.194854276 | 0.1626251 | 0 |
| GH11472 | FBgn0284257 | Ttd14 | -0.194793356 | 0.14478603 | 0 |
| LD29992 | FBgn0027835 | Dp1 | 0.194746516 | 0.15489812 | 0 |
| RE27528 | FBgn0032919 | CG9253 | 0.194724402 | 0.15310733 | 0 |
| LD41874 | FBgn0032783 | CG10237 | -0.194679842 | 0.15472004 | 0 |
| RE38276 | FBgn0266465 | GckIII | -0.194679797 | 0.1246311 | 0 |
| GH28686 | FBgn0086442 | mib2 | 0.194676354 | 0.1525526 | 0 |
| LD21426 | FBgn0031148 | Cbs | 0.194614889 | 0.17456233 | 0 |
| SD17695 | FBgn0039225 | Ets96B | -0.19458626 | 0.31217459 | 0 |
| GH06271 | FBgn0037301 | Mms19 | 0.19443512 | 0.2088032 | 0 |
| SD10926 | FBgn0035001 | Slik | 0.194371021 | 0.12541392 | 0 |
| LD40758 | FBgn0039055 | Rassf | 0.19434667 | 0.12693221 | 0 |
| GH21903 | FBgn0032857 | CG10947 | -0.194303629 | 0.24742933 | 0 |

| | | | | | |
|---|---|---|---|---|---|
| LD05823 | FBgn0037244 | CG14647 | -0.194282467 | 0.19904473 | 0 |
| LD46590 | FBgn0015623 | Cpr | 0.194273637 | 0.17434219 | 0 |
| LD15323 | FBgn0032191 | CG5734 | -0.194163289 | 0.14614309 | 0 |
| GH06357 | FBgn0030276 | Dlic | -0.194081652 | 0.14164577 | 0 |
| RE52723 | FBgn0038371 | Pbp45 | 0.194049606 | 0.25865893 | 0 |
| AT30027 | FBgn0035727 | CG10063 | 0.194040068 | 0.14248667 | 0 |
| RE39037 | FBgn0035533 | Cip4 | 0.193864907 | 0.13970977 | 0 |
| LD13184 | FBgn0265295 | asRNA:CR42871 | 0.193724597 | 0.15667569 | 0 |
| GH17483 | FBgn0030925 | Hayan | -0.193721285 | 0.23069212 | 0 |
| LD27810 | FBgn0032600 | BuGZ | 0.193639579 | 0.21635313 | 0 |
| LD07362 | FBgn0000455 | Dip-C | 0.193569614 | 0.131545 | 0 |
| LD07362 | FBgn0065087 | snmRNA:649 | 0.193569614 | 0.131545 | 0 |
| LD33044 | FBgn0026369 | Sara | -0.193486939 | 0.15310733 | 0 |
| LD47054 | FBgn0011741 | Arp6 | 0.19348449 | 0.13597632 | 0 |
| GH04731 | FBgn0001133 | grau | 0.193385328 | 0.20555225 | 0 |
| LD27069 | FBgn0036353 | CG10171 | 0.193308563 | 0.14614309 | 0 |
| LD20420 | FBgn0030053 | CG12081 | 0.193253152 | 0.12590333 | 0 |
| GH19849 | FBgn0038912 | CG6656 | 0.193238349 | 0.12693221 | 0 |
| LD31522 | FBgn0042134 | Capr | -0.19320849 | 0.18538505 | 0 |
| LD24618 | FBgn0011205 | fbl | 0.193083248 | 0.13356813 | 0 |
| AT05393 | FBgn0035113 | pyx | 0.193077755 | 0.30301105 | 0 |
| RE40955 | FBgn0001168 | h | 0.193077372 | 0.12702453 | 0 |
| SD23896 | FBgn0039702 | Vps16B | -0.193050821 | 0.12947861 | 0 |
| LD20718 | FBgn0030418 | CG4004 | 0.193023624 | 0.16307179 | 0 |
| LD24380 | FBgn0261458 | capt | 0.192924964 | 0.25794301 | 0 |
| RE43755 | FBgn0029905 | Nf-YC | -0.192759203 | 0.19232116 | 0 |
| LD46678 | FBgn0037490 | CG10053 | 0.192565938 | 0.25821219 | 0 |
| RE42025 | FBgn0037364 | Rab23 | -0.19249119 | 0.19895429 | 0 |
| LP07057 | FBgn0004373 | fwd | 0.192486961 | 0.16595428 | 0 |
| RE51966 | FBgn0036873 | CG18294 | -0.19247791 | 0.18468714 | 0 |
| SD07723 | FBgn0261041 | stj | -0.192477119 | 0.56216758 | 0 |
| SD27051 | FBgn0028426 | RNaseZ | -0.192421877 | 0.13175758 | 0 |
| GH16914 | FBgn0025608 | Faf2 | 0.192413586 | 0.16577056 | 0 |
| HL01610 | FBgn0051687 | CG31687 | -0.192321889 | 0.13417211 | 0 |
| GH02751 | FBgn0027603 | Ulp1 | 0.192275621 | 0.18031733 | 0 |
| GH02751 | FBgn0031208 | CG11023 | 0.192275621 | 0.18031733 | 0 |
| LD03060 | FBgn0040271 | Sulf1 | 0.192246213 | 0.1366984 | 0 |
| LD46805 | FBgn0043903 | dome | 0.192092644 | 0.14164577 | 0 |
| RE58428 | FBgn0051869 | CG31869 | -0.192072307 | 0.25348968 | 0 |
| GM13219 | FBgn0039489 | CG5880 | 0.192062711 | 0.16534452 | 0 |
| RE52884 | FBgn0025373 | Fpps | 0.192026491 | 0.15362605 | 0 |
| RE10344 | FBgn0050463 | CG30463 | -0.191940119 | 0.16822308 | 0 |
| GM03003 | FBgn0026083 | tyf | 0.19182655 | 0.13178906 | 0 |
| SD02894 | FBgn0030603 | CG5541 | 0.191825424 | 0.1363073 | 0 |
| RH74005 | FBgn0011695 | EbpIII | -0.191809613 | 0.28754289 | 0 |
| LD42380 | FBgn0023216 | Parg | 0.191763823 | 0.131545 | 0 |
| AT19233 | FBgn0259917 | CG42446 | -0.19175886 | 0.21791239 | 0 |
| RH06304 | FBgn0030272 | CG15201 | 0.191631271 | 0.23906935 | 0 |

| | | | | | |
|---|---|---|---|---|---|
| LP11161 | FBgn0010223 | Galphaf | -0.191518249 | 0.28123934 | 0 |
| RE64116 | FBgn0040319 | Gclc | -0.191462195 | 0.15024149 | 0 |
| AT16994 | FBgn0261588 | pdm3 | -0.191333752 | 0.37582281 | 0 |
| LD21756 | FBgn0003279 | RpL4 | 0.191294201 | 0.33110818 | 0 |
| SD08803 | FBgn0262743 | Fs(2)Ket | -0.191245057 | 0.29009021 | 0 |
| LD12474 | FBgn0028506 | CG4455 | 0.191063232 | 0.13243618 | 0 |
| GM05817 | FBgn0053155 | CG33155 | -0.191055673 | 0.15134248 | 0 |
| GM05817 | FBgn0050481 | mRpL53 | -0.191055673 | 0.15134248 | 0 |
| RH04757 | FBgn0011745 | Arp1 | -0.191054976 | 0.17341214 | 0 |
| LD15904 | FBgn0259785 | pzg | 0.191051227 | 0.13695182 | 0 |
| AT05602 | FBgn0035477 | CG14982 | -0.191028556 | 0.40457678 | 0 |
| LD41821 | FBgn0039434 | TwdlM | -0.19091571 | 0.23049934 | 0 |
| LD32234 | FBgn0065102 | snmRNA:184 | 0.190888759 | 0.18031733 | 0 |
| LD32234 | FBgn0086026 | snoRNA:Me28S-G2596 | 0.190888759 | 0.18031733 | 0 |
| RE57944 | FBgn0052557 | CG32557 | -0.190874163 | 0.16201098 | 0 |
| LD04789 | FBgn0036522 | CG7372 | 0.190834786 | 0.12841406 | 0 |
| LD47795 | FBgn0035464 | PIG-B | 0.190833622 | 0.19635709 | 0 |
| LP02922 | FBgn0030097 | CG12115 | 0.19082839 | 0.40940941 | 0 |
| GH14470 | FBgn0037994 | eIF3d2 | 0.190800182 | 0.14079552 | 0 |
| GM01959 | FBgn0033241 | CG2915 | 0.190788957 | 0.20102435 | 0 |
| LD27504 | FBgn0043070 | MESK2 | -0.190733148 | 0.38055367 | 0 |
| LD13607 | FBgn0029822 | CG12236 | -0.190666454 | 0.18538505 | 0 |
| LD15404 | FBgn0264090 | CG43759 | -0.190528297 | 0.29332909 | 0 |
| AT16856 | FBgn0035999 | CG3552 | -0.190385939 | 0.21862256 | 0 |
| GM12126 | FBgn0039233 | CG7006 | 0.190326311 | 0.20762985 | 0 |
| RE12890 | FBgn0052243 | CG32243 | -0.190264206 | 0.28400619 | 0 |
| LD31211 | FBgn0022942 | Cbp80 | 0.190253044 | 0.20693481 | 0 |
| LD12639 | FBgn0035807 | CG7492 | -0.190115893 | 0.16222819 | 0 |
| RE53044 | FBgn0033867 | Cpr50Ca | -0.19011023 | 0.28762122 | 0 |
| LD09561 | FBgn0011260 | Sema2a | 0.190061313 | 0.14349017 | 0 |
| SD16985 | FBgn0028515 | EndoGI | -0.19005024 | 0.14171917 | 0 |
| GH17919 | FBgn0026375 | RhoGAPp190 | 0.18999546 | 0.37651512 | 0 |
| RE14971 | FBgn0011591 | fng | 0.189934171 | 0.44885159 | 0 |
| LP05937 | FBgn0262617 | CG43143 | -0.189729809 | 0.30224264 | 0 |
| LP02983 | FBgn0032843 | CG10730 | 0.189716074 | 0.24753663 | 0 |
| LD36375 | FBgn0033474 | CG1407 | -0.189670633 | 0.1463064 | 0 |
| GM14215 | FBgn0020653 | Trxr-1 | -0.189599411 | 0.19136517 | 0 |
| LD28135 | FBgn0033316 | Gle1 | 0.189594272 | 0.17711358 | 0 |
| LD13269 | FBgn0058160 | CG40160 | -0.189486824 | 0.32095126 | 0 |
| LD13269 | FBgn0085570 | CR41243 | -0.189486824 | 0.32095126 | 0 |
| LD13269 | FBgn0266762 | CR45227 | -0.189486824 | 0.32095126 | 0 |
| GM06606 | FBgn0041186 | Slbp | -0.189295845 | 0.2706749 | 0 |
| SD02972 | FBgn0020909 | Rtc1 | 0.189262453 | 0.1734013 | 0 |
| RE20374 | FBgn0039052 | CG6733 | 0.189239887 | 0.28758766 | 0 |
| GH07389 | FBgn0011746 | ana | 0.18923883 | 0.23976406 | 0 |
| RH18934 | FBgn0038324 | CG5038 | -0.18923412 | 0.51256081 | 0 |
| RE27895 | FBgn0004620 | GluRIIA | -0.189214321 | 0.6329865 | 0 |
| RE36115 | FBgn0034095 | CG15701 | -0.189140145 | 0.19242579 | 0 |

| | | | | | |
|---|---|---|---|---|---|
| RH09134 | FBgn0030160 | CG9691 | -0.1891367 | 0.63281434 | 0 |
| RE39480 | FBgn0033437 | CG12926 | -0.189124559 | 0.25948197 | 0 |
| LD16635 | FBgn0031638 | CG11927 | 0.189025005 | 0.13112426 | 0 |
| LD03827 | FBgn0031972 | Wwox | 0.189017406 | 0.14294744 | 0 |
| RE47125 | FBgn0030263 | CG2076 | 0.188977159 | 0.21004507 | 0 |
| AT31172 | FBgn0029809 | CG15767 | 0.188974138 | 0.2353666 | 0 |
| AT31172 | FBgn0261610 | CG42699 | 0.188974138 | 0.2353666 | 0 |
| GH04473 | FBgn0259878 | Fs | -0.188959734 | 0.15429476 | 0 |
| SD08263 | FBgn0039120 | Nup98-96 | 0.188928801 | 0.19494466 | 0 |
| RE73548 | FBgn0039523 | CG12885 | -0.188849572 | 0.34349101 | 0 |
| GH10780 | FBgn0031279 | CG3544 | 0.188840568 | 0.29545439 | 0 |
| RH07106 | FBgn0038195 | CG3061 | -0.188826932 | 0.14379958 | 0 |
| LD28370 | FBgn0037073 | Tsr1 | 0.188819977 | 0.15667569 | 0 |
| GH27705 | FBgn0032666 | CG5758 | -0.188791466 | 0.13906458 | 0 |
| RE49429 | FBgn0004394 | pdm2 | 0.188646982 | 0.26307647 | 0 |
| AT23778 | FBgn0028858 | CG10839 | 0.188608435 | 0.15799279 | 0 |
| RE09466 | FBgn0035517 | CG1265 | -0.188543812 | 0.14419172 | 0 |
| GH05569 | FBgn0034660 | Loxl2 | 0.188503066 | 0.307972 | 0 |
| LD29879 | FBgn0035036 | CG4707 | 0.18843765 | 0.22706789 | 0 |
| AT04666 | FBgn0266428 | asRNA:CR43463 | 0.188257727 | 0.16778459 | 0 |
| GH07166 | FBgn0025186 | ari-2 | 0.188146439 | 0.1626251 | 0 |
| SD01287 | FBgn0030581 | CG14408 | -0.188130515 | 0.37294475 | 0 |
| RE04130 | FBgn0030347 | CG15739 | 0.188093158 | 0.27113412 | 0 |
| LD23513 | FBgn0002022 | Catsup | 0.18801145 | 0.1911202 | 0 |
| LD38355 | FBgn0020620 | RN-tre | 0.18784594 | 0.18387894 | 0 |
| GH10344 | FBgn0030634 | CG9164 | -0.187810252 | 0.56066561 | 0 |
| GM13757 | FBgn0037001 | ND-39 | 0.187804891 | 0.26687802 | 0 |
| LD14963 | FBgn0261380 | mRpL37 | -0.187801976 | 0.34069725 | 0 |
| RH24769 | FBgn0032946 | nrv3 | 0.187763581 | 0.44252208 | 0 |
| LD05471 | FBgn0027783 | SMC2 | 0.187749902 | 0.22199756 | 0 |
| LP08348 | FBgn0030334 | Karl | 0.187724852 | 0.23669958 | 0 |
| GH08043 | FBgn0037538 | CG3223 | 0.187642066 | 0.15852146 | 0 |
| GH01546 | FBgn0032820 | fbp | 0.187489562 | 0.22734671 | 0 |
| LD22255 | FBgn0000566 | Eip55E | 0.187440604 | 0.19930016 | 0 |
| GH25723 | FBgn0022935 | D19A | -0.187347624 | 0.15742794 | 0 |
| LD20186 | FBgn0032078 | C1GalTA | -0.187264296 | 0.18499405 | 0 |
| GH28787 | FBgn0005654 | Orc3 | -0.187248632 | 0.13232211 | 0 |
| GH28787 | FBgn0085344 | CG34315 | -0.187248632 | 0.13232211 | 0 |
| LD30622 | FBgn0261109 | mrn | -0.187237525 | 0.14164577 | 0 |
| GH14162 | FBgn0039332 | alrm | 0.187235755 | 0.45302394 | 0 |
| AT15442 | FBgn0031201 | CG12446 | 0.187123952 | 0.37410463 | 0 |
| GM14618 | FBgn0022023 | eIF3h | 0.187122862 | 0.22695902 | 0 |
| AT14373 | FBgn0039360 | CLS | -0.187068916 | 0.15049835 | 0 |
| LD41978 | FBgn0037644 | CG11964 | -0.187051894 | 0.27067599 | 0 |
| LD22648 | FBgn0028692 | Rpn2 | -0.187031141 | 0.13356813 | 0 |
| LD06164 | FBgn0026751 | XRCC1 | 0.186956424 | 0.14005329 | 0 |
| LD23804 | FBgn0039743 | CG7946 | 0.186836873 | 0.14669281 | 0 |
| LP07727 | FBgn0033400 | CG2063 | 0.186796381 | 0.19758154 | 0 |

| | | | | | |
|---|---|---|---|---|---|
| LP06238 | FBgn0083950 | side-VI | -0.186790321 | 0.36395337 | 0 |
| RE08679 | FBgn0264978 | Slh | -0.186762902 | 0.22447805 | 0 |
| GH27154 | FBgn0052082 | IRSp53 | -0.186692133 | 0.28553785 | 0 |
| GH09035 | FBgn0036848 | CG10424 | 0.186621662 | 0.13368437 | 0 |
| LD37852 | FBgn0000064 | Ald | 0.186453698 | 0.1888044 | 0 |
| AT22662 | FBgn0035009 | CG16837 | 0.186350525 | 0.21285991 | 0 |
| LD27847 | FBgn0259791 | bora | 0.186330017 | 0.20884974 | 0 |
| AT17730 | FBgn0037574 | Coq2 | -0.186249675 | 0.13486731 | 0 |
| RH02566 | FBgn0051313 | CG31313 | 0.186144259 | 0.19232116 | 0 |
| LD09819 | FBgn0286027 | Rnmt | 0.186077567 | 0.14601911 | 0 |
| SD06708 | FBgn0015513 | mbc | -0.186024205 | 0.15080952 | 0 |
| LD05265 | FBgn0001202 | hook | 0.185830381 | 0.16146809 | 0 |
| GH27027 | FBgn0036279 | Ncc69 | 0.185808443 | 0.1954325 | 0 |
| LD11847 | FBgn0033555 | RpS15Ab | -0.185759954 | 0.13983076 | 0 |
| LD11847 | FBgn0010198 | RpS15Aa | -0.185759954 | 0.13983076 | 0 |
| LD11847 | FBgn0267014 | CR45458 | -0.185759954 | 0.13983076 | 0 |
| RE28850 | FBgn0035265 | PIG-Wb | -0.185679099 | 0.17059317 | 0 |
| LD32788 | FBgn0027493 | AdSS | 0.185675906 | 0.16825502 | 0 |
| RE55984 | FBgn0024734 | PRL-1 | -0.185613159 | 0.13767703 | 0 |
| GH17175 | FBgn0039780 | PH4alphaNE1 | -0.185583896 | 0.55622948 | 0 |
| LD38710 | FBgn0032154 | mtDNA-helicase | -0.185564913 | 0.14484845 | 0 |
| RH63285 | FBgn0015781 | P5cr | -0.185483787 | 0.13597632 | 0 |
| GH05218 | FBgn0036857 | CG9629 | 0.185418758 | 0.14270036 | 0 |
| LD28173 | FBgn0039767 | CG2218 | 0.185370784 | 0.24736024 | 0 |
| LP09906 | FBgn0004431 | LysX | -0.185320215 | 0.35756558 | 0 |
| LD33829 | FBgn0001139 | gro | 0.185181542 | 0.16476127 | 0 |
| AT12771 | FBgn0032376 | Tsp33B | -0.185103895 | 0.42584477 | 0 |
| LD21642 | FBgn0033845 | mars | 0.185081983 | 0.15356193 | 0 |
| LD28027 | FBgn0037338 | Snm1 | -0.185025581 | 0.1609697 | 0 |
| LD04951 | FBgn0031317 | Charon | -0.184877233 | 0.29410644 | 0 |
| GH04978 | FBgn0027587 | CG7028 | 0.184808983 | 0.14525232 | 0 |
| GH10703 | FBgn0038593 | Vps39 | 0.184773873 | 0.13551782 | 0 |
| LD18774 | FBgn0034277 | OstDelta | 0.184712278 | 0.16323728 | 0 |
| LD25129 | FBgn0013764 | Chi | 0.184646828 | 0.19503434 | 0 |
| GH18372 | FBgn0022959 | yps | -0.184561568 | 0.30110704 | 0 |
| GH12467 | FBgn0033834 | CG4744 | -0.184557969 | 0.41304632 | 0 |
| RE74890 | FBgn0037835 | CG14687 | 0.184531027 | 0.37446557 | 0 |
| SD06977 | FBgn0025684 | MFS18 | -0.184470827 | 0.17827299 | 0 |
| LP05062 | FBgn0066365 | dyl | -0.184457411 | 0.60742739 | 0 |
| RH68439 | FBgn0034885 | Eglp4 | 0.1844423 | 0.22860822 | 0 |
| RE19513 | FBgn0014391 | sun | -0.184380715 | 0.20920946 | 0 |
| LD20239 | FBgn0034530 | Rcd6 | 0.184379374 | 0.40671034 | 0 |
| LP06731 | FBgn0003209 | raw | -0.184309645 | 0.14876696 | 0 |
| LD27967 | FBgn0260862 | Vti1a | -0.184306151 | 0.20256328 | 0 |
| GH03957 | FBgn0050460 | CG30460 | -0.184299536 | 0.29553005 | 0 |
| HL01777 | FBgn0000414 | Dab | 0.184052474 | 0.23451933 | 0 |
| AT15894 | FBgn0040368 | eIF4E7 | -0.183999917 | 0.25867824 | 0 |
| GM14163 | FBgn0031771 | ND-51 | -0.183974015 | 0.15310733 | 0 |

| | | | | | |
|---|---|---|---|---|---|
| AT02541 | FBgn0032785 | CG10026 | 0.183973239 | 0.22598525 | 0 |
| AT15479 | FBgn0051542 | CG31542 | 0.183935086 | 0.37429525 | 0 |
| LD32040 | FBgn0027949 | msb1l | 0.183923844 | 0.16647323 | 0 |
| RE28061 | FBgn0263231 | bel | -0.183804943 | 0.16763613 | 0 |
| GH09628 | FBgn0033289 | CG2121 | -0.183747678 | 0.25270491 | 0 |
| AT06668 | FBgn0036224 | Rpt4R | 0.1836737 | 0.14784952 | 0 |
| RE73487 | FBgn0039730 | CG7903 | 0.183632046 | 0.17107296 | 0 |
| LD36566 | FBgn0031143 | CG1532 | -0.183455561 | 0.2649278 | 0 |
| AT24870 | FBgn0031469 | CG18558 | -0.183418607 | 0.55455148 | 0 |
| GM30768 | FBgn0037482 | CG10055 | -0.183387544 | 0.19421166 | 0 |
| GM01182 | FBgn0037384 | dgrn | -0.183378622 | 0.17147856 | 0 |
| LD24780 | FBgn0039867 | CstF50 | 0.183290195 | 0.14626288 | 0 |
| RE39020 | FBgn0020617 | Rx | -0.183161731 | 0.3639518 | 0 |
| SD02215 | FBgn0051739 | AspRS-m | 0.183136097 | 0.14270036 | 0 |
| GM14467 | FBgn0035233 | Pex10 | -0.183086107 | 0.16546644 | 0 |
| LD39388 | FBgn0034498 | CG16868 | 0.183053943 | 0.21079071 | 0 |
| LD07144 | FBgn0284442 | Mcm3 | -0.183016647 | 0.2412154 | 0 |
| GH15286 | FBgn0039430 | CG5455 | 0.183011766 | 0.14539209 | 0 |
| RE48592 | FBgn0013276 | Hsp70Ab | -0.183003299 | 0.16552302 | 0 |
| RE48592 | FBgn0013275 | Hsp70Aa | -0.183003299 | 0.16552302 | 0 |
| RE48592 | FBgn0051354 | Hsp70Bbb | -0.183003299 | 0.16552302 | 0 |
| RE48592 | FBgn0013278 | Hsp70Bb | -0.183003299 | 0.16552302 | 0 |
| RE48592 | FBgn0013279 | Hsp70Bc | -0.183003299 | 0.16552302 | 0 |
| RE48592 | FBgn0013277 | Hsp70Ba | -0.183003299 | 0.16552302 | 0 |
| RE48592 | FBgn0001230 | Hsp68 | -0.183003299 | 0.16552302 | 0 |
| RE74287 | FBgn0002571 | Mal-A3 | 0.182956984 | 0.29250904 | 0 |
| SD05804 | FBgn0023000 | mth | 0.182936993 | 0.19643218 | 0 |
| LD15941 | FBgn0015477 | Rme-8 | 0.182719719 | 0.14854672 | 0 |
| GH25564 | FBgn0038046 | CG5641 | -0.182709375 | 0.56450689 | 0 |
| RE64027 | FBgn0003447 | sn | 0.182640335 | 0.26339056 | 0 |
| SD09488 | FBgn0003044 | Pcl | -0.182549039 | 0.14601911 | 0 |
| LD29807 | FBgn0031988 | CG8668 | -0.182540098 | 0.1510648 | 0 |
| SD04693 | FBgn0032731 | Swip-1 | -0.182502412 | 0.14248667 | 0 |
| LD13366 | FBgn0033051 | Strica | 0.182462728 | 0.18959093 | 0 |
| LD25605 | FBgn0000084 | AnxB10 | -0.182461046 | 0.23697189 | 0 |
| GH23228 | FBgn0003396 | shn | 0.182315554 | 0.15957938 | 0 |
| GH01515 | FBgn0028399 | TMS1 | 0.182296939 | 0.18458646 | 0 |
| AT27314 | FBgn0033447 | dila | 0.182212157 | 0.26375792 | 0 |
| GM01339 | FBgn0025629 | CG4045 | -0.182167018 | 0.21994469 | 0 |
| LD21619 | FBgn0013763 | Idgf6 | 0.182114814 | 0.19376318 | 0 |
| LD35174 | FBgn0024983 | CG4293 | 0.182112773 | 0.13997338 | 0 |
| AT31332 | FBgn0034800 | CG3788 | 0.182086803 | 0.1382545 | 0 |
| LD09626 | FBgn0264693 | ens | 0.182077075 | 0.15423895 | 0 |
| RE44625 | FBgn0015520 | nonA-l | 0.181877386 | 0.22672346 | 0 |
| LP04687 | FBgn0005660 | Ets21C | 0.181868202 | 0.16549337 | 0 |
| RE15373 | FBgn0030541 | CG11584 | -0.181837038 | 0.16625089 | 0 |
| LD14687 | FBgn0027453 | Dnz1 | 0.181810941 | 0.21998928 | 0 |
| RE64245 | FBgn0027574 | CG5815 | 0.181684231 | 0.14646998 | 0 |

| | | | | | |
|---|---|---|---|---|---|
| GH12207 | FBgn0033079 | Fmo-2 | 0.18165407 | 0.29671934 | 0 |
| RE38860 | FBgn0034160 | CG5550 | 0.181586738 | 0.19780035 | 0 |
| RE39957 | FBgn0052189 | CG32189 | -0.181564441 | 0.20513188 | 0 |
| RE39957 | FBgn0052188 | CG32188 | -0.181564441 | 0.20513188 | 0 |
| HL01241 | FBgn0037607 | CG8036 | -0.181379255 | 0.23919493 | 0 |
| GH10266 | FBgn0000078 | Amy-d | -0.181319845 | 0.17136716 | 0 |
| GH10266 | FBgn0000079 | Amy-p | -0.181319845 | 0.17136716 | 0 |
| GH07619 | FBgn0035294 | Mfap1 | 0.181287577 | 0.18080336 | 0 |
| LD20919 | FBgn0037703 | JHDM2 | 0.181255612 | 0.14465613 | 0 |
| RE11385 | FBgn0039098 | GILT3 | -0.181186129 | 0.17394337 | 0 |
| RH17222 | FBgn0034399 | CG15083 | -0.181122385 | 0.17917133 | 0 |
| AT07671 | FBgn0039343 | CG5111 | 0.181058038 | 0.16898804 | 0 |
| GH08719 | FBgn0038467 | AdSL | -0.180984249 | 0.17107296 | 0 |
| RE58406 | FBgn0023177 | Pp4-19C | 0.18087261 | 0.14164577 | 0 |
| GH11945 | FBgn0039647 | jus | -0.180839959 | 0.3326856 | 0 |
| RE44040 | FBgn0261429 | lncRNA:CR42646 | 0.180669183 | 0.25250442 | 0 |
| GM01014 | FBgn0259711 | CG42365 | -0.180638551 | 0.15625212 | 0 |
| AT19949 | FBgn0031905 | gudu | 0.180477576 | 0.38135654 | 0 |
| LD21924 | FBgn0036886 | CG9300 | 0.180394293 | 0.16625089 | 0 |
| SD12984 | FBgn0043362 | bchs | 0.180334743 | 0.42258529 | 0 |
| LD13649 | FBgn0034117 | CG7997 | 0.180326842 | 0.20074155 | 0 |
| SD25936 | FBgn0031144 | CG1529 | -0.180288619 | 0.14254937 | 0 |
| GH27547 | FBgn0261938 | mtRNApol | 0.180274814 | 0.16963505 | 0 |
| GH15741 | FBgn0003460 | so | -0.180231739 | 0.34356345 | 0 |
| LD30520 | FBgn0038251 | Hexim | 0.180218974 | 0.22550565 | 0 |
| SD04925 | FBgn0038058 | CG5608 | 0.180138396 | 0.16584151 | 0 |
| LD21720 | FBgn0027518 | Wdr24 | -0.180025606 | 0.19232116 | 0 |
| LP03275 | FBgn0015919 | caup | -0.18001411 | 0.25821219 | 0 |
| AT05466 | FBgn0264464 | asRNA:CR43872 | 0.179993578 | 0.17417057 | 0 |
| AT19485 | FBgn0030207 | CG2887 | 0.179986773 | 0.26240018 | 0 |
| SD02118 | FBgn0034599 | hng1 | 0.17997305 | 0.1609697 | 0 |
| RE14575 | FBgn0011770 | Gip | -0.179970072 | 0.17178797 | 0 |
| GM02923 | FBgn0040752 | Prosap | 0.179759159 | 0.28251594 | 0 |
| GH20987 | FBgn0020496 | CtBP | 0.17971183 | 0.16476127 | 0 |
| LD11958 | FBgn0001255 | ImpE3 | 0.17967925 | 0.20639193 | 0 |
| LD47309 | FBgn0020312 | Tmtc3 | -0.179590475 | 0.15310733 | 0 |
| GM01069 | FBgn0028509 | CenG1A | 0.17957272 | 0.17147856 | 0 |
| LD39850 | FBgn0035027 | CG3511 | 0.179473988 | 0.17341214 | 0 |
| LP04851 | FBgn0038943 | CG5391 | 0.179395315 | 0.41551801 | 0 |
| LD43293 | FBgn0032801 | CG10165 | 0.179374635 | 0.19232116 | 0 |
| LP04042 | FBgn0036040 | CG6749 | -0.179373678 | 0.1613223 | 0 |
| RE21555 | FBgn0035998 | CG3437 | -0.179353275 | 0.29260321 | 0 |
| RE37920 | FBgn0052057 | dpr10 | -0.179258407 | 0.21912098 | 0 |
| RH44960 | FBgn0037328 | RpL35A | 0.1792378 | 0.23566452 | 0 |
| LP12305 | FBgn0039045 | CG17119 | 0.17923703 | 0.30329273 | 0 |
| RE23450 | FBgn0031040 | CG14210 | 0.179231168 | 0.16096423 | 0 |
| AT08277 | FBgn0052436 | CG32436 | -0.179181178 | 0.22332897 | 0 |
| GH12815 | FBgn0052026 | CG32026 | 0.179125829 | 0.1431271 | 0 |

| | | | | | |
|---|---|---|---|---|---|
| LD28328 | FBgn0016122 | Acer | 0.178911261 | 0.15049835 | 0 |
| AT07738 | FBgn0030789 | Rrp45 | -0.17890178 | 0.18702248 | 0 |
| LD35413 | FBgn0051211 | CG31211 | 0.17866108 | 0.15032154 | 0 |
| LD03613 | FBgn0024986 | CG3719 | 0.178133961 | 0.20459308 | 0 |
| LD13435 | FBgn0259685 | crb | 0.178119049 | 0.24826642 | 0 |
| GH15189 | FBgn0051032 | CR31032 | 0.177977327 | 0.23170768 | 0 |
| RE29864 | FBgn0030915 | CG6179 | 0.1779754 | 0.25867824 | 0 |
| SD06579 | FBgn0001961 | Arpc1 | 0.177879946 | 0.29328557 | 0 |
| RH09461 | FBgn0036984 | CG13248 | -0.177812835 | 0.21464103 | 0 |
| LD13191 | FBgn0036958 | CG17233 | -0.177802244 | 0.25250442 | 0 |
| RE53796 | FBgn0036998 | CG5969 | -0.177778947 | 0.20979542 | 0 |
| GH23439 | FBgn0023531 | CG32809 | 0.177756538 | 0.27805782 | 0 |
| AT07769 | FBgn0036738 | CG7542 | 0.177635148 | 0.35191106 | 0 |
| GH09360 | FBgn0037242 | CG9855 | -0.177506884 | 0.18040272 | 0 |
| RH07954 | FBgn0014865 | Mtk | 0.177470083 | 0.28274607 | 0 |
| LD06759 | FBgn0001224 | Hsp23 | 0.177330032 | 0.21784977 | 0 |
| RE55712 | FBgn0037018 | CG4042 | -0.177198563 | 0.25989729 | 0 |
| LD43729 | FBgn0011836 | Taf2 | 0.177191154 | 0.14962497 | 0 |
| LP05860 | FBgn0005586 | Rab3 | -0.177188965 | 0.23196332 | 0 |
| RH42524 | FBgn0028837 | CSN6 | 0.177184826 | 0.25511933 | 0 |
| LD21815 | FBgn0011606 | Klp3A | -0.177071926 | 0.16229632 | 0 |
| GH09958 | FBgn0038436 | Gyc89Db | 0.176859927 | 0.49018183 | 0 |
| GH02837 | FBgn0030921 | CG6290 | -0.176852587 | 0.20879833 | 0 |
| LD40702 | FBgn0036695 | Papst2 | 0.176750216 | 0.17898071 | 0 |
| AT23443 | FBgn0033243 | CG14763 | 0.176725192 | 0.19635709 | 0 |
| GH04637 | FBgn0028497 | CG3530 | 0.176715961 | 0.19786515 | 0 |
| GH01486 | FBgn0261573 | CoRest | 0.176704746 | 0.17407606 | 0 |
| RE37758 | FBgn0001170 | H2.0 | -0.176689255 | 0.31301389 | 0 |
| SD20456 | FBgn0260965 | CG42588 | 0.176649371 | 0.21351144 | 0 |
| LD16109 | FBgn0016926 | Pino | -0.176580766 | 0.38571465 | 0 |
| LD14270 | FBgn0022213 | Cse1 | -0.176533015 | 0.1888044 | 0 |
| RE14559 | FBgn0031266 | Sf3b1 | 0.176507947 | 0.16052533 | 0 |
| AT01724 | FBgn0043900 | pygo | 0.176453485 | 0.16476127 | 0 |
| LD43943 | FBgn0013343 | Syx1A | 0.176441584 | 0.15334832 | 0 |
| RH68082 | FBgn0011280 | Obp19d | 0.176433156 | 0.17794044 | 0 |
| LD22423 | FBgn0032728 | Tango6 | 0.176423189 | 0.17593513 | 0 |
| RE72064 | FBgn0035640 | mad2 | -0.176413973 | 0.19638916 | 0 |
| GH10763 | FBgn0033821 | CG10799 | 0.176324579 | 0.16822308 | 0 |
| AT11641 | FBgn0031333 | CG5561 | -0.176272966 | 0.15489812 | 0 |
| AT12538 | FBgn0261955 | kdn | 0.176216246 | 0.15303711 | 0 |
| GH04419 | FBgn0032287 | CG6415 | 0.176187074 | 0.47903487 | 0 |
| GH24192 | FBgn0035867 | CG13671 | -0.17610376 | 0.19843877 | 0 |
| RE52028 | FBgn0038065 | Snx3 | 0.176031417 | 0.1738415 | 0 |
| LD18882 | FBgn0030067 | Rbm13 | 0.175960637 | 0.17147856 | 0 |
| LD45152 | FBgn0031493 | Sf3b2 | 0.175958095 | 0.1888044 | 0 |
| GH08336 | FBgn0051207 | CG31207 | 0.175917437 | 0.45505432 | 0 |
| AT07234 | FBgn0030015 | CG10761 | 0.175903616 | 0.17201904 | 0 |
| LD04722 | FBgn0034158 | CG5522 | 0.175883911 | 0.19116579 | 0 |

| | | | | | |
|---|---|---|---|---|---|
| GH22081 | FBgn0036939 | CG7365 | -0.175854972 | 0.28912309 | 0 |
| RE06996 | FBgn0050411 | CG30411 | 0.175758781 | 0.16394152 | 0 |
| LD41715 | FBgn0037327 | PEK | 0.175680936 | 0.17737524 | 0 |
| LD22322 | FBgn0021764 | sdk | 0.175651596 | 0.16096423 | 0 |
| GH01960 | FBgn0011296 | l(2)efl | -0.175611 | 0.55695269 | 0 |
| LD10153 | FBgn0026777 | Rad23 | -0.175606538 | 0.26514466 | 0 |
| RE07996 | FBgn0036152 | CG6175 | -0.175472224 | 0.14626288 | 0 |
| GH01188 | FBgn0039379 | CG5886 | 0.175451644 | 0.28146115 | 0 |
| GH12103 | FBgn0050021 | metro | 0.175398823 | 0.31903254 | 0 |
| SD07158 | FBgn0033661 | CG13185 | 0.175393011 | 0.17404184 | 0 |
| RH63449 | FBgn0032100 | CG13108 | -0.175383423 | 0.26892566 | 0 |
| LD18251 | FBgn0003502 | Btk29A | 0.175085153 | 0.16307179 | 0 |
| RE20756 | FBgn0051626 | CG31626 | -0.175064857 | 0.37331777 | 0 |
| SD09005 | FBgn0264326 | DNApol-epsilon255 | -0.175055066 | 0.1595985 | 0 |
| AT24031 | FBgn0034383 | CG17821 | -0.17504266 | 0.29410644 | 0 |
| GH01464 | FBgn0035298 | SCOT | 0.175034571 | 0.16825502 | 0 |
| RE68036 | FBgn0031184 | CG14615 | -0.174882752 | 0.23667113 | 0 |
| LD24834 | FBgn0015926 | dah | 0.174851074 | 0.2323332 | 0 |
| GH24985 | FBgn0025378 | CG3795 | -0.174778858 | 0.15324921 | 0 |
| RE73239 | FBgn0036469 | CG18649 | 0.174722945 | 0.2918496 | 0 |
| LD22651 | FBgn0031947 | CG7154 | 0.174718925 | 0.19858587 | 0 |
| LD24832 | FBgn0032467 | CG9934 | 0.174694575 | 0.25016484 | 0 |
| GH23529 | FBgn0051116 | ClC-a | -0.174638171 | 0.2287555 | 0 |
| GH03315 | FBgn0029843 | Nep1 | -0.174607325 | 0.3610233 | 0 |
| RE63504 | FBgn0066084 | RpL41 | -0.174602283 | 0.17512775 | 0 |
| GH08318 | FBgn0039827 | CG1544 | 0.174551075 | 0.29108403 | 0 |
| RE47429 | FBgn0264894 | CG44085 | -0.174531795 | 0.3610233 | 0 |
| LD36562 | FBgn0086655 | jing | 0.174515647 | 0.15705273 | 0 |
| GH10637 | FBgn0003132 | Pp1-13C | 0.174434802 | 0.16017861 | 0 |
| GH10637 | FBgn0004103 | Pp1-87B | 0.174434802 | 0.16017861 | 0 |
| RH35726 | FBgn0029928 | CG3032 | -0.174431232 | 0.16276021 | 0 |
| RE59554 | FBgn0029718 | mRpL30 | -0.174423519 | 0.19838629 | 0 |
| AT01932 | FBgn0029976 | snz | -0.174381691 | 0.17015286 | 0 |
| GH19706 | FBgn0035251 | Vta1 | -0.174358982 | 0.46911308 | 0 |
| GH19706 | FBgn0040634 | CG4186 | -0.174358982 | 0.46911308 | 0 |
| AT14778 | FBgn0031711 | CG6907 | 0.17435884 | 0.19717375 | 0 |
| LD10456 | FBgn0032988 | Tif-IA | -0.174356975 | 0.29026821 | 0 |
| AT01846 | FBgn0024945 | NitFhit | 0.174099706 | 0.16394152 | 0 |
| GM12116 | FBgn0041164 | armi | -0.174097995 | 0.30946795 | 0 |
| LD18231 | FBgn0034617 | Panx | 0.173999679 | 0.17341214 | 0 |
| LD34229 | FBgn0001180 | hb | 0.173996884 | 0.30835636 | 0 |
| GH25257 | FBgn0030933 | CG6470 | -0.173960987 | 0.36847667 | 0 |
| GH14510 | FBgn0002031 | l(2)37Cc | -0.173844547 | 0.2323332 | 0 |
| RE13953 | FBgn0032420 | CG6583 | -0.173687652 | 0.18882961 | 0 |
| LD47989 | FBgn0037657 | hyx | 0.173662302 | 0.17104539 | 0 |
| AT03531 | FBgn0038892 | CG15498 | 0.17366208 | 0.41921911 | 0 |
| RE66734 | FBgn0017566 | ND-75 | 0.173630751 | 0.16930072 | 0 |
| RE56341 | FBgn0036428 | Gbs-70E | 0.173620339 | 0.18265457 | 0 |

| | | | | | |
|---|---|---|---|---|---|
| AT21758 | FBgn0283658 | muc | 0.173610111 | 0.21474126 | 0 |
| AT19038 | FBgn0029751 | CG17764 | 0.173598757 | 0.23147563 | 0 |
| RH44312 | FBgn0037529 | mRpS9 | 0.173580072 | 0.17107296 | 0 |
| RH15369 | FBgn0035903 | CG6765 | -0.17356956 | 0.22197099 | 0 |
| RE24064 | FBgn0003231 | ref(2)P | 0.173546389 | 0.21487795 | 0 |
| LD31217 | FBgn0033972 | Ciao1 | 0.173533518 | 0.18318521 | 0 |
| LD43563 | FBgn0027057 | CSN1b | 0.173434617 | 0.19930016 | 0 |
| SD01152 | FBgn0036821 | CG3961 | 0.173278483 | 0.25086746 | 0 |
| GH14260 | FBgn0037261 | CG9775 | 0.173248876 | 0.17551784 | 0 |
| GH14260 | FBgn0286036 | CR46358 | 0.173248876 | 0.17551784 | 0 |
| GH14845 | FBgn0036337 | AdenoK | -0.173207697 | 0.21346567 | 0 |
| GH03016 | FBgn0028699 | Rh50 | 0.17317827 | 0.35405702 | 0 |
| LD44094 | FBgn0037117 | CG11248 | 0.173021469 | 0.19454951 | 0 |
| RE01745 | FBgn0035844 | CG13676 | -0.172973381 | 0.22672346 | 0 |
| LD03007 | FBgn0029935 | CG4615 | 0.172958342 | 0.20924814 | 0 |
| LP03361 | FBgn0004592 | Eig71Ee | -0.172937703 | 0.3547438 | 0 |
| RE42475 | FBgn0051997 | CG31997 | 0.172906512 | 0.57143701 | 0 |
| LD46760 | FBgn0030038 | CG1440 | 0.172898322 | 0.15032154 | 0 |
| LD47515 | FBgn0033540 | Elp2 | -0.172894907 | 0.22515731 | 0 |
| LD22664 | FBgn0020261 | pcm | 0.172891632 | 0.1626251 | 0 |
| HL07802 | FBgn0087012 | 5-HT2A | 0.172872962 | 0.15694906 | 0 |
| RE03722 | FBgn0052039 | CG32039 | -0.172847662 | 0.2071339 | 0 |
| SD09478 | FBgn0035060 | Eps-15 | -0.172839971 | 0.16387105 | 0 |
| SD04985 | FBgn0261524 | lic | 0.172748187 | 0.27805782 | 0 |
| GM01233 | FBgn0025582 | eIF3e | 0.172746097 | 0.26820425 | 0 |
| RH21749 | FBgn0030294 | Pa1 | -0.172732877 | 0.21366213 | 0 |
| RE38504 | FBgn0036991 | CG5872 | -0.172601673 | 0.18274693 | 0 |
| RE12806 | FBgn0262599 | SmydA-3 | 0.172421144 | 0.16841384 | 0 |
| AT21416 | FBgn0016693 | Past1 | 0.172420553 | 0.16358493 | 0 |
| RE06652 | FBgn0051360 | CG31360 | -0.172393345 | 0.25511933 | 0 |
| LD07014 | FBgn0266667 | Exo70 | -0.172357296 | 0.20179693 | 0 |
| RE74245 | FBgn0025519 | fidipidine | 0.17234599 | 0.18080336 | 0 |
| LD12174 | FBgn0266136 | Gyc76C | 0.172281643 | 0.15385529 | 0 |
| SD07110 | FBgn0029525 | CG18273 | 0.172249809 | 0.16163549 | 0 |
| GH26230 | FBgn0085408 | Shroom | 0.172214017 | 0.20408049 | 0 |
| SD03430 | FBgn0032401 | Plzf | 0.172043953 | 0.21862256 | 0 |
| RE30174 | FBgn0035355 | CG16985 | 0.172005855 | 0.20519738 | 0 |
| RE26764 | FBgn0036888 | CG9330 | -0.171938165 | 0.20373372 | 0 |
| SD10415 | FBgn0029147 | NtR | -0.171880731 | 0.67205817 | 0 |
| LD16247 | FBgn0039938 | Sox102F | 0.171859166 | 0.17862499 | 0 |
| GH12861 | FBgn0032793 | CG10189 | 0.171819414 | 0.37446557 | 0 |
| RE08678 | FBgn0032042 | CG13398 | -0.171797977 | 0.16794426 | 0 |
| GH13807 | FBgn0027083 | MetRS-m | 0.171777994 | 0.22082195 | 0 |
| SD07147 | FBgn0037963 | Cad87A | 0.171772656 | 0.15586736 | 0 |
| LD21659 | FBgn0037943 | CG14722 | 0.171665607 | 0.17201904 | 0 |
| RE66521 | FBgn0264294 | Cyt-b5 | -0.171612503 | 0.19152934 | 0 |
| RE66521 | FBgn0033190 | Gpo2 | -0.171612503 | 0.19152934 | 0 |
| SD05406 | FBgn0015218 | eIF4E1 | -0.171579467 | 0.29545439 | 0 |

| | | | | | |
|---|---|---|---|---|---|
| GM03859 | FBgn0038191 | CG9925 | -0.171541119 | 0.28616681 | 0 |
| GH10944 | FBgn0025626 | CG4281 | 0.171374976 | 0.26066785 | 0 |
| LD08455 | FBgn0052819 | CG32819 | 0.171354748 | 0.31103764 | 0 |
| LD08455 | FBgn0052820 | CG32820 | 0.171354748 | 0.31103764 | 0 |
| LD40028 | FBgn0052423 | shep | -0.171326857 | 0.22332897 | 0 |
| GM01240 | FBgn0086066 | snoRNA:Me5.8S-G74 | 0.171317153 | 0.15698966 | 0 |
| GM01240 | FBgn0266694 | lncRNA:CR45184 | 0.171317153 | 0.15698966 | 0 |
| LD01189 | FBgn0031374 | Wdr62 | 0.171269215 | 0.17529678 | 0 |
| SD01193 | FBgn0001965 | Sos | -0.171238249 | 0.19904473 | 0 |
| LD03170 | FBgn0035761 | RhoGEF4 | 0.17120575 | 0.16321212 | 0 |
| LD27553 | FBgn0014931 | CG2678 | 0.171194493 | 0.20576619 | 0 |
| LD29816 | FBgn0029157 | ssh | -0.171185384 | 0.23101323 | 0 |
| RE58280 | FBgn0004227 | nonA | 0.171175017 | 0.15738589 | 0 |
| RE58280 | FBgn0039979 | CR40282 | 0.171175017 | 0.15738589 | 0 |
| LD43147 | FBgn0013717 | not | 0.171172741 | 0.18499405 | 0 |
| LP11520 | FBgn0039427 | CG5447 | -0.171075467 | 0.25143788 | 0 |
| GH11824 | FBgn0037329 | POLDIP2 | 0.171051212 | 0.18075619 | 0 |
| GH19411 | FBgn0051121 | CG31121 | 0.170924486 | 0.23413506 | 0 |
| LD23968 | FBgn0030007 | alpha-PheRS | -0.170913756 | 0.21606032 | 0 |
| AT01695 | FBgn0062517 | CG16984 | 0.170869859 | 0.34916277 | 0 |
| RE49802 | FBgn0284220 | Top2 | 0.170807606 | 0.22332897 | 0 |
| AT27448 | FBgn0266709 | Zmynd10 | 0.170801433 | 0.22199756 | 0 |
| LD40450 | FBgn0037070 | CG11309 | -0.170717936 | 0.22736386 | 0 |
| RH11587 | FBgn0036839 | SmydA-2 | 0.170551978 | 0.32282288 | 0 |
| RE50942 | FBgn0023170 | RpL39 | 0.17052834 | 0.23983018 | 0 |
| LP09416 | FBgn0010504 | kermit | 0.17052078 | 0.24586808 | 0 |
| GM16979 | FBgn0030520 | Pdcd4 | 0.170483706 | 0.17083669 | 0 |
| LD27070 | FBgn0032721 | CG10602 | -0.170458571 | 0.28473292 | 0 |
| LD43740 | FBgn0051217 | modSP | 0.170421386 | 0.16321212 | 0 |
| GM14375 | FBgn0036574 | elg1 | 0.170393731 | 0.2036043 | 0 |
| AT18617 | FBgn0031546 | CG8851 | -0.170279718 | 0.24719359 | 0 |
| AT24036 | FBgn0034595 | CG15657 | 0.170279342 | 0.24392475 | 0 |
| AT11401 | FBgn0033512 | CG12902 | 0.170195399 | 0.24360361 | 0 |
| AT11401 | FBgn0263271 | CG43397 | 0.170195399 | 0.24360361 | 0 |
| GH12756 | FBgn0263603 | Zn72D | -0.170188973 | 0.1911631 | 0 |
| AT13896 | FBgn0038311 | CG14864 | -0.170083774 | 0.17551784 | 0 |
| LD07917 | FBgn0030245 | CG1637 | 0.170072549 | 0.20879833 | 0 |
| SD03278 | FBgn0029095 | aru | 0.170032069 | 0.3160745 | 0 |
| GH11688 | FBgn0027544 | CG2217 | -0.169964597 | 0.30825767 | 0 |
| GH11688 | FBgn0261704 | CG42740 | -0.169964597 | 0.30825767 | 0 |
| LD32923 | FBgn0250869 | CG42240 | 0.169928297 | 0.1595985 | 0 |
| RH10407 | FBgn0263395 | hppy | -0.169916637 | 0.24828889 | 0 |
| RE69372 | FBgn0038055 | trus | -0.169895419 | 0.27774828 | 0 |
| AT04363 | FBgn0038558 | CG12347 | 0.169879435 | 0.2190254 | 0 |
| LD02044 | FBgn0036298 | nst | 0.16987409 | 0.15624049 | 0 |
| GH28333 | FBgn0040996 | CG12617 | 0.169866068 | 0.44885159 | 0 |
| HL08030 | FBgn0039927 | CG11155 | 0.169818581 | 0.33928392 | 0 |
| GM08821 | FBgn0039831 | CG12054 | 0.169790373 | 0.74880707 | 0 |

| | | | | | |
|---|---|---|---|---|---|
| LD44584 | FBgn0003319 | Sb | -0.169785556 | 0.32282288 | 0 |
| LD29458 | FBgn0025716 | Bap55 | 0.169732619 | 0.23778979 | 0 |
| AT18451 | FBgn0053017 | CG33017 | 0.169729728 | 0.42336335 | 0 |
| RE68566 | FBgn0035146 | CG13893 | 0.169690928 | 0.21741855 | 0 |
| AT23505 | FBgn0033491 | CG18011 | 0.169602164 | 0.17551784 | 0 |
| AT07979 | FBgn0250903 | lmgA | 0.169470328 | 0.16360447 | 0 |
| AT07979 | FBgn0085470 | lmgB | 0.169470328 | 0.16360447 | 0 |
| AT02427 | FBgn0037455 | CG2336 | 0.16943066 | 0.23621839 | 0 |
| RE19116 | FBgn0002609 | E(spl)m3-HLH | -0.169337747 | 0.37845572 | 0 |
| RE42326 | FBgn0030519 | CG11151 | -0.169244503 | 0.18058128 | 0 |
| GH16214 | FBgn0005536 | Mbs | 0.169234726 | 0.16279953 | 0 |
| RE68655 | FBgn0038854 | CG7044 | 0.169170726 | 0.20762985 | 0 |
| RE09574 | FBgn0034567 | CG15651 | -0.169170659 | 0.18959093 | 0 |
| AT25164 | FBgn0243511 | psidin | -0.169133455 | 0.17877971 | 0 |
| LD18710 | FBgn0040212 | Dhap-at | -0.169095835 | 0.17171186 | 0 |
| AT25102 | FBgn0035638 | Tektin-C | 0.169093816 | 0.31052406 | 0 |
| LD10457 | FBgn0011016 | SsRbeta | -0.169025494 | 0.19494466 | 0 |
| SD04526 | FBgn0052452 | CG32452 | 0.168986061 | 0.24250047 | 0 |
| LD41184 | FBgn0031762 | CG9098 | -0.168967771 | 0.1923341 | 0 |
| RE10864 | FBgn0010217 | ATPsynbeta | -0.168909311 | 0.17817627 | 0 |
| GH03795 | FBgn0036035 | CG18178 | 0.168907713 | 0.21782124 | 0 |
| LD41739 | FBgn0039073 | CG4408 | 0.168888169 | 0.19012583 | 0 |
| RH47192 | FBgn0036740 | Vps60 | -0.168819988 | 0.30755546 | 0 |
| AT03573 | FBgn0040092 | lectin-46Cb | 0.168816861 | 0.36847667 | 0 |
| LD23140 | FBgn0001323 | knrl | 0.168615021 | 0.33919374 | 0 |
| RE09547 | FBgn0029785 | RpL35 | -0.168533162 | 0.15774023 | 0 |
| RE16342 | FBgn0035164 | CG13901 | -0.168482431 | 0.38582221 | 0 |
| AT16318 | FBgn0029667 | Gas8 | -0.168476273 | 0.7083456 | 0 |
| RE49877 | FBgn0032329 | Art8 | 0.168459879 | 0.20926155 | 0 |
| GH01059 | FBgn0020497 | emb | 0.16843575 | 0.21117523 | 0 |
| LD45682 | FBgn0036614 | Golgin104 | -0.168411254 | 0.17775002 | 0 |
| GH12794 | FBgn0039054 | Cow | -0.168383428 | 0.40221267 | 0 |
| LD07883 | FBgn0039116 | CG10375 | 0.16828214 | 0.18468714 | 0 |
| GH14494 | FBgn0037433 | CG17919 | 0.168279279 | 0.30194797 | 0 |
| RE30427 | FBgn0041585 | olf186-F | -0.168276083 | 0.20606892 | 0 |
| LD27661 | FBgn0029088 | disp | 0.168255585 | 0.1595985 | 0 |
| LD25913 | FBgn0028380 | fal | -0.168252607 | 0.19245828 | 0 |
| LD34475 | FBgn0028467 | CG11070 | -0.168184507 | 0.24336727 | 0 |
| GH20978 | FBgn0262579 | Ect4 | 0.168171903 | 0.45158951 | 0 |
| AT26678 | FBgn0011743 | Arp53D | 0.168118266 | 0.31677144 | 0 |
| LD09376 | FBgn0003517 | sta | 0.168083492 | 0.38135654 | 0 |
| LD31664 | FBgn0035430 | CG12009 | 0.16805828 | 0.7402025 | 0 |
| SD04793 | FBgn0041789 | Pax | 0.167949664 | 0.17087838 | 0 |
| GH21273 | FBgn0037312 | CG11999 | 0.167899219 | 0.27123661 | 0 |
| GH28721 | FBgn0036876 | CG9451 | 0.167798998 | 0.2591248 | 0 |
| RH09719 | FBgn0033340 | CG13751 | 0.167785591 | 0.21296194 | 0 |
| RH67809 | FBgn0040923 | CG11368 | -0.167724813 | 0.19399445 | 0 |
| GM14547 | FBgn0050055 | lncRNA:CR30055 | -0.167386741 | 0.16096423 | 0 |

| | | | | | |
|---|---|---|---|---|---|
| SD18087 | FBgn0261618 | larp | -0.167285655 | 0.3883281 | 0 |
| LD24318 | FBgn0039712 | CG15514 | 0.167218923 | 0.25143788 | 0 |
| RE14585 | FBgn0034025 | Pgant1 | 0.167188134 | 0.22860822 | 0 |
| RE13191 | FBgn0052176 | CG32176 | 0.167164277 | 0.20380583 | 0 |
| GH09240 | FBgn0261020 | wol | 0.167110247 | 0.17593513 | 0 |
| RE50843 | FBgn0034753 | CG2852 | -0.16707879 | 0.17737524 | 0 |
| GH08894 | FBgn0039849 | CG11334 | 0.167067741 | 0.21464103 | 0 |
| GH09153 | FBgn0039137 | CG13604 | 0.166961972 | 0.2219358 | 0 |
| GM03112 | FBgn0035498 | Fit1 | 0.166900331 | 0.16476127 | 0 |
| RE58623 | FBgn0086532 | Spt-I | 0.166864688 | 0.18268123 | 0 |
| GM23685 | FBgn0051992 | gw | 0.166850622 | 0.36865559 | 0 |
| RH07612 | FBgn0001263 | inaD | -0.166806415 | 0.21312024 | 0 |
| LP02728 | FBgn0030025 | CG2147 | -0.166755236 | 0.24910111 | 0 |
| SD05769 | FBgn0039634 | alpha-Man-Ib | -0.166697744 | 0.21424794 | 0 |
| LD21293 | FBgn0027532 | CG7139 | 0.166642485 | 0.18274693 | 0 |
| RE25263 | FBgn0086710 | RpL30 | -0.166470828 | 0.16322389 | 0 |
| LD45768 | FBgn0037371 | Sym | -0.166432593 | 0.16728274 | 0 |
| LD46336 | FBgn0003512 | Sry-delta | 0.166403959 | 0.22008157 | 0 |
| LD24714 | FBgn0031492 | CG3542 | 0.166388332 | 0.29671934 | 0 |
| GH23865 | FBgn0038536 | CG7655 | 0.166382838 | 0.18468714 | 0 |
| RE43724 | FBgn0284255 | Arpc4 | 0.166373063 | 0.23215197 | 0 |
| RH23043 | FBgn0035046 | ND-19 | -0.166285768 | 0.16620492 | 0 |
| RE54605 | FBgn0038545 | pasi1 | -0.166247038 | 0.33074938 | 0 |
| AT07585 | FBgn0035571 | CG12493 | 0.16618481 | 0.35345047 | 0 |
| RE38958 | FBgn0028953 | CG14478 | -0.166155699 | 0.56989412 | 0 |
| GH07902 | FBgn0020372 | TM4SF | -0.166069953 | 0.17371718 | 0 |
| RE23388 | FBgn0028685 | Rpt4 | 0.166058064 | 0.29012551 | 0 |
| GM04789 | FBgn0037617 | nom | 0.166043788 | 0.22883922 | 0 |
| RH09231 | FBgn0034407 | DptB | -0.166007548 | 0.22005916 | 0 |
| LD29159 | FBgn0031188 | CG14613 | -0.165989066 | 0.2323332 | 0 |
| LD27836 | FBgn0032821 | CdGAPr | 0.165950878 | 0.17405834 | 0 |
| LD23641 | FBgn0039562 | Gp93 | 0.165940297 | 0.25897591 | 0 |
| GH09876 | FBgn0046302 | CG10650 | 0.165900754 | 0.36395337 | 0 |
| GH26183 | FBgn0035241 | CG12105 | -0.165839068 | 0.22451466 | 0 |
| LD15505 | FBgn0031996 | CG8460 | 0.16580329 | 0.18452696 | 0 |
| RE22927 | FBgn0039942 | CG17163 | -0.165801156 | 0.22736386 | 0 |
| LD37169 | FBgn0031037 | CG14207 | 0.165780601 | 0.27527549 | 0 |
| RE47570 | FBgn0035210 | msd5 | 0.165772522 | 0.25086746 | 0 |
| RE63854 | FBgn0002522 | lab | -0.165771996 | 0.41216592 | 0 |
| RH04127 | FBgn0030716 | CG9170 | 0.165749119 | 0.31028912 | 0 |
| LD44970 | FBgn0029118 | Sucb | 0.16562478 | 0.16279953 | 0 |
| LP03102 | FBgn0034989 | CG3356 | -0.165556496 | 0.17417057 | 0 |
| GH15080 | FBgn0000635 | Fas2 | -0.165541231 | 0.17147856 | 0 |
| LP04135 | FBgn0033240 | CG2906 | -0.165490023 | 0.24600826 | 0 |
| GM13876 | FBgn0052056 | scramb1 | -0.165484597 | 0.21322625 | 0 |
| GH26447 | FBgn0262869 | Gfrl | -0.165411094 | 0.51140355 | 0 |
| RH04252 | FBgn0039801 | Npc2h | 0.165381912 | 0.44868134 | 0 |
| RH04252 | FBgn0039800 | Npc2g | 0.165381912 | 0.44868134 | 0 |

| | | | | | |
|---|---|---|---|---|---|
| AT19426 | FBgn0032478 | CG5458 | 0.16524571 | 0.26424444 | 0 |
| LD23767 | FBgn0033902 | eIF3m | 0.165093075 | 0.25086746 | 0 |
| LP09251 | FBgn0037885 | CG17721 | -0.16509262 | 0.27838547 | 0 |
| RH14320 | FBgn0033736 | CG13154 | 0.165031228 | 0.17794044 | 0 |
| LD40094 | FBgn0283724 | Girdin | 0.16496526 | 0.20973394 | 0 |
| RH44796 | FBgn0036262 | CG6910 | 0.164936063 | 0.1911631 | 0 |
| GM02985 | FBgn0033990 | CG10265 | 0.164874465 | 0.2006496 | 0 |
| GM13874 | FBgn0014011 | Rac2 | 0.164816553 | 0.16394152 | 0 |
| GH23377 | FBgn0032699 | CG10383 | 0.164698078 | 0.16822308 | 0 |
| AT17410 | FBgn0032650 | CG7094 | -0.164640261 | 0.17877971 | 0 |
| LD11854 | FBgn0037478 | CG2656 | 0.164606746 | 0.27610621 | 0 |
| AT17414 | FBgn0032176 | CG13127 | 0.164567599 | 0.38251295 | 0 |
| RH50927 | FBgn0037930 | CG14715 | 0.164502605 | 0.23606201 | 0 |
| GH14155 | FBgn0039023 | CG4723 | 0.164494191 | 0.51228718 | 0 |
| LP01609 | FBgn0037094 | CG7611 | 0.164421512 | 0.18274693 | 0 |
| RE72817 | FBgn0036351 | CG14107 | 0.164409727 | 0.36033739 | 0 |
| GH16717 | FBgn0030895 | CG7135 | 0.164343021 | 0.2412154 | 0 |
| RH59970 | FBgn0038017 | CG4115 | -0.164341122 | 0.22992538 | 0 |
| RE14108 | FBgn0026619 | Taz | 0.164325014 | 0.17711358 | 0 |
| HL05936 | FBgn0024542 | Neos | 0.164292923 | 0.20738221 | 0 |
| LD14917 | FBgn0000317 | ck | 0.164275995 | 0.31198099 | 0 |
| RE03814 | FBgn0038454 | CG10324 | 0.164226405 | 0.17331563 | 0 |
| GM07807 | FBgn0260482 | CG32599 | 0.164213905 | 0.38251436 | 0 |
| GM04053 | FBgn0033265 | coil | 0.164204824 | 0.23984813 | 0 |
| LP02760 | FBgn0036643 | Syx8 | -0.16419687 | 0.30110704 | 0 |
| RE46906 | FBgn0030657 | cerv | 0.164161888 | 0.21322625 | 0 |
| LP02363 | FBgn0050394 | CG30394 | 0.164136616 | 0.19449538 | 0 |
| LD11531 | FBgn0004049 | yrt | 0.164106551 | 0.30364662 | 0 |
| RE38979 | FBgn0031494 | CG17219 | 0.164047476 | 0.33063114 | 0 |
| LD29301 | FBgn0026427 | Su(var)2-HP2 | 0.164023597 | 0.20979542 | 0 |
| LP01337 | FBgn0038114 | CG11670 | 0.16398902 | 0.16822308 | 0 |
| RH04006 | FBgn0004465 | Su(P) | -0.163863111 | 0.16476127 | 0 |
| LD31809 | FBgn0264695 | Mhc | 0.163769927 | 0.36971494 | 0 |
| LD10016 | FBgn0030060 | CG2004 | 0.16370008 | 0.22598212 | 0 |
| GH11117 | FBgn0039875 | sip3 | 0.163673843 | 0.26463547 | 0 |
| RE06583 | FBgn0051810 | CG31810 | 0.163664396 | 0.1888044 | 0 |
| RE06583 | FBgn0051809 | CG31809 | 0.163664396 | 0.1888044 | 0 |
| GH26351 | FBgn0036805 | Chmp1 | -0.163650695 | 0.20111941 | 0 |
| GH07919 | FBgn0010263 | Rbp9 | -0.163582996 | 0.4344621 | 0 |
| LD21643 | FBgn0086129 | snama | 0.163577818 | 0.26140631 | 0 |
| LD36162 | FBgn0001223 | Hsp22 | 0.163567835 | 0.17394337 | 0 |
| LD36162 | FBgn0001228 | CG4456 | 0.163567835 | 0.17394337 | 0 |
| GH03502 | FBgn0261285 | Ppcs | 0.163549096 | 0.24120973 | 0 |
| RE15191 | FBgn0028855 | CG15282 | -0.163394098 | 0.29410644 | 0 |
| LD10349 | FBgn0032479 | CG16974 | 0.163332167 | 0.31217459 | 0 |
| LD07740 | FBgn0260392 | CG42518 | 0.163308876 | 0.23774463 | 0 |
| LD07740 | FBgn0260401 | MED9 | 0.163308876 | 0.23774463 | 0 |
| SD02058 | FBgn0053062 | CG33062 | -0.163205057 | 0.30080363 | 0 |

| | | | | | |
|---|---|---|---|---|---|
| RH15319 | FBgn0045038 | Proc | -0.163162485 | 0.2071339 | 0 |
| GH19585 | FBgn0033781 | CG13319 | 0.163142188 | 0.21784977 | 0 |
| GH10531 | FBgn0000038 | nAChRbeta1 | -0.163013357 | 0.31964742 | 0 |
| SD10002 | FBgn0039136 | CG5902 | 0.163004932 | 0.19232116 | 0 |
| GM05329 | FBgn0000181 | bic | -0.162926712 | 0.26069911 | 0 |
| LD30319 | FBgn0033539 | Git | 0.162924041 | 0.18959139 | 0 |
| LD07939 | FBgn0039304 | CG10425 | 0.16286663 | 0.22082195 | 0 |
| LD14895 | FBgn0030434 | Brms1 | -0.16286528 | 0.3639518 | 0 |
| GH26663 | FBgn0052698 | CG32698 | 0.162782463 | 0.44596074 | 0 |
| GM14014 | FBgn0263987 | spoon | 0.162777825 | 0.1840059 | 0 |
| RE54096 | FBgn0037188 | CG7369 | -0.162770774 | 0.27527304 | 0 |
| RE31520 | FBgn0037550 | CG9667 | 0.162737645 | 0.29543016 | 0 |
| RH10862 | FBgn0035534 | mRpS6 | 0.162675146 | 0.22376316 | 0 |
| GM04590 | FBgn0266390 | lncRNA:CR45030 | 0.162658278 | 0.30138677 | 0 |
| GH07383 | FBgn0039210 | CG13625 | 0.162581639 | 0.34349101 | 0 |
| RE49860 | FBgn0033543 | CG12338 | 0.162315809 | 0.39205385 | 0 |
| GH23407 | FBgn0028542 | NimB4 | 0.162314146 | 0.25748944 | 0 |
| AT13107 | FBgn0032089 | Rcd-1r | 0.162191494 | 0.37187434 | 0 |
| AT12780 | FBgn0035069 | CG3611 | 0.16208913 | 0.17264646 | 0 |
| AT09438 | FBgn0034237 | eIF3b | 0.161958609 | 0.21524805 | 0 |
| GM13460 | FBgn0259113 | DNApol-alpha180 | -0.161894042 | 0.37459627 | 0 |
| RE09339 | FBgn0035089 | Phk-3 | -0.16184817 | 0.56275719 | 0 |
| RE21944 | FBgn0037124 | CG14568 | -0.161775566 | 0.36885296 | 0 |
| LD46714 | FBgn0035876 | Pex2 | 0.161774165 | 0.29009021 | 0 |
| LD24048 | FBgn0017457 | U2af38 | 0.161760671 | 0.20152257 | 0 |
| LD01910 | FBgn0013762 | Cdk5 | -0.161686454 | 0.2012105 | 0 |
| LD23102 | FBgn0270924 | zf30C | 0.161670382 | 0.29328557 | 0 |
| GH12581 | FBgn0020415 | Idgf2 | 0.161651144 | 0.19136517 | 0 |
| AT18808 | FBgn0031045 | CG14212 | 0.161644984 | 0.3271688 | 0 |
| LD46284 | FBgn0039858 | CycG | 0.161639331 | 0.1887837 | 0 |
| GM08271 | FBgn0038876 | CG5919 | 0.161619723 | 0.19717375 | 0 |
| LD36616 | FBgn0030930 | Pgant7 | 0.161614999 | 0.1839008 | 0 |
| LD39912 | FBgn0038381 | EndoU | 0.161537532 | 0.24907872 | 0 |
| GH16917 | FBgn0260964 | Vmat | 0.161496188 | 0.40479015 | 0 |
| LD17077 | FBgn0031284 | PGAP2 | 0.161485976 | 0.21524805 | 0 |
| LP10790 | FBgn0263110 | CG43367 | 0.161471663 | 0.18725155 | 0 |
| SD02419 | FBgn0051125 | CG31125 | 0.161457932 | 0.1839008 | 0 |
| RE24170 | FBgn0001108 | DCTN1-p150 | -0.161456339 | 0.17394337 | 0 |
| RE70727 | FBgn0264785 | Hph | -0.16144595 | 0.2323332 | 0 |
| LD16285 | FBgn0010622 | DCTN3-p24 | 0.161314342 | 0.23606201 | 0 |
| SD20954 | FBgn0034126 | jtb | -0.161307369 | 0.31206199 | 0 |
| GM01181 | FBgn0086691 | UK114 | -0.161271804 | 0.25270491 | 0 |
| RE47242 | FBgn0037882 | CG17187 | -0.161200787 | 0.2430466 | 0 |
| AT03259 | FBgn0038488 | m-cup | 0.161116008 | 0.17877971 | 0 |
| LD01372 | FBgn0267390 | dop | 0.161019089 | 0.18075619 | 0 |
| RE57644 | FBgn0039944 | CG17162 | -0.160975507 | 0.21796586 | 0 |
| LD21614 | FBgn0265574 | Cdc5 | 0.160954436 | 0.17917133 | 0 |
| RE55321 | FBgn0003357 | Jon99Ciii | -0.160953705 | 0.18075619 | 0 |

| | | | | | |
|---|---|---|---|---|---|
| RE55321 | FBgn0003356 | Jon99Cii | -0.160953705 | 0.18075619 | 0 |
| RE55321 | FBgn0039778 | Jon99Fi | -0.160953705 | 0.18075619 | 0 |
| RE55321 | FBgn0039777 | Jon99Fii | -0.160953705 | 0.18075619 | 0 |
| RH33950 | FBgn0037200 | CG11109 | -0.160934081 | 0.44351826 | 0 |
| LD24527 | FBgn0037555 | Ada2b | 0.160873277 | 0.23140423 | 0 |
| LD37137 | FBgn0034372 | Gint3 | 0.160872935 | 0.21070801 | 0 |
| LD37137 | FBgn0259202 | CG42306 | 0.160872935 | 0.21070801 | 0 |
| GH08349 | FBgn0283500 | Sac1 | -0.160869221 | 0.56024256 | 0 |
| LP12231 | FBgn0053128 | CG33128 | 0.160785834 | 0.18458646 | 0 |
| RE05533 | FBgn0033368 | CG13743 | -0.160785313 | 0.29408612 | 0 |
| RE36442 | FBgn0030332 | CG9360 | 0.160782317 | 0.31688128 | 0 |
| SD03358 | FBgn0262937 | Rabex-5 | -0.160744466 | 0.19133917 | 0 |
| GH11263 | FBgn0024248 | chico | 0.160740565 | 0.20983362 | 0 |
| HL07808 | FBgn0040765 | luna | -0.160710259 | 0.4047529 | 0 |
| GM27257 | FBgn0262730 | dtn | -0.160706149 | 0.24733094 | 0 |
| LD27791 | FBgn0086711 | mol | 0.160687282 | 0.17722118 | 0 |
| LD02310 | FBgn0035983 | CG4080 | 0.160680159 | 0.20693481 | 0 |
| LD15203 | FBgn0026181 | Rok | 0.160662845 | 0.28911598 | 0 |
| RE09173 | FBgn0032642 | CG5110 | -0.160580296 | 0.19655822 | 0 |
| LD21667 | FBgn0027334 | l(1)G0004 | 0.160522227 | 0.24657701 | 0 |
| LD24308 | FBgn0027280 | l(1)G0193 | 0.160519152 | 0.17551784 | 0 |
| GM02889 | FBgn0038763 | PIG-L | -0.160468773 | 0.20119161 | 0 |
| LD15641 | FBgn0004378 | Klp61F | 0.160459359 | 0.28878272 | 0 |
| SD05495 | FBgn0031886 | Nuf2 | 0.160453424 | 0.20973394 | 0 |
| LP03545 | FBgn0039027 | CG7031 | -0.160442484 | 0.47568595 | 0 |
| GH05949 | FBgn0032032 | CG17294 | -0.160330799 | 0.29080958 | 0 |
| GM18959 | FBgn0038787 | CG4360 | -0.160326864 | 0.3283412 | 0 |
| LD23542 | FBgn0086361 | alph | 0.160323278 | 0.2276499 | 0 |
| LD14510 | FBgn0010292 | bys | 0.160307588 | 0.23661875 | 0 |
| SD05512 | FBgn0032036 | CG13384 | 0.16028108 | 0.18167614 | 0 |
| RH51452 | FBgn0034576 | ND-B14.7 | 0.160219762 | 0.22060168 | 0 |
| RE36634 | FBgn0035523 | Ctl1 | 0.160040433 | 0.17463993 | 0 |
| GM01561 | FBgn0052479 | Usp10 | 0.159998768 | 0.32403871 | 0 |
| RE07751 | FBgn0046687 | Tre1 | -0.159966854 | 0.23428979 | 0 |
| GH17155 | FBgn0034925 | CG5339 | -0.159950888 | 0.22695432 | 0 |
| RE14544 | FBgn0259209 | Mlp60A | -0.159931146 | 0.8194291 | 0 |
| RE66338 | FBgn0050085 | Rif1 | 0.159875505 | 0.27309497 | 0 |
| SD05186 | FBgn0087013 | Karybeta3 | 0.159741412 | 0.17646871 | 0 |
| RH08077 | FBgn0042641 | frc | -0.159680614 | 0.29392156 | 0 |
| RH08077 | FBgn0052174 | CG32174 | -0.159680614 | 0.29392156 | 0 |
| LD11247 | FBgn0052484 | Sk2 | 0.159658071 | 0.21137971 | 0 |
| RH06643 | FBgn0261599 | RpS29 | -0.159597196 | 0.18212145 | 0 |
| LD24649 | FBgn0034908 | CG5543 | 0.15956617 | 0.18449918 | 0 |
| SD23356 | FBgn0000579 | Eno | -0.159448454 | 0.17793042 | 0 |
| GH07464 | FBgn0043576 | PGRP-SC1a | -0.159349423 | 0.30470968 | 0 |
| GH07464 | FBgn0033327 | PGRP-SC1b | -0.159349423 | 0.30470968 | 0 |
| RE46539 | FBgn0039381 | SppL | -0.159241435 | 0.17464104 | 0 |
| LD24373 | FBgn0263933 | ebi | 0.159238959 | 0.21567639 | 0 |

| | | | | | |
|---|---|---|---|---|---|
| RE57845 | FBgn0016032 | lbm | 0.159231899 | 0.63373047 | 0 |
| LD17911 | FBgn0024984 | CG3457 | 0.159230385 | 0.23170768 | 0 |
| RE70055 | FBgn0024326 | Mkk4 | -0.159164143 | 0.23791734 | 0 |
| LP08452 | FBgn0259163 | CG42268 | 0.159150627 | 0.24986172 | 0 |
| RE35348 | FBgn0034392 | MFS15 | -0.158991002 | 0.25948197 | 0 |
| GH18178 | FBgn0032505 | CG16826 | -0.158981014 | 0.21137971 | 0 |
| LD14246 | FBgn0011202 | dia | 0.158942734 | 0.24333048 | 0 |
| LD22201 | FBgn0001104 | Galphai | 0.158933191 | 0.17551784 | 0 |
| RH56553 | FBgn0003714 | tko | -0.158924821 | 0.18882961 | 0 |
| RH04903 | FBgn0005593 | RpL7 | 0.158820999 | 0.3085804 | 0 |
| GH20665 | FBgn0260935 | Vps15 | -0.158813518 | 0.46525228 | 0 |
| GH18886 | FBgn0035032 | ATPsynF | -0.158754069 | 0.17859077 | 0 |
| GH07769 | FBgn0261842 | pre-mod(mdg4)-AE | 0.158609194 | 0.20050793 | 0 |
| GH07769 | FBgn0261844 | pre-mod(mdg4)-V | 0.158609194 | 0.20050793 | 0 |
| GH07769 | FBgn0261845 | pre-mod(mdg4)-U | 0.158609194 | 0.20050793 | 0 |
| GH03529 | FBgn0027594 | drpr | -0.158582745 | 0.19554083 | 0 |
| LD22412 | FBgn0035357 | MEP-1 | 0.158453293 | 0.17923407 | 0 |
| LD22412 | FBgn0250911 | CG42245 | 0.158453293 | 0.17923407 | 0 |
| RE49866 | FBgn0023514 | CG14805 | 0.158293077 | 0.20279766 | 0 |
| LD39166 | FBgn0038755 | Hs6st | 0.158270494 | 0.17530806 | 0 |
| LD12892 | FBgn0032586 | Tpr2 | 0.158259402 | 0.17529678 | 0 |
| AT14909 | FBgn0037777 | CG11722 | -0.158257939 | 0.19232116 | 0 |
| GH22929 | FBgn0034136 | DAT | -0.15823488 | 0.29835009 | 0 |
| LD44220 | FBgn0051223 | CG31223 | 0.158231365 | 0.17819436 | 0 |
| GM08240 | FBgn0052473 | CG32473 | 0.158220945 | 0.24852054 | 0 |
| LD37765 | FBgn0024991 | CG2694 | -0.158187404 | 0.17551784 | 0 |
| HL01508 | FBgn0261108 | Atg13 | 0.158153805 | 0.21689011 | 0 |
| LD29276 | FBgn0038235 | CG8461 | 0.1581453 | 0.2781605 | 0 |
| SD05444 | FBgn0027358 | Tim9b | -0.157969896 | 0.18115038 | 0 |
| SD05444 | FBgn0031041 | CG12788 | -0.157969896 | 0.18115038 | 0 |
| LD31229 | FBgn0037687 | CG8132 | 0.157938001 | 0.22736386 | 0 |
| GH22016 | FBgn0032858 | CG10949 | -0.157927723 | 0.27417488 | 0 |
| LD36006 | FBgn0037336 | CG2519 | 0.157893223 | 0.24556656 | 0 |
| LP05803 | FBgn0037363 | Atg17 | 0.157872077 | 0.20884829 | 0 |
| GH07506 | FBgn0034914 | CG5554 | 0.157858281 | 0.18165 | 0 |
| LD38682 | FBgn0034569 | dgt3 | -0.157845946 | 0.18265457 | 0 |
| LD04107 | FBgn0043884 | mask | 0.157824366 | 0.24519727 | 0 |
| RE12587 | FBgn0030390 | CG15731 | -0.157746142 | 0.29950032 | 0 |
| LD24139 | FBgn0040279 | Osi14 | -0.157624803 | 0.52200292 | 0 |
| GM12157 | FBgn0263346 | CG43427 | 0.157612168 | 0.1762235 | 0 |
| LD28119 | FBgn0037911 | CG10898 | -0.157564833 | 0.26152324 | 0 |
| RH21989 | FBgn0028703 | Nhe3 | -0.157546859 | 0.23621839 | 0 |
| RE70685 | FBgn0051660 | smog | -0.157506724 | 0.29671934 | 0 |
| RE26705 | FBgn0038511 | cysu | 0.157384383 | 0.5354438 | 0 |
| LD15209 | FBgn0029873 | CG3918 | 0.157295436 | 0.2481764 | 0 |
| GH11964 | FBgn0032010 | CG8086 | -0.157242561 | 0.41609944 | 0 |
| SD05277 | FBgn0262866 | S6kII | -0.157127926 | 0.36813914 | 0 |
| RE09708 | FBgn0034437 | CG10051 | -0.15706174 | 0.32095126 | 0 |

| | | | | | |
|---|---|---|---|---|---|
| RH45455 | FBgn0030432 | CG4404 | -0.157054614 | 0.21109531 | 0 |
| LD07457 | FBgn0036305 | CG10984 | 0.157030646 | 0.17889351 | 0 |
| RE11317 | FBgn0052982 | CG32982 | 0.156933293 | 0.17859077 | 0 |
| GH12692 | FBgn0030164 | CG1889 | -0.156884505 | 0.19959552 | 0 |
| LD21748 | FBgn0035993 | Nf-YA | -0.156804358 | 0.32656161 | 0 |
| LP04564 | FBgn0051694 | CG31694 | -0.156780509 | 0.24202703 | 0 |
| GH20219 | FBgn0004618 | gl | -0.156773406 | 0.23434738 | 0 |
| LD15480 | FBgn0024365 | CG12773 | 0.156719633 | 0.2191356 | 0 |
| GH08448 | FBgn0038109 | CG11656 | 0.156687544 | 0.21693903 | 0 |
| GH04440 | FBgn0033376 | CG8777 | 0.1566736 | 0.2282607 | 0 |
| GH19655 | FBgn0030247 | CG15208 | 0.156661445 | 0.33517148 | 0 |
| HL05773 | FBgn0267252 | Ggamma30A | -0.156616806 | 0.49855608 | 0 |
| LD23835 | FBgn0263855 | BubR1 | 0.156594995 | 0.32345633 | 0 |
| LD24865 | FBgn0035421 | CG12034 | 0.156594001 | 0.29328557 | 0 |
| GH04034 | FBgn0037466 | CG1965 | 0.15653756 | 0.19203472 | 0 |
| AT11516 | FBgn0038281 | RpL10Aa | -0.156480676 | 0.35771172 | 0 |
| RH47633 | FBgn0034141 | CG8311 | -0.156425081 | 0.27904632 | 0 |
| GH18819 | FBgn0042712 | HBS1 | 0.156328069 | 0.25250442 | 0 |
| SD22530 | FBgn0030611 | CG15027 | 0.156298335 | 0.20678417 | 0 |
| GH11341 | FBgn0028690 | Rpn5 | -0.156212633 | 0.19232116 | 0 |
| LD29995 | FBgn0026722 | drosha | 0.156110281 | 0.19593884 | 0 |
| RE26454 | FBgn0010453 | Wnt4 | -0.15607234 | 0.4373929 | 0 |
| LD36304 | FBgn0000166 | bcd | 0.156049677 | 0.24806598 | 0 |
| LD44770 | FBgn0032348 | CG4751 | 0.156046363 | 0.18204988 | 0 |
| AT06839 | FBgn0031288 | CG13949 | 0.155961936 | 0.50825782 | 0 |
| LD24353 | FBgn0030891 | Ada3 | 0.155912005 | 0.21285991 | 0 |
| GH19118 | FBgn0262007 | CG42825 | -0.155903009 | 0.23984813 | 0 |
| GH23026 | FBgn0037623 | CG9801 | -0.155880653 | 0.31391251 | 0 |
| LD10524 | FBgn0034528 | CG11180 | 0.155847786 | 0.32355904 | 0 |
| LD18126 | FBgn0023536 | CG3156 | 0.15583647 | 0.26040573 | 0 |
| LD21589 | FBgn0010825 | Gug | 0.155832579 | 0.26188951 | 0 |
| RE54684 | FBgn0050438 | CG30438 | 0.155716184 | 0.25821219 | 0 |
| LD02491 | FBgn0036257 | RhoGAP68F | -0.155659532 | 0.18466997 | 0 |
| GH21728 | FBgn0034258 | eIF3c | 0.155657186 | 0.23906935 | 0 |
| AT19988 | FBgn0039503 | CG14262 | 0.155643265 | 0.26610197 | 0 |
| RE02747 | FBgn0038412 | Zip89B | -0.155608529 | 0.34509821 | 0 |
| RE64188 | FBgn0003023 | otu | -0.155598617 | 0.20762985 | 0 |
| LD08354 | FBgn0039183 | Dis3 | 0.155591641 | 0.19216264 | 0 |
| GH27909 | FBgn0002921 | Atpalpha | -0.155549496 | 0.1888044 | 0 |
| GH27909 | FBgn0038145 | Droj2 | -0.155549496 | 0.1888044 | 0 |
| RE16005 | FBgn0036112 | CG14147 | 0.155521725 | 0.30993577 | 0 |
| LD24810 | FBgn0039140 | Miro | 0.155502596 | 0.21117523 | 0 |
| LP01044 | FBgn0029807 | CG3108 | -0.1555023 | 0.30364662 | 0 |
| RH63812 | FBgn0026872 | CG14777 | -0.155495734 | 0.20884829 | 0 |
| LD36407 | FBgn0035227 | Iml1 | 0.155489538 | 0.22082195 | 0 |
| GH02833 | FBgn0027602 | CG8611 | 0.155488346 | 0.24657701 | 0 |
| GH16395 | FBgn0030749 | AnxB11 | 0.1554017 | 0.22543836 | 0 |
| GH08902 | FBgn0043783 | CG32444 | -0.155388404 | 0.28473292 | 0 |

| | | | | | |
|---|---|---|---|---|---|
| GH01240 | FBgn0004242 | Syt1 | -0.155360522 | 0.3547438 | 0 |
| LD22320 | FBgn0027509 | TBCD | -0.1553395 | 0.21845712 | 0 |
| RE68558 | FBgn0051999 | CG31999 | 0.155321639 | 0.26449625 | 0 |
| GH08429 | FBgn0036627 | Gagr | -0.155317945 | 0.24907872 | 0 |
| RE08669 | FBgn0039857 | RpL6 | 0.155236553 | 0.40340611 | 0 |
| GH12726 | FBgn0001296 | kar | 0.155191304 | 0.2219358 | 0 |
| LD27042 | FBgn0037008 | mTerf3 | -0.155124654 | 0.46004273 | 0 |
| RH07835 | FBgn0013348 | TpnC41C | -0.155094039 | 0.29835009 | 0 |
| LD17594 | FBgn0034708 | Vps35 | 0.154989219 | 0.18206528 | 0 |
| LD12844 | FBgn0262511 | Vha44 | -0.154916955 | 0.21117523 | 0 |
| GH14951 | FBgn0051038 | CG31038 | 0.154798627 | 0.37651512 | 0 |
| RE53508 | FBgn0011787 | mRpL12 | -0.154783146 | 0.25582587 | 0 |
| LD36256 | FBgn0011290 | Taf12 | 0.154737264 | 0.2378597 | 0 |
| AT14266 | FBgn0263453 | asRNA:CR43476 | -0.154697731 | 0.20380583 | 0 |
| RE49709 | FBgn0062442 | Cisd2 | 0.154671235 | 0.21487795 | 0 |
| RE38475 | FBgn0052816 | CG32816 | 0.154607355 | 0.19025194 | 0 |
| LD14901 | FBgn0035397 | CG11486 | 0.154600695 | 0.33497579 | 0 |
| GH02863 | FBgn0033093 | CG3270 | -0.154580768 | 0.19635709 | 0 |
| LD46269 | FBgn0032925 | CG9246 | 0.154531559 | 0.24413419 | 0 |
| GM21396 | FBgn0037569 | tex | 0.154498799 | 0.21267823 | 0 |
| RE58316 | FBgn0015011 | AhcyL2 | -0.154445699 | 0.27904632 | 0 |
| RE65257 | FBgn0024887 | kin17 | 0.154419897 | 0.29545439 | 0 |
| GH11008 | FBgn0036713 | Mip | -0.154407847 | 0.2412154 | 0 |
| RE08455 | FBgn0034733 | CG4752 | 0.154361623 | 0.1975511 | 0 |
| GH04962 | FBgn0027588 | GCS2alpha | 0.154317563 | 0.35471777 | 0 |
| GH12385 | FBgn0030926 | psh | -0.154315828 | 0.21644222 | 0 |
| RE55033 | FBgn0038649 | CG7718 | -0.1542697 | 0.18882961 | 0 |
| LD45581 | FBgn0015903 | apt | 0.154253355 | 0.19171028 | 0 |
| LD11139 | FBgn0039244 | CG11069 | 0.154193691 | 0.57219448 | 0 |
| GH06241 | FBgn0010053 | Jheh1 | 0.154147516 | 0.3162624 | 0 |
| GM03763 | FBgn0025394 | inc | 0.154147097 | 0.31217459 | 0 |
| AT20876 | FBgn0038957 | CG7059 | 0.15409631 | 0.35619541 | 0 |
| LD27166 | FBgn0027086 | IleRS | 0.154027395 | 0.19876252 | 0 |
| LP04014 | FBgn0038113 | CG11668 | 0.153966158 | 0.23613029 | 0 |
| RE51574 | FBgn0000565 | Eip71CD | 0.153962587 | 0.43555986 | 0 |
| LD29352 | FBgn0030625 | CG5877 | 0.153926275 | 0.20179693 | 0 |
| GH28534 | FBgn0035083 | Tina-1 | -0.153887962 | 0.1911631 | 0 |
| LD33329 | FBgn0039350 | jigr1 | 0.153887167 | 0.2088032 | 0 |
| GH25972 | FBgn0036222 | SdhAL | -0.153683297 | 0.19028495 | 0 |
| LD47277 | FBgn0037185 | CG11367 | 0.15368151 | 0.27904632 | 0 |
| LD05287 | FBgn0002441 | l(3)mbt | 0.153502373 | 0.2323332 | 0 |
| LD01127 | FBgn0000253 | Cam | -0.153477388 | 0.37664429 | 0 |
| LD33081 | FBgn0002715 | mei-S332 | 0.153450085 | 0.19904473 | 0 |
| GH21295 | FBgn0266377 | Pde8 | -0.153431907 | 0.37936225 | 0 |
| AT21479 | FBgn0036706 | ND-24L | 0.153427021 | 0.45351921 | 0 |
| RE28669 | FBgn0260439 | Pp2A-29B | 0.153424005 | 0.22734671 | 0 |
| SD05956 | FBgn0025630 | Rtca | -0.153417934 | 0.23330886 | 0 |
| AT10359 | FBgn0029603 | CG14053 | 0.153415804 | 0.5313111 | 0 |

| | | | | | |
|---|---|---|---|---|---|
| RE65105 | FBgn0033978 | Cyp6a23 | -0.153393862 | 0.30080363 | 0 |
| LD14049 | FBgn0262601 | SmB | 0.153304107 | 0.29914002 | 0 |
| HL08191 | FBgn0038925 | Cchl | -0.153285369 | 0.23774463 | 0 |
| LD09823 | FBgn0001341 | Mybbp1A | 0.153250133 | 0.35877075 | 0 |
| LD39271 | FBgn0032597 | CG17904 | 0.153152472 | 0.19571234 | 0 |
| SD07737 | FBgn0032633 | Lrch | -0.153146557 | 0.27581094 | 0 |
| LD23052 | FBgn0039709 | Cad99C | 0.153092312 | 0.307972 | 0 |
| LD32258 | FBgn0051643 | CG31643 | 0.153055703 | 0.21548782 | 0 |
| RH61716 | FBgn0024989 | CG3777 | -0.152913127 | 0.29578352 | 0 |
| LD43171 | FBgn0052564 | CG32564 | 0.152826077 | 0.27113412 | 0 |
| RE05963 | FBgn0039481 | Cpr97Eb | -0.152817863 | 0.19788722 | 0 |
| LD10447 | FBgn0052418 | vito | 0.15279126 | 0.24120973 | 0 |
| GH10154 | FBgn0004622 | TkR99D | -0.152761607 | 0.21781804 | 0 |
| LD28680 | FBgn0051898 | CG31898 | -0.152706518 | 0.31110387 | 0 |
| GM03875 | FBgn0051635 | CG31635 | 0.152688844 | 0.31217459 | 0 |
| LP06586 | FBgn0085311 | CG34282 | 0.152684024 | 0.21862256 | 0 |
| LD47441 | FBgn0014861 | Mcm2 | -0.152619439 | 0.20513188 | 0 |
| GH08442 | FBgn0050069 | CG30069 | -0.15261265 | 0.42344497 | 0 |
| LD19518 | FBgn0038013 | CG10038 | 0.15253591 | 0.26687802 | 0 |
| GH12558 | FBgn0028479 | Mtpalpha | 0.152492114 | 0.40340611 | 0 |
| LD11580 | FBgn0034496 | CG9143 | 0.152429647 | 0.24910111 | 0 |
| AT02774 | FBgn0039850 | CG11333 | 0.152412988 | 0.60237747 | 0 |
| GH18625 | FBgn0029874 | CG3342 | 0.152387394 | 0.21014705 | 0 |
| LP07253 | FBgn0034824 | Klp59C | -0.152377009 | 0.24305287 | 0 |
| LD45447 | FBgn0039601 | CG1523 | 0.152325238 | 0.38507121 | 0 |
| GH14591 | FBgn0261934 | dikar | -0.152172096 | 0.43822876 | 0 |
| GM21394 | FBgn0042094 | Adk3 | -0.152168917 | 0.24360361 | 0 |
| SD09147 | FBgn0034579 | mRpL54 | -0.152156871 | 0.23170768 | 0 |
| LD31238 | FBgn0025627 | CG4194 | 0.152135036 | 0.66044765 | 0 |
| GH03554 | FBgn0266917 | Sf3a1 | 0.152114275 | 0.24963632 | 0 |
| AT15262 | FBgn0040364 | CG11378 | -0.15209254 | 0.27650528 | 0 |
| SD02219 | FBgn0034958 | CG3907 | 0.152079342 | 0.2323332 | 0 |
| RH54455 | FBgn0265778 | PDZ-GEF | 0.15203057 | 0.20102435 | 0 |
| LD44591 | FBgn0022382 | Pka-R2 | -0.151994433 | 0.56127049 | 0 |
| LD09351 | FBgn0038129 | TBC1D5 | 0.151992279 | 0.19609219 | 0 |
| LD24274 | FBgn0036816 | Indy | 0.151872683 | 0.23292417 | 0 |
| GM31840 | FBgn0053236 | Ste:CG33236 | 0.151860222 | 0.32314931 | 0 |
| GM31840 | FBgn0053237 | Ste:CG33237 | 0.151860222 | 0.32314931 | 0 |
| GM31840 | FBgn0053238 | Ste:CG33238 | 0.151860222 | 0.32314931 | 0 |
| GM31840 | FBgn0053239 | Ste:CG33239 | 0.151860222 | 0.32314931 | 0 |
| GM31840 | FBgn0053240 | Ste:CG33240 | 0.151860222 | 0.32314931 | 0 |
| GM31840 | FBgn0053241 | Ste:CG33241 | 0.151860222 | 0.32314931 | 0 |
| GM31840 | FBgn0053242 | Ste:CG33242 | 0.151860222 | 0.32314931 | 0 |
| GM31840 | FBgn0053243 | Ste:CG33243 | 0.151860222 | 0.32314931 | 0 |
| GM31840 | FBgn0053244 | Ste:CG33244 | 0.151860222 | 0.32314931 | 0 |
| GM31840 | FBgn0053245 | Ste:CG33245 | 0.151860222 | 0.32314931 | 0 |
| GM31840 | FBgn0053246 | Ste:CG33246 | 0.151860222 | 0.32314931 | 0 |
| GM31840 | FBgn0053247 | Ste:CG33247 | 0.151860222 | 0.32314931 | 0 |

| | | | | | |
|---|---|---|---|---|---|
| GM31840 | FBgn0259817 | SteXh:CG42398 | 0.151860222 | 0.32314931 | 0 |
| GM31840 | FBgn0259837 | Su(Ste):CR42406 | 0.151860222 | 0.32314931 | 0 |
| GM31840 | FBgn0259838 | Su(Ste):CR42407 | 0.151860222 | 0.32314931 | 0 |
| GM31840 | FBgn0044817 | Ste12DOR | 0.151860222 | 0.32314931 | 0 |
| GM31840 | FBgn0259850 | Su(Ste):CR42419 | 0.151860222 | 0.32314931 | 0 |
| GM31840 | FBgn0267446 | Su(Ste):CR45796 | 0.151860222 | 0.32314931 | 0 |
| RE49852 | FBgn0035335 | mRpL23 | -0.151828519 | 0.30918864 | 0 |
| SD01663 | FBgn0001114 | Glt | 0.151794919 | 0.54577962 | 0 |
| RE73641 | FBgn0037323 | CG2663 | 0.151776716 | 0.29658578 | 0 |
| LD24239 | FBgn0030063 | CG1789 | 0.151716131 | 0.19822213 | 0 |
| RE57183 | FBgn0052029 | Cpr66D | -0.151637143 | 0.44707004 | 0 |
| LD23703 | FBgn0014879 | Set | 0.151586537 | 0.26453289 | 0 |
| GM13382 | FBgn0053554 | Nipped-A | -0.151585343 | 0.42676291 | 0 |
| LD25702 | FBgn0035644 | DNApol-epsilon58 | -0.151553528 | 0.35294946 | 0 |
| LD45688 | FBgn0035831 | CG8038 | -0.151546076 | 0.27610621 | 0 |
| HL08149 | FBgn0005391 | Yp2 | -0.15141966 | 0.22199756 | 0 |
| LD24870 | FBgn0030658 | CG7872 | 0.151396062 | 0.220445 | 0 |
| LD16414 | FBgn0035539 | slow | 0.151375273 | 0.43154461 | 0 |
| GH03360 | FBgn0030051 | spirit | -0.151302806 | 0.41332137 | 0 |
| LD19715 | FBgn0037224 | TwdlF | 0.151284107 | 0.28679696 | 0 |
| AT17975 | FBgn0022709 | Adk1 | 0.151195648 | 0.22533232 | 0 |
| RH74410 | FBgn0037102 | CRIF | 0.151149083 | 0.22529804 | 0 |
| GH10341 | FBgn0085447 | sif | 0.151077239 | 0.27904632 | 0 |
| LD43174 | FBgn0261987 | Pxt | -0.15087777 | 0.23418232 | 0 |
| SD18780 | FBgn0034919 | CG5569 | -0.15084184 | 0.41897567 | 0 |
| AT08504 | FBgn0261570 | CG42684 | -0.150841699 | 0.38913924 | 0 |
| GH10964 | FBgn0027795 | CG14785 | 0.150736683 | 0.20102435 | 0 |
| AT09114 | FBgn0037759 | CG8526 | -0.150699948 | 0.5354438 | 0 |
| AT27340 | FBgn0033952 | Adgf-E | -0.150674318 | 0.38135654 | 0 |
| LD13710 | FBgn0032848 | nesd | -0.150647872 | 0.41052494 | 0 |
| SD27876 | FBgn0035482 | CG14985 | 0.150632259 | 0.22830867 | 0 |
| SD12335 | FBgn0052251 | Claspin | -0.150544356 | 0.25998362 | 0 |
| AT25667 | FBgn0032471 | CG5122 | -0.150535474 | 0.28341838 | 0 |
| LD17515 | FBgn0011829 | Ret | 0.150390922 | 0.21845712 | 0 |
| LD41905 | FBgn0033926 | Arc1 | 0.150351101 | 0.3546485 | 0 |
| LD41905 | FBgn0033927 | CR10102 | 0.150351101 | 0.3546485 | 0 |
| GH02439 | FBgn0027608 | CG2082 | -0.150127743 | 0.2990465 | 0 |
| RH01665 | FBgn0034162 | CG6426 | -0.1500233 | 0.26397981 | 0 |
| LD21213 | FBgn0261596 | RpS24 | 0.150021715 | 0.22672346 | 0 |
| RH08687 | FBgn0040074 | retinin | -0.150013716 | 0.48214468 | 0 |
| LD10743 | FBgn0040372 | G9a | -0.14987817 | 0.20687266 | 0 |
| GH01650 | FBgn0086450 | su(r) | -0.149861552 | 0.21683387 | 0 |
| LP06735 | FBgn0034416 | CG15109 | -0.149831871 | 0.60251362 | 0 |
| LP08709 | FBgn0039476 | CG6271 | -0.149830369 | 0.19822213 | 0 |
| LP08709 | FBgn0039475 | CG6277 | -0.149830369 | 0.19822213 | 0 |
| LD43519 | FBgn0040239 | bc10 | -0.149787535 | 0.37533608 | 0 |
| RH55627 | FBgn0266137 | Dop1R2 | 0.149784547 | 0.19648812 | 0 |
| RE39205 | FBgn0037015 | cmpy | -0.149742285 | 0.29178686 | 0 |

| | | | | | |
|---|---|---|---|---|---|
| SD08108 | FBgn0033244 | CG8726 | 0.149732965 | 0.19232116 | 0 |
| RH25931 | FBgn0040653 | IM4 | -0.149698165 | 0.23434738 | 0 |
| LD35896 | FBgn0261532 | cdm | 0.149566116 | 0.22495288 | 0 |
| LD22376 | FBgn0029737 | CG6903 | -0.149556245 | 0.24981226 | 0 |
| GM16138 | FBgn0262116 | RNASEK | -0.14947779 | 0.25949969 | 0 |
| SD07188 | FBgn0039338 | XNP | 0.149456219 | 0.20461703 | 0 |
| AT20544 | FBgn0001233 | Hsp83 | -0.14937822 | 0.20142104 | 0 |
| LD21610 | FBgn0031876 | Atac1 | 0.149303735 | 0.30138677 | 0 |
| RE74673 | FBgn0015320 | Ubc2 | 0.149245327 | 0.2088032 | 0 |
| RE13370 | FBgn0037181 | CG11370 | 0.149241461 | 0.23778979 | 0 |
| RE45514 | FBgn0038432 | CG14883 | -0.149226707 | 0.24417029 | 0 |
| RE57443 | FBgn0036101 | NijA | 0.149195187 | 0.37690504 | 0 |
| LD24566 | FBgn0024973 | CG2701 | 0.149150187 | 0.25016484 | 0 |
| LP05366 | FBgn0004584 | Rrp1 | -0.149125731 | 0.20379355 | 0 |
| GH05731 | FBgn0030159 | CG9689 | 0.149096858 | 0.53118114 | 0 |
| GH07367 | FBgn0038806 | CG5412 | -0.149046612 | 0.19635709 | 0 |
| RH39293 | FBgn0040299 | Myo28B1 | -0.148832285 | 0.24946171 | 0 |
| AT27896 | FBgn0052396 | CG32396 | 0.148802438 | 0.24754354 | 0 |
| GH11495 | FBgn0267790 | rump | 0.148689557 | 0.20923367 | 0 |
| RE23260 | FBgn0035827 | Srp9 | -0.148655548 | 0.27904632 | 0 |
| RE21614 | FBgn0004569 | aos | 0.148648357 | 0.39226166 | 0 |
| RE71517 | FBgn0261555 | CG42673 | -0.14864578 | 0.40367739 | 0 |
| RE31658 | FBgn0040066 | wds | 0.148511957 | 0.19968924 | 0 |
| RH51511 | FBgn0030467 | CG1764 | -0.148500852 | 0.2677426 | 0 |
| GH14710 | FBgn0260659 | CG42542 | -0.148488933 | 0.22607606 | 0 |
| LD38910 | FBgn0034086 | CG8441 | 0.148473466 | 0.24946171 | 0 |
| LD45221 | FBgn0261786 | mi | 0.148448073 | 0.21683387 | 0 |
| RE32325 | FBgn0038733 | CG11407 | -0.148442105 | 0.26457886 | 0 |
| RE32325 | FBgn0038731 | CG11659 | -0.148442105 | 0.26457886 | 0 |
| RE32325 | FBgn0038730 | CG6300 | -0.148442105 | 0.26457886 | 0 |
| LD09868 | FBgn0035986 | CG4022 | 0.148422632 | 0.30098776 | 0 |
| GH01813 | FBgn0036141 | wls | 0.148362656 | 0.23906935 | 0 |
| LD33138 | FBgn0036920 | CG8004 | 0.148360058 | 0.2287555 | 0 |
| GH02649 | FBgn0035855 | CG7366 | -0.14835473 | 0.27902723 | 0 |
| RE16905 | FBgn0031878 | sip2 | 0.148337403 | 0.29410644 | 0 |
| LD37139 | FBgn0026400 | Noa36 | 0.14833041 | 0.22397302 | 0 |
| LP11673 | FBgn0026869 | Thd1 | 0.14827407 | 0.21783142 | 0 |
| GM01081 | FBgn0032690 | CG10333 | 0.148230336 | 0.29080958 | 0 |
| RE38953 | FBgn0031452 | Cwc25 | 0.148177551 | 0.19562813 | 0 |
| RE45537 | FBgn0036373 | Tgi | -0.148114169 | 0.21689011 | 0 |
| AT27044 | FBgn0031844 | CG13771 | -0.148109719 | 0.37374788 | 0 |
| GH18690 | FBgn0035266 | Gk2 | -0.148099038 | 0.24826642 | 0 |
| LD33761 | FBgn0263979 | Caf1-55 | 0.148079893 | 0.25086746 | 0 |
| AT19549 | FBgn0029989 | CG15333 | 0.14807098 | 0.24097117 | 0 |
| LD15342 | FBgn0051151 | wge | 0.148014109 | 0.22860822 | 0 |
| RE15838 | FBgn0037816 | CG6345 | 0.147982975 | 0.23829121 | 0 |
| LD30158 | FBgn0037021 | CG11399 | -0.147965274 | 0.26140399 | 0 |
| LD16736 | FBgn0016701 | Rab4 | -0.147768136 | 0.37785052 | 0 |

| | | | | | |
|---|---|---|---|---|---|
| RE01736 | FBgn0031005 | Hs3st-B | -0.147603506 | 0.39598601 | 0 |
| RE44251 | FBgn0036331 | CG14117 | -0.147532442 | 0.43161816 | 0 |
| RE09848 | FBgn0030759 | CG13014 | -0.147477887 | 0.35160214 | 0 |
| LD05664 | FBgn0025874 | Meics | 0.14747458 | 0.22082195 | 0 |
| RE59129 | FBgn0023212 | EloB | 0.147456928 | 0.27904632 | 0 |
| GH02315 | FBgn0030799 | CG4872 | 0.147435291 | 0.24244937 | 0 |
| GH07623 | FBgn0039560 | BOD1 | 0.147354424 | 0.27774367 | 0 |
| AT21853 | FBgn0035708 | axed | -0.147275615 | 0.56449163 | 0 |
| GH19141 | FBgn0037709 | CG8199 | -0.147213152 | 0.19717375 | 0 |
| RE24423 | FBgn0038366 | CG4576 | -0.14714952 | 0.45259874 | 0 |
| LD22010 | FBgn0035524 | CG11583 | 0.147131932 | 0.25086746 | 0 |
| LD21089 | FBgn0000289 | cg | 0.147084405 | 0.25710987 | 0 |
| AT18988 | FBgn0028896 | CG17329 | 0.147051536 | 0.25629977 | 0 |
| LD47007 | FBgn0037653 | CG11982 | 0.147004458 | 0.23683887 | 0 |
| LD38070 | FBgn0030969 | Usp39 | 0.14695156 | 0.28620267 | 0 |
| RE18761 | FBgn0038682 | CG5835 | 0.146897442 | 0.43735269 | 0 |
| LD22344 | FBgn0043456 | CG4747 | 0.146856677 | 0.21296194 | 0 |
| GH08377 | FBgn0040060 | yip7 | -0.146850138 | 0.49396123 | 0 |
| RE25320 | FBgn0030985 | Obp18a | 0.146811235 | 0.3155453 | 0 |
| GM16107 | FBgn0053082 | CG33082 | 0.146726168 | 0.30365845 | 0 |
| RH67410 | FBgn0031110 | Obp19b | -0.146698054 | 0.27774828 | 0 |
| LD21248 | FBgn0028671 | Vha100-1 | 0.146650289 | 0.27156398 | 0 |
| LD10325 | FBgn0037106 | CG11307 | 0.146609306 | 0.20055502 | 0 |
| GH09387 | FBgn0032061 | CG9314 | 0.146594706 | 0.28466789 | 0 |
| LD13018 | FBgn0010235 | Klc | -0.146449405 | 0.23069212 | 0 |
| GM13767 | FBgn0031229 | CG3436 | -0.146438537 | 0.307972 | 0 |
| GH24859 | FBgn0036029 | CG16719 | 0.146362287 | 0.52590592 | 0 |
| GM08665 | FBgn0021856 | l(2)k14505 | 0.1462161 | 0.33183546 | 0 |
| GH21465 | FBgn0051017 | PH4alphaNE3 | -0.146208529 | 0.20693481 | 0 |
| GH11210 | FBgn0000370 | crc | 0.146196458 | 0.22944009 | 0 |
| GH11210 | FBgn0284230 | CG46314 | 0.146196458 | 0.22944009 | 0 |
| GH12222 | FBgn0036211 | CG5946 | 0.146149734 | 0.27067267 | 0 |
| SD11801 | FBgn0261838 | pre-mod(mdg4)-Z | 0.146149622 | 0.22277648 | 0 |
| SD11801 | FBgn0261837 | pre-mod(mdg4)-T | 0.146149622 | 0.22277648 | 0 |
| LD10168 | FBgn0000250 | cact | 0.145988023 | 0.24345967 | 0 |
| LD03583 | FBgn0053138 | AGBE | -0.145955841 | 0.29335128 | 0 |
| LP09696 | FBgn0260744 | Tango9 | -0.145902871 | 0.3322274 | 0 |
| RH67819 | FBgn0011769 | Fdx1 | -0.145897569 | 0.21464103 | 0 |
| AT13913 | FBgn0037172 | ND-B14.5AL | 0.145888511 | 0.31204614 | 0 |
| LP09309 | FBgn0031428 | CG9886 | 0.14587134 | 0.23599521 | 0 |
| LD07532 | FBgn0010516 | wal | 0.145819275 | 0.22860822 | 0 |
| RE66761 | FBgn0032397 | Tom70 | -0.145804349 | 0.21650949 | 0 |
| GH16191 | FBgn0051148 | Gba1a | -0.145675858 | 0.30165319 | 0 |
| LD37349 | FBgn0038765 | CG4424 | 0.145634943 | 0.26307647 | 0 |
| LD44720 | FBgn0038256 | CG7530 | 0.145616948 | 0.2838771 | 0 |
| RE71708 | FBgn0039172 | Spase22-23 | -0.145595927 | 0.23635904 | 0 |
| SD07675 | FBgn0010660 | Nup214 | 0.145543405 | 0.21464232 | 0 |
| GM13341 | FBgn0037369 | CG2100 | 0.14549821 | 0.25250442 | 0 |

| | | | | | |
|---|---|---|---|---|---|
| LD06275 | FBgn0038640 | CG7706 | 0.14548953 | 0.27703693 | 0 |
| LD06123 | FBgn0051224 | CG31224 | -0.14546426 | 0.25821219 | 0 |
| LD04137 | FBgn0032901 | sky | -0.145413937 | 0.25821219 | 0 |
| LD04137 | FBgn0263996 | CG43739 | -0.145413937 | 0.25821219 | 0 |
| GM01527 | FBgn0036897 | CG8786 | -0.145366488 | 0.40053145 | 0 |
| AT20169 | FBgn0039139 | Mettl3 | -0.145344598 | 0.52700798 | 0 |
| LD38259 | FBgn0032940 | Mondo | 0.145328201 | 0.23906935 | 0 |
| RE22677 | FBgn0031092 | CG9577 | -0.14530158 | 0.30190742 | 0 |
| SD12357 | FBgn0050187 | CG30187 | 0.145277876 | 0.32437253 | 0 |
| GM06782 | FBgn0036191 | Sugb | 0.145268979 | 0.20693481 | 0 |
| LD43891 | FBgn0028546 | ics | 0.145268275 | 0.23778979 | 0 |
| RE31155 | FBgn0034756 | Cyp6d2 | -0.14523474 | 0.46064534 | 0 |
| LD24980 | FBgn0030877 | Arp8 | 0.145233142 | 0.23048196 | 0 |
| LD24355 | FBgn0034961 | CG3163 | 0.145202896 | 0.23778979 | 0 |
| GH11150 | FBgn0015828 | TfIIEalpha | -0.145157661 | 0.26610197 | 0 |
| LD41157 | FBgn0030013 | GIIIspla2 | -0.14514868 | 0.24829452 | 0 |
| RE40914 | FBgn0260467 | CG7071 | 0.145141501 | 0.28754289 | 0 |
| RE40914 | FBgn0083967 | Muted | 0.145141501 | 0.28754289 | 0 |
| LP10895 | FBgn0038250 | CG3505 | -0.145124211 | 0.32073204 | 0 |
| RE38449 | FBgn0052532 | CG32532 | 0.145013178 | 0.3139793 | 0 |
| LD10749 | FBgn0040929 | CG12659 | 0.144998349 | 0.23734967 | 0 |
| LD08974 | FBgn0262872 | milt | 0.144986117 | 0.22894331 | 0 |
| LD28985 | FBgn0034061 | Ufc1 | -0.144975097 | 0.34994022 | 0 |
| SD10843 | FBgn0264267 | asRNA:CR43767 | -0.144884676 | 0.28279691 | 0 |
| LP03403 | FBgn0026250 | eIF1A | -0.144858758 | 0.29448305 | 0 |
| LP03403 | FBgn0035057 | CG3880 | -0.144858758 | 0.29448305 | 0 |
| GH26613 | FBgn0051064 | CG31064 | 0.144730283 | 0.25352966 | 0 |
| HL07889 | FBgn0050011 | gem | -0.144655932 | 0.30100738 | 0 |
| HL07889 | FBgn0284235 | CG46319 | -0.144655932 | 0.30100738 | 0 |
| RE48767 | FBgn0035622 | TM9SF3 | 0.144624339 | 0.21224705 | 0 |
| AT21785 | FBgn0036075 | CG8065 | 0.144591842 | 0.45220473 | 0 |
| AT04823 | FBgn0069354 | Porin2 | -0.144574784 | 0.35405702 | 0 |
| GH02210 | FBgn0033954 | CG12860 | 0.144527856 | 0.4176333 | 0 |
| AT21495 | FBgn0041337 | Cyp309a2 | 0.144403008 | 0.23829121 | 0 |
| LD09689 | FBgn0284249 | Adf1 | -0.144392974 | 0.21836122 | 0 |
| LD17234 | FBgn0261881 | l(2)35Be | 0.144299009 | 0.23195764 | 0 |
| AT04491 | FBgn0038581 | CG14314 | 0.144298452 | 0.21836122 | 0 |
| RH66629 | FBgn0039305 | CG11858 | 0.144237656 | 0.2658646 | 0 |
| RE17794 | FBgn0038498 | beat-IIa | 0.144236986 | 0.43002019 | 0 |
| RE42506 | FBgn0040007 | RpL38 | -0.144189637 | 0.3551038 | 0 |
| SD14970 | FBgn0036018 | CG3335 | 0.144186838 | 0.31678475 | 0 |
| RH07980 | FBgn0025820 | JTBR | -0.144164532 | 0.24719359 | 0 |
| RE01053 | FBgn0038427 | ema | 0.144151366 | 0.27929923 | 0 |
| LD22577 | FBgn0029113 | Uba2 | 0.14413408 | 0.20926237 | 0 |
| RE20092 | FBgn0039441 | TwdlN | -0.144128827 | 0.47732665 | 0 |
| LD01807 | FBgn0030724 | Nipsnap | -0.144103443 | 0.25352966 | 0 |
| GH07085 | FBgn0032394 | CG6746 | -0.144011631 | 0.29672664 | 0 |
| AT28327 | FBgn0037986 | CG14736 | 0.143974877 | 0.43934245 | 0 |

| | | | | | |
|---|---|---|---|---|---|
| LD45279 | FBgn0038745 | CG4538 | 0.143918074 | 0.21862256 | 0 |
| GH07062 | FBgn0038686 | CG5555 | -0.143907529 | 0.26834543 | 0 |
| LD17001 | FBgn0039644 | CG11897 | 0.14390552 | 0.29328557 | 0 |
| RE53556 | FBgn0002577 | m | -0.143880451 | 0.41053073 | 0 |
| LD21735 | FBgn0028670 | Vha100-2 | -0.143797857 | 0.28988547 | 0 |
| RE60854 | FBgn0035405 | CG15812 | 0.143767981 | 0.25159252 | 0 |
| RE21686 | FBgn0034723 | CG13506 | 0.143691455 | 0.71344353 | 0 |
| LP04971 | FBgn0260770 | CG42568 | 0.143660845 | 0.29428669 | 0 |
| LD28817 | FBgn0037958 | CG6962 | 0.143658295 | 0.33743595 | 0 |
| SD10385 | FBgn0031897 | CG13784 | -0.143652942 | 0.3547438 | 0 |
| LD35151 | FBgn0030321 | CG1703 | -0.143631554 | 0.21312024 | 0 |
| GH03671 | FBgn0261985 | Ptpmeg | 0.143597366 | 0.22147922 | 0 |
| RH03777 | FBgn0030752 | CG9947 | 0.14358238 | 0.33990239 | 0 |
| RE02607 | FBgn0284084 | wg | 0.143508921 | 0.612073 | 0 |
| RE42582 | FBgn0019624 | COX5A | -0.143484256 | 0.28754289 | 0 |
| RE31002 | FBgn0036875 | CG9449 | 0.143464969 | 0.30918864 | 0 |
| AT31219 | FBgn0029745 | Rpn13R | 0.14341604 | 0.31825117 | 0 |
| AT27789 | FBgn0259139 | glo | 0.143325143 | 0.20884829 | 0 |
| GH02203 | FBgn0016078 | wun | 0.143316804 | 0.2353666 | 0 |
| GH10770 | FBgn0037822 | CG14683 | 0.143215178 | 0.26066785 | 0 |
| LD16810 | FBgn0003087 | pim | -0.143201827 | 0.4451242 | 0 |
| GH13548 | FBgn0024179 | wit | 0.14317809 | 0.22736386 | 0 |
| RE28434 | FBgn0036646 | CR18217 | -0.14313952 | 0.2476695 | 0 |
| LD43724 | FBgn0039250 | Mink | 0.143099093 | 0.24366394 | 0 |
| LD44814 | FBgn0037652 | CG11980 | -0.143049147 | 0.24700724 | 0 |
| GH14439 | FBgn0028984 | Spn88Ea | 0.142922756 | 0.34043647 | 0 |
| GH24286 | FBgn0031516 | CG9663 | 0.142912852 | 0.21473034 | 0 |
| LD29844 | FBgn0037521 | CG2993 | 0.142887422 | 0.20884829 | 0 |
| SD01586 | FBgn0003175 | px | -0.142885982 | 0.33866194 | 0 |
| SD03870 | FBgn0032702 | CG10376 | 0.142864988 | 0.20847027 | 0 |
| GH05829 | FBgn0034566 | CG9313 | 0.142798569 | 0.37710008 | 0 |
| LD44235 | FBgn0035713 | velo | 0.14265302 | 0.27578495 | 0 |
| AT13439 | FBgn0033498 | CG12209 | 0.142652804 | 0.28296337 | 0 |
| LP11629 | FBgn0033504 | CAP | 0.142548071 | 0.38272145 | 0 |
| GM05661 | FBgn0036557 | mRpS31 | -0.142543564 | 0.22495288 | 0 |
| LD23292 | FBgn0267488 | Mcr | -0.142478426 | 0.21014705 | 0 |
| LD30472 | FBgn0037900 | CG5276 | 0.142473829 | 0.23695841 | 0 |
| RH48979 | FBgn0030584 | CG14407 | -0.142434076 | 0.23614451 | 0 |
| AT23465 | FBgn0031596 | CG15429 | 0.142396253 | 0.43772294 | 0 |
| SD10366 | FBgn0011661 | Moe | -0.142350827 | 0.25086746 | 0 |
| SD09360 | FBgn0085430 | CG34401 | -0.142335951 | 0.36849518 | 0 |
| LD39491 | FBgn0261238 | Alh | 0.142307579 | 0.23434738 | 0 |
| GH01626 | FBgn0261788 | Ank2 | -0.142286594 | 0.37177187 | 0 |
| GM24483 | FBgn0034511 | GNBP-like3 | 0.142271234 | 0.30755546 | 0 |
| LP01162 | FBgn0035206 | CG9186 | -0.142184149 | 0.21318822 | 0 |
| LD38816 | FBgn0260655 | l(3)76BDm | 0.142150401 | 0.23502135 | 0 |
| GH08670 | FBgn0029755 | Sas10 | 0.142123534 | 0.22851335 | 0 |
| LP02988 | FBgn0033137 | Tsp42Ep | -0.142028336 | 0.31235324 | 0 |

| | | | | | |
|---|---|---|---|---|---|
| RH09802 | FBgn0063491 | GstE9 | -0.141980797 | 0.2658646 | 0 |
| GH12410 | FBgn0064237 | Idgf5 | 0.141973226 | 0.30786949 | 0 |
| LD35257 | FBgn0037489 | CG1234 | 0.141922326 | 0.21609263 | 0 |
| GH04717 | FBgn0032123 | Oatp30B | 0.141771197 | 0.22851335 | 0 |
| LD48007 | FBgn0039904 | Hcf | 0.141765345 | 0.31672867 | 0 |
| GH14130 | FBgn0034997 | CG3376 | 0.141728091 | 0.29835009 | 0 |
| LD44032 | FBgn0037539 | CG10435 | 0.141707228 | 0.20926155 | 0 |
| RE62293 | FBgn0265605 | Ric | -0.141683628 | 0.22598525 | 0 |
| LP07813 | FBgn0033725 | Cpr49Ac | 0.141621752 | 0.30050924 | 0 |
| LD42046 | FBgn0033185 | CG1603 | 0.141552007 | 0.24120973 | 0 |
| SD02110 | FBgn0003278 | RpI135 | 0.141431964 | 0.36120961 | 0 |
| RE56428 | FBgn0024320 | Npc1a | 0.14141799 | 0.21526439 | 0 |
| RE25122 | FBgn0003134 | Pp1alpha-96A | 0.141327732 | 0.29119321 | 0 |
| RE27439 | FBgn0261963 | mid | 0.141318269 | 0.34043647 | 0 |
| GH02866 | FBgn0034704 | CG6758 | -0.141317972 | 0.27711515 | 0 |
| RE57482 | FBgn0020440 | Fak | -0.14127275 | 0.27309497 | 0 |
| AT11816 | FBgn0036654 | CG9692 | -0.141029977 | 0.28847675 | 0 |
| RH08585 | FBgn0262801 | twr | -0.140976884 | 0.22276303 | 0 |
| LD33681 | FBgn0000117 | arm | 0.140869261 | 0.24120973 | 0 |
| GH25366 | FBgn0028970 | betaggt-II | 0.140810296 | 0.29332909 | 0 |
| RE38426 | FBgn0039091 | CG10182 | -0.140778234 | 0.48656864 | 0 |
| RH06121 | FBgn0052085 | CG32085 | -0.140753067 | 0.56419565 | 0 |
| LD02340 | FBgn0003416 | sl | 0.140646376 | 0.31103764 | 0 |
| GH21896 | FBgn0265137 | Spn42Da | 0.140612448 | 0.36641715 | 0 |
| LD47064 | FBgn0033485 | RpLP0-like | 0.140611509 | 0.2803392 | 0 |
| LD38503 | FBgn0035960 | CG4942 | 0.140555725 | 0.26226354 | 0 |
| AT30631 | FBgn0025612 | CG3062 | 0.140411938 | 0.21444299 | 0 |
| RE11908 | FBgn0038387 | blp | -0.140401402 | 0.26312193 | 0 |
| LD28793 | FBgn0039566 | CG4849 | 0.140333995 | 0.22894331 | 0 |
| RH43234 | FBgn0000092 | CG2650 | 0.140326746 | 0.60480402 | 0 |
| RE32990 | FBgn0038390 | Rbf2 | -0.140324338 | 0.60282369 | 0 |
| GM02932 | FBgn0033473 | CG12128 | 0.140300067 | 0.25748944 | 0 |
| GH06306 | FBgn0031294 | IA-2 | -0.140299246 | 0.31825117 | 0 |
| LD21794 | FBgn0027515 | CG7115 | -0.140287212 | 0.21366213 | 0 |
| RE13646 | FBgn0037578 | CG9601 | -0.14026578 | 0.29790263 | 0 |
| GH07774 | FBgn0027568 | Cand1 | 0.140196823 | 0.30470968 | 0 |
| LD45826 | FBgn0033891 | CG8067 | 0.140104711 | 0.25151244 | 0 |
| LD22866 | FBgn0028292 | ric8a | 0.140049732 | 0.21487795 | 0 |
| LD33222 | FBgn0263005 | CG43313 | -0.140046261 | 0.26375792 | 0 |
| SD07655 | FBgn0032456 | MRP | 0.139917776 | 0.2465071 | 0 |
| LD44641 | FBgn0039150 | CG13605 | 0.139913014 | 0.2677426 | 0 |
| RE21922 | FBgn0033257 | sand | 0.139910465 | 0.36885296 | 0 |
| LD27030 | FBgn0033264 | Nup50 | 0.139788596 | 0.2851942 | 0 |
| GH11591 | FBgn0039044 | p53 | 0.139762886 | 0.22929842 | 0 |
| GH23342 | FBgn0025454 | Cyp6g1 | -0.139737375 | 0.33998407 | 0 |
| LD36231 | FBgn0033738 | DUBAI | 0.139683683 | 0.25143788 | 0 |
| LD10092 | FBgn0025807 | Rad9 | -0.139567407 | 0.27527549 | 0 |
| LD16147 | FBgn0031268 | cold | -0.139526033 | 0.38507121 | 0 |

| | | | | | |
|---|---|---|---|---|---|
| RE03558 | FBgn0035522 | CG1273 | 0.139525718 | 0.58240653 | 0 |
| RE58242 | FBgn0052645 | CG32645 | -0.139359182 | 0.40617197 | 0 |
| RE19335 | FBgn0040010 | CG17493 | 0.139321765 | 0.31825117 | 0 |
| LD28033 | FBgn0283521 | lola | 0.139221274 | 0.23337287 | 0 |
| AT20211 | FBgn0036549 | CG10516 | 0.139216991 | 0.32653939 | 0 |
| AT23411 | FBgn0036714 | CG7692 | 0.139101437 | 0.22060168 | 0 |
| RE32514 | FBgn0031213 | galectin | -0.138974267 | 0.37774984 | 0 |
| LD41613 | FBgn0030366 | Usp7 | 0.138962039 | 0.23774463 | 0 |
| GM05760 | FBgn0039663 | CG2321 | -0.138934751 | 0.22005916 | 0 |
| LD47312 | FBgn0036661 | CG9705 | 0.138851528 | 0.21567639 | 0 |
| GM01918 | FBgn0030082 | HP1b | 0.138842572 | 0.27086432 | 0 |
| GM09101 | FBgn0003870 | ttk | -0.138825442 | 0.34274251 | 0 |
| LD17611 | FBgn0053505 | U3-55K | -0.138816004 | 0.4787 | 0 |
| LP04090 | FBgn0051374 | sals | 0.13878488 | 0.21635313 | 0 |
| LP04090 | FBgn0002565 | Lsp2 | 0.13878488 | 0.21635313 | 0 |
| RH59553 | FBgn0025814 | Mgstl | 0.138761037 | 0.30100331 | 0 |
| RH59553 | FBgn0042701 | CR12628 | 0.138761037 | 0.30100331 | 0 |
| GH13253 | FBgn0086445 | l(2)37Cd | 0.13868936 | 0.24946171 | 0 |
| GH04589 | FBgn0032814 | CG10366 | -0.138683175 | 0.27446683 | 0 |
| RE06911 | FBgn0035371 | AhcyL1 | -0.138675475 | 0.2308379 | 0 |
| GM13395 | FBgn0050342 | Prp38 | 0.138637384 | 0.37410463 | 0 |
| SD07605 | FBgn0038360 | CG9590 | 0.138599042 | 0.21845712 | 0 |
| RE38484 | FBgn0066101 | LpR1 | 0.138596453 | 0.34663191 | 0 |
| RE38484 | FBgn0011648 | Mad | 0.138596453 | 0.34663191 | 0 |
| AT17902 | FBgn0035273 | CG12020 | 0.138544452 | 0.21797572 | 0 |
| RE21057 | FBgn0030041 | CG12116 | 0.138485352 | 0.27441429 | 0 |
| RE21057 | FBgn0053223 | CG33223 | 0.138485352 | 0.27441429 | 0 |
| RE27227 | FBgn0027841 | CstF64 | 0.138468963 | 0.22266915 | 0 |
| AT28681 | FBgn0028942 | CG16852 | 0.138468187 | 0.33994384 | 0 |
| LD40944 | FBgn0033581 | CG12391 | 0.138461607 | 0.29099529 | 0 |
| LD41207 | FBgn0010269 | Dsor1 | -0.138423073 | 0.30080363 | 0 |
| RE73934 | FBgn0028398 | Taf10 | 0.138362561 | 0.33110818 | 0 |
| AT11375 | FBgn0031142 | r-cup | -0.138360391 | 0.45159352 | 0 |
| RE26559 | FBgn0025383 | CG14780 | -0.138346996 | 0.42399091 | 0 |
| RE46702 | FBgn0036288 | CG10660 | 0.138306134 | 0.46421815 | 0 |
| GH14462 | FBgn0039697 | CG7834 | 0.138305384 | 0.32237304 | 0 |
| GM02019 | FBgn0041111 | lilli | -0.138144225 | 0.4625759 | 0 |
| SD10213 | FBgn0034243 | Ns2 | 0.138117663 | 0.26687802 | 0 |
| LD23856 | FBgn0033734 | CG8520 | 0.138028343 | 0.37559906 | 0 |
| LD05707 | FBgn0051216 | Naam | 0.138028335 | 0.28703489 | 0 |
| LD22825 | FBgn0028952 | Kebab | 0.137967377 | 0.30609085 | 0 |
| LD12407 | FBgn0262975 | cnc | -0.137963891 | 0.29328557 | 0 |
| GH09688 | FBgn0027844 | CAH1 | 0.137955338 | 0.2947148 | 0 |
| RE70243 | FBgn0032002 | CG8353 | -0.137936403 | 0.39339733 | 0 |
| GM02206 | FBgn0052000 | anne | -0.137925636 | 0.34255324 | 0 |
| RE27803 | FBgn0033195 | CG1360 | 0.137919494 | 0.23606201 | 0 |
| RE71924 | FBgn0086706 | pix | 0.137823179 | 0.26124013 | 0 |
| LD21733 | FBgn0037638 | CG8379 | 0.137817844 | 0.2975402 | 0 |

| | | | | | |
|---|---|---|---|---|---|
| GH04863 | FBgn0035515 | CG14997 | 0.137799881 | 0.32338964 | 0 |
| RE49819 | FBgn0035308 | CG15822 | -0.137764655 | 0.32681987 | 0 |
| RE23085 | FBgn0037976 | Tk | -0.137729953 | 0.23778979 | 0 |
| LD29078 | FBgn0033177 | CG11141 | 0.13769713 | 0.22555325 | 0 |
| LD26511 | FBgn0000617 | e(y)1 | 0.137696711 | 0.26615408 | 0 |
| GH12873 | FBgn0005778 | PpD5 | 0.13765445 | 0.28208194 | 0 |
| RH48101 | FBgn0039043 | CG17121 | -0.137591727 | 0.35250161 | 0 |
| SD07336 | FBgn0032475 | Sfmbt | 0.137583702 | 0.24755362 | 0 |
| LD29039 | FBgn0002873 | mud | 0.137427735 | 0.24102302 | 0 |
| RE38650 | FBgn0033501 | CG12911 | 0.137410655 | 0.24485913 | 0 |
| GH10083 | FBgn0029863 | CG3823 | 0.137410298 | 0.39072093 | 0 |
| AT17761 | FBgn0036082 | CG12362 | 0.137350993 | 0.38272145 | 0 |
| HL03404 | FBgn0263392 | Tet | -0.137109993 | 0.46508385 | 0 |
| LD29485 | FBgn0035264 | Oseg4 | 0.137076162 | 0.26522758 | 0 |
| AT21810 | FBgn0038280 | CG14861 | 0.136975729 | 0.39180956 | 0 |
| GH10333 | FBgn0027553 | NELF-B | 0.13697392 | 0.28123934 | 0 |
| AT09144 | FBgn0038993 | CG13843 | 0.136893734 | 0.31358313 | 0 |
| GH04850 | FBgn0031645 | CG3036 | 0.13686661 | 0.25308468 | 0 |
| AT25917 | FBgn0029987 | CG15330 | -0.136803293 | 0.44373623 | 0 |
| AT26486 | FBgn0030384 | CG2577 | 0.136791468 | 0.32725139 | 0 |
| GH02773 | FBgn0016715 | Reg-2 | -0.136719427 | 0.40945908 | 0 |
| RE13779 | FBgn0266566 | CG45105 | -0.136702232 | 0.50789499 | 0 |
| AT25266 | FBgn0038630 | CG14305 | 0.13666513 | 0.2888356 | 0 |
| SD10447 | FBgn0016081 | fry | 0.136588746 | 0.39339733 | 0 |
| SD05875 | FBgn0259178 | 5PtaseI | -0.136561086 | 0.29529306 | 0 |
| LD08195 | FBgn0040395 | Unc-76 | -0.136555497 | 0.24806598 | 0 |
| RE03865 | FBgn0000659 | fkh | -0.136549977 | 0.36404769 | 0 |
| GH01660 | FBgn0001989 | ND-B17 | 0.136517409 | 0.28916091 | 0 |
| HL04357 | FBgn0264572 | asRNA:CR43944 | 0.136449001 | 0.22082195 | 0 |
| RE66853 | FBgn0015904 | ara | 0.136440618 | 0.4655033 | 0 |
| GH10306 | FBgn0036759 | CG5577 | -0.13643207 | 0.30293062 | 0 |
| GM04645 | FBgn0040259 | Ugt86Da | -0.136391354 | 0.23170768 | 0 |
| GH12904 | FBgn0034722 | Rtf1 | 0.136330847 | 0.272048 | 0 |
| RE58581 | FBgn0013469 | klu | 0.136313413 | 0.28703489 | 0 |
| LD12965 | FBgn0002645 | Map205 | 0.136291442 | 0.22390724 | 0 |
| GH26184 | FBgn0039667 | CG2010 | 0.136269342 | 0.43887794 | 0 |
| LD12690 | FBgn0086371 | poly | -0.136138501 | 0.25439475 | 0 |
| GH14674 | FBgn0064115 | GatC | -0.136137651 | 0.34411592 | 0 |
| GH14674 | FBgn0004407 | DNApol-gamma35 | -0.136137651 | 0.34411592 | 0 |
| LD24482 | FBgn0015664 | Dref | 0.136121571 | 0.23101323 | 0 |
| GM13045 | FBgn0037874 | Tctp | -0.13610472 | 0.41864591 | 0 |
| LD44495 | FBgn0028683 | spt4 | -0.136052851 | 0.24556656 | 0 |
| LD23644 | FBgn0014189 | Hel25E | 0.136029321 | 0.32458788 | 0 |
| LD30953 | FBgn0033095 | CG3409 | 0.13602324 | 0.25250442 | 0 |
| LD30953 | FBgn0266525 | CG45092 | 0.13602324 | 0.25250442 | 0 |
| SD07391 | FBgn0038168 | omd | -0.135988346 | 0.32007204 | 0 |
| RE58962 | FBgn0037129 | CG14565 | 0.135960543 | 0.57265237 | 0 |
| SD04011 | FBgn0030893 | RhoGAP16F | 0.135954315 | 0.22250637 | 0 |

| | | | | | |
|---|---|---|---|---|---|
| LD20162 | FBgn0051633 | CG31633 | -0.135902293 | 0.24582366 | 0 |
| RE43738 | FBgn0003944 | Ubx | 0.135899812 | 0.33784941 | 0 |
| LD10857 | FBgn0030683 | CG8239 | -0.135848696 | 0.22929842 | 0 |
| LD10609 | FBgn0033984 | Lap1 | 0.135777554 | 0.26233948 | 0 |
| LD38082 | FBgn0030912 | CG6023 | -0.13572034 | 0.38555567 | 0 |
| LD45246 | FBgn0038401 | CG5916 | -0.135618991 | 0.27904632 | 0 |
| RE13279 | FBgn0030994 | HP1D3csd | -0.135551728 | 0.43390857 | 0 |
| LP04791 | FBgn0028532 | CG7968 | 0.13554285 | 0.26370926 | 0 |
| RE72132 | FBgn0031607 | CG15440 | 0.135532151 | 0.24960785 | 0 |
| RE68879 | FBgn0039943 | CG17168 | 0.135523637 | 0.2323332 | 0 |
| GH02220 | FBgn0027615 | CG6404 | -0.135421821 | 0.23048196 | 0 |
| LD47606 | FBgn0264302 | CG43780 | 0.135396119 | 0.25086746 | 0 |
| LD47606 | FBgn0264303 | CG43781 | 0.135396119 | 0.25086746 | 0 |
| AT26438 | FBgn0031749 | CG14000 | -0.135367488 | 0.24417029 | 0 |
| LD43649 | FBgn0033916 | Usp20-33 | -0.135334454 | 0.25336796 | 0 |
| LD04591 | FBgn0039492 | CG6051 | 0.135269203 | 0.22199756 | 0 |
| LD14173 | FBgn0036090 | CG8009 | 0.135220781 | 0.31586471 | 0 |
| RE29544 | FBgn0032082 | CG18088 | 0.1351928 | 0.28911598 | 0 |
| LD15160 | FBgn0283657 | Tlk | -0.135028445 | 0.30993577 | 0 |
| GH12946 | FBgn0052548 | CG32548 | -0.135007248 | 0.37153414 | 0 |
| RH56202 | FBgn0002643 | mam | 0.134941578 | 0.38135654 | 0 |
| RE71127 | FBgn0032144 | CG17633 | 0.134929836 | 0.38667969 | 0 |
| LD41907 | FBgn0010309 | pigeon | -0.134917569 | 0.25270491 | 0 |
| AT15718 | FBgn0037115 | CG11249 | -0.134886263 | 0.22787212 | 0 |
| LD10463 | FBgn0013746 | alien | 0.134876202 | 0.25489759 | 0 |
| LD31194 | FBgn0015778 | rin | 0.134780651 | 0.36212941 | 0 |
| RH68724 | FBgn0052695 | CG32695 | 0.134730903 | 0.48277258 | 0 |
| RE21021 | FBgn0263510 | nclb | 0.134535292 | 0.29332909 | 0 |
| RE65343 | FBgn0022724 | Taf8 | 0.134481359 | 0.29135153 | 0 |
| RE32413 | FBgn0033590 | CG13217 | 0.134450384 | 0.48619427 | 0 |
| RE01752 | FBgn0259099 | DCX-EMAP | -0.134412119 | 0.30418103 | 0 |
| SD22308 | FBgn0030237 | CG15209 | 0.134315037 | 0.46023118 | 0 |
| GH05975 | FBgn0031814 | retm | -0.134287724 | 0.27882139 | 0 |
| AT13932 | FBgn0050462 | ste24c | 0.134275107 | 0.40810689 | 0 |
| AT13932 | FBgn0050461 | CG30461 | 0.134275107 | 0.40810689 | 0 |
| RE03918 | FBgn0026415 | Idgf4 | 0.134252729 | 0.39490947 | 0 |
| GH11731 | FBgn0039797 | eIF4H2 | -0.134235097 | 0.32169954 | 0 |
| LD06546 | FBgn0011703 | RnrL | -0.134186721 | 0.34043647 | 0 |
| LP02831 | FBgn0033124 | Tsp42Ec | -0.134180924 | 0.38251295 | 0 |
| LD13917 | FBgn0027338 | Kap-alpha3 | 0.13409062 | 0.31301389 | 0 |
| LD41490 | FBgn0263929 | jvl | 0.134084304 | 0.2908404 | 0 |
| SD05285 | FBgn0003660 | Syb | 0.134060952 | 0.42747108 | 0 |
| LD24203 | FBgn0014340 | mof | 0.134036554 | 0.26066785 | 0 |
| LD30049 | FBgn0026570 | CG5704 | 0.134008028 | 0.24604725 | 0 |
| AT23209 | FBgn0039752 | CG15530 | -0.133996916 | 0.56066561 | 0 |
| RE63289 | FBgn0040385 | CG12496 | 0.133975323 | 0.23541966 | 0 |
| RE65766 | FBgn0037892 | mRpL40 | -0.133927922 | 0.29214334 | 0 |
| SD03973 | FBgn0033996 | CG11807 | -0.133923529 | 0.22771972 | 0 |

| | | | | | |
|---|---|---|---|---|---|
| RH55416 | FBgn0052758 | CG32758 | -0.133909993 | 0.40479015 | 0 |
| GH12543 | FBgn0259795 | loopin-1 | -0.133885516 | 0.35636665 | 0 |
| LD26412 | FBgn0031633 | Fnta | -0.133863808 | 0.29250904 | 0 |
| RE16782 | FBgn0003334 | Scm | -0.133846653 | 0.23048196 | 0 |
| HL01082 | FBgn0026721 | fat-spondin | -0.133832612 | 0.50705399 | 0 |
| LD11641 | FBgn0024947 | NTPase | 0.133789414 | 0.25250442 | 0 |
| LD20082 | FBgn0014029 | Sep2 | 0.133720036 | 0.27711515 | 0 |
| RE53265 | FBgn0023083 | fray | -0.133582026 | 0.23413506 | 0 |
| RH04289 | FBgn0037873 | SdhC | -0.133481233 | 0.24806598 | 0 |
| GH27888 | FBgn0032713 | CG17323 | 0.133403119 | 0.74043457 | 0 |
| GM16226 | FBgn0032703 | CG10343 | 0.133348497 | 0.22754963 | 0 |
| AT23419 | FBgn0052252 | asRNA:CR32252 | 0.133037064 | 0.31049626 | 0 |
| GM13948 | FBgn0037351 | RpL13A | 0.133031121 | 0.42689889 | 0 |
| LD16566 | FBgn0036309 | Hip1 | 0.13297421 | 0.3883281 | 0 |
| RE30678 | FBgn0031851 | Aatf | 0.132942234 | 0.24417029 | 0 |
| LP04383 | FBgn0038299 | Spn88Eb | 0.132919556 | 0.27805782 | 0 |
| AT28833 | FBgn0037899 | RpL24-like | 0.132896735 | 0.24372705 | 0 |
| SD05207 | FBgn0267828 | Fatp | 0.132880644 | 0.29250904 | 0 |
| LP08544 | FBgn0033062 | Ars2 | 0.13283885 | 0.22929842 | 0 |
| RH63365 | FBgn0037469 | Dpck | 0.132814684 | 0.33831061 | 0 |
| RH08291 | FBgn0025583 | IM2 | -0.13281021 | 0.42681424 | 0 |
| SD03177 | FBgn0260003 | Dys | 0.132805968 | 0.30092567 | 0 |
| LP03146 | FBgn0033848 | CG13330 | -0.132671674 | 0.30080363 | 0 |
| RH56317 | FBgn0053169 | CG33169 | -0.132656213 | 0.27510038 | 0 |
| LP05695 | FBgn0033297 | Mal-A8 | 0.132463498 | 0.26424444 | 0 |
| RH05415 | FBgn0040637 | CG11458 | -0.132413885 | 0.41907068 | 0 |
| LD32106 | FBgn0010408 | RpS9 | 0.132350496 | 0.25865893 | 0 |
| GH28730 | FBgn0035499 | Chd64 | 0.132299477 | 0.27391484 | 0 |
| RE28982 | FBgn0011722 | Tig | -0.132279678 | 0.23172711 | 0 |
| AT20635 | FBgn0029643 | CG14420 | -0.132276968 | 0.24315669 | 0 |
| LD43109 | FBgn0267351 | Topors | 0.132171963 | 0.23719307 | 0 |
| RH67967 | FBgn0032465 | CG12404 | -0.132129849 | 0.29545439 | 0 |
| LD10408 | FBgn0003607 | Su(var)205 | 0.132118382 | 0.28600853 | 0 |
| LD43554 | FBgn0030612 | CG5599 | -0.132109415 | 0.267185 | 0 |
| LD12131 | FBgn0030003 | CG2116 | 0.132091871 | 0.24338343 | 0 |
| LD28662 | FBgn0003415 | skd | 0.132090433 | 0.25250442 | 0 |
| RE73615 | FBgn0051145 | CG31145 | -0.13206992 | 0.42280478 | 0 |
| LD41848 | FBgn0029914 | CG4558 | 0.132032207 | 0.23064179 | 0 |
| RH61722 | FBgn0029639 | CG14419 | -0.132023997 | 0.33919374 | 0 |
| LD30271 | FBgn0041171 | ago | 0.131999637 | 0.24338343 | 0 |
| GH16384 | FBgn0030777 | CG9672 | -0.131989313 | 0.24893482 | 0 |
| SD07139 | FBgn0038872 | Nelf-A | -0.131956344 | 0.4047529 | 0 |
| GH01248 | FBgn0053107 | CG33107 | -0.131946948 | 0.34457495 | 0 |
| AT26312 | FBgn0259794 | sinah | -0.131903408 | 0.31811714 | 0 |
| LD41276 | FBgn0052207 | hpRNA:CR32207 | -0.13178156 | 0.27557713 | 0 |
| RH30069 | FBgn0023530 | CG3740 | -0.131684784 | 0.37872757 | 0 |
| LD45430 | FBgn0020279 | lig | 0.131528234 | 0.24910111 | 0 |
| LD39967 | FBgn0035763 | CG8602 | 0.131510922 | 0.31622897 | 0 |

| | | | | | |
|---|---|---|---|---|---|
| GH27092 | FBgn0033698 | CG8858 | -0.131510332 | 0.31996025 | 0 |
| RH19248 | FBgn0040551 | CG11686 | 0.131503377 | 0.26364213 | 0 |
| AT25373 | FBgn0051287 | CG31287 | -0.131485239 | 0.27703693 | 0 |
| AT01812 | FBgn0033860 | S-Lap5 | 0.131471234 | 0.410397 | 0 |
| GH03876 | FBgn0011740 | alpha-Man-IIa | -0.131449482 | 0.28208194 | 0 |
| GH23768 | FBgn0030742 | CG9919 | 0.13143657 | 0.37774984 | 0 |
| SD07787 | FBgn0033038 | CG7791 | 0.131412043 | 0.39149549 | 0 |
| RE18426 | FBgn0033153 | Gadd45 | 0.131358043 | 0.58332023 | 0 |
| RE40156 | FBgn0037503 | CG14598 | -0.131229422 | 0.59911737 | 0 |
| LD12613 | FBgn0035281 | Cpr62Bc | -0.131225927 | 0.25656875 | 0 |
| RE16722 | FBgn0039896 | yellow-h | 0.131182034 | 0.27527304 | 0 |
| LD43412 | FBgn0042180 | CG18870 | -0.131136721 | 0.24981226 | 0 |
| LD12115 | FBgn0031379 | CG7289 | -0.131128588 | 0.32345633 | 0 |
| RE33807 | FBgn0031643 | CG3008 | 0.130999221 | 0.24685736 | 0 |
| GH07092 | FBgn0036031 | CG6761 | -0.130950808 | 0.27121609 | 0 |
| RE55238 | FBgn0004657 | mys | 0.130929953 | 0.24446919 | 0 |
| RE54693 | FBgn0037996 | CG4830 | -0.130734507 | 0.29328557 | 0 |
| GH08251 | FBgn0036091 | CG18628 | 0.130726447 | 0.29328557 | 0 |
| LD46952 | FBgn0266064 | GlyS | 0.130661824 | 0.34170455 | 0 |
| AT11556 | FBgn0039374 | CG17770 | -0.130613594 | 0.50652933 | 0 |
| AT11556 | FBgn0011273 | Acam | -0.130613594 | 0.50652933 | 0 |
| SD20854 | FBgn0040342 | CG3706 | 0.130607671 | 0.29835009 | 0 |
| LP06338 | FBgn0015010 | Ag5r | 0.130561365 | 0.41242951 | 0 |
| RE57757 | FBgn0000541 | E(bx) | -0.130535535 | 0.35050206 | 0 |
| LD23303 | FBgn0039117 | tst | -0.130358129 | 0.26375792 | 0 |
| RE53774 | FBgn0035207 | Herc4 | 0.13034879 | 0.2766162 | 0 |
| LD38226 | FBgn0031664 | CG8892 | -0.130326095 | 0.24417029 | 0 |
| LD37819 | FBgn0035909 | ergic53 | 0.130284813 | 0.38278109 | 0 |
| GH15688 | FBgn0263911 | COX8 | 0.130272053 | 0.26808258 | 0 |
| AT26616 | FBgn0034846 | CG9863 | 0.130254487 | 0.59512639 | 0 |
| AT12644 | FBgn0052087 | CG32087 | -0.130178137 | 0.30476758 | 0 |
| RE50345 | FBgn0036638 | CG13033 | -0.130119338 | 0.45761209 | 0 |
| GH01278 | FBgn0033669 | PI31 | -0.130106015 | 0.27156398 | 0 |
| RE35738 | FBgn0037166 | CG11426 | 0.130086074 | 0.3155453 | 0 |
| LD28419 | FBgn0028468 | rtet | 0.130054911 | 0.330418 | 0 |
| LD19810 | FBgn0027356 | Amph | -0.130044282 | 0.28758766 | 0 |
| LD45302 | FBgn0003449 | snf | 0.129980933 | 0.31916907 | 0 |
| LD07624 | FBgn0034360 | CG10927 | 0.129977339 | 0.35619541 | 0 |
| LD12912 | FBgn0035772 | Sh3beta | -0.129954596 | 0.36404769 | 0 |
| LD28038 | FBgn0025678 | CaBP1 | 0.129946279 | 0.28822565 | 0 |
| GH23934 | FBgn0040992 | CG10570 | 0.129933989 | 0.3267099 | 0 |
| GH23934 | FBgn0260228 | CG42502 | 0.129933989 | 0.3267099 | 0 |
| LD24159 | FBgn0002284 | Prosbeta6 | 0.129900609 | 0.25454796 | 0 |
| SD03848 | FBgn0260634 | eIF4G2 | -0.129809707 | 0.32681987 | 0 |
| GH09263 | FBgn0027088 | GlyRS | -0.129777224 | 0.29096007 | 0 |
| LD39323 | FBgn0250786 | Chd1 | -0.129733086 | 0.27045007 | 0 |
| GM02567 | FBgn0036460 | CG5114 | -0.12972387 | 0.25358171 | 0 |
| LD19406 | FBgn0034013 | unc-5 | 0.129691313 | 0.34401092 | 0 |

| | | | | | |
|---|---|---|---|---|---|
| AT08405 | FBgn0266432 | CG45062 | 0.129613667 | 0.29914002 | 0 |
| LD25151 | FBgn0037446 | Zif | 0.129503402 | 0.30663023 | 0 |
| LD35918 | FBgn0003654 | sw | 0.129474418 | 0.34570469 | 0 |
| LD35918 | FBgn0053497 | Sdic2 | 0.129474418 | 0.34570469 | 0 |
| LD35918 | FBgn0067861 | Sdic1 | 0.129474418 | 0.34570469 | 0 |
| AT04449 | FBgn0051128 | CG31128 | 0.129392052 | 0.51401758 | 0 |
| RE39287 | FBgn0001086 | fzy | 0.129361226 | 0.25159252 | 0 |
| GM10015 | FBgn0035452 | CG10359 | -0.12934592 | 0.3883281 | 0 |
| GH10905 | FBgn0003053 | peb | -0.129328206 | 0.56750708 | 0 |
| RE44773 | FBgn0038828 | CG17270 | -0.129319596 | 0.26651248 | 0 |
| LP05936 | FBgn0032208 | Ufd4 | 0.129252098 | 0.23778979 | 0 |
| LD14392 | FBgn0027053 | CSN5 | 0.129168595 | 0.26097892 | 0 |
| RE31401 | FBgn0029798 | CG4078 | 0.129128725 | 0.24338343 | 0 |
| LD21207 | FBgn0061476 | Zwilch | -0.129093201 | 0.27955803 | 0 |
| GH17676 | FBgn0053278 | CG33278 | 0.12908396 | 0.46704143 | 0 |
| LD06532 | FBgn0035121 | Tudor-SN | 0.129050602 | 0.33866194 | 0 |
| LD16661 | FBgn0029006 | Smurf | -0.129011819 | 0.44573816 | 0 |
| GH11066 | FBgn0039325 | CG10560 | 0.129009496 | 0.33976676 | 0 |
| LD43457 | FBgn0050440 | CG30440 | 0.128995357 | 0.36865559 | 0 |
| LD23779 | FBgn0026317 | Tsc1 | 0.128977742 | 0.51677309 | 0 |
| LD41405 | FBgn0015589 | Apc | 0.128918903 | 0.32747679 | 0 |
| GH18574 | FBgn0261562 | CG42676 | 0.128914266 | 0.30100738 | 0 |
| SD07467 | FBgn0086895 | pea | 0.128852922 | 0.40945908 | 0 |
| LD37010 | FBgn0020510 | Abi | 0.128771164 | 0.28758766 | 0 |
| RE19290 | FBgn0020258 | ppk | 0.128763179 | 0.43651754 | 0 |
| GH23574 | FBgn0029117 | Surf1 | -0.128751984 | 0.25900832 | 0 |
| RE30473 | FBgn0039068 | CG13827 | -0.128751801 | 0.45889729 | 0 |
| LD15494 | FBgn0036117 | CG6321 | 0.128747249 | 0.34419536 | 0 |
| LD45758 | FBgn0036333 | MICAL-like | -0.12873931 | 0.29544216 | 0 |
| LD23745 | FBgn0025825 | HDAC3 | 0.128670583 | 0.34069725 | 0 |
| LD23745 | FBgn0266539 | CG45100 | 0.128670583 | 0.34069725 | 0 |
| RH44949 | FBgn0016650 | Lgr1 | -0.12864947 | 0.5274825 | 0 |
| GH20547 | FBgn0260480 | CG32295 | 0.128585823 | 0.45433285 | 0 |
| LD29083 | FBgn0032234 | gny | 0.128569354 | 0.31443438 | 0 |
| GH01266 | FBgn0027572 | CG5009 | 0.128490661 | 0.29332909 | 0 |
| RE05926 | FBgn0022800 | Cad96Ca | 0.128451723 | 0.40940941 | 0 |
| LD37618 | FBgn0036735 | Edc3 | 0.128424283 | 0.28020454 | 0 |
| GH09258 | FBgn0038952 | CG7069 | -0.128421307 | 0.35114145 | 0 |
| GH24458 | FBgn0034985 | CG3328 | -0.128410301 | 0.24662826 | 0 |
| LD10783 | FBgn0015808 | ScpX | -0.128401602 | 0.26424444 | 0 |
| LD10783 | FBgn0032715 | CG17597 | -0.128401602 | 0.26424444 | 0 |
| AT31812 | FBgn0052588 | CG32588 | -0.128353513 | 0.375197 | 0 |
| RE21313 | FBgn0036994 | CG5199 | -0.128342786 | 0.41639809 | 0 |
| GH27779 | FBgn0051663 | CG31663 | 0.12832243 | 0.43887794 | 0 |
| LD39811 | FBgn0037241 | CG14646 | 0.128300594 | 0.25250442 | 0 |
| RE11562 | FBgn0259979 | CG17337 | -0.128192992 | 0.29331016 | 0 |
| GH27591 | FBgn0003360 | sesB | -0.12818563 | 0.27904632 | 0 |
| LD44221 | FBgn0051549 | CG31549 | 0.128145486 | 0.29488289 | 0 |

| | | | | | |
|---|---|---|---|---|---|
| GH13775 | FBgn0259176 | bun | 0.128097892 | 0.28122998 | 0 |
| LD18032 | FBgn0051678 | CG31678 | 0.128046328 | 0.25352966 | 0 |
| AT13882 | FBgn0029596 | CG14054 | -0.128008295 | 0.24525282 | 0 |
| RE52038 | FBgn0026755 | Ugt37b1 | -0.127995842 | 0.30993577 | 0 |
| RE24718 | FBgn0030723 | dpr18 | 0.127989028 | 0.28383357 | 0 |
| RE55753 | FBgn0031315 | CG14341 | 0.127973826 | 0.27156398 | 0 |
| RH68173 | FBgn0011274 | Dif | 0.127970312 | 0.34356345 | 0 |
| RE57840 | FBgn0035084 | CG15861 | 0.127968786 | 0.24719359 | 0 |
| GH19286 | FBgn0035557 | CG11353 | -0.127955287 | 0.41650931 | 0 |
| AT10573 | FBgn0032054 | Uba4 | 0.127933483 | 0.26058224 | 0 |
| RE16764 | FBgn0266579 | tau | -0.127859357 | 0.55944204 | 0 |
| LD39940 | FBgn0050089 | CG30089 | -0.127828498 | 0.57121226 | 0 |
| LD25514 | FBgn0035890 | CG13667 | 0.12778465 | 0.25086746 | 0 |
| LD22902 | FBgn0027378 | MRG15 | 0.127776144 | 0.33866194 | 0 |
| LD29280 | FBgn0019957 | ND-42 | 0.127674441 | 0.27859447 | 0 |
| GM08326 | FBgn0063492 | GstE8 | 0.127625292 | 0.28754289 | 0 |
| RE48511 | FBgn0034974 | CG16786 | -0.127593535 | 0.50688795 | 0 |
| RE13143 | FBgn0039111 | PTPMT1 | -0.127540858 | 0.26424444 | 0 |
| RE27292 | FBgn0034366 | Atg7 | 0.127452267 | 0.28299558 | 0 |
| RE47050 | FBgn0260934 | par-1 | -0.127450888 | 0.25744851 | 0 |
| RH44935 | FBgn0030292 | CG11752 | 0.127450519 | 0.35000557 | 0 |
| LP11750 | FBgn0029826 | CG6041 | 0.12741882 | 0.43335785 | 0 |
| HL04706 | FBgn0260005 | wtrw | -0.127417535 | 0.60534837 | 0 |
| RE40567 | FBgn0265351 | nac | 0.127375637 | 0.31677144 | 0 |
| LD23683 | FBgn0039335 | Vps33B | 0.127322362 | 0.32418356 | 0 |
| RE01490 | FBgn0031734 | CG11147 | -0.127292685 | 0.60742739 | 0 |
| SD05480 | FBgn0032521 | CG7110 | 0.127265575 | 0.26424444 | 0 |
| SD08037 | FBgn0039385 | CG5913 | -0.127254305 | 0.25159252 | 0 |
| GH18971 | FBgn0029799 | CG15772 | -0.127156633 | 0.50684673 | 0 |
| SD06318 | FBgn0033224 | Nop17l | -0.127152689 | 0.25189448 | 0 |
| LD23605 | FBgn0038463 | CG3534 | 0.127032046 | 0.25489759 | 0 |
| GH23590 | FBgn0042092 | CG13773 | -0.127024912 | 0.36614213 | 0 |
| LD30037 | FBgn0016070 | smg | 0.127017836 | 0.37463606 | 0 |
| LP08365 | FBgn0033875 | CG6357 | 0.126936842 | 0.24417029 | 0 |
| GH18028 | FBgn0023507 | D2hgdh | -0.126888022 | 0.36703978 | 0 |
| RE60237 | FBgn0083228 | Frq2 | -0.126862524 | 0.74067491 | 0 |
| GM08688 | FBgn0040075 | rept | 0.126797949 | 0.28758766 | 0 |
| SD02536 | FBgn0020930 | Dgkepsilon | 0.126765864 | 0.25847808 | 0 |
| RH33060 | FBgn0035022 | CG11413 | 0.126764892 | 0.30993649 | 0 |
| AT25463 | FBgn0037751 | topi | -0.126666435 | 0.30100738 | 0 |
| RE67729 | FBgn0035545 | CG12607 | -0.126649446 | 0.59935081 | 0 |
| RE10855 | FBgn0040020 | MED21 | -0.126646057 | 0.27815421 | 0 |
| RE63354 | FBgn0020655 | ArfGAP1 | 0.126638071 | 0.34812568 | 0 |
| LD37992 | FBgn0265935 | coro | 0.1266185 | 0.31844787 | 0 |
| RE34143 | FBgn0086698 | frtz | 0.126604315 | 0.27698881 | 0 |
| GH18902 | FBgn0039563 | CG4951 | -0.126593232 | 0.42214097 | 0 |
| LD25083 | FBgn0035918 | Cdc6 | 0.126575536 | 0.40940941 | 0 |
| GH16343 | FBgn0037756 | CG8507 | -0.126568968 | 0.34713139 | 0 |

| | | | | | |
|---|---|---|---|---|---|
| RE46682 | FBgn0030576 | CG15890 | -0.126559297 | 0.57209443 | 0 |
| LD03548 | FBgn0040347 | mus81 | 0.126524044 | 0.24542373 | 0 |
| LP01559 | FBgn0028514 | CG4793 | 0.126494413 | 0.30755546 | 0 |
| LD47466 | FBgn0041604 | dlp | 0.126485063 | 0.33728419 | 0 |
| LD24721 | FBgn0032640 | Sgt | 0.126478064 | 0.39339733 | 0 |
| AT06247 | FBgn0051050 | CG31050 | -0.126448456 | 0.24813057 | 0 |
| SD11991 | FBgn0031640 | Mon1 | 0.126431927 | 0.27774828 | 0 |
| AT21004 | FBgn0039498 | CG17991 | -0.126425191 | 0.307972 | 0 |
| RE71384 | FBgn0005649 | Rox8 | 0.126415137 | 0.25696915 | 0 |
| RE28720 | FBgn0040942 | CG12643 | 0.126409078 | 0.43887794 | 0 |
| GH17076 | FBgn0032358 | Ppt2 | -0.126391401 | 0.2829206 | 0 |
| LD24756 | FBgn0030329 | prtp | 0.12632577 | 0.3610233 | 0 |
| LD29665 | FBgn0035025 | uri | 0.126322445 | 0.25873823 | 0 |
| RE44043 | FBgn0039900 | Syt7 | 0.126225248 | 0.31005535 | 0 |
| SD22171 | FBgn0083962 | CG34126 | 0.126142394 | 0.25569722 | 0 |
| LD22081 | FBgn0034629 | Acox57D-d | -0.126058257 | 0.28294178 | 0 |
| LP02169 | FBgn0033266 | Socs44A | -0.126018768 | 0.25471465 | 0 |
| LD11211 | FBgn0031285 | CG3662 | 0.126001872 | 0.24946171 | 0 |
| SD03168 | FBgn0050147 | Hil | 0.125997141 | 0.27838547 | 0 |
| SD07608 | FBgn0003079 | Raf | 0.125789602 | 0.24981226 | 0 |
| GH04835 | FBgn0042086 | Tsp42Eb | -0.12577854 | 0.6049282 | 0 |
| GH04835 | FBgn0050160 | CG30160 | -0.12577854 | 0.6049282 | 0 |
| LP05220 | FBgn0262126 | Sec24CD | -0.125669731 | 0.35114145 | 0 |
| RE60459 | FBgn0250754 | CG42232 | -0.125636947 | 0.72296584 | 0 |
| GH02505 | FBgn0003249 | Rh3 | 0.125616525 | 0.29785901 | 0 |
| AT03240 | FBgn0035014 | CG13581 | -0.125526417 | 0.31964356 | 0 |
| SD08329 | FBgn0035142 | Hipk | 0.12550485 | 0.35969384 | 0 |
| RH57845 | FBgn0034103 | prim | -0.12549675 | 0.28014595 | 0 |
| GH02459 | FBgn0036286 | CG10616 | -0.125430792 | 0.2829206 | 0 |
| GH07929 | FBgn0027791 | O-fut2 | -0.125386971 | 0.44521723 | 0 |
| RE21692 | FBgn0036258 | eIF3l | 0.125385872 | 0.43962207 | 0 |
| GH28013 | FBgn0034394 | CG15096 | 0.125347465 | 0.35079094 | 0 |
| GH09052 | FBgn0037386 | CG1208 | 0.125347212 | 0.48048527 | 0 |
| LD28127 | FBgn0035425 | CG17746 | 0.125260664 | 0.31049626 | 0 |
| GH10777 | FBgn0028484 | Ack | 0.125259581 | 0.25821219 | 0 |
| GM09977 | FBgn0035830 | CG8209 | 0.125222174 | 0.33491792 | 0 |
| RH58777 | FBgn0016726 | RpL29 | -0.125173245 | 0.29135153 | 0 |
| LD10005 | FBgn0028980 | tant | 0.125109817 | 0.24960785 | 0 |
| LD27356 | FBgn0031598 | Vps53 | 0.125074985 | 0.29328557 | 0 |
| LD10351 | FBgn0034267 | CG4984 | -0.12502959 | 0.32839578 | 0 |
| LD46494 | FBgn0031611 | FIG4 | -0.125010699 | 0.48048527 | 0 |
| LD05345 | FBgn0034858 | eIF2Bdelta | -0.124996019 | 0.40746238 | 0 |
| SD16815 | FBgn0051619 | nolo | 0.124948183 | 0.468209 | 0 |
| RH42690 | FBgn0029887 | CG3198 | 0.12494064 | 0.33483119 | 0 |
| LD09078 | FBgn0025463 | Bap60 | 0.12488436 | 0.25907018 | 0 |
| LD47264 | FBgn0029831 | CG5966 | -0.124764611 | 0.26848 | 0 |
| GH03108 | FBgn0027599 | CG2790 | 0.124731488 | 0.28391388 | 0 |
| LD40216 | FBgn0278604 | dmt | -0.124730887 | 0.27309497 | 0 |

| | | | | | |
|---|---|---|---|---|---|
| GH10422 | FBgn0036237 | viaf | -0.124703654 | 0.25698584 | 0 |
| LD31537 | FBgn0031820 | Daxx | -0.124672009 | 0.32826856 | 0 |
| GH08848 | FBgn0040298 | Myt1 | -0.124662868 | 0.30786949 | 0 |
| GM08694 | FBgn0086408 | stl | 0.124567329 | 0.42487759 | 0 |
| LD24347 | FBgn0029629 | eIF3g1 | 0.124456529 | 0.2918756 | 0 |
| LD24478 | FBgn0030973 | CG7332 | 0.124411502 | 0.2677426 | 0 |
| SD22430 | FBgn0031389 | CG4259 | 0.124408563 | 0.43578782 | 0 |
| AT12172 | FBgn0264844 | asRNA:CR44052 | 0.124405223 | 0.38667402 | 0 |
| LP03829 | FBgn0036945 | Ssk | 0.124391888 | 0.71775913 | 0 |
| GH03475 | FBgn0038771 | CG4390 | 0.124361398 | 0.35150163 | 0 |
| HL02049 | FBgn0004516 | Gad1 | -0.124355904 | 0.35298275 | 0 |
| RE22928 | FBgn0030669 | CG9240 | 0.124275195 | 0.29393334 | 0 |
| RE53486 | FBgn0030788 | Sap30 | 0.124243985 | 0.28320138 | 0 |
| LD35502 | FBgn0030091 | CG7065 | 0.124236796 | 0.25304991 | 0 |
| GH01562 | FBgn0037956 | CG6959 | 0.124233485 | 0.4139621 | 0 |
| GH09868 | FBgn0025790 | TBPH | 0.124196679 | 0.33183546 | 0 |
| SD05887 | FBgn0034854 | Golgin245 | 0.124148219 | 0.26083523 | 0 |
| RH13158 | FBgn0011604 | Iswi | -0.124137321 | 0.29338401 | 0 |
| LD38352 | FBgn0263143 | vret | -0.124099228 | 0.33987527 | 0 |
| RE68504 | FBgn0037770 | Art4 | -0.124090813 | 0.25439475 | 0 |
| RE32936 | FBgn0036282 | Smyd4-2 | 0.124087373 | 0.4420803 | 0 |
| RE25030 | FBgn0011818 | oaf | -0.124054536 | 0.38788796 | 0 |
| RE54270 | FBgn0038319 | mRpL9 | -0.124051414 | 0.30830485 | 0 |
| RE33542 | FBgn0035921 | CG13305 | -0.123964803 | 0.31605389 | 0 |
| SD04572 | FBgn0016976 | stnA | 0.123927058 | 0.28299558 | 0 |
| SD04572 | FBgn0016975 | stnB | 0.123927058 | 0.28299558 | 0 |
| AT31705 | FBgn0033317 | CG8635 | -0.123903343 | 0.43004772 | 0 |
| SD02021 | FBgn0035588 | CG10672 | -0.123841399 | 0.29066719 | 0 |
| LD24461 | FBgn0027951 | MTA1-like | 0.123778198 | 0.26011542 | 0 |
| AT29119 | FBgn0051803 | CG31803 | 0.123731144 | 0.39903152 | 0 |
| LD30683 | FBgn0032214 | CG4968 | -0.123650312 | 0.3793285 | 0 |
| LD05256 | FBgn0034371 | SP2637 | 0.123649135 | 0.33227921 | 0 |
| LD05057 | FBgn0263602 | Tasp1 | 0.123639638 | 0.26814996 | 0 |
| AT23634 | FBgn0038453 | Arl6IP1 | 0.123539706 | 0.25818104 | 0 |
| LD22119 | FBgn0051729 | CG31729 | 0.123531736 | 0.25348051 | 0 |
| SD05291 | FBgn0030993 | Mec2 | 0.123520329 | 0.47568595 | 0 |
| RE04404 | FBgn0033072 | CG17994 | 0.123405506 | 0.25455646 | 0 |
| LD13761 | FBgn0031868 | Rat1 | 0.123401511 | 0.26188951 | 0 |
| AT02481 | FBgn0052572 | CG32572 | 0.123336901 | 0.29658578 | 0 |
| LD48015 | FBgn0035945 | CG5026 | 0.123312099 | 0.25897591 | 0 |
| GH12182 | FBgn0032797 | Hasp | 0.123244571 | 0.56750708 | 0 |
| GH24292 | FBgn0019972 | Drice | 0.123226289 | 0.28758766 | 0 |
| LD26991 | FBgn0034674 | CG9304 | 0.12316333 | 0.26514466 | 0 |
| LD45641 | FBgn0004169 | up | 0.123136222 | 0.28754289 | 0 |
| GH13492 | FBgn0259244 | CG42342 | -0.123128384 | 0.56767477 | 0 |
| GH08269 | FBgn0041775 | tral | -0.123038033 | 0.35619541 | 0 |
| RE24343 | FBgn0052843 | Dh31-R | -0.122985932 | 0.29671934 | 0 |
| RE66587 | FBgn0039161 | CG13606 | -0.122898243 | 0.29056081 | 0 |

| | | | | | |
|---|---|---|---|---|---|
| LD27203 | FBgn0050101 | CG30101 | -0.122860517 | 0.32224933 | 0 |
| RH63170 | FBgn0085195 | CG34166 | 0.12285884 | 0.26248252 | 0 |
| AT18054 | FBgn0040755 | CG17580 | 0.122690476 | 0.62869413 | 0 |
| LD24835 | FBgn0036980 | RhoBTB | 0.122638321 | 0.34271691 | 0 |
| GH13040 | FBgn0265512 | mlt | -0.122619115 | 0.31581229 | 0 |
| LD25907 | FBgn0002973 | numb | 0.122608643 | 0.29319322 | 0 |
| RE36324 | FBgn0004400 | rhi | 0.122588901 | 0.46742542 | 0 |
| LD45195 | FBgn0035370 | Non2 | 0.122555279 | 0.34777011 | 0 |
| GM02293 | FBgn0024814 | Clc | 0.122533337 | 0.34975848 | 0 |
| GM14222 | FBgn0086712 | Egm | 0.122458015 | 0.29332909 | 0 |
| LD32354 | FBgn0025676 | CkIIalpha-i3 | 0.122442459 | 0.28812883 | 0 |
| RH28890 | FBgn0033983 | ADPS | -0.122328789 | 0.33482596 | 0 |
| AT05453 | FBgn0001084 | fy | -0.122249334 | 0.26375792 | 0 |
| RH03791 | FBgn0031318 | CG4887 | 0.122209976 | 0.29332909 | 0 |
| RE57116 | FBgn0036108 | Cpr67Fa1 | -0.122199701 | 0.27578495 | 0 |
| RE57116 | FBgn0036109 | Cpr67Fa2 | -0.122199701 | 0.27578495 | 0 |
| GM06026 | FBgn0037074 | CG7324 | 0.122173637 | 0.28754289 | 0 |
| AT07549 | FBgn0029993 | CG1571 | 0.122149818 | 0.57261017 | 0 |
| AT26009 | FBgn0035196 | CG9129 | 0.122140866 | 0.29335128 | 0 |
| RE30177 | FBgn0028931 | CG16863 | -0.12208378 | 0.36642713 | 0 |
| LD36116 | FBgn0003656 | sws | 0.122073367 | 0.29056081 | 0 |
| LD13504 | FBgn0267585 | lncRNA:CR45923 | 0.122018739 | 0.44813254 | 0 |
| LP09584 | FBgn0032075 | Tsp29Fb | 0.122013271 | 0.59783502 | 0 |
| SD02269 | FBgn0021796 | Tor | 0.121992481 | 0.37804643 | 0 |
| LD12127 | FBgn0050497 | CG30497 | -0.121989011 | 0.66120595 | 0 |
| LD12127 | FBgn0266526 | CG45093 | -0.121989011 | 0.66120595 | 0 |
| LP04536 | FBgn0051781 | lncRNA:CR31781 | -0.121982281 | 0.54155484 | 0 |
| RH45008 | FBgn0040931 | CG9034 | 0.121969516 | 0.39564083 | 0 |
| LD24340 | FBgn0030551 | CG11674 | -0.121965413 | 0.36161041 | 0 |
| LD28427 | FBgn0000447 | Dhod | 0.12191079 | 0.29168881 | 0 |
| AT09390 | FBgn0032503 | CG16825 | -0.121871415 | 0.38902664 | 0 |
| LD40680 | FBgn0050467 | CG30467 | 0.121824089 | 0.32681987 | 0 |
| SD02332 | FBgn0050382 | CG30382 | -0.12173434 | 0.3066096 | 0 |
| SD02332 | FBgn0263121 | Prosalpha1 | -0.12173434 | 0.3066096 | 0 |
| GH04950 | FBgn0265194 | Trpm | 0.121696187 | 0.27903828 | 0 |
| GM03761 | FBgn0263396 | sqd | 0.121672797 | 0.59982904 | 0 |
| RE30146 | FBgn0035104 | CG13875 | 0.121669801 | 0.26307647 | 0 |
| RH06462 | FBgn0035726 | CG9953 | 0.121644193 | 0.34731796 | 0 |
| LD16982 | FBgn0259734 | Nost | -0.121633001 | 0.26375028 | 0 |
| GH03720 | FBgn0262473 | Tl | 0.12151057 | 0.28192759 | 0 |
| RE33150 | FBgn0034520 | lms | -0.121503484 | 0.30456668 | 0 |
| LD19131 | FBgn0030710 | CG8924 | 0.12148495 | 0.26853246 | 0 |
| GH11566 | FBgn0023529 | CG2918 | -0.121349411 | 0.28208194 | 0 |
| SD25413 | FBgn0011573 | Cdc37 | 0.121342198 | 0.27703693 | 0 |
| RE63280 | FBgn0038660 | CG14291 | 0.121315319 | 0.36866368 | 0 |
| GH04687 | FBgn0061359 | CG33671 | -0.121315029 | 0.32332024 | 0 |
| GH04687 | FBgn0061360 | CG33672 | -0.121315029 | 0.32332024 | 0 |
| GH14865 | FBgn0031702 | fusl | -0.121257805 | 0.31989649 | 0 |

| | | | | | |
|---|---|---|---|---|---|
| RE74001 | FBgn0026431 | Grip75 | 0.121238111 | 0.36203856 | 0 |
| AT20011 | FBgn0036652 | CG13032 | 0.121213901 | 0.41331428 | 0 |
| LD43225 | FBgn0036249 | CG11560 | 0.121181906 | 0.31511681 | 0 |
| LD43460 | FBgn0026761 | Trap1 | -0.121158738 | 0.29410644 | 0 |
| SD04106 | FBgn0010380 | AP-1-2beta | 0.121109015 | 0.26604496 | 0 |
| RE23081 | FBgn0250843 | Prosalpha6 | 0.120998795 | 0.3505826 | 0 |
| RE20019 | FBgn0039232 | sosie | -0.120984717 | 0.55177617 | 0 |
| GH02051 | FBgn0003889 | betaTub85D | 0.120982501 | 0.49579399 | 0 |
| SD17025 | FBgn0037071 | CG7632 | -0.12095051 | 0.28383357 | 0 |
| LD23434 | FBgn0036685 | CG6664 | -0.120947381 | 0.29168881 | 0 |
| LP06780 | FBgn0032973 | CG6675 | -0.120937341 | 0.50815974 | 0 |
| RE11886 | FBgn0015790 | Rab11 | 0.12093039 | 0.26615408 | 0 |
| LD22716 | FBgn0036846 | MESR6 | 0.120836133 | 0.26397981 | 0 |
| LD03509 | FBgn0000611 | exd | 0.120834959 | 0.32345633 | 0 |
| RE08710 | FBgn0037678 | CG16749 | 0.120790537 | 0.61374063 | 0 |
| LD42063 | FBgn0030030 | CG1636 | 0.12065137 | 0.2767948 | 0 |
| RE27192 | FBgn0086680 | vvl | -0.12064918 | 0.27904632 | 0 |
| RH66343 | FBgn0033672 | rho-7 | -0.120581634 | 0.26935415 | 0 |
| LD38634 | FBgn0037544 | CG11035 | 0.120566068 | 0.41957824 | 0 |
| LD39082 | FBgn0032729 | L2HGDH | 0.120538492 | 0.30869977 | 0 |
| RE54409 | FBgn0038307 | mRpS10 | -0.120480376 | 0.36035079 | 0 |
| LD03052 | FBgn0030838 | CG5445 | 0.120443789 | 0.27903828 | 0 |
| RH04924 | FBgn0086348 | se | -0.120427861 | 0.28997789 | 0 |
| AT24862 | FBgn0033113 | Spn42Dc | 0.120241028 | 0.35796898 | 0 |
| GM14481 | FBgn0030241 | feo | -0.120238399 | 0.34494598 | 0 |
| LD31969 | FBgn0034418 | CG15118 | -0.120108098 | 0.26834543 | 0 |
| LD46221 | FBgn0034312 | CG10916 | 0.120031684 | 0.36983196 | 0 |
| GH06451 | FBgn0086443 | AsnRS | -0.119973345 | 0.32054678 | 0 |
| LD47944 | FBgn0035641 | CG5568 | -0.119954435 | 0.35079094 | 0 |
| LP12157 | FBgn0283451 | br | 0.119932283 | 0.32290021 | 0 |
| LD46604 | FBgn0036801 | MYPT-75D | -0.119882505 | 0.30975761 | 0 |
| LD18960 | FBgn0034300 | CG5098 | 0.119871999 | 0.36934556 | 0 |
| RE70528 | FBgn0039237 | CG13640 | -0.119861875 | 0.31912746 | 0 |
| LP03218 | FBgn0034990 | CG11406 | -0.119811292 | 0.39664352 | 0 |
| AT13740 | FBgn0037454 | CG1137 | 0.119740409 | 0.34598163 | 0 |
| GH21865 | FBgn0035709 | eIF4E4 | -0.11972755 | 0.29135153 | 0 |
| RE19540 | FBgn0015754 | Lis-1 | -0.119649873 | 0.26463547 | 0 |
| GH26995 | FBgn0040832 | CG8012 | 0.119568019 | 0.31861085 | 0 |
| AT28705 | FBgn0031951 | r2d2 | 0.11954373 | 0.3218739 | 0 |
| RH04586 | FBgn0038681 | Cyp12a4 | -0.119536393 | 0.30755546 | 0 |
| LD24633 | FBgn0250746 | Prosbeta7 | 0.119522676 | 0.33866194 | 0 |
| HL01517 | FBgn0034876 | wmd | 0.119407391 | 0.28674941 | 0 |
| GH19061 | FBgn0033339 | Sec31 | 0.119403718 | 0.31110387 | 0 |
| LD44914 | FBgn0039854 | CG1635 | 0.119398763 | 0.44088664 | 0 |
| LD22945 | FBgn0265189 | sud1 | -0.119386563 | 0.31049391 | 0 |
| GH02222 | FBgn0033320 | CG8586 | 0.119375893 | 0.38772065 | 0 |
| LD46082 | FBgn0020224 | Cbl | 0.119370859 | 0.2671491 | 0 |
| AT17674 | FBgn0032175 | CG13131 | -0.119254462 | 0.40705659 | 0 |

| | | | | | |
|---|---|---|---|---|---|
| LD13525 | FBgn0040823 | dpr6 | 0.119242359 | 0.36612989 | 0 |
| RH49745 | FBgn0027836 | Dgp-1 | 0.119145663 | 0.38855327 | 0 |
| LD12946 | FBgn0030703 | MSBP | -0.119111839 | 0.40333031 | 0 |
| AT26738 | FBgn0053208 | Mical | -0.119101375 | 0.53045256 | 0 |
| RE06553 | FBgn0034382 | CG18609 | 0.119094672 | 0.53139066 | 0 |
| LD42828 | FBgn0010356 | Taf5 | 0.119079795 | 0.39605958 | 0 |
| LD45235 | FBgn0003257 | r-l | 0.119010472 | 0.30092567 | 0 |
| AT18160 | FBgn0034527 | CG9945 | -0.118986988 | 0.31646308 | 0 |
| RE68460 | FBgn0000061 | al | -0.118969978 | 0.29914002 | 0 |
| RE48727 | FBgn0039003 | wfs1 | -0.118968357 | 0.31570137 | 0 |
| LD43055 | FBgn0037814 | CG6325 | -0.118916796 | 0.37559906 | 0 |
| AT29114 | FBgn0036036 | CG14174 | -0.1189028 | 0.3505826 | 0 |
| RH56266 | FBgn0039584 | beat-VI | -0.118895378 | 0.30301105 | 0 |
| RE17417 | FBgn0036891 | CG9372 | 0.118866096 | 0.33074938 | 0 |
| RE22235 | FBgn0261647 | Axud1 | 0.118798301 | 0.27423935 | 0 |
| SD01727 | FBgn0039252 | CG11771 | 0.118747956 | 0.30443808 | 0 |
| AT31732 | FBgn0038219 | CG14839 | 0.118731527 | 0.30663023 | 0 |
| GM13517 | FBgn0036691 | beg | 0.118719751 | 0.42306953 | 0 |
| GH02020 | FBgn0035734 | CG14823 | -0.118716288 | 0.2976745 | 0 |
| AT24407 | FBgn0051974 | CG31974 | 0.118608416 | 0.61605959 | 0 |
| LD28410 | FBgn0027342 | fz4 | -0.118583064 | 0.33147651 | 0 |
| GM12182 | FBgn0039687 | Naa40 | -0.118546341 | 0.3883281 | 0 |
| LD34217 | FBgn0010333 | Rac1 | 0.118515067 | 0.26675152 | 0 |
| GH27470 | FBgn0039205 | CG13623 | -0.118500128 | 0.48113099 | 0 |
| RE53745 | FBgn0028708 | Mst85C | 0.11848744 | 0.29119748 | 0 |
| LD46502 | FBgn0022984 | qkr58E-3 | 0.118456927 | 0.29410644 | 0 |
| RE62692 | FBgn0033814 | CG4670 | -0.11844562 | 0.71953576 | 0 |
| RE34144 | FBgn0267384 | Ubc7 | 0.118405214 | 0.43875144 | 0 |
| LD22971 | FBgn0038316 | CG6276 | -0.118404646 | 0.32355904 | 0 |
| LP07188 | FBgn0034240 | MESR4 | 0.11840013 | 0.2767948 | 0 |
| LD02105 | FBgn0043458 | CG12084 | 0.118397904 | 0.40644068 | 0 |
| RE29404 | FBgn0051146 | Nlg1 | -0.11834921 | 0.52205335 | 0 |
| LD34766 | FBgn0010422 | TfIIS | 0.11833536 | 0.34170455 | 0 |
| AT02241 | FBgn0004406 | tam | 0.118304904 | 0.36532391 | 0 |
| LD35329 | FBgn0264959 | Src42A | 0.118276297 | 0.29250904 | 0 |
| AT08349 | FBgn0264560 | garz | -0.118212838 | 0.26967233 | 0 |
| LD27878 | FBgn0023416 | Ac3 | 0.11813506 | 0.28621512 | 0 |
| GH16672 | FBgn0025680 | cry | -0.118127804 | 0.375197 | 0 |
| GH09808 | FBgn0051523 | CG31523 | 0.118079357 | 0.28754289 | 0 |
| RE15976 | FBgn0034201 | CG17290 | -0.118071744 | 0.4235086 | 0 |
| RE72677 | FBgn0038439 | Cad89D | -0.117992622 | 0.34994022 | 0 |
| LD22943 | FBgn0002914 | Myb | 0.117889125 | 0.28669623 | 0 |
| RE09630 | FBgn0039131 | CG12268 | -0.117776539 | 0.30296611 | 0 |
| RE18318 | FBgn0034389 | Mctp | -0.117767645 | 0.29792775 | 0 |
| LD23646 | FBgn0023129 | aay | -0.117740656 | 0.41649923 | 0 |
| LD02975 | FBgn0030672 | CG9281 | -0.117704616 | 0.28967916 | 0 |
| RE59626 | FBgn0037972 | CG10005 | 0.117699781 | 0.72734227 | 0 |
| RE02671 | FBgn0003480 | spn-B | -0.117684039 | 0.45317518 | 0 |

| | | | | | |
|---|---|---|---|---|---|
| GH15472 | FBgn0030717 | CG8931 | 0.117603 | 0.31217459 | 0 |
| GM14238 | FBgn0038964 | Nop56 | 0.117544372 | 0.35619541 | 0 |
| LD34905 | FBgn0029763 | Usp16-45 | 0.117445056 | 0.27929923 | 0 |
| RE39992 | FBgn0031432 | Cyp309a1 | 0.117433122 | 0.38902664 | 0 |
| RE53354 | FBgn0013749 | Arf102F | -0.11742375 | 0.28911598 | 0 |
| LD24449 | FBgn0001970 | Pgant35A | 0.11739634 | 0.36885296 | 0 |
| LD11447 | FBgn0004611 | Plc21C | 0.117386122 | 0.29410644 | 0 |
| LP11057 | FBgn0052302 | CG32302 | 0.117381355 | 0.33254606 | 0 |
| SD24901 | FBgn0035059 | CG3894 | -0.117332757 | 0.29578352 | 0 |
| LD46868 | FBgn0034046 | tun | 0.117331833 | 0.42280478 | 0 |
| SD19826 | FBgn0039487 | gb | 0.117265969 | 0.36994545 | 0 |
| LD29226 | FBgn0034598 | CG4266 | 0.117257834 | 0.31586471 | 0 |
| RE32966 | FBgn0039420 | CG6154 | -0.117218427 | 0.29007656 | 0 |
| LD05615 | FBgn0037205 | BoYb | 0.117153985 | 0.2766162 | 0 |
| LD34582 | FBgn0051961 | TBCC | 0.117077114 | 0.31912746 | 0 |
| AT19932 | FBgn0029877 | CG17717 | -0.117034941 | 0.36033739 | 0 |
| LD43688 | FBgn0053113 | Rtnl1 | 0.116994924 | 0.27815421 | 0 |
| RE67565 | FBgn0040382 | CG5273 | 0.116956653 | 0.28615679 | 0 |
| RE08220 | FBgn0033130 | Tsp42Ei | -0.116907495 | 0.58625109 | 0 |
| RE33114 | FBgn0064225 | RpL5 | -0.116850138 | 0.32437253 | 0 |
| RE62842 | FBgn0266724 | Trs20 | -0.116790163 | 0.34270683 | 0 |
| RE07882 | FBgn0037428 | Osi18 | -0.11676232 | 0.47474134 | 0 |
| RH02452 | FBgn0000715 | FMRFa | -0.116728157 | 0.37294475 | 0 |
| LD15891 | FBgn0003016 | osp | 0.11672682 | 0.38666161 | 0 |
| LD45365 | FBgn0040294 | POSH | -0.116703857 | 0.32362322 | 0 |
| AT24194 | FBgn0011286 | RyR | 0.116686552 | 0.74274998 | 0 |
| RE33778 | FBgn0039970 | CG17508 | -0.116670297 | 0.58332023 | 0 |
| GH04125 | FBgn0028988 | Spn42Dd | 0.116668894 | 0.39339733 | 0 |
| RE16970 | FBgn0037470 | Tailor | 0.116662601 | 0.39197671 | 0 |
| LD12605 | FBgn0040001 | FASN3 | -0.116653152 | 0.64364316 | 0 |
| AT24116 | FBgn0030280 | CG11106 | -0.116644745 | 0.41242951 | 0 |
| RE13587 | FBgn0034008 | CG8152 | -0.116638865 | 0.335791 | 0 |
| LD45403 | FBgn0002044 | swm | 0.116498685 | 0.38251295 | 0 |
| LD34222 | FBgn0030358 | CG10362 | 0.116457092 | 0.45026411 | 0 |
| SD03142 | FBgn0011297 | l(2)not | -0.116422263 | 0.46172055 | 0 |
| LD48010 | FBgn0037696 | GstZ1 | 0.11637825 | 0.31677144 | 0 |
| RH67033 | FBgn0034328 | IM23 | -0.116308379 | 0.45433285 | 0 |
| RH44637 | FBgn0034877 | levy | -0.116284831 | 0.29331016 | 0 |
| LD15729 | FBgn0024330 | MED6 | 0.116279246 | 0.32095126 | 0 |
| RE63441 | FBgn0027055 | CSN3 | 0.116266408 | 0.35183241 | 0 |
| LD04179 | FBgn0036827 | CG6843 | 0.116261518 | 0.3547438 | 0 |
| SD05105 | FBgn0015279 | Pi3K92E | -0.116190796 | 0.29289815 | 0 |
| LD18435 | FBgn0011725 | twin | -0.116189027 | 0.61004629 | 0 |
| LD20892 | FBgn0033402 | Myd88 | 0.116165692 | 0.2976745 | 0 |
| GM01793 | FBgn0037156 | CG11523 | -0.116145771 | 0.30755546 | 0 |
| SD05472 | FBgn0014073 | Tie | -0.116073083 | 0.59726688 | 0 |
| LD23816 | FBgn0267378 | sau | 0.116066415 | 0.375197 | 0 |
| RE20929 | FBgn0262699 | fzr | -0.116054079 | 0.28912309 | 0 |

| | | | | | |
|---|---|---|---|---|---|
| LD44376 | FBgn0027085 | LeuRS-m | -0.116030491 | 0.35009244 | 0 |
| RE53127 | FBgn0038132 | CG15887 | -0.116023072 | 0.40617197 | 0 |
| GH01619 | FBgn0014009 | Rab2 | 0.116011081 | 0.36935223 | 0 |
| LD31216 | FBgn0037293 | CG12007 | -0.115967912 | 0.33866194 | 0 |
| LD12439 | FBgn0039710 | dgt1 | 0.115958738 | 0.31701299 | 0 |
| LD14533 | FBgn0035211 | CG2211 | -0.115924111 | 0.27904632 | 0 |
| AT04225 | FBgn0003310 | S | -0.115887461 | 0.36971494 | 0 |
| RE04933 | FBgn0010247 | Parp | -0.115866215 | 0.38135654 | 0 |
| RE42756 | FBgn0035526 | CG1316 | -0.115854751 | 0.36263426 | 0 |
| RE35615 | FBgn0037122 | CG14570 | 0.115831558 | 0.27904632 | 0 |
| LD08644 | FBgn0030834 | CG8675 | 0.115804906 | 0.28251594 | 0 |
| GH07831 | FBgn0031779 | CG9175 | 0.115786374 | 0.30224264 | 0 |
| AT04446 | FBgn0024912 | agt | 0.115783148 | 0.36161041 | 0 |
| RE53955 | FBgn0003205 | Ras85D | 0.115765277 | 0.42026159 | 0 |
| LD34262 | FBgn0011207 | pelo | -0.115747191 | 0.28182098 | 0 |
| LD46404 | FBgn0035393 | CG16753 | 0.115733086 | 0.32095126 | 0 |
| RH39663 | FBgn0261602 | RpL8 | 0.115731977 | 0.47263843 | 0 |
| RE16761 | FBgn0029690 | CG6414 | -0.115699382 | 0.28128819 | 0 |
| RE17845 | FBgn0032782 | Rab9 | -0.115690994 | 0.3034516 | 0 |
| SD08189 | FBgn0039525 | CG5646 | -0.115642151 | 0.2975402 | 0 |
| RE04347 | FBgn0087035 | AGO2 | -0.115588995 | 0.28450931 | 0 |
| GH25818 | FBgn0036497 | Ran-like | 0.11558723 | 0.39552963 | 0 |
| GH23636 | FBgn0037998 | Cog1 | -0.115581676 | 0.36847667 | 0 |
| GH20022 | FBgn0027081 | ThrRS | 0.115549537 | 0.2843977 | 0 |
| RE12410 | FBgn0034295 | CG10911 | -0.11554918 | 0.38114458 | 0 |
| RE29225 | FBgn0001150 | gt | -0.115548627 | 0.38571465 | 0 |
| RH65975 | FBgn0015298 | Srp19 | -0.115354176 | 0.29332909 | 0 |
| RH62264 | FBgn0034485 | CG11099 | 0.115302996 | 0.38996816 | 0 |
| GH14621 | FBgn0031808 | CG9507 | -0.115274296 | 0.32887229 | 0 |
| AT07541 | FBgn0040333 | brv3 | -0.115202964 | 0.32742498 | 0 |
| RE14381 | FBgn0267365 | l(2)SH0834 | 0.115180656 | 0.2975402 | 0 |
| LD18893 | FBgn0260945 | Atg1 | 0.1151327 | 0.2848657 | 0 |
| GH19222 | FBgn0037512 | CG2616 | 0.115120243 | 0.35710349 | 0 |
| LD41392 | FBgn0020018 | Ppox | 0.115091175 | 0.27851375 | 0 |
| RH56796 | FBgn0035179 | CG12038 | 0.115080706 | 0.60309107 | 0 |
| GM01838 | FBgn0011336 | Stt3B | 0.115020812 | 0.39660162 | 0 |
| RH04334 | FBgn0032538 | CG16885 | 0.115014765 | 0.30262017 | 0 |
| AT01495 | FBgn0030661 | CG8105 | 0.114973298 | 0.39662446 | 0 |
| RE39515 | FBgn0047114 | CG31142 | -0.114876418 | 0.30878207 | 0 |
| RE70412 | FBgn0030617 | CG9095 | 0.114872278 | 0.56705417 | 0 |
| LD03446 | FBgn0027932 | Akap200 | -0.114866298 | 0.6944496 | 0 |
| SD05640 | FBgn0014141 | cher | -0.11485297 | 0.39157917 | 0 |
| SD09114 | FBgn0030501 | BthD | 0.114819311 | 0.34777011 | 0 |
| RH08458 | FBgn0031940 | CG7214 | -0.114793597 | 0.36885296 | 0 |
| LP05330 | FBgn0032187 | CG4839 | -0.114782852 | 0.39339733 | 0 |
| LD14125 | FBgn0027548 | nito | 0.114734574 | 0.29790263 | 0 |
| RE06930 | FBgn0030794 | CG13005 | -0.114674588 | 0.28703489 | 0 |
| LD32555 | FBgn0037670 | Ibf1 | 0.114652087 | 0.33728419 | 0 |

| | | | | | |
|---|---|---|---|---|---|
| LD21432 | FBgn0035601 | Uev1A | -0.114627216 | 0.39339733 | 0 |
| RE70039 | FBgn0030900 | Him | -0.114598329 | 0.51140355 | 0 |
| RE20784 | FBgn0030465 | CG15743 | 0.114547542 | 0.31164827 | 0 |
| SD02555 | FBgn0031622 | CG3251 | -0.114491661 | 0.43026786 | 0 |
| LP08352 | FBgn0039705 | Atg16 | 0.114362995 | 0.29099529 | 0 |
| GH28004 | FBgn0039722 | Capa | 0.11434534 | 0.50184892 | 0 |
| LD23604 | FBgn0030943 | Nup35 | 0.114314541 | 0.45015388 | 0 |
| AT31060 | FBgn0037462 | sunz | 0.114308302 | 0.30657727 | 0 |
| LD24702 | FBgn0027500 | spd-2 | -0.114291784 | 0.2958322 | 0 |
| GH01837 | FBgn0038878 | CG3301 | 0.114290724 | 0.53326482 | 0 |
| AT19513 | FBgn0015221 | Fer2LCH | 0.114244357 | 0.29168881 | 0 |
| RE74917 | FBgn0261575 | tobi | 0.114230365 | 0.37093907 | 0 |
| SD08161 | FBgn0036640 | nxf2 | 0.114213616 | 0.29096007 | 0 |
| RH14061 | FBgn0259110 | mmd | -0.114180324 | 0.31253644 | 0 |
| RH48239 | FBgn0031437 | Arpc5 | -0.11418001 | 0.35175496 | 0 |
| HL01018 | FBgn0263200 | Galt | 0.114151914 | 0.37872757 | 0 |
| RE54443 | FBgn0052767 | CG32767 | 0.114081025 | 0.31511681 | 0 |
| RE28280 | FBgn0032284 | CG7294 | 0.114031089 | 0.29671934 | 0 |
| RE27547 | FBgn0044011 | Spn43Ad | -0.114007776 | 0.36532391 | 0 |
| RE65121 | FBgn0019662 | qm | 0.113978688 | 0.3246684 | 0 |
| LD03212 | FBgn0034688 | CG11474 | 0.113923759 | 0.28683795 | 0 |
| LD11117 | FBgn0051092 | LpR2 | 0.113914019 | 0.28500043 | 0 |
| RH29440 | FBgn0261792 | snRNP-U1-C | 0.113888256 | 0.32588723 | 0 |
| LD37788 | FBgn0045862 | btz | -0.113795514 | 0.32567485 | 0 |
| LP04163 | FBgn0051472 | sgll | -0.11370676 | 0.31825117 | 0 |
| GH25793 | FBgn0243516 | Vrp1 | 0.113673529 | 0.32296998 | 0 |
| GH08002 | FBgn0261714 | Cpn | -0.113647625 | 0.29328557 | 0 |
| RH57193 | FBgn0052214 | CG32214 | -0.113623808 | 0.34240724 | 0 |
| RH57193 | FBgn0062928 | hpRNA:CR33940 | -0.113623808 | 0.34240724 | 0 |
| LD16082 | FBgn0026376 | Rgl | 0.113623802 | 0.28320138 | 0 |
| LD25139 | FBgn0033304 | Cyp6a13 | 0.113597207 | 0.31840362 | 0 |
| SD05126 | FBgn0033783 | CG17019 | -0.113573974 | 0.29250904 | 0 |
| SD01276 | FBgn0003638 | su(w[a]) | 0.113538138 | 0.32172684 | 0 |
| SD01276 | FBgn0266569 | CG42259 | 0.113538138 | 0.32172684 | 0 |
| AT25429 | FBgn0039329 | CG10669 | -0.113526635 | 0.38451222 | 0 |
| RE40275 | FBgn0039355 | CG4730 | 0.113514426 | 0.47322827 | 0 |
| HL08052 | FBgn0002641 | mal | 0.113489789 | 0.29448305 | 0 |
| RE67917 | FBgn0035679 | CG10467 | 0.113434078 | 0.35183241 | 0 |
| RE67917 | FBgn0035678 | CG10469 | 0.113434078 | 0.35183241 | 0 |
| RE39606 | FBgn0011305 | Rsf1 | 0.113370797 | 0.33998407 | 0 |
| RE68569 | FBgn0058006 | CG40006 | -0.113364097 | 0.35959979 | 0 |
| LP12057 | FBgn0028686 | Rpt3 | -0.113341357 | 0.31811714 | 0 |
| LP01645 | FBgn0002531 | Lcp1 | 0.113329435 | 0.48920218 | 0 |
| RE08694 | FBgn0003256 | rl | -0.113327867 | 0.46030056 | 0 |
| RE50040 | FBgn0039160 | CG5510 | 0.113305638 | 0.38995622 | 0 |
| LD22396 | FBgn0000257 | car | 0.113135459 | 0.30470968 | 0 |
| GM05702 | FBgn0039689 | CIA30 | -0.113092042 | 0.2908404 | 0 |
| AT04231 | FBgn0032474 | DnaJ-H | -0.113038225 | 0.43839377 | 0 |

| | | | | | |
|---|---|---|---|---|---|
| AT22075 | FBgn0031517 | CG15406 | 0.113029554 | 0.48013701 | 0 |
| LD46621 | FBgn0040235 | c12.1 | 0.112991016 | 0.29168881 | 0 |
| LD46023 | FBgn0032262 | CG7384 | 0.112958398 | 0.44214976 | 0 |
| LD35207 | FBgn0033166 | Eaf | 0.112942908 | 0.28703489 | 0 |
| AT19436 | FBgn0023171 | rnh1 | 0.11290615 | 0.32271909 | 0 |
| SD04165 | FBgn0261383 | IntS6 | 0.112840823 | 0.31905024 | 0 |
| HL02936 | FBgn0030345 | CG1847 | 0.112797035 | 0.29168881 | 0 |
| LD05080 | FBgn0030240 | CG2202 | -0.112731848 | 0.35360544 | 0 |
| AT15112 | FBgn0286075 | DNAlig3 | 0.112713583 | 0.29033759 | 0 |
| RE35772 | FBgn0016978 | snRNP-U1-70K | -0.112668513 | 0.38276825 | 0 |
| LD24034 | FBgn0035205 | Ctr9 | 0.112655669 | 0.3155453 | 0 |
| LD09947 | FBgn0000083 | AnxB9 | 0.112640993 | 0.31912746 | 0 |
| LD02291 | FBgn0040513 | kappaB-Ras | 0.112636326 | 0.30080363 | 0 |
| GH21880 | FBgn0032618 | CG31743 | -0.112568088 | 0.37832142 | 0 |
| GH21880 | FBgn0085342 | CG34313 | -0.112568088 | 0.37832142 | 0 |
| LD20519 | FBgn0030653 | CG7860 | 0.112510222 | 0.31566227 | 0 |
| LD18443 | FBgn0035270 | CG13933 | -0.112500923 | 0.3066096 | 0 |
| LD06593 | FBgn0032084 | CG13101 | -0.112460819 | 0.38251436 | 0 |
| LD40495 | FBgn0032216 | Usp14 | -0.112428483 | 0.32418356 | 0 |
| GH27383 | FBgn0037676 | CG8861 | 0.112427725 | 0.33908067 | 0 |
| LD46437 | FBgn0261270 | SelD | 0.112405847 | 0.39602344 | 0 |
| RE09303 | FBgn0013433 | beat-Ia | -0.112396778 | 0.7508361 | 0 |
| GM01994 | FBgn0027335 | Rip11 | -0.112354054 | 0.34663191 | 0 |
| RE62779 | FBgn0022700 | Cht4 | 0.112325615 | 0.35573779 | 0 |
| RE14453 | FBgn0014184 | Oda | 0.112325511 | 0.43814365 | 0 |
| GH05039 | FBgn0000711 | flw | 0.112318052 | 0.36724764 | 0 |
| RH38110 | FBgn0039666 | Diedel | -0.112278409 | 0.34904751 | 0 |
| SD06366 | FBgn0085443 | spri | -0.112260488 | 0.36870325 | 0 |
| LD17773 | FBgn0036423 | CG3919 | 0.112230835 | 0.34663191 | 0 |
| RE43130 | FBgn0038130 | CG8630 | -0.112140543 | 0.42701482 | 0 |
| HL01702 | FBgn0028743 | Dhit | 0.112083563 | 0.57813855 | 0 |
| RE21618 | FBgn0014857 | His3.3A | 0.111986281 | 0.46030056 | 0 |
| SD08653 | FBgn0004132 | boca | 0.111896338 | 0.36212941 | 0 |
| LD30316 | FBgn0030354 | Upf1 | -0.111813477 | 0.37294475 | 0 |
| LD45559 | FBgn0035398 | Cht7 | 0.111797527 | 0.58742676 | 0 |
| RH01578 | FBgn0035510 | Cpr64Aa | -0.111767014 | 0.35347549 | 0 |
| LD46305 | FBgn0030813 | CG4949 | -0.11155635 | 0.30194797 | 0 |
| GM01555 | FBgn0004908 | Arl2 | -0.111541121 | 0.36870325 | 0 |
| LD40728 | FBgn0032256 | RluA-2 | 0.111511691 | 0.31566227 | 0 |
| AT12403 | FBgn0029702 | CG15572 | -0.111445804 | 0.33795166 | 0 |
| SD16537 | FBgn0031006 | rictor | -0.111399944 | 0.37294475 | 0 |
| AT21612 | FBgn0040291 | Roc1b | 0.111378571 | 0.38251436 | 0 |
| AT02150 | FBgn0025702 | Srpk79D | 0.111377914 | 0.38451222 | 0 |
| LD36528 | FBgn0033782 | sug | -0.111316246 | 0.45877745 | 0 |
| GH07821 | FBgn0039555 | mRpS22 | 0.111273431 | 0.37251549 | 0 |
| GH15583 | FBgn0041096 | rols | 0.111268128 | 0.67659661 | 0 |
| AT19678 | FBgn0036882 | CG9279 | 0.111238144 | 0.40248241 | 0 |
| LD33178 | FBgn0051301 | CG31301 | 0.111195951 | 0.3403667 | 0 |

| | | | | | |
|---|---|---|---|---|---|
| RH69567 | FBgn0005672 | spi | 0.111182538 | 0.40340611 | 0 |
| GH11762 | FBgn0030593 | CG9512 | 0.11118228 | 0.67167761 | 0 |
| SD08625 | FBgn0035073 | CG16896 | 0.111179866 | 0.31360581 | 0 |
| RE63851 | FBgn0013531 | MED20 | -0.111126201 | 0.36845117 | 0 |
| RE57384 | FBgn0033656 | S2P | 0.111105447 | 0.3639518 | 0 |
| GH10778 | FBgn0264086 | CG43755 | 0.110968903 | 0.35241103 | 0 |
| GH02288 | FBgn0029175 | sotv | 0.110946333 | 0.30470968 | 0 |
| LD05963 | FBgn0032499 | Uvrag | -0.110871639 | 0.35604019 | 0 |
| LD24362 | FBgn0036402 | CG6650 | 0.110798336 | 0.3160745 | 0 |
| LD43863 | FBgn0031895 | CG4497 | 0.110748676 | 0.30830485 | 0 |
| GH27039 | FBgn0033791 | Drl-2 | -0.11073837 | 0.39374565 | 0 |
| GM27203 | FBgn0036155 | CG6163 | -0.110691109 | 0.343932 | 0 |
| GM27203 | FBgn0260441 | RpS12 | -0.110691109 | 0.343932 | 0 |
| LD41209 | FBgn0036487 | Prp31 | 0.110689224 | 0.40836759 | 0 |
| SD07714 | FBgn0011570 | cpb | -0.110688147 | 0.39925094 | 0 |
| RE57454 | FBgn0032088 | CG13102 | 0.110682934 | 0.44738846 | 0 |
| LD37358 | FBgn0029833 | MCTS1 | 0.110665269 | 0.53620282 | 0 |
| SD03610 | FBgn0033388 | CG8046 | 0.110623254 | 0.3597264 | 0 |
| GH11073 | FBgn0261597 | RpS26 | 0.11052064 | 0.58085754 | 0 |
| GH11073 | FBgn0039792 | CG2267 | 0.11052064 | 0.58085754 | 0 |
| RE57134 | FBgn0031716 | CG14015 | -0.110513726 | 0.31905024 | 0 |
| RE58349 | FBgn0029838 | CG4666 | 0.110511809 | 0.66139187 | 0 |
| RE73739 | FBgn0052205 | hpRNA:CR32205 | -0.110462648 | 0.31043584 | 0 |
| RE73739 | FBgn0036872 | CG12519 | -0.110462648 | 0.31043584 | 0 |
| RE47719 | FBgn0016983 | smid | 0.110423627 | 0.37937674 | 0 |
| AT08706 | FBgn0034007 | ND-51L2 | -0.110422055 | 0.4101661 | 0 |
| RE36103 | FBgn0003206 | Ras64B | -0.110390275 | 0.33644471 | 0 |
| RE33041 | FBgn0031467 | Cpr23B | -0.110353132 | 0.40522779 | 0 |
| GH01206 | FBgn0031514 | CG3332 | 0.110328227 | 0.3547438 | 0 |
| GH13342 | FBgn0030269 | CDK2AP1 | -0.110321228 | 0.35177097 | 0 |
| GH13342 | FBgn0264863 | asRNA:CR44054 | -0.110321228 | 0.35177097 | 0 |
| GH25884 | FBgn0031310 | Vps29 | 0.110317774 | 0.32653939 | 0 |
| RE03440 | FBgn0026630 | nes | -0.110310548 | 0.34021082 | 0 |
| GM14040 | FBgn0039260 | Smg6 | 0.110302414 | 0.29420649 | 0 |
| AT21561 | FBgn0051910 | CG31910 | 0.110300476 | 0.48048208 | 0 |
| RE44287 | FBgn0033645 | CG13196 | -0.11028574 | 0.32332024 | 0 |
| LD31742 | FBgn0261862 | whd | 0.110258246 | 0.37234057 | 0 |
| RE35033 | FBgn0033392 | CG8027 | 0.110246653 | 0.30935459 | 0 |
| LD31822 | FBgn0031678 | CG31918 | 0.110181502 | 0.33825867 | 0 |
| SD09045 | FBgn0040466 | Dlip2 | 0.110160739 | 0.35688979 | 0 |
| GH15479 | FBgn0037064 | CG9389 | -0.110148072 | 0.31217459 | 0 |
| RH40246 | FBgn0039529 | CG5612 | -0.110079058 | 0.37533608 | 0 |
| RE52086 | FBgn0040351 | CG11638 | -0.11005156 | 0.39498124 | 0 |
| SD09850 | FBgn0035039 | Adck | -0.11002823 | 0.33110818 | 0 |
| AT12460 | FBgn0028897 | CG4935 | 0.110007452 | 0.32229354 | 0 |
| LD36016 | FBgn0035063 | CG3594 | 0.109983381 | 0.38538296 | 0 |
| AT16953 | FBgn0036217 | CG5906 | 0.109917247 | 0.47068547 | 0 |
| LP05669 | FBgn0050029 | CR30029 | -0.109770276 | 0.33928392 | 0 |

| | | | | | |
|---|---|---|---|---|---|
| LP09923 | FBgn0262081 | Csk | -0.109708696 | 0.31531092 | 0 |
| AT23338 | FBgn0034196 | CG15605 | 0.109624209 | 0.63569444 | 0 |
| LD23540 | FBgn0025457 | Bub3 | 0.109567926 | 0.33110818 | 0 |
| RE02772 | FBgn0040623 | Spase12 | 0.109487414 | 0.4141373 | 0 |
| LD16270 | FBgn0010300 | brat | -0.109484582 | 0.45026411 | 0 |
| GH01937 | FBgn0034069 | CG8401 | 0.109440418 | 0.50296341 | 0 |
| LD01659 | FBgn0030833 | CG8915 | 0.10943046 | 0.468209 | 0 |
| GH10708 | FBgn0036199 | Bmcp | -0.109312454 | 0.33398053 | 0 |
| AT29766 | FBgn0033369 | CG8197 | -0.109227145 | 0.40221267 | 0 |
| RE30346 | FBgn0264906 | lncRNA:CR44097 | -0.109161703 | 0.31964742 | 0 |
| RE52007 | FBgn0004379 | Klp67A | 0.108996582 | 0.33976676 | 0 |
| RE52135 | FBgn0011824 | CG4038 | -0.108986433 | 0.32650577 | 0 |
| LD07519 | FBgn0259916 | CG42445 | 0.108949102 | 0.43726284 | 0 |
| GH10833 | FBgn0032229 | CG5045 | 0.108931003 | 0.31511681 | 0 |
| GH20904 | FBgn0033869 | Cpr50Cb | -0.108906213 | 0.61238802 | 0 |
| LD37690 | FBgn0039636 | Atg14 | 0.108874129 | 0.37347593 | 0 |
| GH05443 | FBgn0011701 | repo | -0.108861864 | 0.44586101 | 0 |
| SD22372 | FBgn0030996 | CG14194 | 0.108842656 | 0.34349101 | 0 |
| LD28757 | FBgn0037240 | Cont | 0.108737704 | 0.32747679 | 0 |
| LP09838 | FBgn0262887 | lncRNA:CR43242 | -0.108725108 | 0.45330723 | 0 |
| RE72116 | FBgn0033480 | mRpL42 | 0.108687568 | 0.3923685 | 0 |
| GM22991 | FBgn0010549 | l(2)03659 | -0.108668481 | 0.3835278 | 0 |
| LD14807 | FBgn0037137 | Nopp140 | 0.108668057 | 0.36070583 | 0 |
| GH01453 | FBgn0031737 | obst-E | 0.108667588 | 0.43962207 | 0 |
| GH10640 | FBgn0036512 | CG16979 | -0.108654245 | 0.30080363 | 0 |
| RE16861 | FBgn0031635 | tank | 0.108649527 | 0.38555567 | 0 |
| GH13022 | FBgn0035915 | S-Lap1 | -0.108639997 | 0.35191106 | 0 |
| LP01886 | FBgn0038783 | CG4367 | 0.108635827 | 0.49093682 | 0 |
| LD27552 | FBgn0011737 | Wee1 | 0.108631283 | 0.3218739 | 0 |
| RH62530 | FBgn0033593 | Listericin | -0.10861616 | 0.32084759 | 0 |
| LD36294 | FBgn0034491 | Hsl | 0.108609134 | 0.30755546 | 0 |
| RE72862 | FBgn0032943 | Tsp39D | -0.108532642 | 0.33105043 | 0 |
| SD03378 | FBgn0031574 | TTLL4B | 0.108436379 | 0.35079094 | 0 |
| RH02253 | FBgn0004240 | DptA | 0.108406063 | 0.32084759 | 0 |
| LD20901 | FBgn0026150 | ApepP | 0.108385402 | 0.307972 | 0 |
| LD09907 | FBgn0001186 | Hex-A | -0.108294775 | 0.37504611 | 0 |
| LD41803 | FBgn0039182 | CG5728 | 0.108286293 | 0.36988327 | 0 |
| RE15383 | FBgn0014127 | barr | -0.108278582 | 0.32282288 | 0 |
| GH17311 | FBgn0030061 | CG1785 | 0.10821507 | 0.39499648 | 0 |
| RE16136 | FBgn0035551 | CG7465 | 0.10817636 | 0.60840684 | 0 |
| GH10971 | FBgn0020381 | Dredd | 0.108160126 | 0.36847667 | 0 |
| RE57896 | FBgn0013432 | bcn92 | -0.108155748 | 0.32345633 | 0 |
| LD29847 | FBgn0086357 | Sec61alpha | 0.108047228 | 0.50779269 | 0 |
| GH01072 | FBgn0036811 | MED11 | -0.108015958 | 0.55347949 | 0 |
| LD47713 | FBgn0263278 | Fic | -0.107942318 | 0.34994022 | 0 |
| RH38368 | FBgn0038879 | CG17298 | -0.107927064 | 0.49704094 | 0 |
| SD04949 | FBgn0266672 | Sec8 | 0.107699397 | 0.50714837 | 0 |
| RE43931 | FBgn0031918 | CG6055 | 0.1075919 | 0.51838673 | 0 |

| | | | | | |
|---|---|---|---|---|---|
| RE23041 | FBgn0034011 | CG8160 | -0.107591629 | 0.39598601 | 0 |
| RH01011 | FBgn0032402 | CG14945 | 0.107578794 | 0.39197671 | 0 |
| RH45738 | FBgn0040775 | CG12158 | -0.10757776 | 0.54856666 | 0 |
| GH25035 | FBgn0034638 | CG10433 | -0.107515442 | 0.40698474 | 0 |
| GM01757 | FBgn0042112 | mRpL36 | 0.10750871 | 0.38135654 | 0 |
| RE55472 | FBgn0264291 | Det | 0.107432292 | 0.43959925 | 0 |
| RE01906 | FBgn0020906 | Jon25Bi | -0.107380494 | 0.43749234 | 0 |
| RE41041 | FBgn0039197 | CG17780 | 0.107364977 | 0.63419496 | 0 |
| AT26096 | FBgn0036206 | CG5964 | 0.107355873 | 0.53333403 | 0 |
| RE06169 | FBgn0051272 | CG31272 | -0.107336483 | 0.37918289 | 0 |
| RE11703 | FBgn0053969 | CG33969 | -0.107290921 | 0.3835278 | 0 |
| LD07728 | FBgn0035902 | CG6683 | 0.107156854 | 0.37702988 | 0 |
| AT19571 | FBgn0030623 | PPYR1 | 0.10714422 | 0.58704537 | 0 |
| RE47667 | FBgn0031660 | mRpL28 | 0.107140657 | 0.38912035 | 0 |
| RE07451 | FBgn0035141 | Cypl | 0.107103366 | 0.31028912 | 0 |
| GH11578 | FBgn0036016 | CG3306 | 0.107028152 | 0.39558212 | 0 |
| RE67391 | FBgn0026409 | Mpcp2 | 0.10698416 | 0.42364786 | 0 |
| RE67391 | FBgn0034497 | Mpcp1 | 0.10698416 | 0.42364786 | 0 |
| RE33612 | FBgn0031322 | CG5001 | -0.106968316 | 0.35205829 | 0 |
| LD40940 | FBgn0031484 | CG3165 | -0.106964272 | 0.31801668 | 0 |
| GH05922 | FBgn0003118 | pnt | 0.106950452 | 0.50766723 | 0 |
| RE22964 | FBgn0040080 | pins | -0.106913751 | 0.31581229 | 0 |
| LD15224 | FBgn0034750 | CG3732 | 0.106898056 | 0.33275745 | 0 |
| SD08412 | FBgn0051642 | CG31642 | 0.106851902 | 0.34091275 | 0 |
| LD42289 | FBgn0032149 | CG4036 | 0.106824279 | 0.36971494 | 0 |
| AT08210 | FBgn0032894 | CG14402 | -0.106806474 | 0.35447946 | 0 |
| GH01560 | FBgn0030259 | CG1545 | 0.106795326 | 0.49287764 | 0 |
| LD43328 | FBgn0033359 | CG8213 | -0.106771676 | 0.67051055 | 0 |
| GH03958 | FBgn0002937 | ninaB | 0.106765391 | 0.47408961 | 0 |
| GH10642 | FBgn0033448 | hebe | 0.106754459 | 0.3883281 | 0 |
| GM13959 | FBgn0025366 | Ip259 | 0.106693683 | 0.37752507 | 0 |
| RE16762 | FBgn0015737 | Hmu | 0.106619281 | 0.41685591 | 0 |
| LD04710 | FBgn0036741 | CG7510 | 0.106606084 | 0.34975894 | 0 |
| RE11711 | FBgn0034707 | MED16 | 0.106543388 | 0.45923276 | 0 |
| LD24223 | FBgn0014133 | bif | 0.106524971 | 0.31432201 | 0 |
| GH05918 | FBgn0033033 | scaf | -0.106522899 | 0.53542074 | 0 |
| LD16174 | FBgn0003882 | tub | 0.106493426 | 0.35177097 | 0 |
| RH37294 | FBgn0039723 | CG15522 | -0.106490769 | 0.40623544 | 0 |
| GH02901 | FBgn0027601 | pdgy | 0.106464341 | 0.36395337 | 0 |
| LD26447 | FBgn0037024 | tzn | 0.106446021 | 0.49428172 | 0 |
| LD42588 | FBgn0010583 | dock | 0.106365354 | 0.30993577 | 0 |
| RH17413 | FBgn0030011 | Gbeta5 | 0.106341453 | 0.47277506 | 0 |
| RE13447 | FBgn0040230 | dbo | 0.10626694 | 0.32041354 | 0 |
| LD41231 | FBgn0037025 | Spc105R | -0.106203886 | 0.33919374 | 0 |
| LD04190 | FBgn0051163 | SKIP | 0.106113695 | 0.37633652 | 0 |
| RE14595 | FBgn0086472 | RpS25 | -0.106097458 | 0.34916277 | 0 |
| GH28477 | FBgn0034214 | CG6550 | 0.106059961 | 0.38448688 | 0 |
| RE62270 | FBgn0028968 | gammaCOP | -0.10602898 | 0.36847667 | 0 |

| | | | | | |
|---|---|---|---|---|---|
| LP05733 | FBgn0028533 | CG7953 | 0.106027032 | 0.34419536 | 0 |
| RH68619 | FBgn0051324 | CG31324 | -0.105995091 | 0.40916709 | 0 |
| GH10480 | FBgn0037643 | skap | 0.105962455 | 0.38296185 | 0 |
| RE10234 | FBgn0035131 | mthl9 | -0.105932794 | 0.52195992 | 0 |
| GH07711 | FBgn0028992 | sds22 | -0.105924701 | 0.36847667 | 0 |
| GH04518 | FBgn0053963 | lncRNA:CR33963 | 0.105891607 | 0.48526469 | 0 |
| GH04518 | FBgn0011837 | Tis11 | 0.105891607 | 0.48526469 | 0 |
| LD45391 | FBgn0000464 | Lar | 0.105711054 | 0.3939846 | 0 |
| SD17528 | FBgn0039713 | RpS8 | 0.105681826 | 0.4101661 | 0 |
| RE26382 | FBgn0010409 | RpL18A | 0.105662993 | 0.60358513 | 0 |
| AT04640 | FBgn0259729 | CG42383 | 0.10565815 | 0.57946324 | 0 |
| LD45157 | FBgn0000180 | bib | 0.105654261 | 0.61208607 | 0 |
| RE04882 | FBgn0004780 | Ccp84Ad | -0.105622591 | 0.53608002 | 0 |
| RE04882 | FBgn0004779 | Ccp84Ae | -0.105622591 | 0.53608002 | 0 |
| RE74211 | FBgn0005648 | Pabp2 | 0.105622572 | 0.3160745 | 0 |
| GM14138 | FBgn0031194 | CG17598 | -0.105576667 | 0.45460496 | 0 |
| RH02901 | FBgn0031538 | CG3246 | -0.105569007 | 0.51140355 | 0 |
| SD10988 | FBgn0046756 | pncr017:3R | 0.105506982 | 0.31586471 | 0 |
| RE39465 | FBgn0266417 | ringer | 0.105498081 | 0.31678803 | 0 |
| GH14327 | FBgn0035871 | BI-1 | 0.105467463 | 0.33476179 | 0 |
| RH55750 | FBgn0040837 | CG8620 | -0.105465829 | 0.3218739 | 0 |
| RE46560 | FBgn0039751 | CG1983 | 0.105443123 | 0.3597264 | 0 |
| SD21818 | FBgn0021750 | SerRS-m | -0.10543559 | 0.31444456 | 0 |
| RE71294 | FBgn0033753 | Cyp301a1 | 0.105418094 | 0.36971494 | 0 |
| LD09950 | FBgn0017572 | Mo25 | 0.105341303 | 0.34975848 | 0 |
| LD12396 | FBgn0039163 | CG5515 | 0.105335644 | 0.42856874 | 0 |
| LD04521 | FBgn0000063 | Mps1 | -0.105314084 | 0.33403681 | 0 |
| GH09250 | FBgn0021872 | Xbp1 | -0.105269566 | 0.36161041 | 0 |
| RE61993 | FBgn0030518 | CG11134 | 0.105195506 | 0.38781426 | 0 |
| GM13293 | FBgn0003227 | rec | 0.105158687 | 0.36890521 | 0 |
| AT19876 | FBgn0050065 | CG30065 | -0.105064356 | 0.45210484 | 0 |
| AT15052 | FBgn0032609 | CG13280 | 0.105057265 | 0.36813914 | 0 |
| LD42024 | FBgn0031195 | CG17600 | 0.105056159 | 0.40852778 | 0 |
| LD25561 | FBgn0266582 | ND-30 | 0.105056099 | 0.42292976 | 0 |
| LD05521 | FBgn0037878 | CG6693 | 0.105053492 | 0.34120473 | 0 |
| RE44079 | FBgn0036915 | Prp3 | 0.105050954 | 0.33362643 | 0 |
| AT29350 | FBgn0036784 | CG5103 | -0.105001085 | 0.51930145 | 0 |
| LD35843 | FBgn0034027 | CG8187 | 0.104985677 | 0.32278903 | 0 |
| GM03621 | FBgn0051368 | CG31368 | 0.104945795 | 0.36971494 | 0 |
| LD21713 | FBgn0026576 | CG5991 | 0.104915053 | 0.38782292 | 0 |
| LD24232 | FBgn0030945 | Ing3 | 0.104874355 | 0.34581821 | 0 |
| RE72989 | FBgn0027341 | Gfat1 | 0.104861435 | 0.5160646 | 0 |
| LD23308 | FBgn0037710 | CG9393 | 0.104820086 | 0.32073204 | 0 |
| RE62581 | FBgn0032987 | RpL21 | -0.104815943 | 0.39615894 | 0 |
| SD09735 | FBgn0031519 | Fign | 0.104807443 | 0.37374788 | 0 |
| GM13207 | FBgn0019830 | colt | 0.104788303 | 0.35120297 | 0 |
| RH24774 | FBgn0031500 | CG17221 | -0.104722124 | 0.31811714 | 0 |
| LD37206 | FBgn0032259 | CG6144 | 0.104677619 | 0.40945908 | 0 |

| | | | | | |
|---|---|---|---|---|---|
| RE40614 | FBgn0035953 | CG5087 | 0.104477883 | 0.35345047 | 0 |
| LD08638 | FBgn0020270 | mre11 | 0.104449848 | 0.38272145 | 0 |
| GH26270 | FBgn0038922 | Idh3b | 0.10442932 | 0.41123404 | 0 |
| RH40610 | FBgn0021906 | RFeSP | 0.10420771 | 0.43786983 | 0 |
| SD02261 | FBgn0262516 | Trpml | -0.104175245 | 0.34664489 | 0 |
| SD02261 | FBgn0261361 | CG42638 | -0.104175245 | 0.34664489 | 0 |
| AT03104 | FBgn0020647 | KrT95D | -0.104169108 | 0.37603083 | 0 |
| LD45245 | FBgn0038056 | CG5961 | -0.104157963 | 0.38251436 | 0 |
| LD31161 | FBgn0265998 | Doa | -0.104144376 | 0.4690427 | 0 |
| SD09318 | FBgn0003969 | vap | 0.104143768 | 0.33976676 | 0 |
| RE27549 | FBgn0022985 | qkr58E-2 | 0.104125706 | 0.37477926 | 0 |
| RE50669 | FBgn0036791 | CG7271 | -0.104086762 | 0.34713139 | 0 |
| GH23955 | FBgn0037679 | Aduk | -0.104063266 | 0.43822876 | 0 |
| GH12682 | FBgn0031865 | Nha1 | 0.10404805 | 0.42117179 | 0 |
| RE41093 | FBgn0035942 | ValRS-m | -0.104037768 | 0.38128171 | 0 |
| RE52283 | FBgn0037448 | CG15186 | -0.104003235 | 0.37294475 | 0 |
| LD29481 | FBgn0000376 | crm | 0.103980266 | 0.42107197 | 0 |
| SD10251 | FBgn0026015 | Top3beta | 0.10397737 | 0.42280478 | 0 |
| GH22740 | FBgn0031713 | CG7277 | -0.103973228 | 0.48156985 | 0 |
| RE49731 | FBgn0031351 | CG14352 | 0.103954808 | 0.34190553 | 0 |
| LD44083 | FBgn0003687 | Tbp | -0.103950369 | 0.35771172 | 0 |
| GH12243 | FBgn0283509 | Phm | -0.103943624 | 0.33777794 | 0 |
| LD10920 | FBgn0265974 | ttv | -0.103939785 | 0.42705205 | 0 |
| GH04238 | FBgn0034920 | CG5597 | -0.103918072 | 0.37627069 | 0 |
| LD29875 | FBgn0034304 | CG5742 | -0.10388506 | 0.48313356 | 0 |
| AT01380 | FBgn0033963 | CG12857 | -0.103866003 | 0.33866194 | 0 |
| LD24841 | FBgn0033737 | Nup54 | 0.103783374 | 0.43335785 | 0 |
| LD35854 | FBgn0034933 | CG3735 | 0.1037487 | 0.32282288 | 0 |
| LD30122 | FBgn0029687 | Vap33 | -0.10372064 | 0.42061149 | 0 |
| AT07829 | FBgn0034838 | CG12782 | 0.103701703 | 0.44253792 | 0 |
| GM04983 | FBgn0033715 | CG8490 | 0.103600849 | 0.31861085 | 0 |
| SD04516 | FBgn0030605 | ND-B18 | 0.103587095 | 0.41089595 | 0 |
| AT25567 | FBgn0039204 | CG6607 | -0.103576554 | 0.49647104 | 0 |
| LD34806 | FBgn0051849 | CG31849 | 0.103566184 | 0.35688979 | 0 |
| GH24739 | FBgn0036820 | Grx1 | -0.103524257 | 0.36618192 | 0 |
| RE09163 | FBgn0036819 | Dysb | -0.103503896 | 0.43087554 | 0 |
| LD02061 | FBgn0039907 | lgs | 0.10350281 | 0.38917984 | 0 |
| RE42193 | FBgn0264712 | CG1172 | 0.103436741 | 0.32121856 | 0 |
| RE47447 | FBgn0039670 | CG7567 | -0.103430377 | 0.37533608 | 0 |
| SD09011 | FBgn0261954 | east | 0.103400276 | 0.48998316 | 0 |
| SD21513 | FBgn0030761 | CG9784 | -0.103382259 | 0.50147119 | 0 |
| GH05862 | FBgn0027785 | NP15.6 | 0.103370159 | 0.39536176 | 0 |
| RE39068 | FBgn0035936 | Tsp66E | -0.103354243 | 0.34581096 | 0 |
| LD44506 | FBgn0038855 | CG5745 | -0.103327396 | 0.5695879 | 0 |
| LD03564 | FBgn0011586 | e(r) | -0.10332096 | 0.36803712 | 0 |
| AT19392 | FBgn0031462 | CG2964 | -0.103309337 | 0.43942913 | 0 |
| RH47720 | FBgn0040227 | eIF3d1 | -0.103303855 | 0.43578782 | 0 |
| GH27433 | FBgn0038237 | Pde6 | -0.103286707 | 0.34994022 | 0 |

| | | | | | |
|---|---|---|---|---|---|
| RE66370 | FBgn0037683 | CG18473 | -0.103241586 | 0.54357267 | 0 |
| LD40326 | FBgn0261353 | Ccdc56 | 0.103189023 | 0.40688164 | 0 |
| LD40326 | FBgn0261381 | mtTFB1 | 0.103189023 | 0.40688164 | 0 |
| GH09510 | FBgn0052354 | CG32354 | -0.103108765 | 0.400006 | 0 |
| GM04721 | FBgn0030753 | rngo | -0.103069091 | 0.32543083 | 0 |
| LD40504 | FBgn0039270 | PQBP1 | 0.102890834 | 0.521131 | 0 |
| LD29090 | FBgn0037554 | DNApol-iota | 0.102888929 | 0.37804643 | 0 |
| GH13177 | FBgn0264305 | CG43783 | 0.102863942 | 0.41211126 | 0 |
| RE64050 | FBgn0038261 | CG14856 | 0.102779455 | 0.38066293 | 0 |
| RE56935 | FBgn0026255 | clumsy | -0.102768527 | 0.54254366 | 0 |
| SD07645 | FBgn0261839 | pre-mod(mdg4)-Y | 0.102709724 | 0.34904751 | 0 |
| SD07645 | FBgn0261843 | pre-mod(mdg4)-W | 0.102709724 | 0.34904751 | 0 |
| LP10803 | FBgn0033344 | CG8252 | -0.102678935 | 0.44131655 | 0 |
| RH41593 | FBgn0261608 | RpL37A | -0.102561935 | 0.34297995 | 0 |
| GH14561 | FBgn0037231 | Vps24 | -0.10251475 | 0.35302499 | 0 |
| LD24895 | FBgn0034962 | MAN1 | 0.102487152 | 0.35710349 | 0 |
| LP08350 | FBgn0003950 | unc | -0.102440601 | 0.47240791 | 0 |
| LD24616 | FBgn0259212 | cno | -0.102430943 | 0.66510973 | 0 |
| RE06383 | FBgn0041160 | comm2 | -0.102379071 | 0.61209673 | 0 |
| SD10311 | FBgn0037622 | CG8202 | -0.102273896 | 0.36423299 | 0 |
| RE02292 | FBgn0040348 | CG3703 | 0.102240705 | 0.35954543 | 0 |
| RE15501 | FBgn0250820 | meigo | 0.102236303 | 0.39333702 | 0 |
| RE70942 | FBgn0003358 | Jon99Ci | 0.102169741 | 0.41639809 | 0 |
| SD02826 | FBgn0031703 | CG12512 | 0.102165736 | 0.36731441 | 0 |
| HL01284 | FBgn0010350 | CdsA | 0.102063474 | 0.41483181 | 0 |
| LD40801 | FBgn0002891 | mus205 | -0.102041032 | 0.36870325 | 0 |
| LD36817 | FBgn0033874 | CG6347 | -0.102017785 | 0.60452986 | 0 |
| LD16191 | FBgn0038829 | CG17271 | 0.101990659 | 0.4026238 | 0 |
| SD21019 | FBgn0033756 | CG17760 | 0.101957988 | 0.44693039 | 0 |
| RH19665 | FBgn0002789 | Mp20 | -0.10191244 | 0.78816984 | 0 |
| GM02143 | FBgn0014411 | Vps26 | 0.101841455 | 0.35647622 | 0 |
| RE45089 | FBgn0005564 | Shal | -0.101771226 | 0.46514252 | 0 |
| LD23634 | FBgn0028474 | CG4119 | 0.101732285 | 0.32725139 | 0 |
| RE35653 | FBgn0031659 | CG14043 | 0.101721629 | 0.36070583 | 0 |
| RE67573 | FBgn0063670 | CG40228 | 0.101709499 | 0.38782292 | 0 |
| GH01161 | FBgn0265464 | Traf6 | 0.101697564 | 0.39339733 | 0 |
| RH68464 | FBgn0030737 | CG9914 | 0.101678438 | 0.54936559 | 0 |
| LD44732 | FBgn0035520 | CG11586 | 0.101652466 | 0.34384335 | 0 |
| LD04530 | FBgn0000384 | cta | -0.101627744 | 0.52275108 | 0 |
| LD04530 | FBgn0058005 | CR40005 | -0.101627744 | 0.52275108 | 0 |
| GH13823 | FBgn0039219 | CG13630 | -0.101622013 | 0.39339733 | 0 |
| RH02823 | FBgn0031459 | CG2862 | -0.101610524 | 0.33837141 | 0 |
| LP12144 | FBgn0285947 | RpS10b | 0.101610366 | 0.34425978 | 0 |
| RE09269 | FBgn0030981 | CG14191 | 0.101590017 | 0.36885296 | 0 |
| GM02568 | FBgn0033884 | CG13344 | -0.10155799 | 0.32725772 | 0 |
| LD43876 | FBgn0025549 | unc-119 | -0.101550114 | 0.5164049 | 0 |
| RE61424 | FBgn0033421 | CG1888 | 0.101427446 | 0.33141556 | 0 |
| LD41918 | FBgn0284254 | Impbeta11 | -0.101413495 | 0.40515451 | 0 |

| | | | | | |
|---|---|---|---|---|---|
| GH12663 | FBgn0027539 | lili | 0.101391702 | 0.35166517 | 0 |
| RH34523 | FBgn0085438 | CG34409 | -0.101284673 | 0.48214468 | 0 |
| GM08142 | FBgn0266848 | wap | 0.101226431 | 0.40379867 | 0 |
| RH38463 | FBgn0040688 | CG12483 | -0.101222539 | 0.37459627 | 0 |
| GH04802 | FBgn0027589 | CG1688 | -0.10121647 | 0.59945145 | 0 |
| LP07986 | FBgn0000109 | Aprt | 0.101139258 | 0.44629412 | 0 |
| LD27988 | FBgn0028341 | Ptpmeg2 | 0.101061681 | 0.33784959 | 0 |
| LD22655 | FBgn0031566 | CG2818 | 0.100898279 | 0.3551038 | 0 |
| GM09915 | FBgn0034194 | CG15611 | -0.100878129 | 0.43026786 | 0 |
| LD23846 | FBgn0038984 | AdipoR | -0.100793044 | 0.47419245 | 0 |
| RE06807 | FBgn0032816 | Nf-YB | 0.100758477 | 0.51340267 | 0 |
| LD18468 | FBgn0260780 | wisp | 0.100757353 | 0.36885296 | 0 |
| LD45273 | FBgn0266101 | CG44838 | 0.100700941 | 0.3883281 | 0 |
| GH19680 | FBgn0039872 | salt | -0.100696374 | 0.35205829 | 0 |
| SD05642 | FBgn0265102 | Oseg1 | -0.100693591 | 0.34693537 | 0 |
| RH19136 | FBgn0051326 | CG31326 | 0.100630172 | 0.58411484 | 0 |
| LD10578 | FBgn0003423 | slgA | -0.10062602 | 0.37177187 | 0 |
| GM27819 | FBgn0036052 | CG10809 | 0.100622346 | 0.41252617 | 0 |
| RH52788 | FBgn0014454 | Acp1 | 0.100566285 | 0.36813914 | 0 |
| HL01428 | FBgn0000299 | Col4a1 | -0.100565905 | 0.75669451 | 0 |
| RE25188 | FBgn0010741 | Pfdn2 | -0.100537112 | 0.45026411 | 0 |
| LD17482 | FBgn0038325 | Atg4b | 0.10049891 | 0.3547438 | 0 |
| RH46857 | FBgn0000500 | Dsk | 0.100455648 | 0.53948777 | 0 |
| GM01885 | FBgn0034215 | Mtap | 0.100439113 | 0.37074408 | 0 |
| RE51891 | FBgn0033989 | CG7639 | -0.100429916 | 0.34471615 | 0 |
| LD21722 | FBgn0024909 | Taf7 | -0.100429231 | 0.34170455 | 0 |
| LD26833 | FBgn0036697 | rogdi | 0.100400821 | 0.40340611 | 0 |
| LD38464 | FBgn0001215 | Hrb98DE | 0.100393382 | 0.35347549 | 0 |
| GH03163 | FBgn0027598 | cindr | -0.10038647 | 0.35649586 | 0 |
| RH01553 | FBgn0025866 | CalpB | 0.10037831 | 0.37459627 | 0 |
| LD47479 | FBgn0036850 | Gem2 | 0.100364596 | 0.34494598 | 0 |
| GH07209 | FBgn0025712 | CG13920 | 0.100311097 | 0.62120043 | 0 |
| SD10847 | FBgn0036987 | CG5274 | 0.100294933 | 0.43923068 | 0 |
| LD42275 | FBgn0030687 | CG17209 | -0.100293017 | 0.4344621 | 0 |
| GM14972 | FBgn0028707 | Mt2 | 0.100280003 | 0.64469975 | 0 |
| RE74350 | FBgn0015756 | RpL9 | -0.100279422 | 0.35054609 | 0 |
| GH01229 | FBgn0038585 | Non3 | 0.100256552 | 0.40901478 | 0 |
| GH13919 | FBgn0261439 | SdhA | -0.100248229 | 0.33110818 | 0 |
| GH12350 | FBgn0026144 | CBP | -0.100239128 | 0.36303405 | 0 |
| RE01528 | FBgn0035006 | CG4563 | -0.100238228 | 0.39676786 | 0 |
| GH19142 | FBgn0032774 | CG17549 | 0.10018892 | 0.60010256 | 0 |
| GH02816 | FBgn0026593 | CG5707 | 0.100176453 | 0.44868134 | 0 |
| GH14923 | FBgn0016696 | Pitslre | 0.100153457 | 0.40819607 | 0 |
| GH01991 | FBgn0030163 | CG1791 | -0.099981562 | 0.42449526 | 0 |
| LP01585 | FBgn0031224 | CG11454 | -0.099949499 | 0.33254606 | 0 |
| RE18748 | FBgn0031419 | CG15390 | 0.09989452 | 0.38829968 | 0 |
| LD38235 | FBgn0037536 | CG2698 | 0.099892671 | 0.34354567 | 0 |
| RH72686 | FBgn0031466 | CG3119 | -0.099830376 | 0.52433469 | 0 |

| | | | | | |
|---|---|---|---|---|---|
| LD24839 | FBgn0032635 | CG15141 | 0.099805396 | 0.44868134 | 0 |
| AT17603 | FBgn0039347 | CG5071 | -0.099799465 | 0.55511879 | 0 |
| LD09945 | FBgn0023521 | CG3587 | 0.099791529 | 0.38582221 | 0 |
| RE69116 | FBgn0017418 | ari-1 | 0.099777652 | 0.38363218 | 0 |
| RE53401 | FBgn0037504 | CG1142 | 0.099701273 | 0.40448993 | 0 |
| GH28859 | FBgn0036813 | Atg3 | 0.09967576 | 0.3885625 | 0 |
| GH01077 | FBgn0038271 | UQCR-C1 | -0.099615976 | 0.53539893 | 0 |
| GH11140 | FBgn0038358 | CG4525 | -0.099505763 | 0.41052439 | 0 |
| RH54928 | FBgn0035544 | CG15021 | -0.099419876 | 0.33866194 | 0 |
| LP11508 | FBgn0260799 | p120ctn | -0.09940582 | 0.36724764 | 0 |
| LP11508 | FBgn0039336 | CG4553 | -0.09940582 | 0.36724764 | 0 |
| LD21606 | FBgn0030628 | CG9114 | 0.099365704 | 0.3394885 | 0 |
| AT14088 | FBgn0002709 | mei-218 | -0.099342404 | 0.43026786 | 0 |
| AT27980 | FBgn0039359 | RpL27 | 0.099307978 | 0.36070583 | 0 |
| RE05370 | FBgn0035192 | CG9194 | 0.099302333 | 0.40103942 | 0 |
| RH40211 | FBgn0031888 | Pvf2 | -0.099278718 | 0.43658434 | 0 |
| RE18101 | FBgn0043841 | vir-1 | 0.099207262 | 0.51833128 | 0 |
| GH21948 | FBgn0028507 | CG3793 | -0.099195392 | 0.34091275 | 0 |
| RE42228 | FBgn0042205 | CG18764 | 0.099167581 | 0.44577837 | 0 |
| LD46389 | FBgn0034313 | CG5726 | -0.099163231 | 0.51375619 | 0 |
| GH19763 | FBgn0037721 | CG9427 | 0.09914329 | 0.4101661 | 0 |
| RE18415 | FBgn0038389 | CG5516 | 0.099080559 | 0.39480753 | 0 |
| RE18828 | FBgn0036629 | GluRS-m | 0.098996008 | 0.34349101 | 0 |
| SD10511 | FBgn0029079 | icln | -0.098971086 | 0.47560044 | 0 |
| GH22843 | FBgn0000039 | nAChRalpha2 | -0.098957504 | 0.60480402 | 0 |
| RH66048 | FBgn0033627 | CG13204 | -0.098957273 | 0.40175048 | 0 |
| GH22035 | FBgn0005614 | trpl | 0.098955678 | 0.40243057 | 0 |
| LD22771 | FBgn0037944 | CG6923 | 0.098908618 | 0.35954543 | 0 |
| LD44026 | FBgn0032197 | CG5694 | -0.098839765 | 0.40528755 | 0 |
| LD34444 | FBgn0034009 | CG8155 | 0.098793738 | 0.34401092 | 0 |
| RH26504 | FBgn0037009 | CG5104 | -0.098706642 | 0.38463262 | 0 |
| LD45889 | FBgn0005655 | PCNA | -0.098655727 | 0.35771172 | 0 |
| GH14524 | FBgn0005775 | Con | 0.098631874 | 0.48526469 | 0 |
| GM12270 | FBgn0010621 | CCT5 | -0.098625643 | 0.40819607 | 0 |
| LD09912 | FBgn0031871 | Fgop2 | 0.098619819 | 0.38582221 | 0 |
| LD03534 | FBgn0032053 | mRpL51 | 0.098585004 | 0.50652933 | 0 |
| LD03179 | FBgn0043455 | CG5986 | 0.098553561 | 0.35347303 | 0 |
| AT09434 | FBgn0031849 | CG11327 | -0.098536409 | 0.55244011 | 0 |
| LD04071 | FBgn0036518 | RhoGAP71E | 0.098488669 | 0.35876322 | 0 |
| LD29166 | FBgn0037134 | CG7407 | 0.098452332 | 0.37374788 | 0 |
| LD44187 | FBgn0038472 | CG3995 | 0.098421434 | 0.42856874 | 0 |
| LP08864 | FBgn0037456 | CG1138 | -0.098413113 | 0.47263843 | 0 |
| GH25466 | FBgn0036698 | CG7724 | -0.09839854 | 0.36724764 | 0 |
| AT07117 | FBgn0037829 | CG14691 | -0.098337219 | 0.44917217 | 0 |
| AT21457 | FBgn0051294 | CG31294 | 0.098332867 | 0.47956863 | 0 |
| GH03113 | FBgn0025878 | wrapper | 0.098276162 | 0.70202124 | 0 |
| GH03873 | FBgn0030873 | CG15814 | 0.098259749 | 0.34494598 | 0 |
| LD35705 | FBgn0004597 | CycC | 0.098248937 | 0.36890521 | 0 |

| | | | | | |
|---|---|---|---|---|---|
| LD06016 | FBgn0032016 | Mettl14 | 0.098157291 | 0.35612816 | 0 |
| RE20268 | FBgn0037376 | Hat1 | -0.098148777 | 0.45015388 | 0 |
| LD01470 | FBgn0086362 | spn-F | 0.098136935 | 0.4322029 | 0 |
| LD24550 | FBgn0038588 | CG7156 | 0.098121778 | 0.34255324 | 0 |
| GH11587 | FBgn0031129 | CG1324 | 0.098085395 | 0.50054141 | 0 |
| SD07614 | FBgn0033247 | Nup44A | -0.09804135 | 0.35932238 | 0 |
| SD09294 | FBgn0035471 | Sc2 | 0.097985822 | 0.49579399 | 0 |
| RE02264 | FBgn0028436 | ECSIT | 0.09797234 | 0.39426214 | 0 |
| GH04035 | FBgn0036157 | CG7560 | 0.09796573 | 0.69569828 | 0 |
| GH20310 | FBgn0025335 | CG4585 | -0.097927279 | 0.35062898 | 0 |
| LD22701 | FBgn0004856 | Bx42 | 0.097922766 | 0.34994022 | 0 |
| RH34065 | FBgn0052160 | lncRNA:CR32160 | -0.097893568 | 0.38463262 | 0 |
| GH03691 | FBgn0000406 | Cyt-b5-r | 0.097888715 | 0.52246578 | 0 |
| LD26258 | FBgn0035379 | spz5 | -0.097883846 | 0.66347298 | 0 |
| GH08338 | FBgn0010406 | RNaseX25 | -0.097868285 | 0.42715664 | 0 |
| RE04580 | FBgn0030980 | CG7406 | -0.097824736 | 0.38667969 | 0 |
| LD25526 | FBgn0001105 | Gbeta13F | 0.097721221 | 0.36971494 | 0 |
| RH63089 | FBgn0041629 | Hexo2 | -0.097671217 | 0.59746344 | 0 |
| RE35371 | FBgn0036094 | CG14153 | -0.097632018 | 0.483335 | 0 |
| RE46710 | FBgn0004509 | Fur1 | -0.097575833 | 0.42613926 | 0 |
| GH07590 | FBgn0038321 | CG6218 | 0.09745513 | 0.45127659 | 0 |
| AT20942 | FBgn0033442 | CG1690 | -0.097435274 | 0.39157917 | 0 |
| LP03636 | FBgn0003046 | Pcp | -0.097430062 | 0.44252208 | 0 |
| GH25431 | FBgn0031127 | CG1835 | 0.097344695 | 0.63539356 | 0 |
| AT02829 | FBgn0031244 | CG11601 | 0.097330235 | 0.4005413 | 0 |
| LP11807 | FBgn0052793 | CG32793 | 0.097308458 | 0.55511879 | 0 |
| LD41606 | FBgn0015295 | Shark | -0.097304189 | 0.3940746 | 0 |
| LD16331 | FBgn0030391 | Rab40 | -0.097297808 | 0.43618956 | 0 |
| RE37654 | FBgn0029994 | Ldsdh1 | 0.097283314 | 0.63748581 | 0 |
| RE31839 | FBgn0035422 | RpL28 | 0.097276579 | 0.56666839 | 0 |
| LD10883 | FBgn0031216 | Zir | 0.097242147 | 0.60773166 | 0 |
| LD18233 | FBgn0037540 | Pbp95 | 0.097189306 | 0.34846053 | 0 |
| SD03914 | FBgn0263490 | mld | -0.097153957 | 0.39913672 | 0 |
| LD34623 | FBgn0036005 | pall | 0.097153006 | 0.35000557 | 0 |
| RE30523 | FBgn0011710 | Sep1 | 0.097018203 | 0.42276183 | 0 |
| RE37196 | FBgn0034391 | CG15080 | -0.097013818 | 0.48526469 | 0 |
| LD24627 | FBgn0051133 | Slimp | 0.09700026 | 0.38251295 | 0 |
| LD09921 | FBgn0026616 | alpha-Man-IIb | 0.096948669 | 0.36334792 | 0 |
| LD47425 | FBgn0043010 | Fsn | -0.096946111 | 0.36245059 | 0 |
| LD27581 | FBgn0036341 | Syx13 | -0.09693456 | 0.40221267 | 0 |
| LD19086 | FBgn0037724 | Fst | -0.096925162 | 0.49320822 | 0 |
| GH25976 | FBgn0036623 | CG4729 | 0.096910714 | 0.36059633 | 0 |
| LD38452 | FBgn0263108 | BtbVII | -0.096893074 | 0.40870454 | 0 |
| RE63030 | FBgn0035879 | GAPcenA | -0.096883861 | 0.41907068 | 0 |
| GM14243 | FBgn0036812 | Nufip | 0.096876342 | 0.44491731 | 0 |
| LD08201 | FBgn0029067 | Dd | 0.096847239 | 0.41963347 | 0 |
| LD43047 | FBgn0037924 | CG14712 | 0.096817009 | 0.48038085 | 0 |
| LD31377 | FBgn0024833 | AP-1mu | 0.096796692 | 0.3939846 | 0 |

| | | | | | |
|---|---|---|---|---|---|
| LD13959 | FBgn0025355 | SuUR | 0.096795506 | 0.39581893 | 0 |
| LD13959 | FBgn0266540 | CG45101 | 0.096795506 | 0.39581893 | 0 |
| LD21457 | FBgn0039836 | CG1750 | 0.096749693 | 0.46087433 | 0 |
| GH11908 | FBgn0046297 | lncRNA:CR9284 | 0.096624959 | 0.4441678 | 0 |
| RH14104 | FBgn0004783 | Ccp84Aa | 0.096562327 | 0.485618 | 0 |
| GH18986 | FBgn0038658 | CG14292 | -0.096488053 | 0.42125778 | 0 |
| LD36173 | FBgn0035589 | CHMP2B | 0.096438398 | 0.40094113 | 0 |
| SD17207 | FBgn0039613 | CG14527 | -0.096374383 | 0.40340611 | 0 |
| AT26401 | FBgn0032971 | ttm3 | -0.096366663 | 0.36733173 | 0 |
| RE14391 | FBgn0035771 | Sec63 | 0.096313107 | 0.36575027 | 0 |
| GM14292 | FBgn0029715 | CG11444 | 0.096312186 | 0.44131655 | 0 |
| LD31571 | FBgn0037566 | mRpL1 | 0.096283757 | 0.49346112 | 0 |
| GM02445 | FBgn0030678 | CG11679 | 0.096249846 | 0.45302394 | 0 |
| AT12437 | FBgn0050284 | CG30284 | 0.096177862 | 0.58996985 | 0 |
| LD06929 | FBgn0035721 | CG9948 | 0.096152874 | 0.57029141 | 0 |
| RH13652 | FBgn0034725 | CG6044 | 0.096145769 | 0.63776901 | 0 |
| RE26528 | FBgn0036592 | CG13049 | 0.096145683 | 0.48243833 | 0 |
| LP04958 | FBgn0040736 | IM3 | 0.096145682 | 0.38679813 | 0 |
| GH27401 | FBgn0033973 | HPS1 | -0.096144758 | 0.3546485 | 0 |
| RE52151 | FBgn0030035 | PIG-T | 0.096077922 | 0.36847667 | 0 |
| SD02507 | FBgn0263079 | CG43338 | -0.096047267 | 0.38251295 | 0 |
| GH06222 | FBgn0032156 | CG13124 | -0.096010159 | 0.59400557 | 0 |
| GM14611 | FBgn0032407 | Pex19 | 0.095989664 | 0.37728963 | 0 |
| RE39879 | FBgn0020643 | Lcp65Ab2 | -0.095956139 | 0.52148082 | 0 |
| RE39879 | FBgn0020644 | Lcp65Ab1 | -0.095956139 | 0.52148082 | 0 |
| RE30424 | FBgn0040323 | GNBP1 | 0.095929412 | 0.36969092 | 0 |
| GH10293 | FBgn0039228 | CG6980 | -0.09590923 | 0.35504469 | 0 |
| LD02192 | FBgn0037844 | CG4570 | 0.095894267 | 0.48017271 | 0 |
| GH26411 | FBgn0031023 | CG14200 | -0.095843869 | 0.47390358 | 0 |
| RH02904 | FBgn0284248 | Cyt-c-p | -0.095824291 | 0.54765386 | 0 |
| LD28068 | FBgn0039977 | CG17454 | 0.095823122 | 0.36600685 | 0 |
| GM22844 | FBgn0031985 | mon2 | 0.095822933 | 0.39118797 | 0 |
| LD27358 | FBgn0034626 | CG10795 | -0.095809801 | 0.39339733 | 0 |
| RE29623 | FBgn0031505 | ND-B14.5B | 0.095771996 | 0.38226482 | 0 |
| LD37882 | FBgn0050290 | Ppcdc | 0.095752784 | 0.46087433 | 0 |
| LD37882 | FBgn0260222 | CG42496 | 0.095752784 | 0.46087433 | 0 |
| AT21944 | FBgn0043865 | SA-2 | 0.095751266 | 0.36642713 | 0 |
| GM23845 | FBgn0037562 | CG11671 | 0.095750385 | 0.3881886 | 0 |
| RE08746 | FBgn0000229 | bsk | -0.09571401 | 0.60269094 | 0 |
| RE02574 | FBgn0027073 | CG4302 | 0.095668707 | 0.43176226 | 0 |
| RE67475 | FBgn0267486 | Ptp36E | -0.095661736 | 0.53139066 | 0 |
| LD44813 | FBgn0031498 | CG17260 | 0.095592009 | 0.35688979 | 0 |
| LD23129 | FBgn0264691 | Lst8 | -0.095553411 | 0.36404769 | 0 |
| RH65663 | FBgn0039557 | CG12259 | 0.095549733 | 0.37294475 | 0 |
| LD35121 | FBgn0039356 | CG5039 | -0.095353239 | 0.49839847 | 0 |
| LD21907 | FBgn0038252 | BigH1 | -0.095323162 | 0.43517385 | 0 |
| LD05893 | FBgn0030364 | Lsm12 | -0.095319172 | 0.47965207 | 0 |
| RE52048 | FBgn0015550 | tap | -0.095271953 | 0.45761209 | 0 |

| | | | | | |
|---|---|---|---|---|---|
| LP05719 | FBgn0033542 | CG12309 | 0.095247544 | 0.42145924 | 0 |
| AT27166 | FBgn0036924 | hale | 0.09512912 | 0.50325549 | 0 |
| LD42227 | FBgn0023522 | CG11596 | 0.095097542 | 0.37317796 | 0 |
| GH17465 | FBgn0030066 | CG1885 | 0.095047739 | 0.42895101 | 0 |
| LD09786 | FBgn0034217 | Lhr | 0.095032619 | 0.41072974 | 0 |
| SD17630 | FBgn0261859 | CG42788 | -0.095023737 | 0.37664429 | 0 |
| RH61745 | FBgn0030829 | CG12998 | 0.094997554 | 0.43004772 | 0 |
| GH11740 | FBgn0036553 | CG17027 | -0.09492177 | 0.38248754 | 0 |
| LD27979 | FBgn0266053 | Patr-1 | 0.094917472 | 0.38461723 | 0 |
| RE15313 | FBgn0051559 | CG31559 | 0.09490157 | 0.58180172 | 0 |
| GH03826 | FBgn0033459 | CG12744 | 0.094899953 | 0.51450559 | 0 |
| GM02640 | FBgn0000568 | Eip75B | -0.094893157 | 0.67523158 | 0 |
| RE39031 | FBgn0038774 | CG5023 | 0.094853402 | 0.410397 | 0 |
| RH50824 | FBgn0052137 | CG32137 | 0.094752935 | 0.4655014 | 0 |
| SD19581 | FBgn0038951 | CG5380 | 0.094747434 | 0.51668314 | 0 |
| GH23727 | FBgn0260795 | NaPi-III | -0.094744457 | 0.35617561 | 0 |
| LD34343 | FBgn0027492 | wdb | 0.094636262 | 0.39499648 | 0 |
| GH20107 | FBgn0041712 | yellow-d | -0.094563084 | 0.48529408 | 0 |
| LD30525 | FBgn0036892 | Lon | 0.094529809 | 0.40593563 | 0 |
| LD43716 | FBgn0015360 | oxt | 0.094427512 | 0.41157851 | 0 |
| LD01190 | FBgn0020412 | JIL-1 | 0.094422888 | 0.40947408 | 0 |
| RE13534 | FBgn0030410 | Aven | 0.09440886 | 0.42692336 | 0 |
| SD22389 | FBgn0038827 | Fancd2 | -0.094356147 | 0.35601985 | 0 |
| RE57682 | FBgn0034224 | insb | -0.094284 | 0.57988153 | 0 |
| GH23839 | FBgn0037044 | CG10585 | -0.094263774 | 0.38507121 | 0 |
| SD01637 | FBgn0034181 | CG8963 | 0.094186478 | 0.40655523 | 0 |
| LP10820 | FBgn0025624 | CG4025 | 0.094130251 | 0.3939846 | 0 |
| RE33981 | FBgn0250830 | CG12547 | -0.094045018 | 0.50874492 | 0 |
| HL02883 | FBgn0037279 | CG1129 | 0.093960922 | 0.37992892 | 0 |
| GH11459 | FBgn0039081 | Irk2 | 0.093955491 | 0.5620425 | 0 |
| AT01906 | FBgn0031275 | GABA-B-R3 | -0.093950494 | 0.37477926 | 0 |
| RE16882 | FBgn0013750 | Arf51F | -0.093934779 | 0.39219627 | 0 |
| LD22634 | FBgn0035811 | CG12262 | 0.093924229 | 0.42613926 | 0 |
| LD17744 | FBgn0260759 | CG42557 | -0.093884684 | 0.44868134 | 0 |
| LD17744 | FBgn0260760 | CG42558 | -0.093884684 | 0.44868134 | 0 |
| RH08414 | FBgn0029837 | Tsp5D | 0.093879393 | 0.39662446 | 0 |
| RE30552 | FBgn0263598 | Vha68-2 | -0.09372569 | 0.38251436 | 0 |
| GH01429 | FBgn0038546 | CG7379 | 0.093660626 | 0.36718327 | 0 |
| GH04025 | FBgn0016698 | Ptpa | 0.093653997 | 0.40819607 | 0 |
| SD02478 | FBgn0037120 | CG11247 | -0.093619194 | 0.47206345 | 0 |
| SD27303 | FBgn0052582 | lncRNA:CR32582 | 0.093609783 | 0.37597259 | 0 |
| LD19671 | FBgn0262103 | Sik3 | 0.093600181 | 0.41089595 | 0 |
| LD28763 | FBgn0030964 | Pvf1 | -0.093584043 | 0.39430382 | 0 |
| GM01846 | FBgn0036990 | mRpL15 | -0.093461522 | 0.45923276 | 0 |
| RE05239 | FBgn0002933 | ng1 | -0.093379694 | 0.40217921 | 0 |
| RE05239 | FBgn0010294 | ng2 | -0.093379694 | 0.40217921 | 0 |
| RE09301 | FBgn0037936 | CG6908 | -0.093341637 | 0.60714017 | 0 |
| GH18222 | FBgn0050359 | Mal-A5 | 0.093212181 | 0.44738846 | 0 |

| | | | | | |
|---|---|---|---|---|---|
| RH14088 | FBgn0032518 | RpL24 | 0.093197734 | 0.42014376 | 0 |
| GH25868 | FBgn0036489 | CG7011 | 0.093189487 | 0.3939846 | 0 |
| LP03126 | FBgn0267727 | Pen | -0.093146121 | 0.36724764 | 0 |
| SD03244 | FBgn0004057 | Zw | 0.093095101 | 0.39686379 | 0 |
| RE01052 | FBgn0030340 | CG15740 | -0.093090203 | 0.61194988 | 0 |
| RE56180 | FBgn0030089 | AP-1gamma | 0.093073026 | 0.36412204 | 0 |
| LD14015 | FBgn0035715 | CG10103 | 0.093029947 | 0.37424588 | 0 |
| RH43346 | FBgn0085370 | Pde11 | 0.09300706 | 0.42884219 | 0 |
| SD08605 | FBgn0031490 | CG17264 | 0.093001952 | 0.48048527 | 0 |
| LD03515 | FBgn0037655 | Kcmf1 | 0.092946047 | 0.54324174 | 0 |
| LD39211 | FBgn0033236 | CG14764 | 0.092938961 | 0.3693818 | 0 |
| LD40663 | FBgn0086358 | Tab2 | 0.092892776 | 0.47773771 | 0 |
| LD19634 | FBgn0036386 | CG8833 | -0.092820149 | 0.47213318 | 0 |
| GH23626 | FBgn0030589 | CG9519 | 0.092787971 | 0.68281253 | 0 |
| LD05623 | FBgn0046692 | Stlk | -0.092718461 | 0.37116135 | 0 |
| LP07257 | FBgn0031004 | CG7992 | -0.092712766 | 0.38822579 | 0 |
| SD08002 | FBgn0260779 | GatA | 0.092696741 | 0.37187434 | 0 |
| RE34075 | FBgn0038527 | CG14324 | -0.092690105 | 0.56767477 | 0 |
| AT16729 | FBgn0037195 | CG11226 | 0.092672503 | 0.5956691 | 0 |
| GH13231 | FBgn0040343 | CG3713 | -0.092666509 | 0.41551801 | 0 |
| LD47509 | FBgn0010220 | Dbp45A | 0.09264728 | 0.37064794 | 0 |
| AT24389 | FBgn0037239 | Sfxn1-3 | -0.092629321 | 0.37872757 | 0 |
| LD04782 | FBgn0034072 | Dg | 0.092625276 | 0.43791662 | 0 |
| GH09173 | FBgn0020545 | kraken | 0.092608211 | 0.43026786 | 0 |
| LD18613 | FBgn0032955 | CG2201 | -0.092572021 | 0.37722091 | 0 |
| RH06755 | FBgn0003861 | trp | -0.09249732 | 0.50019526 | 0 |
| GH02982 | FBgn0024997 | CG2681 | -0.092488431 | 0.43772294 | 0 |
| GH07188 | FBgn0027578 | CG14526 | 0.092365921 | 0.36733173 | 0 |
| LP01106 | FBgn0261811 | pico | 0.092321746 | 0.44663971 | 0 |
| GH27912 | FBgn0028582 | lqf | -0.092321493 | 0.38829968 | 0 |
| LD06034 | FBgn0002121 | l(2)gl | 0.09228061 | 0.38582199 | 0 |
| LD32459 | FBgn0034084 | CG8435 | 0.092219187 | 0.59548176 | 0 |
| GH23891 | FBgn0037230 | CG9780 | 0.092193645 | 0.39366894 | 0 |
| AT16114 | FBgn0032094 | CG12439 | 0.092193316 | 0.63878328 | 0 |
| GH08116 | FBgn0015039 | Cyp9b2 | 0.092186551 | 0.72762315 | 0 |
| GH22104 | FBgn0029791 | CG4096 | -0.092181076 | 0.42092985 | 0 |
| AT18413 | FBgn0029656 | CG10793 | 0.092139366 | 0.44712988 | 0 |
| AT28104 | FBgn0028868 | CG4701 | 0.092126357 | 0.44868134 | 0 |
| GH22889 | FBgn0038919 | CG17843 | 0.092116459 | 0.70788092 | 0 |
| RH31482 | FBgn0030571 | CG14414 | 0.092095093 | 0.40940941 | 0 |
| LD26817 | FBgn0082598 | akirin | 0.092084413 | 0.38582221 | 0 |
| LD44422 | FBgn0036063 | CG6674 | 0.092033474 | 0.47068547 | 0 |
| GH07145 | FBgn0036545 | GXIVsPLA2 | -0.09192728 | 0.48445544 | 0 |
| RE41364 | FBgn0027066 | Eb1 | 0.09190693 | 0.45133957 | 0 |
| RH13728 | FBgn0030828 | CG5162 | -0.091890481 | 0.46514252 | 0 |
| LP08411 | FBgn0030396 | CG2556 | 0.09188279 | 0.39979556 | 0 |
| LD18062 | FBgn0260398 | Pbp49 | 0.091858026 | 0.40870454 | 0 |
| RH47707 | FBgn0003867 | tsl | 0.091856915 | 0.6944496 | 0 |

| | | | | | |
|---|---|---|---|---|---|
| GH10915 | FBgn0036126 | Irbp18 | -0.091841304 | 0.5123227 | 0 |
| RE66843 | FBgn0285937 | Rab1 | -0.091833189 | 0.40688164 | 0 |
| LD30968 | FBgn0038296 | CG6752 | 0.091832918 | 0.43267322 | 0 |
| RH12261 | FBgn0032150 | CG13123 | -0.091792819 | 0.48523766 | 0 |
| GH13765 | FBgn0040350 | CG3690 | 0.091780288 | 0.63748581 | 0 |
| RH63144 | FBgn0061362 | CG33785 | -0.091770727 | 0.46847056 | 0 |
| RH63144 | FBgn0061361 | CG33786 | -0.091770727 | 0.46847056 | 0 |
| RH49505 | FBgn0037749 | CG9471 | 0.091749151 | 0.48592281 | 0 |
| LD29155 | FBgn0038268 | CG3631 | -0.091681812 | 0.55948171 | 0 |
| GH03014 | FBgn0042135 | CG18812 | -0.091624729 | 0.50628892 | 0 |
| LP10505 | FBgn0038873 | CG5892 | 0.091554161 | 0.37936225 | 0 |
| GH28883 | FBgn0030997 | CG7990 | 0.091420993 | 0.37458406 | 0 |
| GH10029 | FBgn0052549 | CG32549 | 0.091380676 | 0.49226865 | 0 |
| LD33361 | FBgn0010762 | simj | 0.091267621 | 0.45483857 | 0 |
| SD21917 | FBgn0039833 | CG15564 | 0.091259253 | 0.39359496 | 0 |
| RE64552 | FBgn0030300 | Sk1 | 0.091227762 | 0.40221267 | 0 |
| GH12014 | FBgn0029753 | CG4198 | -0.091207025 | 0.42681424 | 0 |
| LD03691 | FBgn0039465 | Tsp97E | 0.09118923 | 0.36885296 | 0 |
| GM14169 | FBgn0026326 | Mad1 | 0.091160305 | 0.39552963 | 0 |
| LD30573 | FBgn0037530 | EMC1 | 0.09110335 | 0.43339125 | 0 |
| GM13047 | FBgn0002590 | RpS5a | 0.09107699 | 0.45761209 | 0 |
| AT28243 | FBgn0037170 | Trxr-2 | -0.091049392 | 0.42950889 | 0 |
| GH26238 | FBgn0260470 | SP555 | 0.091040441 | 0.39333702 | 0 |
| GH26238 | FBgn0260451 | CG14042 | 0.091040441 | 0.39333702 | 0 |
| SD07795 | FBgn0022764 | Sin3A | 0.091031953 | 0.3939846 | 0 |
| LD47926 | FBgn0039039 | lmd | -0.091012278 | 0.61080847 | 0 |
| LD14730 | FBgn0051673 | CG31673 | 0.090988152 | 0.4496302 | 0 |
| RH01575 | FBgn0032652 | CG6870 | -0.090987958 | 0.42284175 | 0 |
| RE14437 | FBgn0032515 | loqs | 0.090981049 | 0.4130867 | 0 |
| GH03003 | FBgn0037553 | CG18249 | -0.090890486 | 0.55917167 | 0 |
| LD29295 | FBgn0031044 | MKP-4 | 0.090878244 | 0.44855728 | 0 |
| LD20458 | FBgn0266347 | nAChRalpha4 | -0.0908514 | 0.52318403 | 0 |
| LD23744 | FBgn0037099 | CG7173 | -0.090644657 | 0.63793196 | 0 |
| AT30562 | FBgn0031108 | CG15459 | -0.09062141 | 0.59614743 | 0 |
| RE01412 | FBgn0026432 | Grip163 | 0.090613683 | 0.40543771 | 0 |
| LD13915 | FBgn0033995 | GPHR | -0.090603551 | 0.40819607 | 0 |
| RE67551 | FBgn0264895 | RapGAP1 | 0.090587117 | 0.39817928 | 0 |
| RE18294 | FBgn0037408 | NPFR | -0.090582296 | 0.46705356 | 0 |
| RE50726 | FBgn0031708 | CG7382 | 0.090576177 | 0.52418589 | 0 |
| HL03177 | FBgn0267435 | chp | 0.090562946 | 0.51114596 | 0 |
| LP10938 | FBgn0034710 | CG3292 | -0.090535816 | 0.74370858 | 0 |
| RE32166 | FBgn0032248 | Bug22 | -0.090533536 | 0.65498342 | 0 |
| GH22881 | FBgn0031859 | CG17377 | -0.090445977 | 0.49826051 | 0 |
| AT22782 | FBgn0051826 | CG31826 | 0.090408087 | 0.58302715 | 0 |
| LP05464 | FBgn0250851 | CG33981 | 0.090406102 | 0.40705659 | 0 |
| LP05464 | FBgn0028741 | fab1 | 0.090406102 | 0.40705659 | 0 |
| LP08751 | FBgn0040367 | CG11382 | 0.090402152 | 0.47568595 | 0 |
| RE52162 | FBgn0051760 | CG31760 | -0.090398034 | 0.40581357 | 0 |

| | | | | | |
|---|---|---|---|---|---|
| RE12138 | FBgn0040363 | CG11384 | -0.090358582 | 0.48928658 | 0 |
| GH07688 | FBgn0027569 | cert | 0.090354999 | 0.37832142 | 0 |
| GH07816 | FBgn0032935 | Atg18b | 0.090314787 | 0.42676291 | 0 |
| LP02990 | FBgn0259818 | CG42399 | 0.090258663 | 0.44157108 | 0 |
| GH13192 | FBgn0033065 | Cyp6w1 | 0.090252651 | 0.54968015 | 0 |
| RH42893 | FBgn0038181 | CG9297 | 0.090123701 | 0.37459627 | 0 |
| RE06354 | FBgn0038034 | Cyp9f3 | -0.089999128 | 0.69553036 | 0 |
| RE12209 | FBgn0029506 | Tsp42Ee | -0.089975464 | 0.48577999 | 0 |
| LD39232 | FBgn0263050 | CG43345 | 0.089970674 | 0.4344621 | 0 |
| LD39232 | FBgn0263051 | CG43346 | 0.089970674 | 0.4344621 | 0 |
| AT18037 | FBgn0035384 | CG2113 | 0.089945005 | 0.61374063 | 0 |
| GH17442 | FBgn0033502 | CG12910 | 0.089823909 | 0.47068547 | 0 |
| GH09946 | FBgn0033799 | GLaz | -0.089778988 | 0.60417516 | 0 |
| LD12037 | FBgn0031458 | aph-1 | 0.089752342 | 0.44532787 | 0 |
| GH03514 | FBgn0028662 | VhaPPA1-1 | -0.089728921 | 0.37558581 | 0 |
| RE17056 | FBgn0036580 | PDCD-5 | 0.089726076 | 0.51140355 | 0 |
| LD39079 | FBgn0260748 | CG5004 | 0.089712092 | 0.41812284 | 0 |
| AT13267 | FBgn0003997 | hid | 0.089702861 | 0.59937435 | 0 |
| RH28218 | FBgn0032008 | CG14277 | -0.089678836 | 0.48583439 | 0 |
| LD09733 | FBgn0037634 | hng2 | 0.089668435 | 0.44823738 | 0 |
| RE37995 | FBgn0039588 | mIF2 | 0.089654723 | 0.42984502 | 0 |
| RE37995 | FBgn0058068 | CR40068 | 0.089654723 | 0.42984502 | 0 |
| AT22204 | FBgn0032483 | CG15482 | 0.089629115 | 0.50642541 | 0 |
| RE54341 | FBgn0037065 | CG12974 | 0.089576744 | 0.43077684 | 0 |
| LD06557 | FBgn0034430 | mip40 | 0.089548268 | 0.38272145 | 0 |
| SD02490 | FBgn0035145 | MED14 | 0.089524987 | 0.39339733 | 0 |
| LD44815 | FBgn0030963 | CG7101 | 0.089507555 | 0.40643756 | 0 |
| LD34464 | FBgn0026238 | gus | -0.089453348 | 0.44868134 | 0 |
| GH14313 | FBgn0034792 | YME1L | 0.089410973 | 0.45287985 | 0 |
| RE26442 | FBgn0040507 | ACXD | -0.089408427 | 0.49425278 | 0 |
| LD08826 | FBgn0033547 | CG12935 | -0.089311772 | 0.43376409 | 0 |
| LP07235 | FBgn0015571 | alpha-Est3 | -0.089261329 | 0.49293909 | 0 |
| GH04120 | FBgn0028499 | CG7985 | 0.089255502 | 0.41638474 | 0 |
| GH26080 | FBgn0034687 | CG11475 | -0.089225973 | 0.40617197 | 0 |
| GH05095 | FBgn0028494 | CG6424 | 0.089212525 | 0.40744996 | 0 |
| GH05095 | FBgn0284231 | CG46315 | 0.089212525 | 0.40744996 | 0 |
| RE22703 | FBgn0265187 | CG44252 | -0.089203861 | 0.51105834 | 0 |
| GH14476 | FBgn0035034 | CG3565 | 0.08916846 | 0.68006514 | 0 |
| RE53348 | FBgn0263447 | asRNA:CR43470 | 0.089005503 | 0.67670719 | 0 |
| HL07956 | FBgn0026741 | mRpL18 | -0.088988284 | 0.49704094 | 0 |
| LD23193 | FBgn0022069 | Nnp-1 | 0.088925986 | 0.43116435 | 0 |
| LD24837 | FBgn0036187 | RIOK1 | 0.088909478 | 0.37862901 | 0 |
| AT09126 | FBgn0050271 | CG30271 | -0.088867316 | 0.47294983 | 0 |
| GH21490 | FBgn0034247 | CG6484 | -0.088865468 | 0.40080031 | 0 |
| GH01881 | FBgn0014018 | Rel | -0.088840725 | 0.4655033 | 0 |
| AT23096 | FBgn0029501 | Crtp | -0.088646077 | 0.43154461 | 0 |
| GH04113 | FBgn0033423 | CG1809 | 0.08855778 | 0.4371378 | 0 |
| LP03268 | FBgn0033853 | CG6145 | -0.088552904 | 0.49154345 | 0 |

| | | | | | |
|---|---|---|---|---|---|
| LD23523 | FBgn0032671 | JMJD4 | 0.088518969 | 0.55877041 | 0 |
| RH38008 | FBgn0031561 | IM33 | 0.088300544 | 0.72375908 | 0 |
| LP01967 | FBgn0031321 | Tgt | 0.088267734 | 0.40243057 | 0 |
| GH17337 | FBgn0027621 | Pfrx | 0.088200449 | 0.42169192 | 0 |
| SD07020 | FBgn0053100 | eIF4EHP | -0.088198839 | 0.52641863 | 0 |
| GH05321 | FBgn0035670 | CG10472 | 0.088197401 | 0.48135302 | 0 |
| LD19190 | FBgn0029713 | CG11436 | -0.088136073 | 0.45519247 | 0 |
| GH01085 | FBgn0001404 | egh | 0.08809627 | 0.40406672 | 0 |
| GM10122 | FBgn0028434 | Ercc1 | -0.088081169 | 0.49895529 | 0 |
| RE31004 | FBgn0260008 | UQCR-11 | -0.088076432 | 0.56091028 | 0 |
| RE16407 | FBgn0013954 | FK506-bp2 | 0.08804675 | 0.41252617 | 0 |
| RE38972 | FBgn0019936 | RpS20 | -0.088039808 | 0.5630283 | 0 |
| AT03279 | FBgn0023415 | Acp32CD | 0.087965315 | 0.38902664 | 0 |
| AT28366 | FBgn0035143 | Ppm1 | 0.087934888 | 0.53005697 | 0 |
| GM12605 | FBgn0030468 | CG1622 | 0.08793406 | 0.38839659 | 0 |
| RE73208 | FBgn0032022 | CG14275 | 0.08790909 | 0.49425278 | 0 |
| LP02621 | FBgn0266756 | btsz | -0.087825579 | 0.73772784 | 0 |
| RE39081 | FBgn0041156 | exex | -0.087814472 | 0.43791662 | 0 |
| RE72392 | FBgn0038029 | GstD11 | 0.087801815 | 0.53152622 | 0 |
| LD11109 | FBgn0040324 | Ephrin | -0.087758294 | 0.53326482 | 0 |
| AT11276 | FBgn0039501 | TTLL6B | 0.087719606 | 0.64223488 | 0 |
| GH19382 | FBgn0034137 | CG4945 | 0.087717162 | 0.67181984 | 0 |
| RH04437 | FBgn0038819 | Cpr92F | 0.087695411 | 0.54219341 | 0 |
| AT03621 | FBgn0036072 | CG6628 | 0.087695216 | 0.64411479 | 0 |
| LD20464 | FBgn0260722 | lncRNA:CR42549 | 0.087694865 | 0.5868984 | 0 |
| LD39658 | FBgn0028425 | JhI-21 | 0.087656159 | 0.45308696 | 0 |
| LD21681 | FBgn0013548 | l(2)dtl | 0.087638529 | 0.44459953 | 0 |
| AT08135 | FBgn0034477 | CG13872 | 0.0876239 | 0.42681424 | 0 |
| LD12022 | FBgn0005683 | pie | -0.087549759 | 0.44428137 | 0 |
| GH11567 | FBgn0036038 | defl | -0.087547102 | 0.49845238 | 0 |
| RE73673 | FBgn0031682 | CG5828 | -0.087546183 | 0.38582221 | 0 |
| LD15458 | FBgn0001941 | ifc | 0.087527906 | 0.41331428 | 0 |
| RE58715 | FBgn0051221 | CG31221 | 0.087487227 | 0.43898429 | 0 |
| LD41433 | FBgn0023512 | eIF2Bepsilon | 0.087450277 | 0.42805285 | 0 |
| RH64745 | FBgn0004045 | Yp1 | 0.087430492 | 0.58332023 | 0 |
| LD23881 | FBgn0039019 | HP1c | 0.087331243 | 0.44981111 | 0 |
| RE51076 | FBgn0052499 | Cda4 | 0.087322486 | 0.70415705 | 0 |
| GH25733 | FBgn0037962 | CG6971 | 0.087269091 | 0.39339733 | 0 |
| RE40534 | FBgn0033226 | CG1882 | 0.087249391 | 0.43822876 | 0 |
| GH12430 | FBgn0030796 | CG4829 | 0.087194 | 0.8305917 | 0 |
| RE13549 | FBgn0039049 | CG6726 | 0.087105254 | 0.4427422 | 0 |
| LD20590 | FBgn0011335 | vers | 0.087028737 | 0.4344621 | 0 |
| GM13434 | FBgn0030048 | CG12112 | 0.087014318 | 0.43618956 | 0 |
| GH14092 | FBgn0031375 | erm | -0.08697549 | 0.5602318 | 0 |
| GH18129 | FBgn0025830 | IntS8 | -0.08687268 | 0.39366894 | 0 |
| GM16166 | FBgn0260767 | CG42565 | -0.086837973 | 0.44131655 | 0 |
| GM16166 | FBgn0034759 | CG13511 | -0.086837973 | 0.44131655 | 0 |
| GM16166 | FBgn0034758 | CG13510 | -0.086837973 | 0.44131655 | 0 |

| | | | | | |
|---|---|---|---|---|---|
| LD23602 | FBgn0036641 | Smn | 0.086817511 | 0.45923276 | 0 |
| LP08566 | FBgn0030748 | Traf-like | 0.086792887 | 0.49334127 | 0 |
| LP11788 | FBgn0039527 | CG5639 | -0.086713002 | 0.48493795 | 0 |
| GM14815 | FBgn0033260 | Cul4 | -0.086615539 | 0.42145924 | 0 |
| LD11064 | FBgn0003321 | sbr | 0.086586309 | 0.39339733 | 0 |
| LD20933 | FBgn0015019 | CCT3 | -0.086571346 | 0.41394421 | 0 |
| GH24640 | FBgn0037896 | ninaG | 0.086559735 | 0.45051871 | 0 |
| RE51969 | FBgn0034590 | Magi | 0.086540695 | 0.38987282 | 0 |
| RE36989 | FBgn0031676 | senju | -0.086533024 | 0.49867738 | 0 |
| GH11862 | FBgn0031995 | CG8475 | -0.086455308 | 0.48354193 | 0 |
| RE59371 | FBgn0033481 | CG12920 | 0.086448198 | 0.43887794 | 0 |
| LD46328 | FBgn0053094 | Synd | 0.086371041 | 0.41089595 | 0 |
| GH20281 | FBgn0036806 | Cyp12c1 | -0.08630357 | 0.51386113 | 0 |
| GH18520 | FBgn0038344 | obe | 0.08624684 | 0.40479587 | 0 |
| RE23490 | FBgn0261799 | dsx-c73A | -0.086245965 | 0.48802258 | 0 |
| AT19278 | FBgn0284244 | l(2)k05911 | 0.086166631 | 0.5700488 | 0 |
| LD28808 | FBgn0037834 | Art1 | -0.086131835 | 0.39662446 | 0 |
| LD39959 | FBgn0035848 | Uxs | 0.086120897 | 0.42681424 | 0 |
| LD21601 | FBgn0250791 | alphaSnap | 0.086090358 | 0.48539549 | 0 |
| LD41421 | FBgn0030323 | Vlet | -0.086066718 | 0.58302715 | 0 |
| RE39488 | FBgn0261933 | SmD1 | -0.08601864 | 0.42894764 | 0 |
| RE59171 | FBgn0031710 | Vps52 | -0.085987306 | 0.44464685 | 0 |
| SD02553 | FBgn0038478 | cal1 | -0.085975626 | 0.45794217 | 0 |
| RE60364 | FBgn0029881 | pigs | 0.085904644 | 0.41561017 | 0 |
| RE41194 | FBgn0030929 | CG15043 | -0.085879313 | 0.70367871 | 0 |
| RH51096 | FBgn0031609 | CG15443 | -0.085832184 | 0.42117179 | 0 |
| RH54729 | FBgn0267033 | mamo | -0.085811081 | 0.46222895 | 0 |
| LD29477 | FBgn0086253 | rumi | 0.085743252 | 0.44000308 | 0 |
| GH08893 | FBgn0039265 | CG11790 | -0.085712116 | 0.43839377 | 0 |
| SD05618 | FBgn0267978 | ap | 0.085711859 | 0.63262065 | 0 |
| GM21150 | FBgn0030447 | CG2200 | -0.085618638 | 0.50444486 | 0 |
| AT07595 | FBgn0039821 | CG15556 | 0.085581899 | 0.56000711 | 0 |
| LD31422 | FBgn0264607 | CaMKII | -0.085573806 | 0.46890529 | 0 |
| AT04835 | FBgn0032525 | Hsp60D | -0.085531604 | 0.48592281 | 0 |
| RH37427 | FBgn0266722 | Trs33 | 0.085459342 | 0.43339125 | 0 |
| LP08941 | FBgn0030955 | CG6891 | 0.085458475 | 0.59911737 | 0 |
| AT12351 | FBgn0036437 | CG5048 | 0.085384271 | 0.44458159 | 0 |
| RE61693 | FBgn0010287 | Trf | 0.085375032 | 0.42951104 | 0 |
| LD46533 | FBgn0032643 | GCS2beta | 0.085355431 | 0.44868134 | 0 |
| LD27413 | FBgn0034002 | CG8079 | 0.085304201 | 0.40448993 | 0 |
| AT02326 | FBgn0033322 | CG8584 | 0.085279616 | 0.49676491 | 0 |
| SD05267 | FBgn0263601 | mib1 | -0.085230726 | 0.39886621 | 0 |
| LD21545 | FBgn0033475 | CG12129 | -0.085223643 | 0.50138518 | 0 |
| SD02279 | FBgn0024250 | brk | -0.085199494 | 0.63569444 | 0 |
| RH47207 | FBgn0283450 | Glo1 | 0.085130169 | 0.52195992 | 0 |
| RE54265 | FBgn0035777 | CG8563 | 0.085079317 | 0.5060605 | 0 |
| AT31112 | FBgn0032269 | w-cup | 0.085070494 | 0.50766723 | 0 |
| RE65163 | FBgn0032656 | CG5674 | -0.08496642 | 0.44538287 | 0 |

| | | | | | |
|---|---|---|---|---|---|
| LD41462 | FBgn0030456 | CG4332 | 0.084879684 | 0.46238255 | 0 |
| GH06679 | FBgn0030481 | CG1662 | 0.0848695 | 0.41906431 | 0 |
| RE19849 | FBgn0051675 | CG31675 | 0.084807323 | 0.49640161 | 0 |
| LP04889 | FBgn0036762 | CG7430 | 0.084783628 | 0.4577751 | 0 |
| GH13594 | FBgn0037939 | CG14718 | -0.084735587 | 0.44157108 | 0 |
| AT11581 | FBgn0033680 | CG13186 | 0.084700939 | 0.59804839 | 0 |
| RE41712 | FBgn0038491 | CG5292 | -0.084684821 | 0.43523511 | 0 |
| RE58741 | FBgn0037979 | GCC185 | -0.084639782 | 0.49334127 | 0 |
| AT30481 | FBgn0032637 | CG5050 | 0.084630974 | 0.41157851 | 0 |
| RE14116 | FBgn0002736 | mago | -0.084623267 | 0.50842938 | 0 |
| RE40159 | FBgn0035583 | CG13704 | 0.084612168 | 0.69780063 | 0 |
| GH05104 | FBgn0039151 | CG13607 | -0.084604487 | 0.5996356 | 0 |
| GH13848 | FBgn0035497 | CG14995 | -0.084541319 | 0.46859987 | 0 |
| LP08082 | FBgn0037894 | Ranbp9 | 0.084491116 | 0.4068281 | 0 |
| GH17088 | FBgn0030688 | CG8952 | 0.084409088 | 0.59701303 | 0 |
| LD30746 | FBgn0061492 | loj | -0.08440228 | 0.69165887 | 0 |
| RE26043 | FBgn0023001 | melt | -0.084302377 | 0.72226794 | 0 |
| LD02379 | FBgn0015776 | nrv1 | -0.084290781 | 0.57430725 | 0 |
| LD22034 | FBgn0039099 | GILT2 | 0.084285647 | 0.47795046 | 0 |
| RE59992 | FBgn0035462 | IntS10 | 0.08426445 | 0.40539538 | 0 |
| RE02247 | FBgn0003975 | vg | 0.084223254 | 0.50711874 | 0 |
| LD24026 | FBgn0015834 | eIF3i | 0.084149685 | 0.52406854 | 0 |
| LD29806 | FBgn0032724 | CG10428 | 0.084123479 | 0.42276183 | 0 |
| GM25129 | FBgn0266042 | asRNA:CR44807 | 0.084095213 | 0.5466194 | 0 |
| RE54280 | FBgn0004666 | sim | -0.084084534 | 0.49397432 | 0 |
| LD39624 | FBgn0034971 | CG3209 | 0.084076148 | 0.4047529 | 0 |
| HL07915 | FBgn0033724 | CG8501 | 0.084065262 | 0.65315239 | 0 |
| RE69682 | FBgn0030806 | CG13001 | 0.084055149 | 0.42306953 | 0 |
| RE01054 | FBgn0037429 | Osi19 | -0.084019238 | 0.65712854 | 0 |
| RE59221 | FBgn0036833 | CG3819 | 0.084010589 | 0.4176333 | 0 |
| GH05530 | FBgn0267726 | CG46059 | -0.084000262 | 0.70559789 | 0 |
| RE56495 | FBgn0014877 | Roe1 | 0.083945628 | 0.53487304 | 0 |
| AT07424 | FBgn0029750 | CG3323 | 0.083937169 | 0.49596037 | 0 |
| LP01608 | FBgn0033887 | St4 | 0.083913431 | 0.48172108 | 0 |
| LD23656 | FBgn0033935 | Sin1 | -0.083909795 | 0.47660089 | 0 |
| LP01489 | FBgn0026179 | siz | -0.083853967 | 0.45662682 | 0 |
| LD45288 | FBgn0260855 | Sec22 | 0.083831432 | 0.42681424 | 0 |
| LD21896 | FBgn0030805 | wus | -0.083808713 | 0.44489043 | 0 |
| LD23243 | FBgn0011766 | E2f1 | -0.083778316 | 0.48315697 | 0 |
| GH01369 | FBgn0030961 | CG7058 | -0.083666646 | 0.66102494 | 0 |
| RE68357 | FBgn0035722 | CG10075 | -0.083652687 | 0.60019681 | 0 |
| RE03173 | FBgn0001189 | hfw | 0.08358931 | 0.52296023 | 0 |
| RH29859 | FBgn0026873 | MED18 | -0.083558894 | 0.4153913 | 0 |
| LD04047 | FBgn0032130 | CG3838 | 0.083557754 | 0.42125778 | 0 |
| AT30085 | FBgn0028513 | CG9254 | 0.083543506 | 0.59783502 | 0 |
| RE43077 | FBgn0001330 | kz | 0.083540669 | 0.47768899 | 0 |
| SD02883 | FBgn0266669 | Sec3 | 0.083535374 | 0.42884219 | 0 |
| RE62247 | FBgn0000233 | btd | -0.083518838 | 0.71134843 | 0 |

| | | | | | |
|---|---|---|---|---|---|
| LD22131 | FBgn0020386 | Pdk1 | 0.083510492 | 0.413586 | 0 |
| LD08703 | FBgn0028387 | chm | 0.083502814 | 0.44870155 | 0 |
| RH68195 | FBgn0032195 | CG4908 | -0.08347457 | 0.49067383 | 0 |
| SD09259 | FBgn0039674 | CG1907 | 0.083442985 | 0.4770526 | 0 |
| RE15216 | FBgn0053217 | CG33217 | -0.083379706 | 0.58228008 | 0 |
| HL07907 | FBgn0263077 | CG43340 | 0.08335966 | 0.69997899 | 0 |
| AT16542 | FBgn0036482 | CG13457 | 0.083341073 | 0.48316831 | 0 |
| RE19570 | FBgn0034744 | Vps20 | -0.083288943 | 0.42676291 | 0 |
| AT25933 | FBgn0052686 | CG32686 | 0.083278947 | 0.44782334 | 0 |
| GH14210 | FBgn0038803 | CG5191 | 0.083246661 | 0.42459935 | 0 |
| AT02512 | FBgn0023545 | PIG-K | -0.083236207 | 0.41630253 | 0 |
| LD43670 | FBgn0023510 | Rbcn-3B | -0.083213711 | 0.40698557 | 0 |
| LD03453 | FBgn0004921 | Ggamma1 | 0.083201952 | 0.54968015 | 0 |
| GM03427 | FBgn0038679 | CG6040 | 0.083201359 | 0.65386861 | 0 |
| SD08175 | FBgn0013753 | Bgb | 0.083195164 | 0.47331034 | 0 |
| LD24646 | FBgn0051453 | pch2 | 0.083149633 | 0.45127659 | 0 |
| AT01876 | FBgn0035730 | CG17744 | 0.083132318 | 0.65231546 | 0 |
| RH09485 | FBgn0062961 | lncRNA:CR33942 | 0.083102413 | 0.48842433 | 0 |
| LD01705 | FBgn0032198 | eEF1delta | 0.083094095 | 0.49180042 | 0 |
| RE28512 | FBgn0028938 | CG16886 | -0.08308422 | 0.62723084 | 0 |
| RE10615 | FBgn0003008 | or | 0.08305603 | 0.4754937 | 0 |
| GH05767 | FBgn0033187 | PIG-G | -0.083039293 | 0.43087418 | 0 |
| RE08605 | FBgn0033170 | sPLA2 | 0.082999043 | 0.51270727 | 0 |
| LD33880 | FBgn0032430 | CG6388 | -0.08298201 | 0.47068547 | 0 |
| RE25723 | FBgn0030864 | CG8173 | 0.082971337 | 0.45330723 | 0 |
| GH09123 | FBgn0038292 | CG3987 | 0.082949275 | 0.57449724 | 0 |
| AT02742 | FBgn0038960 | CG13855 | -0.082942201 | 0.50815974 | 0 |
| SD03094 | FBgn0036448 | mop | 0.082825741 | 0.55917167 | 0 |
| LD30178 | FBgn0039728 | CG7896 | 0.08280775 | 0.42276183 | 0 |
| RE35481 | FBgn0051102 | CG31102 | -0.082783521 | 0.63329257 | 0 |
| LD19356 | FBgn0037358 | elm | -0.082778363 | 0.51833128 | 0 |
| RH61375 | FBgn0003462 | Sod1 | -0.082730487 | 0.84422131 | 0 |
| LD07058 | FBgn0036536 | CG12713 | 0.082642599 | 0.56089775 | 0 |
| LD31220 | FBgn0259174 | Nedd4 | -0.082622121 | 0.52195992 | 0 |
| LP02603 | FBgn0031910 | CG15818 | 0.082615028 | 0.66405539 | 0 |
| GH25462 | FBgn0052450 | CG32450 | 0.082584718 | 0.64746163 | 0 |
| SD07613 | FBgn0033205 | CG2064 | -0.082558618 | 0.57999504 | 0 |
| GH14349 | FBgn0036680 | Cpr73D | 0.082421688 | 0.51157454 | 0 |
| LD31474 | FBgn0030718 | ND-20 | 0.082385717 | 0.47918187 | 0 |
| GM02602 | FBgn0260944 | Rbp1 | 0.082366267 | 0.53255122 | 0 |
| RE24635 | FBgn0032282 | CG7299 | 0.082357212 | 0.48539549 | 0 |
| LD44982 | FBgn0034351 | rswl | 0.082334842 | 0.45433285 | 0 |
| GH27380 | FBgn0039844 | CG1607 | 0.082302251 | 0.64133825 | 0 |
| GH27361 | FBgn0036742 | CG7497 | -0.08225005 | 0.72458946 | 0 |
| GH11326 | FBgn0000121 | Arr2 | -0.082209316 | 0.4176333 | 0 |
| LD23642 | FBgn0025720 | Ate1 | 0.082206934 | 0.53800958 | 0 |
| LD45876 | FBgn0025836 | RhoGAP1A | 0.082174599 | 0.44577837 | 0 |
| GH04692 | FBgn0052573 | CG32573 | 0.082108447 | 0.42963885 | 0 |

| | | | | | |
|---|---|---|---|---|---|
| GH04692 | FBgn0260238 | CG42512 | 0.082108447 | 0.42963885 | 0 |
| RE19904 | FBgn0037609 | CG9773 | 0.082105266 | 0.49872991 | 0 |
| LD24265 | FBgn0033879 | CG6543 | 0.082095064 | 0.53394739 | 0 |
| GH13023 | FBgn0260386 | mtg | -0.082031228 | 0.46047599 | 0 |
| LD29161 | FBgn0026199 | myo | -0.082004497 | 0.44917217 | 0 |
| RE14405 | FBgn0032297 | CG17124 | -0.082000834 | 0.51237278 | 0 |
| GH15416 | FBgn0053143 | CG33143 | -0.081967368 | 0.4420803 | 0 |
| LP05296 | FBgn0261509 | haf | -0.081917399 | 0.52195992 | 0 |
| LD38004 | FBgn0035388 | CG2162 | -0.081907747 | 0.49704094 | 0 |
| RE50350 | FBgn0037573 | CG7483 | 0.081904365 | 0.52533004 | 0 |
| RH49821 | FBgn0034856 | yellow-d2 | -0.08189637 | 0.68732773 | 0 |
| LP12095 | FBgn0038973 | Pebp1 | -0.081804711 | 0.56235406 | 0 |
| LP02978 | FBgn0036952 | CG6933 | 0.081787967 | 0.69331385 | 0 |
| RE57643 | FBgn0039507 | mrt | 0.081760728 | 0.42701482 | 0 |
| AT30249 | FBgn0001217 | Hsc70-2 | -0.081756672 | 0.55622948 | 0 |
| GM14585 | FBgn0033912 | RpS23 | -0.081735947 | 0.43189074 | 0 |
| LD46853 | FBgn0004838 | Hrb27C | 0.081722366 | 0.49911647 | 0 |
| RE03226 | FBgn0031117 | GstT3 | 0.081667431 | 0.42195731 | 0 |
| RE43922 | FBgn0034350 | CG5189 | 0.081657883 | 0.42951104 | 0 |
| RE43922 | FBgn0050120 | CG30120 | 0.081657883 | 0.42951104 | 0 |
| RE28276 | FBgn0050158 | CG30158 | -0.081653142 | 0.52493274 | 0 |
| LD40460 | FBgn0005278 | Sam-S | 0.081589195 | 0.48445105 | 0 |
| LD29964 | FBgn0025865 | Cortactin | 0.081565397 | 0.43154461 | 0 |
| RE44302 | FBgn0013347 | TfIIA-S | 0.081428643 | 0.52133054 | 0 |
| LD40063 | FBgn0032895 | twit | -0.081415071 | 0.42715664 | 0 |
| SD08664 | FBgn0028675 | Sur | -0.081396426 | 0.45904015 | 0 |
| GM26647 | FBgn0010412 | RpS19a | 0.081395494 | 0.57950445 | 0 |
| RE30687 | FBgn0034945 | CG10904 | 0.081385965 | 0.51549011 | 0 |
| LD32747 | FBgn0031610 | CG15436 | 0.081322022 | 0.47168974 | 0 |
| RE14730 | FBgn0036889 | CG14100 | 0.081296555 | 0.45317518 | 0 |
| LD22815 | FBgn0262026 | CG42837 | 0.081278029 | 0.60251021 | 0 |
| LD22815 | FBgn0265178 | CG44243 | 0.081278029 | 0.60251021 | 0 |
| GM24986 | FBgn0030545 | CG11590 | 0.081274426 | 0.43000457 | 0 |
| LD44321 | FBgn0052264 | CG32264 | -0.081165015 | 0.63267966 | 0 |
| GH26007 | FBgn0035440 | CG14969 | -0.081101583 | 0.44633941 | 0 |
| RH65683 | FBgn0032139 | CG13116 | 0.081080993 | 0.64097638 | 0 |
| RE36623 | FBgn0267398 | Yeti | -0.081030922 | 0.49432901 | 0 |
| GH01760 | FBgn0016691 | ATPsynO | 0.080993712 | 0.5354438 | 0 |
| RE43371 | FBgn0040393 | CG14265 | -0.080991508 | 0.53659879 | 0 |
| LD02194 | FBgn0026378 | Rep | 0.080906308 | 0.45297056 | 0 |
| LD42637 | FBgn0030122 | Aladin | 0.080888429 | 0.5158442 | 0 |
| LD46661 | FBgn0010303 | hep | -0.080870903 | 0.51477535 | 0 |
| RH38923 | FBgn0033465 | Etf-QO | 0.080817069 | 0.60358513 | 0 |
| RE02096 | FBgn0036781 | CG13699 | 0.080788816 | 0.49704094 | 0 |
| LD36576 | FBgn0250753 | kra | -0.080694604 | 0.44459953 | 0 |
| RE65748 | FBgn0034433 | EndoB | 0.080687311 | 0.47277506 | 0 |
| GH10614 | FBgn0027552 | CG10863 | -0.080549648 | 0.49014766 | 0 |
| SD27034 | FBgn0034694 | CG6613 | -0.080513328 | 0.42276183 | 0 |

| | | | | | |
|---|---|---|---|---|---|
| AT26735 | FBgn0036895 | ms(3)76Cc | 0.08050426 | 0.51458349 | 0 |
| GH11730 | FBgn0037169 | CG11404 | -0.080484761 | 0.54219341 | 0 |
| SD02734 | FBgn0050493 | CG30493 | -0.080461328 | 0.63075595 | 0 |
| RE54994 | FBgn0030189 | CG2909 | -0.080437579 | 0.57075432 | 0 |
| RE12101 | FBgn0024273 | WASp | 0.080428965 | 0.4306595 | 0 |
| RE26825 | FBgn0036538 | CG15715 | -0.080393833 | 0.4550091 | 0 |
| GM03914 | FBgn0262733 | Src64B | 0.080365245 | 0.53021623 | 0 |
| LD28489 | FBgn0051793 | CG31793 | -0.080241725 | 0.43176226 | 0 |
| LD24240 | FBgn0030114 | CG17754 | -0.080147096 | 0.43501544 | 0 |
| RE22146 | FBgn0037543 | CG10903 | 0.080122913 | 0.50642541 | 0 |
| RE63284 | FBgn0036537 | CG18081 | -0.080078355 | 0.42692336 | 0 |
| LD28966 | FBgn0263998 | Ack-like | -0.08006879 | 0.47414835 | 0 |
| LD46935 | FBgn0037182 | ArfGAP3 | 0.080044974 | 0.68844193 | 0 |
| SD20015 | FBgn0067622 | LSm-4 | -0.08001932 | 0.50628892 | 0 |
| SD20015 | FBgn0032240 | CG17768 | -0.08001932 | 0.50628892 | 0 |
| RE38659 | FBgn0000559 | eEF2 | 0.079999118 | 0.53032948 | 0 |
| LD03419 | FBgn0014020 | Rho1 | 0.079966099 | 0.57029141 | 0 |
| LD10758 | FBgn0259824 | Hip14 | 0.079937239 | 0.51326969 | 0 |
| LD27322 | FBgn0038672 | CG6005 | 0.0799306 | 0.45026411 | 0 |
| LD18955 | FBgn0011742 | Arp2 | -0.07984026 | 0.50357769 | 0 |
| RE64759 | FBgn0031619 | CG3355 | -0.079752072 | 0.55130538 | 0 |
| LP11175 | FBgn0004590 | Eig71Ec | 0.079699216 | 0.55986286 | 0 |
| LD39612 | FBgn0030768 | CG9723 | 0.079691946 | 0.47573073 | 0 |
| RE40776 | FBgn0037614 | TMEM216 | 0.07968998 | 0.43232503 | 0 |
| RE55125 | FBgn0039930 | CG11077 | 0.079687651 | 0.46030056 | 0 |
| RE56892 | FBgn0050418 | nord | 0.079643609 | 0.42827003 | 0 |
| GM10264 | FBgn0261995 | CG42813 | -0.079600272 | 0.50130965 | 0 |
| LP05346 | FBgn0033541 | CG12934 | -0.079585933 | 0.68400985 | 0 |
| RE35182 | FBgn0037409 | Osi24 | -0.079574238 | 0.49386315 | 0 |
| LD13864 | FBgn0033029 | Not3 | -0.079505023 | 0.49596037 | 0 |
| RE03551 | FBgn0038118 | timeout | 0.079488378 | 0.48204887 | 0 |
| SD03629 | FBgn0037391 | CG2017 | 0.079468517 | 0.46236205 | 0 |
| RE09176 | FBgn0026088 | CG14818 | 0.079423006 | 0.4344621 | 0 |
| HL08157 | FBgn0030078 | CG10970 | 0.079270467 | 0.65299747 | 0 |
| HL08157 | FBgn0265413 | CG44325 | 0.079270467 | 0.65299747 | 0 |
| RH12281 | FBgn0053110 | CG33110 | -0.079241206 | 0.72196449 | 0 |
| LD47639 | FBgn0034067 | CG8399 | -0.079239607 | 0.58717508 | 0 |
| GM21055 | FBgn0035166 | JMJD5 | 0.079221537 | 0.4344621 | 0 |
| GH11818 | FBgn0034887 | St1 | 0.079183554 | 0.46879983 | 0 |
| SD28142 | FBgn0036372 | Abp1 | 0.079159476 | 0.4344621 | 0 |
| GH02855 | FBgn0039280 | Mocs2 | -0.079069806 | 0.55700655 | 0 |
| GH02855 | FBgn0260229 | CG42503 | -0.079069806 | 0.55700655 | 0 |
| LD24631 | FBgn0050122 | CG30122 | 0.079015546 | 0.4655033 | 0 |
| LD43873 | FBgn0259243 | Pka-R1 | 0.078971183 | 0.44332218 | 0 |
| SD16956 | FBgn0038861 | CG7009 | -0.078950324 | 0.82837799 | 0 |
| GH26796 | FBgn0038037 | Cyp9f2 | -0.078950132 | 0.75378493 | 0 |
| RE42878 | FBgn0266410 | CG45050 | -0.078881505 | 0.50684673 | 0 |
| LD13901 | FBgn0266487 | CG45086 | -0.078828746 | 0.56766945 | 0 |

| | | | | | |
|---|---|---|---|---|---|
| LP10313 | FBgn0041092 | tai | -0.078791171 | 0.67876001 | 0 |
| LD33876 | FBgn0032222 | CG5037 | 0.078777267 | 0.43070215 | 0 |
| RE23541 | FBgn0031395 | CG10874 | 0.078776115 | 0.44489043 | 0 |
| AT22101 | FBgn0038709 | CG15025 | 0.078707252 | 0.43087418 | 0 |
| LD03769 | FBgn0032243 | Klp31E | 0.078687778 | 0.7105132 | 0 |
| HL01728 | FBgn0035989 | CG3967 | -0.07867278 | 0.52605755 | 0 |
| LD21134 | FBgn0031738 | CG9171 | -0.078667424 | 0.4949169 | 0 |
| SD08447 | FBgn0029957 | CG12155 | -0.07859762 | 0.43189074 | 0 |
| LD30602 | FBgn0259735 | CG42389 | -0.078582105 | 0.60237747 | 0 |
| RE60792 | FBgn0027101 | Dyrk3 | -0.078538405 | 0.57219448 | 0 |
| RE40293 | FBgn0036929 | CG7668 | -0.078537027 | 0.48574254 | 0 |
| LD12394 | FBgn0004889 | tws | -0.078527762 | 0.44252208 | 0 |
| RE41492 | FBgn0085360 | CG34331 | 0.078509089 | 0.50628892 | 0 |
| GM23677 | FBgn0037989 | ATP8B | -0.078473261 | 0.8266986 | 0 |
| GH10201 | FBgn0030148 | CG3106 | -0.07845589 | 0.51140355 | 0 |
| GH11067 | FBgn0037842 | CG6567 | 0.078381019 | 0.57713962 | 0 |
| GH07373 | FBgn0243486 | rdo | 0.078369201 | 0.53882295 | 0 |
| SD05384 | FBgn0034113 | CG8060 | -0.078355915 | 0.5700143 | 0 |
| AT26975 | FBgn0035970 | CG4483 | -0.078332187 | 0.62004082 | 0 |
| GH19789 | FBgn0037988 | CG14740 | -0.078313458 | 0.64116333 | 0 |
| LD43650 | FBgn0027786 | Mtch | -0.078288368 | 0.46464953 | 0 |
| RE71995 | FBgn0037416 | Osi9 | -0.078260866 | 0.7346144 | 0 |
| RE10861 | FBgn0036377 | CG10710 | -0.078248819 | 0.55108285 | 0 |
| RH03850 | FBgn0034471 | Obp56e | 0.078241045 | 0.50842938 | 0 |
| AT23078 | FBgn0034033 | CG8204 | 0.078219939 | 0.45243277 | 0 |
| RH21402 | FBgn0001218 | Hsc70-3 | -0.078210339 | 0.43651754 | 0 |
| RE25373 | FBgn0260943 | Rbp6 | -0.078179588 | 0.53998527 | 0 |
| GM02565 | FBgn0050377 | CG30377 | -0.078127447 | 0.43286313 | 0 |
| RH67628 | FBgn0034622 | CG15666 | 0.078109839 | 0.5826711 | 0 |
| RE06985 | FBgn0025631 | moody | 0.078064216 | 0.45494449 | 0 |
| GM12295 | FBgn0029915 | CG14434 | -0.078063593 | 0.53059171 | 0 |
| LD16387 | FBgn0039067 | wda | -0.078048189 | 0.45877745 | 0 |
| LD29556 | FBgn0039207 | CG5789 | -0.07796677 | 0.47768899 | 0 |
| SD21394 | FBgn0036814 | CG14073 | 0.077951632 | 0.59384083 | 0 |
| LD04613 | FBgn0002567 | Rab32 | 0.077943648 | 0.45051693 | 0 |
| GH26994 | FBgn0034402 | CG15084 | -0.077938362 | 0.48522619 | 0 |
| LD33778 | FBgn0038551 | Odj | 0.077924567 | 0.49293909 | 0 |
| GH14758 | FBgn0035695 | CG10226 | 0.077915097 | 0.51915168 | 0 |
| GH12023 | FBgn0037439 | CG10286 | -0.077892771 | 0.51340267 | 0 |
| RE10324 | FBgn0035428 | CG14960 | -0.077886237 | 0.5946724 | 0 |
| LD03002 | FBgn0261789 | SmD2 | 0.0778694 | 0.56666839 | 0 |
| RE16596 | FBgn0015025 | CkIIalpha-i1 | -0.077850227 | 0.61927718 | 0 |
| LD19058 | FBgn0024921 | Tnpo | 0.077829095 | 0.50711874 | 0 |
| GH04877 | FBgn0087005 | rtp | 0.077823776 | 0.4814305 | 0 |
| LD29015 | FBgn0033000 | CG14464 | -0.077784493 | 0.52055144 | 0 |
| GM01209 | FBgn0036710 | CG6479 | 0.077768822 | 0.44227735 | 0 |
| GH13165 | FBgn0034957 | CG3121 | -0.077746296 | 0.53937384 | 0 |
| GH21622 | FBgn0036687 | CG6652 | -0.077712236 | 0.45923276 | 0 |

| | | | | | |
|---|---|---|---|---|---|
| GH06180 | FBgn0261609 | eIF2alpha | -0.077676206 | 0.52533004 | 0 |
| LP02252 | FBgn0035199 | tfc | -0.07765545 | 0.49201375 | 0 |
| RH42017 | FBgn0025625 | Sik2 | -0.077640888 | 0.54104339 | 0 |
| LD31163 | FBgn0036248 | ssp | 0.077588357 | 0.47978982 | 0 |
| RE09212 | FBgn0035624 | Eaf6 | 0.077575841 | 0.52386816 | 0 |
| RE31380 | FBgn0035094 | CG9380 | 0.077517252 | 0.47885962 | 0 |
| GH13992 | FBgn0086656 | shrb | -0.077504578 | 0.48048527 | 0 |
| RE41391 | FBgn0031776 | Pfdn1 | -0.077496571 | 0.45593476 | 0 |
| RE26079 | FBgn0011762 | DNApol-alpha50 | -0.077478764 | 0.55754859 | 0 |
| HL08032 | FBgn0285962 | CG46338 | -0.077446886 | 0.64968823 | 0 |
| GH27987 | FBgn0032577 | CG13244 | 0.077437362 | 0.49043505 | 0 |
| LD22033 | FBgn0027513 | ana2 | -0.077384449 | 0.51930145 | 0 |
| GH02584 | FBgn0029526 | CR18166 | 0.077342958 | 0.47276184 | 0 |
| GH08474 | FBgn0039635 | CG11876 | -0.077315804 | 0.6471999 | 0 |
| AT08673 | FBgn0284250 | Oaz | 0.077298802 | 0.64988699 | 0 |
| LD10305 | FBgn0051522 | CG31522 | 0.077218619 | 0.49704094 | 0 |
| GH03511 | FBgn0031191 | Cp110 | -0.077130958 | 0.52148082 | 0 |
| LD47477 | FBgn0033740 | dgt5 | 0.077103661 | 0.54085803 | 0 |
| AT05149 | FBgn0034327 | CG14505 | 0.077072806 | 0.50343313 | 0 |
| GH15471 | FBgn0026059 | Mhcl | 0.07706864 | 0.47414835 | 0 |
| SD01621 | FBgn0039016 | Dcr-1 | -0.077039161 | 0.49182323 | 0 |
| LD30246 | FBgn0031018 | CG12200 | -0.077035077 | 0.52724918 | 0 |
| LP05416 | FBgn0028534 | CG7916 | 0.076977168 | 0.67100423 | 0 |
| GH24265 | FBgn0015036 | Cyp4ae1 | -0.076904796 | 0.47616901 | 0 |
| RE14386 | FBgn0038613 | Vha100-4 | -0.07689515 | 0.52406246 | 0 |
| RE22911 | FBgn0037669 | Ibf2 | 0.076866102 | 0.50655576 | 0 |
| LD44821 | FBgn0262598 | mtsh | -0.076863219 | 0.58180172 | 0 |
| RH26422 | FBgn0085359 | CG34330 | -0.076836376 | 0.4688245 | 0 |
| LD14031 | FBgn0039923 | MED26 | 0.076836256 | 0.63855331 | 0 |
| GH11671 | FBgn0004580 | Cbp53E | 0.076834562 | 0.55996416 | 0 |
| LD45843 | FBgn0036837 | CG18135 | 0.076807221 | 0.51915168 | 0 |
| LD45115 | FBgn0030801 | Rcp | -0.076792678 | 0.4625759 | 0 |
| GH09983 | FBgn0039802 | dj-1beta | -0.076777282 | 0.46265346 | 0 |
| RE52922 | FBgn0040928 | CR15345 | 0.076757869 | 0.62327293 | 0 |
| LD05920 | FBgn0030966 | CG7280 | 0.07673719 | 0.44577837 | 0 |
| LD35051 | FBgn0051637 | CG31637 | -0.076733788 | 0.49324251 | 0 |
| GH13793 | FBgn0261276 | Opa1 | 0.076717163 | 0.44489043 | 0 |
| LP05376 | FBgn0086611 | Lcp65Ag3 | 0.076712365 | 0.44794024 | 0 |
| GH14517 | FBgn0034720 | Liprin-gamma | -0.076674721 | 0.61374063 | 0 |
| GH25986 | FBgn0285943 | knon | 0.076672485 | 0.50652933 | 0 |
| LD40792 | FBgn0011640 | lark | 0.076454189 | 0.47552935 | 0 |
| LP06872 | FBgn0026565 | Ass | -0.076417669 | 0.64988699 | 0 |
| RE73235 | FBgn0267376 | SelR | 0.076379321 | 0.49324251 | 0 |
| GH08526 | FBgn0037202 | Ssl1 | 0.076376936 | 0.59173688 | 0 |
| RE73432 | FBgn0040383 | CG5254 | 0.076293541 | 0.61840137 | 0 |
| RE18653 | FBgn0039757 | RpS7 | 0.076242848 | 0.6064626 | 0 |
| AT24650 | FBgn0047334 | BG642312 | 0.076110183 | 0.5620425 | 0 |
| SD17376 | FBgn0086782 | amn | -0.076061309 | 0.59496777 | 0 |

| | | | | | |
|---|---|---|---|---|---|
| RH40749 | FBgn0039480 | Cpr97Ea | -0.076004516 | 0.49647104 | 0 |
| RE32027 | FBgn0030230 | Rph | -0.075891335 | 0.57941879 | 0 |
| RE24382 | FBgn0013751 | Awh | -0.075874056 | 0.88328086 | 0 |
| RH56103 | FBgn0036488 | CG6878 | 0.075854074 | 0.46125218 | 0 |
| LD04004 | FBgn0063493 | GstE7 | 0.075833479 | 0.56127049 | 0 |
| GM03174 | FBgn0037121 | Rpb8 | 0.075826515 | 0.71610168 | 0 |
| LD46465 | FBgn0034814 | CG9890 | 0.075783195 | 0.45026411 | 0 |
| GH15825 | FBgn0034816 | CG3085 | 0.075719422 | 0.5123227 | 0 |
| LD33386 | FBgn0033233 | Kdm4A | 0.075675565 | 0.45351921 | 0 |
| RE11283 | FBgn0039805 | Cpr100A | -0.075602499 | 0.58996985 | 0 |
| RE47284 | FBgn0024995 | CG2680 | 0.075557132 | 0.4929071 | 0 |
| GH10478 | FBgn0030183 | CG15309 | -0.075518624 | 0.47828857 | 0 |
| GH12636 | FBgn0019643 | AANAT1 | -0.075474648 | 0.58659051 | 0 |
| LD30873 | FBgn0086676 | spin | 0.07547456 | 0.4929185 | 0 |
| RE15779 | FBgn0016684 | NaPi-T | -0.075454426 | 0.52148082 | 0 |
| LP06937 | FBgn0038638 | CG7702 | -0.075317948 | 0.70340304 | 0 |
| GH02384 | FBgn0042175 | CG18858 | 0.075299477 | 0.51682826 | 0 |
| GH02384 | FBgn0051683 | CG31683 | 0.075299477 | 0.51682826 | 0 |
| LD23619 | FBgn0039226 | Ude | 0.075244647 | 0.67563785 | 0 |
| RE74901 | FBgn0031036 | CG14220 | 0.075236712 | 0.45593254 | 0 |
| SD02907 | FBgn0032339 | Wdr59 | 0.075234561 | 0.51876735 | 0 |
| GH06635 | FBgn0029861 | CG3815 | 0.075099268 | 0.4577751 | 0 |
| LD22862 | FBgn0012051 | CalpA | 0.075032569 | 0.84292044 | 0 |
| RE38869 | FBgn0032382 | Mal-B2 | 0.074995898 | 0.55944204 | 0 |
| LD47230 | FBgn0034661 | CG4386 | 0.074990197 | 0.62511938 | 0 |
| LD35507 | FBgn0032258 | CG7456 | 0.074982264 | 0.45616529 | 0 |
| RH02365 | FBgn0035470 | Ccz1 | 0.074929586 | 0.49357117 | 0 |
| AT14658 | FBgn0029725 | CG2871 | -0.074908801 | 0.46568684 | 0 |
| RH62928 | FBgn0046999 | CG6429 | 0.074877276 | 0.67405248 | 0 |
| RE07278 | FBgn0052204 | CG32204 | -0.074847291 | 0.65941293 | 0 |
| LD33348 | FBgn0026207 | mbo | 0.074818082 | 0.52724918 | 0 |
| LD04718 | FBgn0034417 | CG15117 | -0.07477234 | 0.56886935 | 0 |
| RE27952 | FBgn0005626 | ple | 0.074770624 | 0.5551644 | 0 |
| GH24669 | FBgn0039006 | Cyp6d4 | 0.074757842 | 0.70130204 | 0 |
| SD04292 | FBgn0033460 | Sec24AB | -0.07475141 | 0.45255085 | 0 |
| RE18450 | FBgn0002021 | l(2)37Bb | 0.07464439 | 0.55653115 | 0 |
| AT11165 | FBgn0031485 | CG9643 | 0.07462987 | 0.46483688 | 0 |
| SD09295 | FBgn0037831 | Cap-H2 | 0.074604082 | 0.5130467 | 0 |
| GM06363 | FBgn0263745 | CR43670 | -0.074588275 | 0.60209852 | 0 |
| LD44887 | FBgn0002901 | mus304 | 0.074583909 | 0.45961469 | 0 |
| SD04054 | FBgn0004374 | neb | -0.074542457 | 0.51140355 | 0 |
| GH06546 | FBgn0030521 | CtsB1 | 0.074534219 | 0.48928658 | 0 |
| RE12122 | FBgn0022943 | Cbp20 | 0.074521786 | 0.47918187 | 0 |
| GH18645 | FBgn0003710 | tipE | -0.074518043 | 0.59897104 | 0 |
| RH58567 | FBgn0017414 | cag | -0.07443401 | 0.67290219 | 0 |
| RH25447 | FBgn0030976 | CG7378 | -0.074407061 | 0.63267966 | 0 |
| LD22910 | FBgn0028476 | Usp1 | 0.07438655 | 0.45433285 | 0 |
| RH50933 | FBgn0050503 | CG30503 | 0.074368002 | 0.60269094 | 0 |

| | | | | | |
|---|---|---|---|---|---|
| RE56970 | FBgn0052103 | CG32103 | -0.074365422 | 0.45917583 | 0 |
| AT08566 | FBgn0037254 | CG14651 | 0.074355781 | 0.62452508 | 0 |
| AT08566 | FBgn0265373 | CG44314 | 0.074355781 | 0.62452508 | 0 |
| RE28421 | FBgn0033899 | CG13016 | 0.074291558 | 0.56237351 | 0 |
| GH08340 | FBgn0032689 | CG10413 | 0.07427682 | 0.51163144 | 0 |
| LP01188 | FBgn0035238 | CG12104 | 0.074267355 | 0.57713962 | 0 |
| RE63672 | FBgn0033755 | ClC-b | 0.074192773 | 0.5130467 | 0 |
| LD03829 | FBgn0025777 | homer | 0.074191832 | 0.53326481 | 0 |
| LD18692 | FBgn0034354 | GstE11 | 0.074175841 | 0.48842433 | 0 |
| HL04319 | FBgn0038223 | Afti | -0.074089177 | 0.53019604 | 0 |
| AT07420 | FBgn0034242 | CG14480 | 0.07408024 | 0.47240791 | 0 |
| AT07420 | FBgn0263029 | CG43324 | 0.07408024 | 0.47240791 | 0 |
| LD42756 | FBgn0030174 | CG15312 | 0.07400041 | 0.47068631 | 0 |
| GH13461 | FBgn0003748 | Treh | -0.073969314 | 0.50628892 | 0 |
| LP09144 | FBgn0004855 | RpII15 | 0.073908891 | 0.60237747 | 0 |
| RE13747 | FBgn0031601 | Dim1 | -0.07386017 | 0.70697617 | 0 |
| GH14779 | FBgn0038972 | CG7054 | 0.073847384 | 0.56528149 | 0 |
| GM14553 | FBgn0001612 | Grip91 | -0.073837851 | 0.53009356 | 0 |
| RE45701 | FBgn0030701 | CG16952 | -0.073827283 | 0.46222895 | 0 |
| LD24696 | FBgn0033101 | CG9436 | 0.073824411 | 0.51901403 | 0 |
| LD45324 | FBgn0038570 | Prx5 | 0.073773287 | 0.48669963 | 0 |
| LD45324 | FBgn0038571 | CG7215 | 0.073773287 | 0.48669963 | 0 |
| RE07958 | FBgn0035064 | TyrRS-m | -0.073757991 | 0.51337412 | 0 |
| RE60835 | FBgn0028919 | CG16865 | 0.073750141 | 0.49093682 | 0 |
| LD44443 | FBgn0000404 | CycA | -0.073653659 | 0.47828857 | 0 |
| GH01066 | FBgn0010222 | Nmdmc | -0.07364894 | 0.52555804 | 0 |
| LD20229 | FBgn0016034 | mael | 0.07363248 | 0.5130467 | 0 |
| GH26622 | FBgn0013305 | Nmda1 | -0.073516071 | 0.4867474 | 0 |
| AT03090 | FBgn0034546 | CG13442 | -0.07350837 | 0.5870756 | 0 |
| RE51744 | FBgn0035755 | CG14830 | 0.073445116 | 0.67579802 | 0 |
| RE18068 | FBgn0037856 | Leash | 0.073420365 | 0.46551707 | 0 |
| LD31082 | FBgn0040206 | krz | 0.07341823 | 0.45877745 | 0 |
| SD07261 | FBgn0003507 | srp | -0.073402495 | 0.65683532 | 0 |
| LD23587 | FBgn0028475 | Hrd3 | 0.073394925 | 0.51317355 | 0 |
| LP07781 | FBgn0038915 | CG17819 | 0.073314607 | 0.74392699 | 0 |
| AT13310 | FBgn0250817 | lncRNA:CR40465 | -0.07330283 | 0.63075595 | 0 |
| RH08962 | FBgn0038834 | RpS30 | -0.073298512 | 0.5946724 | 0 |
| GH21672 | FBgn0037734 | trbd | 0.07327322 | 0.46457207 | 0 |
| AT09752 | FBgn0052064 | S-Lap4 | 0.073271864 | 0.75215614 | 0 |
| AT09752 | FBgn0038567 | CG14316 | 0.073271864 | 0.75215614 | 0 |
| RE41906 | FBgn0020912 | Ptx1 | -0.073262658 | 0.75785286 | 0 |
| LD32155 | FBgn0029944 | Dok | 0.073205211 | 0.54470443 | 0 |
| SD07852 | FBgn0011708 | Syx5 | 0.07317434 | 0.50815974 | 0 |
| GM14307 | FBgn0032871 | CG2611 | -0.073172117 | 0.58929993 | 0 |
| LD32755 | FBgn0010097 | gammaTub37C | 0.07315897 | 0.47918187 | 0 |
| GH16485 | FBgn0035170 | dpr20 | -0.07309675 | 0.63927327 | 0 |
| LD10416 | FBgn0264744 | CG44002 | 0.073087839 | 0.52021846 | 0 |
| RH09956 | FBgn0031563 | CG10031 | 0.073016028 | 0.80407614 | 0 |

| | | | | | |
|---|---|---|---|---|---|
| GH27795 | FBgn0265045 | Strn-Mlck | -0.072956651 | 0.50947991 | 0 |
| AT28768 | FBgn0030319 | CG2533 | 0.072955442 | 0.56237351 | 0 |
| LD45607 | FBgn0032166 | CG4619 | 0.072931229 | 0.47080905 | 0 |
| LD32539 | FBgn0262656 | Myc | 0.072909761 | 0.61403286 | 0 |
| LD09158 | FBgn0262515 | VhaAC45 | -0.072860639 | 0.53326482 | 0 |
| RE24507 | FBgn0035547 | CG15022 | -0.072792964 | 0.57237973 | 0 |
| RE45347 | FBgn0039596 | CG10000 | 0.072730461 | 0.49138671 | 0 |
| RH05388 | FBgn0037144 | CG7458 | 0.072637184 | 0.61060702 | 0 |
| LD10322 | FBgn0038721 | subdued | 0.072613184 | 0.47068547 | 0 |
| SD10395 | FBgn0029507 | Tsp42Ed | 0.072550517 | 0.50766723 | 0 |
| GH10711 | FBgn0027550 | CG6495 | -0.072543187 | 0.70541114 | 0 |
| RE59088 | FBgn0032235 | CG5096 | -0.072457769 | 0.7089224 | 0 |
| GM13037 | FBgn0037901 | CG6744 | -0.072420027 | 0.51607831 | 0 |
| GH13760 | FBgn0032250 | holn1 | 0.072385249 | 0.59127991 | 0 |
| GH14368 | FBgn0000543 | ecd | 0.072295323 | 0.49704094 | 0 |
| RE14402 | FBgn0029885 | CG3224 | -0.072251682 | 0.54985807 | 0 |
| RE40745 | FBgn0024732 | Drep1 | -0.072229514 | 0.72174238 | 0 |
| LD36057 | FBgn0004878 | cas | -0.072213452 | 0.60358513 | 0 |
| LD26837 | FBgn0035631 | Txl | 0.072127622 | 0.52533004 | 0 |
| GH23064 | FBgn0030524 | CG10993 | 0.072099262 | 0.59945145 | 0 |
| AT13713 | FBgn0039083 | CG10177 | 0.07209347 | 0.70285413 | 0 |
| AT14762 | FBgn0051418 | CG31418 | -0.072085123 | 0.50766723 | 0 |
| AT15089 | FBgn0003900 | twi | -0.072022248 | 0.74203925 | 0 |
| LD35711 | FBgn0262716 | Arp3 | 0.072020442 | 0.48038085 | 0 |
| RE50332 | FBgn0035285 | CG12025 | -0.07201485 | 0.5123227 | 0 |
| GM08958 | FBgn0029870 | Marf | -0.072001572 | 0.53322848 | 0 |
| LD19028 | FBgn0086683 | Spf45 | 0.071957006 | 0.51270727 | 0 |
| LD21120 | FBgn0027784 | Prp18 | -0.071948702 | 0.47359183 | 0 |
| LD03247 | FBgn0032884 | Pomp | 0.071843249 | 0.60029887 | 0 |
| AT30415 | FBgn0030386 | CG2574 | 0.071820576 | 0.58394695 | 0 |
| RE21276 | FBgn0031885 | Mnn1 | -0.071789858 | 0.50628892 | 0 |
| AT23073 | FBgn0261683 | B9d2 | 0.071786677 | 0.50680256 | 0 |
| AT23073 | FBgn0261697 | tctn | 0.071786677 | 0.50680256 | 0 |
| GH10361 | FBgn0041191 | Rheb | -0.071740874 | 0.51185589 | 0 |
| GH16479 | FBgn0025724 | beta'COP | -0.071726083 | 0.5274825 | 0 |
| LD20793 | FBgn0022770 | Peritrophin-A | -0.071615092 | 0.64065743 | 0 |
| GH11358 | FBgn0034705 | CG11170 | -0.071557098 | 0.73818379 | 0 |
| LD16456 | FBgn0016685 | Nlp | -0.071466146 | 0.53608002 | 0 |
| SD22712 | FBgn0038974 | CG5377 | -0.071450706 | 0.4929071 | 0 |
| LD29969 | FBgn0042127 | CG18789 | -0.071370201 | 0.48529408 | 0 |
| LD29969 | FBgn0042125 | CG18787 | -0.071370201 | 0.48529408 | 0 |
| RE74511 | FBgn0000100 | RpLP0 | 0.071284637 | 0.50766723 | 0 |
| RE05389 | FBgn0038470 | CG18213 | -0.071215758 | 0.70015384 | 0 |
| RE35406 | FBgn0002566 | lt | -0.071167047 | 0.71253998 | 0 |
| SD09502 | FBgn0036463 | Reck | 0.071160755 | 0.51337412 | 0 |
| GH11294 | FBgn0033917 | SmydA-1 | -0.071157088 | 0.52533004 | 0 |
| LD42896 | FBgn0261278 | grp | -0.071125752 | 0.53045256 | 0 |
| SD16010 | FBgn0034647 | pirk | -0.071124034 | 0.70788092 | 0 |

| | | | | | |
|---|---|---|---|---|---|
| RE17954 | FBgn0002521 | pho | -0.071109163 | 0.6381511 | 0 |
| GM24338 | FBgn0052266 | CG32266 | -0.071102429 | 0.54486855 | 0 |
| AT13773 | FBgn0037879 | scpr-C | 0.071068776 | 0.60452986 | 0 |
| AT13773 | FBgn0037889 | scpr-A | 0.071068776 | 0.60452986 | 0 |
| AT13773 | FBgn0037888 | scpr-B | 0.071068776 | 0.60452986 | 0 |
| RE07473 | FBgn0034010 | CG8157 | 0.071025892 | 0.61620649 | 0 |
| GH24360 | FBgn0033900 | CysRS-m | 0.070966656 | 0.56419565 | 0 |
| GH13245 | FBgn0034518 | CG13430 | 0.070963376 | 0.5405103 | 0 |
| GH13245 | FBgn0034519 | CG18065 | 0.070963376 | 0.5405103 | 0 |
| LD47668 | FBgn0037443 | Dmtn | 0.070911524 | 0.52087078 | 0 |
| LD10434 | FBgn0030878 | CG6769 | 0.070887163 | 0.61196621 | 0 |
| GM06493 | FBgn0037880 | CG17726 | 0.07086779 | 0.49138671 | 0 |
| LP01770 | FBgn0000157 | Dll | 0.070757401 | 0.62384379 | 0 |
| SD04409 | FBgn0040336 | Seipin | -0.070743925 | 0.53045256 | 0 |
| LD10543 | FBgn0035267 | CG13921 | 0.070739274 | 0.48669963 | 0 |
| RE13669 | FBgn0031792 | CG13983 | -0.070706356 | 0.57992443 | 0 |
| GH08045 | FBgn0038877 | CG3308 | 0.070698594 | 0.58586946 | 0 |
| SD03002 | FBgn0029685 | CG2938 | 0.070683582 | 0.49845238 | 0 |
| LD44171 | FBgn0002872 | mu2 | 0.070656016 | 0.48771665 | 0 |
| GH17679 | FBgn0032651 | Oli | 0.07058984 | 0.8397362 | 0 |
| LD47616 | FBgn0035414 | CG14965 | 0.070567719 | 0.49456407 | 0 |
| SD11486 | FBgn0035861 | CG7213 | -0.070553281 | 0.65924877 | 0 |
| GH20621 | FBgn0032494 | CG5945 | 0.070513944 | 0.65276285 | 0 |
| LD31570 | FBgn0004957 | por | 0.070493473 | 0.50990391 | 0 |
| SD28072 | FBgn0038893 | Archease | -0.070466521 | 0.56925579 | 0 |
| SD01279 | FBgn0003139 | PpV | -0.07044798 | 0.5700488 | 0 |
| LP03575 | FBgn0052191 | CG32191 | 0.070421393 | 0.62949402 | 0 |
| GH28665 | FBgn0036043 | CG8177 | 0.070409659 | 0.48523766 | 0 |
| AT02529 | FBgn0050156 | CG30156 | -0.070398118 | 0.51140355 | 0 |
| RE74758 | FBgn0036102 | CG12523 | -0.070394976 | 0.51898428 | 0 |
| LD34461 | FBgn0036928 | Tom20 | 0.070389247 | 0.49579399 | 0 |
| LD35637 | FBgn0026403 | Ndg | -0.070345133 | 0.57813855 | 0 |
| GH16885 | FBgn0053548 | SmydA-8 | -0.070329076 | 0.53045256 | 0 |
| AT06826 | FBgn0029705 | CG12693 | 0.070320337 | 0.58989477 | 0 |
| GM08320 | FBgn0052425 | CG32425 | 0.070315872 | 0.49587261 | 0 |
| AT28101 | FBgn0031292 | CG15824 | -0.070256266 | 0.5996356 | 0 |
| LD23181 | FBgn0027506 | EDTP | 0.070253249 | 0.56984459 | 0 |
| GH10050 | FBgn0034577 | cpa | -0.070241444 | 0.52724918 | 0 |
| LP02712 | FBgn0023550 | FarO | -0.070241046 | 0.61579639 | 0 |
| LD18761 | FBgn0027868 | Nup107 | 0.070177347 | 0.56683332 | 0 |
| AT07283 | FBgn0037781 | Fancl | -0.070167489 | 0.59945145 | 0 |
| LD23630 | FBgn0037746 | CG8478 | 0.070102722 | 0.50251481 | 0 |
| RE38147 | FBgn0036551 | CG17029 | 0.07007192 | 0.59873026 | 0 |
| GH16592 | FBgn0041710 | yellow-f | -0.070039223 | 0.6153161 | 0 |
| GH26260 | FBgn0004852 | Ac76E | 0.070025602 | 0.5130467 | 0 |
| RE33604 | FBgn0000346 | comt | 0.069980189 | 0.59191735 | 0 |
| LD38433 | FBgn0039341 | CG5112 | 0.069966277 | 0.59940307 | 0 |
| SD24632 | FBgn0031894 | CG4496 | -0.069936186 | 0.49845238 | 0 |

| | | | | | |
|---|---|---|---|---|---|
| LD44273 | FBgn0032880 | TM9SF2 | 0.06992611 | 0.53287857 | 0 |
| LD36546 | FBgn0013334 | Sap47 | 0.069915033 | 0.52976763 | 0 |
| LD21637 | FBgn0037442 | godzilla | 0.06991026 | 0.50911135 | 0 |
| LP05693 | FBgn0016917 | Stat92E | 0.069894807 | 0.52598204 | 0 |
| GH23382 | FBgn0036934 | sNPF-R | -0.069882661 | 0.53326482 | 0 |
| GH27120 | FBgn0033529 | CG17765 | 0.069860631 | 0.55378047 | 0 |
| RE18093 | FBgn0005617 | msl-1 | 0.069820172 | 0.49192318 | 0 |
| SD01613 | FBgn0036702 | CG6512 | 0.069770964 | 0.5386271 | 0 |
| AT05383 | FBgn0036093 | CG14154 | -0.069733873 | 0.71610168 | 0 |
| LD31024 | FBgn0037199 | CG11137 | 0.069732688 | 0.52088072 | 0 |
| GH22841 | FBgn0002036 | CG10561 | 0.069696212 | 0.51140355 | 0 |
| RH54244 | FBgn0041713 | yellow-c | 0.069694992 | 0.62002199 | 0 |
| AT13807 | FBgn0034774 | CG13526 | 0.069644513 | 0.50013064 | 0 |
| LP12093 | FBgn0013997 | Nrx-IV | 0.069632763 | 0.72711601 | 0 |
| LD24737 | FBgn0026713 | l(1)G0007 | 0.069456948 | 0.54357267 | 0 |
| GH11867 | FBgn0051344 | CG31344 | -0.069430502 | 0.52055144 | 0 |
| GH13296 | FBgn0026060 | Mipp2 | 0.069415704 | 0.5087503 | 0 |
| RE06730 | FBgn0038837 | KaiR1D | -0.069233309 | 0.83526512 | 0 |
| GH07689 | FBgn0034132 | S-Lap8 | -0.069226306 | 0.55511879 | 0 |
| SD07915 | FBgn0033121 | Cyp6u1 | -0.069219428 | 0.49647104 | 0 |
| RE55842 | FBgn0037750 | Whamy | -0.069215792 | 0.50049226 | 0 |
| RE07060 | FBgn0031070 | CG12702 | 0.069171572 | 0.5160646 | 0 |
| RH35480 | FBgn0026372 | RpL23A | 0.06916436 | 0.71610168 | 0 |
| LP04931 | FBgn0039848 | Loxl1 | 0.069135842 | 0.66478522 | 0 |
| RE03680 | FBgn0034138 | RpS15 | -0.069112639 | 0.51424726 | 0 |
| RE63163 | FBgn0002569 | Mal-A2 | 0.069080278 | 0.61060702 | 0 |
| SD21096 | FBgn0039407 | CG14544 | -0.069066027 | 0.5405103 | 0 |
| GH02096 | FBgn0034534 | maf-S | 0.068995082 | 0.49596037 | 0 |
| RE47112 | FBgn0032192 | CG5731 | -0.068987319 | 0.55877041 | 0 |
| RE08425 | FBgn0036213 | RpL10Ab | -0.068971419 | 0.53326482 | 0 |
| LD27852 | FBgn0035147 | Gale | 0.068961673 | 0.51218464 | 0 |
| LD19544 | FBgn0032870 | CG2608 | 0.068934649 | 0.49653741 | 0 |
| RH49488 | FBgn0031882 | Rab30 | -0.0689107 | 0.57573901 | 0 |
| GH01133 | FBgn0261986 | RASSF8 | 0.068895842 | 0.52200292 | 0 |
| AT04117 | FBgn0035263 | CG12035 | -0.068890772 | 0.51185589 | 0 |
| RE25175 | FBgn0035619 | CG10592 | -0.068881186 | 0.82364941 | 0 |
| GM01362 | FBgn0038704 | CG5316 | 0.068845929 | 0.65210401 | 0 |
| LD46840 | FBgn0028833 | Dak1 | -0.068844951 | 0.49845238 | 0 |
| GH14903 | FBgn0052333 | CG32333 | -0.068814983 | 0.64428139 | 0 |
| GH25860 | FBgn0050273 | CG30273 | 0.068707911 | 0.53045256 | 0 |
| GH25860 | FBgn0050269 | CG30269 | 0.068707911 | 0.53045256 | 0 |
| GH25860 | FBgn0004888 | Scsalpha | 0.068707911 | 0.53045256 | 0 |
| RE38067 | FBgn0066114 | GlcAT-I | 0.068692129 | 0.53984651 | 0 |
| RE29678 | FBgn0037128 | CG14572 | 0.068666206 | 0.53587727 | 0 |
| RE53092 | FBgn0035359 | CG1143 | -0.068642333 | 0.55662043 | 0 |
| RE22492 | FBgn0037427 | Osi17 | -0.068635994 | 0.48790255 | 0 |
| RH10371 | FBgn0036601 | CG13063 | 0.068631806 | 0.67285881 | 0 |
| RH10371 | FBgn0036600 | CG13043 | 0.068631806 | 0.67285881 | 0 |

| | | | | | |
|---|---|---|---|---|---|
| AT18267 | FBgn0052574 | Twdlalpha | 0.068521998 | 0.75690104 | 0 |
| LD31990 | FBgn0034142 | CG8306 | -0.068515029 | 0.80152746 | 0 |
| LD47721 | FBgn0037920 | CG14710 | 0.068467981 | 0.51386113 | 0 |
| LD11744 | FBgn0030735 | CG3632 | 0.068437875 | 0.51807014 | 0 |
| LD44919 | FBgn0034186 | CG8950 | 0.068412819 | 0.59966849 | 0 |
| RE52392 | FBgn0036173 | CG7394 | -0.068400982 | 0.52641863 | 0 |
| AT31519 | FBgn0028531 | CG15286 | 0.068387263 | 0.65868587 | 0 |
| LD06837 | FBgn0028700 | RfC38 | -0.06828592 | 0.56070417 | 0 |
| GM01721 | FBgn0000052 | Pfas | 0.068275619 | 0.5702947 | 0 |
| RE09982 | FBgn0030598 | CG9503 | -0.068252603 | 0.6308489 | 0 |
| LD17730 | FBgn0030088 | Arfrp1 | 0.068242904 | 0.5695879 | 0 |
| LD47384 | FBgn0015372 | RabX1 | -0.068239267 | 0.57075432 | 0 |
| GH07001 | FBgn0028560 | sut4 | 0.068220504 | 0.8409258 | 0 |
| RE72002 | FBgn0032949 | Lamp1 | -0.068195634 | 0.63733953 | 0 |
| RE06140 | FBgn0028648 | mRpL50 | 0.068153269 | 0.54691517 | 0 |
| GH12192 | FBgn0003071 | Pfk | -0.068128055 | 0.56126345 | 0 |
| SD02391 | FBgn0029688 | lva | -0.068123481 | 0.55130538 | 0 |
| GM16372 | FBgn0036188 | CG7339 | 0.06811874 | 0.55179338 | 0 |
| GH26723 | FBgn0032796 | CG10188 | 0.06806861 | 0.52087078 | 0 |
| LD21241 | FBgn0031030 | Tao | 0.06801929 | 0.52218436 | 0 |
| RH05209 | FBgn0033308 | CG8736 | -0.067969167 | 0.58180172 | 0 |
| RE44923 | FBgn0030352 | sicily | -0.067903735 | 0.57341402 | 0 |
| GH09289 | FBgn0003721 | Tm1 | -0.067902069 | 0.60169005 | 0 |
| RE35556 | FBgn0030433 | mRpL49 | -0.067793712 | 0.49911647 | 0 |
| RE26725 | FBgn0050403 | CG30403 | 0.067756315 | 0.58228008 | 0 |
| GH19816 | FBgn0261244 | inaE | 0.06775526 | 0.52133054 | 0 |
| RE64858 | FBgn0035685 | CG13297 | -0.067739377 | 0.67001488 | 0 |
| RH31391 | FBgn0260941 | app | 0.067721895 | 0.56528149 | 0 |
| LD31138 | FBgn0036689 | CG7730 | -0.067658821 | 0.49352768 | 0 |
| RE67445 | FBgn0035082 | CG2811 | -0.067654309 | 0.50956169 | 0 |
| LP02833 | FBgn0038135 | CG8773 | 0.067639671 | 0.57941879 | 0 |
| GH19047 | FBgn0035091 | CG3829 | -0.067564187 | 0.77665449 | 0 |
| GM23249 | FBgn0035802 | Pura | -0.067480307 | 0.55103743 | 0 |
| RH34302 | FBgn0040575 | CG15922 | -0.067467706 | 0.69946427 | 0 |
| GH13884 | FBgn0032677 | CG5790 | -0.067460786 | 0.63539356 | 0 |
| GH24154 | FBgn0011656 | Mef2 | 0.067441241 | 0.49425278 | 0 |
| LD21888 | FBgn0061469 | Ube3a | 0.067425624 | 0.51131122 | 0 |
| GH25580 | FBgn0039157 | Myo95E | -0.067417768 | 0.5196847 | 0 |
| AT27111 | FBgn0036520 | CG13449 | -0.067374419 | 0.59945145 | 0 |
| LD22358 | FBgn0266580 | Gp210 | -0.06736231 | 0.53224171 | 0 |
| RE39629 | FBgn0037087 | CG7519 | 0.067361687 | 0.5354438 | 0 |
| LD21358 | FBgn0051363 | Jupiter | 0.067330802 | 0.61374063 | 0 |
| GH23259 | FBgn0036422 | CG3868 | -0.067323587 | 0.83278542 | 0 |
| RH63179 | FBgn0259728 | CG42382 | -0.0673189 | 0.70062518 | 0 |
| LD29218 | FBgn0033859 | fand | 0.067311745 | 0.54053228 | 0 |
| LP08717 | FBgn0002564 | Lsp1gamma | 0.067240335 | 0.52493274 | 0 |
| LD38760 | FBgn0034657 | LBR | 0.067201936 | 0.76663015 | 0 |
| GH14389 | FBgn0052529 | Hers | 0.067193579 | 0.59804839 | 0 |

| | | | | | |
|---|---|---|---|---|---|
| SD05230 | FBgn0264495 | gpp | 0.06717075 | 0.53139066 | 0 |
| RE70477 | FBgn0263030 | CG43325 | -0.067119898 | 0.56759299 | 0 |
| RE70477 | FBgn0263031 | CG43326 | -0.067119898 | 0.56759299 | 0 |
| GH28353 | FBgn0030892 | CG7206 | -0.067033715 | 0.54097209 | 0 |
| GH25582 | FBgn0015379 | dod | 0.067031153 | 0.59400557 | 0 |
| GH25582 | FBgn0262782 | Mdh1 | 0.067031153 | 0.59400557 | 0 |
| RH14840 | FBgn0000723 | FER | 0.067021699 | 0.71796792 | 0 |
| AT11646 | FBgn0038051 | CG17207 | -0.066898286 | 0.52363431 | 0 |
| GM20557 | FBgn0032595 | CG17996 | 0.066893188 | 0.72336385 | 0 |
| LD25378 | FBgn0002778 | mnd | 0.066889597 | 0.52195992 | 0 |
| LD26789 | FBgn0031051 | Ranbp21 | 0.066781017 | 0.59226191 | 0 |
| RE65321 | FBgn0032421 | crok | -0.066757503 | 0.56081877 | 0 |
| LP02583 | FBgn0035666 | Jon65Aii | -0.066757169 | 0.66268185 | 0 |
| LP02583 | FBgn0035667 | Jon65Ai | -0.066757169 | 0.66268185 | 0 |
| RE45276 | FBgn0032921 | Mpp6 | -0.066725331 | 0.53638065 | 0 |
| LD26422 | FBgn0035102 | CG7049 | 0.066622322 | 0.51140355 | 0 |
| GH14608 | FBgn0030103 | Obp8a | 0.066599052 | 0.65276285 | 0 |
| GH12731 | FBgn0039580 | Gfat2 | 0.066594703 | 0.63934365 | 0 |
| GH17930 | FBgn0050350 | CG30350 | -0.066567777 | 0.56251972 | 0 |
| LP01469 | FBgn0038466 | CG8907 | 0.066553635 | 0.64605999 | 0 |
| GH25251 | FBgn0031925 | Cyp4d21 | 0.066495909 | 0.53032948 | 0 |
| RE02647 | FBgn0017567 | ND-23 | -0.06647598 | 0.5422612 | 0 |
| LD41224 | FBgn0032876 | Cen | 0.066423693 | 0.54091047 | 0 |
| GH23975 | FBgn0038799 | MFS9 | 0.066405799 | 0.62949402 | 0 |
| AT27794 | FBgn0052296 | Mrtf | -0.066400364 | 0.69852926 | 0 |
| LP10386 | FBgn0025615 | Torsin | -0.066375517 | 0.69356879 | 0 |
| LD18740 | FBgn0031908 | CG5177 | 0.066371674 | 0.75595695 | 0 |
| LD10274 | FBgn0035110 | thoc7 | 0.066203866 | 0.59639617 | 0 |
| GM08106 | FBgn0037016 | CG13252 | 0.066115446 | 0.59512639 | 0 |
| RH53050 | FBgn0010770 | ppan | 0.066093848 | 0.66469202 | 0 |
| RE01809 | FBgn0031307 | MFS3 | 0.066064959 | 0.80442213 | 0 |
| RE01809 | FBgn0267281 | lncRNA:CR45717 | 0.066064959 | 0.80442213 | 0 |
| GH24554 | FBgn0085452 | CG34423 | 0.066052667 | 0.54794397 | 0 |
| GH01761 | FBgn0053191 | CG33191 | -0.066042204 | 0.70020287 | 0 |
| LD13350 | FBgn0024555 | flfl | 0.065999283 | 0.52754514 | 0 |
| GH08407 | FBgn0038607 | hmw | -0.065983353 | 0.65040006 | 0 |
| GH03375 | FBgn0033453 | CG1667 | -0.065921774 | 0.51318789 | 0 |
| RE29708 | FBgn0023549 | Mct1 | -0.065871847 | 0.54464515 | 0 |
| RE53017 | FBgn0036317 | CG10948 | -0.06575762 | 0.60162735 | 0 |
| GM12762 | FBgn0031661 | Gmd | -0.065730453 | 0.63620149 | 0 |
| RE12402 | FBgn0037592 | CG11737 | 0.065696159 | 0.61717198 | 0 |
| LD21709 | FBgn0025674 | CycK | -0.065646733 | 0.51833128 | 0 |
| LD46333 | FBgn0032516 | Ing5 | 0.065642153 | 0.50766723 | 0 |
| RE54419 | FBgn0015838 | Vang | 0.06564094 | 0.56908072 | 0 |
| RH11815 | FBgn0052220 | Csas | 0.065538634 | 0.5996356 | 0 |
| RE69013 | FBgn0039914 | mav | 0.065514803 | 0.63539356 | 0 |
| LD27389 | FBgn0275436 | PheRS-m | 0.065417756 | 0.58332023 | 0 |
| LD03322 | FBgn0086784 | stmA | 0.065388309 | 0.51424726 | 0 |

| | | | | | |
|---|---|---|---|---|---|
| RE66796 | FBgn0033667 | reb | -0.065361621 | 0.61499827 | 0 |
| AT14419 | FBgn0037147 | CG7140 | -0.065350564 | 0.66455385 | 0 |
| GH09925 | FBgn0025726 | unc-13 | 0.065321589 | 0.54495811 | 0 |
| RE70601 | FBgn0051015 | PH4alphaPV | -0.065233334 | 0.62110021 | 0 |
| RE08174 | FBgn0031232 | CG11617 | 0.065134537 | 0.63761196 | 0 |
| LD34308 | FBgn0003444 | smo | 0.065098846 | 0.57149269 | 0 |
| LP05863 | FBgn0030098 | CG12057 | -0.065065701 | 0.58270393 | 0 |
| AT07821 | FBgn0034840 | CG3124 | -0.06506533 | 0.68866504 | 0 |
| GH20375 | FBgn0029091 | Chs2 | -0.065054478 | 0.63761196 | 0 |
| RH04607 | FBgn0031544 | CG17593 | -0.064895241 | 0.54846518 | 0 |
| LD44022 | FBgn0001179 | hay | 0.064878971 | 0.51496355 | 0 |
| AT08471 | FBgn0004173 | Mst84Db | 0.064816452 | 0.71274102 | 0 |
| LD11462 | FBgn0035766 | eco | -0.064789073 | 0.54848488 | 0 |
| GH15104 | FBgn0002909 | mus312 | 0.064784296 | 0.51458349 | 0 |
| GH15104 | FBgn0259203 | CG42307 | 0.064784296 | 0.51458349 | 0 |
| GH12714 | FBgn0005777 | PpD3 | 0.064759656 | 0.5620425 | 0 |
| GH17344 | FBgn0031287 | CG4291 | -0.064750717 | 0.54145977 | 0 |
| RE61589 | FBgn0039051 | CG17109 | -0.064731327 | 0.5615961 | 0 |
| LP07267 | FBgn0034582 | Cht9 | -0.064703603 | 0.52533004 | 0 |
| LD18667 | FBgn0053213 | CG33213 | 0.064676783 | 0.51164038 | 0 |
| LD24460 | FBgn0035916 | GAPsec | 0.064612204 | 0.64810895 | 0 |
| SD03208 | FBgn0037250 | CG1074 | -0.064586803 | 0.52406246 | 0 |
| LD39455 | FBgn0039623 | CG1951 | -0.064491107 | 0.55937022 | 0 |
| LD06894 | FBgn0033401 | CG1968 | -0.064412725 | 0.56078401 | 0 |
| AT26814 | FBgn0263234 | Phae1 | 0.064407436 | 0.73668522 | 0 |
| LD24657 | FBgn0035088 | CG3776 | 0.064389617 | 0.6713123 | 0 |
| LD26737 | FBgn0030431 | CG4407 | -0.064334916 | 0.54453623 | 0 |
| GH06377 | FBgn0033807 | AQP | 0.064322449 | 0.52055144 | 0 |
| RE10515 | FBgn0031766 | CG9117 | 0.064320646 | 0.59614743 | 0 |
| GH07959 | FBgn0031970 | CG7227 | 0.064317305 | 0.86797103 | 0 |
| GH12382 | FBgn0031436 | ND-B17.2 | -0.064294128 | 0.6415074 | 0 |
| LD34544 | FBgn0038306 | Art3 | 0.064292314 | 0.53045256 | 0 |
| SD18376 | FBgn0034963 | Not11 | 0.064185655 | 0.6867439 | 0 |
| GH07444 | FBgn0035519 | CG1309 | -0.064151061 | 0.56066561 | 0 |
| SD06892 | FBgn0026086 | Adar | 0.064127382 | 0.63055916 | 0 |
| GH23641 | FBgn0052833 | CG32833 | 0.064097584 | 0.58302715 | 0 |
| GH23641 | FBgn0052834 | CG32834 | 0.064097584 | 0.58302715 | 0 |
| RE22315 | FBgn0086856 | CG11555 | -0.064082509 | 0.56644277 | 0 |
| SD13785 | FBgn0039633 | CG11873 | -0.064078886 | 0.73353773 | 0 |
| LD47413 | FBgn0052075 | CG32075 | 0.064070446 | 0.59873026 | 0 |
| LP12178 | FBgn0033362 | CG8172 | 0.064064409 | 0.71207813 | 0 |
| GM04363 | FBgn0264821 | asRNA:CR44029 | 0.064059567 | 0.59826826 | 0 |
| LD19312 | FBgn0034087 | clu | -0.064030826 | 0.56089775 | 0 |
| AT12189 | FBgn0270925 | CG4836 | 0.06399947 | 0.55630347 | 0 |
| RE29832 | FBgn0037737 | Pnn | -0.063939385 | 0.55630347 | 0 |
| AT15471 | FBgn0031410 | CG17237 | 0.063866386 | 0.66747259 | 0 |
| RH56762 | FBgn0039694 | fig | -0.063859581 | 0.66996924 | 0 |
| GH04632 | FBgn0260657 | CG42540 | -0.063857201 | 0.83365222 | 0 |

| | | | | | |
|---|---|---|---|---|---|
| RE60058 | FBgn0025833 | CG8910 | -0.063784244 | 0.52498489 | 0 |
| GH04568 | FBgn0037720 | CG8312 | -0.063730837 | 0.64282997 | 0 |
| GH14654 | FBgn0034335 | GstE1 | 0.063707787 | 0.59873026 | 0 |
| SD04652 | FBgn0033929 | Tfb1 | -0.06368404 | 0.57807986 | 0 |
| RE25803 | FBgn0036953 | CG17145 | -0.063617744 | 0.66102494 | 0 |
| RE14259 | FBgn0010520 | Bka | 0.063597713 | 0.58545935 | 0 |
| GH01592 | FBgn0034229 | CG4847 | 0.063565694 | 0.53487304 | 0 |
| GH23390 | FBgn0038924 | CG6028 | -0.063550782 | 0.54856666 | 0 |
| RH64511 | FBgn0069973 | CG40485 | -0.063532674 | 0.69303332 | 0 |
| SD22440 | FBgn0039129 | RpS19b | -0.063523912 | 0.69692368 | 0 |
| LP01932 | FBgn0032644 | CG5131 | 0.063464252 | 0.6867439 | 0 |
| RE35715 | FBgn0020235 | ATPsyngamma | 0.063460385 | 0.63516293 | 0 |
| AT20383 | FBgn0267326 | Ntl | 0.063454136 | 0.68287198 | 0 |
| GM14344 | FBgn0036212 | CG11597 | 0.063438765 | 0.56711281 | 0 |
| GH26186 | FBgn0025879 | Timp | 0.063428237 | 0.67259652 | 0 |
| LD10431 | FBgn0260960 | Baldspot | 0.063419227 | 0.60773166 | 0 |
| SD15809 | FBgn0033907 | mRpS16 | -0.063281646 | 0.52724918 | 0 |
| GH20378 | FBgn0037931 | CG18476 | -0.063225296 | 0.71774153 | 0 |
| LD44162 | FBgn0037109 | MED1 | 0.063147022 | 0.55301786 | 0 |
| GM15606 | FBgn0032252 | loh | 0.063115583 | 0.55218332 | 0 |
| GH12359 | FBgn0038809 | CG16953 | 0.063101805 | 0.52498489 | 0 |
| GH08468 | FBgn0010612 | ATPsynG | 0.063094308 | 0.59746344 | 0 |
| GH08468 | FBgn0038092 | beat-Vb | 0.063094308 | 0.59746344 | 0 |
| RE62284 | FBgn0032409 | Ced-12 | 0.06305678 | 0.55279134 | 0 |
| RE02049 | FBgn0052694 | CG32694 | -0.063034812 | 0.90665555 | 0 |
| RE30213 | FBgn0039915 | Gat | -0.062969208 | 0.75373984 | 0 |
| GH03683 | FBgn0283535 | Vha26 | -0.062960466 | 0.53539893 | 0 |
| RE25260 | FBgn0050349 | CG30349 | -0.062945461 | 0.57236982 | 0 |
| GH17672 | FBgn0037387 | CG1213 | 0.06294477 | 0.53326482 | 0 |
| GH10322 | FBgn0024944 | Oamb | -0.062842862 | 0.630677 | 0 |
| GH08860 | FBgn0027560 | Tps1 | 0.062840979 | 0.63075595 | 0 |
| SD18306 | FBgn0050010 | CG30010 | 0.062840576 | 0.55518066 | 0 |
| RE17834 | FBgn0004101 | bs | -0.062830586 | 0.61601092 | 0 |
| AT04467 | FBgn0036568 | ATPsynbetaL | -0.062821648 | 0.71631551 | 0 |
| LD29185 | FBgn0037551 | Gie | -0.062736293 | 0.67523158 | 0 |
| GH03263 | FBgn0030100 | CG12106 | 0.062711019 | 0.67447417 | 0 |
| RE01586 | FBgn0003285 | rst | -0.062710465 | 0.65890838 | 0 |
| SD10848 | FBgn0040305 | MTF-1 | -0.062672906 | 0.55996416 | 0 |
| GH13168 | FBgn0038930 | CG5778 | 0.062598705 | 0.65761341 | 0 |
| GH25014 | FBgn0039936 | Gyf | 0.062590005 | 0.66170242 | 0 |
| LP02841 | FBgn0039593 | Sid | 0.062527931 | 0.74043457 | 0 |
| SD02518 | FBgn0053116 | CG33116 | -0.062521484 | 0.55112556 | 0 |
| GH09096 | FBgn0052066 | CG32066 | -0.062490422 | 0.54464515 | 0 |
| AT19803 | FBgn0040376 | AANATL7 | 0.062478975 | 0.61410296 | 0 |
| RE15472 | FBgn0000137 | ase | 0.062451359 | 0.55130538 | 0 |
| GM25163 | FBgn0284408 | trol | -0.062439492 | 0.63612855 | 0 |
| GH02452 | FBgn0045770 | S-Lap3 | -0.062376972 | 0.8099408 | 0 |
| LD23533 | FBgn0027655 | htt | 0.062364951 | 0.62327293 | 0 |

| | | | | | |
|---|---|---|---|---|---|
| LD34154 | FBgn0031881 | MME1 | 0.062320665 | 0.58061153 | 0 |
| SD08737 | FBgn0038519 | Prx3 | 0.062317744 | 0.53994009 | 0 |
| AT11648 | FBgn0039376 | CG14354 | 0.062296417 | 0.53608002 | 0 |
| GM08777 | FBgn0261574 | kug | 0.06228354 | 0.6675533 | 0 |
| LD03595 | FBgn0022029 | l(2)k01209 | -0.062258713 | 0.67909927 | 0 |
| LP05266 | FBgn0265271 | CG14717 | 0.062258255 | 0.77369211 | 0 |
| RE29709 | FBgn0034262 | swi2 | -0.062247085 | 0.58177873 | 0 |
| GH13914 | FBgn0031723 | CG7251 | -0.062235203 | 0.5410973 | 0 |
| GM01519 | FBgn0032923 | CG9248 | -0.062081612 | 0.54627099 | 0 |
| SD07354 | FBgn0003495 | spz | -0.06206113 | 0.52964617 | 0 |
| AT19107 | FBgn0036125 | Iyd | -0.061926638 | 0.66519803 | 0 |
| LP02557 | FBgn0052631 | CG32631 | -0.061889837 | 0.55816722 | 0 |
| GH10049 | FBgn0000206 | boss | 0.061877476 | 0.66102494 | 0 |
| RE36920 | FBgn0032429 | CG5446 | -0.061847457 | 0.62110021 | 0 |
| LD10588 | FBgn0270927 | betaGlu | 0.061810006 | 0.57255039 | 0 |
| RE11961 | FBgn0036099 | CG11811 | -0.06180886 | 0.58332023 | 0 |
| AT14090 | FBgn0037506 | Mics1 | 0.06177675 | 0.7057677 | 0 |
| AT09635 | FBgn0036329 | CG11262 | -0.061771745 | 0.53500525 | 0 |
| LD40776 | FBgn0034646 | Rae1 | -0.061756337 | 0.55622948 | 0 |
| LD09850 | FBgn0032863 | Cdc23 | 0.061743458 | 0.60358513 | 0 |
| GH11727 | FBgn0034948 | PPP1R15 | 0.061737599 | 0.5551644 | 0 |
| LD11851 | FBgn0003124 | polo | 0.061730125 | 0.58887916 | 0 |
| LD21615 | FBgn0029672 | CG2875 | 0.061720588 | 0.62551521 | 0 |
| LD04957 | FBgn0039994 | conu | -0.06170248 | 0.63605373 | 0 |
| RE11768 | FBgn0039459 | IntS12 | 0.061657198 | 0.57003352 | 0 |
| SD07234 | FBgn0011828 | Pxn | 0.061646485 | 0.88575032 | 0 |
| RE28824 | FBgn0034968 | RpL12 | 0.06158628 | 0.58332023 | 0 |
| GH05294 | FBgn0038946 | rdhB | 0.061561641 | 0.58558362 | 0 |
| RH02211 | FBgn0011455 | ND-SGDH | -0.061550487 | 0.71344353 | 0 |
| RE69226 | FBgn0035513 | Cpr64Ad | -0.061541931 | 0.54350573 | 0 |
| GH13855 | FBgn0085427 | CG34398 | -0.061534395 | 0.63793196 | 0 |
| RE46687 | FBgn0035737 | Cpr65Ec | -0.061505134 | 0.75764133 | 0 |
| LD47970 | FBgn0036301 | CG4069 | 0.061458697 | 0.58061153 | 0 |
| LD15939 | FBgn0039017 | CG6985 | -0.061457439 | 0.57261017 | 0 |
| RH07074 | FBgn0033515 | Ir47a | 0.061418423 | 0.56081877 | 0 |
| RH07074 | FBgn0265141 | Ir47b | 0.061418423 | 0.56081877 | 0 |
| SD11312 | FBgn0033450 | Lsm11 | -0.061401396 | 0.65315239 | 0 |
| RE07885 | FBgn0024897 | b6 | 0.061378188 | 0.68437858 | 0 |
| RE60404 | FBgn0030373 | moon | 0.061377605 | 0.60237747 | 0 |
| LD02646 | FBgn0033292 | Cyp4ad1 | 0.061372947 | 0.53326482 | 0 |
| LD37258 | FBgn0039126 | CG13601 | -0.061348249 | 0.55877041 | 0 |
| LD23959 | FBgn0069242 | eca | 0.061339556 | 0.5446744 | 0 |
| LD16296 | FBgn0031319 | CG4896 | -0.061318054 | 0.70792416 | 0 |
| AT05261 | FBgn0083960 | CG34124 | 0.061235452 | 0.69454841 | 0 |
| RH02379 | FBgn0038172 | Adgf-D | 0.061135595 | 0.57246588 | 0 |
| GH06489 | FBgn0002862 | Mst87F | 0.061130511 | 0.71810828 | 0 |
| RH59219 | FBgn0052817 | CG32817 | -0.061128291 | 0.58219562 | 0 |
| RH59219 | FBgn0029524 | CG3176 | -0.061128291 | 0.58219562 | 0 |

| | | | | | |
|---|---|---|---|---|---|
| LD18757 | FBgn0039115 | CG10214 | 0.061127932 | 0.64983042 | 0 |
| LD20439 | FBgn0019932 | SamDC | -0.061116314 | 0.68799293 | 0 |
| RE22711 | FBgn0033958 | jef | -0.061067916 | 0.55279134 | 0 |
| RE09008 | FBgn0034717 | CG5819 | 0.061037735 | 0.68107497 | 0 |
| SD02223 | FBgn0000719 | fog | 0.061022121 | 0.63620149 | 0 |
| AT21741 | FBgn0034842 | Prosbeta5R1 | 0.060997825 | 0.64428139 | 0 |
| SD10481 | FBgn0034271 | Vps50 | 0.060997128 | 0.53781013 | 0 |
| GH14910 | FBgn0028540 | CG9008 | 0.06097819 | 0.62452508 | 0 |
| GM03782 | FBgn0267823 | Gmer | 0.060936131 | 0.5946724 | 0 |
| LP09580 | FBgn0001089 | Gal | 0.060867356 | 0.6592 | 0 |
| LD02287 | FBgn0036254 | CG5645 | 0.060861872 | 0.55129267 | 0 |
| LD09559 | FBgn0050499 | CG30499 | 0.060791244 | 0.62949402 | 0 |
| RH61844 | FBgn0035020 | CG13585 | 0.06078693 | 0.58974392 | 0 |
| GM14828 | FBgn0030369 | Cyp318a1 | -0.060769705 | 0.63149088 | 0 |
| GH21145 | FBgn0042119 | Cpr65Au | 0.060740659 | 0.7087949 | 0 |
| SD12606 | FBgn0024957 | Irp-1B | -0.060728999 | 0.59614743 | 0 |
| RE41968 | FBgn0038767 | trem | 0.060672749 | 0.5446744 | 0 |
| LD17449 | FBgn0002524 | lace | 0.060666114 | 0.56085473 | 0 |
| RE62785 | FBgn0033471 | CG12134 | 0.060660271 | 0.57834171 | 0 |
| GH21048 | FBgn0031039 | Shawn | 0.060626531 | 0.70347091 | 0 |
| GH21048 | FBgn0031038 | Tyler | 0.060626531 | 0.70347091 | 0 |
| LD22579 | FBgn0001990 | wek | 0.060611374 | 0.55294141 | 0 |
| GH16625 | FBgn0029514 | 312 | -0.060555825 | 0.55454826 | 0 |
| AT17667 | FBgn0037461 | CG15177 | -0.060543987 | 0.5704794 | 0 |
| LD18835 | FBgn0038941 | CG7080 | -0.060542198 | 0.55754859 | 0 |
| LP05025 | FBgn0030578 | CG5347 | 0.060518532 | 0.56016313 | 0 |
| AT26889 | FBgn0017556 | Prosalpha4T2 | 0.060480214 | 0.55455148 | 0 |
| SD11113 | FBgn0034246 | Dcr-2 | 0.060478309 | 0.58996985 | 0 |
| LD23958 | FBgn0010265 | RpS13 | -0.060468629 | 0.6494646 | 0 |
| GH07093 | FBgn0038797 | Dic2 | 0.060391062 | 0.73573878 | 0 |
| GH13872 | FBgn0034580 | Cht8 | 0.060355499 | 0.78738786 | 0 |
| GM14480 | FBgn0001128 | Gpdh1 | -0.060344751 | 0.60209852 | 0 |
| RE17222 | FBgn0051109 | CG31109 | 0.060329009 | 0.54155484 | 0 |
| RE18856 | FBgn0038312 | Zip88E | 0.060326772 | 0.55010499 | 0 |
| GH06026 | FBgn0027583 | CG7601 | 0.060169847 | 0.59340524 | 0 |
| RE37138 | FBgn0052406 | PVRAP | -0.060138402 | 0.65061803 | 0 |
| GH11415 | FBgn0027546 | CG4766 | 0.060092503 | 0.84479527 | 0 |
| LD12826 | FBgn0012034 | AcCoAS | -0.060083194 | 0.54464515 | 0 |
| RE68347 | FBgn0039467 | CG14253 | 0.060035593 | 0.73326242 | 0 |
| RE68347 | FBgn0284229 | CG46313 | 0.060035593 | 0.73326242 | 0 |
| LD29917 | FBgn0262028 | lncRNA:CR42839 | -0.059973328 | 0.63364025 | 0 |
| GH20226 | FBgn0038652 | CG7720 | 0.059970447 | 0.77895895 | 0 |
| RE01338 | FBgn0039167 | CG17786 | 0.059851971 | 0.61343941 | 0 |
| GH01142 | FBgn0034198 | CG11400 | 0.059833107 | 0.75008294 | 0 |
| RE49431 | FBgn0036942 | CG7328 | 0.059775368 | 0.56235406 | 0 |
| RE43942 | FBgn0259170 | alpha-Man-Ia | 0.059729808 | 0.55130538 | 0 |
| GH27752 | FBgn0031476 | CG8813 | -0.059663583 | 0.66366905 | 0 |
| AT10635 | FBgn0034506 | CG13870 | -0.059602927 | 0.58996985 | 0 |

| | | | | | |
|---|---|---|---|---|---|
| RE49094 | FBgn0003733 | tor | 0.059559947 | 0.61914843 | 0 |
| RE57102 | FBgn0036612 | CG4998 | 0.059548122 | 0.58302715 | 0 |
| LD36161 | FBgn0024958 | Irp-1A | 0.059528418 | 0.59438716 | 0 |
| RH70154 | FBgn0032472 | CG9928 | -0.059520806 | 0.5673378 | 0 |
| GM14862 | FBgn0001308 | Khc | 0.059483514 | 0.74370858 | 0 |
| GH10914 | FBgn0004837 | Su(H) | -0.059473218 | 0.71796792 | 0 |
| GH22106 | FBgn0027654 | jdp | 0.059425066 | 0.70792416 | 0 |
| GH09339 | FBgn0038301 | CG6654 | 0.059404786 | 0.66005747 | 0 |
| LD31278 | FBgn0031909 | CG5181 | 0.059375258 | 0.57430725 | 0 |
| RE52596 | FBgn0031668 | CG31917 | 0.059336553 | 0.65040006 | 0 |
| RE52596 | FBgn0265968 | Tfb5 | 0.059336553 | 0.65040006 | 0 |
| GH09410 | FBgn0028974 | xmas-2 | 0.059282014 | 0.66412331 | 0 |
| RH42520 | FBgn0037715 | CG9399 | 0.059280374 | 0.60926429 | 0 |
| LD25232 | FBgn0015222 | Fer1HCH | 0.059113501 | 0.63613238 | 0 |
| RE39769 | FBgn0034425 | CG11906 | 0.059078114 | 0.5946724 | 0 |
| LD22559 | FBgn0038552 | Alg1 | -0.059076668 | 0.65069747 | 0 |
| RE12144 | FBgn0035427 | ckd | 0.059047935 | 0.6590961 | 0 |
| LD25556 | FBgn0285925 | Fas1 | 0.058951926 | 0.62578275 | 0 |
| GH08205 | FBgn0034871 | CG3906 | 0.058931844 | 0.7600784 | 0 |
| LD05405 | FBgn0051759 | CG31759 | 0.058920356 | 0.56760757 | 0 |
| LD05574 | FBgn0015024 | CkIalpha | 0.058881651 | 0.63104366 | 0 |
| RE29957 | FBgn0039969 | Fis1 | -0.058848488 | 0.64255682 | 0 |
| GH10486 | FBgn0033249 | CG11191 | -0.058839607 | 0.66503649 | 0 |
| RE17115 | FBgn0036105 | Blos4 | 0.05882571 | 0.55294141 | 0 |
| RE04224 | FBgn0034204 | CG10953 | 0.058802206 | 0.65520568 | 0 |
| GH19182 | FBgn0035917 | Zasp66 | -0.058794121 | 0.65209811 | 0 |
| GH20565 | FBgn0025573 | PpN58A | 0.058742732 | 0.71646669 | 0 |
| LD39576 | FBgn0039870 | CG1896 | 0.058712562 | 0.58536138 | 0 |
| RE74560 | FBgn0038766 | CG4854 | 0.058671683 | 0.63516293 | 0 |
| GH19275 | FBgn0035756 | unc-13-4A | -0.058667 | 0.83819637 | 0 |
| GM10839 | FBgn0026262 | bip2 | -0.058640572 | 0.64689723 | 0 |
| GH07976 | FBgn0033367 | PPO2 | 0.058589976 | 0.71610168 | 0 |
| LP11016 | FBgn0026090 | CG14812 | 0.05840095 | 0.55437845 | 0 |
| GH23965 | FBgn0030398 | Cpr11B | 0.058305576 | 0.62534909 | 0 |
| GH28833 | FBgn0051248 | CG31248 | 0.058291345 | 0.71134843 | 0 |
| GH07049 | FBgn0036106 | CG6409 | 0.058261046 | 0.59099743 | 0 |
| RE68725 | FBgn0034135 | Syn2 | -0.058206212 | 0.85108952 | 0 |
| LD05512 | FBgn0003141 | pr | 0.058180189 | 0.65299747 | 0 |
| LD46272 | FBgn0029936 | CG4617 | -0.058171893 | 0.5813999 | 0 |
| LD09536 | FBgn0261479 | nero | -0.058170357 | 0.64924683 | 0 |
| LD16446 | FBgn0033372 | CG13742 | 0.058145379 | 0.6308412 | 0 |
| LD15002 | FBgn0022361 | Pur-alpha | 0.058117474 | 0.80662381 | 0 |
| RH44662 | FBgn0011227 | ox | -0.058113244 | 0.56126345 | 0 |
| GM03781 | FBgn0032079 | CG31886 | -0.058103479 | 0.71914677 | 0 |
| LD18758 | FBgn0034763 | RYBP | 0.058016256 | 0.56419565 | 0 |
| RH45340 | FBgn0083167 | Neb-cGP | -0.058016205 | 0.62260022 | 0 |
| GM02147 | FBgn0030719 | eIF5 | 0.057999142 | 0.61780569 | 0 |
| GM02147 | FBgn0284228 | CG46312 | 0.057999142 | 0.61780569 | 0 |

| | | | | | |
|---|---|---|---|---|---|
| AT29460 | FBgn0260453 | CG17140 | 0.057944826 | 0.62586044 | 0 |
| AT29460 | FBgn0260454 | CG17139 | 0.057944826 | 0.62586044 | 0 |
| RE53819 | FBgn0036023 | CG18179 | -0.05788199 | 0.80991433 | 0 |
| RE53819 | FBgn0036024 | CG18180 | -0.05788199 | 0.80991433 | 0 |
| AT15655 | FBgn0035483 | Mul1 | -0.057825236 | 0.57263883 | 0 |
| LD41023 | FBgn0034727 | mRpS29 | -0.057770457 | 0.572498 | 0 |
| AT20781 | FBgn0031774 | CG9147 | 0.057705494 | 0.66837183 | 0 |
| SD08167 | FBgn0030808 | RhoGAP15B | 0.057662516 | 0.57065145 | 0 |
| RE25411 | FBgn0033570 | ND-B14 | 0.057631735 | 0.6189734 | 0 |
| GM08026 | FBgn0259937 | Nop60B | -0.057612688 | 0.59966849 | 0 |
| GM08026 | FBgn0082935 | snoRNA:Me28S-G1083a | -0.057612688 | 0.59966849 | 0 |
| GM08026 | FBgn0082934 | snoRNA:Me28S-G1083b | -0.057612688 | 0.59966849 | 0 |
| GM08026 | FBgn0082933 | snoRNA:Me28S-G1083c | -0.057612688 | 0.59966849 | 0 |
| GM08026 | FBgn0082932 | snoRNA:Me28S-G1083d | -0.057612688 | 0.59966849 | 0 |
| GM08026 | FBgn0261973 | snoRNA:Or-CD9a | -0.057612688 | 0.59966849 | 0 |
| GM08026 | FBgn0261974 | snoRNA:Or-CD9b | -0.057612688 | 0.59966849 | 0 |
| AT01821 | FBgn0029084 | gom | 0.057592588 | 0.6777421 | 0 |
| GH08420 | FBgn0027563 | CG9631 | -0.057569832 | 0.66120595 | 0 |
| LD23740 | FBgn0014028 | SdhB | 0.057490907 | 0.61060702 | 0 |
| GH02974 | FBgn0000077 | amx | 0.057401629 | 0.57219448 | 0 |
| SD08670 | FBgn0039218 | Rpb10 | 0.057376954 | 0.62659876 | 0 |
| AT03911 | FBgn0264297 | CG43775 | -0.05726137 | 0.60926429 | 0 |
| AT03911 | FBgn0264298 | CG43776 | -0.05726137 | 0.60926429 | 0 |
| AT03911 | FBgn0264299 | CG43777 | -0.05726137 | 0.60926429 | 0 |
| GH26702 | FBgn0035598 | CG4669 | 0.057253443 | 0.61696801 | 0 |
| AT21490 | FBgn0036704 | CG6497 | 0.057147726 | 0.7105132 | 0 |
| LD14743 | FBgn0003514 | sqh | 0.057127939 | 0.63391225 | 0 |
| GH09166 | FBgn0028993 | scro | 0.056988371 | 0.68189609 | 0 |
| HL06052 | FBgn0053203 | CG33203 | 0.056981481 | 0.6299384 | 0 |
| LD22548 | FBgn0027508 | Tnks | 0.05685651 | 0.61753324 | 0 |
| RH10188 | FBgn0052553 | lncRNA:CR32553 | 0.056854579 | 0.66317477 | 0 |
| SD03412 | FBgn0260475 | CG30059 | 0.056797163 | 0.5700488 | 0 |
| SD03412 | FBgn0033836 | CG18278 | 0.056797163 | 0.5700488 | 0 |
| LD26757 | FBgn0041582 | tamo | 0.05678317 | 0.67926481 | 0 |
| RE17836 | FBgn0038277 | RpS5b | 0.056767606 | 0.6770583 | 0 |
| AT14346 | FBgn0039707 | Prtl99C | -0.056750177 | 0.72612808 | 0 |
| RH56745 | FBgn0036590 | CG13065 | -0.056728647 | 0.7351973 | 0 |
| SD10877 | FBgn0032000 | CG8372 | 0.056661211 | 0.57576423 | 0 |
| RE66227 | FBgn0033081 | geminin | -0.056609552 | 0.57027683 | 0 |
| LD22449 | FBgn0036299 | Tsf2 | -0.056539739 | 0.61067189 | 0 |
| LD29476 | FBgn0015791 | Rab14 | 0.056508593 | 0.65895948 | 0 |
| GH12891 | FBgn0261260 | mgl | 0.056456045 | 0.71632885 | 0 |
| RE20382 | FBgn0019925 | Surf4 | 0.056418171 | 0.63761196 | 0 |
| LD47549 | FBgn0038738 | CG4572 | 0.056393989 | 0.5831082 | 0 |
| LD12755 | FBgn0086686 | l(3)L1231 | 0.056385231 | 0.68911606 | 0 |
| RE17611 | FBgn0036825 | RpL26 | 0.05637125 | 0.78406297 | 0 |
| LD30995 | FBgn0037472 | CG10098 | -0.056327007 | 0.60844177 | 0 |
| SD03117 | FBgn0004397 | Vinc | 0.056325099 | 0.57579748 | 0 |

| | | | | | |
|---|---|---|---|---|---|
| SD02148 | FBgn0028343 | Ankle2 | 0.056320493 | 0.65209811 | 0 |
| GH12641 | FBgn0025592 | Gk1 | 0.056290619 | 0.58332023 | 0 |
| LD33485 | FBgn0001995 | mRpL4 | -0.05626183 | 0.63569444 | 0 |
| HL02449 | FBgn0036783 | CheA75a | -0.056210285 | 0.6308412 | 0 |
| LD11885 | FBgn0001078 | ftz-f1 | 0.056152074 | 0.65712854 | 0 |
| LD11885 | FBgn0267601 | lncRNA:CR45939 | 0.056152074 | 0.65712854 | 0 |
| AT20659 | FBgn0034786 | CG13531 | 0.056145233 | 0.59873026 | 0 |
| RE16440 | FBgn0034503 | MED8 | -0.056131129 | 0.62551521 | 0 |
| GH14326 | FBgn0034703 | CG3045 | 0.05604002 | 0.57988153 | 0 |
| GM01172 | FBgn0036291 | CG10681 | 0.056036805 | 0.5946724 | 0 |
| LD22858 | FBgn0031696 | Bub1 | 0.05596364 | 0.5831082 | 0 |
| AT01479 | FBgn0038258 | CG7362 | -0.055961333 | 0.64428139 | 0 |
| SD11716 | FBgn0263316 | Mrp4 | 0.055950795 | 0.69456206 | 0 |
| RH05666 | FBgn0028544 | CG16884 | 0.055924511 | 0.73322951 | 0 |
| LD30661 | FBgn0035586 | Fitm | 0.055879624 | 0.73128205 | 0 |
| AT16669 | FBgn0031085 | CG9570 | -0.055828088 | 0.77846913 | 0 |
| LD37985 | FBgn0035111 | Dis3l2 | 0.055767485 | 0.57807986 | 0 |
| GH09825 | FBgn0030883 | CG7772 | -0.055684486 | 0.7089224 | 0 |
| LD29815 | FBgn0032859 | Arpc2 | -0.055668485 | 0.66009873 | 0 |
| LP09631 | FBgn0039620 | wat | 0.055616376 | 0.67174105 | 0 |
| RE23981 | FBgn0029646 | CG14423 | -0.055519639 | 0.76708087 | 0 |
| LD23647 | FBgn0032454 | CG5787 | 0.055497375 | 0.60063579 | 0 |
| GM25059 | FBgn0036589 | CG13067 | 0.055457308 | 0.82686762 | 0 |
| RE64280 | FBgn0032210 | CYLD | 0.055452487 | 0.74321414 | 0 |
| GH21666 | FBgn0050090 | CG30090 | 0.055450462 | 0.71438632 | 0 |
| LD01635 | FBgn0032120 | CG33298 | -0.055418505 | 0.59139012 | 0 |
| SD04477 | FBgn0001297 | kay | 0.055403704 | 0.77665449 | 0 |
| GH16843 | FBgn0052485 | CG32485 | -0.055382857 | 0.73822829 | 0 |
| LD45244 | FBgn0034451 | TBCB | -0.055373597 | 0.66120595 | 0 |
| LD29223 | FBgn0026192 | par-6 | -0.055367128 | 0.64418025 | 0 |
| SD04019 | FBgn0046763 | CG17278 | -0.055355691 | 0.7743912 | 0 |
| LD10028 | FBgn0037330 | mRpL44 | 0.055298251 | 0.62949402 | 0 |
| LD23405 | FBgn0032408 | CG6712 | 0.055282026 | 0.60237747 | 0 |
| LD17235 | FBgn0013325 | RpL11 | 0.055277374 | 0.62578275 | 0 |
| AT11044 | FBgn0031187 | Usp2 | 0.055242554 | 0.73554257 | 0 |
| AT11044 | FBgn0039463 | CG18472 | 0.055242554 | 0.73554257 | 0 |
| RE44624 | FBgn0051715 | CG31715 | 0.055194295 | 0.59982904 | 0 |
| RE54691 | FBgn0034938 | CG3803 | -0.055181832 | 0.67163573 | 0 |
| GM01628 | FBgn0035829 | HP4 | 0.055178776 | 0.68590315 | 0 |
| LD22957 | FBgn0010315 | CycD | 0.055169199 | 0.58660837 | 0 |
| LD40170 | FBgn0031420 | Atxn7 | 0.055112609 | 0.64924683 | 0 |
| RE54405 | FBgn0023496 | Lip1 | -0.055104922 | 0.85243914 | 0 |
| GH07105 | FBgn0053995 | CG33995 | 0.055082374 | 0.6299384 | 0 |
| GH07105 | FBgn0264742 | CG44000 | 0.055082374 | 0.6299384 | 0 |
| GH07105 | FBgn0264743 | CG44001 | 0.055082374 | 0.6299384 | 0 |
| LP07116 | FBgn0039470 | CG6296 | 0.055046921 | 0.77583645 | 0 |
| RE06748 | FBgn0032059 | PrBP | 0.055005668 | 0.70379538 | 0 |
| LD29377 | FBgn0035853 | UbcE2M | 0.054981676 | 0.61374063 | 0 |

| | | | | | |
|---|---|---|---|---|---|
| HL07902 | FBgn0036768 | CG7402 | -0.054949908 | 0.72237475 | 0 |
| RE48009 | FBgn0002985 | odd | -0.054849879 | 0.78816984 | 0 |
| GH02250 | FBgn0041102 | ocn | 0.05480746 | 0.69301704 | 0 |
| LD21771 | FBgn0003655 | swa | -0.054765352 | 0.62436909 | 0 |
| LD18112 | FBgn0032997 | CG17486 | -0.054761108 | 0.76272184 | 0 |
| GH10002 | FBgn0038675 | CG6013 | 0.054758086 | 0.60088213 | 0 |
| LD40777 | FBgn0036184 | PCID2 | 0.054632244 | 0.59605473 | 0 |
| RE05591 | FBgn0266711 | EloC | -0.054628766 | 0.64603483 | 0 |
| RE55001 | FBgn0062440 | EMRE | 0.054577902 | 0.72296584 | 0 |
| GM13116 | FBgn0034420 | CG10737 | 0.054555493 | 0.67579802 | 0 |
| AT12544 | FBgn0029703 | CG12692 | 0.054487197 | 0.76425756 | 0 |
| LD04578 | FBgn0011826 | Pp2B-14D | -0.054481496 | 0.59471571 | 0 |
| GH15539 | FBgn0004368 | Ptp4E | 0.054474768 | 0.61025385 | 0 |
| GH25778 | FBgn0263397 | Ih | -0.054321427 | 0.67194867 | 0 |
| RE44901 | FBgn0010590 | Prosbeta1 | 0.054309006 | 0.58332023 | 0 |
| LD25626 | FBgn0046685 | Wsck | 0.05430734 | 0.58993327 | 0 |
| SD14939 | FBgn0036534 | DCP2 | 0.054275273 | 0.63748581 | 0 |
| GH17639 | FBgn0266723 | Trs31 | -0.054272131 | 0.63765433 | 0 |
| RE30975 | FBgn0034940 | CG16787 | -0.054178576 | 0.65473862 | 0 |
| LD35981 | FBgn0030959 | CG6961 | 0.054142912 | 0.58332023 | 0 |
| GH09810 | FBgn0011576 | Cyp4d2 | 0.054108605 | 0.73818379 | 0 |
| GH12788 | FBgn0260012 | pds5 | -0.054106706 | 0.74304581 | 0 |
| RE31310 | FBgn0052476 | mthl14 | 0.054096147 | 0.60873523 | 0 |
| SD14874 | FBgn0032024 | CG14273 | 0.054089455 | 0.75299807 | 0 |
| RE57452 | FBgn0030394 | Cpr11A | -0.054064765 | 0.84184002 | 0 |
| LD19391 | FBgn0002673 | twe | 0.054039303 | 0.68341559 | 0 |
| GH03827 | FBgn0034844 | CG9861 | -0.05401588 | 0.59731197 | 0 |
| GH03748 | FBgn0259700 | CG42354 | -0.054004755 | 0.63834771 | 0 |
| GH03748 | FBgn0259699 | CG42353 | -0.054004755 | 0.63834771 | 0 |
| GH26575 | FBgn0037727 | CG8358 | -0.053961796 | 0.73119662 | 0 |
| GH26058 | FBgn0034263 | CG10934 | 0.053941018 | 0.65093878 | 0 |
| GH26058 | FBgn0034264 | Dlish | 0.053941018 | 0.65093878 | 0 |
| RE30049 | FBgn0030928 | CG15044 | -0.053909374 | 0.63410391 | 0 |
| LD40852 | FBgn0020255 | Ran | 0.053885264 | 0.67659661 | 0 |
| RH17539 | FBgn0032840 | sNPF | -0.053797774 | 0.73719326 | 0 |
| AT02388 | FBgn0034532 | CG13436 | -0.05376987 | 0.62452508 | 0 |
| RE58951 | FBgn0038986 | sit | 0.053748776 | 0.70614655 | 0 |
| GM14337 | FBgn0051908 | CG31908 | -0.053695744 | 0.60767971 | 0 |
| LP03104 | FBgn0036417 | CG7906 | 0.053611971 | 0.68647409 | 0 |
| LD24661 | FBgn0266671 | Sec6 | -0.05349761 | 0.72694226 | 0 |
| AT16002 | FBgn0035217 | FucTD | -0.05346949 | 0.65209811 | 0 |
| LD11394 | FBgn0030539 | CG1368 | -0.053392185 | 0.63104366 | 0 |
| GH16463 | FBgn0023525 | CG3191 | 0.053383901 | 0.60458241 | 0 |
| AT13539 | FBgn0031253 | CG11885 | 0.053364232 | 0.60773166 | 0 |
| GH02671 | FBgn0261279 | lqfR | -0.053345798 | 0.59352005 | 0 |
| SD08423 | FBgn0030502 | tth | -0.053338136 | 0.70843295 | 0 |
| GM09909 | FBgn0260484 | HIP | 0.053266103 | 0.76562949 | 0 |
| GM09909 | FBgn0029676 | HIP-R | 0.053266103 | 0.76562949 | 0 |

| | | | | | |
|---|---|---|---|---|---|
| LD24586 | FBgn0052409 | CG32409 | 0.053226971 | 0.74447794 | 0 |
| GH06439 | FBgn0013770 | Cp1 | 0.053168013 | 0.61411163 | 0 |
| RE60714 | FBgn0037217 | CG14636 | -0.053141626 | 0.73039335 | 0 |
| RE12757 | FBgn0260243 | E(var)3-9 | -0.05313276 | 0.65498342 | 0 |
| AT14664 | FBgn0012344 | Dh44 | 0.05310727 | 0.73554257 | 0 |
| AT07409 | FBgn0028944 | Semp1 | -0.053058683 | 0.59955054 | 0 |
| LD26050 | FBgn0027499 | wde | 0.052984361 | 0.5946724 | 0 |
| LD35060 | FBgn0032014 | CG7840 | 0.052961079 | 0.61184309 | 0 |
| GH20124 | FBgn0038225 | soti | -0.052935256 | 0.80473998 | 0 |
| GM13065 | FBgn0032620 | CG12288 | 0.052912748 | 0.66928654 | 0 |
| LD45860 | FBgn0029133 | REG | 0.052895235 | 0.61552841 | 0 |
| LD25880 | FBgn0051441 | CG31441 | 0.052881705 | 0.66878059 | 0 |
| LD07439 | FBgn0024291 | Sirt1 | 0.052811563 | 0.61374063 | 0 |
| LD22216 | FBgn0002780 | mod | 0.052805497 | 0.63516293 | 0 |
| SD07933 | FBgn0004050 | z | -0.0527421 | 0.70079235 | 0 |
| RH40480 | FBgn0005658 | Ets65A | 0.05271393 | 0.60706427 | 0 |
| LD47540 | FBgn0284246 | l(2)k09848 | 0.052676682 | 0.71210731 | 0 |
| GH12381 | FBgn0039419 | CG12290 | 0.052627508 | 0.63539356 | 0 |
| GH06388 | FBgn0033820 | CG4716 | 0.052614015 | 0.62260022 | 0 |
| RE37786 | FBgn0263133 | mEFG1 | -0.052583456 | 0.62260022 | 0 |
| GH10101 | FBgn0051475 | CG31475 | 0.052570687 | 0.7490882 | 0 |
| AT27014 | FBgn0010258 | Rbp4 | -0.05254638 | 0.71610168 | 0 |
| LD25077 | FBgn0039004 | Nup133 | 0.052518756 | 0.66476062 | 0 |
| RE14694 | FBgn0000032 | Acph-1 | -0.052453401 | 0.62152317 | 0 |
| AT05390 | FBgn0036729 | CG13733 | -0.05244684 | 0.72575842 | 0 |
| LD12153 | FBgn0024558 | Dph5 | -0.052357598 | 0.59731197 | 0 |
| AT05371 | FBgn0031722 | CG14011 | -0.052346773 | 0.67954061 | 0 |
| AT12601 | FBgn0035168 | cep290 | -0.052317662 | 0.66366905 | 0 |
| RE52979 | FBgn0037935 | CG6834 | -0.052228729 | 0.73818379 | 0 |
| GH21710 | FBgn0036405 | CG6833 | -0.052194658 | 0.65743493 | 0 |
| LD45730 | FBgn0002791 | mr | -0.052172346 | 0.62807781 | 0 |
| GH02880 | FBgn0023517 | Pgam5 | 0.05214861 | 0.70355841 | 0 |
| GH14842 | FBgn0040281 | Aplip1 | 0.052133136 | 0.65072121 | 0 |
| LD47188 | FBgn0260789 | mxc | -0.052120162 | 0.68045534 | 0 |
| GH26015 | FBgn0033679 | CG8888 | 0.052117137 | 0.78119705 | 0 |
| GH03852 | FBgn0000442 | Pkg21D | -0.052040284 | 0.75680551 | 0 |
| LD15527 | FBgn0039186 | CG5746 | 0.052009426 | 0.63075595 | 0 |
| GH13756 | FBgn0027582 | CG6230 | 0.051997357 | 0.63833628 | 0 |
| RE64551 | FBgn0015546 | spel1 | 0.051979044 | 0.60635127 | 0 |
| AT19883 | FBgn0036327 | CG10748 | -0.051959829 | 0.74392699 | 0 |
| RH45888 | FBgn0037313 | CG1161 | 0.051895216 | 0.64937219 | 0 |
| RE33833 | FBgn0032404 | RpL7-like | 0.051792646 | 0.63075595 | 0 |
| RE33833 | FBgn0085193 | CG34164 | 0.051792646 | 0.63075595 | 0 |
| LD14109 | FBgn0039132 | AP-1sigma | -0.0517793 | 0.76604496 | 0 |
| RE64348 | FBgn0001147 | gsb-n | 0.051773432 | 0.71579289 | 0 |
| AT01616 | FBgn0036925 | schuy | -0.051730354 | 0.86280439 | 0 |
| LD07651 | FBgn0037933 | Ho | 0.051712825 | 0.67214703 | 0 |
| GH04989 | FBgn0029709 | CHOp24 | 0.051703211 | 0.70503123 | 0 |

| | | | | | |
|---|---|---|---|---|---|
| AT26376 | FBgn0038125 | CG8141 | -0.051699046 | 0.85789581 | 0 |
| GM29372 | FBgn0032298 | CG6724 | 0.051693709 | 0.64810895 | 0 |
| RE22432 | FBgn0005427 | ewg | -0.051669347 | 0.62730277 | 0 |
| GH11112 | FBgn0027547 | CG1927 | 0.051612216 | 0.65386861 | 0 |
| LD33276 | FBgn0040346 | CG3704 | 0.051577669 | 0.63262065 | 0 |
| HL01743 | FBgn0037163 | laza | 0.051560037 | 0.65520568 | 0 |
| RH11377 | FBgn0032013 | Scgalpha | -0.051526136 | 0.71614595 | 0 |
| LD26005 | FBgn0020369 | Rpt6 | -0.051519069 | 0.63577062 | 0 |
| SD04329 | FBgn0003742 | tra2 | -0.051494724 | 0.65436806 | 0 |
| RE36966 | FBgn0036509 | CG7739 | 0.051488155 | 0.67458238 | 0 |
| SD02572 | FBgn0015905 | ast | -0.051486638 | 0.64915473 | 0 |
| GM14426 | FBgn0039509 | bigmax | 0.051464559 | 0.73818379 | 0 |
| GH13968 | FBgn0030344 | Nrd1 | -0.05144257 | 0.60251021 | 0 |
| AT07222 | FBgn0034689 | CG2921 | 0.051407165 | 0.69831057 | 0 |
| AT07222 | FBgn0037964 | CG14731 | 0.051407165 | 0.69831057 | 0 |
| AT21729 | FBgn0034459 | TTLL6A | 0.051406101 | 0.64746163 | 0 |
| GH13879 | FBgn0032889 | CG9331 | 0.051374258 | 0.70635098 | 0 |
| GM14559 | FBgn0284245 | eEF1alpha1 | -0.051323557 | 0.70843295 | 0 |
| LD11291 | FBgn0031631 | CG3225 | 0.051251085 | 0.61343941 | 0 |
| LD45776 | FBgn0045073 | Stim | 0.051230816 | 0.66102494 | 0 |
| GH07456 | FBgn0026373 | RpII33 | 0.05121925 | 0.6834709 | 0 |
| GH09439 | FBgn0027054 | CSN4 | 0.051202514 | 0.67313338 | 0 |
| SD04748 | FBgn0027079 | ValRS | 0.051158257 | 0.64868773 | 0 |
| GH02025 | FBgn0033710 | CG17739 | -0.051157951 | 0.73702091 | 0 |
| RE05288 | FBgn0031183 | CG14621 | -0.051145383 | 0.72535658 | 0 |
| LD05365 | FBgn0261266 | zuc | 0.051092133 | 0.72174238 | 0 |
| AT09463 | FBgn0036941 | CG7335 | 0.051083739 | 0.70374009 | 0 |
| AT09463 | FBgn0259741 | CG42395 | 0.051083739 | 0.70374009 | 0 |
| LD35420 | FBgn0034618 | CG9485 | -0.051081275 | 0.61499827 | 0 |
| RE22286 | FBgn0037020 | Pex14 | -0.051056561 | 0.60630427 | 0 |
| SD09427 | FBgn0036733 | U4-U6-60K | -0.051056421 | 0.66659034 | 0 |
| GH14656 | FBgn0046294 | CG12699 | 0.051010257 | 0.67579802 | 0 |
| AT11343 | FBgn0052022 | CG32022 | 0.050951367 | 0.66659034 | 0 |
| LD46306 | FBgn0030529 | Clic | -0.050947636 | 0.7089224 | 0 |
| GH02636 | FBgn0041150 | hoe1 | -0.050938759 | 0.80473998 | 0 |
| RH61560 | FBgn0041174 | Vhl | -0.050922798 | 0.72858694 | 0 |
| LD21768 | FBgn0259985 | Mppe | 0.050910789 | 0.65269004 | 0 |
| LD40282 | FBgn0030800 | Nprl2 | 0.050830733 | 0.69456206 | 0 |
| LP04941 | FBgn0030101 | CG12118 | -0.050806615 | 0.61948987 | 0 |
| RE33420 | FBgn0036906 | CG14102 | -0.050802552 | 0.63262065 | 0 |
| RE51539 | FBgn0039473 | CG17191 | -0.050801738 | 0.72681454 | 0 |
| AT04926 | FBgn0035380 | CG9970 | -0.050747397 | 0.6396014 | 0 |
| RE04259 | FBgn0001259 | in | 0.050735712 | 0.75752473 | 0 |
| RE34039 | FBgn0032101 | CG9586 | 0.050723306 | 0.63774559 | 0 |
| LD23569 | FBgn0034894 | sigmar | 0.050695262 | 0.64898489 | 0 |
| GM14877 | FBgn0038976 | Pfdn5 | -0.050690004 | 0.61067189 | 0 |
| RE06505 | FBgn0037730 | CG9444 | 0.050659289 | 0.7508361 | 0 |
| LP10549 | FBgn0034543 | galla-1 | -0.050657314 | 0.64605999 | 0 |

| | | | | | |
|---|---|---|---|---|---|
| RH26041 | FBgn0262728 | Pal2 | -0.050641306 | 0.64616221 | 0 |
| SD01562 | FBgn0034400 | CG15099 | -0.05063076 | 0.68438067 | 0 |
| LD46144 | FBgn0026566 | CG1307 | 0.050592092 | 0.67523158 | 0 |
| LD42916 | FBgn0032018 | CG7806 | 0.050578842 | 0.78816984 | 0 |
| LD27084 | FBgn0039664 | CG2006 | 0.050562585 | 0.73822829 | 0 |
| RE37229 | FBgn0086356 | tum | 0.050550071 | 0.67407955 | 0 |
| RH66269 | FBgn0017578 | Max | 0.050491509 | 0.61961874 | 0 |
| AT05209 | FBgn0033933 | CG10104 | 0.050489583 | 0.64658932 | 0 |
| LD08659 | FBgn0033117 | CG3358 | 0.050443948 | 0.68437858 | 0 |
| RH07667 | FBgn0038070 | CG6753 | -0.050427513 | 0.71201737 | 0 |
| RE39378 | FBgn0037533 | CD98hc | -0.05040255 | 0.70202124 | 0 |
| LD21269 | FBgn0035878 | CG7182 | 0.050343598 | 0.69503389 | 0 |
| RE27406 | FBgn0250757 | CG42235 | 0.05031969 | 0.61662647 | 0 |
| LD15403 | FBgn0086370 | sra | 0.050230927 | 0.65666725 | 0 |
| LD04933 | FBgn0034176 | ste24a | -0.050211442 | 0.62659876 | 0 |
| AT15161 | FBgn0052437 | CG32437 | 0.050158942 | 0.71530616 | 0 |
| LD04507 | FBgn0004573 | 5-HT7 | -0.050057324 | 0.63737895 | 0 |
| GH25405 | FBgn0033915 | CG8485 | 0.050030871 | 0.61499827 | 0 |
| GH02976 | FBgn0027600 | obst-B | -0.050026237 | 0.73117965 | 0 |
| AT03617 | FBgn0031529 | CG9662 | 0.050015779 | 0.67817051 | 0 |
| GH17475 | FBgn0032025 | CG7778 | -0.049904579 | 0.6419413 | 0 |
| GH15759 | FBgn0038398 | sxe2 | -0.049872404 | 0.74360435 | 0 |
| LD03394 | FBgn0010607 | l(2)05714 | 0.049859813 | 0.67259652 | 0 |
| LD03161 | FBgn0033375 | CG8078 | 0.049797557 | 0.71610168 | 0 |
| SD07421 | FBgn0027779 | VhaSFD | 0.049795267 | 0.62502208 | 0 |
| AT26201 | FBgn0035584 | CG17030 | -0.049777237 | 0.79228271 | 0 |
| LD33318 | FBgn0016697 | Prosalpha5 | -0.049745004 | 0.62586044 | 0 |
| RE24541 | FBgn0039958 | CG12567 | -0.049736062 | 0.68542466 | 0 |
| GH23463 | FBgn0004364 | 18w | 0.049725609 | 0.73898129 | 0 |
| GH12584 | FBgn0023179 | amon | -0.049653673 | 0.83241529 | 0 |
| LD09927 | FBgn0026257 | cav | 0.049597503 | 0.62370623 | 0 |
| GH08991 | FBgn0038388 | CG4287 | 0.049561087 | 0.67100423 | 0 |
| LD45253 | FBgn0085188 | CG34159 | 0.049387111 | 0.67476925 | 0 |
| GH27148 | FBgn0005671 | Vha55 | -0.049369616 | 0.72702922 | 0 |
| GM04269 | FBgn0031479 | Prx6005 | -0.049337884 | 0.62120043 | 0 |
| LD17939 | FBgn0030858 | IntS2 | 0.049336728 | 0.65774198 | 0 |
| LD34576 | FBgn0053503 | Cyp12d1-d | 0.049302745 | 0.6934758 | 0 |
| LD34576 | FBgn0050489 | Cyp12d1-p | 0.049302745 | 0.6934758 | 0 |
| AT29464 | FBgn0035943 | CG5653 | 0.049287484 | 0.80722509 | 0 |
| GH02581 | FBgn0023535 | arg | 0.049276508 | 0.66881994 | 0 |
| RH19679 | FBgn0040793 | CG7630 | -0.049186658 | 0.65725076 | 0 |
| LD36183 | FBgn0003076 | Pgm | -0.049130864 | 0.69692368 | 0 |
| RE60936 | FBgn0261854 | aPKC | 0.049051911 | 0.67009677 | 0 |
| SD03793 | FBgn0028894 | GMF | -0.048988211 | 0.66606074 | 0 |
| AT13857 | FBgn0039787 | CG9702 | 0.048964161 | 0.72174238 | 0 |
| RH02789 | FBgn0031822 | Phf5a | -0.048847211 | 0.66510973 | 0 |
| RH35689 | FBgn0016123 | Alp4 | 0.048830394 | 0.63656931 | 0 |
| RE53485 | FBgn0025800 | Smox | -0.048802118 | 0.64428139 | 0 |

| | | | | | |
|---|---|---|---|---|---|
| RE43246 | FBgn0285958 | Fuca | 0.048770078 | 0.82050364 | 0 |
| RE43246 | FBgn0036170 | CG11714 | 0.048770078 | 0.82050364 | 0 |
| GH17972 | FBgn0031491 | alpha4GT1 | 0.048734353 | 0.70130204 | 0 |
| RE46304 | FBgn0002899 | mus301 | 0.048720835 | 0.65069747 | 0 |
| RE67967 | FBgn0035582 | CG13705 | 0.048718105 | 0.71610168 | 0 |
| LD14594 | FBgn0004463 | RpIII128 | 0.048653288 | 0.74055331 | 0 |
| LD29455 | FBgn0001301 | kel | 0.048633682 | 0.73284139 | 0 |
| AT12764 | FBgn0033767 | CG13148 | 0.048609771 | 0.75215614 | 0 |
| GH06154 | FBgn0037057 | CG10512 | -0.04858232 | 0.69179173 | 0 |
| LD28084 | FBgn0014163 | fax | 0.048578364 | 0.91406259 | 0 |
| LD21404 | FBgn0023519 | mRpL16 | 0.048498539 | 0.72008127 | 0 |
| AT03047 | FBgn0028381 | Decay | 0.048488166 | 0.78402117 | 0 |
| HL02043 | FBgn0037084 | Syx6 | -0.048487455 | 0.62791269 | 0 |
| GH18514 | FBgn0033574 | Spn47C | -0.048485908 | 0.81060184 | 0 |
| RE23942 | FBgn0031001 | CG7884 | -0.048461652 | 0.82439188 | 0 |
| RH05112 | FBgn0035501 | CG1299 | -0.04845559 | 0.63419496 | 0 |
| RH04612 | FBgn0004403 | RpS14a | -0.048390413 | 0.6415074 | 0 |
| RH04612 | FBgn0004404 | RpS14b | -0.048390413 | 0.6415074 | 0 |
| LD46584 | FBgn0025725 | alphaCOP | -0.048388177 | 0.78339445 | 0 |
| SD07768 | FBgn0039774 | CDase | 0.048380198 | 0.72113867 | 0 |
| GH10330 | FBgn0035008 | CG3494 | 0.048344643 | 0.76481021 | 0 |
| LD12948 | FBgn0035461 | ntc | 0.048330362 | 0.71344353 | 0 |
| LP03115 | FBgn0266448 | CG45078 | 0.048272839 | 0.68501741 | 0 |
| SD10289 | FBgn0002174 | l(2)tid | -0.048243406 | 0.63780646 | 0 |
| LD12453 | FBgn0028991 | seq | -0.048237285 | 0.69407656 | 0 |
| LD47579 | FBgn0003525 | stg | 0.048234582 | 0.66827158 | 0 |
| GH24564 | FBgn0037766 | Teh1 | -0.048215169 | 0.74872476 | 0 |
| LD45836 | FBgn0037299 | Vps37B | 0.048146516 | 0.64252972 | 0 |
| GH26429 | FBgn0038603 | PKD | -0.04811644 | 0.74880707 | 0 |
| RE67845 | FBgn0032398 | CG6766 | 0.048106068 | 0.67285881 | 0 |
| LD24245 | FBgn0011763 | Dp | 0.04809913 | 0.66796118 | 0 |
| GH10354 | FBgn0261456 | hpo | 0.048060772 | 0.68045633 | 0 |
| LD29554 | FBgn0032717 | CG10600 | -0.048004055 | 0.65381313 | 0 |
| LD24833 | FBgn0037339 | Pi4KIIalpha | 0.047997439 | 0.65040006 | 0 |
| RE67261 | FBgn0003866 | tsh | 0.047990907 | 0.69311102 | 0 |
| GH15921 | FBgn0032495 | CG16820 | 0.047989746 | 0.73470242 | 0 |
| LD33331 | FBgn0039696 | Rnb | -0.047984257 | 0.64313511 | 0 |
| LD39409 | FBgn0024227 | aurB | 0.047981034 | 0.68799293 | 0 |
| LD19727 | FBgn0027865 | Tsp96F | 0.047899972 | 0.7743912 | 0 |
| LD44801 | FBgn0002306 | sas | 0.047836634 | 0.75659211 | 0 |
| GM04011 | FBgn0031777 | CG9154 | -0.04783234 | 0.6867439 | 0 |
| RE35807 | FBgn0033720 | CG13160 | -0.047800286 | 0.64350491 | 0 |
| LD32068 | FBgn0035213 | CG2199 | -0.047760835 | 0.65498342 | 0 |
| LP10945 | FBgn0039058 | VhaAC39-2 | 0.047727741 | 0.79228271 | 0 |
| AT31822 | FBgn0283625 | CG46298 | -0.047719853 | 0.68972272 | 0 |
| RE17910 | FBgn0052237 | CG32237 | -0.047701702 | 0.78183059 | 0 |
| GH04176 | FBgn0010620 | CG10939 | 0.04764144 | 0.63748581 | 0 |
| LD15005 | FBgn0036000 | CG3434 | 0.047600465 | 0.74230624 | 0 |

| | | | | | |
|---|---|---|---|---|---|
| LP10436 | FBgn0003890 | betaTub97EF | 0.0475974 | 0.65520568 | 0 |
| RE14181 | FBgn0017430 | Nelf-E | -0.047548773 | 0.66156276 | 0 |
| GM12790 | FBgn0259228 | C3G | 0.047547455 | 0.71914677 | 0 |
| LD40049 | FBgn0033943 | CG12869 | -0.047430085 | 0.75646603 | 0 |
| GH22690 | FBgn0039911 | CG1909 | 0.047398291 | 0.67259652 | 0 |
| LD26673 | FBgn0046776 | CR14033 | 0.047364552 | 0.63834771 | 0 |
| GH28372 | FBgn0034712 | CG3264 | 0.047331114 | 0.74331829 | 0 |
| GH28372 | FBgn0283480 | Alp2 | 0.047331114 | 0.74331829 | 0 |
| RE66104 | FBgn0013303 | Nca | 0.047314813 | 0.69384831 | 0 |
| RE31679 | FBgn0031654 | Jon25Bii | -0.04730078 | 0.67194867 | 0 |
| GH07312 | FBgn0027575 | GABA-B-R2 | -0.047241327 | 0.80097632 | 0 |
| AT01776 | FBgn0034907 | CG5539 | -0.047224857 | 0.72110813 | 0 |
| LD31045 | FBgn0035997 | phol | 0.047206041 | 0.64249481 | 0 |
| LD19266 | FBgn0029092 | ced-6 | -0.04715716 | 0.70863841 | 0 |
| RE06394 | FBgn0031830 | COX5B | 0.047145174 | 0.63748581 | 0 |
| LD47568 | FBgn0036263 | thoc6 | -0.047139827 | 0.66412331 | 0 |
| RE36204 | FBgn0030587 | CG9522 | -0.047111026 | 0.73772784 | 0 |
| LP07767 | FBgn0030816 | CG16700 | -0.047095677 | 0.75900853 | 0 |
| LD37858 | FBgn0038455 | CG14907 | -0.047036634 | 0.75275195 | 0 |
| LD37858 | FBgn0015351 | CG14906 | -0.047036634 | 0.75275195 | 0 |
| GH19521 | FBgn0266456 | CG45081 | 0.04700282 | 0.65062289 | 0 |
| LD22928 | FBgn0027950 | MBD-like | 0.046965179 | 0.76491456 | 0 |
| AT01806 | FBgn0030620 | CG5662 | 0.046883435 | 0.69096385 | 0 |
| LD10516 | FBgn0261268 | Cul3 | 0.046870764 | 0.67563785 | 0 |
| GH09623 | FBgn0039464 | CG6330 | 0.046870226 | 0.66741248 | 0 |
| LD23842 | FBgn0016041 | Tom40 | -0.046859202 | 0.68437858 | 0 |
| GH22833 | FBgn0035948 | CG5644 | 0.046791394 | 0.69692368 | 0 |
| RE15531 | FBgn0003057 | CG10598 | -0.04673269 | 0.74112273 | 0 |
| RE03989 | FBgn0037934 | CG6830 | -0.046731922 | 0.82001815 | 0 |
| RE37895 | FBgn0036229 | CG7248 | -0.046711299 | 0.6941614 | 0 |
| RE66826 | FBgn0039902 | Zip102B | -0.046710862 | 0.77146686 | 0 |
| GH04205 | FBgn0038749 | Xport-A | 0.046675214 | 0.70634951 | 0 |
| GH04205 | FBgn0260234 | Xport-B | 0.046675214 | 0.70634951 | 0 |
| RE67859 | FBgn0037184 | CG14450 | -0.046648413 | 0.65609442 | 0 |
| LD14839 | FBgn0260942 | bond | -0.046603715 | 0.70614655 | 0 |
| LD34429 | FBgn0052179 | Krn | 0.046591189 | 0.67834327 | 0 |
| GH19431 | FBgn0083969 | CG34133 | 0.046582026 | 0.75752473 | 0 |
| RE63575 | FBgn0050000 | GstT1 | -0.046572244 | 0.72612808 | 0 |
| GH07066 | FBgn0027579 | mino | 0.046568362 | 0.67436037 | 0 |
| LP05421 | FBgn0034221 | CG10764 | 0.046566487 | 0.65878894 | 0 |
| LD35132 | FBgn0260399 | gwl | 0.046553844 | 0.68641234 | 0 |
| LD17340 | FBgn0031815 | frj | -0.046531018 | 0.67167761 | 0 |
| SD06362 | FBgn0031022 | CG12204 | 0.046503596 | 0.71250408 | 0 |
| GH03106 | FBgn0014396 | tim | -0.046498403 | 0.67405248 | 0 |
| LD31064 | FBgn0026080 | Tip60 | -0.046450507 | 0.824186 | 0 |
| RE24439 | FBgn0035859 | CG13678 | -0.046431951 | 0.65287735 | 0 |
| RE24439 | FBgn0035858 | CG13674 | -0.046431951 | 0.65287735 | 0 |
| AT31745 | FBgn0032449 | CG17036 | 0.046425015 | 0.72812911 | 0 |

| | | | | | |
|---|---|---|---|---|---|
| SD10604 | FBgn0033387 | CG8008 | -0.046422021 | 0.65386861 | 0 |
| RE16777 | FBgn0029686 | Glos | 0.046417213 | 0.64928446 | 0 |
| LD21953 | FBgn0031090 | Rab35 | 0.046385933 | 0.68400985 | 0 |
| LD47387 | FBgn0037235 | CG1103 | -0.046383036 | 0.67523158 | 0 |
| LD44791 | FBgn0028424 | JhI-26 | 0.046325933 | 0.90542596 | 0 |
| GH20441 | FBgn0028870 | CG4691 | 0.046324494 | 0.66354741 | 0 |
| LD40103 | FBgn0029924 | CG4586 | 0.04632134 | 0.7089224 | 0 |
| AT01314 | FBgn0033628 | CG13203 | -0.046287394 | 0.72008127 | 0 |
| RE11224 | FBgn0039897 | CG1674 | -0.04619884 | 0.87986369 | 0 |
| LP01681 | FBgn0086368 | tw | -0.046168531 | 0.90197438 | 0 |
| RE33302 | FBgn0037708 | CG9386 | -0.04611672 | 0.66082443 | 0 |
| LD15470 | FBgn0026620 | tacc | 0.046046117 | 0.65895948 | 0 |
| AT04690 | FBgn0038218 | CG14841 | 0.046043103 | 0.7779676 | 0 |
| LD03945 | FBgn0034476 | Toll-7 | 0.046017334 | 0.6900713 | 0 |
| GH21941 | FBgn0263983 | CG43732 | 0.045988458 | 0.76660426 | 0 |
| GM09085 | FBgn0037999 | CG4860 | 0.045937047 | 0.751557 | 0 |
| RE11134 | FBgn0260986 | mei-38 | 0.045921018 | 0.67476925 | 0 |
| GM21080 | FBgn0036342 | CG11279 | -0.045919392 | 0.74331829 | 0 |
| GH20208 | FBgn0032072 | CG9483 | 0.045908211 | 0.76442259 | 0 |
| AT09290 | FBgn0038000 | CG10014 | 0.045902926 | 0.80787858 | 0 |
| GH09383 | FBgn0039213 | atl | 0.045882912 | 0.80278052 | 0 |
| LD28902 | FBgn0040236 | c11.1 | -0.045861229 | 0.66120595 | 0 |
| GM14851 | FBgn0036277 | CG10418 | 0.045812978 | 0.70792416 | 0 |
| LD11036 | FBgn0022786 | Hira | 0.045730526 | 0.67303541 | 0 |
| LD25753 | FBgn0038818 | Nep4 | 0.045680964 | 0.73822829 | 0 |
| LD39243 | FBgn0031896 | CG4502 | 0.045680311 | 0.65093878 | 0 |
| AT16577 | FBgn0034834 | LS2 | 0.045624592 | 0.79897121 | 0 |
| LD21969 | FBgn0001491 | l(1)10Bb | 0.045623536 | 0.69853819 | 0 |
| GH19726 | FBgn0030403 | CG1824 | 0.045591067 | 0.73702091 | 0 |
| LD03886 | FBgn0035689 | CG7376 | 0.045508279 | 0.71210731 | 0 |
| AT07685 | FBgn0033020 | COX4L | -0.045492487 | 0.70229545 | 0 |
| LD22649 | FBgn0285913 | red | -0.045391125 | 0.70792416 | 0 |
| RE01048 | FBgn0031357 | mRpL48 | 0.045343855 | 0.76229565 | 0 |
| LD47462 | FBgn0263697 | Uba3 | -0.045340696 | 0.74112273 | 0 |
| LD44733 | FBgn0034345 | CG5174 | 0.045308014 | 0.71591203 | 0 |
| SD06787 | FBgn0037222 | CG14642 | 0.045244079 | 0.80623065 | 0 |
| LD09551 | FBgn0035232 | CG12099 | 0.04523151 | 0.65315239 | 0 |
| AT29209 | FBgn0034739 | CG3927 | 0.045215673 | 0.67659661 | 0 |
| AT29209 | FBgn0034740 | nsr | 0.045215673 | 0.67659661 | 0 |
| AT29209 | FBgn0032898 | CR9337 | 0.045215673 | 0.67659661 | 0 |
| RE61564 | FBgn0033527 | CG11777 | -0.045178731 | 0.69187073 | 0 |
| LD20358 | FBgn0037019 | Pex16 | 0.045170198 | 0.66817438 | 0 |
| SD07650 | FBgn0032077 | CG9515 | -0.045158106 | 0.79049929 | 0 |
| SD07650 | FBgn0032076 | CG9510 | -0.045158106 | 0.79049929 | 0 |
| RH58440 | FBgn0029839 | CG4660 | 0.045145352 | 0.68085911 | 0 |
| GH27237 | FBgn0030099 | CG12056 | -0.045142304 | 0.75468121 | 0 |
| AT10149 | FBgn0014395 | tilB | 0.045135122 | 0.80404698 | 0 |
| LD21504 | FBgn0027526 | Ublcp1 | 0.045106151 | 0.70792416 | 0 |

| | | | | | |
|---|---|---|---|---|---|
| RE42624 | FBgn0033716 | Den1 | 0.045100656 | 0.74122585 | 0 |
| LD40292 | FBgn0035400 | CG11537 | -0.045059144 | 0.67970498 | 0 |
| LD01958 | FBgn0030452 | MFS10 | 0.04501553 | 0.7163269 | 0 |
| GH02377 | FBgn0031515 | CG9664 | 0.045002245 | 0.79497485 | 0 |
| GH14384 | FBgn0029161 | slmo | -0.044982178 | 0.67259652 | 0 |
| GH14384 | FBgn0085208 | CG34179 | -0.044982178 | 0.67259652 | 0 |
| LD39445 | FBgn0003862 | trx | -0.044953489 | 0.81546069 | 0 |
| LD14783 | FBgn0036777 | CG7341 | 0.044950132 | 0.7440048 | 0 |
| LD46519 | FBgn0004638 | drk | 0.044913933 | 0.66902685 | 0 |
| GH14774 | FBgn0035769 | CTCF | 0.04490928 | 0.6675533 | 0 |
| LD24220 | FBgn0023081 | gek | -0.044854332 | 0.80049362 | 0 |
| GH04680 | FBgn0035620 | CG5150 | -0.04484461 | 0.71610168 | 0 |
| LD02743 | FBgn0033194 | Vps13 | 0.044824828 | 0.70997929 | 0 |
| AT20767 | FBgn0030536 | CG11068 | -0.044790706 | 0.77895895 | 0 |
| GH11711 | FBgn0051872 | CG31872 | -0.044764806 | 0.67670719 | 0 |
| LP04651 | FBgn0037714 | CG9396 | -0.044641543 | 0.68931927 | 0 |
| LP04651 | FBgn0003261 | Rm62 | -0.044641543 | 0.68931927 | 0 |
| SD26403 | FBgn0026085 | dgt4 | -0.044619996 | 0.7600784 | 0 |
| LD35137 | FBgn0023407 | B4 | -0.044612452 | 0.7346144 | 0 |
| AT22207 | FBgn0035486 | Gr64d | -0.044605756 | 0.88783257 | 0 |
| AT22207 | FBgn0045476 | Gr64e | -0.044605756 | 0.88783257 | 0 |
| LD46233 | FBgn0032295 | CG12299 | 0.044546158 | 0.74880707 | 0 |
| LD09790 | FBgn0025809 | Paf-AHalpha | 0.044433663 | 0.6693745 | 0 |
| LD42595 | FBgn0037680 | pasi2 | 0.044346254 | 0.72828251 | 0 |
| RE59757 | FBgn0026239 | gukh | -0.0443376 | 0.68341559 | 0 |
| RE33235 | FBgn0005596 | yem | 0.044323769 | 0.664447 | 0 |
| RE62495 | FBgn0030102 | SmydA-9 | -0.044312495 | 0.85108952 | 0 |
| RH22417 | FBgn0030511 | CG11158 | -0.044307069 | 0.69173384 | 0 |
| GM31526 | FBgn0026089 | CG14817 | -0.04428535 | 0.67447417 | 0 |
| RH46075 | FBgn0034290 | CG5773 | 0.044244127 | 0.77362173 | 0 |
| RE57564 | FBgn0034232 | CG4866 | 0.044186982 | 0.67163573 | 0 |
| AT02364 | FBgn0051161 | CG31161 | 0.044167535 | 0.72702922 | 0 |
| RE24941 | FBgn0038095 | Cyp304a1 | 0.044120769 | 0.80665486 | 0 |
| LD23922 | FBgn0028369 | kirre | 0.043947876 | 0.7046166 | 0 |
| LD15606 | FBgn0038100 | Paip2 | -0.043911399 | 0.71344353 | 0 |
| GH06247 | FBgn0036239 | Pop2 | 0.04389765 | 0.73908154 | 0 |
| LD42147 | FBgn0261988 | Gprk2 | -0.043886786 | 0.78367742 | 0 |
| RE01652 | FBgn0037354 | CG12171 | -0.04384875 | 0.66413256 | 0 |
| RE01652 | FBgn0051548 | CG31548 | -0.04384875 | 0.66413256 | 0 |
| GH01474 | FBgn0031174 | CG1486 | 0.043842726 | 0.72694226 | 0 |
| RE23864 | FBgn0263852 | Dad1 | -0.043718995 | 0.6971039 | 0 |
| GH12661 | FBgn0052262 | CG32262 | 0.043703439 | 0.692482 | 0 |
| AT15593 | FBgn0013810 | Dhc36C | -0.04366667 | 0.72920516 | 0 |
| LD32974 | FBgn0026148 | CG12253 | -0.043656964 | 0.67275497 | 0 |
| GH27892 | FBgn0032055 | CG13091 | -0.043649979 | 0.67817051 | 0 |
| LD41235 | FBgn0283649 | elgi | 0.043613465 | 0.66398691 | 0 |
| GH14950 | FBgn0033134 | Tsp42El | 0.043589241 | 0.80609956 | 0 |
| LD21131 | FBgn0010226 | GstS1 | 0.043500106 | 0.67954061 | 0 |

| | | | | | |
|---|---|---|---|---|---|
| LD11871 | FBgn0011204 | cue | 0.043491596 | 0.71844404 | 0 |
| AT02643 | FBgn0051867 | CG31867 | 0.043471916 | 0.87049229 | 0 |
| AT29287 | FBgn0029964 | CG1409 | 0.043453275 | 0.67579802 | 0 |
| SD07008 | FBgn0032815 | CG10462 | -0.043444994 | 0.71796792 | 0 |
| AT16639 | FBgn0052703 | Erk7 | 0.043424522 | 0.70792416 | 0 |
| RE17505 | FBgn0250862 | CG42237 | -0.043421997 | 0.75900853 | 0 |
| RE31140 | FBgn0038337 | CG6125 | 0.043386216 | 0.69625016 | 0 |
| RE55080 | FBgn0037603 | CG11753 | 0.043368446 | 0.75461324 | 0 |
| RE17456 | FBgn0045063 | fdl | 0.043346135 | 0.67954061 | 0 |
| LD23890 | FBgn0035028 | Start1 | 0.043297219 | 0.67009677 | 0 |
| LD41760 | FBgn0044323 | Cka | 0.043285683 | 0.67163573 | 0 |
| GH19969 | FBgn0024187 | syd | -0.043278152 | 0.70618162 | 0 |
| RE58783 | FBgn0030162 | CG1986 | -0.043259883 | 0.67855177 | 0 |
| RE36587 | FBgn0044871 | Gos28 | 0.043239888 | 0.70541114 | 0 |
| RE35789 | FBgn0039259 | CG11781 | 0.043212176 | 0.77360345 | 0 |
| GH19532 | FBgn0030087 | CG7766 | -0.043209907 | 0.70245147 | 0 |
| LD22682 | FBgn0010382 | CycE | -0.043160663 | 0.69173384 | 0 |
| RE36786 | FBgn0266668 | Exo84 | 0.043006477 | 0.67340286 | 0 |
| LD18018 | FBgn0036826 | arx | 0.042997924 | 0.75181102 | 0 |
| RE14195 | FBgn0038072 | CG6225 | -0.042985205 | 0.74092387 | 0 |
| RE37047 | FBgn0024913 | Actbeta | 0.042971688 | 0.82879464 | 0 |
| LP03188 | FBgn0063127 | lncRNA:CR33938 | 0.042804782 | 0.80991433 | 0 |
| LD38087 | FBgn0037374 | jagn | 0.042771648 | 0.67954061 | 0 |
| RE55560 | FBgn0040366 | CG11398 | -0.042738129 | 0.70457747 | 0 |
| RE18364 | FBgn0051989 | Cap-D3 | -0.042735555 | 0.79007757 | 0 |
| LD45352 | FBgn0024362 | Naa30A | -0.042731135 | 0.6704492 | 0 |
| SD17344 | FBgn0040964 | CG18661 | 0.04261835 | 0.68045633 | 0 |
| SD17344 | FBgn0015316 | Try29F | 0.04261835 | 0.68045633 | 0 |
| GH21184 | FBgn0035980 | mRRF1 | 0.042605249 | 0.72724963 | 0 |
| RH07413 | FBgn0029594 | CG14806 | -0.042514872 | 0.68797776 | 0 |
| GH18838 | FBgn0261561 | CG42675 | 0.042450442 | 0.75900853 | 0 |
| AT30951 | FBgn0038200 | CG9920 | 0.042435976 | 0.83421415 | 0 |
| LD24073 | FBgn0036579 | CG5027 | 0.042430162 | 0.75388221 | 0 |
| RH22029 | FBgn0028862 | dao | 0.042396604 | 0.71430254 | 0 |
| GH21160 | FBgn0082585 | sprt | 0.042337664 | 0.71620412 | 0 |
| GH25848 | FBgn0036236 | CG6931 | 0.042322377 | 0.67610965 | 0 |
| GH14048 | FBgn0038598 | CG7131 | 0.042312051 | 0.75680551 | 0 |
| LD24440 | FBgn0011327 | Uch-L5 | -0.04225041 | 0.75819037 | 0 |
| LP02089 | FBgn0032039 | CG13385 | 0.042231317 | 0.67965876 | 0 |
| LP10922 | FBgn0036975 | CG5618 | 0.042095076 | 0.702698 | 0 |
| LD47671 | FBgn0031818 | CG9536 | -0.042084053 | 0.72366837 | 0 |
| LD23660 | FBgn0005695 | gcl | -0.042082808 | 0.7105132 | 0 |
| GH14137 | FBgn0267253 | CG32700 | -0.04198192 | 0.75008294 | 0 |
| RE62711 | FBgn0052442 | Arv1 | 0.041975385 | 0.72595991 | 0 |
| GH15335 | FBgn0260635 | Diap1 | 0.041953156 | 0.80430094 | 0 |
| AT26083 | FBgn0051226 | CG31226 | 0.041936899 | 0.86574262 | 0 |
| GH26380 | FBgn0028916 | CG33090 | 0.041936385 | 0.69068143 | 0 |
| GM02310 | FBgn0029980 | CG10778 | 0.041928288 | 0.70843295 | 0 |

| | | | | | |
|---|---|---|---|---|---|
| LD04267 | FBgn0013773 | Cyp6a22 | 0.041861785 | 0.67934041 | 0 |
| RH69713 | FBgn0034538 | CG16799 | -0.041794724 | 0.70717874 | 0 |
| RE70470 | FBgn0010383 | Cyp18a1 | 0.041734206 | 0.83365222 | 0 |
| RE66546 | FBgn0039112 | SdhD | 0.041712436 | 0.71253998 | 0 |
| AT03646 | FBgn0025455 | CycT | -0.041694528 | 0.73110648 | 0 |
| AT18217 | FBgn0026174 | SkpD | -0.041613402 | 0.8194291 | 0 |
| AT18217 | FBgn0026175 | SkpC | -0.041613402 | 0.8194291 | 0 |
| AT18217 | FBgn0031074 | SkpE | -0.041613402 | 0.8194291 | 0 |
| RH57268 | FBgn0039505 | CG5934 | 0.041605415 | 0.72165482 | 0 |
| AT22312 | FBgn0030839 | CG5613 | 0.041598066 | 0.737114 | 0 |
| AT18730 | FBgn0037572 | CG11698 | 0.041545526 | 0.82467864 | 0 |
| RE54143 | FBgn0001206 | Hmr | -0.041465535 | 0.70630672 | 0 |
| LP06112 | FBgn0039264 | CG11786 | 0.041463837 | 0.67954061 | 0 |
| RE40459 | FBgn0032907 | CG9272 | 0.041445974 | 0.76560865 | 0 |
| SD03837 | FBgn0030362 | regucalcin | -0.04138878 | 0.68314977 | 0 |
| RE16720 | FBgn0040339 | MED22 | 0.041357134 | 0.70541114 | 0 |
| RE24409 | FBgn0038294 | Mf | 0.041277912 | 0.87167543 | 0 |
| AT27353 | FBgn0035952 | CG5280 | 0.041258255 | 0.74334213 | 0 |
| LD43488 | FBgn0259736 | CG42390 | -0.041215928 | 0.72096436 | 0 |
| AT27514 | FBgn0261823 | Asx | 0.041150952 | 0.79788248 | 0 |
| GM14315 | FBgn0265174 | PIG-V | 0.04109656 | 0.68880174 | 0 |
| GH23534 | FBgn0020493 | Dad | -0.041050767 | 0.72522764 | 0 |
| LD40224 | FBgn0032509 | CG6523 | -0.040984687 | 0.76857044 | 0 |
| GH14443 | FBgn0030258 | CG1552 | 0.040983067 | 0.71610168 | 0 |
| GH20492 | FBgn0053519 | Unc-89 | -0.040949855 | 0.70503123 | 0 |
| SD14418 | FBgn0032780 | CG13085 | -0.04090583 | 0.78228048 | 0 |
| RE69679 | FBgn0063498 | GstE2 | 0.040899966 | 0.72113867 | 0 |
| LP01827 | FBgn0030747 | CG4301 | 0.040891004 | 0.68799293 | 0 |
| SD22572 | FBgn0260856 | Membrin | -0.040873461 | 0.75574821 | 0 |
| SD16138 | FBgn0036745 | CG7484 | -0.040796127 | 0.71238036 | 0 |
| GH19385 | FBgn0033294 | Mal-A4 | 0.040780645 | 0.70247199 | 0 |
| LD38670 | FBgn0030435 | CG4645 | 0.040728204 | 0.71903546 | 0 |
| LP05521 | FBgn0034288 | CG5084 | 0.040709124 | 0.70202124 | 0 |
| LD12750 | FBgn0030351 | CG1840 | 0.040699206 | 0.70808537 | 0 |
| LD22157 | FBgn0014469 | Cyp4e2 | -0.040668218 | 0.69096207 | 0 |
| RE54577 | FBgn0035043 | CG4781 | -0.040630112 | 0.84936695 | 0 |
| LP11751 | FBgn0023508 | Ocrl | -0.040558833 | 0.77583645 | 0 |
| SD04273 | FBgn0030289 | GCS1 | -0.040531716 | 0.71620412 | 0 |
| AT11049 | FBgn0035097 | CG13405 | 0.040517143 | 0.68797776 | 0 |
| LD32469 | FBgn0265298 | SC35 | 0.040458857 | 0.7276348 | 0 |
| LD37115 | FBgn0004391 | shtd | 0.040442813 | 0.69156295 | 0 |
| SD05989 | FBgn0038108 | CG7518 | -0.040422309 | 0.78977013 | 0 |
| GH20028 | FBgn0039165 | CG6204 | 0.04039717 | 0.76475904 | 0 |
| LD29716 | FBgn0052756 | CG32756 | -0.040375127 | 0.77725376 | 0 |
| LD24448 | FBgn0000173 | ben | 0.040336505 | 0.75659211 | 0 |
| LD16038 | FBgn0031227 | CG3709 | 0.040325134 | 0.69407656 | 0 |
| RE19358 | FBgn0032189 | Ripalpha | 0.04026296 | 0.74067491 | 0 |
| RE58855 | FBgn0003380 | Sh | 0.040258126 | 0.9059394 | 0 |

| | | | | | |
|---|---|---|---|---|---|
| RH34338 | FBgn0035469 | CG14977 | 0.040238388 | 0.71430254 | 0 |
| RE34120 | FBgn0022987 | qkr54B | 0.040225674 | 0.70906987 | 0 |
| RE57120 | FBgn0033017 | CG10465 | -0.040163575 | 0.76310556 | 0 |
| RE35023 | FBgn0038646 | CG7715 | -0.040151751 | 0.77362173 | 0 |
| LP09339 | FBgn0000451 | ect | -0.040123318 | 0.90245143 | 0 |
| GM21916 | FBgn0051278 | CG31278 | -0.040002175 | 0.69665888 | 0 |
| HL08111 | FBgn0003410 | sina | 0.039996733 | 0.72953692 | 0 |
| AT13736 | FBgn0036728 | UQCR-Q | 0.039965038 | 0.70291938 | 0 |
| LD11371 | FBgn0033551 | CG7222 | -0.03995021 | 0.80023376 | 0 |
| LP07821 | FBgn0033047 | CG7882 | -0.039927775 | 0.74112273 | 0 |
| LD23667 | FBgn0267849 | Syx7 | 0.03988655 | 0.70792416 | 0 |
| LP08817 | FBgn0037174 | CG14457 | 0.039848167 | 0.82957707 | 0 |
| GH25390 | FBgn0017561 | Ork1 | -0.039839147 | 0.76262842 | 0 |
| RE61046 | FBgn0025936 | Eph | -0.039752954 | 0.79506425 | 0 |
| GH11417 | FBgn0032049 | Bace | -0.039749516 | 0.70808537 | 0 |
| RH06833 | FBgn0266599 | Hsc70-4 | 0.039727871 | 0.70408868 | 0 |
| RH07236 | FBgn0031942 | CG7203 | -0.039699061 | 0.70247199 | 0 |
| GH01576 | FBgn0026252 | msk | -0.039687062 | 0.69724355 | 0 |
| GH01974 | FBgn0039024 | CG4721 | 0.0396698 | 0.8228587 | 0 |
| GH23743 | FBgn0035092 | Nplp1 | -0.03964533 | 0.70808537 | 0 |
| RE26296 | FBgn0034819 | CG9877 | 0.039641605 | 0.7552332 | 0 |
| AT11857 | FBgn0267730 | lncRNA:CR46061 | 0.039626734 | 0.79788248 | 0 |
| RE16690 | FBgn0035865 | CG7201 | 0.039578239 | 0.79828119 | 0 |
| RE22456 | FBgn0011692 | pav | -0.039554177 | 0.72877135 | 0 |
| LD24671 | FBgn0028427 | Ilk | 0.039533612 | 0.75038121 | 0 |
| LD22180 | FBgn0026149 | BCL7-like | 0.039520099 | 0.71113647 | 0 |
| RE14390 | FBgn0034143 | CG8303 | 0.039468037 | 0.7258219 | 0 |
| RE49158 | FBgn0034484 | CG11044 | -0.039430603 | 0.8192996 | 0 |
| RE19804 | FBgn0035692 | Sf3b6 | -0.039422885 | 0.74872476 | 0 |
| RE51713 | FBgn0037359 | MED27 | 0.039347785 | 0.70997929 | 0 |
| SD07805 | FBgn0064766 | CG7600 | 0.039244853 | 0.74335301 | 0 |
| RE40051 | FBgn0038042 | Scgbeta | 0.039242711 | 0.79695477 | 0 |
| GH01042 | FBgn0051988 | CG31988 | 0.039219485 | 0.69692368 | 0 |
| GH01042 | FBgn0051624 | CG31624 | 0.039219485 | 0.69692368 | 0 |
| AT10619 | FBgn0051639 | CG31639 | -0.039198474 | 0.70792416 | 0 |
| LP06384 | FBgn0030746 | CG9981 | -0.039169931 | 0.74072856 | 0 |
| LP06384 | FBgn0004907 | 14-3-3zeta | -0.039169931 | 0.74072856 | 0 |
| GH24702 | FBgn0014010 | Rab5 | -0.039168922 | 0.70146536 | 0 |
| RH27607 | FBgn0265101 | Sgt1 | 0.039124154 | 0.7170824 | 0 |
| RE23580 | FBgn0038437 | Sdhaf3 | -0.039049066 | 0.80049362 | 0 |
| AT28434 | FBgn0034121 | CG6262 | -0.03902506 | 0.86813326 | 0 |
| SD10469 | FBgn0033913 | CG8468 | -0.038963289 | 0.8845372 | 0 |
| GH01053 | FBgn0041707 | 7B2 | 0.038961209 | 0.71614595 | 0 |
| LD32148 | FBgn0027494 | RpS10a | 0.038961017 | 0.74370858 | 0 |
| GH17154 | FBgn0040228 | DCTN5-p25 | -0.038956793 | 0.70015384 | 0 |
| AT15210 | FBgn0002707 | mei-9 | 0.038940762 | 0.75373984 | 0 |
| LD43683 | FBgn0031097 | obst-A | 0.038722966 | 0.9089514 | 0 |
| AT08090 | FBgn0038018 | Tim17a1 | 0.038709514 | 0.77429856 | 0 |

| | | | | | |
|---|---|---|---|---|---|
| GH13859 | FBgn0050281 | CG30281 | -0.038665169 | 0.82050364 | 0 |
| SD05477 | FBgn0265726 | Nna1 | -0.038589194 | 0.75613967 | 0 |
| RE41896 | FBgn0259226 | CG42326 | -0.038569669 | 0.71796792 | 0 |
| LD47995 | FBgn0034494 | CG10444 | 0.038566067 | 0.72942072 | 0 |
| RE02926 | FBgn0036467 | CG12310 | -0.038515903 | 0.74121638 | 0 |
| LD20971 | FBgn0030803 | CG4880 | 0.038455702 | 0.74880707 | 0 |
| RH42281 | FBgn0031821 | KFase | 0.038453667 | 0.82086322 | 0 |
| SD02658 | FBgn0015282 | Rpt2 | -0.038428448 | 0.79748454 | 0 |
| GH10824 | FBgn0002938 | ninaC | -0.038415421 | 0.7643795 | 0 |
| LD27185 | FBgn0035906 | GstO2 | -0.038412601 | 0.73436132 | 0 |
| AT05239 | FBgn0030721 | CG12698 | 0.038403456 | 0.80310836 | 0 |
| AT26906 | FBgn0030215 | CG15296 | -0.038392101 | 0.78359071 | 0 |
| RE17141 | FBgn0035344 | Cyp4d20 | 0.03834772 | 0.74703605 | 0 |
| LD42058 | FBgn0036389 | ssp2 | -0.038302967 | 0.70792416 | 0 |
| GH12664 | FBgn0051025 | Ppi1 | 0.038295972 | 0.81546069 | 0 |
| RE04226 | FBgn0051708 | DIP-zeta | -0.038213976 | 0.74274998 | 0 |
| SD26982 | FBgn0032819 | CG10463 | 0.038176101 | 0.76719873 | 0 |
| LD41827 | FBgn0029819 | Usp30 | -0.038161154 | 0.71614595 | 0 |
| SD04942 | FBgn0000247 | ca | -0.038145483 | 0.73322951 | 0 |
| GH02741 | FBgn0034649 | PIG-M | 0.038105247 | 0.76031937 | 0 |
| GH02741 | FBgn0259727 | CG42381 | 0.038105247 | 0.76031937 | 0 |
| GH02741 | FBgn0259725 | CG42379 | 0.038105247 | 0.76031937 | 0 |
| GH02741 | FBgn0259726 | CG42380 | 0.038105247 | 0.76031937 | 0 |
| GH11973 | FBgn0261258 | rgn | 0.038102257 | 0.79497485 | 0 |
| GH18955 | FBgn0030157 | CG1468 | -0.037986865 | 0.82593113 | 0 |
| AT22116 | FBgn0029661 | CG16781 | 0.037979988 | 0.75766719 | 0 |
| GM20271 | FBgn0028932 | CG16890 | -0.037932838 | 0.83334335 | 0 |
| RE44303 | FBgn0034706 | CG11275 | 0.037828113 | 0.74880707 | 0 |
| SD08430 | FBgn0036662 | CG9706 | -0.037793869 | 0.70882981 | 0 |
| RE08107 | FBgn0053993 | CG33993 | 0.037755264 | 0.77929936 | 0 |
| RH18819 | FBgn0040389 | mRpL14 | 0.037752049 | 0.71254997 | 0 |
| RE17942 | FBgn0040208 | Kat60 | 0.037734477 | 0.7189053 | 0 |
| AT05246 | FBgn0038286 | CG6966 | -0.037706285 | 0.81837817 | 0 |
| GH23389 | FBgn0033287 | CG8701 | 0.037584012 | 0.72753023 | 0 |
| GH26507 | FBgn0030018 | slpr | -0.037551253 | 0.90650548 | 0 |
| GH09436 | FBgn0036116 | CG7888 | 0.037538721 | 0.7606236 | 0 |
| SD08006 | FBgn0030554 | CG1434 | 0.037520635 | 0.72120914 | 0 |
| GM08921 | FBgn0031145 | Ntf-2 | 0.037493709 | 0.76745448 | 0 |
| RE40325 | FBgn0010357 | betaTry | -0.03747587 | 0.87047102 | 0 |
| RE40325 | FBgn0050031 | CG30031 | -0.03747587 | 0.87047102 | 0 |
| RE40325 | FBgn0010358 | deltaTry | -0.03747587 | 0.87047102 | 0 |
| RE40325 | FBgn0050025 | CG30025 | -0.03747587 | 0.87047102 | 0 |
| RE40325 | FBgn0010359 | gammaTry | -0.03747587 | 0.87047102 | 0 |
| RE40325 | FBgn0003863 | alphaTry | -0.03747587 | 0.87047102 | 0 |
| RE51791 | FBgn0031663 | CG8891 | -0.037415335 | 0.74122585 | 0 |
| RE08812 | FBgn0033762 | ZnT49B | 0.037414975 | 0.74365742 | 0 |
| GH13369 | FBgn0003317 | sax | 0.037408165 | 0.73668522 | 0 |
| LD05344 | FBgn0031651 | mRpL24 | 0.037406057 | 0.80884662 | 0 |

| | | | | | |
|---|---|---|---|---|---|
| GH07242 | FBgn0025643 | CG3588 | 0.037361777 | 0.74183198 | 0 |
| RE27306 | FBgn0046875 | Obp83g | -0.037345194 | 0.8194291 | 0 |
| GH21655 | FBgn0019952 | Orct | 0.037275553 | 0.8059566 | 0 |
| RH28596 | FBgn0027793 | CG14787 | 0.037193562 | 0.87388986 | 0 |
| LD24257 | FBgn0020370 | TppII | 0.037108413 | 0.81417767 | 0 |
| GH21132 | FBgn0265262 | Vha68-1 | -0.037102982 | 0.76745448 | 0 |
| AT22183 | FBgn0035140 | CG17180 | -0.037088353 | 0.72753023 | 0 |
| RE64209 | FBgn0262717 | Skeletor | 0.037054413 | 0.7440048 | 0 |
| AT27602 | FBgn0024188 | san | -0.036933704 | 0.75287661 | 0 |
| GH03980 | FBgn0014000 | Hf | 0.036928227 | 0.78575402 | 0 |
| LP09690 | FBgn0036831 | CG6839 | -0.036927927 | 0.79788248 | 0 |
| GH26340 | FBgn0037502 | wa-cup | 0.036863279 | 0.86823692 | 0 |
| GH05582 | FBgn0027585 | CG8740 | 0.036831769 | 0.79736797 | 0 |
| RH06812 | FBgn0266186 | Vamp7 | 0.036780892 | 0.77429856 | 0 |
| HL03474 | FBgn0028526 | CG15293 | 0.036762679 | 0.86726761 | 0 |
| LD17329 | FBgn0266916 | Cenp-C | 0.036710884 | 0.73218417 | 0 |
| LD08717 | FBgn0029134 | Prosbeta5 | -0.036706108 | 0.74625465 | 0 |
| LP10861 | FBgn0035165 | CG13887 | -0.036638616 | 0.7454045 | 0 |
| RE06767 | FBgn0000318 | cl | -0.036585743 | 0.73822829 | 0 |
| LD21354 | FBgn0024222 | IKKbeta | -0.03658091 | 0.77583645 | 0 |
| LD15555 | FBgn0052446 | Atox1 | -0.036528828 | 0.775882 | 0 |
| LD35289 | FBgn0036096 | CG8003 | -0.036497978 | 0.80819092 | 0 |
| LD38671 | FBgn0039528 | dsd | 0.036481964 | 0.77623996 | 0 |
| RE54692 | FBgn0039882 | CG11576 | -0.036471535 | 0.72734227 | 0 |
| RE56803 | FBgn0053181 | CG33181 | -0.036431809 | 0.79422756 | 0 |
| GM04226 | FBgn0263006 | SERCA | -0.036310721 | 0.76443644 | 0 |
| LP01629 | FBgn0039760 | CG9682 | -0.036302458 | 0.74392699 | 0 |
| SD08787 | FBgn0263106 | DnaJ-1 | -0.036290385 | 0.7237343 | 0 |
| RE62710 | FBgn0038462 | CG17556 | 0.036274081 | 0.75576044 | 0 |
| RE62710 | FBgn0038461 | CG3678 | 0.036274081 | 0.75576044 | 0 |
| HL05815 | FBgn0011603 | ine | 0.036242982 | 0.82531199 | 0 |
| AT18606 | FBgn0031856 | CG11322 | -0.036203467 | 0.81425029 | 0 |
| AT28638 | FBgn0050267 | asRNA:CR30267 | 0.036163925 | 0.76595602 | 0 |
| GH21247 | FBgn0033229 | CG12822 | -0.036113932 | 0.72355512 | 0 |
| LP03070 | FBgn0040505 | Alk | 0.036106948 | 0.7440048 | 0 |
| LD32943 | FBgn0026196 | nop5 | 0.036106431 | 0.75646603 | 0 |
| SD03917 | FBgn0265140 | Meltrin | -0.036095218 | 0.72366837 | 0 |
| SD03592 | FBgn0034628 | Acox57D-p | -0.036034594 | 0.73822829 | 0 |
| RE64043 | FBgn0052373 | CG32373 | 0.035994957 | 0.86036844 | 0 |
| LP08456 | FBgn0040723 | CG5011 | -0.035874124 | 0.75215614 | 0 |
| LP08456 | FBgn0263081 | CG43349 | -0.035874124 | 0.75215614 | 0 |
| LD19982 | FBgn0036059 | nudE | 0.03587024 | 0.76631077 | 0 |
| AT19280 | FBgn0085405 | CG34376 | 0.035840634 | 0.79574682 | 0 |
| LD37839 | FBgn0032292 | EMC3 | 0.035810195 | 0.76912216 | 0 |
| RE42844 | FBgn0036058 | CG6707 | -0.035782838 | 0.75659211 | 0 |
| AT03788 | FBgn0250824 | CG33655 | 0.03566401 | 0.73822829 | 0 |
| GH07296 | FBgn0039190 | CG5762 | 0.035662926 | 0.81715471 | 0 |
| GM01315 | FBgn0034946 | CG3065 | -0.035657921 | 0.74112273 | 0 |

| | | | | | |
|---|---|---|---|---|---|
| LP01487 | FBgn0000567 | Eip74EF | 0.035565525 | 0.74177921 | 0 |
| SD08216 | FBgn0015359 | CG2034 | 0.035532293 | 0.74880707 | 0 |
| GH08311 | FBgn0013972 | Gycalpha99B | 0.035513394 | 0.84184002 | 0 |
| GH28809 | FBgn0023215 | Mnt | -0.035511863 | 0.90331456 | 0 |
| AT22559 | FBgn0051802 | CG31802 | 0.035509364 | 0.75299807 | 0 |
| LP09975 | FBgn0026404 | Dronc | 0.035436108 | 0.73128205 | 0 |
| LD24634 | FBgn0020305 | dbe | 0.035426293 | 0.77113187 | 0 |
| SD17959 | FBgn0259108 | futsch | 0.035401456 | 0.78868788 | 0 |
| RE01329 | FBgn0004110 | tin | 0.035384338 | 0.79506425 | 0 |
| RE27758 | FBgn0031231 | mRpL10 | -0.035304682 | 0.8099408 | 0 |
| GH27042 | FBgn0003390 | shf | -0.035301813 | 0.80207278 | 0 |
| LD20362 | FBgn0053229 | CG33229 | -0.035230105 | 0.76002645 | 0 |
| AT13019 | FBgn0069913 | CG4681 | 0.03522983 | 0.86823692 | 0 |
| LD22804 | FBgn0035432 | ZnT63C | 0.035187956 | 0.73719326 | 0 |
| LD27216 | FBgn0031233 | Tbc1d15-17 | 0.035092417 | 0.74408605 | 0 |
| RH55021 | FBgn0033558 | CG12344 | 0.035048725 | 0.73702091 | 0 |
| LD27667 | FBgn0033356 | CG8229 | 0.035020413 | 0.74426874 | 0 |
| RE34669 | FBgn0028746 | CG18508 | 0.034975298 | 0.75299807 | 0 |
| LD43152 | FBgn0010441 | pll | -0.034956538 | 0.74686036 | 0 |
| RE34324 | FBgn0029836 | CG5928 | 0.034955333 | 0.74344296 | 0 |
| GH12549 | FBgn0035914 | CG6282 | 0.034892929 | 0.8338329 | 0 |
| LD35341 | FBgn0032305 | CG6700 | 0.034871418 | 0.7454045 | 0 |
| LD23157 | FBgn0036974 | eRF1 | 0.03481983 | 0.75373984 | 0 |
| RE63168 | FBgn0036726 | QIL1 | -0.03481774 | 0.82339725 | 0 |
| SD01241 | FBgn0032886 | CG9328 | 0.03481225 | 0.76382068 | 0 |
| RH06639 | FBgn0032536 | Ance-3 | -0.034799258 | 0.77103327 | 0 |
| RE63406 | FBgn0033380 | Phax | -0.034798062 | 0.737114 | 0 |
| RE45259 | FBgn0037180 | CG14452 | 0.034777621 | 0.77570532 | 0 |
| RE45259 | FBgn0037178 | CG12546 | 0.034777621 | 0.77570532 | 0 |
| AT15039 | FBgn0037325 | CG12147 | 0.034686839 | 0.79828119 | 0 |
| GH14734 | FBgn0036015 | CG3088 | 0.034683576 | 0.74392699 | 0 |
| AT12527 | FBgn0040696 | CG18675 | 0.034665322 | 0.74177921 | 0 |
| RE03780 | FBgn0037204 | CG11131 | -0.034663693 | 0.81707667 | 0 |
| SD02418 | FBgn0261111 | BHD | 0.034569879 | 0.76382068 | 0 |
| RH26557 | FBgn0033067 | CG11211 | -0.034534091 | 0.80261258 | 0 |
| RE60462 | FBgn0052736 | CG32736 | -0.03453274 | 0.77429856 | 0 |
| RE60462 | FBgn0259204 | CG42308 | -0.03453274 | 0.77429856 | 0 |
| RE42781 | FBgn0052102 | CG32102 | 0.034526352 | 0.79828119 | 0 |
| LD34436 | FBgn0037608 | mRpL19 | 0.034514184 | 0.81266421 | 0 |
| AT21454 | FBgn0033479 | PIG-N | 0.03447996 | 0.74043457 | 0 |
| LD30731 | FBgn0031043 | Naa20A | 0.03445042 | 0.79897121 | 0 |
| LD22679 | FBgn0027507 | CG1344 | 0.034294528 | 0.7656592 | 0 |
| LD15658 | FBgn0040009 | CG17490 | -0.034292708 | 0.84079443 | 0 |
| RE49934 | FBgn0038737 | CG11447 | -0.034283001 | 0.77583645 | 0 |
| LD18447 | FBgn0032486 | mRF1 | -0.034280484 | 0.76031937 | 0 |
| LD17019 | FBgn0039158 | CG6182 | -0.034278549 | 0.78402117 | 0 |
| RE35233 | FBgn0029913 | Cht11 | 0.034222936 | 0.74344296 | 0 |
| LP06848 | FBgn0039251 | Trf4-2 | 0.034210588 | 0.7705541 | 0 |

| | | | | | |
|---|---|---|---|---|---|
| GH05994 | FBgn0015032 | Cyp4c3 | 0.034200767 | 0.87763697 | 0 |
| AT30094 | FBgn0051717 | CG31717 | -0.03419094 | 0.75595695 | 0 |
| RE37471 | FBgn0037773 | CG5359 | 0.034174014 | 0.75659211 | 0 |
| RH10246 | FBgn0038678 | mRpL55 | -0.034168005 | 0.79686702 | 0 |
| RE28759 | FBgn0032688 | CG15160 | 0.034167011 | 0.76031937 | 0 |
| RE26329 | FBgn0026324 | Taf10b | 0.034161008 | 0.77502283 | 0 |
| RE18408 | FBgn0035696 | Best2 | 0.03413894 | 0.75576044 | 0 |
| LD38128 | FBgn0036842 | CG3797 | 0.034135482 | 0.73862998 | 0 |
| AT18469 | FBgn0039450 | CG5484 | -0.03412201 | 0.74931637 | 0 |
| GH21176 | FBgn0034269 | CG6406 | 0.03412155 | 0.75038121 | 0 |
| SD09866 | FBgn0261363 | PPO3 | 0.034014156 | 0.75646603 | 0 |
| RE24638 | FBgn0039303 | CG11857 | 0.034006399 | 0.73818379 | 0 |
| GM23435 | FBgn0064116 | CG33713 | 0.033951896 | 0.77739978 | 0 |
| GM23435 | FBgn0064117 | SLIRP1 | 0.033951896 | 0.77739978 | 0 |
| LD43682 | FBgn0002552 | lin | -0.033947833 | 0.78738786 | 0 |
| RE34633 | FBgn0032116 | Mco1 | 0.033942191 | 0.83010914 | 0 |
| GH01975 | FBgn0003068 | per | -0.033937914 | 0.7454045 | 0 |
| AT28470 | FBgn0034031 | CG12963 | 0.033931504 | 0.80226572 | 0 |
| GH03731 | FBgn0085423 | CG34394 | 0.033904465 | 0.74183198 | 0 |
| AT23266 | FBgn0033357 | Tom7 | 0.033894116 | 0.76031937 | 0 |
| AT30494 | FBgn0032492 | Prosalpha6T | -0.033883693 | 0.74941378 | 0 |
| RE71183 | FBgn0036340 | SRm160 | -0.033778628 | 0.79476858 | 0 |
| RE59052 | FBgn0030887 | CG6867 | 0.033744364 | 0.78963579 | 0 |
| LD26402 | FBgn0042185 | MCU | -0.033741389 | 0.77858478 | 0 |
| GM14561 | FBgn0036847 | CNPYb | 0.033690897 | 0.775882 | 0 |
| RE36854 | FBgn0028693 | Rpn12 | 0.033678152 | 0.76745412 | 0 |
| AT01448 | FBgn0045035 | tefu | -0.033644895 | 0.75008294 | 0 |
| SD06791 | FBgn0033901 | O-fut1 | 0.033549285 | 0.79049929 | 0 |
| RE26667 | FBgn0032692 | CG15161 | -0.033523261 | 0.8485472 | 0 |
| LD29081 | FBgn0026313 | X11L | 0.033496196 | 0.76824763 | 0 |
| LD29596 | FBgn0000251 | cad | -0.033438986 | 0.76520502 | 0 |
| LD16074 | FBgn0265192 | Snp | 0.033335468 | 0.77789689 | 0 |
| RE28524 | FBgn0034986 | mRpS17 | 0.033275982 | 0.79475668 | 0 |
| SD27779 | FBgn0038590 | CG12320 | 0.03327001 | 0.7916694 | 0 |
| LD38533 | FBgn0024698 | Cpsf160 | 0.033267401 | 0.74474402 | 0 |
| RE74626 | FBgn0039908 | Asator | 0.033265093 | 0.85180474 | 0 |
| RH44531 | FBgn0032487 | Ski6 | 0.033263888 | 0.81707667 | 0 |
| RH44902 | FBgn0032249 | TBC1D16 | 0.033246467 | 0.82643222 | 0 |
| GH15170 | FBgn0037396 | CG11459 | -0.03321377 | 0.80430094 | 0 |
| GH15170 | FBgn0037397 | CR1075 | -0.03321377 | 0.80430094 | 0 |
| AT24439 | FBgn0034423 | tbrd-2 | 0.033209256 | 0.83365222 | 0 |
| LD37002 | FBgn0039212 | Syx18 | -0.033188785 | 0.75717197 | 0 |
| LD10526 | FBgn0087008 | e(y)3 | 0.033139741 | 0.86766483 | 0 |
| SD10157 | FBgn0003475 | spir | 0.033124576 | 0.75285742 | 0 |
| GH28507 | FBgn0033702 | CG8854 | -0.03309673 | 0.79748454 | 0 |
| LP06368 | FBgn0005670 | Cyp4d1 | 0.033091263 | 0.77623996 | 0 |
| RE40412 | FBgn0033100 | CG3420 | -0.033088622 | 0.80308212 | 0 |
| SD03467 | FBgn0266670 | Sec5 | -0.032997814 | 0.85064331 | 0 |

| | | | | | |
|---|---|---|---|---|---|
| LD20554 | FBgn0042178 | CG32165 | -0.032993262 | 0.77887541 | 0 |
| LD20554 | FBgn0042177 | CG32164 | -0.032993262 | 0.77887541 | 0 |
| SD12705 | FBgn0032811 | CG10268 | 0.032986662 | 0.77929936 | 0 |
| LD14312 | FBgn0041702 | CG15107 | -0.032929998 | 0.79676163 | 0 |
| LD31957 | FBgn0030809 | Ubr1 | -0.032915281 | 0.88541246 | 0 |
| RE26656 | FBgn0037414 | Osi7 | 0.032899618 | 0.97289186 | 0 |
| RE71854 | FBgn0031957 | TwdlE | -0.032827194 | 0.79694099 | 0 |
| AT21573 | FBgn0032522 | CG16848 | -0.03279302 | 0.7634483 | 0 |
| RE50056 | FBgn0030507 | CG11164 | 0.03275451 | 0.80014786 | 0 |
| RE54930 | FBgn0047092 | lncRNA:CR32730 | -0.032745538 | 0.89810972 | 0 |
| LD05816 | FBgn0052672 | Atg8a | 0.032590457 | 0.75332027 | 0 |
| RE01527 | FBgn0038474 | mRpS11 | -0.032558648 | 0.77848437 | 0 |
| AT11423 | FBgn0035768 | CG14834 | 0.032509865 | 0.84184002 | 0 |
| RE49227 | FBgn0040360 | CG14626 | -0.032482691 | 0.86445543 | 0 |
| LP06969 | FBgn0035300 | CG1139 | 0.032467168 | 0.82846125 | 0 |
| GH19049 | FBgn0266673 | Sec10 | 0.032397344 | 0.78183059 | 0 |
| RE43020 | FBgn0035429 | CG12017 | -0.03236771 | 0.75752473 | 0 |
| RE62456 | FBgn0042118 | Cpr65Ax2 | -0.032332282 | 0.84833378 | 0 |
| RE62456 | FBgn0086900 | Cpr65Ax1 | -0.032332282 | 0.84833378 | 0 |
| RH41663 | FBgn0034803 | CG9849 | 0.032306201 | 0.77847474 | 0 |
| LD42035 | FBgn0031320 | CG5126 | 0.032275193 | 0.75646603 | 0 |
| LD09043 | FBgn0027620 | Acf | -0.032238605 | 0.7506759 | 0 |
| GH16481 | FBgn0015034 | Cyp4e1 | 0.032211826 | 0.76425756 | 0 |
| GH19567 | FBgn0036773 | CG13698 | -0.032208391 | 0.78738786 | 0 |
| GH24939 | FBgn0032285 | CG17108 | -0.032174027 | 0.82072908 | 0 |
| RE38876 | FBgn0035160 | hng3 | -0.032171123 | 0.809388 | 0 |
| GH22047 | FBgn0031381 | Npc2a | -0.032161589 | 0.8266986 | 0 |
| GH27792 | FBgn0023526 | CG2865 | -0.032159085 | 0.8220452 | 0 |
| GM01211 | FBgn0034521 | Mgat1 | -0.032155391 | 0.77570532 | 0 |
| RH19795 | FBgn0039703 | CG7829 | -0.032121412 | 0.8491373 | 0 |
| RE10196 | FBgn0032845 | CG10747 | -0.032067805 | 0.86907327 | 0 |
| LD25692 | FBgn0086444 | l(2)37Cb | 0.032065893 | 0.75306535 | 0 |
| RE36877 | FBgn0034093 | CG15706 | -0.031965792 | 0.75369084 | 0 |
| RE14725 | FBgn0029868 | ND-B16.6 | 0.031944891 | 0.76031937 | 0 |
| HL03025 | FBgn0036890 | CG9368 | -0.031882222 | 0.79289038 | 0 |
| RE45833 | FBgn0032200 | CG5676 | 0.031848298 | 0.75994748 | 0 |
| LD46228 | FBgn0005694 | Aef1 | 0.031797453 | 0.75646603 | 0 |
| SD02517 | FBgn0016794 | dos | -0.031766279 | 0.77604805 | 0 |
| GH01404 | FBgn0027375 | RecQ5 | -0.031707965 | 0.81546069 | 0 |
| LD23637 | FBgn0039249 | CG11168 | -0.031619979 | 0.82713362 | 0 |
| LD38046 | FBgn0030187 | lpod | -0.031536165 | 0.76443644 | 0 |
| RE18506 | FBgn0040318 | HGTX | -0.031521253 | 0.8326161 | 0 |
| RE64386 | FBgn0036565 | CG5235 | -0.03148044 | 0.82349892 | 0 |
| RE41295 | FBgn0027291 | l(1)G0156 | 0.031468522 | 0.77848437 | 0 |
| AT30334 | FBgn0032291 | CG17118 | 0.031411186 | 0.82643222 | 0 |
| AT29994 | FBgn0028561 | sut3 | 0.031410977 | 0.86445543 | 0 |
| LD41522 | FBgn0039835 | mRpL32 | 0.031388325 | 0.82203015 | 0 |
| RE70153 | FBgn0052091 | CG32091 | -0.031379528 | 0.80991433 | 0 |

| | | | | | |
|---|---|---|---|---|---|
| LD32282 | FBgn0003557 | Su(dx) | -0.031368776 | 0.78183059 | 0 |
| RH27559 | FBgn0033817 | GstE14 | -0.031341389 | 0.90544935 | 0 |
| SD20666 | FBgn0008635 | betaCOP | 0.031299427 | 0.81635674 | 0 |
| GM14157 | FBgn0033691 | CG8860 | -0.031263855 | 0.85356356 | 0 |
| GH28366 | FBgn0033347 | CG8248 | -0.03123506 | 0.7916694 | 0 |
| AT13860 | FBgn0032464 | Vha68-3 | 0.031191035 | 0.8158978 | 0 |
| LD47924 | FBgn0000392 | cup | -0.031171499 | 0.77697185 | 0 |
| LD47924 | FBgn0085339 | CG34310 | -0.031171499 | 0.77697185 | 0 |
| RE63690 | FBgn0264711 | CG43980 | -0.031138989 | 0.80660681 | 0 |
| RE70141 | FBgn0052654 | Sec16 | 0.031121797 | 0.78687753 | 0 |
| LD28549 | FBgn0036306 | CG10973 | 0.031121442 | 0.80330987 | 0 |
| AT09395 | FBgn0053098 | CG33098 | -0.031116055 | 0.77877962 | 0 |
| GH13214 | FBgn0026679 | IntS4 | 0.031092971 | 0.77583645 | 0 |
| RH08557 | FBgn0004047 | Yp3 | 0.031062589 | 0.77274159 | 0 |
| RE26983 | FBgn0261976 | Psf2 | 0.03102545 | 0.83769067 | 0 |
| AT11549 | FBgn0085454 | CG34425 | 0.030915114 | 0.78977013 | 0 |
| GH17451 | FBgn0038105 | yellow-f2 | -0.030839855 | 0.8099408 | 0 |
| LD43819 | FBgn0032961 | CG1416 | -0.030721027 | 0.77583645 | 0 |
| RH02475 | FBgn0034588 | CG9394 | 0.030710175 | 0.84479527 | 0 |
| GH18334 | FBgn0038708 | CG6255 | -0.030707153 | 0.84878506 | 0 |
| LD27537 | FBgn0266452 | CTPsyn | 0.030703087 | 0.78501524 | 0 |
| LD22578 | FBgn0034452 | Oseg6 | -0.030657362 | 0.82679057 | 0 |
| AT08189 | FBgn0034658 | Grx1t | 0.030653161 | 0.82344805 | 0 |
| AT16061 | FBgn0031132 | CG15450 | 0.030637229 | 0.82342777 | 0 |
| LD11626 | FBgn0022772 | Orc1 | -0.030596151 | 0.84063592 | 0 |
| LD44381 | FBgn0010113 | hdc | -0.030495916 | 0.79095333 | 0 |
| RE47911 | FBgn0033348 | Spt | 0.030469337 | 0.76935806 | 0 |
| LP06572 | FBgn0039685 | Obp99b | -0.030448613 | 0.8556048 | 0 |
| LD43276 | FBgn0036828 | CG6841 | 0.030350379 | 0.81569022 | 0 |
| RE41314 | FBgn0039756 | CG9743 | -0.030308611 | 0.82643222 | 0 |
| RH03361 | FBgn0033027 | TpnC4 | -0.03030074 | 0.88066118 | 0 |
| RE50565 | FBgn0036462 | mRpL39 | 0.030231484 | 0.80665486 | 0 |
| RE59557 | FBgn0043001 | Chrac-16 | -0.030179086 | 0.77583645 | 0 |
| GH27419 | FBgn0036910 | Cyp305a1 | -0.030174528 | 0.88205076 | 0 |
| LP11159 | FBgn0053490 | CG33490 | -0.030149672 | 0.8409258 | 0 |
| LD11162 | FBgn0259733 | tal-AA | 0.030136848 | 0.86989242 | 0 |
| LD11162 | FBgn0259730 | tal-1A | 0.030136848 | 0.86989242 | 0 |
| LD11162 | FBgn0259731 | tal-2A | 0.030136848 | 0.86989242 | 0 |
| LD11162 | FBgn0259732 | tal-3A | 0.030136848 | 0.86989242 | 0 |
| SD05240 | FBgn0032446 | CG5780 | 0.030131705 | 0.80736526 | 0 |
| RE19845 | FBgn0003482 | spn-D | 0.030105521 | 0.79228271 | 0 |
| GH03607 | FBgn0038569 | CG7218 | 0.03010137 | 0.80819092 | 0 |
| RE18886 | FBgn0039873 | Smvt | 0.030065234 | 0.84505819 | 0 |
| LD36412 | FBgn0034368 | zda | -0.030025026 | 0.78816984 | 0 |
| GM14109 | FBgn0001942 | eIF4A | -0.02995903 | 0.78402117 | 0 |
| GH28796 | FBgn0035268 | CG8001 | 0.029937562 | 0.775882 | 0 |
| LD24561 | FBgn0010100 | Acon | 0.029873931 | 0.86813326 | 0 |
| SD16838 | FBgn0032034 | Rcd4 | -0.02983395 | 0.8102681 | 0 |

| | | | | | |
|---|---|---|---|---|---|
| RE11035 | FBgn0031548 | CG8852 | -0.029819461 | 0.77623996 | 0 |
| LD46418 | FBgn0029121 | Sras | -0.029786131 | 0.77052699 | 0 |
| LP10887 | FBgn0031249 | CG11911 | -0.029783903 | 0.83817966 | 0 |
| RH14406 | FBgn0030884 | CG6847 | -0.029697854 | 0.78808935 | 0 |
| LP11549 | FBgn0032966 | CG1421 | 0.02967066 | 0.88066118 | 0 |
| GH26896 | FBgn0040965 | lncRNA:CR13130 | 0.029640359 | 0.85180474 | 0 |
| LD47843 | FBgn0040011 | Slmap | -0.029632925 | 0.77887541 | 0 |
| LD16544 | FBgn0030218 | CG1628 | 0.029628706 | 0.8158978 | 0 |
| LD29885 | FBgn0027496 | epsilonCOP | -0.02961417 | 0.85512244 | 0 |
| GH01933 | FBgn0033793 | CG3955 | 0.029613027 | 0.81000343 | 0 |
| LD17053 | FBgn0037491 | CG1227 | -0.029541229 | 0.81707667 | 0 |
| AT23224 | FBgn0032090 | CG9582 | 0.029521019 | 0.80049362 | 0 |
| RE01453 | FBgn0030234 | CG15211 | -0.029465107 | 0.90968483 | 0 |
| GH28569 | FBgn0085450 | Snoo | 0.029449787 | 0.88979506 | 0 |
| GH26458 | FBgn0028888 | CG4168 | 0.029390336 | 0.85150759 | 0 |
| LD43561 | FBgn0023527 | CG3071 | 0.029323232 | 0.77794782 | 0 |
| RE01983 | FBgn0042696 | Nfl | -0.029291495 | 0.82690228 | 0 |
| LP05023 | FBgn0283476 | Dhc16F | 0.029242743 | 0.82531199 | 0 |
| AT13606 | FBgn0020280 | laf | 0.029220272 | 0.79816815 | 0 |
| LD21289 | FBgn0022893 | Df31 | 0.029219383 | 0.78752591 | 0 |
| AT11443 | FBgn0031372 | CG7295 | -0.02921187 | 0.8326161 | 0 |
| RE17991 | FBgn0285948 | RpL27A | 0.029146688 | 0.79897121 | 0 |
| HL01463 | FBgn0015663 | Dot | -0.029123229 | 0.89360343 | 0 |
| LP04865 | FBgn0026379 | Pten | -0.029120487 | 0.89010723 | 0 |
| LD33652 | FBgn0029512 | Aos1 | -0.029103126 | 0.80991433 | 0 |
| AT05678 | FBgn0029539 | CG11663 | 0.029085887 | 0.91093986 | 0 |
| RH66336 | FBgn0003515 | st | 0.029076168 | 0.824186 | 0 |
| RH51528 | FBgn0085256 | CG34227 | -0.029052674 | 0.86011954 | 0 |
| LD44824 | FBgn0259978 | vlc | 0.029046933 | 0.80407614 | 0 |
| LD23191 | FBgn0001124 | Got1 | -0.029045926 | 0.79539583 | 0 |
| LD20870 | FBgn0259199 | Snup | 0.028996042 | 0.86572919 | 0 |
| LD20870 | FBgn0259200 | CG42304 | 0.028996042 | 0.86572919 | 0 |
| GH13952 | FBgn0032242 | CG5355 | 0.02896349 | 0.80167761 | 0 |
| GH25188 | FBgn0263658 | lncRNA:CR43649 | -0.028921107 | 0.87887309 | 0 |
| GH08773 | FBgn0027562 | CG10345 | -0.028895018 | 0.81007863 | 0 |
| GH27756 | FBgn0030814 | CG4955 | 0.028866449 | 0.88891487 | 0 |
| AT30002 | FBgn0285963 | CG46339 | -0.028827135 | 0.78648364 | 0 |
| GH24781 | FBgn0004784 | inaC | -0.028674571 | 0.91347899 | 0 |
| RE69756 | FBgn0285971 | prg | 0.028584562 | 0.78102854 | 0 |
| GH03051 | FBgn0046114 | Gclm | -0.02856669 | 0.81417767 | 0 |
| LD01950 | FBgn0262689 | asRNA:CR43159 | -0.028552981 | 0.82167017 | 0 |
| GH02603 | FBgn0038160 | CG9759 | 0.028496139 | 0.81707667 | 0 |
| LD39664 | FBgn0024975 | CG2712 | -0.028476907 | 0.8099408 | 0 |
| LD21772 | FBgn0021761 | Nup154 | 0.028472722 | 0.81859024 | 0 |
| GH24271 | FBgn0034943 | Fmo-1 | -0.028428282 | 0.80028254 | 0 |
| GM14348 | FBgn0031848 | Nse1 | 0.0283581 | 0.79748454 | 0 |
| LD24904 | FBgn0010348 | Arf79F | -0.028253005 | 0.81857797 | 0 |
| RE21901 | FBgn0035760 | CG8607 | 0.028223484 | 0.79444163 | 0 |

| | | | | | |
|---|---|---|---|---|---|
| GH04611 | FBgn0039155 | Kal1 | -0.028209563 | 0.8308951 | 0 |
| RH39904 | FBgn0032143 | CG4017 | 0.028201733 | 0.8326161 | 0 |
| SD01736 | FBgn0033905 | CG18324 | 0.028143171 | 0.86726761 | 0 |
| RH04014 | FBgn0038820 | CG4000 | 0.028138199 | 0.82785528 | 0 |
| RE27538 | FBgn0035508 | CG15005 | -0.028128705 | 0.79802346 | 0 |
| LD46238 | FBgn0031893 | MICU1 | -0.028116662 | 0.84973261 | 0 |
| RE02644 | FBgn0035360 | CG1246 | -0.028092125 | 0.84784159 | 0 |
| GH09040 | FBgn0032986 | CG3262 | -0.028042899 | 0.80860773 | 0 |
| GH11218 | FBgn0031724 | CG18266 | -0.028041483 | 0.7972368 | 0 |
| GH14143 | FBgn0003965 | v | 0.02801378 | 0.7973342 | 0 |
| GM01318 | FBgn0036318 | Wbp2 | -0.027998031 | 0.80049362 | 0 |
| AT10515 | FBgn0032345 | CG14921 | 0.02798427 | 0.79422756 | 0 |
| GH16758 | FBgn0040985 | CG6115 | 0.02794552 | 0.82643222 | 0 |
| SD03549 | FBgn0000588 | esc | -0.027937531 | 0.79880976 | 0 |
| LD32717 | FBgn0283682 | Ge-1 | 0.027905871 | 0.8122472 | 0 |
| RH29724 | FBgn0031858 | CG17378 | 0.02789113 | 0.82531199 | 0 |
| RH17496 | FBgn0031021 | ND-18 | 0.027853223 | 0.8708313 | 0 |
| LD39025 | FBgn0038126 | CG8483 | -0.027807551 | 0.87924987 | 0 |
| LD42296 | FBgn0035077 | CG9083 | 0.027747178 | 0.79519944 | 0 |
| SD06390 | FBgn0032167 | CG5853 | 0.027736761 | 0.82727028 | 0 |
| RE16081 | FBgn0260987 | vtd | -0.027562789 | 0.85899932 | 0 |
| GH05567 | FBgn0010019 | Cyp4g1 | 0.02754747 | 0.85031271 | 0 |
| RH31634 | FBgn0010388 | Dro | -0.027508349 | 0.79748454 | 0 |
| LD29208 | FBgn0030268 | Klp10A | 0.027418573 | 0.79444163 | 0 |
| GH06403 | FBgn0036020 | CG8336 | 0.027414556 | 0.79476858 | 0 |
| RE60154 | FBgn0038005 | Cyp313a5 | 0.02738642 | 0.91093986 | 0 |
| AT01837 | FBgn0031296 | CG4415 | -0.027353749 | 0.80406448 | 0 |
| RE51073 | FBgn0031257 | CG4133 | -0.027324893 | 0.82458082 | 0 |
| RE56801 | FBgn0010421 | TfIIFbeta | -0.027316581 | 0.81549464 | 0 |
| HL05577 | FBgn0033886 | Rpn13 | -0.027252488 | 0.86623995 | 0 |
| RE07960 | FBgn0047178 | CG32147 | 0.027241998 | 0.84738555 | 0 |
| LP01766 | FBgn0040732 | CG16926 | 0.02723068 | 0.86354745 | 0 |
| LD47433 | FBgn0033802 | CG17724 | 0.027172545 | 0.80908024 | 0 |
| LD47433 | FBgn0053182 | Kdm4B | 0.027172545 | 0.80908024 | 0 |
| LD06574 | FBgn0267330 | KdelR | 0.02713959 | 0.80651168 | 0 |
| LD03464 | FBgn0033713 | CG8841 | 0.027134472 | 0.81546069 | 0 |
| RE54950 | FBgn0020660 | eIF4B | 0.027089285 | 0.8326161 | 0 |
| RE20037 | FBgn0001112 | Gld | -0.027076946 | 0.90380589 | 0 |
| RE57174 | FBgn0036030 | CG6767 | 0.027021589 | 0.8556048 | 0 |
| LD11409 | FBgn0030512 | Nadsyn | -0.026986124 | 0.84784159 | 0 |
| LP01149 | FBgn0031048 | CG12237 | 0.026977213 | 0.8266986 | 0 |
| RH14214 | FBgn0037546 | mAChR-B | 0.026943123 | 0.83784969 | 0 |
| LD25803 | FBgn0030594 | CG9509 | -0.026925957 | 0.80665486 | 0 |
| AT26658 | FBgn0046873 | Pif2 | 0.026924754 | 0.89306065 | 0 |
| AT03665 | FBgn0051088 | CG31088 | 0.026857118 | 0.80860773 | 0 |
| LD40095 | FBgn0046247 | CG5938 | 0.026826203 | 0.79848192 | 0 |
| LD28120 | FBgn0259834 | out | 0.026824716 | 0.81660353 | 0 |
| LD30817 | FBgn0013269 | FK506-bp1 | 0.026811376 | 0.80441002 | 0 |

| | | | | | |
|---|---|---|---|---|---|
| RH31677 | FBgn0034996 | CG13575 | -0.026806123 | 0.82113907 | 0 |
| GH20554 | FBgn0034634 | CG10494 | -0.026783752 | 0.80574102 | 0 |
| AT31162 | FBgn0037827 | CG4073 | -0.026758363 | 0.79897121 | 0 |
| GH03216 | FBgn0051774 | fred | -0.026756518 | 0.87121655 | 0 |
| RE38148 | FBgn0039354 | Lgr3 | -0.026726354 | 0.86557857 | 0 |
| LD46175 | FBgn0037912 | sea | 0.026697721 | 0.80404698 | 0 |
| RE09259 | FBgn0086519 | Cpr47Eg | -0.026696347 | 0.89935525 | 0 |
| RE36503 | FBgn0038198 | Npc2b | 0.02668595 | 0.80074563 | 0 |
| RH38789 | FBgn0041229 | Gr93a | 0.026624491 | 0.80101804 | 0 |
| GH15431 | FBgn0032974 | CG3651 | -0.026624058 | 0.80049362 | 0 |
| LD46766 | FBgn0033728 | Cpr49Ae | 0.026584933 | 0.8255577 | 0 |
| RH66170 | FBgn0029148 | NHP2 | 0.026570062 | 0.86726761 | 0 |
| LD30826 | FBgn0030890 | CG7536 | 0.026562651 | 0.79952723 | 0 |
| AT17751 | FBgn0031171 | CG1801 | 0.026562333 | 0.87621309 | 0 |
| RE13070 | FBgn0002431 | hyd | 0.026539127 | 0.85922221 | 0 |
| RE16431 | FBgn0032480 | Edem2 | -0.026537759 | 0.8099408 | 0 |
| LD30894 | FBgn0262169 | magu | 0.026530758 | 0.8145127 | 0 |
| LD20432 | FBgn0033019 | CG10395 | -0.02646775 | 0.81925609 | 0 |
| LD03569 | FBgn0034654 | eIF3k | 0.026443504 | 0.87197988 | 0 |
| GH27640 | FBgn0034032 | CG8195 | 0.026440784 | 0.79946844 | 0 |
| LD19945 | FBgn0027864 | Ogg1 | 0.026423844 | 0.86623995 | 0 |
| AT13819 | FBgn0031295 | CG4375 | -0.026392056 | 0.85108952 | 0 |
| RE61267 | FBgn0030592 | CG9514 | -0.026348621 | 0.80473998 | 0 |
| GM10731 | FBgn0004876 | cdi | -0.026277723 | 0.8091114 | 0 |
| LP06328 | FBgn0033203 | CG2070 | -0.026266037 | 0.83784969 | 0 |
| RE56868 | FBgn0261387 | CG17528 | -0.026259924 | 0.85150759 | 0 |
| SD04590 | FBgn0033313 | Cirl | -0.02625955 | 0.81546069 | 0 |
| LD29662 | FBgn0036332 | CG11261 | -0.026169468 | 0.86458069 | 0 |
| GM25382 | FBgn0034362 | CG5323 | -0.02612042 | 0.82203015 | 0 |
| RE28995 | FBgn0028411 | Nxt1 | -0.026070029 | 0.87924987 | 0 |
| GH26685 | FBgn0260972 | alc | -0.025974427 | 0.80404698 | 0 |
| LP02515 | FBgn0035951 | CG5068 | 0.025968605 | 0.80681168 | 0 |
| AT24316 | FBgn0000287 | salr | -0.025948767 | 0.82846125 | 0 |
| RH35331 | FBgn0030556 | mRNA-cap | 0.025941081 | 0.80473998 | 0 |
| AT31252 | FBgn0005779 | PpD6 | 0.025912647 | 0.87764308 | 0 |
| GH25957 | FBgn0029663 | CG10804 | 0.025868265 | 0.83334335 | 0 |
| RE02181 | FBgn0265191 | Glycogenin | 0.025824589 | 0.84205895 | 0 |
| RE14171 | FBgn0031141 | CG1304 | -0.025793537 | 0.89655877 | 0 |
| RE14171 | FBgn0011834 | Ser6 | -0.025793537 | 0.89655877 | 0 |
| GH23780 | FBgn0031228 | ND-15 | 0.025772426 | 0.81489401 | 0 |
| LD19782 | FBgn0025381 | rush | 0.025694448 | 0.8136797 | 0 |
| AT29981 | FBgn0034483 | CG16894 | 0.02565866 | 0.82680014 | 0 |
| GH11935 | FBgn0032015 | Ostgamma | -0.025654758 | 0.80787858 | 0 |
| RE55660 | FBgn0259247 | laccase2 | 0.02564839 | 0.84130841 | 0 |
| AT19257 | FBgn0052181 | CG32181 | 0.02564044 | 0.86643241 | 0 |
| RH44664 | FBgn0030733 | UQCR-14 | 0.025638345 | 0.81211294 | 0 |
| RH44664 | FBgn0039576 | UQCR-14L | 0.025638345 | 0.81211294 | 0 |
| AT16405 | FBgn0031785 | CG13991 | -0.025630038 | 0.89776331 | 0 |

| | | | | | |
|---|---|---|---|---|---|
| GH03839 | FBgn0063923 | Kaz1-ORFB | 0.025618469 | 0.88773912 | 0 |
| RE04143 | FBgn0029503 | CHORD | 0.025613684 | 0.83800136 | 0 |
| GH03922 | FBgn0027592 | MED15 | -0.025568122 | 0.87240285 | 0 |
| LD26546 | FBgn0028327 | l(1)G0320 | 0.025550282 | 0.84973261 | 0 |
| LD04457 | FBgn0034321 | CG14502 | 0.025519712 | 0.8628743 | 0 |
| RE25453 | FBgn0037481 | MAGE | -0.025516644 | 0.83321633 | 0 |
| LD21616 | FBgn0000662 | fl(2)d | -0.025510459 | 0.83695778 | 0 |
| RE46684 | FBgn0265901 | asRNA:CR44690 | 0.025491511 | 0.83689195 | 0 |
| GM06875 | FBgn0039056 | CenB1A | 0.025420197 | 0.80787858 | 0 |
| GH14433 | FBgn0031461 | daw | 0.02539811 | 0.8920299 | 0 |
| RE17806 | FBgn0037537 | CG2767 | 0.025387733 | 0.8446935 | 0 |
| SD09476 | FBgn0261445 | sgl | 0.025352747 | 0.80753121 | 0 |
| GH12064 | FBgn0034951 | CG3860 | -0.025323627 | 0.83160436 | 0 |
| LD03592 | FBgn0034913 | Snap29 | 0.025313504 | 0.8266174 | 0 |
| GH02484 | FBgn0039543 | CG12428 | 0.025172605 | 0.8099408 | 0 |
| HL08167 | FBgn0263220 | Hk | -0.025153552 | 0.87759485 | 0 |
| SD15982 | FBgn0260743 | GC1 | -0.025147015 | 0.82525114 | 0 |
| LD29420 | FBgn0031391 | CG11723 | 0.025068888 | 0.81798186 | 0 |
| AT08390 | FBgn0045827 | CG13245 | -0.025012254 | 0.82573313 | 0 |
| AT08390 | FBgn0264364 | CG43816 | -0.025012254 | 0.82573313 | 0 |
| RE13652 | FBgn0031068 | Alr | -0.024969033 | 0.8920299 | 0 |
| AT03323 | FBgn0004581 | bgcn | 0.024849669 | 0.8194291 | 0 |
| RE14272 | FBgn0033658 | Twdlbeta | -0.024771945 | 0.82184061 | 0 |
| GH04919 | FBgn0037372 | CG2091 | 0.024754357 | 0.88356772 | 0 |
| GH03694 | FBgn0028500 | Rich | 0.024732402 | 0.824186 | 0 |
| LD24887 | FBgn0029158 | Las | 0.024727241 | 0.85897093 | 0 |
| LD04986 | FBgn0026401 | Nipped-B | -0.024709629 | 0.87451154 | 0 |
| GH28342 | FBgn0033439 | CG1773 | 0.024693649 | 0.84184002 | 0 |
| GH28342 | FBgn0033440 | CG10459 | 0.024693649 | 0.84184002 | 0 |
| RE35766 | FBgn0038923 | mRpL35 | -0.024681273 | 0.8305917 | 0 |
| LD24950 | FBgn0029857 | wuho | 0.024681012 | 0.87451154 | 0 |
| LD23613 | FBgn0000405 | CycB | -0.024640451 | 0.8338329 | 0 |
| LD27103 | FBgn0022338 | dnk | 0.024622696 | 0.81925609 | 0 |
| AT12494 | FBgn0032511 | ND-B22 | 0.024559319 | 0.82612861 | 0 |
| GH05836 | FBgn0033951 | CG10139 | 0.024540306 | 0.86797103 | 0 |
| GH01330 | FBgn0086384 | Mer | 0.024534408 | 0.8338329 | 0 |
| LD47508 | FBgn0038149 | GILT1 | 0.02451188 | 0.86614799 | 0 |
| LD13909 | FBgn0263398 | CG6364 | 0.024440939 | 0.88035815 | 0 |
| GH14812 | FBgn0000422 | Ddc | 0.024411698 | 0.84350912 | 0 |
| GH08353 | FBgn0036931 | CG14183 | -0.024405205 | 0.82634875 | 0 |
| LD14485 | FBgn0034223 | Tes | 0.024403649 | 0.84401075 | 0 |
| SD04977 | FBgn0025815 | Mcm6 | -0.024397407 | 0.82612861 | 0 |
| AT21585 | FBgn0029710 | CG3568 | -0.024351012 | 0.82815736 | 0 |
| AT01047 | FBgn0038539 | Atg8b | 0.024339953 | 0.82050364 | 0 |
| RE44650 | FBgn0031247 | CG11562 | 0.02426248 | 0.8338329 | 0 |
| LD44762 | FBgn0262518 | Rab8 | 0.024249485 | 0.81961472 | 0 |
| GH25131 | FBgn0038173 | Adgf-C | -0.024242548 | 0.8266986 | 0 |
| AT29875 | FBgn0002579 | RpL36 | 0.024157016 | 0.82785528 | 0 |

| | | | | | |
|---|---|---|---|---|---|
| AT10852 | FBgn0037191 | CG14448 | -0.024018624 | 0.88980781 | 0 |
| SD19739 | FBgn0032196 | CG5708 | 0.023986602 | 0.83239844 | 0 |
| SD19739 | FBgn0285910 | VhaAC39-1 | 0.023986602 | 0.83239844 | 0 |
| GH02992 | FBgn0034693 | CG11073 | -0.023977216 | 0.86354745 | 0 |
| LD28566 | FBgn0014092 | CG4278 | 0.023952387 | 0.84648831 | 0 |
| GH12037 | FBgn0035346 | CG1146 | 0.023934821 | 0.8220452 | 0 |
| LD44138 | FBgn0010877 | l(3)05822 | 0.023874588 | 0.86011954 | 0 |
| GH17339 | FBgn0025382 | Rab27 | 0.02387435 | 0.89606178 | 0 |
| AT13777 | FBgn0033953 | CG12861 | 0.023848118 | 0.9059394 | 0 |
| GH24474 | FBgn0032800 | CG10137 | -0.023821428 | 0.84051293 | 0 |
| LD33980 | FBgn0015772 | Nak | -0.02379843 | 0.84079443 | 0 |
| LD44234 | FBgn0023174 | Prosbeta2 | 0.023640569 | 0.84877535 | 0 |
| RE72625 | FBgn0036660 | CG13025 | 0.023467439 | 0.88563388 | 0 |
| RE31221 | FBgn0034788 | CG13532 | 0.02341312 | 0.88787168 | 0 |
| AT20887 | FBgn0040063 | yip3 | 0.023405611 | 0.88823481 | 0 |
| GH04666 | FBgn0019960 | Mitofilin | 0.023302998 | 0.84051293 | 0 |
| LD14731 | FBgn0040773 | COX7C | -0.023278481 | 0.85283586 | 0 |
| LD41911 | FBgn0266420 | Ote | 0.023188217 | 0.84184002 | 0 |
| GH14206 | FBgn0031585 | CG2955 | 0.023169252 | 0.88547239 | 0 |
| GH28327 | FBgn0259221 | CG42321 | 0.023118051 | 0.83334335 | 0 |
| AT15673 | FBgn0030725 | CG8958 | -0.023083312 | 0.93653996 | 0 |
| GH26638 | FBgn0264502 | CG43901 | 0.022983371 | 0.83321633 | 0 |
| HL01677 | FBgn0031695 | Cyp4ac3 | -0.022911576 | 0.86639186 | 0 |
| GM19936 | FBgn0261790 | SmE | -0.022894083 | 0.83124048 | 0 |
| RH04232 | FBgn0030158 | CG9686 | -0.022886346 | 0.88035815 | 0 |
| LD34980 | FBgn0029943 | Atg5 | 0.022822933 | 0.84789117 | 0 |
| GH02471 | FBgn0263111 | cac | -0.022768958 | 0.9216045 | 0 |
| GH01674 | FBgn0040349 | CG3699 | -0.022735017 | 0.83872919 | 0 |
| SD04201 | FBgn0034063 | CG8389 | 0.022713289 | 0.85256792 | 0 |
| LD23217 | FBgn0030065 | CG12075 | -0.022710718 | 0.90120063 | 0 |
| GH04831 | FBgn0037251 | CG9804 | -0.022710433 | 0.85635658 | 0 |
| LD08534 | FBgn0250837 | dUTPase | 0.022696005 | 0.86838856 | 0 |
| HL01444 | FBgn0028543 | NimB2 | -0.022672033 | 0.90702594 | 0 |
| RH01479 | FBgn0033208 | mRpL52 | -0.022587145 | 0.87621309 | 0 |
| RE66017 | FBgn0263663 | lncRNA:CR43654 | -0.022548736 | 0.82934131 | 0 |
| GH08937 | FBgn0030853 | ND-24 | -0.022524256 | 0.8652082 | 0 |
| LD27861 | FBgn0003612 | Su(var)2-10 | -0.022510459 | 0.85533784 | 0 |
| RE58063 | FBgn0036396 | CG17359 | -0.022504971 | 0.84346583 | 0 |
| GH05923 | FBgn0021800 | Reph | 0.022503426 | 0.83847249 | 0 |
| RE47096 | FBgn0000015 | Abd-B | 0.022401344 | 0.83906973 | 0 |
| RE36848 | FBgn0032346 | Csl4 | -0.022356557 | 0.86797103 | 0 |
| RH22958 | FBgn0033032 | kune | 0.022337793 | 0.84353178 | 0 |
| LD06293 | FBgn0027095 | Manf | 0.022279227 | 0.88563388 | 0 |
| HL08076 | FBgn0262508 | CG43078 | 0.022187526 | 0.84184002 | 0 |
| GH12052 | FBgn0053556 | form3 | 0.022138265 | 0.84878506 | 0 |
| GM01970 | FBgn0037815 | Rrp46 | -0.022134614 | 0.88982638 | 0 |
| RE04467 | FBgn0030763 | CG9782 | 0.022124053 | 0.8523069 | 0 |
| RH30783 | FBgn0032719 | CG17321 | -0.02205737 | 0.89660208 | 0 |

| | | | | | |
|---|---|---|---|---|---|
| SD12145 | FBgn0267350 | PI4KIIIalpha | 0.02205153 | 0.86454134 | 0 |
| GH28656 | FBgn0035770 | pst | -0.022036256 | 0.85068505 | 0 |
| RE51929 | FBgn0028645 | beat-Ib | 0.021915573 | 0.88982638 | 0 |
| LD36129 | FBgn0038582 | CG7988 | -0.021901373 | 0.84109298 | 0 |
| AT18706 | FBgn0052601 | betaNACtes3 | -0.021832564 | 0.91118289 | 0 |
| AT18706 | FBgn0052598 | betaNACtes6 | -0.021832564 | 0.91118289 | 0 |
| AT18706 | FBgn0030538 | betaNACtes1 | -0.021832564 | 0.91118289 | 0 |
| RE50009 | FBgn0037342 | CG2931 | -0.021759617 | 0.87291713 | 0 |
| SD13982 | FBgn0032213 | CG5390 | -0.021700763 | 0.86072829 | 0 |
| LP12385 | FBgn0030494 | Cpr12A | -0.021648649 | 0.93279605 | 0 |
| RH49308 | FBgn0085468 | ND-MWFE | -0.021620061 | 0.84111405 | 0 |
| RE03721 | FBgn0029665 | CG14270 | 0.021578311 | 0.87924987 | 0 |
| RE56519 | FBgn0000246 | c(3)G | 0.021562699 | 0.87103703 | 0 |
| HL07933 | FBgn0266451 | fau | 0.021525267 | 0.88274711 | 0 |
| LD11455 | FBgn0034422 | CG7137 | 0.021523366 | 0.84785235 | 0 |
| RE12054 | FBgn0265003 | koi | 0.021509664 | 0.85150759 | 0 |
| RE42418 | FBgn0004636 | Rap1 | -0.021473534 | 0.92048566 | 0 |
| GH11985 | FBgn0031697 | CG14024 | -0.021468291 | 0.87147571 | 0 |
| GM04934 | FBgn0040285 | Scamp | 0.021453556 | 0.85698892 | 0 |
| RE18374 | FBgn0051005 | qless | -0.021398068 | 0.84249406 | 0 |
| AT28291 | FBgn0032366 | CG14930 | 0.021358944 | 0.92554929 | 0 |
| GH09630 | FBgn0032512 | Bdp1 | 0.021347133 | 0.89362315 | 0 |
| LD28980 | FBgn0039828 | CG1542 | 0.021310833 | 0.90535831 | 0 |
| LD32311 | FBgn0266450 | Kr-h1 | -0.021267845 | 0.92650613 | 0 |
| LD32311 | FBgn0266445 | CG45075 | -0.021267845 | 0.92650613 | 0 |
| LD07912 | FBgn0022981 | rpk | -0.021267745 | 0.84048484 | 0 |
| RE35358 | FBgn0003065 | CG2150 | 0.021265065 | 0.89362315 | 0 |
| RE40670 | FBgn0032218 | CG5381 | 0.021210025 | 0.90197438 | 0 |
| LD27862 | FBgn0037027 | HIPP1 | -0.021203517 | 0.86011954 | 0 |
| RE17884 | FBgn0261953 | TfAP-2 | -0.021199403 | 0.88507259 | 0 |
| LD38718 | FBgn0004106 | Cdk1 | -0.021171689 | 0.87197988 | 0 |
| LD23375 | FBgn0037875 | ZnT86D | -0.021099047 | 0.8491373 | 0 |
| RH31685 | FBgn0051321 | CG31321 | -0.021087573 | 0.85332858 | 0 |
| SD07678 | FBgn0040388 | boi | 0.021078526 | 0.85102165 | 0 |
| RE11532 | FBgn0030291 | CG1738 | 0.021076034 | 0.84140794 | 0 |
| GH24644 | FBgn0033309 | Lnpk | 0.021063441 | 0.8446935 | 0 |
| RE19416 | FBgn0034001 | mRpL41 | -0.021055174 | 0.85478697 | 0 |
| LD20087 | FBgn0031344 | CG7420 | -0.021033377 | 0.88231166 | 0 |
| LD15196 | FBgn0031904 | CG5149 | -0.021025529 | 0.8571093 | 0 |
| AT26187 | FBgn0030120 | CG17440 | 0.021014835 | 0.87924987 | 0 |
| SD02996 | FBgn0263706 | CG43658 | -0.021011515 | 0.86557857 | 0 |
| LP07407 | FBgn0034659 | CG4021 | -0.020963707 | 0.88844362 | 0 |
| GH15894 | FBgn0035090 | CG2736 | 0.020931213 | 0.88549613 | 0 |
| AT26524 | FBgn0031235 | CG13693 | -0.020926587 | 0.91093986 | 0 |
| LP03531 | FBgn0052284 | CG32284 | -0.020912589 | 0.89877582 | 0 |
| RE09178 | FBgn0037895 | CG6723 | -0.020886782 | 0.84434985 | 0 |
| LD23107 | FBgn0000146 | aub | 0.020753693 | 0.87047102 | 0 |
| RE35124 | FBgn0028955 | CG8788 | -0.020708539 | 0.84651118 | 0 |

| | | | | | |
|---|---|---|---|---|---|
| RE35124 | FBgn0265313 | CG44286 | -0.020708539 | 0.84651118 | 0 |
| LD32220 | FBgn0036330 | CG11263 | -0.02067999 | 0.89129152 | 0 |
| LD12677 | FBgn0033010 | Atf6 | -0.020645612 | 0.86572919 | 0 |
| RE29909 | FBgn0264384 | lncRNA:CR43836 | -0.02062369 | 0.87621309 | 0 |
| SD01117 | FBgn0034791 | EMC8-9 | -0.0206233 | 0.85150759 | 0 |
| RE23984 | FBgn0011584 | Trp1 | -0.020616567 | 0.88246686 | 0 |
| AT01605 | FBgn0037826 | CG14689 | -0.020608108 | 0.92511011 | 0 |
| RE34245 | FBgn0024184 | unc-4 | 0.020601776 | 0.85628447 | 0 |
| GH01346 | FBgn0038032 | CG10096 | 0.020543633 | 0.84643351 | 0 |
| GH15456 | FBgn0003116 | pn | 0.020452329 | 0.85154456 | 0 |
| LP02768 | FBgn0025469 | slv | 0.020414538 | 0.91093986 | 0 |
| RE02065 | FBgn0032247 | CG5188 | -0.020388033 | 0.86445543 | 0 |
| RE01590 | FBgn0032168 | CG13126 | 0.020381584 | 0.8545044 | 0 |
| RE11751 | FBgn0030654 | CG15643 | 0.02036121 | 0.84785235 | 0 |
| GH13631 | FBgn0003093 | Pkc98E | 0.020322713 | 0.85446511 | 0 |
| RE03009 | FBgn0035610 | Lkr | -0.020320973 | 0.87129521 | 0 |
| GH11272 | FBgn0030555 | Fbxl4 | 0.020206474 | 0.86989242 | 0 |
| GH20168 | FBgn0038349 | AOX3 | -0.020112741 | 0.85928831 | 0 |
| AT01267 | FBgn0286199 | shps | 0.020086758 | 0.89810972 | 0 |
| RE14222 | FBgn0039818 | CG11318 | 0.019997795 | 0.90665555 | 0 |
| LD35003 | FBgn0038772 | mdlc | 0.019984095 | 0.86283366 | 0 |
| LP06762 | FBgn0031832 | CG9596 | 0.019973551 | 0.85070543 | 0 |
| GH17085 | FBgn0040345 | CG3708 | 0.01987848 | 0.86797103 | 0 |
| AT26578 | FBgn0034066 | CG8397 | 0.019857955 | 0.87986369 | 0 |
| LD16402 | FBgn0026380 | Prosbeta3 | -0.019800769 | 0.87970829 | 0 |
| LD08427 | FBgn0024811 | Crk | 0.019771019 | 0.90989009 | 0 |
| GH06387 | FBgn0035540 | Syx17 | -0.019753318 | 0.85626769 | 0 |
| LP11089 | FBgn0000153 | b | 0.019747937 | 0.90097006 | 0 |
| RE63456 | FBgn0039406 | RpL34a | -0.019690244 | 0.87896939 | 0 |
| LD21957 | FBgn0038471 | CG5220 | -0.019636641 | 0.89283296 | 0 |
| GH02938 | FBgn0030260 | CG1537 | -0.019573877 | 0.88994695 | 0 |
| GM14421 | FBgn0030757 | CG9902 | 0.019561574 | 0.85652135 | 0 |
| RE23308 | FBgn0003345 | sd | -0.019546653 | 0.90142322 | 0 |
| SD08852 | FBgn0033766 | Nup188 | 0.019449374 | 0.88205076 | 0 |
| LD42267 | FBgn0032685 | CG10211 | 0.019410568 | 0.87610397 | 0 |
| LD34542 | FBgn0039698 | CG7789 | -0.019361002 | 0.88458768 | 0 |
| GH14073 | FBgn0035710 | SP1173 | 0.019303687 | 0.93143597 | 0 |
| RE49422 | FBgn0036278 | CrzR | -0.019252104 | 0.89067908 | 0 |
| GH10077 | FBgn0036725 | CG18265 | 0.01918459 | 0.887763 | 0 |
| RE13129 | FBgn0030914 | CG6106 | -0.019129132 | 0.87908346 | 0 |
| LD36705 | FBgn0024432 | Dlc90F | -0.019120707 | 0.89306065 | 0 |
| LD17717 | FBgn0004828 | His3.3B | 0.019058432 | 0.86149564 | 0 |
| GH01304 | FBgn0034716 | Oatp58Dc | -0.019031603 | 0.91237593 | 0 |
| RE67638 | FBgn0038234 | mRpL11 | 0.019007516 | 0.8782569 | 0 |
| RH17440 | FBgn0029765 | CG16756 | 0.019000342 | 0.89349898 | 0 |
| LD36454 | FBgn0038424 | CG17565 | 0.018999767 | 0.86181064 | 0 |
| LD40977 | FBgn0026749 | Yippee | 0.018991345 | 0.90385307 | 0 |
| AT12418 | FBgn0034793 | asrij | 0.018947831 | 0.87282882 | 0 |

| | | | | | |
|---|---|---|---|---|---|
| LP10490 | FBgn0053523 | CG33523 | 0.018925047 | 0.87859568 | 0 |
| LD24878 | FBgn0035204 | CG2277 | 0.018881058 | 0.87469163 | 0 |
| RE52350 | FBgn0036556 | CG5830 | 0.018822367 | 0.86445543 | 0 |
| GH03570 | FBgn0032223 | GATAd | -0.018777508 | 0.87388986 | 0 |
| LD38389 | FBgn0023175 | Prosalpha7 | 0.018755341 | 0.88982638 | 0 |
| GH25012 | FBgn0038354 | CG5404 | -0.018736483 | 0.87147571 | 0 |
| AT19020 | FBgn0031301 | CG14339 | 0.018711753 | 0.91991168 | 0 |
| SD17676 | FBgn0039788 | Rpt6R | -0.018671358 | 0.86907327 | 0 |
| SD02805 | FBgn0050372 | Asap | -0.018646599 | 0.86557857 | 0 |
| LD37196 | FBgn0087021 | Spc25 | -0.018572524 | 0.87621309 | 0 |
| RE71584 | FBgn0260444 | Not10 | 0.018569962 | 0.86445543 | 0 |
| GH08867 | FBgn0002936 | ninaA | -0.018549368 | 0.86623995 | 0 |
| GH12267 | FBgn0039089 | beat-IV | 0.018465427 | 0.93279605 | 0 |
| AT01661 | FBgn0038266 | CG3610 | -0.018429526 | 0.86512977 | 0 |
| LD21829 | FBgn0086605 | CG9853 | -0.01841088 | 0.90665555 | 0 |
| AT09142 | FBgn0032968 | CG11634 | -0.018400266 | 0.92030066 | 0 |
| RE29446 | FBgn0033523 | CG12895 | 0.018389301 | 0.88787168 | 0 |
| AT29184 | FBgn0051875 | CG31875 | -0.018380582 | 0.88534492 | 0 |
| LD36501 | FBgn0030854 | CG8289 | -0.018338126 | 0.86800214 | 0 |
| LD30416 | FBgn0284251 | l(2)05287 | 0.018306581 | 0.88917806 | 0 |
| GH26248 | FBgn0031406 | Send1 | -0.018270711 | 0.92951171 | 0 |
| RE17665 | FBgn0030675 | CG8191 | -0.018232033 | 0.91314897 | 0 |
| GH04753 | FBgn0027590 | GstE12 | 0.01822554 | 0.90385307 | 0 |
| RE27904 | FBgn0047135 | CG32276 | -0.01821328 | 0.89950889 | 0 |
| RE47936 | FBgn0040022 | Set1 | 0.018159997 | 0.87146057 | 0 |
| LD26873 | FBgn0032979 | Clamp | -0.018120157 | 0.87986369 | 0 |
| SD07707 | FBgn0033482 | CG1371 | 0.018034138 | 0.87329598 | 0 |
| RE16553 | FBgn0015591 | AstA | -0.017995226 | 0.9059394 | 0 |
| RH01675 | FBgn0029932 | CG4607 | -0.017966574 | 0.87451154 | 0 |
| SD04017 | FBgn0033434 | CG1902 | 0.017957668 | 0.87424896 | 0 |
| RE60735 | FBgn0030847 | CG12991 | 0.017940477 | 0.92030066 | 0 |
| RE55559 | FBgn0036062 | CG6685 | -0.017922469 | 0.90197438 | 0 |
| RE23862 | FBgn0261394 | Prosalpha3 | 0.017848683 | 0.87257179 | 0 |
| GH14380 | FBgn0029810 | CG12239 | 0.01783334 | 0.9059394 | 0 |
| GH03581 | FBgn0030350 | SelG | 0.017831031 | 0.87388986 | 0 |
| LD25345 | FBgn0027070 | CG17322 | 0.017807988 | 0.89283296 | 0 |
| GH08548 | FBgn0038224 | ATPsynE | 0.017744374 | 0.88780495 | 0 |
| LD35644 | FBgn0037262 | MED31 | 0.017743363 | 0.8920299 | 0 |
| RE26969 | FBgn0030485 | CG1998 | 0.017737268 | 0.89979935 | 0 |
| GM20805 | FBgn0000115 | Arl1 | 0.017718805 | 0.87373732 | 0 |
| GM01751 | FBgn0034734 | CG4554 | 0.017693874 | 0.90542596 | 0 |
| LP03039 | FBgn0039229 | Saf-B | -0.017671673 | 0.94993242 | 0 |
| RH03863 | FBgn0033777 | CG17574 | -0.017561316 | 0.87197988 | 0 |
| LD32934 | FBgn0000499 | dsh | -0.017537237 | 0.92898521 | 0 |
| AT27178 | FBgn0039246 | CG10845 | -0.017532414 | 0.89457804 | 0 |
| GM02110 | FBgn0278608 | Dsp1 | 0.017508004 | 0.94082661 | 0 |
| LD32631 | FBgn0086350 | tef | 0.017506552 | 0.87924987 | 0 |
| GH27226 | FBgn0053987 | lncRNA:CR33987 | 0.017475249 | 0.88740312 | 0 |

| | | | | | |
|---|---|---|---|---|---|
| SD03277 | FBgn0030674 | CG8184 | 0.017463136 | 0.88066118 | 0 |
| RE64145 | FBgn0024196 | robl | -0.017213818 | 0.87885599 | 0 |
| GH10666 | FBgn0037023 | CG3698 | 0.017196934 | 0.8845372 | 0 |
| LD41384 | FBgn0029152 | Mkrn1 | 0.017167552 | 0.88309811 | 0 |
| AT06885 | FBgn0250821 | CG14644 | 0.017147489 | 0.87424896 | 0 |
| LD27406 | FBgn0033918 | CG8531 | 0.017131425 | 0.88097648 | 0 |
| LD25830 | FBgn0039272 | CG11836 | -0.017080606 | 0.90245143 | 0 |
| RE56061 | FBgn0051176 | CG31176 | 0.017077455 | 0.88699056 | 0 |
| SD10722 | FBgn0035375 | Pgant6 | -0.017074775 | 0.89140016 | 0 |
| LD16419 | FBgn0036921 | RhoGDI | 0.017013408 | 0.91394786 | 0 |
| GH16988 | FBgn0264087 | Slob | -0.016944882 | 0.91703282 | 0 |
| RE47547 | FBgn0051431 | CG31431 | -0.016941389 | 0.91092064 | 0 |
| AT30546 | FBgn0039214 | puf | -0.016917217 | 0.9355655 | 0 |
| LD07633 | FBgn0046706 | Haspin | -0.016908507 | 0.90530603 | 0 |
| RE11719 | FBgn0029681 | CG15239 | 0.01689272 | 0.89776331 | 0 |
| SD17342 | FBgn0033550 | CG12341 | 0.016892487 | 0.88066118 | 0 |
| RE12030 | FBgn0053145 | GalT1 | -0.016788693 | 0.87764308 | 0 |
| LD07775 | FBgn0264922 | smt3 | -0.016681968 | 0.91093986 | 0 |
| SD03652 | FBgn0250850 | rig | 0.016666194 | 0.87658653 | 0 |
| LD15753 | FBgn0043069 | MESK4 | 0.016649579 | 0.91445332 | 0 |
| LD15753 | FBgn0051274 | CG31274 | 0.016649579 | 0.91445332 | 0 |
| LP06628 | FBgn0038484 | CG5246 | 0.016589875 | 0.91093986 | 0 |
| GH15945 | FBgn0039184 | CG6432 | 0.016543238 | 0.88482046 | 0 |
| AT13663 | FBgn0029747 | CG5062 | -0.016491812 | 0.9338671 | 0 |
| RE43153 | FBgn0029804 | CG3097 | -0.0164806 | 0.9533955 | 0 |
| LD38689 | FBgn0033225 | TTLL12 | 0.016429894 | 0.91400719 | 0 |
| LD40196 | FBgn0260639 | gammaTub23C | 0.016414564 | 0.88024811 | 0 |
| RE08019 | FBgn0026417 | Hus1-like | 0.016400817 | 0.90385307 | 0 |
| LD14453 | FBgn0004811 | fs(2)ltoPP43 | -0.01636407 | 0.89810972 | 0 |
| GH09147 | FBgn0027558 | Pgant3 | -0.016354041 | 0.91181756 | 0 |
| RH49123 | FBgn0027790 | GV1 | -0.016323029 | 0.88790715 | 0 |
| RE03006 | FBgn0027259 | Kmn1 | 0.016302492 | 0.89306065 | 0 |
| HL05736 | FBgn0039734 | Tace | -0.016253513 | 0.92408276 | 0 |
| RE37929 | FBgn0039852 | nyo | 0.016226904 | 0.89306065 | 0 |
| LD27655 | FBgn0033021 | CG10417 | 0.016226194 | 0.8920299 | 0 |
| GH04581 | FBgn0040754 | CG17059 | 0.016224158 | 0.88284578 | 0 |
| RE55959 | FBgn0039632 | Cul5 | 0.016205346 | 0.88891487 | 0 |
| GH22307 | FBgn0053202 | dpr11 | -0.016165401 | 0.89810972 | 0 |
| RE33457 | FBgn0035742 | CG8629 | -0.016155195 | 0.92176728 | 0 |
| RE22192 | FBgn0039238 | CG7016 | -0.01613738 | 0.90245143 | 0 |
| RH43162 | FBgn0051973 | Cda5 | -0.016129075 | 0.90523818 | 0 |
| LD46538 | FBgn0036039 | Naa60 | 0.016122946 | 0.90197438 | 0 |
| LD22374 | FBgn0033005 | CG3107 | -0.016110362 | 0.90266065 | 0 |
| RH16169 | FBgn0042201 | Nplp3 | -0.016070244 | 0.92461708 | 0 |
| LD11861 | FBgn0033076 | CG15233 | -0.016062984 | 0.89776331 | 0 |
| LD40493 | FBgn0038805 | TFAM | 0.015990397 | 0.88791063 | 0 |
| GH09326 | FBgn0051183 | CG31183 | 0.015940708 | 0.90466018 | 0 |
| AT19640 | FBgn0033579 | CG13229 | 0.015822466 | 0.90663518 | 0 |

| | | | | | |
|---|---|---|---|---|---|
| RE08808 | FBgn0033942 | Cpr51A | -0.01580958 | 0.88948914 | 0 |
| LD43101 | FBgn0000319 | Chc | -0.015759889 | 0.91917651 | 0 |
| LD16125 | FBgn0000577 | en | -0.015759437 | 0.91381667 | 0 |
| RH69723 | FBgn0036926 | CG7646 | -0.015755531 | 0.93016829 | 0 |
| GH06435 | FBgn0034144 | CG5089 | 0.015509631 | 0.96039957 | 0 |
| GH07181 | FBgn0036367 | CG10116 | 0.015508675 | 0.91168789 | 0 |
| GH28769 | FBgn0035857 | CG8006 | 0.015458176 | 0.93425703 | 0 |
| GH12469 | FBgn0051233 | CG31233 | -0.015408645 | 0.90197438 | 0 |
| RH13070 | FBgn0034361 | mRpS28 | -0.01533339 | 0.93279605 | 0 |
| RE41484 | FBgn0034553 | CG9993 | -0.015281715 | 0.93461496 | 0 |
| RE07460 | FBgn0003060 | CG9757 | 0.015274285 | 0.88980781 | 0 |
| RE25483 | FBgn0033519 | CG11825 | 0.015245934 | 0.90646665 | 0 |
| LD44960 | FBgn0015766 | Msr-110 | -0.015238949 | 0.91314897 | 0 |
| LD44960 | FBgn0065097 | snmRNA:357 | -0.015238949 | 0.91314897 | 0 |
| LD25772 | FBgn0036515 | AIMP2 | -0.01522633 | 0.91768741 | 0 |
| RH45818 | FBgn0026562 | SPARC | 0.015220003 | 0.93639037 | 0 |
| GH27420 | FBgn0032484 | kek4 | -0.015208605 | 0.9511587 | 0 |
| GH26628 | FBgn0035649 | CG10483 | 0.015176895 | 0.89554415 | 0 |
| SD10607 | FBgn0022960 | vimar | 0.015096476 | 0.90097006 | 0 |
| LD35371 | FBgn0032236 | mRpS7 | 0.01505032 | 0.90665555 | 0 |
| GH26280 | FBgn0085479 | CG34450 | 0.015034813 | 0.91296713 | 0 |
| GH18152 | FBgn0020416 | Idgf1 | 0.015032404 | 0.92652628 | 0 |
| RE58902 | FBgn0031926 | CG6739 | 0.015019957 | 0.88979506 | 0 |
| SD01790 | FBgn0038318 | CG6236 | -0.015016394 | 0.89058653 | 0 |
| GH14445 | FBgn0037950 | HisCl1 | -0.014974489 | 0.90266065 | 0 |
| GH03029 | FBgn0035528 | CG15012 | -0.014944424 | 0.92825215 | 0 |
| RE05231 | FBgn0050197 | CG30197 | -0.014940875 | 0.91108137 | 0 |
| RE69157 | FBgn0264815 | Pde1c | -0.014902979 | 0.89606178 | 0 |
| RH60875 | FBgn0039759 | CG9733 | 0.014850759 | 0.89760459 | 0 |
| RE41571 | FBgn0031000 | Muc18B | -0.014837622 | 0.90865737 | 0 |
| RH40237 | FBgn0260861 | Trs23 | -0.014837586 | 0.92012928 | 0 |
| RE49527 | FBgn0039886 | CG2003 | 0.014821475 | 0.89810972 | 0 |
| LD34214 | FBgn0047199 | CG31517 | -0.014815705 | 0.91447405 | 0 |
| LD17927 | FBgn0030177 | CG2972 | -0.014756567 | 0.90097006 | 0 |
| RE24163 | FBgn0036395 | CG17361 | 0.014736231 | 0.89638651 | 0 |
| GM01305 | FBgn0037848 | Tsp86D | 0.014701615 | 0.89457804 | 0 |
| LD04844 | FBgn0030309 | CG1572 | 0.014658696 | 0.92825215 | 0 |
| GH20077 | FBgn0033971 | CG10209 | 0.014626302 | 0.90487612 | 0 |
| AT04844 | FBgn0053322 | CG33322 | 0.014600693 | 0.93167131 | 0 |
| LD47289 | FBgn0034529 | FAM21 | -0.014536107 | 0.90197438 | 0 |
| LD04449 | FBgn0033686 | Hen1 | 0.014409489 | 0.94962083 | 0 |
| AT01413 | FBgn0039010 | CG4907 | -0.014399337 | 0.93925707 | 0 |
| RE35745 | FBgn0283680 | IP3K2 | 0.014371055 | 0.91400719 | 0 |
| AT09608 | FBgn0050491 | CG30491 | 0.014165198 | 0.9682755 | 0 |
| LD44370 | FBgn0032929 | Mcm10 | -0.014092282 | 0.91093986 | 0 |
| RE57126 | FBgn0034755 | CG3746 | 0.014047699 | 0.90679166 | 0 |
| SD01203 | FBgn0022288 | l(2)09851 | -0.0140299 | 0.91503649 | 0 |
| LD24056 | FBgn0011760 | ctp | -0.0140177 | 0.91093986 | 0 |

| | | | | | |
|---|---|---|---|---|---|
| LD08709 | FBgn0035627 | Sse | 0.014006429 | 0.90542596 | 0 |
| RE17113 | FBgn0035856 | CG13679 | 0.014004898 | 0.91237593 | 0 |
| LD03462 | FBgn0083992 | Mkp | -0.014003641 | 0.90542596 | 0 |
| RH40165 | FBgn0035287 | CG13937 | -0.013963876 | 0.90079274 | 0 |
| RH49358 | FBgn0001216 | Hsc70-1 | -0.013838107 | 0.92137792 | 0 |
| GH15507 | FBgn0044826 | Pak3 | 0.013837291 | 0.91037648 | 0 |
| LP09811 | FBgn0025620 | CG13360 | -0.013799246 | 0.92313526 | 0 |
| RE01069 | FBgn0027052 | STUB1 | -0.013794404 | 0.91466152 | 0 |
| AT10229 | FBgn0033238 | azot | 0.013719393 | 0.96374114 | 0 |
| AT10229 | FBgn0050378 | CG30378 | 0.013719393 | 0.96374114 | 0 |
| RE07883 | FBgn0034261 | HPS4 | -0.013707613 | 0.90542596 | 0 |
| LD34846 | FBgn0259168 | mnb | -0.013660247 | 0.92774252 | 0 |
| LD14505 | FBgn0022238 | lolal | 0.013646681 | 0.9059394 | 0 |
| RH16027 | FBgn0259219 | CG42319 | -0.013616029 | 0.90646665 | 0 |
| RE57135 | FBgn0029588 | CG14798 | 0.013600901 | 0.91280507 | 0 |
| HL05903 | FBgn0000303 | ChAT | 0.013594805 | 0.91108137 | 0 |
| AT10439 | FBgn0030370 | Uch-L5R | -0.013501779 | 0.91118289 | 0 |
| GH08582 | FBgn0004583 | ex | 0.013498122 | 0.92176728 | 0 |
| LD30423 | FBgn0010215 | alpha-Cat | -0.013497034 | 0.92243354 | 0 |
| HL01392 | FBgn0266446 | CG45076 | -0.013493876 | 0.91867235 | 0 |
| LP04610 | FBgn0004511 | dy | -0.013454501 | 0.93354727 | 0 |
| RE60081 | FBgn0034821 | CG9876 | -0.013415467 | 0.91917651 | 0 |
| LP02196 | FBgn0026141 | Cdlc2 | 0.013340095 | 0.92951171 | 0 |
| AT15424 | FBgn0035888 | CG7120 | -0.013329463 | 0.9258529 | 0 |
| AT27322 | FBgn0031440 | CG15395 | 0.013327978 | 0.93056384 | 0 |
| LD35950 | FBgn0039627 | CG11837 | -0.013314599 | 0.90530603 | 0 |
| GH20973 | FBgn0004919 | gol | 0.013233015 | 0.94546589 | 0 |
| LP11145 | FBgn0035791 | CG8539 | 0.013183456 | 0.96067641 | 0 |
| AT09955 | FBgn0032481 | CG16972 | 0.013127419 | 0.90663518 | 0 |
| SD01985 | FBgn0261113 | Xrp1 | -0.013117119 | 0.91085515 | 0 |
| GH22991 | FBgn0263118 | tx | 0.01310919 | 0.95999575 | 0 |
| GH05754 | FBgn0035076 | Ance-5 | -0.013079178 | 0.93639037 | 0 |
| LD17129 | FBgn0000183 | BicD | 0.013056702 | 0.91400719 | 0 |
| GH06474 | FBgn0034664 | CG4377 | -0.013005071 | 0.93565977 | 0 |
| LD40134 | FBgn0025574 | Pli | -0.01299738 | 0.91093986 | 0 |
| GH11924 | FBgn0033794 | CG13326 | 0.012949082 | 0.94929057 | 0 |
| LD37974 | FBgn0032052 | PIG-U | 0.012918017 | 0.91093986 | 0 |
| LD16949 | FBgn0000147 | aurA | -0.012867381 | 0.91093986 | 0 |
| RE44811 | FBgn0031655 | Marcal1 | 0.012839006 | 0.91400719 | 0 |
| GH18995 | FBgn0034937 | fzr2 | 0.012827675 | 0.93565977 | 0 |
| AT11889 | FBgn0267689 | CG46025 | 0.012808401 | 0.94410075 | 0 |
| AT10577 | FBgn0000489 | Pka-C3 | -0.01277406 | 0.9576828 | 0 |
| RE35169 | FBgn0030261 | CG15203 | -0.012741021 | 0.95824709 | 0 |
| RH14578 | FBgn0022027 | Vps25 | -0.012736565 | 0.91093986 | 0 |
| LD25271 | FBgn0030944 | CG6617 | 0.012714723 | 0.9089514 | 0 |
| GH05991 | FBgn0030624 | CG9106 | -0.01260997 | 0.91506865 | 0 |
| AT26626 | FBgn0035704 | Vps8 | -0.01256293 | 0.91917651 | 0 |
| GH07575 | FBgn0034296 | CG10912 | -0.01255707 | 0.9232851 | 0 |

| | | | | | |
|---|---|---|---|---|---|
| AT05866 | FBgn0037296 | Prosbeta2R2 | -0.012547416 | 0.91466152 | 0 |
| RE18252 | FBgn0032587 | CG5953 | -0.012340217 | 0.93475125 | 0 |
| AT16007 | FBgn0052081 | CG32081 | 0.012330419 | 0.93016829 | 0 |
| LP02570 | FBgn0033968 | hui | 0.012282145 | 0.91466152 | 0 |
| AT11468 | FBgn0038361 | CG9589 | 0.012279539 | 0.94623838 | 0 |
| SD08134 | FBgn0032698 | CG10336 | 0.012226688 | 0.91181756 | 0 |
| LP11564 | FBgn0051638 | CG31638 | 0.012191304 | 0.93275981 | 0 |
| GH16240 | FBgn0020304 | drongo | -0.012188086 | 0.93247171 | 0 |
| GH28556 | FBgn0259112 | CR42254 | 0.012171949 | 0.91326811 | 0 |
| LD28626 | FBgn0037377 | CG1218 | 0.012140115 | 0.91308313 | 0 |
| GH14208 | FBgn0036260 | Rh7 | -0.012080092 | 0.94887773 | 0 |
| AT04270 | FBgn0038565 | CG7794 | 0.012011373 | 0.94684132 | 0 |
| RE35907 | FBgn0033608 | CG13220 | 0.011965181 | 0.93279605 | 0 |
| LD43687 | FBgn0013563 | Pex1 | -0.011965014 | 0.93167131 | 0 |
| LP04804 | FBgn0033048 | CG7881 | 0.011900418 | 0.91314897 | 0 |
| RE63660 | FBgn0038842 | hdly | -0.011879694 | 0.92963642 | 0 |
| RH18149 | FBgn0038460 | CG18622 | 0.011852714 | 0.91980156 | 0 |
| GH16403 | FBgn0263241 | Mocs1 | -0.011842395 | 0.9217333 | 0 |
| AT30660 | FBgn0051093 | CG31093 | 0.011838443 | 0.94362349 | 0 |
| AT07710 | FBgn0037547 | CG7910 | -0.011809872 | 0.91768741 | 0 |
| GH12638 | FBgn0035985 | Cpr67B | -0.011803435 | 0.96155876 | 0 |
| GH17657 | FBgn0034127 | CG7848 | 0.011775327 | 0.92412728 | 0 |
| RE31829 | FBgn0025808 | Rad17 | -0.011774303 | 0.93064929 | 0 |
| GH15686 | FBgn0036319 | Ent3 | 0.011671496 | 0.92487351 | 0 |
| RE10167 | FBgn0036470 | EAChm | 0.0116587 | 0.97075205 | 0 |
| LD21503 | FBgn0027527 | Osi6 | 0.01159755 | 0.98964917 | 0 |
| LD28404 | FBgn0038586 | CG7168 | 0.011553928 | 0.93279605 | 0 |
| RE29729 | FBgn0004915 | TfIIB | -0.011546736 | 0.92527126 | 0 |
| LD22238 | FBgn0067864 | Patj | -0.011528007 | 0.92552391 | 0 |
| GM01939 | FBgn0030744 | CG9992 | -0.011527929 | 0.92843205 | 0 |
| RH08189 | FBgn0037712 | CG16789 | 0.011501375 | 0.92408276 | 0 |
| RH07540 | FBgn0034743 | RpS16 | 0.011494257 | 0.92408276 | 0 |
| GH02310 | FBgn0034718 | wdp | -0.011476565 | 0.92374488 | 0 |
| LD44962 | FBgn0003479 | spn-A | 0.011464808 | 0.91644566 | 0 |
| LD16326 | FBgn0285950 | RpL19 | 0.011387875 | 0.92413751 | 0 |
| LD21956 | FBgn0283473 | S6KL | -0.011383745 | 0.93549306 | 0 |
| SD26038 | FBgn0038302 | CG4210 | 0.011327692 | 0.93179591 | 0 |
| LD29238 | FBgn0033015 | d4 | -0.011126044 | 0.9326671 | 0 |
| LD37668 | FBgn0032204 | CG4953 | -0.011053772 | 0.93465877 | 0 |
| LD04582 | FBgn0003159 | CG2841 | 0.011029261 | 0.93461496 | 0 |
| GH28095 | FBgn0033544 | CG7220 | -0.010932705 | 0.94887773 | 0 |
| RE44586 | FBgn0033259 | CG11210 | -0.010920119 | 0.92708882 | 0 |
| RE42071 | FBgn0036133 | CG7638 | -0.01091451 | 0.93465877 | 0 |
| RE74918 | FBgn0037424 | Osi15 | -0.010890032 | 0.98323589 | 0 |
| RH05746 | FBgn0035280 | Cpr62Bb | 0.010879342 | 0.96493746 | 0 |
| GH03576 | FBgn0083068 | CG33947 | -0.010759492 | 0.92408276 | 0 |
| LD21841 | FBgn0037980 | DCAF12 | 0.010730227 | 0.92444647 | 0 |
| SD22791 | FBgn0025621 | CG16989 | -0.010728168 | 0.94690947 | 0 |

| | | | | | |
|---|---|---|---|---|---|
| LD03714 | FBgn0005411 | U2af50 | -0.010718623 | 0.92708882 | 0 |
| LP02895 | FBgn0052475 | mthl8 | -0.010712017 | 0.95471326 | 0 |
| AT12292 | FBgn0029812 | Prosbeta2R1 | 0.010681301 | 0.92793545 | 0 |
| GH06923 | FBgn0032341 | Reps | -0.010668032 | 0.92657171 | 0 |
| RE04768 | FBgn0034636 | twz | -0.010661265 | 0.9258529 | 0 |
| SD09178 | FBgn0024329 | Mekk1 | -0.010589527 | 0.92843205 | 0 |
| GH07785 | FBgn0035420 | CG14967 | 0.010511192 | 0.93461496 | 0 |
| AT17875 | FBgn0037276 | CG17387 | -0.010466081 | 0.9438365 | 0 |
| AT09606 | FBgn0036785 | CG13700 | 0.010465086 | 0.94690947 | 0 |
| SD23366 | FBgn0033052 | SCAP | 0.010456181 | 0.93565977 | 0 |
| RE11449 | FBgn0065080 | snoRNA:Me18S-A1374 | 0.010427488 | 0.9348892 | 0 |
| RH02303 | FBgn0037635 | CG9837 | 0.010426467 | 0.9537815 | 0 |
| LD25288 | FBgn0031398 | CG10880 | 0.01035699 | 0.9326671 | 0 |
| LD39062 | FBgn0038917 | CG6678 | -0.010354392 | 0.92652628 | 0 |
| RE15265 | FBgn0036272 | CG4300 | -0.010260584 | 0.95825654 | 0 |
| LD21069 | FBgn0034083 | lbk | 0.010232617 | 0.94887773 | 0 |
| RE13557 | FBgn0085354 | CG34325 | 0.010172713 | 0.94690947 | 0 |
| RE61847 | FBgn0044020 | Roc2 | 0.010117349 | 0.93055481 | 0 |
| LP01070 | FBgn0039754 | CG9747 | -0.010104046 | 0.95560094 | 0 |
| RE53289 | FBgn0031692 | TpnC25D | 0.01009837 | 0.95100888 | 0 |
| RE32113 | FBgn0037069 | Cpr78Cc | -0.009988639 | 0.9440305 | 0 |
| SD22984 | FBgn0011224 | heph | 0.009978671 | 0.95999575 | 0 |
| RE40268 | FBgn0284225 | CG46309 | 0.00994766 | 0.95096416 | 0 |
| SD06355 | FBgn0028572 | qtc | 0.00992801 | 0.96171549 | 0 |
| LD21741 | FBgn0035639 | kri | 0.009878813 | 0.93016829 | 0 |
| LD40846 | FBgn0031054 | Ssu72 | 0.009858303 | 0.93016829 | 0 |
| AT03371 | FBgn0035724 | CG10064 | 0.009857954 | 0.95059213 | 0 |
| LD03255 | FBgn0039846 | PNPase | -0.00981222 | 0.93476615 | 0 |
| SD03319 | FBgn0034936 | CG2970 | -0.009791955 | 0.9326671 | 0 |
| AT04875 | FBgn0267795 | Frl | -0.009751502 | 0.96067641 | 0 |
| SD07148 | FBgn0028687 | Rpt1 | -0.009725994 | 0.93722323 | 0 |
| RE02160 | FBgn0038916 | dnd | -0.009713374 | 0.9592585 | 0 |
| LD36273 | FBgn0030026 | sni | -0.009695854 | 0.93960811 | 0 |
| LD05321 | FBgn0032513 | CG6565 | -0.009669765 | 0.9326671 | 0 |
| GH27269 | FBgn0037252 | CG14650 | -0.009651439 | 0.93653996 | 0 |
| GH09754 | FBgn0038179 | CG9312 | -0.009643212 | 0.93599797 | 0 |
| AT20029 | FBgn0033868 | S-Lap7 | 0.009581016 | 0.9533955 | 0 |
| RE25364 | FBgn0038526 | CG14327 | -0.009520057 | 0.94987734 | 0 |
| GH17980 | FBgn0037975 | CG3397 | 0.009514103 | 0.95999575 | 0 |
| GH17980 | FBgn0037974 | CG12224 | 0.009514103 | 0.95999575 | 0 |
| LD08815 | FBgn0035959 | CG4911 | -0.009501805 | 0.93639037 | 0 |
| RE40372 | FBgn0051446 | CG31446 | 0.009481994 | 0.96095906 | 0 |
| GH12970 | FBgn0036671 | CG9951 | 0.009448606 | 0.96274656 | 0 |
| AT17606 | FBgn0034671 | CG13494 | -0.009416816 | 0.9342149 | 0 |
| AT09528 | FBgn0030092 | fh | -0.009403708 | 0.93461496 | 0 |
| LD44361 | FBgn0015602 | BEAF-32 | -0.009402126 | 0.93461496 | 0 |
| RE08173 | FBgn0031975 | Tg | 0.009391204 | 0.93799411 | 0 |
| GH01139 | FBgn0035134 | CG1231 | -0.009390938 | 0.9648049 | 0 |

| | | | | | |
|---|---|---|---|---|---|
| LP12023 | FBgn0063083 | lncRNA:CR33948 | -0.00925587 | 0.96274656 | 0 |
| LD16758 | FBgn0261403 | sxc | 0.009247917 | 0.94688691 | 0 |
| LD15349 | FBgn0032455 | Pih1D1 | 0.009231996 | 0.93657815 | 0 |
| SD09672 | FBgn0026370 | SRPK | 0.009159183 | 0.93657815 | 0 |
| GH16433 | FBgn0032135 | GlcAT-S | 0.009158426 | 0.93516513 | 0 |
| RE26879 | FBgn0034499 | Cpr56F | -0.009094843 | 0.94929057 | 0 |
| SD10476 | FBgn0265190 | PIG-S | 0.009032969 | 0.9361651 | 0 |
| RE42104 | FBgn0004197 | Ser | 0.008970804 | 0.97037855 | 0 |
| GH01528 | FBgn0023489 | Pph13 | -0.008956613 | 0.94663726 | 0 |
| RE01730 | FBgn0034709 | Swim | -0.008949913 | 0.93527105 | 0 |
| RE63043 | FBgn0003575 | su(sable) | 0.008890704 | 0.93992437 | 0 |
| LD04388 | FBgn0036734 | CG7564 | 0.008796054 | 0.9438365 | 0 |
| GH04442 | FBgn0032428 | CG6405 | -0.008731813 | 0.94986929 | 0 |
| SD06557 | FBgn0036668 | Zcchc7 | 0.008716165 | 0.93657815 | 0 |
| LD31673 | FBgn0021760 | chb | -0.008712951 | 0.95811951 | 0 |
| GH25737 | FBgn0029666 | CG10803 | -0.008688704 | 0.94473307 | 0 |
| GH26392 | FBgn0052278 | CG32278 | 0.008652021 | 0.9576828 | 0 |
| LD23674 | FBgn0030136 | RpS28b | 0.008646608 | 0.94915336 | 0 |
| LD33852 | FBgn0004797 | mdy | -0.008636445 | 0.94379781 | 0 |
| GH19329 | FBgn0002563 | Lsp1beta | 0.008625451 | 0.93931801 | 0 |
| LD11760 | FBgn0037757 | CG8516 | 0.008592408 | 0.94158624 | 0 |
| LD21102 | FBgn0026315 | Ugt35a | -0.008575193 | 0.94887773 | 0 |
| AT20190 | FBgn0038210 | CG3199 | -0.008376805 | 0.96364089 | 0 |
| RE57556 | FBgn0032524 | CG9267 | -0.008338514 | 0.96017135 | 0 |
| RE09075 | FBgn0033378 | tsu | -0.008308188 | 0.9508919 | 0 |
| GH20109 | FBgn0035781 | CG8560 | -0.00828469 | 0.96071608 | 0 |
| RE56714 | FBgn0038260 | CG14855 | -0.00821882 | 0.94626611 | 0 |
| HL07966 | FBgn0004623 | Gbeta76C | 0.008210863 | 0.94276854 | 0 |
| GH17414 | FBgn0266346 | CngB | -0.008194068 | 0.94529701 | 0 |
| GH26207 | FBgn0035574 | RhoGEF64C | -0.008130345 | 0.96637136 | 0 |
| GH14469 | FBgn0259936 | Uhg3 | 0.008108125 | 0.94542114 | 0 |
| GH14469 | FBgn0082936 | snoRNA:Me28S-C2645c | 0.008108125 | 0.94542114 | 0 |
| LD07767 | FBgn0014906 | Hydr2 | 0.008094072 | 0.94887773 | 0 |
| RH03891 | FBgn0035817 | CG7409 | 0.008092835 | 0.9698582 | 0 |
| GH03057 | FBgn0024889 | Kap-alpha1 | -0.008041193 | 0.94405174 | 0 |
| LD46912 | FBgn0032017 | CG7810 | 0.007986624 | 0.94410075 | 0 |
| LD16783 | FBgn0040373 | CG3038 | -0.007918147 | 0.96317502 | 0 |
| RH22541 | FBgn0031299 | CG4629 | 0.007881087 | 0.96067641 | 0 |
| RE04975 | FBgn0051716 | Cnot4 | -0.007871324 | 0.94790481 | 0 |
| RE27818 | FBgn0039469 | TwdlC | -0.007830369 | 0.94757494 | 0 |
| RE01373 | FBgn0028697 | RpL15 | -0.00775688 | 0.95900786 | 0 |
| LD09531 | FBgn0260771 | Larp7 | -0.007728906 | 0.9511587 | 0 |
| AT04622 | FBgn0083956 | CG34120 | 0.007688181 | 0.95864157 | 0 |
| RE69204 | FBgn0051646 | DIP-theta | -0.007660548 | 0.95100888 | 0 |
| RH18307 | FBgn0038680 | Cyp12a5 | 0.007627178 | 0.95655667 | 0 |
| GH10882 | FBgn0032768 | CG17564 | -0.007540706 | 0.96736754 | 0 |
| SD10043 | FBgn0259876 | Cap-G | -0.007507758 | 0.96019973 | 0 |
| RE37682 | FBgn0038564 | CG7785 | 0.007434399 | 0.94899393 | 0 |

| | | | | | |
|---|---|---|---|---|---|
| RE65233 | FBgn0263216 | CG43386 | 0.007431517 | 0.95100888 | 0 |
| RE28175 | FBgn0004066 | Prosalpha4 | 0.007405388 | 0.96020624 | 0 |
| LD13253 | FBgn0016031 | lama | 0.007395619 | 0.94887773 | 0 |
| LD10695 | FBgn0024728 | Slip1 | 0.007370882 | 0.9592585 | 0 |
| GH09594 | FBgn0037215 | beta-Man | -0.007327083 | 0.95820869 | 0 |
| RH22042 | FBgn0039109 | CG10365 | -0.007160424 | 0.96374114 | 0 |
| LD44179 | FBgn0036639 | CG4229 | 0.007154121 | 0.96120581 | 0 |
| LP04475 | FBgn0038110 | CG8031 | -0.007036816 | 0.95822927 | 0 |
| SD01502 | FBgn0266369 | Mtp | -0.006959954 | 0.96039957 | 0 |
| RE66446 | FBgn0036510 | SCCRO | 0.006955484 | 0.9576828 | 0 |
| LD46811 | FBgn0038313 | CG4338 | -0.006953877 | 0.96039957 | 0 |
| RE35159 | FBgn0037165 | CG11437 | -0.006939467 | 0.97314841 | 0 |
| GH05625 | FBgn0004237 | Hrb87F | 0.006928369 | 0.9558823 | 0 |
| GH04057 | FBgn0030562 | CG9400 | 0.006827595 | 0.96964342 | 0 |
| GH18521 | FBgn0031526 | CG8838 | 0.006805466 | 0.97012297 | 0 |
| GH11706 | FBgn0038208 | CG14355 | -0.006751984 | 0.96841558 | 0 |
| LD37859 | FBgn0039300 | RpS27 | -0.006716708 | 0.95900786 | 0 |
| AT04306 | FBgn0051912 | asRNA:CR31912 | 0.00666122 | 0.9698582 | 0 |
| GH15938 | FBgn0040907 | mRpL33 | 0.006628246 | 0.95655667 | 0 |
| RE54230 | FBgn0050339 | CG30339 | 0.006541677 | 0.9576828 | 0 |
| RH44244 | FBgn0028879 | CG15270 | 0.006494747 | 0.96286126 | 0 |
| RH03940 | FBgn0002626 | RpL32 | 0.006335953 | 0.97122792 | 0 |
| RE69232 | FBgn0035587 | Gdap1 | 0.006309325 | 0.96329968 | 0 |
| SD08157 | FBgn0243512 | puc | 0.006272829 | 0.96401742 | 0 |
| RE08556 | FBgn0032069 | LManVI | -0.006269747 | 0.96236709 | 0 |
| LD11831 | FBgn0266376 | lncRNA:CR45018 | 0.006232743 | 0.96325604 | 0 |
| AT05691 | FBgn0052119 | CG32119 | 0.00622207 | 0.96564525 | 0 |
| RE52725 | FBgn0020269 | mspo | 0.006199358 | 0.96736754 | 0 |
| GH08772 | FBgn0031857 | LUBEL | 0.006170116 | 0.95862087 | 0 |
| HL01263 | FBgn0025637 | SkpA | -0.00613412 | 0.97347383 | 0 |
| LD33266 | FBgn0037116 | Als2 | 0.006109886 | 0.96096814 | 0 |
| GH06385 | FBgn0267794 | lncRNA:CR43174 | -0.006061311 | 0.96374114 | 0 |
| RH05091 | FBgn0034331 | CG15067 | 0.006035278 | 0.96158752 | 0 |
| RE41180 | FBgn0267728 | otk2 | 0.006015199 | 0.96120581 | 0 |
| LD22567 | FBgn0050404 | Tango11 | -0.006004516 | 0.96014221 | 0 |
| AT09129 | FBgn0038438 | Der-2 | 0.005954066 | 0.96317502 | 0 |
| RE30881 | FBgn0033749 | achi | -0.005948194 | 0.96699821 | 0 |
| LD44304 | FBgn0010328 | woc | -0.005938663 | 0.96120581 | 0 |
| LD30634 | FBgn0037671 | ATP6AP2 | -0.005787528 | 0.97243084 | 0 |
| RH16768 | FBgn0038425 | CG14881 | -0.005774557 | 0.96552578 | 0 |
| LD47553 | FBgn0038965 | mats | 0.005767037 | 0.96067641 | 0 |
| GH27293 | FBgn0032399 | CG6785 | -0.005694692 | 0.96699821 | 0 |
| SD02622 | FBgn0037082 | CG5664 | -0.005651583 | 0.96699821 | 0 |
| GH16413 | FBgn0028902 | Tektin-A | 0.005631124 | 0.96736754 | 0 |
| RE07994 | FBgn0039868 | CG11563 | 0.005619401 | 0.96849092 | 0 |
| SD24044 | FBgn0036581 | MED10 | 0.005611013 | 0.96736754 | 0 |
| RE71227 | FBgn0030863 | CG8188 | -0.005574641 | 0.96318994 | 0 |
| GH15157 | FBgn0031047 | Rcd-1 | 0.005564665 | 0.96701931 | 0 |

| | | | | | |
|---|---|---|---|---|---|
| AT17867 | FBgn0039532 | Mtl | -0.005502036 | 0.96584927 | 0 |
| SD04658 | FBgn0038524 | sll | -0.005344583 | 0.96493746 | 0 |
| RE60135 | FBgn0035675 | CG6610 | -0.005341886 | 0.97174109 | 0 |
| SD08549 | FBgn0039741 | CG7943 | -0.005327811 | 0.96364089 | 0 |
| GH26786 | FBgn0005633 | fln | 0.005311753 | 0.97289186 | 0 |
| LP03924 | FBgn0010295 | ng3 | -0.005274299 | 0.97594539 | 0 |
| AT14148 | FBgn0037298 | CG2604 | 0.005195894 | 0.97122792 | 0 |
| SD07295 | FBgn0032365 | CG14929 | -0.005158186 | 0.96720208 | 0 |
| AT03044 | FBgn0031304 | CG4552 | 0.005107673 | 0.96564525 | 0 |
| LD26327 | FBgn0036948 | CG7298 | -0.004889551 | 0.97122792 | 0 |
| LD37956 | FBgn0000667 | Actn | 0.004860511 | 0.9698582 | 0 |
| RE31721 | FBgn0032700 | CG10338 | 0.004851163 | 0.96701931 | 0 |
| RE04135 | FBgn0259977 | Tdc1 | -0.004849646 | 0.97072101 | 0 |
| RE56733 | FBgn0052230 | ND-MLRQ | -0.004809964 | 0.96913306 | 0 |
| SD05646 | FBgn0024314 | Plap | 0.004803577 | 0.9698582 | 0 |
| AT31406 | FBgn0031772 | CG13994 | -0.004799708 | 0.97075205 | 0 |
| AT13486 | FBgn0034601 | CG4286 | -0.004760403 | 0.98114755 | 0 |
| AT07592 | FBgn0262002 | CG42820 | -0.004730861 | 0.97289186 | 0 |
| RH59310 | FBgn0029975 | spidey | -0.004673924 | 0.9698582 | 0 |
| GH15883 | FBgn0042710 | Hex-t2 | -0.00464056 | 0.97858942 | 0 |
| GH15448 | FBgn0026261 | bonsai | 0.004633951 | 0.97075205 | 0 |
| LP02703 | FBgn0035550 | CG11349 | 0.00462921 | 0.9698582 | 0 |
| LD07634 | FBgn0015925 | csul | -0.004621959 | 0.97075205 | 0 |
| RH17470 | FBgn0039031 | Gbp3 | 0.004599556 | 0.96829644 | 0 |
| RE14560 | FBgn0002775 | msl-3 | 0.00456863 | 0.9698582 | 0 |
| RE02417 | FBgn0024491 | Bin1 | -0.004519165 | 0.97672685 | 0 |
| GH08303 | FBgn0036764 | CG5535 | 0.004465429 | 0.9767595 | 0 |
| LD08331 | FBgn0001316 | klar | -0.004414594 | 0.97780493 | 0 |
| SD01508 | FBgn0031011 | CG8034 | -0.00441145 | 0.97319961 | 0 |
| RE12073 | FBgn0283438 | CG46280 | -0.00435375 | 0.97159758 | 0 |
| AT27686 | FBgn0034667 | comr | -0.004259416 | 0.98250772 | 0 |
| LD05703 | FBgn0260749 | Utx | 0.004255152 | 0.97495035 | 0 |
| LD32260 | FBgn0260464 | CG9288 | -0.004240365 | 0.97672685 | 0 |
| LD32260 | FBgn0259721 | CG42375 | -0.004240365 | 0.97672685 | 0 |
| GH12537 | FBgn0035187 | Trh | -0.004174651 | 0.9855849 | 0 |
| LD28458 | FBgn0051365 | CG31365 | -0.004042857 | 0.97381238 | 0 |
| AT17820 | FBgn0053233 | CG33233 | 0.003907585 | 0.98226035 | 0 |
| AT27946 | FBgn0267457 | asRNA:CR45807 | -0.003906648 | 0.97804058 | 0 |
| AT10032 | FBgn0035823 | eIF4E5 | 0.003881242 | 0.98573652 | 0 |
| RE63320 | FBgn0038246 | CG14853 | 0.003879951 | 0.97594539 | 0 |
| GH20501 | FBgn0033778 | Balat | 0.003833882 | 0.99118771 | 0 |
| GH09296 | FBgn0027657 | glob1 | -0.003816548 | 0.98254152 | 0 |
| LD36177 | FBgn0032956 | Cul2 | -0.003785383 | 0.97755313 | 0 |
| GH13256 | FBgn0025352 | Thiolase | 0.003722837 | 0.98639087 | 0 |
| GH06717 | FBgn0028491 | CG2930 | -0.003633164 | 0.98409905 | 0 |
| RH56938 | FBgn0261556 | CG42674 | -0.003625593 | 0.97594539 | 0 |
| LD42233 | FBgn0038890 | CG7956 | 0.00362361 | 0.98113385 | 0 |
| GH26112 | FBgn0051528 | CG31528 | 0.003612617 | 0.97594539 | 0 |

| | | | | | |
|---|---|---|---|---|---|
| GH24371 | FBgn0051343 | CG31343 | -0.003557133 | 0.9853898 | 0 |
| LD26077 | FBgn0004177 | mts | 0.003543463 | 0.98094689 | 0 |
| LD27045 | FBgn0260049 | flr | 0.003522323 | 0.97648231 | 0 |
| GH16729 | FBgn0031683 | CG4230 | -0.003509298 | 0.97617258 | 0 |
| AT21693 | FBgn0033286 | CG2127 | -0.003503132 | 0.98399426 | 0 |
| SD03464 | FBgn0023506 | Es2 | -0.003380756 | 0.98077731 | 0 |
| GH13116 | FBgn0040502 | CG8343 | -0.003354083 | 0.98409905 | 0 |
| RE52036 | FBgn0001235 | hth | 0.00330966 | 0.98076518 | 0 |
| AT13591 | FBgn0033056 | CG7856 | 0.003265633 | 0.98396235 | 0 |
| AT11204 | FBgn0003388 | shd | 0.003241517 | 0.98741361 | 0 |
| LD21568 | FBgn0001197 | His2Av | 0.003222914 | 0.98113385 | 0 |
| RH73910 | FBgn0032202 | REPTOR-BP | 0.003166057 | 0.98571096 | 0 |
| LD28933 | FBgn0261722 | fwe | -0.003084391 | 0.98400803 | 0 |
| AT26093 | FBgn0250845 | CG1288 | -0.003079876 | 0.98908848 | 0 |
| SD16302 | FBgn0024994 | Ugalt | 0.003024444 | 0.98627542 | 0 |
| RE16127 | FBgn0037627 | CG13318 | -0.003006564 | 0.98713005 | 0 |
| RE73323 | FBgn0033615 | CG7741 | 0.002961238 | 0.98710517 | 0 |
| GH03623 | FBgn0036619 | Cpr72Ec | -0.002944148 | 0.98323589 | 0 |
| RE35104 | FBgn0030670 | Pis | -0.002915402 | 0.98258487 | 0 |
| GH27858 | FBgn0031240 | CG3345 | 0.00289552 | 0.98920864 | 0 |
| GH13495 | FBgn0039311 | CG10513 | 0.002886788 | 0.98174073 | 0 |
| SD23839 | FBgn0025638 | Roc1a | 0.002883508 | 0.98466611 | 0 |
| LD01461 | FBgn0036505 | CG7945 | 0.002821099 | 0.98409905 | 0 |
| SD07884 | FBgn0027490 | D12 | 0.002809793 | 0.98323589 | 0 |
| RH72196 | FBgn0032596 | Prosbeta4 | -0.002759186 | 0.98323589 | 0 |
| GM21019 | FBgn0030057 | Ppt1 | -0.002730667 | 0.98573652 | 0 |
| GH22474 | FBgn0027087 | HisRS | 0.002628212 | 0.98409905 | 0 |
| RH42867 | FBgn0031417 | CG3597 | -0.002624679 | 0.9932331 | 0 |
| RE58921 | FBgn0004924 | Top1 | 0.002585645 | 0.98881297 | 0 |
| LD21410 | FBgn0028664 | VhaM9.7-c | 0.002583108 | 0.98920864 | 0 |
| LD31675 | FBgn0032378 | CycY | -0.002575068 | 0.98323589 | 0 |
| RE51688 | FBgn0000097 | aop | -0.002379305 | 0.98611641 | 0 |
| RE36938 | FBgn0038700 | CG3734 | -0.002345548 | 0.98964917 | 0 |
| GM02257 | FBgn0034879 | Rrp4 | -0.002325883 | 0.98950196 | 0 |
| GH12561 | FBgn0025681 | CG3558 | -0.002204202 | 0.98573652 | 0 |
| AT22982 | FBgn0034175 | ste24b | 0.002202192 | 0.98964917 | 0 |
| GM25741 | FBgn0030460 | CG2453 | 0.002190684 | 0.98835285 | 0 |
| RE34924 | FBgn0033571 | Rpb5 | -0.002176408 | 0.98964917 | 0 |
| RE23690 | FBgn0033921 | tej | 0.002129991 | 0.98901247 | 0 |
| RH23455 | FBgn0033204 | CG2065 | -0.002041975 | 0.98920864 | 0 |
| AT11715 | FBgn0033757 | muskelin | -0.002014028 | 0.98734307 | 0 |
| RE59279 | FBgn0035996 | CG3448 | 0.001996329 | 0.98936378 | 0 |
| GH13883 | FBgn0029896 | CG3168 | -0.001959449 | 0.99091247 | 0 |
| GM13912 | FBgn0034050 | bug | 0.00195776 | 0.98964917 | 0 |
| RE37350 | FBgn0262115 | CG17683 | -0.001951652 | 0.99091247 | 0 |
| LD37221 | FBgn0031060 | CG14231 | -0.001939821 | 0.9890083 | 0 |
| GH07387 | FBgn0029720 | CG3009 | -0.001935327 | 0.98901247 | 0 |
| RE03738 | FBgn0038720 | CG6231 | -0.001932327 | 0.98832288 | 0 |

| | | | | | |
|---|---|---|---|---|---|
| GH11496 | FBgn0037552 | CG7800 | -0.001884053 | 0.99087184 | 0 |
| GM23968 | FBgn0000426 | SmF | 0.001865312 | 0.99091247 | 0 |
| RH61753 | FBgn0069923 | CG41128 | -0.001859541 | 0.99006081 | 0 |
| LP01144 | FBgn0035234 | CG12003 | 0.001859466 | 0.99091247 | 0 |
| AT13732 | FBgn0260459 | CG14106 | -0.001836314 | 0.99419077 | 0 |
| LD24014 | FBgn0046222 | Wdr33 | 0.001822966 | 0.99064621 | 0 |
| AT12574 | FBgn0001280 | janA | 0.001743213 | 0.99079786 | 0 |
| GM04716 | FBgn0050343 | CG30343 | -0.001708711 | 0.9918048 | 0 |
| RE65335 | FBgn0029531 | CG13362 | -0.001672405 | 0.99468458 | 0 |
| RE32029 | FBgn0051634 | Oatp26F | 0.001657827 | 0.99424478 | 0 |
| AT08601 | FBgn0036905 | ms(3)76Ca | -0.001641114 | 0.99468458 | 0 |
| AT08601 | FBgn0260434 | CG42529 | -0.001641114 | 0.99468458 | 0 |
| GM02712 | FBgn0039274 | CG11920 | -0.001620503 | 0.99189341 | 0 |
| RH31195 | FBgn0031558 | CG16704 | 0.001561845 | 0.99131493 | 0 |
| RE21802 | FBgn0030151 | CG1354 | -0.001559695 | 0.99189341 | 0 |
| RE11685 | FBgn0033128 | Tsp42Eg | 0.001513474 | 0.99248365 | 0 |
| GH16993 | FBgn0034883 | Eglp2 | 0.001503043 | 0.99374612 | 0 |
| RH07250 | FBgn0038980 | Octbeta1R | -0.001501687 | 0.99223541 | 0 |
| RE41410 | FBgn0036214 | CG7264 | -0.001490226 | 0.9934543 | 0 |
| GM01569 | FBgn0038167 | Lkb1 | 0.001447453 | 0.99190607 | 0 |
| GH22857 | FBgn0033812 | Pex13 | -0.001422589 | 0.99189341 | 0 |
| GH13361 | FBgn0030999 | Mur18B | -0.001415527 | 0.99331572 | 0 |
| SD09786 | FBgn0050456 | CG30456 | 0.001301387 | 0.99223541 | 0 |
| LD22317 | FBgn0033381 | GstE13 | -0.001279928 | 0.99248365 | 0 |
| GH05851 | FBgn0265597 | rad | 0.001233405 | 0.99421248 | 0 |
| HL04910 | FBgn0033633 | Smyd4-3 | -0.001224575 | 0.99325731 | 0 |
| GH20963 | FBgn0036403 | CG6661 | -0.001201956 | 0.99300737 | 0 |
| GH02759 | FBgn0010213 | Sod2 | 0.001183982 | 0.993873 | 0 |
| RE13542 | FBgn0036765 | CG7408 | 0.001137393 | 0.99300737 | 0 |
| GH14121 | FBgn0037610 | CG8043 | -0.00107678 | 0.99298349 | 0 |
| GH14121 | FBgn0037611 | CG11755 | -0.00107678 | 0.99298349 | 0 |
| SD05886 | FBgn0035437 | Strip | 0.001031282 | 0.99325731 | 0 |
| LD31235 | FBgn0036908 | TORIP | 0.001000185 | 0.99681678 | 0 |
| RE28911 | FBgn0266492 | SclB | -0.000991209 | 0.99350269 | 0 |
| RE28911 | FBgn0266491 | SclA | -0.000991209 | 0.99350269 | 0 |
| GH04243 | FBgn0031589 | CG3714 | 0.000961312 | 0.993873 | 0 |
| LP04011 | FBgn0004167 | kst | 0.000906184 | 0.99469675 | 0 |
| SD22208 | FBgn0003274 | RpLP2 | -0.000904365 | 0.9943569 | 0 |
| LD34777 | FBgn0015247 | Diap2 | -0.000859683 | 0.99421248 | 0 |
| RE03646 | FBgn0020307 | dve | 0.000841035 | 0.99694366 | 0 |
| LD16921 | FBgn0026160 | tna | -0.000783832 | 0.99759549 | 0 |
| LD48009 | FBgn0037370 | CG1236 | 0.000578005 | 0.99681678 | 0 |
| HL01494 | FBgn0035392 | CG1271 | -0.000575191 | 0.99756219 | 0 |
| GH26991 | FBgn0030028 | Corp | 0.000571166 | 0.99759549 | 0 |
| RH36417 | FBgn0050361 | mtt | 0.000490285 | 0.99729549 | 0 |
| LD37279 | FBgn0037606 | CG8032 | -0.00046514 | 0.99694366 | 0 |
| GH10451 | FBgn0011581 | Ms | 0.00042074 | 0.99759549 | 0 |
| GM13604 | FBgn0086134 | Prosalpha2 | -0.000420695 | 0.99759549 | 0 |

| | | | | | |
|---|---|---|---|---|---|
| GM05459 | FBgn0025639 | Hmt4-20 | 0.000395055 | 0.99759549 | 0 |
| RE70149 | FBgn0039188 | Golgin84 | 0.000381377 | 0.99759549 | 0 |
| GH01354 | FBgn0087039 | Sbp2 | 0.000309053 | 0.99809856 | 0 |
| AT07524 | FBgn0036161 | CG7551 | -0.000300675 | 0.99908841 | 0 |
| RE54776 | FBgn0037849 | CG4596 | 0.000265305 | 0.99921695 | 0 |
| RE26660 | FBgn0035625 | Blimp-1 | -0.00019366 | 0.99908841 | 0 |
| LD19812 | FBgn0263251 | vnc | 0.000124398 | 0.99927567 | 0 |
| LD19284 | FBgn0022710 | Ac13E | -0.000115535 | 0.99965972 | 0 |
| RE29988 | FBgn0052576 | CG32576 | 9.43158E-05 | 0.99965049 | 0 |
| RE17452 | FBgn0032938 | CG8671 | 7.62812E-05 | 0.99965972 | 0 |
| GH02426 | FBgn0053111 | CG33111 | -7.31185E-05 | 0.99965972 | 0 |
| LD31204 | FBgn0035866 | Arl5 | -5.40822E-05 | 0.99965972 | 0 |
| GH08677 | FBgn0005322 | nmd | -4.25703E-05 | 0.99966894 | 0 |

**Table S2A**. Flies used in this study.

| Gene name | CG number | Transgene | Source* (stock number) |
|---|---|---|---|
| Reporter and GAL4 | | | |
| sGMCA | | P{w[+mC]=sGMCA} | Kiehart et al., 2000 |
| sChMCA | | P{w[+mC]=sChMCA} | BDSC(35519, 35520, 35521, ref) |
| tGPH (PIP3 reporter) | | P{w[+mC]=tGPH} | BDSC(8164) |
| matalpha4-GAL-VP16 | | P{w[+mC]=matalpha4-GAL-VP16}V2H | BDSC (7062) |
| matalpha4-GAL-VP16 | | P{w[+mC]=matalpha4-GAL-VP16}V37 | BDSC (7063) |
| His2Av-mRFP1 and MCP-EGFP | | P{w[+mC]=His2Av-mRFP1}II.2; P{w[+mC]=nos-MCP.EGFP}2 | BDSC(60340) |
| MS2-lacZ | | P{hbP2-MS2-lacZ} | BDSC(60338) |
| RNAi and mutant | | | |
| ImpL2 | CG15009 | P{y[+t7.7] v[+t1.8]=TRiP.HMC03863} | BDSC (55855) |
| Link | CG13333 | P{y[+t7.7] v[+t1.8]=TRiP.HMJ21610} | NIG-FLY (HMJ21610) |
| Thor | CG8846 | P{y[+t7.7] v[+t1.8]=TRiP.GL01034} | BDSC (36815) |
| Dtg | CG6234 | P{y[+t7.7] v[+t1.8]=TRiP.HMC06344} | BDSC (67242) |
| Geko | CG13695 | 13695R-3 | NIG-FLY (13695R-3) |
| CG43693 | CG43693 | P{y[+t7.7] v[+t1.8]=TRiP.HMJ22481} | BDSC (60341) |
| Egfr | CG10079 | P{y[+t7.7] v[+t1.8]=TRiP.HMS05003} | BDSC (60012) |
| Ama | CG2198 | P{y[+t7.7] v[+t1.8]=TRiP.HMS00297} | BDSC (33416) |
| l(3)neo38 | CG6930 | P{y[+t7.7] v[+t1.8]=TRiP.HMC03292} | BDSC (51515) |
| Dpn | CG8704 | P{y[+t7.7] v[+t1.8]=TRiP.HMC03154} | BDSC (51440) |
| Gp150 | CG5820 | P{y[+t7.7] v[+t1.8]=TRiP.HMS00395} | BDSC (32400) |
| Nullo | CG14426 | P{y[+t7.7] v[+t1.8]=TRiP.HMS02297} | BDSC (41905) |
| Danr | CG13651 | P{y[+t7.7] v[+t1.8]=TRiP.HMC05981} | BDSC (65125) |
| Jbug | CG30092 | P{y[+t7.7] v[+t1.8]=TRiP.HMS01990} | BDSC (39070) |
| P32 | CG6459 | P{y[+t7.7] v[+t1.8]=TRiP.HMS01059} | BDSC (34585) |
| CG1598 | CG1598 | P{y[+t7.7] v[+t1.8]=TRiP.HMC05685} | BDSC (64650) |
| CG3652 | CG3652 | P{y[+t7.7] v[+t1.8]=TRiP.HMJ23083} | NIG-FLY (HMJ23083) |
| CG4960 | CG4960 | 4960R-1 | NIG-FLY (4960R-1) |
| ReepB | CG8331 | P{y[+t7.7] v[+t1.8]=TRiP.HMJ23959} | BDSC (62476) |
| CG14545 | CG14545 | P{y[+t7.7] v[+t1.8]=TRiP.GL00470} | BDSC (35625) |
| Pex11 | CG8315 | P{y[+t7.7] v[+t1.8]=TRiP.HMS02576} | BDSC (42883) |
| Rpl23 | CG3661 | 3661R-2 | NIG-FLY (3661R-2) |
| Cyp6a9 | CG10246 | P{y[+t7.7] v[+t1.8]=TRiP.HMC06193} | BDSC (65927) |
| Exu | CG8994 | P{y[+t7.7] v[+t1.8]=TRiP.GL01244} | BDSC (41816) |
| Nacalpha | CG8759 | 8759R-3 | NIG-FLY (8759R-3) |
| CG31075 | CG31075 | P{y[+t7.7] v[+t1.8]=TRiP.HMC03055} | BDSC (50654) |
| GstD2 | CG4181 | P{y[+t7.7] v[+t1.8]=TRiP.HMS05960} | BDSC (77361) |
| GstD1 | CG10045 | P{ y[+t7.7] v[+t1.8]=TRiP.HMS05925} | BDSC (77338) |
| Dhd | CG4193 | P{y[+t7.7] v[+t1.8]=TRiP.HMC06450} | BDSC (67345) |
| Cyp6a19 | CG10243 | P{y[+t7.7] v[+t1.8]=TRiP.HMC05648} | BDSC (64613) |
| Akt1 | CG4006 | P{ y[+t7.7] v[+t1.8]=TRiP.HM04007} | BDSC (31701) |
| FoxO | CG3143 | P{ y[+t7.7] v[+t1.8]=TRiP.HMS00793} | BDSC (32993) |
| Ilp4 | CG6736 | TI{w[+mW.hs]=TI}Ilp4[1] | BDSC (30883) |
| Reptor | CG13624 | P{ y[+t7.7] v[+t1.8]=TRiP.GL01180} | BDSC (43960) |
| Tor | CG5092 | P{ y[+t7.7] v[+t1.8]=TRiP.HMS01114} | BDSC (34639) |
| InR | CG18402 | P{PZ}InR$^{05545}$ | BDSC (11661) |
| InR | CG18402 | P{ y[+t7.7] v[+t1.8]=TRiP.HMS03166} | BDSC (51518) |
| InR | CG18402 | P{ y[+t7.7] v[+t1.8]=TRiP.GL00139} | BDSC (35251) |
| Chico | CG5686 | P{ y[+t7.7] v[+t1.8]=TRiP.GL00525} | BDSC (36788) |
| Pi3K21B | CG2699 | P{y[+t7.7] v[+t1.8]=TRiP.GL01028} | BDSC (36810) |
| Girdin | CG12734 | P{y[+t7.7] v[+t1.8]=TRiP.HMS05821} | BDSC(67960) |
| chichadee | CG9553 | P{y[+t7.7] v[+t1.8]=TRiP.HMS00550} | BDSC(34523) |
| chichadee | CG9553 | Chic[221] | BDSC(4892) |
| RpII140 | CG3180 | RpII140[wimp] | BDSC(5874) |

* BDSC = Bloomington Drosophila Stock Center, NIG-FLY = National Institute of Genetics

**Table S2B**. Primers used for q-PCR

| Name | Sequence 5' -> 3' |
|---|---|
| RpL32-F | ATGCTAAGCTGTCGCACAAATG |
| RpL32-R | GTTCGATCCGTAACCGATGT |
| GAPDH-F | TAAATTCGACTCGACTCACGGT |
| GAPDH-R | CTCCACCACATACTCGGCTC |
| Impl2-F | CCTCATTAAGTACGGGATAC |
| Impl2-R | CTTCTGATCTCCGAGATCAAG |
| Link-F | CAGAATGCCCTGTACTACAACC |
| Link-R | ATCAGCAGGATAGCCAGAGGT |
| Thor-F | CAGATGCCCGAGGTGTACTC |
| Thor-R | GGAGTGGTGGAGTAGAGGGTT |
| dtg-F | GCTCTGCGTGATCGTCTTCT |
| dtg-R | ACTGGATGGGAAGTATTGTGGT |
| CG43693-F | TACTGCGTCCACATTCTGGTC |
| CG43693-R | GGCGAATCCCATCATCGGG |
| EGFR-F | TGCATCGGCACTAAATCTCGG |
| EGFR-R | GGAAGCTGAGGTCCAAATTCTC |
| Dpn-F | TGTCCAGAAGTTCAAGACGGG |
| Dpn-R | ATGCCATCCATTTGGCTTACA |
| GP150-F | TCTGACGGAAAACGAGCTGAA |
| GP150-R | CGTCTCCACGGGTAAATCCTTT |
| Nullo-F | ATGGGCAGCACACATTCCG |
| Nullo-R | AGTGCAGAAGATGCCAGGATT |
| Jbug-F | GGGCACAAACGAAACCGAAA |
| Jbug-R | GGCAACCTGGAGATAACGATTT |
| Inx3-F | ACGGCCATTCTGTTCACTTGT |
| Inx3-R | CGTTGATGCAACTTATGGGATCG |
| P32-F | CATGTAACCGTGTTGAATCTCCA |
| P32-R | TCGGCCACTATCTCCTCGG |
| CG1598-F | CGACTTGCAGTTCTAGCCTAGC |
| CG1598-R | GCCGGATCTGTGGATATGATTAG |
| CG3652-F | TTCGGGCCGTAGGCATTAAG |
| CG3652-R | GGTCCCAATCTCGTAGCAGAC |
| ReepB-F | TGGGTCACGTTTGGAATCTTC |
| ReepB-R | GGAAAGCACACTTCAGCAGC |
| CG14545-F | CCGTCTCATCATCGAGCGAA |
| CG14545-R | TCGGTGTACATAGCCCAAGC |
| Rpl23-F | GGAGGCAAGTTCCGCATCTC |
| Rpl23-R | GGACGGCGATCACGTACAG |
| Pex11-F | GACAAAATCGCCCGACTTATTCA |
| Pex11-R | TCGGAATGTGCTCAGGATGTA |
| Cyp6a9-F | GAGTGCAGACCAGTCGATCAT |
| Cyp6a9-R | CAGTAGAAGCCTGCGAAGGG |

| | |
|---|---|
| Exu-F | CTGGCTTGAGCAACTGAAGAC |
| Exu-R | TCATGTAGGGAATGAACTTGCG |
| Nacalpha-F | CGTTAGCTGCACTCCTGATTT |
| Nacalpha-R | ATGTTCGACTGGTGTAACATGC |
| CG31075-F | ATCGACCTGGCTGTGAAAGC |
| CG31075-R | GATCCATCAGGGCACAAAGTT |
| GstD2-F | GACTTTTACTACATGCCAGGTGG |
| GstD2-R | GCAACAGATAGTCATCCTTGCC |
| GstD1-F2 | GGTGACTCCCTTACCGTAGC |
| GstD1-R2 | TTGGCGTACTTGCTGATCTCG |
| dhd-F | CGACGACAAGCTAATCGTGC |
| dhd-R | ACCGCCTTGCTGGAGTATTT |
| Cyp6a19-F | TTCAGGAAACAGACCGACGG |
| Cyp6a19-R | GCGAATTAGCACGGTTTTTACAA |
| FoxO-F | CATGGGGAAATCTATCCTATGCG |
| FoxO-R | ACTCAGTGTCAATCGTTTGTCG |
| Chico-F1 | GCGCACTCACCTTATGACCA |
| Chico-R1 | GCACACGAATGTCAGGGATTT |
| Tor-F | AACGATGTGTCCGTCATGGAG |
| Tor-R | GCCTCTGAGCACCTCAAAAG |
| InR-F | GAAGTGGAGACGACGGGTAAA |
| InR-R | TCGCGCTGTTGTCGATTGTT |
| Pi3k21B-F | ACAAGTGGCAGCAACTTTATGA |
| Pi3k21B-R | CATCCTTTAGCAACCAGAGAGC |
| Reptor-F | CCATCATCCCAGTCCTCTACA |
| Reptor-R | AGTGCATCTAGTCCAAAGTCTGA |
| Akt1-F | CCCAGCGTTACATCGGGTC |
| Akt1-R | GCTCGCCCCTCTTCATCAG |
| Girdin-F | CAGCGTCTGGCTGCAAATC |
| Girdin-R | GGTTTGACCAAGCTCCTCCTC |
| Chic-F | TGCACTGCATGAAGACAACA |
| Chic-R | GTTTCTCTACCACGGAAGCG |
